# Hyperparasitism and the evolution of parasite virulence

**DOI:** 10.1101/2023.01.18.524570

**Authors:** Jason Wood, Ben Ashby

## Abstract

Hyperparasites (species which parasitise other parasites) are common in natural populations, affecting many parasitic taxa, including: eukaryotic parasites; bacterial and fungal pathogens. Hyperparasitism is therefore likely to shape the ecology and evolution of many host-parasite systems, representing a promising method for biocontrol (e.g., treating antimicrobial resistant infections). However, the eco-evolutionary consequences of hyperparasitism have received little attention. We use a host-parasite-hyperparasite model to explore how introducing a hyperparasite drives the evolution of parasite virulence, and what impact this has on the host population. We show when the introduction of a hyperparasite selects for higher or lower parasite virulence, and the changes in virulence experienced by the host population. Crucially, we show that variation in the direct effects of hyperparasites on virulence and transmission, and the probability of co-transmission, can lead to a previously unseen hysteresis effect, whereby small shifts in hyperparasite characteristics can lead to sudden shifts in parasite virulence. We also show that hyperparasites can induce diversification in parasite virulence, leading to the coexistence of high and low virulence strains. Our results show hyperparasites can have dramatic effects on the evolution of parasite virulence, and that the use of hyperparasites in biocontrol should be approached with caution.

## Introduction

Hyperparasitism, wherein parasitic organisms are themselves parasitised by another species, is ubiquitous in the natural world. Hyperparasitism has been observed across many taxa, including bacterial (Greer 2005; Ashelford, Day, and Fry 2003) and fungal (Mikhailov, Simdyanov, and Aleoshin 2016) pathogens, parasitic worms (Mohan et al. 2020) and other eukaryotic parasites (Wendling et al. 2017). For example, many human bacterial pathogens are parasitised by bacteriophages, including *E. Coli* (e.g., phage O157:H7 (Munns et al. 2015)); *Salmonella typhimurium* (e.g., phage *ϕ* AB2 (Berchieri, Lovell, and Barrow 1991)); and *Klebsiella pneumoniae* (e.g., phage B5055 (Chhibber, Kaur, and Kumari 2008)). Hyperparasitism is also common among fungal plant pathogens, such as the powdery mildew fungus *Podosphaera plantaginis* with the hyperparasitic fungus *Ampelomyces* spp. (Parratt and Laine 2018), or the chestnut blight fungus *Cryphonectria parasitica* (Nuss 2005) which can be infected by *Cryphonecria hypovirus 1* (Parratt and Laine 2016).

Hyperparasites are understood to play a major role in the ecology of parasites, and have been shown to influence both the early epidemic dynamics and the overwintering success (Parratt and Laine 2018) of plant pathogens. They have also been implicated as a driver of seasonal epidemics of cholera (Faruque et al. 2005). In addition to ecological effects on the parasite, hyperparasites can have significant effects on virulence, either inducing hypovirulence (a reduction in the virulence experienced by the host, e.g., by reducing the population size of the parasite) or hypervirulence (an increase in the virulence experienced by the host, e.g., by causing the parasite to release toxins or by introducing virulence factors). Hyperparasite-induced changes in virulence have been observed in many bacterial pathogens. For example, a phage protein enhances motility (and hence virulence) of *E. coli* (Kakkanat et al. 2017); the temperate phage PHB09 reduces the virulence of *Bordetella bronchiseptica* both *in vivo* and *in vitro* (Chen et al. 2020); the phage CTX*ϕ* encodes the cholera toxin within *Vibrio cholerae* (Waldor and Mekalanos 1996); and the phage *ϕ*CDHM1 interferes with quorum sensing in *Clostridium difficile* (Hargreaves, Kropinski, and Clokie 2014).

Due to their ecological and virulence-mediating effects, hyperparasites long have been considered as possible sources of biological control (biocontrol) for many infectious diseases (Holtappels et al. 2021; Obradovic et al. 2004), including in the agricultural and food industries and in the treatment of chronic or antimicrobial resistant infections in humans (Gordillo Altamirano and Barr 2019). Indeed, phage therapy has long been used as an alternative to antibiotics in some countries (Schooley et al. 2017; Ferriol-González and Domingo-Calap 2021). However, little is known about the evolutionary implications of hyperparasitism for parasites, and so their use as agents of biocontrol in novel settings could lead to unexpected outcomes for important traits such as parasite transmission and virulence. For instance, (Prospero et al. 2021) speculated that the introduction of hyperparasites which reduce parasite virulence (hypovirulence) might select for higher virulence in chestnut blight fungus, which has been observed experimentally (Bryner and Rigling 2012).

Understanding the interplay of host, parasite and hyperparasite ecological and evolutionary dynamics is crucial not only to the development of novel agents of biocontrol, but also for understanding their role in natural ecosystems. Theoretical studies of hyperparasites are rare and have in the past mainly focussed on their ecological consequences (Taylor et al. 1998; A. Yu. Morozov, Robin, and Franc 2007). Yet recently there has been renewed theoretical interest in hyperparasitism in an evolutionary context, in the form of theoretical models of hyperparasite evolution (Northrup et al. 2021) and parasite-hyperparasite coevolution (Sandhu et al. 2021). Two findings are of particular interest. First, Sandhu et al. (2021), who considered parasite-hyperparasite coevolution, observed that the introduction of hyperparasites always increases parasite virulence, but in almost all scenarios decreases average host mortality. Second, Northrup et al. (2021) observed that when hyperparasites are more readily co-transmitted with evolutionarily static parasites, this selects for less harm by the hyperparasite due to an increased link between its fitness and parasite transmission (similar to virulence evolution in vertically transmitted parasites). These findings highlight the importance of understanding hyperparasitism in both an ecological and an evolutionary context.

Here, we further investigate the eco-evolutionary consequences of introducing a hyperparasite for parasite virulence and the resulting net impact on the host population. We show how the introduction of hyperparasites causes changes in parasite virulence and whether this leads to a net positive or negative effect on the host population depending on the relative transmission rate and virulence of hyperparasitised parasites. We also show how small changes in the effects of the hyperparasite on parasite transmission and virulence can lead to large shifts in the evolution of virulence due to hysteresis. Finally, we show that the hyperparasite can induce diversification in parasite virulence, leading to the coexistence of a relatively high and low virulence strains.

## Methods

### Model description

We consider a well-mixed population of asexual hosts, parasites, and hyperparasites, where *S* is the density of uninfected (susceptible) hosts, *I* is the density of hosts only infected by the parasite (parasitised) and *H* is the density of hosts infected by both the parasite and the hyperparasite (hyperparasitised). Hosts reproduce at a baseline per-capita rate *b*, subject to density-dependent crowding *qN*, with *q* > 0 and *N* = *S* + *I* + *H*. Parasite transmission is density-dependent, with parasitised and hyperparasitised hosts having parasite transmission rates of *β* and *ηβ* to susceptible hosts, respectively, where *β* is the baseline transmission rate, *η* > 1 implies “hypertransmission” (the hyperparasite increases parasite transmissibility) and *η* < 1 implies “hypotransmission” (the hyperparasite decreases parasite transmissibility). Hyperparasites are co-transmitted with parasites to susceptible hosts with probability *ρ*. Hyperparasite infection of parasitised hosts is also density-dependent, with transmission rate σ. All hosts experience a natural mortality rate *d*, with parasitised and hyperparasitised hosts experiencing additional mortality due to disease at rates *α* and *λα*, respectively. Thus, when *λ* < 1 the hyperparasite induces hypovirulence (the hyperparasite decreases the disease-associated mortality rate) and when *λ* > 1 the hyperparasite induces hypervirulence (the hyperparasite increases the disease-associated mortality rate). Both parasitised and hyperparasitised hosts recover from infection at rate *γ*, with no lasting immunity (see Table 1 for a full summary of model parameters and their default values for the analysis).

**Table 1:** Model parameters and default values.

| Parameter | Description | Default value |
| --- | --- | --- |
| $b$ | Natural birth rate of hosts | 2.0 |
| $c$ | Curvature parameter for transmission-virulence trade-off | −60 |
| $d$ | Natural mortality rate of hosts | 0.5 |
| $q$ | Strength of density dependent competition | 0.1 |
| $\alpha$ | Parasite virulence | n/a |
| $\alpha_{max}$ | Maximum parasite virulence | 5.0 |
| $\beta$ | Parasite transmission rate | n/a |
| $\beta_0$ | Scale factor in virulence-transmission trade-off | $\sqrt{5}$ |
| $\beta_s$ | Non-linear transmission scale factor | 0.5 |
| $\beta_l$ | Linear transmission scale factor | 1.0 |
| $\gamma$ | Host recovery rate | 0.5 |
| $\eta$ | Hyperparasite transmission modifier | n/a |
| $\lambda$ | Hyperparasite virulence modifier | n/a |
| $\rho$ | Probability of hyperparasite co-transmission | 0.5 |
| $\sigma$ | Hyperparasite transmission rate | 4.0 |

The ecological dynamics for a monomorphic population are described by the following three ordinary differential equations:

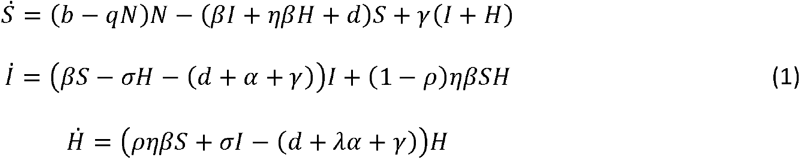

We explore the evolution of parasite virulence under a standard transmission-virulence trade-off with diminishing returns, such that *β* = *β*(*α*), *β′*(*α*) > 0 and *β ″*(*α*) < 0. We consider the following simple trade-off between virulence and transmission,

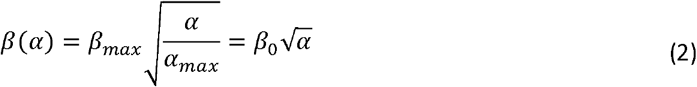

where virulence increases quadratically with the transmission rate of the parasite. To check the generality of our results we consider a second trade-off,

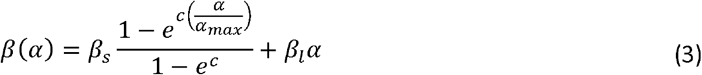

which allows us to explore trade-offs with different curvatures.

We assume that mutations are sufficiently rare so that the ecological dynamics in Equation 1 reach a stable endemic equilibrium (with either the host and parasite, or all three species present; see *Supplementary materials*) before a new mutant arises, and that mutations have small phenotypic effects. Note that since the endemic equilibrium of Equation 1 is analytically intractable, in practice we find the endemic equilibrium and verify its stability numerically. The invasion dynamics of a rare mutant parasite with transmission rate *β*_*m*_ and virulence *α*_*m*_ are then,

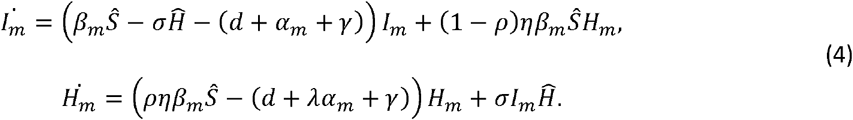

where 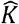 indicates the equilibrium density for population *K* ∈ {*S, H*}. The invasion fitness *w*(*α*_*m*_)is given by the largest eigenvalue of the next generation matrix (Hurford, Cownden, and Day 2010) (see *Supplementary materials*),

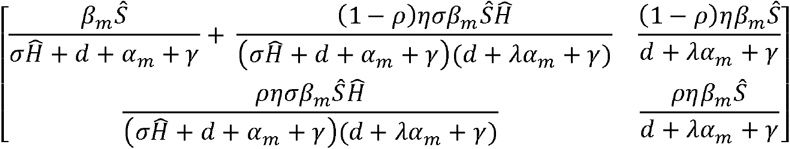

which we omit here for the sake of brevity. We derive the fitness gradient 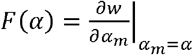 and hence calculate singular strategies, *α\**, that satisfy *F*(*α\**) = 0. To distinguish between direct (*λ*) and evolutionary effects of the hyperparasite on virulence we refer to (*α*\*) as intrinsic virulence. We determine the evolutionary stability of a singular strategy by the sign of 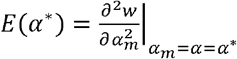 (negative values indicate evolutionary stability), and convergence stability by numerically approximating the derivative 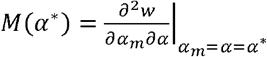 and checking if the inequality *E*(*α\**) + *M*(*α\**) < 0 holds. If a singular strategy is both evolutionarily and convergence stable then it is a continuously stable strategy (an evolutionary attractor); if it is convergence stable but not evolutionarily stable then it is an evolutionary branching point; and if it is convergence unstable then it is an evolutionary repeller.

### Simulations

We complement our numerical analysis with simulations of the evolutionary dynamics of our system, which relax the adaptive dynamics assumptions of continuous traits and separate ecological and evolutionary timescales. To perform our simulations, we first create a discretised trait space for the parasite. We then initialise the hyperparasite-free system at its eco-evolutionary attractor, and introduce hyperparasites at an arbitrarily low population density. We use a fourth order Runga-Kutta method to solve the ordinary differential equations over a long time period, stopping when the populations have relatively small changes in size, or the time threshold has been reached.

After removing phenotypes that fall below an arbitrary threshold, we choose one of the extant parasite phenotypes (using a weighted probability based on parasite density) and introduce a rare mutant a small phenotypic distance away at a low frequency. We then use this new population as the initial condition for our ordinary differential equation system, which we again evaluate using a fourth order Runga-Kutta method. All code used to produce the figures within this paper is available within the Supplementary Material and on GitHub.

### Measuring the impact of the hyperparasite on the host population

We assume that the hyperparasite is introduced into a well-established host-parasite system, such that the parasite is initially at its continuously stable strategy *α*_0_ (see *Supplementary materials*) and the system is at equilibrium. In addition to exploring the evolutionary implications for parasite virulence we also consider the ecological implications for the host. Specifically, we consider two metrics to encapsulate the impact on the host population following the introduction of the hyperparasite, namely: the impact on the host population size 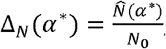, and the impact on the average mortality rate of infected individuals 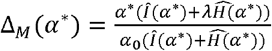, where *N*_0_ corresponds to the steady state of the host population before the introduction of the hyperparasite (with ancestral virulence *α*_0_), and 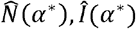 and 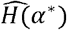 are the steady states following the introduction of the hyperparasite evaluated at a continuously stable strategy, *α\**. For brevity, we refer to Δ_*M*_ (*α\**) as the relative average virulence. When Δ_*M*_ (*α\**) > 1 the hyperparasite leads to an increase in the average virulence experienced by infected hosts, whereas when Δ_*M*_(*α\**) < 1 the average virulence experienced by infected hosts decreases.

## Results

### Hyperparasite-induced shifts in virulence evolution

Following the introduction of the hyperparasite, we see pronounced shifts in the evolution of virulence depending on the extent to which hyperparasites modify parasite transmission (*η*) and/or virulence (*λ*), and the extent to which co-transmission of the hyperparasite (*ρ*) occurs. When the hyperparasite induces hypovirulence (a direct reduction in the disease-associated mortality rate, *λ* < 1), the parasite always experiences selection for higher intrinsic virulence (Fig. 1A). However, when the hyperparasite either has no direct effect on virulence (when *λ* = 1; Fig. 1B) or induces hypervirulence (when *λ* = 1; Fig. 1C) this is no longer the case. When the hyperparasite strongly reduces parasite transmission (*η* ≪ 1), the parasite evolves higher intrinsic virulence, and when the hyperparasite increases transmission (*η* > 1), intrinsic virulence either remains virtually unchanged (*λ* = 1; Fig. 1B) or evolves to lower levels (*η* > 1; Fig. 1C). Yet at intermediate values of *η* < 1 (hypotransmission), the behaviour of the system becomes more complicated, as there is a bistable region which becomes more prominent as the virulence modifier (*λ*) or probability of co-transmission (*ρ*) increase (Fig. 1B-C).

**Figure 1:**
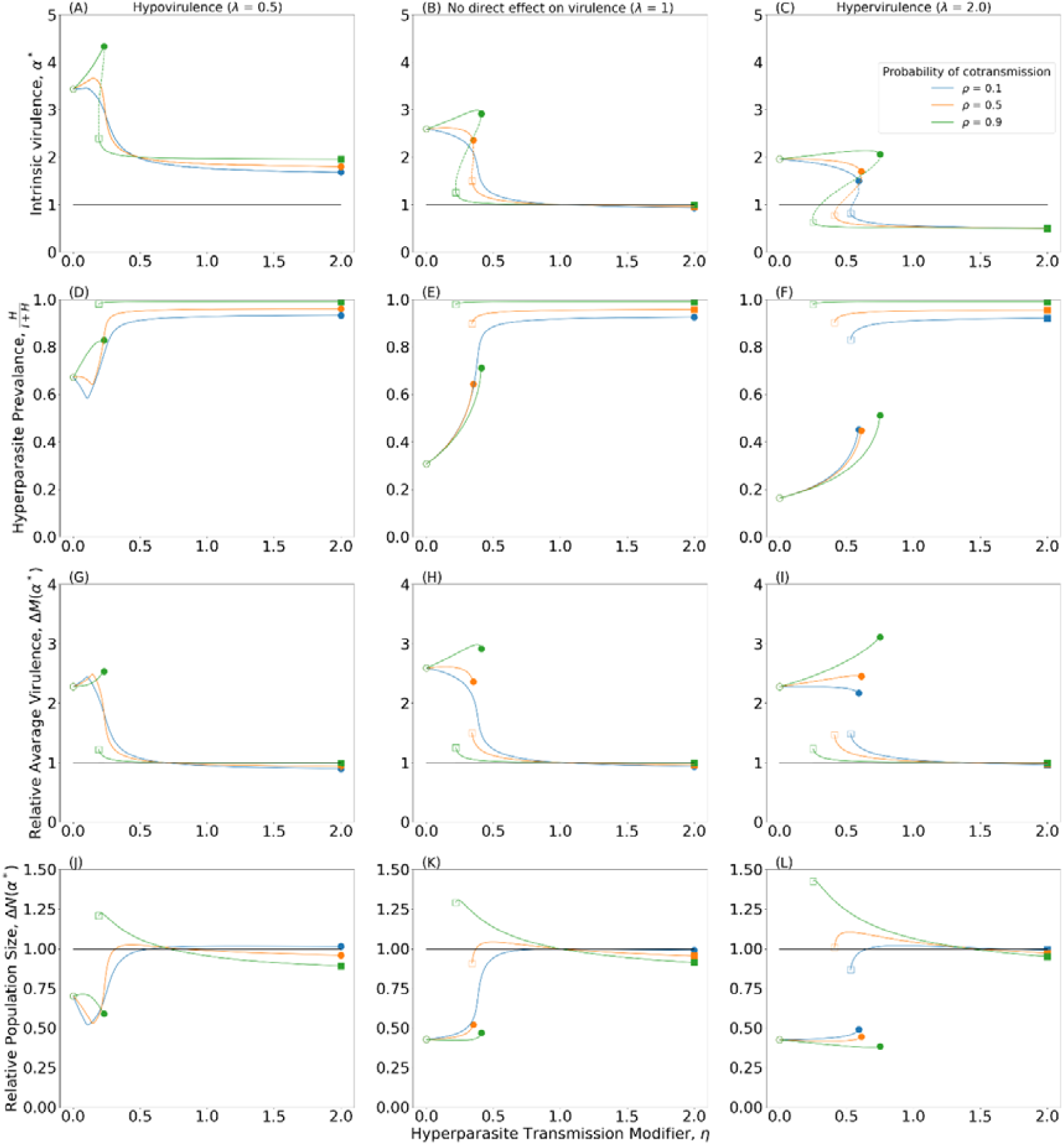
Evolutionary consequences for parasite virulence (A-C) (solid: evolutionary attractors, dashed: evolutionary repellers),the prevalence of the hyperparasite (D-E), relative average mortality (G-I) and relative population size (J-L), as the direct effects of the hyperparasite on parasite transmission hypotransmission: ; hypertransmission: and virulence vary. Evolutionary attractors are both convergence stable and evolutionarily stable, and are therefore continuously stable strategies (CSSs). Left hand column -hypovirulence ; Central column -no effect on virulence ; Right hand column -hypervirulence . All panels contain three sets of curves showing the evolutionary endpoint as the probability of hyperparasite co-transmission varies: (blue), (orange), (green). The start and end of each continuous set of evolutionary attractors are shown with empty and filled shapes. The black line indicates the ancestral state prior to the introduction of the hyperparasite. Here the trade-off used in Equation (2) is used.

The bistable region means that the system has two evolutionary attractors, and therefore small changes in the underlying ecology (e.g., direct effects of the hyperparasite on parasite virulence or transmission) may cause sudden shifts between high and low virulence states (Fig. 1A-C). Crucially, these shifts in virulence may be difficult to reverse as the switching points towards higher or lower intrinsic virulence are not the same (i.e., there is a “hysteresis effect”; Fig. 2). For example, suppose the hyperparasite causes a hypervirulence (*λ* > 1; Fig. 1C), co-transmits with high probability (*ρ* ≈ 1), and reduces the transmissibility of the parasite (*η* = 0.5). Selection would then favour a reduction in intrinsic virulence relative to the absence of the hyperparasite (*α\** < *α*_0_)(Fig. 2). A small reduction in the transmission modifier below a critical threshold, 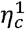, would suddenly shift selection for increased intrinsic virulence (*α\** > *α*_0_), but a reversion in the transmission modifier to its initial value 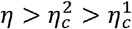 would not lead to a drop in intrinsic virulence until a second critical threshold, 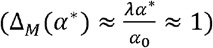 is breached (Fig. 2). Thus, relatively small changes in the effects of the hyperparasite on parasite transmission can lead to large changes in the evolution of virulence that may be difficult to reverse.

**Figure 2:**
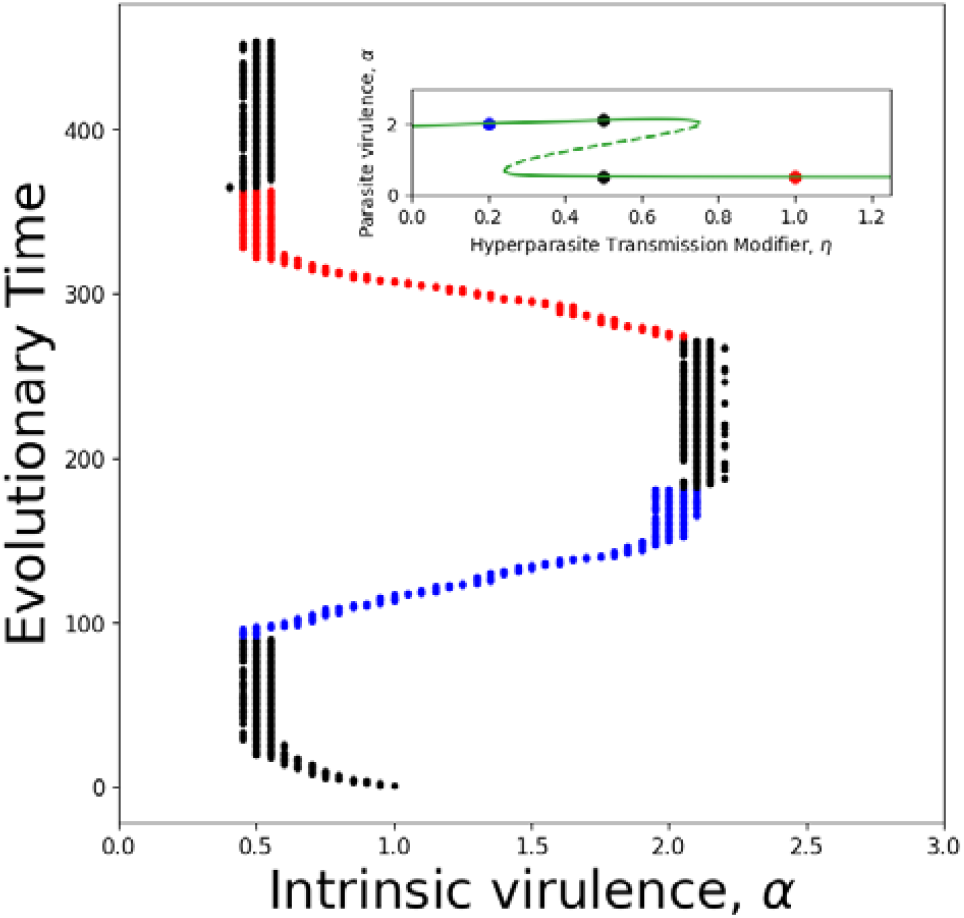
Evolutionary simulations showing the hysteresis effect observed in Fig. 1 due to variation in the direct effects of the hyperparasite on parasite transmission, . A relatively small shift (blue) from an initial value of (black) leads to a sharp increase in intrinsic virulence. However, a reversion to the initial value of (black) does not lead to a return to lower intrinsic virulence. Instead, intrinsic virulence does not change significantly until a critical threshold is reached (red). Parameters as in Table 1 except. Included inset figure replicates Fig. 1C whens, coloured dots indicated predicted for respective value of . Here the trade-off described in Equation (2) is used.

The key driver of the bistable region (and hence the hysteresis effect) is hyperparasite prevalence. The upper virulence branch (Fig. 1A-C) corresponds to relatively low hyperparasite prevalence (Fig. 1D-F), whereas the lower branch corresponds to relatively high prevalence. While there is a numerical feedback between parasite virulence and hyperparasite prevalence, one can understand the link between the two (and hence the hysteresis effect) when viewed through the lens of life-history strategies. Specifically, higher intrinsic virulence is an adaptation to shift reproductive output (transmission) before the parasite is likely to be infected. For example, when the hyperparasite fully prevents parasite transmission(*η* = 0), hyperparasitised hosts make no contribution to parasite fitness and so high intrinsic virulence and transmissibility is advantageous because all transmission must occur before the parasite is infected. Similarly, when the hyperparasite significantly reduces but does not prevent parasite transmission (0 < *η* ≪ 1), most onward transmission occurs before being hyperparasitised. Yet this strategy of shifting reproductive output (transmission) earlier in the parasite’s typical lifespan is only advantageous provided hyperparasite prevalence is relatively low so that sufficient transmission can occur before encountering the hyperparasite.

If the hyperparasite is sufficiently common (prevalence increases with the transmission modifier (*η*) and probability of co-transmission (*ρ*)), then the parasite does not reap the benefits of high transmission before being hyperparasitised. Thus, intrinsic virulence shifts to the lower branch of the bistable region where most parasites are hyperparasitised and there is little change in average virulence 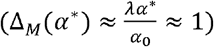. The bistable region therefore exists due to a positive feedback loop between hyperparasite prevalence and parasite virulence (e.g., higher initial hyperparasite prevalence selects for lower intrinsic virulence, which in turn increases hyperparasite prevalence). Note that the key driver of intrinsic virulence along the lower branch appears to be the direct effects of the hyperparasite on virulence (*λ*). Intuitively, if most parasites are hyperparasitised, then *η* (hypo/hyper-transmission) and *ρ* (co-transmission) are close to simple scaling factors for parasite fitness, whereas *λ* (hypo/hyper-virulence) affects the parasite’s infectious period and so is the key driver of selection along this branch.

### Effects of hyperparasitism on the host population

Although the parasite may evolve higher or lower intrinsic virulence following the introduction of the hyperparasite, whether this has a net positive or negative effect for the host population depends on not only the new level of virulence, but also the effects on parasite prevalence, the prevalence of hyperparasites, and the effects of hyperparasitism on virulence. We therefore consider the ecological consequences of the hyperparasite on the host following evolutionary shifts in parasite virulence.

Intuitively, when the hyperparasite selects for higher intrinsic virulence (*α\** > *α*_0_) while not inducing hypovirulence (*λ* ≥ 1), relative average virulence is always higher (Fig. 1H-I), and the resulting host population size is lower (Fig. 1K-L). This only occurs when hyperparasitism strongly reduces transmission(*η* ≪ 1). However, the net impact on hosts is less straightforward when the hyperparasite induces hypovirulence while selecting for higher intrinsic virulence (Fig. 1G, J), or induces hypervirulence while selecting for lower intrinsic virulence (Fig. 1I, L). For example, the latter scenario can lead to little effect on the mortality rate of infected hosts (Fig. 1I) but a marked increase in the host population size (Fig. 1L).

Crucially, the bistability of intrinsic virulence has a striking effect on the host population, with of relatively small shifts in the direct effects of hyperparasites on transmission (*η*) or on the probability co-transmission (*ρ*) causing substantial changes in relative average virulence (Fig. 1H-I) and the host population size (Fig. 1K-L). Typically, infected hosts either experience a significant increase in relative average virulence (for sufficiently low values of *η*), or there is little net effect. This suggests that the introduction of a hyperparasite rarely improves the outcome of infection on average, even if it is beneficial at the population level by reducing the proportion of the population that is infected.

We also see that as the probability of coinfections (*ρ*) increases, the effects of introducing the hyperparasites on the relative average virulence and host population size typically grow stronger. Note that while relative average virulence is close to one for sufficiently large values of *η* in Fig. 1G-I, it does not tend to one (in fact, all the curves pass through 1 before the point *η* = 2; see also Fig. S2, S5).

### Hyperparasite-induced diversification in parasite virulence

In the absence of the hyperparasite, parasite virulence always evolves to a continuously stable strategy due to the trade-off with transmission, which results in diminishing returns for the parasite. The introduction of the hyperparasite either shifts the continuously stable strategy to a new value (as in Fig. 1) or causes the parasite population to diversify through evolutionary branching into two strains, one with relatively high intrinsic virulence which is less hyperparasitised (green branch in Fig. 3) and the other with relatively low intrinsic virulence which is more hyperparasitised (orange branch in Fig. 3). Evolutionary branching occurs when a singular strategy is convergence stable, but is not evolutionarily stable. This does not occur for the quadratic trade-off in Equation 2 as the curvature of the trade-off is too strong. We therefore explored an alternative trade-off (Equation 3) where the curvature is weaker. Note that the second trade-off produces qualitatively similar results to those discussed above (Fig. S2).

**Figure 3.**
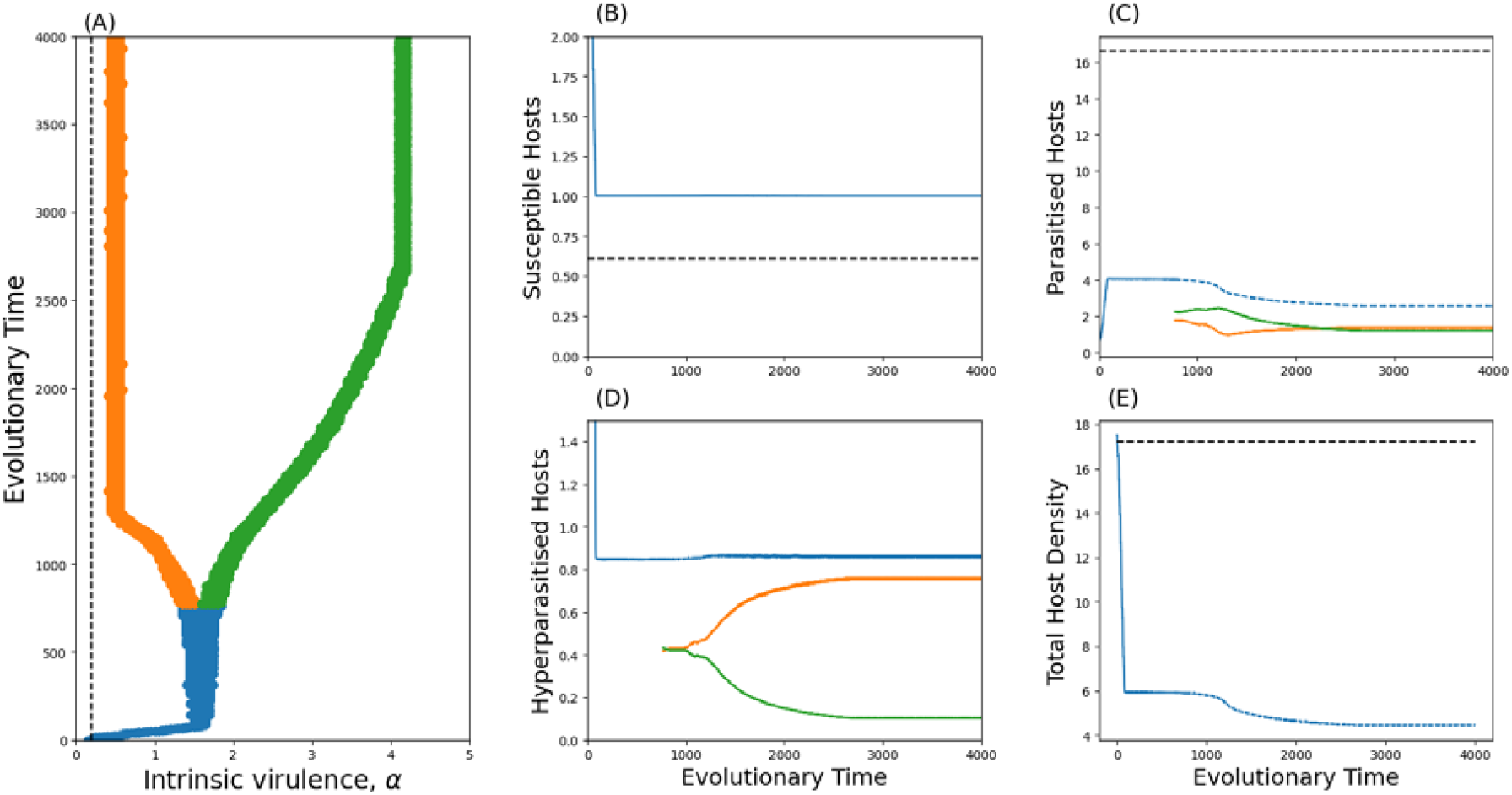
Evolutionary branching in parasite virulence following the introduction of the hyperparasite. Parasite virulence in the absence of the hyperparasite is shown by the dashed line in (A). The parasite branches into relatively high (green) and low (orange) virulence strains. (B-E) Densities of the host, parasite, hyperparasite and total populations: total (blue), high-virulence strain (green), low-virulence strain (orange). The density of the host, parasite and total populations in the absence of the hyperparasite are plotted with a dashed black line. Parameters as in Table 1 except Here the trade-off described in Equation (3) is used.

The more virulent “short-lived” strain essentially prioritises earlier reproductive output (transmission), infecting as many hosts as possible before being infected by the hyperparasite. The less virulent “long-lived” strain instead prioritises transmission over the length of the whole infection. The contrast in strategies can be seen in Fig. 3B-D where the hyperparasite is more commonly associated with the less virulent strain. In general, branching is most likely to occur when the hyperparasite induces hypervirulence (*λ* > 1) and strong hypotransmission (*η* ≪ 1), and the probability of co-transmission is not too high (*ρ* ≪ 1) (Fig. 4).

**Figure 4.**
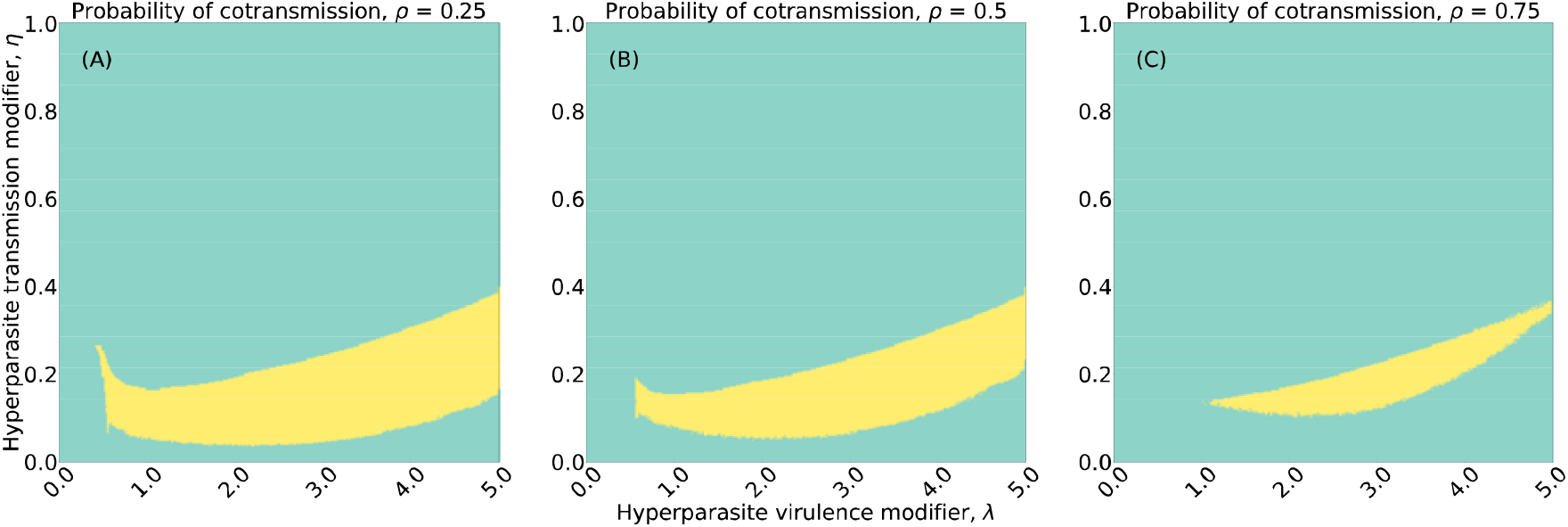
Regions where evolutionary branching occurs (yellow) for the parasite following the introduction of the hyperparasite, as the direct effects on transmission (η) and virulence (λ) vary. Parameters as in Table 1 except d = 0.1, σ = 0.4, γ = 0.1. Here the trade-off described in Equation (3) is used.

However, even though one of the parasite strains is intrinsically less virulent than the other, both are more virulent than the ancestral strain in the absence of the hyperparasite (Fig. 3A). The more virulent strain is rarely infected by the hyperparasite (Fig. 3D), as the host either dies prior to encountering the hyperparasite, or dies after being hyperparasitised having produced few new hyperparasitised individuals (due to relatively lower infectivity). Despite the parasite branching into two distinct strategies that are both intrinsically more virulent than the ancestral state, the overall impact on host and parasite population sizes is minimal (Fig. 3B-D). Additionally, following the introduction of the hyperparasite, there is initially a drastic reduction in host population density, with a further but much smaller decrease in following branching.

## Discussion

Hyperparasites are abundant in nature and are promising sources of biological control in industry (e.g., food production) and public health (e.g., phage therapy to combat antimicrobial resistance). Yet the eco-evolutionary effects of hyperparasitism are poorly understood. In this study, we have theoretically explored the impact of hyperparasites on the evolution of parasite virulence and the consequences for the host population. We have shown that the introduction of hyperparasites can lead to either higher or lower levels of intrinsic virulence depending on the direct effects of the hyperparasitism. Crucially, we have shown that relatively small changes in the ecological effects of hyperparasites can cause large shifts in both evolved and ecologically realised virulence, which can be difficult to reverse due to hysteresis. Although a hyperparasite may cause selection for higher intrinsic virulence, we have shown that this is not necessarily negative for the host population overall, as the hyperparasite can suppress parasite prevalence and may mitigate virulence in infected hosts. Finally, our model shows that the introduction of a hyperparasite can induce diversification in the parasite population, with the potential for both strains to evolve to higher levels of intrinsic virulence than in the absence of the hyperparasite. Overall, our results suggest that the introduction of hyperparasites can have a strong impact on the evolution of parasite virulence and that the nature of the outcome depends crucially on how the hyperparasite directly affects parasite virulence and transmission.

Higher intrinsic virulence typically evolves when the hyperparasite dramatically reduces the transmission rate of the parasite (*η* ≪ 1) and/or the hyperparasite induces hypovirulence (*λ* < 1),yet the reasons differ. Strong hypotransmission selects for higher intrinsic virulence (*α\**) because hyperparasitism results in a significant loss in new infections, and so parasites experience selection to infect as many hosts as possible prior to being hyperparasitised. In contrast, when the hyperparasite induces hypovirulence (*λ* < 1), selection favours an increase in intrinsic virulence because hyperparasitism effectively reduces the costs of higher virulence. This can be trivially understood by considering a hyperparasite that completely mitigates virulence (*λ* = 0), which greatly benefits the parasite by extending the infectious period. Conversely, when the hyperparasite causes hypervirulence (*λ* > 1) it can select for lower intrinsic virulence provided there is not also strong reduction in transmission. Note that we can also conclude that selection for higher intrinsic virulence is driven by hyperparasite prevalence rather than being an adaptation to counteract the direct effects of the hyperparasite on transmission (*η*), as higher intrinsic virulence evolves even when the hyperparasite fully prevents onwards transmission (*η* = 0).

The hysteresis effect in our model suggests that not only might relatively small changes in hyperparasite traits cause large evolutionary changes in the parasite, but that these might be difficult to reverse. Hysteresis arises because for certain regions of parameter space the system is bistable, meaning that the parasite can evolve to have relatively high or low intrinsic virulence depending on the initial level of virulence. The bistability can be understood in terms of a positive feedback loop between hyperparasite prevalence and parasite virulence, with high prevalence selecting for low virulence, and vice versa. When the hyperparasite causes a significant reduction in transmission (*η* ≪ 1) it is very costly to the parasite and so selection favours higher intrinsic virulence as this shifts most transmission to before being hyperparasitised (upper branch in Fig. 1A-C).The hyperparasite is therefore at relatively low prevalence (Fig. 1D-F). As the effects of the hyperparasite on parasite transmission weaken (*η* increases), the hyperparasite may become more C). The common if it can co-transmit well with the parasite (high *ρ*), resulting in a gradual increase in hyperparasite prevalence (Fig. 1D-F). Eventually a threshold is reached where the risk of hyperparasitism is sufficiently high and the cost sufficiently low that selection favours a large reduction in intrinsic virulence (lower branch in Fig. 1A-C), which feeds back to further increase hyperparasite prevalence (Fig. 1D-F). This positive feedback, resulting in high hyperparasite prevalence, is crucial to the hysteresis effect. If the hyperparasite once again reduces parasite transmission more strongly (*η* decreases), the system does not return to the high virulence state because hyperparasite prevalence remains high (Fig. 1D-F). As *η* continues to fall so too does hyperparasite prevalence, until eventually the risk of hyperparasitism is sufficiently low and the cost sufficiently high that selection favours a large increase in intrinsic virulence (upper branch in Fig. 1A-C).

The hysteresis effect has especially important implications for biocontrol, as a relatively small change in the hyperparasite can have a large effect on the evolution of virulence and on the host population. It might be possible to reverse sudden shifts in virulence by removing the biocontrol for a period of time, as selection would then push virulence towards the ancestral state, but the rate at which this occurs will be system-specific. While hysteresis is well documented in the ecological literature (e.g., the spruce budworm (Ludwig, Jones, and Holling 1978)), there are fewer examples in evolutionary models (Kisdi and Geritz 1999; Ronce and Kirkpatrick 2001; Prado et al. 2009; Berdahl et al. 2015; Fortelius et al. 2015; Osmond and Klausmeier 2017; Toivonen and Fromhage 2019). For example, Prado *et al*. 2009 observed hysteresis when considering the evolution of host sociality and pathogen virulence within contact networks. Prado *et al*. observed that the cycling behaviour they see within their system, has hysteresis-like behaviour where selection for or against host sociality does not occur until the parasite passes critical thresholds.

Our model reveals that the introduction of a hyperparasite can cause disruptive selection leading to diversification into relatively high and low virulence parasite strains (although the “low” virulence strain may still be more virulent than the ancestral strain, as in Fig. 3A). Branching typically requires trade-offs that are close to linear (Bowers et al. 2005), such as the trade-off presented in Equation 3, as strongly diminishing returns have a balancing effect on selection. Branching also typically requires the hyperparasite to directly reduce the transmission rate of the parasite, which facilitates the existence of high and low virulence phenotypes by creating distinct ecological niches. The high virulence phenotype is rarely infected by the hyperparasite, favouring a “live fast, die young” strategy, while the less virulent phenotype has a longer infectious period to mitigate the burden of the hyperparasite.

Our study is closely related to previous theoretical explorations of evolution in host-parasite-hyperparasite systems (Sandhu et al. 2021; Northrup et al. 2021). Sandhu et al (2021) also explored the effects of hyperparasitism on the evolution of parasite virulence, but in contrast to our study found that the introduction of a hyperparasite always selects for increased virulence and generally reduces the average mortality rate of the host. Furthermore, Sandhu et al (2021) did not observe a hysteresis effect, nor did they find diversification in virulence. The differences in our results are likely due to several crucial differences in our assumptions. Here, we modelled both hypo- and hyper-virulence and transmission, whereas Sandhu et al (2021) focused on hypo-virulence and transmission. When we restrict our parameter space to hypo-virulence (*λ* < 1) and transmission (*η* < 1) we regain the key finding from Sandhu et al (2021) that higher virulence always evolves (Fig. 1A). However, this does not hold when there is hypervirulence (*λ* > 1 Fig. 1C). The hysteresis effect produced by the bistable region is also more prominent when there is hypervirulence, which may explain why it has not previously been observed. Recovery is possible for many infectious diseases and moreover, one would not expect the hyperparasite to always co-transmit with the parasite (*ρ* < 1). When we remove recovery from our model (*γ* = 0), and assume that the hyperparasite is always co-transmitted with the parasite (*ρ* = 1), as in Sandhu et al (2021), the bistable region is still present (Fig. S3), and so these differences in model assumptions do not explain the lack of bistability in Sandhu et al (2021). Notably, Sandhu et al (2021) allowed superinfection (replacement of parasite strains) mediated by the hyperparasite and assumed that the transmissibility of the hyperparasite (*σ*) was linked to the infectivity of the parasite, such that more infectious parasites also generated more infections by hyperparasites. These effects were not present in our model. As the bistable region in our model represents a change in parasite strategies due to contrasts in hyperparasite prevalence, with more virulent parasites evolving when hyperparasite prevalence is lower, the positive relationship between parasite transmission/virulence and hyperparasite infectivity may explain the lack of bistability in Sandhu et al (2021).

Although models of hyperparasitism are relatively rare, especially in an evolutionary context, some of our key findings are mirrored in models of other tripartite systems. For example, multiple studies have found that introducing an additional species to a host-parasite system can lead to evolutionary branching in the host (Best 2018) or the parasite (Kisdi, Geritz, and Boldin 2013; Best 2018; Smith and Ashby 2022; Morozov and Best 2012) populations. In a related study that also has relevance to biocontrol, Smith and Ashby (2022) explore how the introduction of a tolerance-conferring defensive symbiont affects the evolution of parasite virulence. They show that even if the defensive symbiont is initially beneficial to the host population, in the long-term it is costly because it always selects for higher parasite virulence. More generally, recent theory has highlighted how biocontrol agents can lead to unintuitive ecological and evolutionary outcomes (Ashby and King 2017; Rafaluk-Mohr et al. 2018; King and Bonsall 2017; Nelson and May 2020). Our results similarly emphasise the complex eco-evolutionary outcomes that can arise following the introduction of a hyperparasite, with potentially disastrous consequences for the host population.

In established host-parasite-hyperparasite systems, it is difficult to separate the ecological and evolutionary consequences of the hyperparasite. Few empirical studies have therefore explored the effects of hyperparasitism on parasite evolution (Parratt and Laine 2016), and those that do often focus on consequences for antimicrobial resistance (Chan et al. 2016; Burmeister et al. 2020), the acquisition of virulence factors from hyperparasites (Miao and Miller 1999), or pleiotropic effects on virulence due to selection for resistance against hyperparasitism (Castledine et al. 2022). For example, (Evans et al. 2010) showed that strains of the bacteria *Erwinia carotovora ssp. atroseptica (Eca)* resistant to the hyperparasite *ϕ* AT1 were less likely to produce rot in potato tubers; Casteldine et al. showed that evolution of phage resistance in *Pseudomonas aeruginosa* coincided with the loss of virulence in vitro and in vivo; and (Scanlan and Buckling 2012) showed that coevolution between the bacteria *P. fluourescens* and a lytic phage (⍰2) selects for a mucoid phenotype, which is a virulence factor in both lung infections of cystic fibrosis patients and in plant infections. Clearly, the available empirical evidence suggests that hyperparasites can indeed have significant evolutionary effects on parasite virulence, although the precise effects may depend on pleiotropy between resistance and virulence. Our model did not consider resistance to hyperparasitism but understanding how pleiotropy with resistance affects virulence evolution is a critical direction for future theoretical work (Sandhu et al. 2021). Additionally, were the hyperparasite to die out at any point, the parasite would always return to the ancestral level of virulence prior to the introduction of the hyperparasite (Fig. S4).

We focused our investigation on the evolution of virulence, but the evolutionary dynamics of the hyperparasite are also likely to be important (Sandhu et al. 2021; Northrup et al. 2021), especially for driving additional eco-evolutionary feedback loops (Ashby et al. 2019) . However, in certain cases the hyperparasite might behave as if it is evolutionarily static (e.g., the repeated application of a particular biocontrol to an agricultural crop), in which case the model analysed here will be especially applicable. Still, future theoretical work should consider how our findings are affected by coevolution with the parasite and/or the host (Buckingham and Ashby 2022), and whether the hysteresis in our model could lead to fluctuating selection as in Prado et al. (2009). Additionally, the joint dynamics of resistance and virulence evolution deserve further scrutiny, which could be explored using a resource-allocation model where the parasite can either allocate resources to transmission or defence, with the overall resource “budget” depending on the level of virulence.

Overall, we have shown that the introduction of hyperparasites can have dramatic effects on the evolutionary dynamics of parasite virulence. These effects can lead to selection for higher or lower virulence, or to evolutionary branching. The effect is critically dependent on the within-host effects of the hyperparasite on transmission and virulence, and on the probability of co-transmission. We have also shown how relatively small changes in hyperparasite traits can have dramatic consequences for virulence evolution, which has important implications for the use of hyperparasites as agents of biocontrol.

## Supporting information

Code_base

## Acknowledgements

JW is supported by a scholarship from the EPSRC Centre for Doctoral Training in Statistical Applied Mathematics at Bath (SAMBa), under the project EP/L015684/1. BA is supported by the Natural Environment Research Council (grant numbers NE/N014979/1 and NE/V003909/1). We acknowledge the support of the Natural Sciences and Engineering Research Council of Canada (NSERC). Nous remercions le Conseil de recherches en sciences naturelles et en génie du Canada (CRSNG) de son soutien.

