## Supplementary material for "Hyperparasitism and the evolution of parasite virulence": Code_base: analysis.sh

```
#!/bin/bash
mkdir outputs
cd outputs
mkdir single_evo
cd ../data
mkdir alpha_evo
cp tracker_file_base_version.csv tracker_file.csv
cd ../
mkdir supplementary_figures
pip install -r requirements.txt
cd H-P-HP/
python3 setup.py build_ext --inplace
python3 ad_dyn_hysteresis.py
python3 eta_evo.py
# The file linear_heatmap_res_scalar.py is commented out as it takes ~1 day to run on
# a local machine
# python3 linear_heatmap_res_scalar.py 
python3 visualise_branching_matrices.py
python3 branching_with_linear.py
```
