## Supplementary material for "Hyperparasitism and the evolution of parasite virulence": Code_base: ad_dyn_hysteresis.py

```
"""
This file demonstrates the consequences of the hysteresis affect caused by the introduction
of hyperparasites. This file is used to create Fig. 2.
"""
#We import the libraries we need
import sys, os, shutil
import runner
import numpy as np
import matplotlib.pyplot as plt
import pandas as pd
#These local libraries are used to reduce the length of the file
sys.path.append("../src/Models/")

import singular_strategy as ss

sys.path.append("../src/zero_finder/")

import zero_finder_utilities as zf

#This function calculates the gradient vector for a given parameter set
def calculate_alpha_gradient_vector(sol, resolution, beta_max, alpha_max, alpha_min_res, alpha_max_res, sigma_max, sigma, b, q, d, rho, eta, gamma, lam, c1, c2, hyper, seed):
    
    alpha_grad_vec = []
    alpha_val = []
    
    for i in range(resolution+1):
        
        #calculate the value of alpha
        alpha = alpha_min_res + (alpha_max_res - alpha_min_res) * (i / resolution)
        alpha_val.append(alpha)

        #and corresponding value of beta and it's derivatives
        beta = zf.beta_func(beta_max, alpha, alpha_max, sigma, sigma_max, c1, c2)
        
        dbetadalpha = zf.dbetadalpha_func(
            beta_max, alpha, alpha_max, sigma, sigma_max, c1, c2
        )

        #Solve the system to ecological steady state
        y = sol.eco_steady_state(
            beta, alpha, sigma, b, q, d, rho, eta, gamma, lam, c1, c2, hyper, seed
        )

        S = y[0]
        I = y[1]
        H = y[2]

        #Determine which version of the gradient we need. If the parasite is extinct just add an
        #arbitrary value
        if (I == 0 and H == 0):
            alpha_grad = 1*(alpha <= alpha_max/2) - 1*(alpha> alpha_max/2)
        elif (H == 0):
            alpha_grad = dbetadalpha*S/(d + alpha + gamma) - (beta*S)/((d + alpha + gamma)**2)
        elif (I == 0):
            alpha_grad = dbetadalpha*S/(d + lam*alpha + gamma) - (beta*S)/((d + lam*alpha + gamma)**2)
        else:
#             if rho != 1:
            alpha_grad = ss.fitness_gradient_alpha(
                beta, alpha, sigma, dbetadalpha, H, S, eta, lam, gamma, d, rho
            )
#             else:
#                 if (beta*S - sigma*H - (d + alpha)) > (eta*beta*S - (d + alpha*lam)):
#                 alpha_grad = dbetadalpha*S/(sigma*H + d + alpha + gamma)  - beta*S/((sigma*H + d + alpha + gamma)**2)
#                 else:
#                     alpha_grad = eta*dbetadalpha*S/(lam*alpha + d + gamma) - eta*beta*S/((lam*alpha + d + gamma)**2)
        #Convert the value into a sign
        alpha_grad = (alpha_grad > 0)*1 + (alpha_grad < 0)*-1
        alpha_grad_vec.append(alpha_grad)
        
    return alpha_grad_vec, alpha_val

#Iterate through the gradient vector to find any sign changes
def return_zeros(grad_vec, param_space):
    
    poss_attractors = []
    poss_repellers = []
    for i,val in enumerate(grad_vec[:-1]):
        if val == 1 and grad_vec[i+1] == -1:
            poss_attractors.append([param_space[i], param_space[i+1]])
        elif val == -1 and grad_vec[i+1] == 1:
            poss_repellers.append([param_space[i], param_space[i+1]])
        elif val == 0 and grad_vec[i+1] == -1:
            poss_attractors.append([param_space[i], param_space[i+1]])
            
    return poss_attractors, poss_repellers 
    
#This function calculates the singular strategy using the above functions to calculate the gradient and then find any sign changes
def recusive_sing_strat_function(sol, depth, iter_max, resolution, beta_max, alpha_max, alpha_min_res, alpha_max_res, sigma_max, sigma, b, q, d, rho, eta, gamma, lam, c1, c2, hyper, seed):
    
    
    sing_strats = []
    
    #We fill these lists with attractors and repellers
    attractors = []
    repellers = []
    
    #If we have resolved sufficiently we do one final resolution before exiting
    if depth == iter_max:
        #Calculating the alpha gradient vector
        alpha_grad_vec, alpha_val = calculate_alpha_gradient_vector(sol, resolution, beta_max, alpha_max, alpha_min_res, alpha_max_res, sigma_max, sigma, b, q, d, rho, eta, gamma, lam,c1, c2, hyper, seed)
        
        #Identifying singular strategies
        poss_attractors, poss_repellers = return_zeros(alpha_grad_vec,alpha_val)
#         if len(poss_attractors) > 2:
#             print(f"On run {eta} on rho {rho} with depth {depth}, and there are {len(poss_attractors)} possible singular strategies     ", end = "\r")
        #Allocating them to their respective places
        for zeros in poss_attractors:
            #We take the lower element as the midpoint can sometimes be outside the bounds of the hyperparasites existance
            attractors.append(zeros[0])
        for zeros in poss_repellers:
            repellers.append(zeros[0])
            
    #If we are not at our maximum depth we find possible positions of the singular strategy
    else:
        #First we calculate the alpha gradient
        alpha_grad_vec, alpha_val = calculate_alpha_gradient_vector(sol, resolution, beta_max, alpha_max, alpha_min_res, alpha_max_res, sigma_max, sigma, b, q, d, rho, eta, gamma, lam,c1, c2, hyper, seed)
        
        #Check for any sign changes using our function
        poss_attractors, poss_repellers = return_zeros(alpha_grad_vec,alpha_val)
#         if len(poss_attractors) > 2:
#             print(f"On run {eta} on rho {rho} with depth {depth}, and there are {len(poss_attractors)} possible singular strategies     ")
#             for pair in poss_attractors:
#                 print(pair[1] - pair[0])
        #Check bounds for any attractor regions
        if depth == 0:
            if alpha_grad_vec[0] == -1:
                attractors.append(alpha_val[0])
            if alpha_grad_vec[-1] == 1:
                attractors.append(alpha_val[-1])
                
        #Rerun the process of finding singular strategies for each of our possible attractors and repellers
        for pair in poss_attractors:
            temp_attractors, temp_repellers = recusive_sing_strat_function(sol, depth+1, iter_max, resolution, beta_max, alpha_max, pair[0], pair[1], sigma_max, sigma, b, q, d, rho, eta, gamma, lam, c1, c2, hyper, seed)
            
            for val in temp_attractors:
                attractors.append(val)
                
        for pair in poss_repellers:
            temp_attractors, temp_repellers = recusive_sing_strat_function(sol, depth+1, iter_max, resolution, beta_max, alpha_max, pair[0], pair[1], sigma_max, sigma, b, q, d, rho, eta, gamma, lam, c1, c2, hyper, seed)
                
            for val in temp_repellers:
                repellers.append(val)
                
    #We then clean up the attractors in case there are any singular strategies very close together
    if depth == 0:
        attractors = zf.clean_up_sing_strats(attractors)
        repellers = zf.clean_up_sing_strats(repellers)
                
    return attractors, repellers


#Parameters used in the system
rho = 0.9

lam = 2.0

b = 2.0
q = 0.1
d = 0.5
gamma = 0.5

seed = 100
resolution = 100
sol = runner.PySolver()

evo_etas = [0.5, 0.2, 0.5, 1.0, 0.5]
    
plotting_alphas = []
plotting_evos = []
alpha_init = 20
sigma_init = 100

alpha_max = 5
sigma_max = 4
beta_max = 5

sigma_min = 0.0
sigma_max_range = sigma_max

sigma = sigma_max

hyper = 1.0
c1 = 0.75
c2 = 2.25


beta_max = beta_max
alpha_max_param = alpha_max 
sigma_max_param = sigma_max

iter_max = 3
res_scalar = 4
depth_max = iter_max
resolution = 100
param_res = 200

eta_min = 0.0
eta_max = 1.25
etas = [eta_min + (eta_max - eta_min)*i/param_res for i in range(param_res + 1)]


#Here we create our temporary lists to store the values of the singular strategies
plotting_attractors_hyper = []
plotting_attractors_no_hyper = []

plotting_repellers_hyper = []
plotting_repellers_no_hyper = []

plotting_etas_attractors_hyper = []
plotting_etas_attractors_no_hyper = []

plotting_etas_repellers_hyper = []
plotting_etas_repellers_no_hyper = []

for eta_t in etas:
    eta = round(eta_t,4)
    print(f"On run {eta} on rho {rho}, and lam {lam}  ", end = "\r")

    alpha_max_res = alpha_max
    alpha_min_res = 0.0

    alpha_grad_vec, alpha_val = calculate_alpha_gradient_vector(sol, resolution, beta_max, alpha_max, alpha_min_res, alpha_max_res, sigma_max, sigma, b, q, d, rho, eta, gamma, lam, c1, c2, hyper, seed)


    alpha_grad_no_hyper_vec, alpha_val = calculate_alpha_gradient_vector(sol, resolution, beta_max, alpha_max, alpha_min_res, alpha_max_res, sigma_max, sigma, b, q, d, rho, eta, gamma, lam, c1, c2, 0.0, seed)

    #Calculating the singular strategies when the hyperparasite is present and absent
    attractors_hyper, repellers_hyper = recusive_sing_strat_function(sol, 0, iter_max, resolution, beta_max, alpha_max, alpha_min_res, alpha_max_res, sigma_max, sigma, b, q, d, rho, eta, gamma, lam, c1, c2, hyper, seed)

    attractors_no_hyper, repellers_no_hyper = recusive_sing_strat_function(sol, 0, iter_max, resolution, beta_max, alpha_max, alpha_min_res, alpha_max_res, sigma_max, sigma, b, q, d, rho, eta, gamma, lam, c1, c2, 0.0, seed)

    #Storing these values in our lists from before
    for val in attractors_hyper:
        plotting_etas_attractors_hyper.append(eta)
        plotting_attractors_hyper.append(val)

    for val in attractors_no_hyper:
        plotting_etas_attractors_no_hyper.append(eta)
        plotting_attractors_no_hyper.append(val)

    for val in repellers_hyper:
        plotting_etas_repellers_hyper.append(eta)
        plotting_repellers_hyper.append(val)

    for val in repellers_no_hyper:
        plotting_etas_repellers_no_hyper.append(eta)
        plotting_repellers_no_hyper.append(val)

    for i, val in enumerate(alpha_val):
        if val >= attractors_no_hyper[0]:
            break

#Adding these lists to our stored dictionaries
plotting_dict_hyper = {}
plotting_dict_no_hyper = {}

plotting_dict_hyper["a"] = plotting_etas_attractors_hyper
plotting_dict_no_hyper["a"] = plotting_etas_attractors_no_hyper

plotting_dict_hyper["r"] = plotting_etas_repellers_hyper
plotting_dict_no_hyper["r"] = plotting_etas_repellers_no_hyper

plotting_dict_attractors = plotting_attractors_hyper
plotting_dict_attractors_no_hyper = plotting_attractors_no_hyper

plotting_dict_repellers = plotting_repellers_hyper
plotting_dict_repellers_no_hyper = plotting_repellers_no_hyper


eta_mins = {}
eta_maxs = {}

temp_etas = plotting_dict_hyper["r"]
if len(temp_etas) > 0:
    eta_mins[rho] = min(temp_etas)
    eta_maxs[rho] = max(temp_etas)

#As we have a hystersis effect within our plots it can be difficult to visualise smoothly
alpha_lower_plotting = {}
alpha_upper_plotting = {}
alpha_attractors = {}
eta_attractors = {}
temp_alphas = plotting_dict_repellers
eta_attractors[rho] = {}
alpha_attractors[rho] = {}
eta_attractors[rho]['1'] = []
eta_attractors[rho]['2'] = []
alpha_attractors[rho]['1'] = []
alpha_attractors[rho]['2'] = []

#This section checks if there are any repellers, meaning we have the hysteresis effect present.
if len(temp_alphas) > 0:
    alpha_lower_plotting[rho] = min(temp_alphas)
    alpha_upper_plotting[rho] = max(temp_alphas)

    flag = True
    paired_list = []
    for i, val in enumerate(plotting_dict_attractors):
        paired_list.append([val, plotting_dict_hyper["a"][i]])
    sorted_attractors = sorted(paired_list, key = lambda x: x[0], reverse = True)
    temp_attractors_1 = []
    temp_attractors_2 = []
    temp_etas_1 = []
    temp_etas_2 = []

    #This section checks to see if we have jumped from one branch of the hysteresis affect to the other
    for i, val in enumerate(sorted_attractors[:-1]):
        if flag and abs(sorted_attractors[i+1][0] - val[0]) < 0.1:
            temp_attractors_1.append(val[0])
            temp_etas_1.append(val[1])
        elif flag:
            flag = False
            temp_attractors_1.append(val[0])
            temp_etas_1.append(val[1])
        else:
            temp_attractors_2.append(val[0])
            temp_etas_2.append(val[1])

    temp_attractors_2.append(sorted_attractors[-1][0])
    temp_etas_2.append(sorted_attractors[-1][1])

    paired_list_1 = []
    for i, val in enumerate(temp_etas_1):
        paired_list_1.append([val, temp_attractors_1[i]])

    paired_list_2 = []
    for i, val in enumerate(temp_etas_2):
        paired_list_2.append([val, temp_attractors_2[i]])

    #We sort the values to be in terms of eta again once they have been properly seperated
    sorted_list_1 = sorted(paired_list_1, key = lambda x: x[0])
    sorted_list_2 = sorted(paired_list_2, key = lambda x: x[0])

    temp_attractors_1 = [val[1] for val in sorted_list_1]
    temp_attractors_2 = [val[1] for val in sorted_list_2]

    temp_etas_1 = [val[0] for val in sorted_list_1]
    temp_etas_2 = [val[0] for val in sorted_list_2]

    alpha_attractors[rho]['1'] = temp_attractors_1
    alpha_attractors[rho]['2'] = temp_attractors_2
    eta_attractors[rho]['1'] = temp_etas_1
    eta_attractors[rho]['2'] = temp_etas_2
else:
    alpha_attractors[rho]['1'] = plotting_dict_attractors
    eta_attractors[rho]['1'] = plotting_dict_hyper["a"]

# print(alpha_attractors[rho]['1'])

evo_step_cutoff = 90

#We initialise the system with the hyperparasites absent to find the natural ESS of the parasite
sol.alpha_ad_dyn(beta_max, alpha_max, sigma_max, b, q, d, rho, 0.0, gamma, 0.0, c1, c2, 0.0, seed, alpha_init, sigma_init, H_density = 0.0)
df = pd.read_csv(f"../data/alpha_evo/data_set{seed}.csv")

evo_steps = df["Evolutionary_step"].values
dft = df[df["Evolutionary_step"]==evo_steps[-1]]

pops = []
for val in dft["Trait_index_1"].values:
    df_pop = dft[dft["Trait_index_1"]==val][["Density_of_parasite"]]
    pops.append((df_pop["Density_of_parasite"].iloc[0])/(sum(dft["Density_of_parasite"].values)))
    
#Calculating the average trait value
alpha_weights = []
for i, val in enumerate(pops):
    alpha_weights.append(val*dft["Trait_index_1"].values[i])
alpha_mean = sum(alpha_weights)

#Rounding this to the closest integer to use as our initial conditions
alpha_init = round(alpha_mean)

host_density = dft["Density_of_Hosts"].iloc[0]
para_density = sum(dft["Density_of_parasite"].values)
hyper_density = 0.1

#We iterate through the list of etas, showing the consequences of the hysteresis affect
for eta in evo_etas:
    
    sol.alpha_ad_dyn(beta_max, alpha_max, sigma_max, b, q, d, rho, eta, gamma, lam, c1, c2, hyper, seed, alpha_init, sigma_init, S_density = host_density, I_density = para_density, H_density = hyper_density)
    df = pd.read_csv(f"../data/alpha_evo/data_set{seed}.csv")

    seed += 1
    df = df[df["Evolutionary_step"]<=evo_step_cutoff]
    alpha_coords = df["alpha_val"].values
    evo_steps = df["Evolutionary_step"].values
    
    plotting_alphas.append(alpha_coords)
    plotting_evos.append(evo_steps)
    
    dft = df[df["Evolutionary_step"]==evo_step_cutoff]
    para_densities = dft["Density_of_parasite"].values
    hyper_densities = dft["Density_of_hyperparasite"].values
    trait_inds = dft["Trait_index_1"].values
    alpha_init = 0
    max_density = 0
    for i, den in enumerate(para_densities):
        if (den + hyper_densities[i]) > max_density:
            alpha_init = trait_inds[i]
    
    host_density = dft["Density_of_Hosts"].iloc[0]
    para_density = sum(dft["Density_of_parasite"].values)
    hyper_density = sum(dft["Density_of_hyperparasite"].values)
    
del sol


#Here we visualise the plots by sequencing them after each other using
#this upshift
up_shift = []
for i in range(len(etas)):
    up_shift.append(i*(evo_step_cutoff + 1))
    
#Colours associated with the values of eta
colours = {0.5:"black", 0.2:"blue", 1.0:"red"}
n = 8
#Creating and saving the figure
fig, ax = plt.subplots(figsize = (n, n))
left, bottom, width, height = [0.4, 0.7, 0.45, 0.10]
ax_new = fig.add_axes([left, bottom, width, height])
    
tagged_labels = []
for i,eta in enumerate(evo_etas):
    
    alphas = plotting_alphas[i]
    evos_temp = plotting_evos[i]
    
    evo_steps = []
    for step in evos_temp:
        evo_steps.append(step + up_shift[i])
        
    if eta not in tagged_labels:
        ax.scatter(alphas, evo_steps, marker = ".", label = fr"$\eta$ = {eta}", c = colours[eta])
        tagged_labels.append(eta)
    else:
        ax.scatter(alphas, evo_steps, marker = ".", c = colours[eta])
        
ax_new.plot(eta_attractors[rho]['1'],alpha_attractors[rho]['1'], c = "tab:green")
ax_new.plot(eta_attractors[rho]['2'],alpha_attractors[rho]['2'], c = "tab:green")
#This checks if we need to plot the repellers and then links the repellers to the start and end of the attractors
#as appropriate
if len(plotting_dict_hyper["r"]) > 0:
    repeller_etas = [eta_attractors[rho]["2"][0]]
    for val in plotting_dict_hyper["r"]:
        repeller_etas.append(val)
    repeller_etas.append(eta_attractors[rho]["1"][-1])

    repeller_alphas = [alpha_attractors[rho]['2'][0]]
    for val in plotting_dict_repellers:
        repeller_alphas.append(val)

    repeller_alphas.append(alpha_attractors[rho]["1"][-1])
else:
    repeller_etas = plotting_dict_hyper["r"]
    repeller_alphas = plotting_dict_repellers

ax_new.plot(repeller_etas, repeller_alphas,  c = "tab:green", linestyle='dashed')
ax_new.set_xlim([0,etas[-1]])
for i,val in enumerate(plotting_etas_attractors_hyper):
    if val in evo_etas:
        print(val, plotting_attractors_hyper[i])
        ax_new.scatter(val, plotting_attractors_hyper[i], c = colours[val])
        
# fig.legend(
#    fontsize = 2*n,
#    loc="upper right",
#    bbox_to_anchor=(0.5,0.5)
# )
ax.set_xlabel(r"Intrinsic virulence, $\alpha$", fontsize = 28)
ax.set_xlim([0,3])
ax.tick_params(axis='both', which='major', labelsize=12)
ax.set_ylabel("Evolutionary Time", fontsize = 28)

ax_new.set_xlabel(r"Hyperparasite Transmission Modifier, $\eta$", fontsize = 10)
ax_new.set_ylim([0,3])
ax_new.set_ylabel(r"Parasite virulence, $\alpha$", fontsize = 10)

# ax.legend(loc='center left', bbox_to_anchor=(0.6, 0.85), fontsize = 14)
plt.savefig("../supplementary_figures/ad_dyn_figure.png", bbox_inches='tight')
plt.savefig("../supplementary_figures/ad_dyn_figure.pdf", bbox_inches='tight')
plt.close()
```
