## Supplementary material for "Hyperparasitism and the evolution of parasite virulence": Code_base: branching_with_linear.py

```
""" This file produces a single plot showing branching for a given value of 
eta and lambda. This file uses the pandas, numpy, math, and matplotlib 
libraries. This library also uses a custom runner library created using
cyton and local cpp files.
"""

#Importing libraries needed to perform the analysis
import pandas as pd
import numpy as np
from math import exp
import runner
import matplotlib.pyplot as plt

#Invoking the cpp based solver
sol = runner.PySolver()

#Creating the panel labels we will use in our figures
panel_labels = ["(A)", "(B)", "(C)", "(D)", "(E)"]

#Parameters of the system
b = 2.0
q = 0.1
d = 0.1
gamma = 0.1
sigma = 0.4
sigma_max = sigma
c1 = 1.0

alpha_init = 3
sigma_init = 100
hyper = 1.0
beta_scalar = 1.0
rho = 0.5

#The specific values of eta and lambda chosen for this plot
eta = 0.15
lam = 1.0

#The seed used to track the simulation, and the number of evolutionary steps
seed = 10000
evo_step_size = 4000

#The limits of the parameter space for the parasite
alpha_min = 0.0
alpha_max = 5.0

#The parameters of the trade-off
beta_max = 0.5
beta_lin = 1.0
c2 = -60

#Performing the simulation in the absence of the hyperparasite to get a baseline
seed_for_baseline = 10
rho_baseline = eta_baseline = lam_baseline = 0.0
sol.alpha_ad_dyn_v4(beta_max, beta_lin, alpha_max, alpha_min, sigma_max, b, q, d, rho_baseline, eta_baseline, gamma, lam_baseline, c1, c2, beta_scalar, 0.0, seed_for_baseline, alpha_init, sigma_init)

#Reading in the data associated with the baseline simulation
df_baseline = pd.read_csv(f"../data/alpha_evo/data_set{seed_for_baseline}.csv")
evo_steps = df_baseline["Evolutionary_step"].values

#Calculate the proportion of the population with each trait value
dft = df_baseline[df_baseline["Evolutionary_step"]==evo_steps[-1]]
pops = []
for val in dft["Trait_index_1"].values:
    df_pop = dft[dft["Trait_index_1"]==val][["Density_of_parasite"]]
    pops.append((df_pop["Density_of_parasite"].iloc[0])/(sum(dft["Density_of_parasite"].values)))
    
#Calculating the average trait value
alpha_weights = []
for i, val in enumerate(pops):
    alpha_weights.append(val*dft["Trait_index_1"].values[i])
alpha_mean = sum(alpha_weights)

#Rounding this to the closest integer to use as our initial conditions
alpha_init = round(alpha_mean)

#The densities we initiate our populations with, taken from the end of the simulation
#without the hyperparasite present
host_density = dft["Density_of_Hosts"].iloc[0]
para_density = sum(dft["Density_of_parasite"].values)

#Initialising the hyperparasites with a low density
hyper_density = 0.1

#Performing the simulations with the hyperparasite present
sol.alpha_ad_dyn_v4(beta_max, beta_lin, alpha_max, alpha_min, sigma_max, b, q, d, rho, eta, gamma, lam, c1, c2, beta_scalar, hyper, seed, alpha_init, sigma_init, evo_step_size, S_density = host_density, I_density = para_density, H_density = hyper_density)

#Reading in the data associated with hyperparasite simulation
df = pd.read_csv(f"../data/alpha_evo/data_set{seed}.csv")
alpha_vals = df["alpha_val"].values
evo_steps = df["Evolutionary_step"].values

#Finding the point at which branching occurs
branching_flag = False
for step in set(evo_steps):
    dft = df[df["Evolutionary_step"]==step]
    trait_temps = sorted(dft["Trait_index_1"].values)
    for i, val in enumerate(trait_temps[:-1]):
        if trait_temps[i+1] - val > 1:
            branching_step = step
            branching_flag = True
            break
    if branching_flag:
        break

#Creating a calculating the various populations of hosts, parasites and 
#hyperparasites, before and after the branching event
S_vals = []
I_vals_pre_branch = []
H_vals_pre_branch = []
I_branch_1 = []
H_branch_1 = []
I_branch_2 = []
H_branch_2 = []
alpha_vals_pre_branch = []
alpha_branch_1 = []
alpha_branch_2 = []

total_pre_branch = []
total_post_branch = []

evo_steps_pre_branch_sim = []
evo_steps_branch_1 = []
evo_steps_branch_2 = []
evo_steps_unique = [i for i in set(evo_steps)]

I_post_branch_total = []
H_post_branch_total = []
for step in evo_steps_unique:
    dft = df[df["Evolutionary_step"]==step]
    S_vals.append(dft["Density_of_Hosts"].iloc[0])
    if step < branching_step:
        I_vals_pre_branch.append(sum(dft["Density_of_parasite"].values))
        H_vals_pre_branch.append(sum(dft["Density_of_hyperparasite"].values))
        total_pre_branch.append(S_vals[-1] + I_vals_pre_branch[-1] + H_vals_pre_branch[-1])
        for val in dft["alpha_val"].values:
            alpha_vals_pre_branch.append(val)
        for val in dft["Evolutionary_step"].values:
            evo_steps_pre_branch_sim.append(val)
    else:
        trait_temps = sorted(dft["Trait_index_1"].values)
        branch_1_inds = []
        for i, val in enumerate(trait_temps[:-1]):
            if trait_temps[i+1] - val > 1:
                branch_1_inds.append(val)
                break
            else:
                branch_1_inds.append(val)
        dft_b1 = dft[dft["Trait_index_1"].isin(branch_1_inds)]
        dft_b2 = dft[~dft["Trait_index_1"].isin(branch_1_inds)]
        I_branch_1.append(sum(dft_b1["Density_of_parasite"].values))
        H_branch_1.append(sum(dft_b1["Density_of_hyperparasite"].values))
        I_branch_2.append(sum(dft_b2["Density_of_parasite"].values))
        H_branch_2.append(sum(dft_b2["Density_of_hyperparasite"].values))
        I_post_branch_total.append(I_branch_1[-1] + I_branch_2[-1])
        H_post_branch_total.append(H_branch_1[-1] + H_branch_2[-1])
        
        total_post_branch.append(S_vals[-1] + I_post_branch_total[-1] + H_post_branch_total[-1])
        
        for val in dft_b1["alpha_val"].values:
            alpha_branch_1.append(val)
        for val in dft_b2["alpha_val"].values:
            alpha_branch_2.append(val)
            
        for val in dft_b1["Evolutionary_step"].values:
            evo_steps_branch_1.append(val)
        for val in dft_b2["Evolutionary_step"].values:
            evo_steps_branch_2.append(val)
evo_steps_pre_branch = []
evo_steps_post_branch = []
for val in evo_steps_unique:
    if val < branching_step:
        evo_steps_pre_branch.append(val)
    else:
        evo_steps_post_branch.append(val)
        
#Rerunning the simulations without the hyperparasite to track the population of hosts and
#parasites when the hyperparasite is absent
sol.alpha_ad_dyn_v4(beta_max, beta_lin, alpha_max, alpha_min, sigma_max, b, q, d, rho, eta, gamma, lam, c1, c2, beta_scalar, 0.0, seed + 1, alpha_init, sigma_init, evo_step_size)

df_2 = pd.read_csv(f"../data/alpha_evo/data_set{seed+1}.csv")

evo_steps_2 = df_2["Evolutionary_step"].values

dft = df_2[df_2["Evolutionary_step"]==evo_steps_2[-1]]
pops = []
for val in dft["alpha_val"].values:
    df_pop = dft[dft["alpha_val"]==val][["Density_of_parasite","Density_of_hyperparasite"]]
    pops.append((df_pop["Density_of_parasite"].iloc[0])/(sum(dft["Density_of_parasite"].values)))
    
alpha_weights = []
for i, val in enumerate(pops):
    alpha_weights.append(val*dft["alpha_val"].values[i])
alpha_mean = sum(alpha_weights)

host_density = [dft["Density_of_Hosts"].iloc[0] for i in range(evo_steps_2[-1] + 1)]
para_density = [sum(dft["Density_of_parasite"].values) for i in range(evo_steps_2[-1] + 1)]
alpha_means = [alpha_mean for i in range(evo_steps_2[-1]+1)]
evo_plotting_mean = [i for i in range(evo_steps_2[-1]+1)]

#Creating the figures
fig = plt.figure(figsize = (20, 10))
plt.subplots_adjust(wspace=0.3, hspace = 0.2)
gs = fig.add_gridspec(2,3)
ax = [fig.add_subplot(gs[:, 0]), fig.add_subplot(gs[0,1]), fig.add_subplot(gs[0,2]), fig.add_subplot(gs[1,1]), fig.add_subplot(gs[1,2])]

#Making the various subfigures
ax[0].scatter(alpha_vals_pre_branch, evo_steps_pre_branch_sim)
ax[0].scatter(alpha_branch_1, evo_steps_branch_1)
ax[0].scatter(alpha_branch_2, evo_steps_branch_2)
ax[0].plot(alpha_means, evo_plotting_mean, "k--")
ax[0].set_xlim([0,alpha_max])
ax[0].set_ylim([0, max(evo_steps)])
ax[0].set_xlabel(r"Intrinsic virulence, $\alpha$", fontsize = 20)
ax[0].set_ylabel("Evolutionary Time", fontsize = 20)
ax[0].text(0.02,1.01,panel_labels[0], transform=ax[0].transAxes, fontsize = 18)

ax[1].plot(evo_steps_unique, S_vals)
ax[1].plot(evo_plotting_mean, host_density, "k--")
ax[1].set_xticks([0,1000,2000,3000,4000])
ax[1].set_xlim([0, max(evo_steps)])
ax[1].set_ylim([0, 2])
ax[1].set_ylabel(r"Susceptible Hosts", fontsize = 18)
ax[1].text(0.02,1.05,panel_labels[1], transform=ax[1].transAxes, fontsize = 18)

max_I_values = [max(I_vals_pre_branch), max(I_branch_1), max(I_branch_2)]
max_H_values = [max(H_vals_pre_branch), max(H_branch_1), max(H_branch_2)]
ax[2].plot(evo_steps_pre_branch, I_vals_pre_branch)
ax[2].plot(evo_plotting_mean, para_density, "k--")
ax[2].set_xticks([0,1000,2000,3000,4000])
ax[2].plot(evo_steps_post_branch, I_branch_1)
ax[2].plot(evo_steps_post_branch, I_branch_2)
ax[2].plot(evo_steps_post_branch, I_post_branch_total, "C0--")
ax[2].set_xlim([0, max(evo_steps)])

ax[2].set_ylabel(r"Parasitised Hosts", fontsize = 18)
ax[2].text(0.02,1.05,panel_labels[2], transform=ax[2].transAxes, fontsize = 18)

ax[3].plot(evo_steps_pre_branch, H_vals_pre_branch)
ax[3].set_xticks([0,1000,2000,3000,4000])
ax[3].plot(evo_steps_post_branch, H_branch_1)
ax[3].plot(evo_steps_post_branch, H_branch_2)
ax[3].plot(evo_steps_post_branch, H_post_branch_total, "C0--")
ax[3].set_xlim([0, max(evo_steps)])
ax[3].set_xlabel("Evolutionary Time", fontsize = 18)

ax[3].set_ylim([0, 1.5])
ax[3].set_ylabel(r"Hyperparasitised Hosts", fontsize = 18)
ax[3].text(0.02,1.05,panel_labels[3], transform=ax[3].transAxes, fontsize = 18)

ax[4].plot(evo_steps_pre_branch, total_pre_branch)
ax[4].set_xticks([0,1000,2000,3000,4000])
ax[4].plot(evo_steps_post_branch, total_post_branch, "C0--")
ax[4].plot([evo_steps_2[0], evo_steps_2[-1]], [host_density[-1] + para_density[-1], host_density[-1] + para_density[-1]], "k--")
ax[4].set_ylabel("Total Host Density", fontsize = 18)
ax[4].set_xlabel("Evolutionary Time", fontsize = 18)
ax[4].text(0.02,1.05,panel_labels[4], transform=ax[4].transAxes, fontsize = 18)
#Saving the figure as a png and a pdf
plt.savefig("../supplementary_figures/branching_fig.pdf", bbox_inches = "tight")
plt.savefig("../supplementary_figures/branching_fig.png", bbox_inches = "tight")
plt.close()
```
