## Supplementary material for "Hyperparasitism and the evolution of parasite virulence": Code_base: disease_density_for_different_alphas.py

```
"""This file creates a folder with a unique 4 word title in the single_evo folder
located within the outputs folder. We then calculate the singular strategies
for a given set of parameters and a specified trade-off. Just for personal use.
"""
import sys, os, shutil
import runner
import numpy as np
from math import sqrt, exp, dist, floor, ceil, isnan
import plotly.graph_objects as go
from time import sleep
import matplotlib.pyplot as plt
import seaborn as sn
import imageio
import multiprocessing as mp
import pandas as pd

#These local libraries are used to reduce the length of the file
sys.path.append("../src/Models/")

import singular_strategy as ss

sys.path.append("../src/zero_finder/")

import zero_finder_utilities as zf

rho = 0.1
lam = 0.5
eta =  0.25

b = 2.0
q = 0.1
d = 0.5
gamma = 0.5

hyper = 1.0
c1 = 0.75
c2 = 2.25
b_base = 1
q_base = 1

alpha_max = 5
sigma_max = 4
beta_max = 5

sigma_min = 0.0
sigma_max_range = sigma_max

sigma = sigma_max

iter_max = 3
res_scalar = 4
depth_max = iter_max
resolution = 100
param_res = 200

seed = 10000

#Initiating the CPP solver
sol = runner.PySolver()

alphas = [i*alpha_max/resolution for i in range(resolution + 1)]

total_infected_density = []
hyperparasite_props = []
for alpha in alphas:
    
    beta = zf.beta_func(beta_max, alpha, alpha_max, sigma, sigma_max, c1, c2)
    
    #Solve the system to ecological steady state
    y = sol.eco_steady_state(
        beta, alpha, sigma, b, q, d, rho, eta, gamma, lam, c1, c2, hyper, seed
    )

    S = y[0]
    I = y[1]
    H = y[2]
    total_infected_density.append(I + H)
    if H == 0:
        hyperparasite_props.append(0)
    else:
        hyperparasite_props.append(H/(total_infected_density[-1]))
    
plt.plot(alphas,total_infected_density)
plt.plot([0.5, 0.5], [0, 10], "k--")
plt.plot([2.0, 2.0], [0, 10], "k--")
plt.ylabel("Density of Infecteds")
plt.xlabel(r"Parasite Virulence, $\alpha$")
plt.savefig("../supplementary_figures/infected_density.png", bbox_inches = "tight")
plt.close()

plt.plot(alphas,hyperparasite_props)
plt.plot([0.5, 0.5], [0, 1.0], "k--")
plt.plot([2.0, 2.0], [0, 1.0], "k--")
plt.ylabel("Proportion of hyperparasites")
plt.xlabel(r"Parasite Virulence, $\alpha$")
plt.savefig("../supplementary_figures/hyperparasite_proportions.png", bbox_inches = "tight")
plt.close()
```
