## Supplementary material for "Hyperparasitism and the evolution of parasite virulence": Code_base: ecological_heatmaps.py

```
""" 
This file creates Fig. S1, the ecological heatmaps, parameters below can be changed
to investigate different scenarios.
"""

#Libraries used in the analysis
import sys, os, shutil
import runner
import numpy as np
from math import sqrt, exp, dist, floor, ceil, isnan
import plotly.graph_objects as go
from time import sleep
import matplotlib.pyplot as plt
import seaborn as sn
import imageio
import multiprocessing as mp
import pandas as pd

#Initialising C++ solver
sol = runner.PySolver()

#Parameters used to make grid
lams = [0.5, 1.0, 2.0]
etas = [0.5, 1.0, 2.0]

#Parameters which are not varied
b = 2.0
q = 0.1
d = 0.1
gamma = 0.1
hyper = 1.0
c1 = 0.75
c2 = 2.25
rho = 0.5

b_base = 1
q_base = 1

alpha_max = 5
sigma_max = 4.0
beta_max = 5

sigma = 0.4

resolution = 100

#Parameters swept over
param_res = 400

alphas = [alpha_max*i/(param_res) for i in range (param_res + 1)]
betas = [beta_max*i/(param_res) for i in range (param_res + 1)]

seed = 10000

#Labels for sub figures
panel_labels = [["(A)", "(B)", "(C)"],["(D)", "(E)", "(F)"], ["(G)","(H)","(I)"]]

#Initialising and creating list of figures
fig = plt.figure(figsize = (100, 100))
gs = fig.add_gridspec(3,3)
ax = []

for i in range(3):
    temp_ax = []
    for j in range(3):
        temp_ax.append(fig.add_subplot(gs[i,j]))
    ax.append(temp_ax)
    
#Looping through parameters of interest
for eta_tracker, eta in enumerate(etas):
    for lam_tracker, lam in enumerate(lams):
        
        #Creating empty matrix which will be filled with outputs
        current_mat = []
        for beta_val in betas:
            temp_row = []
            for alpha in alphas:
                
                #Population densities in absence of hyperparasite
                y = sol.eco_steady_state(
                    beta_val, alpha, sigma, b, q, d, rho, eta, gamma, lam, c1, c2, 0.0, seed
                )
                
                #Populations if hyperparasite invades
                y = sol.eco_steady_state(
                    beta_val, alpha, sigma, b, q, d, rho, eta, gamma, lam, c1, c2, hyper, seed, y[0], y[1], 4
                )

                #Outcome population sizes
                S = y[0]
                I = y[1]
                H = y[2]
                
                #Boolean labels determining which scenario occured
                if S > 1e-3 and I < 1e-3 and H < 1e-3:
                    score = 1
                elif S > 1e-3 and I > 1e-3 and H < 1e-3:
                    score = 2
                elif S > 1e-3 and I > 1e-3 and H > 1e-3:
                    score = 3
                #This scenario shouldn't happen, so if it does we flag it and throw an error
                else:
                    print(S,I,H)
                    
                    #Undefined variable to cause error
                    flop
                    
                temp_row.append(score)
            current_mat.append(temp_row)
        
        ax[eta_tracker][lam_tracker].imshow(current_mat, origin = "lower", cmap = "tab10")
        
        ticks_temp = []
        for k in range(len(current_mat)):
            ticks_temp.append(k)

        #Creating ticks for plotting
        alpha_ticks = [0.0, 1.0, 2.0, 3.0, 4.0, 5.0]
        beta_ticks = [0.0, 1.0, 2.0, 3.0, 4.0, 5.0]
        ticks = [i for i in ticks_temp if alphas[i] in alpha_ticks]
        
#         print(f"eta {eta}, lam {lam}")
#         print(current_mat[0][-1])
#         print(current_mat[-1][-1])
#         print(current_mat[int(param_res/2)][int(param_res/2)])
#         print("")
        ax[eta_tracker][lam_tracker].set_yticks(ticks = ticks, labels = beta_ticks, fontsize = 2*resolution/3)
        ax[eta_tracker][lam_tracker].set_xticks(ticks = ticks, labels = alpha_ticks, fontsize = 2*resolution/3, rotation=45)
        ax[eta_tracker][lam_tracker].set_title(fr"$\eta$ = {eta} and $\lambda$ = {lam}", fontsize = 2*resolution/3)
        
        #Adding labels where appropriate
        if lam_tracker == 0 and eta_tracker == 1:
            ax[eta_tracker][lam_tracker].set_ylabel(r"Parasite transmission, $\beta$", fontsize = 2*resolution/3)
        if lam_tracker == 1 and eta_tracker == 2:
            ax[eta_tracker][lam_tracker].set_xlabel(r"Parasite virulence , $\alpha$", fontsize = 2*resolution/3)
        ax[eta_tracker][lam_tracker].text(0.05,0.9,panel_labels[eta_tracker][lam_tracker], transform=ax[eta_tracker][lam_tracker].transAxes, fontsize = 100)    
        
#Saving Figures as pngs and pdfs
plt.savefig(f"../supplementary_figures/ecological_heatmaps.pdf", bbox_inches = "tight")
plt.savefig(f"../supplementary_figures/ecological_heatmaps.png", bbox_inches = "tight")
plt.close()
```
