## Supplementary material for "Hyperparasitism and the evolution of parasite virulence": Code_base: eta_evo.py

```
"""This file creates a folder with a unique 4 word title in the single_evo folder
located within the outputs folder. We then calculate the singular strategies
for a given set of parameters and a specified trade-off. This file is used to make
Fig. 1 and Fig. S5.
"""
import sys, os, shutil
import runner
import numpy as np
from math import sqrt, exp, dist, floor, ceil, isnan
import plotly.graph_objects as go
from time import sleep
import matplotlib.pyplot as plt
import seaborn as sn
import imageio
import multiprocessing as mp
import pandas as pd

#These local libraries are used to reduce the length of the file
sys.path.append("../src/Models/")

import singular_strategy as ss

sys.path.append("../src/zero_finder/")

import zero_finder_utilities as zf

sys.path.append("../src/figures/")

import figure_utilities as ff

from datetime import datetime

#This creates a list of words and cleans them for use later
with open("../data/words.txt", encoding='utf8') as f:
    lines = f.readlines()
    
lines_cleaned = []

for line in lines:
    if line[0] != "#" and "www" not in line and line[0] != " ":
        lines_cleaned.append(line[:-1])

#This function calculates the gradient vector for a given parameter set
def calculate_alpha_gradient_vector(sol, resolution, beta_max, alpha_max, alpha_min_res, alpha_max_res, sigma_max, sigma, b, q, d, rho, eta, gamma, lam, c1, c2, hyper, seed):
    
    alpha_grad_vec = []
    alpha_val = []
    
    for i in range(resolution+1):
        
        #calculate the value of alpha
        alpha = alpha_min_res + (alpha_max_res - alpha_min_res) * (i / resolution)
        alpha_val.append(alpha)

        #and corresponding value of beta and it's derivatives
        beta = zf.beta_func(beta_max, alpha, alpha_max, sigma, sigma_max, c1, c2)
        
        dbetadalpha = zf.dbetadalpha_func(
            beta_max, alpha, alpha_max, sigma, sigma_max, c1, c2
        )

        #Solve the system to ecological steady state
        y = sol.eco_steady_state(
            beta, alpha, sigma, b, q, d, rho, eta, gamma, lam, c1, c2, hyper, seed
        )

        S = y[0]
        I = y[1]
        H = y[2]
        
        if y[3] != 1:
            print(S, I, H, rho, lam, eta, alpha)
            raise Exception("Solver did not exit early due to convergence")
        #Determine which version of the gradient we need. If the parasite is extinct just add an
        #arbitrary value
        if (I == 0 and H == 0):
            alpha_grad = 1*(alpha <= alpha_max/2) - 1*(alpha> alpha_max/2)
        elif (H == 0):
            alpha_grad = dbetadalpha*S/(d + alpha + gamma) - (beta*S)/((d + alpha + gamma)**2)
#         elif (I == 0):
#             alpha_grad = dbetadalpha*S/(d + lam*alpha + gamma) - (beta*S)/((d + lam*alpha + gamma)**2)
        else:
#             if rho != 1:
            alpha_grad = ss.fitness_gradient_alpha(
                beta, alpha, sigma, dbetadalpha, H, S, eta, lam, gamma, d, rho
            )
#             else:
#                 if (beta*S - sigma*H - (d + alpha)) > (eta*beta*S - (d + alpha*lam)):
#                 alpha_grad = dbetadalpha*S/(sigma*H + d + alpha + gamma)  - beta*S/((sigma*H + d + alpha + gamma)**2)
#                 else:
#                     alpha_grad = eta*dbetadalpha*S/(lam*alpha + d + gamma) - eta*beta*S/((lam*alpha + d + gamma)**2)
        #Convert the value into a sign
        alpha_grad = (alpha_grad > 0)*1 + (alpha_grad < 0)*-1
        alpha_grad_vec.append(alpha_grad)
        
    return alpha_grad_vec, alpha_val

#Iterate through the gradient vector to find any sign changes
def return_zeros(grad_vec, param_space):
    
    poss_attractors = []
    poss_repellers = []
    for i,val in enumerate(grad_vec[:-1]):
        if val == 1 and grad_vec[i+1] == -1:
            poss_attractors.append([param_space[i], param_space[i+1]])
        elif val == -1 and grad_vec[i+1] == 1:
            poss_repellers.append([param_space[i], param_space[i+1]])
        elif val == 0 and grad_vec[i+1] == -1:
            poss_attractors.append([param_space[i], param_space[i+1]])
            
    return poss_attractors, poss_repellers 
    
#This function calculates the singular strategy using the above functions to calculate the gradient and then find any sign changes
def recusive_sing_strat_function(sol, depth, iter_max, resolution, beta_max, alpha_max, alpha_min_res, alpha_max_res, sigma_max, sigma, b, q, d, rho, eta, gamma, lam, c1, c2, hyper, seed):
    
    
    sing_strats = []
    
    #We fill these lists with attractors and repellers
    attractors = []
    repellers = []
    
    #If we have resolved sufficiently we do one final resolution before exiting
    if depth == iter_max:
        #Calculating the alpha gradient vector
        alpha_grad_vec, alpha_val = calculate_alpha_gradient_vector(sol, resolution, beta_max, alpha_max, alpha_min_res, alpha_max_res, sigma_max, sigma, b, q, d, rho, eta, gamma, lam,c1, c2, hyper, seed)
        
        #Identifying singular strategies
        poss_attractors, poss_repellers = return_zeros(alpha_grad_vec,alpha_val)
#         if len(poss_attractors) > 2:
#             print(f"On run {eta} on rho {rho} with depth {depth}, and there are {len(poss_attractors)} possible singular strategies     ", end = "\r")
        #Allocating them to their respective places
        for zeros in poss_attractors:
            #We take the lower element as the midpoint can sometimes be outside the bounds of the hyperparasites existance
            attractors.append(zeros[0])
        for zeros in poss_repellers:
            repellers.append(zeros[0])
            
    #If we are not at our maximum depth we find possible positions of the singular strategy
    else:
        #First we calculate the alpha gradient
        alpha_grad_vec, alpha_val = calculate_alpha_gradient_vector(sol, resolution, beta_max, alpha_max, alpha_min_res, alpha_max_res, sigma_max, sigma, b, q, d, rho, eta, gamma, lam,c1, c2, hyper, seed)
        
        #Check for any sign changes using our function
        poss_attractors, poss_repellers = return_zeros(alpha_grad_vec,alpha_val)
#         if len(poss_attractors) > 2:
#             print(f"On run {eta} on rho {rho} with depth {depth}, and there are {len(poss_attractors)} possible singular strategies     ")
#             for pair in poss_attractors:
#                 print(pair[1] - pair[0])
        #Check bounds for any attractor regions
        if depth == 0:
            if alpha_grad_vec[0] == -1:
                attractors.append(alpha_val[0])
            if alpha_grad_vec[-1] == 1:
                attractors.append(alpha_val[-1])
                
        #Rerun the process of finding singular strategies for each of our possible attractors and repellers
        for pair in poss_attractors:
            temp_attractors, temp_repellers = recusive_sing_strat_function(sol, depth+1, iter_max, resolution, beta_max, alpha_max, pair[0], pair[1], sigma_max, sigma, b, q, d, rho, eta, gamma, lam, c1, c2, hyper, seed)
            
            for val in temp_attractors:
                attractors.append(val)
                
        for pair in poss_repellers:
            temp_attractors, temp_repellers = recusive_sing_strat_function(sol, depth+1, iter_max, resolution, beta_max, alpha_max, pair[0], pair[1], sigma_max, sigma, b, q, d, rho, eta, gamma, lam, c1, c2, hyper, seed)
                
            for val in temp_repellers:
                repellers.append(val)
                
    #We then clean up the attractors in case there are any singular strategies very close together
    if depth == 0:
        attractors = zf.clean_up_sing_strats(attractors)
        repellers = zf.clean_up_sing_strats(repellers)
                
    return attractors, repellers

#This function works through the singular strategies and calculates the ecological consequences of them
def produce_suplimentary_data(
    sol,
    plotting_sing_strats_hyper,
    plotting_hyper,
    plotting_sing_strats_no_hyper,
    plotting_no_hyper,
    beta_max,
    alpha_max,
    sigma_max,
    sigma,
    c1,
    c2,
    b,
    q,
    d,
    rho,
    gamma,
    lam,
    hyper,
    seed,
):

    death_measure_1 = []

    death_measure_2 = []

    relative_pop_size = []

    relative_infected_proportion = []

    hyperparasite_densities = []
    infected_densities = []
    host_densities = []
    
    pop_virulences = []
    #We work through the attractors calculated by our singular strategy functions
    for i, alpha in enumerate(plotting_sing_strats_hyper):

        #We get the associated value of eta
        eta = plotting_hyper[i]
        
        no_hyper_ind = plotting_no_hyper.index(eta)
        
        #We find alpha in the absence of hyperparasites
        alpha_no_hyper = plotting_sing_strats_no_hyper[no_hyper_ind]

        #Calculate associated beta values
        beta_val = zf.beta_func(beta_max, alpha, alpha_max, sigma, sigma_max, c1, c2)
        beta_no_hyper = zf.beta_func(beta_max, alpha_no_hyper, alpha_max, sigma, sigma_max, c1, c2)

        #Find our populations when hyperparasites are present
        y = sol.eco_steady_state(
            beta_val, alpha, sigma, b, q, d, rho, eta, gamma, lam, c1, c2, hyper, seed
        )

        S = y[0]
        I = y[1]
        H = y[2]
        
        host_densities.append(S)
        infected_densities.append(I)
        hyperparasite_densities.append(H)
        
        N = S + I + H 

        #And absent
        y_no_hyper = sol.eco_steady_state(
            beta_no_hyper, alpha_no_hyper, sigma, b, q, d, rho, eta, gamma, lam, c1, c2, 0.0, seed
        )

        S_no_hyper = y_no_hyper[0]
        I_no_hyper = y_no_hyper[1]
        H_no_hyper = y_no_hyper[2]

        N_no_hyper = S_no_hyper + I_no_hyper + H_no_hyper

        #Calculate the relative rates of deaths and affects on population sizes
        deaths_1 = ((d*(S + I + H) + alpha*I + lam*alpha*H)/(S+I+H))/((d*(S_no_hyper + I_no_hyper) + alpha_no_hyper*I_no_hyper)/(S_no_hyper + I_no_hyper))

        if I + H == 0:
            deaths_2 = 0
            pop_virulences.append(0)
        else:
            deaths_2 = ((alpha*(I + lam*H))/(I+H))/alpha_no_hyper
            pop_virulences.append(alpha*((I + lam*H)/(I + H)))

        death_measure_1.append(deaths_1)
        death_measure_2.append(deaths_2)
        
        relative_pop_size.append(N/N_no_hyper)

        if I_no_hyper + H_no_hyper == 0 and I + H != 0:
            relative_infected_proportion.append(10)
        elif I_no_hyper + H_no_hyper == 0 and I + H == 0:
            relative_infected_proportion.append(0)
        else:
            relative_infected_proportion.append(((I + H)/N)/((I_no_hyper + H_no_hyper)/N_no_hyper))
            
    return death_measure_1, death_measure_2, relative_pop_size, relative_infected_proportion, host_densities, infected_densities, hyperparasite_densities, pop_virulences

def calculate_covergence_stability(sol, attractors, beta_max, alpha_max, sigma_max, sigma, b, q, d, rho, eta, gamma, lam, c1, c2, hyper, seed, ):
    
    convergence_stabilities = []
    
    derivative_distance = 1e-1
    
    for alpha_t in attractors:
        alpha = alpha_t - derivative_distance/2
        alpha_2 = alpha_t + derivative_distance/2
        
        beta = zf.beta_func(beta_max, alpha, alpha_max, sigma, sigma_max, c1, c2)
        
        beta_2 = zf.beta_func(beta_max, alpha_2, alpha_max, sigma, sigma_max, c1, c2)
        dbetadalpha = zf.dbetadalpha_func(
            beta_max, alpha, alpha_max, sigma, sigma_max, c1, c2
        )
        
        dbetadalpha_2 = zf.dbetadalpha_func(
            beta_max, alpha_2, alpha_max, sigma, sigma_max, c1, c2
        )

        #Solve the system to ecological steady state
        y = sol.eco_steady_state(
            beta, alpha, sigma, b, q, d, rho, eta, gamma, lam, c1, c2, hyper, seed
        )
        
        y_2 = sol.eco_steady_state(
            beta_2, alpha_2, sigma, b, q, d, rho, eta, gamma, lam, c1, c2, hyper, seed
        )
        
        S = y[0]
        I = y[1]
        H = y[2]
        
        S_2 = y_2[0]
        I_2 = y_2[1]
        H_2 = y_2[2]

        #Determine which version of the gradient we need. If the parasite is extinct just add an
        #arbitrary value
        if (I == 0 and H == 0):
            alpha_grad = 1*(alpha <= alpha_max/2) - 1*(alpha> alpha_max/2)
        elif (H == 0):
            alpha_grad = dbetadalpha*S/(d + alpha + gamma) - (beta*S)/((d + alpha + gamma)**2)
        elif (I == 0):
            alpha_grad = dbetadalpha*S/(d + lam*alpha + gamma) - (beta*S)/((d + lam*alpha + gamma)**2)
        else:
            alpha_grad = ss.fitness_gradient_alpha(
                beta, alpha, sigma, dbetadalpha, H, S, eta, lam, gamma, d, rho
            )

        if (I_2 == 0 and H_2 == 0):
            alpha_grad_res = 1*(alpha <= alpha_max/2) - 1*(alpha> alpha_max/2)
        elif (H_2 == 0):
            alpha_grad_res = dbetadalpha*S_2/(d + alpha + gamma) - (beta*S_2)/((d + alpha + gamma)**2)
        elif (I_2 == 0):
            alpha_grad_res = dbetadalpha*S_2/(d + lam*alpha + gamma) - (beta*S_2)/((d + lam*alpha + gamma)**2)
        else:
            alpha_grad_res = ss.fitness_gradient_alpha(
                beta, alpha, sigma, dbetadalpha, H_2, S_2, eta, lam, gamma, d, rho
            )
        
        if (I == 0 and H == 0):
            alpha_grad_mut = 1*(alpha <= alpha_max/2) - 1*(alpha> alpha_max/2)
        elif (H == 0):
            alpha_grad_mut = dbetadalpha_2*S/(d + alpha_2 + gamma) - (beta_2*S)/((d + alpha_2 + gamma)**2)
        elif (I == 0):
            alpha_grad_mut = dbetadalpha_2*S/(d + lam*alpha_2 + gamma) - (beta_2*S)/((d + lam*alpha_2 + gamma)**2)
        else:
            alpha_grad_mut = ss.fitness_gradient_alpha(
                beta_2, alpha_2, sigma, dbetadalpha_2, H, S, eta, lam, gamma, d, rho
            )
        dfitdresident = (alpha_grad_res - alpha_grad)/derivative_distance
        dfitdmut = (alpha_grad_mut - alpha_grad)/derivative_distance
        
        convergence_stabilities.append((dfitdmut + dfitdresident))
#         convergence_stabilities.append((-1))
    return convergence_stabilities
# Parameters for the system we wish to investigate
rhos = [0.1, 0.5, 0.9]
# rhos = [0.0, 1.0]
# rhos = [1.0]
# rhos = [0.1, 0.5, 0.9, 1.0]
lams = [0.5, 1, 2.0]
# lams = [0.0, 0.5, 1.0, 1.5, 2.0]
# rhos = [1.0]
b = 2.0
q = 0.1
d = 0.1
gamma = 0.1
# gamma = 1.0
hyper = 1.0
c1 = 0.75
c2 = 2.25
b_base = 1
q_base = 1

alpha_max = 5
sigma_max = 0.4
beta_max = 5

sigma_min = 0.0
sigma_max_range = sigma_max

sigma = sigma_max

iter_max = 3
res_scalar = 4
depth_max = iter_max
resolution = 100
param_res = 200

seed = 10000

#Initiating the CPP solver
sol = runner.PySolver()

#Creating dictionaries we will use to easily plot our singular strategies later
plotting_dict_attractors = {}
plotting_dict_attractors_no_hyper = {}

plotting_dict_repellers = {}
plotting_dict_repellers_no_hyper = {}

plotting_dict_hyper = {}
plotting_dict_no_hyper = {}
plotting_dict_deaths_1 = {}
plotting_dict_deaths_2 = {}
plotting_dict_pops = {}
plotting_dict_inf = {}
plotting_dict_hosts = {}
plotting_dict_paras = {}
plotting_dict_hypers = {}
plotting_dict_hyper_prop = {}
plotting_dict_pop_virulences = {}
#Creating lists to add plots to for the legend
labels_base = []
labels = []
lines1 = []
lines2 = []

#Here we create a random string of 4 words which we use to create a unique folder
inds = np.random.randint(len(lines_cleaned),size = 4)

filekey = ""
for ind in inds:
    filekey += lines_cleaned[ind] + "_"

filekey += "alpha_figs"

filekey = filekey.replace("'", "").replace(".", "")

output_folder = "../outputs/single_evo/" + filekey

os.makedirs(output_folder)

#We add a text file containing all of the parameters to ensure easy reproduction
with open(f"{output_folder}/parameters.txt", "w") as f:
    f.write("The parameters for the system where:\n")
    f.write(f"b = {b}\n")
    f.write(f"q = {q}\n")
    f.write(f"d = {d}\n")
    for i, rho in enumerate(rhos):
        f.write(f"rho_{i} = {rho}\n")
    for i, lam in enumerate(lams):
        f.write(f"lam_{i} = {lam}\n")
    f.write(f"gamma = {gamma}\n")
    f.write(f"eta = varying\n")
    f.write(f"c1 = {c1}\n")
    f.write(f"c2 = {c2}\n")
    f.write(f"beta_max = {beta_max}\n")
    f.write(f"alpha_max = {alpha_max}\n")
    f.write(f"sigma_max = {sigma_max}\n")
    f.write(f"resolution = {resolution}\n")
    f.write(f"param_res = {param_res}\n")
    f.write(f"iter_max = {iter_max}\n")
    f.write(f"res_scalar = {res_scalar}\n")
    f.write(f"sigma = {sigma}\n")

f.close()

#We create two base figures, one for the evolutionary consequences
fig1 = plt.figure(figsize = (40, 44))
gs1 = fig1.add_gridspec(4,len(lams))
ax1 = []

#And one for the ecological consequences
# fig2 = plt.figure(figsize = (40, 33))
# gs2 = fig2.add_gridspec(3,len(lams))
# gs2 = fig2.add_gridspec(3,len(lams))
#ax2 = []

#Flavour text to add to the readibility of the plots
texts = [["(A)", "(B)", "(C)"],["(D)", "(E)", "(F)"], ["(G)","(H)","(I)"], ["(J)", "(K)", "(L)"]]

titles = ["Hypovirulence ", "No direct effect on virulence ", "Hypervirulence "]
# titles = ["Hypovirulence ", "Hypovirulence ", "No effect on virulence ", "Hypervirulence ", "Hypervirulence "]

#Structuring the figures so we can access subplots
for i in range(4):
    temp_ax = []
    for j in range(len(lams)):
        temp_ax.append(fig1.add_subplot(gs1[i,j]))
    ax1.append(temp_ax)
    
# for i in range(3):
#     temp_ax = []
#     for j in range(len(lams)):
#         temp_ax.append(fig2.add_subplot(gs2[i,j]))
#     #ax2.append(temp_ax)
    
# for i in range(3):
#     temp_ax = []
#     for j in range(len(lams)):
#         temp_ax.append(fig2.add_subplot(gs2[i,j]))
    #ax2.append(temp_ax)
    
    
sigma_res = [sigma_max*(i/100) for i in range(101)]

for i, val in enumerate(sigma_res):
    if val >= sigma:
        sigma_init = i
        break
        
alpha_init = 10
#We initialise the system with the hyperparasites absent to find the natural ESS of the parasite
sol.alpha_ad_dyn(beta_max, alpha_max, sigma_max, b, q, d, rho, 0.0, gamma, 0.0, c1, c2, 0.0, seed, alpha_init, sigma_init, H_density = 0.0)
df = pd.read_csv(f"../data/alpha_evo/data_set{seed}.csv")

evo_steps = df["Evolutionary_step"].values
dft = df[df["Evolutionary_step"]==evo_steps[-1]]

pops = []
for val in dft["Trait_index_1"].values:
    df_pop = dft[dft["Trait_index_1"]==val][["Density_of_parasite"]]
    pops.append((df_pop["Density_of_parasite"].iloc[0])/(sum(dft["Density_of_parasite"].values)))
    
#Calculating the average trait value
alpha_weights = []
for i, val in enumerate(pops):
    alpha_weights.append(val*dft["Trait_index_1"].values[i])
alpha_mean = sum(alpha_weights)

#Rounding this to the closest integer to use as our initial conditions
alpha_init = round(alpha_mean)

alpha_mean_base = alpha_init*alpha_max/100

host_density = dft["Density_of_Hosts"].iloc[0]
para_density = sum(dft["Density_of_parasite"].values)
hyper_density = 0.1

#Here we loop through our different values of eta, rho and lambda. The lam_tracker allows us to access
#the correct figure at the same time
for lam_tracker, lam in enumerate(lams):
    for rho in rhos:
        

#         iter_max = 2
#         res_scalar = 4
#         depth_max = iter_max
#         resolution = 100

        beta_max = beta_max
        alpha_max_param = alpha_max 
        sigma_max_param = sigma_max

        param_res = 200

        eta_min = 0.0
        eta_max = 2.0
        etas = [eta_min + (eta_max - eta_min)*i/param_res for i in range(param_res + 1)]

        
        #Here we create our temporary lists to store the values of the singular strategies
        plotting_attractors_hyper = []
        plotting_attractors_no_hyper = []

        plotting_repellers_hyper = []
        plotting_repellers_no_hyper = []

        plotting_etas_attractors_hyper = []
        plotting_etas_attractors_no_hyper = []

        plotting_etas_repellers_hyper = []
        plotting_etas_repellers_no_hyper = []

        for eta_t in etas:
            eta = round(eta_t,4)
            print(f"On run {eta} on rho {rho}, and lam {lam}  ", end = "\r")

            alpha_max_res = alpha_max
            alpha_min_res = 0.0

            alpha_grad_vec, alpha_val = calculate_alpha_gradient_vector(sol, resolution, beta_max, alpha_max, alpha_min_res, alpha_max_res, sigma_max, sigma, b, q, d, rho, eta, gamma, lam, c1, c2, hyper, seed)


            alpha_grad_no_hyper_vec, alpha_val = calculate_alpha_gradient_vector(sol, resolution, beta_max, alpha_max, alpha_min_res, alpha_max_res, sigma_max, sigma, b, q, d, rho, eta, gamma, lam, c1, c2, 0.0, seed)

            #Calculating the singular strategies when the hyperparasite is present and absent
            attractors_hyper, repellers_hyper = recusive_sing_strat_function(sol, 0, iter_max, resolution, beta_max, alpha_max, alpha_min_res, alpha_max_res, sigma_max, sigma, b, q, d, rho, eta, gamma, lam, c1, c2, hyper, seed)

            attractors_no_hyper, repellers_no_hyper = recusive_sing_strat_function(sol, 0, iter_max, resolution, beta_max, alpha_max, alpha_min_res, alpha_max_res, sigma_max, sigma, b, q, d, rho, eta, gamma, lam, c1, c2, 0.0, seed)

            #Storing these values in our lists from before
            for val in attractors_hyper:
                plotting_etas_attractors_hyper.append(eta)
                plotting_attractors_hyper.append(val)

            for val in attractors_no_hyper:
                plotting_etas_attractors_no_hyper.append(eta)
                plotting_attractors_no_hyper.append(val)

            for val in repellers_hyper:
                plotting_etas_repellers_hyper.append(eta)
                plotting_repellers_hyper.append(val)

            for val in repellers_no_hyper:
                plotting_etas_repellers_no_hyper.append(eta)
                plotting_repellers_no_hyper.append(val)

            for i, val in enumerate(alpha_val):
                if val >= attractors_no_hyper[0]:
                    break

        #Adding these lists to our stored dictionaries
        plotting_dict_hyper[rho] = {}
        plotting_dict_no_hyper[rho] = {}

        plotting_dict_hyper[rho]["a"] = plotting_etas_attractors_hyper
        plotting_dict_no_hyper[rho]["a"] = plotting_etas_attractors_no_hyper

        plotting_dict_hyper[rho]["r"] = plotting_etas_repellers_hyper
        plotting_dict_no_hyper[rho]["r"] = plotting_etas_repellers_no_hyper

        plotting_dict_attractors[rho] = plotting_attractors_hyper
        plotting_dict_attractors_no_hyper[rho] = plotting_attractors_no_hyper

        plotting_dict_repellers[rho] = plotting_repellers_hyper
        plotting_dict_repellers_no_hyper[rho] = plotting_repellers_no_hyper
    
#         #Calculating the ecological consequences of a parasite with those trait values
#         death_measure_1_attractor, death_measure_2_attractor, relative_pop_size_attractor, relative_infected_proportion_attractor, host_density_attractor, para_density_attractor, hyper_density_attractor, pop_virulence_attractor = produce_suplimentary_data(
#             sol,
#             plotting_attractors_hyper,
#             plotting_etas_attractors_hyper,
#             plotting_attractors_no_hyper,
#             plotting_etas_attractors_no_hyper,
#             beta_max,
#             alpha_max,
#             sigma_max,
#             sigma,
#             c1,
#             c2,
#             b,
#             q,
#             d,
#             rho,
#             gamma,
#             lam,
#             hyper,
#             seed,
#           )

#         death_measure_1_repeller, death_measure_2_repeller, relative_pop_size_repeller, relative_infected_proportion_repeller, host_density_repeller, para_density_repeller, hyper_density_repeller, pop_virulence_repeller = produce_suplimentary_data(
#             sol,
#             plotting_repellers_hyper,
#             plotting_etas_repellers_hyper,
#             plotting_attractors_no_hyper,
#             plotting_etas_attractors_no_hyper,
#             beta_max,
#             alpha_max,
#             sigma_max,
#             sigma,
#             c1,
#             c2,
#             b,
#             q,
#             d,
#             rho,
#             gamma,
#             lam,
#             hyper,
#             seed,
#           )

        #Storing that information
        plotting_dict_deaths_1[rho] = {}
        plotting_dict_deaths_2[rho] = {}
        plotting_dict_pops[rho] = {}
        plotting_dict_inf[rho] = {}

#         plotting_dict_deaths_1[rho]["a"] = death_measure_1_attractor
#         plotting_dict_deaths_2[rho]["a"] = death_measure_2_attractor
#         plotting_dict_pops[rho]["a"] = relative_pop_size_attractor
#         plotting_dict_inf[rho]["a"] = relative_infected_proportion_attractor

#         plotting_dict_deaths_1[rho]["r"] = death_measure_1_repeller
#         plotting_dict_deaths_2[rho]["r"] = death_measure_2_repeller
#         plotting_dict_pops[rho]["r"] = relative_pop_size_repeller
#         plotting_dict_inf[rho]["r"] = relative_infected_proportion_repeller

        plotting_dict_hosts[rho] = {}
        plotting_dict_paras[rho] = {}
        plotting_dict_hypers[rho] = {}
        plotting_dict_hyper_prop[rho] = {}
        plotting_dict_pop_virulences[rho] = {}
        
#         plotting_dict_hosts[rho]["a"] = host_density_attractor
#         plotting_dict_paras[rho]["a"] = para_density_attractor
#         plotting_dict_hypers[rho]["a"] = hyper_density_attractor

#         plotting_dict_hosts[rho]["r"] = host_density_repeller
#         plotting_dict_paras[rho]["r"] = para_density_repeller
#         plotting_dict_hypers[rho]["r"] = hyper_density_repeller


                        

    xlabel = r"Hyperparasite Transmission Modifier, $\eta$"
    colours = ["tab:blue", "tab:orange", "tab:green", "tab:red", "tab:gray"]

    
    eta_mins = {}
    eta_maxs = {}
    for rho in rhos:
        temp_etas = plotting_dict_hyper[rho]["r"]
        if len(temp_etas) > 0:
            eta_mins[rho] = min(temp_etas)
            eta_maxs[rho] = max(temp_etas)

    #As we have a hystersis effect within our plots it can be difficult to visualise smoothly
    alpha_lower_plotting = {}
    alpha_upper_plotting = {}
    alpha_attractors = {}
    eta_attractors = {}
    for rho in rhos:
        temp_alphas = plotting_dict_repellers[rho]
        eta_attractors[rho] = {}
        alpha_attractors[rho] = {}
        eta_attractors[rho]['1'] = []
        eta_attractors[rho]['2'] = []
        alpha_attractors[rho]['1'] = []
        alpha_attractors[rho]['2'] = []
        
        #This section checks if there are any repellers, meaning we have the hysteresis effect present.
        if len(temp_alphas) > 0:
            alpha_lower_plotting[rho] = min(temp_alphas)
            alpha_upper_plotting[rho] = max(temp_alphas)
            
            flag = True
            paired_list = []
            for i, val in enumerate(plotting_dict_attractors[rho]):
                paired_list.append([val, plotting_dict_hyper[rho]["a"][i]])
            sorted_attractors = sorted(paired_list, key = lambda x: x[0], reverse = True)
            temp_attractors_1 = []
            temp_attractors_2 = []
            temp_etas_1 = []
            temp_etas_2 = []
            
            #This section checks to see if we have jumped from one branch of the hysteresis affect to the other
            for i, val in enumerate(sorted_attractors[:-1]):
                if flag and abs(sorted_attractors[i+1][0] - val[0]) < 0.1:
                    temp_attractors_1.append(val[0])
                    temp_etas_1.append(val[1])
                elif flag:
                    flag = False
                    temp_attractors_1.append(val[0])
                    temp_etas_1.append(val[1])
                else:
                    temp_attractors_2.append(val[0])
                    temp_etas_2.append(val[1])
                    
            temp_attractors_2.append(sorted_attractors[-1][0])
            temp_etas_2.append(sorted_attractors[-1][1])
            
            paired_list_1 = []
            for i, val in enumerate(temp_etas_1):
                paired_list_1.append([val, temp_attractors_1[i]])
             
            paired_list_2 = []
            for i, val in enumerate(temp_etas_2):
                paired_list_2.append([val, temp_attractors_2[i]])
                
            #We sort the values to be in terms of eta again once they have been properly seperated
            sorted_list_1 = sorted(paired_list_1, key = lambda x: x[0])
            sorted_list_2 = sorted(paired_list_2, key = lambda x: x[0])
            
            temp_attractors_1 = [val[1] for val in sorted_list_1]
            temp_attractors_2 = [val[1] for val in sorted_list_2]
            
            temp_etas_1 = [val[0] for val in sorted_list_1]
            temp_etas_2 = [val[0] for val in sorted_list_2]
            
            alpha_attractors[rho]['1'] = temp_attractors_1
            alpha_attractors[rho]['2'] = temp_attractors_2
            eta_attractors[rho]['1'] = temp_etas_1
            eta_attractors[rho]['2'] = temp_etas_2
            
            temp_convergences_1 = calculate_covergence_stability(sol, temp_attractors_1, beta_max, alpha_max, sigma_max, sigma, b, q, d, rho, eta, gamma, lam, c1, c2, hyper, seed)
            temp_convergences_2 = calculate_covergence_stability(sol, temp_attractors_2, beta_max, alpha_max, sigma_max, sigma, b, q, d, rho, eta, gamma, lam, c1, c2, hyper, seed)
            
#             for i, val in enumerate(temp_convergences_1):
#                 if val > 0:
#                     print(val, temp_etas_1[i], temp_attractors_1[i], seed, "type 1")
#                     j = 0
#                     while ((j/100)*alpha_max < temp_attractors_1[i]):
#                         j += 1
#                     j += -1
#                     sol.alpha_ad_dyn(beta_max, alpha_max, sigma_max, b, q, d, rho, temp_etas_1[i], gamma, lam, c1, c2, hyper, seed, j, sigma_init, S_density = host_density, I_density = para_density, H_density = hyper_density)
#                     seed += 1
                    
#             for i, val in enumerate(temp_convergences_2):
#                 if val > 0:
#                     print(val, temp_etas_2[i], temp_attractors_2[i], seed, "type 2")
#                     j = 0
#                     while ((j/100)*alpha_max < temp_attractors_2[i]):
#                         j += 1
#                     j += -1
#                     sol.alpha_ad_dyn(beta_max, alpha_max, sigma_max, b, q, d, rho, temp_etas_2[i], gamma, lam, c1, c2, hyper, seed, j, sigma_init, S_density = host_density, I_density = para_density, H_density = hyper_density)
#                     seed += 1
                    
            #We calculate the ecological consequences of the introduction of a hyperparasite
            plotting_dict_deaths_1[rho]["1"], plotting_dict_deaths_2[rho]["1"], plotting_dict_pops[rho]["1"], plotting_dict_inf[rho]["1"], host_density_attractor, para_density_attractor, plotting_dict_hypers[rho]["1"], plotting_dict_pop_virulences[rho]["1"] = produce_suplimentary_data(
            sol,
            temp_attractors_1,
            temp_etas_1,
            plotting_attractors_no_hyper,
            plotting_etas_attractors_no_hyper,
            beta_max,
            alpha_max,
            sigma_max,
            sigma,
            c1,
            c2,
            b,
            q,
            d,
            rho,
            gamma,
            lam,
            hyper,
            seed,
          )
            temp_list_hyper_prop = []
            for i, val in enumerate(para_density_attractor):
                H = plotting_dict_hypers[rho]["1"][i]
                temp_list_hyper_prop.append(H/(H + val))
            plotting_dict_hyper_prop[rho]["1"] = temp_list_hyper_prop
            
            plotting_dict_deaths_1[rho]["2"], plotting_dict_deaths_2[rho]["2"], plotting_dict_pops[rho]["2"], plotting_dict_inf[rho]["2"], host_density_attractor, para_density_attractor, plotting_dict_hypers[rho]["2"], plotting_dict_pop_virulences[rho]["2"] = produce_suplimentary_data(
            sol,
            temp_attractors_2,
            temp_etas_2,
            plotting_attractors_no_hyper,
            plotting_etas_attractors_no_hyper,
            beta_max,
            alpha_max,
            sigma_max,
            sigma,
            c1,
            c2,
            b,
            q,
            d,
            rho,
            gamma,
            lam,
            hyper,
            seed,
          )
            temp_list_hyper_prop = []
            for i, val in enumerate(para_density_attractor):
                H = plotting_dict_hypers[rho]["2"][i]
                temp_list_hyper_prop.append(H/(H + val))
            plotting_dict_hyper_prop[rho]["2"] = temp_list_hyper_prop
        else:
            alpha_attractors[rho]['1'] = plotting_dict_attractors[rho]
            eta_attractors[rho]['1'] = plotting_dict_hyper[rho]["a"]

            plotting_dict_deaths_1[rho]["1"], plotting_dict_deaths_2[rho]["1"], plotting_dict_pops[rho]["1"], plotting_dict_inf[rho]["1"], host_density_attractor, para_density_attractor, plotting_dict_hypers[rho]["1"], plotting_dict_pop_virulences[rho]["1"] = produce_suplimentary_data(
            sol,
            plotting_dict_attractors[rho],
            plotting_dict_hyper[rho]["a"],
            plotting_attractors_no_hyper,
            plotting_etas_attractors_no_hyper,
            beta_max,
            alpha_max,
            sigma_max,
            sigma,
            c1,
            c2,
            b,
            q,
            d,
            rho,
            gamma,
            lam,
            hyper,
            seed,
          )
            temp_list_hyper_prop = []
            for i, val in enumerate(para_density_attractor):
                H = plotting_dict_hypers[rho]["1"][i]
                temp_list_hyper_prop.append(H/(H + val))
            plotting_dict_hyper_prop[rho]["1"] = temp_list_hyper_prop
            
            plotting_dict_deaths_1[rho]["2"] = []
            plotting_dict_deaths_2[rho]["2"] = []
            plotting_dict_pops[rho]["2"] = []
            plotting_dict_inf[rho]["2"] = []
            plotting_dict_hypers[rho]["2"] = []
            plotting_dict_pop_virulences[rho]["2"] = []
            plotting_dict_hyper_prop[rho]["2"] = []
    #Here we do 3 dummy plots to create the legend we will use later
    for i, rho in enumerate(rhos):
        if lam_tracker == 0:
            labels_base.append(fr"$\rho$ = {rho}")
            labels.append(labels_base[i])
            l1 = ax1[0][lam_tracker].plot(eta_attractors[rho]['1'], alpha_attractors[rho]['1'], c = f"{colours[i]}")
            lines1.append(l1)
    
#     for i, rho in enumerate(rhos):
#         if lam_tracker == 0:
#             l1 = ax1[3][lam_tracker].plot(eta_attractors[rho]['1'], plotting_dict_deaths_2[rho]["1"], c = f"{colours[i]}")
#             lines2.append(l1)        
    
    #First we plot the evolved levels of virulence
    for i, rho in enumerate(rhos):
        if lam_tracker == 0:
            
            ax1[0][lam_tracker].plot(eta_attractors[rho]['1'], alpha_attractors[rho]['1'], c = f"{colours[i]}")
            
            ax1[0][lam_tracker].plot(eta_attractors[rho]['2'], alpha_attractors[rho]['2'], c = f"{colours[i]}")
            
            #This marks the begining and end of the first section of the plot
            ax1[0][lam_tracker].scatter(eta_attractors[rho]['1'][-1], alpha_attractors[rho]['1'][-1], edgecolors = f"{colours[i]}", marker = "o", facecolors= f"{colours[i]}", s = 200)
            ax1[0][lam_tracker].scatter(eta_attractors[rho]['1'][0], alpha_attractors[rho]['1'][0], edgecolors = f"{colours[i]}", marker = "o", facecolors="none", s = 200)
            
            #If there is a hysteresis effect this will work, else we'll except straight through
            try:
                ax1[0][lam_tracker].scatter(eta_attractors[rho]['2'][-1], alpha_attractors[rho]['2'][-1], edgecolors = f"{colours[i]}", marker = "s", facecolors= f"{colours[i]}", s = 200)
                ax1[0][lam_tracker].scatter(eta_attractors[rho]['2'][0], alpha_attractors[rho]['2'][0], edgecolors = f"{colours[i]}", marker = "s", facecolors="none", s = 200)
            except:
                pass
            
            #This checks if we need to plot the repellers and then links the repellers to the start and end of the attractors
            #as appropriate
            if len(plotting_dict_hyper[rho]["r"]) > 0:
                repeller_etas = [eta_attractors[rho]["2"][0]]
                for val in plotting_dict_hyper[rho]["r"]:
                    repeller_etas.append(val)
                repeller_etas.append(eta_attractors[rho]["1"][-1])

                repeller_alphas = [alpha_attractors[rho]['2'][0]]
                for val in plotting_dict_repellers[rho]:
                    repeller_alphas.append(val)

                repeller_alphas.append(alpha_attractors[rho]["1"][-1])
            else:
                repeller_etas = plotting_dict_hyper[rho]["r"]
                repeller_alphas = plotting_dict_repellers[rho]
                
            ax1[0][lam_tracker].plot(repeller_etas, repeller_alphas,  c = f"{colours[i]}", linestyle='dashed')
        else:
            ax1[0][lam_tracker].plot(eta_attractors[rho]['1'], alpha_attractors[rho]['1'], c = f"{colours[i]}")
            ax1[0][lam_tracker].plot(eta_attractors[rho]['2'], alpha_attractors[rho]['2'], c = f"{colours[i]}")
            ax1[0][lam_tracker].scatter(eta_attractors[rho]['1'][0], alpha_attractors[rho]['1'][0], edgecolors = f"{colours[i]}", marker = "o", facecolors="none", s = 200)
            ax1[0][lam_tracker].scatter(eta_attractors[rho]['1'][-1], alpha_attractors[rho]['1'][-1], edgecolors = f"{colours[i]}", marker = "o", facecolors= f"{colours[i]}", s = 200)
            try:
                ax1[0][lam_tracker].scatter(eta_attractors[rho]['2'][0], alpha_attractors[rho]['2'][0], edgecolors = f"{colours[i]}", marker = "s", facecolors = "none", s = 200)
                ax1[0][lam_tracker].scatter(eta_attractors[rho]['2'][-1], alpha_attractors[rho]['2'][-1], edgecolors = f"{colours[i]}", marker = "s", facecolors= f"{colours[i]}", s = 200)
            except:
                pass
            if len(plotting_dict_hyper[rho]["r"]) > 0:
                repeller_etas = [eta_attractors[rho]["2"][0]]
                for val in plotting_dict_hyper[rho]["r"]:
                    repeller_etas.append(val)
                repeller_etas.append(eta_attractors[rho]["1"][-1])

                repeller_alphas = [alpha_attractors[rho]['2'][0]]
                for val in plotting_dict_repellers[rho]:
                    repeller_alphas.append(val)

                repeller_alphas.append(alpha_attractors[rho]["1"][-1])
            else:
                repeller_etas = plotting_dict_hyper[rho]["r"]
                repeller_alphas = plotting_dict_repellers[rho]
                
            ax1[0][lam_tracker].plot(repeller_etas, repeller_alphas,  c = f"{colours[i]}", linestyle='dashed')


    ax1[0][lam_tracker].plot(plotting_dict_no_hyper[rho]["a"],plotting_dict_attractors_no_hyper[rho], c = "k")
    ax1[0][lam_tracker].plot(plotting_dict_no_hyper[rho]["r"],plotting_dict_repellers_no_hyper[rho], c = "k")
    ax1[0][lam_tracker].text(0.03, 1.01,texts[0][lam_tracker], transform=ax1[0][lam_tracker].transAxes, fontsize = 30)
    
    ax1[0][lam_tracker].set_xlabel("", fontsize = 0)
#     ax1[0][lam_tracker].set_ylim([0, alpha_max_res])
    ax1[0][lam_tracker].set_ylim([0, 2.5])
    ax1[0][lam_tracker].tick_params(axis='both', which='major', labelsize=34)
    if lam_tracker == 0:
        ax1[0][lam_tracker].set_ylabel(r"Intrinsic virulence, $\alpha^*$", fontsize = 34)
    ax1[0][lam_tracker].set_title(fr"{titles[lam_tracker]}($\lambda$ = {lam})", fontsize = 28)
    
    #First we plot the evolved levels of virulence
#     for i, rho in enumerate(rhos):
#         if lam_tracker == 0:
            
#             #ax2[2][lam_tracker].plot(eta_attractors[rho]['1'], alpha_attractors[rho]['1'], c = f"{colours[i]}")
            
#             #ax2[2][lam_tracker].plot(eta_attractors[rho]['2'], alpha_attractors[rho]['2'], c = f"{colours[i]}")
            
#             #This marks the begining and end of the first section of the plot
#             #ax2[2][lam_tracker].scatter(eta_attractors[rho]['1'][-1], alpha_attractors[rho]['1'][-1], edgecolors = f"{colours[i]}", marker = "o", facecolors= f"{colours[i]}", s = 200)
#             #ax2[2][lam_tracker].scatter(eta_attractors[rho]['1'][0], alpha_attractors[rho]['1'][0], edgecolors = f"{colours[i]}", marker = "o", facecolors="none", s = 200)
            
#             #If there is a hysteresis effect this will work, else we'll except straight through
# #             try:
#                 #ax2[2][lam_tracker].scatter(eta_attractors[rho]['2'][-1], alpha_attractors[rho]['2'][-1], edgecolors = f"{colours[i]}", marker = "s", facecolors= f"{colours[i]}", s = 200)
#                 #ax2[2][lam_tracker].scatter(eta_attractors[rho]['2'][0], alpha_attractors[rho]['2'][0], edgecolors = f"{colours[i]}", marker = "s", facecolors="none", s = 200)
# #             except:
# #                 pass
            
#             #This checks if we need to plot the repellers and then links the repellers to the start and end of the attractors
#             #as appropriate
#             if len(plotting_dict_hyper[rho]["r"]) > 0:
#                 repeller_etas = [eta_attractors[rho]["2"][0]]
#                 for val in plotting_dict_hyper[rho]["r"]:
#                     repeller_etas.append(val)
#                 repeller_etas.append(eta_attractors[rho]["1"][-1])

#                 repeller_alphas = [alpha_attractors[rho]['2'][0]]
#                 for val in plotting_dict_repellers[rho]:
#                     repeller_alphas.append(val)

#                 repeller_alphas.append(alpha_attractors[rho]["1"][-1])
#             else:
#                 repeller_etas = plotting_dict_hyper[rho]["r"]
#                 repeller_alphas = plotting_dict_repellers[rho]
                
#             #ax2[2][lam_tracker].plot(repeller_etas, repeller_alphas,  c = f"{colours[i]}", linestyle='dashed')
#         else:
#             #ax2[2][lam_tracker].plot(eta_attractors[rho]['1'], alpha_attractors[rho]['1'], c = f"{colours[i]}")
#             #ax2[2][lam_tracker].plot(eta_attractors[rho]['2'], alpha_attractors[rho]['2'], c = f"{colours[i]}")
#             #ax2[2][lam_tracker].scatter(eta_attractors[rho]['1'][0], alpha_attractors[rho]['1'][0], edgecolors = f"{colours[i]}", marker = "o", facecolors="none", s = 200)
#             #ax2[2][lam_tracker].scatter(eta_attractors[rho]['1'][-1], alpha_attractors[rho]['1'][-1], edgecolors = f"{colours[i]}", marker = "o", facecolors= f"{colours[i]}", s = 200)
#             try:
#                 #ax2[2][lam_tracker].scatter(eta_attractors[rho]['2'][0], alpha_attractors[rho]['2'][0], edgecolors = f"{colours[i]}", marker = "s", facecolors = "none", s = 200)
#                 #ax2[2][lam_tracker].scatter(eta_attractors[rho]['2'][-1], alpha_attractors[rho]['2'][-1], edgecolors = f"{colours[i]}", marker = "s", facecolors= f"{colours[i]}", s = 200)
#             except:
#                 pass
#             if len(plotting_dict_hyper[rho]["r"]) > 0:
#                 repeller_etas = [eta_attractors[rho]["2"][0]]
#                 for val in plotting_dict_hyper[rho]["r"]:
#                     repeller_etas.append(val)
#                 repeller_etas.append(eta_attractors[rho]["1"][-1])

#                 repeller_alphas = [alpha_attractors[rho]['2'][0]]
#                 for val in plotting_dict_repellers[rho]:
#                     repeller_alphas.append(val)

#                 repeller_alphas.append(alpha_attractors[rho]["1"][-1])
#             else:
#                 repeller_etas = plotting_dict_hyper[rho]["r"]
#                 repeller_alphas = plotting_dict_repellers[rho]
                
            #ax2[2][lam_tracker].plot(repeller_etas, repeller_alphas,  c = f"{colours[i]}", linestyle='dashed')


    #ax2[2][lam_tracker].plot(plotting_dict_no_hyper[rho]["a"],plotting_dict_attractors_no_hyper[rho], c = "k")
    #ax2[2][lam_tracker].plot(plotting_dict_no_hyper[rho]["r"],plotting_dict_repellers_no_hyper[rho], c = "k")
#     #ax2[2][lam_tracker].text(0.03, 1.01,texts[0][lam_tracker], transform=ax1[0][lam_tracker].transAxes, fontsize = 30)
    
    #ax2[2][lam_tracker].set_xlabel("", fontsize = 0)
    #ax2[2][lam_tracker].set_ylim([0, 1.1*alpha_max_res])
    #ax2[2][lam_tracker].tick_params(axis='both', which='major', labelsize=34)
#     if lam_tracker == 0:
        #ax2[2][lam_tracker].set_ylabel(r"Evolved virulence, $\alpha$", fontsize = 34)
#     for i, rho in enumerate(rhos):
#         ax1[1][lam_tracker].plot(eta_attractors[rho]['1'], plotting_dict_pop_virulences[rho]["1"], c = f"{colours[i]}")
#         ax1[1][lam_tracker].plot(eta_attractors[rho]['2'], plotting_dict_pop_virulences[rho]["2"], c = f"{colours[i]}")
#         ax1[1][lam_tracker].scatter(eta_attractors[rho]['1'][0], plotting_dict_pop_virulences[rho]['1'][0], edgecolors = f"{colours[i]}", marker = "o", facecolors="none", s = 200)
#         ax1[1][lam_tracker].scatter(eta_attractors[rho]['1'][-1], plotting_dict_pop_virulences[rho]['1'][-1], edgecolors = f"{colours[i]}", marker = "o", facecolors= f"{colours[i]}", s = 200)
#         try:
#             ax1[1][lam_tracker].scatter(eta_attractors[rho]['2'][0], plotting_dict_pop_virulences[rho]['2'][0], edgecolors = f"{colours[i]}", marker = "s", facecolors= "none", s = 200)
#             ax1[1][lam_tracker].scatter(eta_attractors[rho]['2'][-1], plotting_dict_pop_virulences[rho]['2'][-1], edgecolors = f"{colours[i]}", marker = "s", facecolors= f"{colours[i]}", s = 200)
#         except:
#             pass
#     if lam_tracker == 1:
#         ax1[1][lam_tracker].set_xlabel(xlabel, fontsize = 34)
#     else:
#         ax1[1][lam_tracker].set_xlabel("", fontsize = 0)
        
#     ax1[1][lam_tracker].plot(plotting_dict_no_hyper[rho]["a"],plotting_dict_attractors_no_hyper[rho], c = "k")
#     ax1[1][lam_tracker].set_ylim([0, alpha_max_res])
#     ax1[1][lam_tracker].tick_params(axis='both', which='major', labelsize=34)
#     ax1[1][lam_tracker].text(0.03, 1.01,texts[1][lam_tracker], transform=ax1[1][lam_tracker].transAxes, fontsize = 30)
        
    #Here we plot the affect on the population of infected hosts
    for i, rho in enumerate(rhos):
        ax1[2][lam_tracker].plot(eta_attractors[rho]['1'], plotting_dict_deaths_2[rho]["1"], c = f"{colours[i]}")
        ax1[2][lam_tracker].plot(eta_attractors[rho]['2'], plotting_dict_deaths_2[rho]["2"], c = f"{colours[i]}")
        ax1[2][lam_tracker].scatter(eta_attractors[rho]['1'][0], plotting_dict_deaths_2[rho]['1'][0], edgecolors = f"{colours[i]}", marker = "o", facecolors="none", s = 200)
        ax1[2][lam_tracker].scatter(eta_attractors[rho]['1'][-1], plotting_dict_deaths_2[rho]['1'][-1], edgecolors = f"{colours[i]}", marker = "o", facecolors= f"{colours[i]}", s = 200)
        try:
            ax1[2][lam_tracker].scatter(eta_attractors[rho]['2'][0], plotting_dict_deaths_2[rho]['2'][0], edgecolors = f"{colours[i]}", marker = "s", facecolors= "none", s = 200)
            ax1[2][lam_tracker].scatter(eta_attractors[rho]['2'][-1], plotting_dict_deaths_2[rho]['2'][-1], edgecolors = f"{colours[i]}", marker = "s", facecolors= f"{colours[i]}", s = 200)
        except:
            pass

    ax1[2][lam_tracker].plot(plotting_dict_hyper[rho]["a"], [1 for val in plotting_dict_hyper[rho]["a"]], c = "k")
    ax1[2][lam_tracker].plot(plotting_dict_hyper[rho]["r"], [1 for val in plotting_dict_hyper[rho]["r"]], c = "k")

    
    ax1[2][lam_tracker].set_xlabel("", fontsize = 0)
    ax1[2][lam_tracker].set_ylim([0.0, 8.0])
    ax1[2][lam_tracker].text(0.03, 1.01,texts[2][lam_tracker], transform=ax1[2][lam_tracker].transAxes, fontsize = 30)
    ax1[2][lam_tracker].tick_params(axis='both', which='major', labelsize=34)
    if lam_tracker == 0:
        ax1[2][lam_tracker].set_ylabel(r"Relative Avarage Virulence, $\Delta M(\alpha^*)$", fontsize = 34)
        
#     ax1[2][lam_tracker].set_title(fr"{titles[lam_tracker]}($\lambda$ = {lam})", fontsize = 34)
    
    #Here we plot the affect on the overall host population
    for i, rho in enumerate(rhos):
        ax1[3][lam_tracker].plot(eta_attractors[rho]['1'], plotting_dict_pops[rho]["1"], c = f"{colours[i]}")
        ax1[3][lam_tracker].plot(eta_attractors[rho]['2'], plotting_dict_pops[rho]["2"], c = f"{colours[i]}")
        ax1[3][lam_tracker].scatter(eta_attractors[rho]['1'][0], plotting_dict_pops[rho]['1'][0], edgecolors = f"{colours[i]}", marker = "o", facecolors="none", s = 200)
        ax1[3][lam_tracker].scatter(eta_attractors[rho]['1'][-1], plotting_dict_pops[rho]['1'][-1], edgecolors = f"{colours[i]}", marker = "o", facecolors= f"{colours[i]}", s = 200)
        try:
            ax1[3][lam_tracker].scatter(eta_attractors[rho]['2'][-1], plotting_dict_pops[rho]['2'][-1], edgecolors = f"{colours[i]}", marker = "s", facecolors= f"{colours[i]}", s = 200)
            ax1[3][lam_tracker].scatter(eta_attractors[rho]['2'][0], plotting_dict_pops[rho]['2'][0], edgecolors = f"{colours[i]}", marker = "s", facecolors="none", s = 200)
        except:
            pass
        

    ax1[3][lam_tracker].plot(plotting_dict_hyper[rho]["a"], [1 for val in plotting_dict_hyper[rho]["a"]], c = "k")
    ax1[3][lam_tracker].plot(plotting_dict_hyper[rho]["r"], [1 for val in plotting_dict_hyper[rho]["r"]], c = "k")


#     if lam_tracker == 1:
        #ax2[2][lam_tracker].set_xlabel(xlabel, fontsize = 34)
#     else:
        #ax2[2][lam_tracker].set_xlabel("", fontsize = 0)
        
    ax1[3][lam_tracker].set_ylim([0.0, 1.1])
    ax1[3][lam_tracker].tick_params(axis='both', which='major', labelsize=34)
    ax1[3][lam_tracker].text(0.03, 1.01,texts[3][lam_tracker], transform=ax1[3][lam_tracker].transAxes, fontsize = 30)
    #ax2[2][lam_tracker].text(0.03, 1.01,texts[2][lam_tracker], transform=#ax2[2][lam_tracker].transAxes, fontsize = 30)
    
    if lam_tracker == 0:
        ax1[3][lam_tracker].set_ylabel(r"Relative Population Size, $\Delta N(\alpha^*)$", fontsize = 34)
    
#     for i, rho in enumerate(rhos):
#         ax1[1][lam_tracker].plot(eta_attractors[rho]['1'], plotting_dict_hypers[rho]["1"], c = f"{colours[i]}")
#         ax1[1][lam_tracker].plot(eta_attractors[rho]['2'], plotting_dict_hypers[rho]["2"], c = f"{colours[i]}")
#         ax1[1][lam_tracker].scatter(eta_attractors[rho]['1'][0], plotting_dict_hypers[rho]['1'][0], edgecolors = f"{colours[i]}", marker = "o", facecolors="none", s = 200)
#         ax1[1][lam_tracker].scatter(eta_attractors[rho]['1'][-1], plotting_dict_hypers[rho]['1'][-1], edgecolors = f"{colours[i]}", marker = "o", facecolors= f"{colours[i]}", s = 200)
#         try:
#             ax1[1][lam_tracker].scatter(eta_attractors[rho]['2'][-1], plotting_dict_hypers[rho]['2'][-1], edgecolors = f"{colours[i]}", marker = "s", facecolors= f"{colours[i]}", s = 200)
#             ax1[1][lam_tracker].scatter(eta_attractors[rho]['2'][0], plotting_dict_hypers[rho]['2'][0], edgecolors = f"{colours[i]}", marker = "s", facecolors="none", s = 200)
#         except:
#             pass
        
    for i, rho in enumerate(rhos):
        ax1[1][lam_tracker].plot(eta_attractors[rho]['1'], plotting_dict_hyper_prop[rho]["1"], c = f"{colours[i]}")
        ax1[1][lam_tracker].plot(eta_attractors[rho]['2'], plotting_dict_hyper_prop[rho]["2"], c = f"{colours[i]}")
        ax1[1][lam_tracker].scatter(eta_attractors[rho]['1'][0], plotting_dict_hyper_prop[rho]['1'][0], edgecolors = f"{colours[i]}", marker = "o", facecolors="none", s = 200)
        ax1[1][lam_tracker].scatter(eta_attractors[rho]['1'][-1], plotting_dict_hyper_prop[rho]['1'][-1], edgecolors = f"{colours[i]}", marker = "o", facecolors= f"{colours[i]}", s = 200)
        try:
            ax1[1][lam_tracker].scatter(eta_attractors[rho]['2'][-1], plotting_dict_hyper_prop[rho]['2'][-1], edgecolors = f"{colours[i]}", marker = "s", facecolors= f"{colours[i]}", s = 200)
            ax1[1][lam_tracker].scatter(eta_attractors[rho]['2'][0], plotting_dict_hyper_prop[rho]['2'][0], edgecolors = f"{colours[i]}", marker = "s", facecolors="none", s = 200)
        except:
            pass
        
    if lam_tracker == 0:
        ax1[1][lam_tracker].set_ylabel(r"Hyperparasite Prevalance, $\frac{H}{I + H}$", fontsize = 34)
        
    
    if lam_tracker == 1:
        ax1[3][lam_tracker].set_xlabel(xlabel, fontsize = 34)
    else:
        ax1[1][lam_tracker].set_xlabel("", fontsize = 0)
        
    ax1[1][lam_tracker].set_ylim([0,1])
    ax1[1][lam_tracker].tick_params(axis='both', which='major', labelsize=34)
    ax1[1][lam_tracker].text(0.03, 1.01,texts[1][lam_tracker], transform=ax1[1][lam_tracker].transAxes, fontsize = 30)
#     #ax2[2][lam_tracker].text(0.03, 1.01,texts[2][lam_tracker], transform=#ax2[2][lam_tracker].transAxes, fontsize = 30)
#After this we save the figures in the appropriate folder

# fig2.legend(lines2,
#    labels=labels,
#    title='Probability of cotransmission',
#    fontsize = 24,
#    loc="upper right",
#    title_fontsize=26,
#    bbox_to_anchor=(0.90,0.85)
#    )

# plt.savefig(f"{output_folder}/effects_of_hyperparasitism.png", bbox_inches='tight')
# plt.close()

fig1.legend(lines1,
   labels=labels,
   title='Probability of cotransmission',
   fontsize = 24,
   loc="upper right",
   title_fontsize=26,
   bbox_to_anchor=(0.90,0.85)
   )

plt.savefig(f"{output_folder}/evolved_virulences.png", bbox_inches='tight')
plt.close()
del sol
```
