## Supplementary material for "Hyperparasitism and the evolution of parasite virulence": Code_base: hyperparasite_dies_out_plot.py

```
"""
This file demonstrates the consequences of the hyperparasite dying out after the parasite
has reached an ESS. This file is used to create Fig. S4.
"""
#We import the libraries we need
import sys, os, shutil
import runner
import numpy as np
import matplotlib.pyplot as plt
import pandas as pd

#Parameters used in the system
rho = 0.9

lam = 2.0
eta = 0.3

b = 2.0
q = 0.1
d = 0.5
gamma = 0.5

seed = 100
resolution = 100
sol = runner.PySolver()
    
plotting_alphas = []
plotting_evos = []
alpha_init = 20
sigma_init = 100

alpha_max = 5
sigma_max = 4
beta_max = 5


hyper = 1.0
c1 = 0.75
c2 = 2.25

#We initialise the system with the hyperparasites absent to find the natural ESS of the parasite
sol.alpha_ad_dyn(beta_max, alpha_max, sigma_max, b, q, d, rho, 0.0, gamma, 0.0, c1, c2, 0.0, seed, alpha_init, sigma_init, H_density = 0.0)
df = pd.read_csv(f"../data/alpha_evo/data_set{seed}.csv")

evo_steps = df["Evolutionary_step"].values
dft = df[df["Evolutionary_step"]==evo_steps[-1]]

pops = []
for val in dft["Trait_index_1"].values:
    df_pop = dft[dft["Trait_index_1"]==val][["Density_of_parasite"]]
    pops.append((df_pop["Density_of_parasite"].iloc[0])/(sum(dft["Density_of_parasite"].values)))
    
#Calculating the average trait value
alpha_weights = []
for i, val in enumerate(pops):
    alpha_weights.append(val*dft["Trait_index_1"].values[i])
alpha_mean = sum(alpha_weights)

#Rounding this to the closest integer to use as our initial conditions
alpha_init = round(alpha_mean)

host_density = dft["Density_of_hyperparasite"].iloc[0]
para_density = sum(dft["Density_of_parasite"].values)
hyper_density = 0.1

for i in range(2):
    sol.alpha_ad_dyn(beta_max, alpha_max, sigma_max, b, q, d, rho, eta, gamma, lam, c1, c2, hyper, seed, alpha_init, sigma_init, S_density = host_density, I_density = para_density, H_density = hyper_density)
    df = pd.read_csv(f"../data/alpha_evo/data_set{seed}.csv")

    seed += 1

    alpha_coords = df["alpha_val"].values
    evo_steps = df["Evolutionary_step"].values

    plotting_alphas.append(alpha_coords)
    plotting_evos.append(evo_steps)

    dft = df[df["Evolutionary_step"]==evo_steps[-1]]
    para_densities = dft["Density_of_parasite"].values
    hyper_densities = dft["Density_of_hyperparasite"].values
    trait_inds = dft["Trait_index_1"].values
    alpha_init = 0
    max_density = 0
    for i, den in enumerate(para_densities):
        if (den + hyper_densities[i]) > max_density:
            alpha_init = trait_inds[i]

    host_density = dft["Density_of_hyperparasite"].iloc[0]
    para_density = sum(dft["Density_of_parasite"].values)
    hyper_density = 0
    


#Here we visualise the plots by sequencing them after each other using
#this upshift
up_shift = []
for i in range(2):
    up_shift.append(i*1001)
    
#Colours associated with the values of eta
colours = {0.5:"black", 0.2:"blue", 1.0:"red"}

#Creating and saving the figure
fig, ax = plt.subplots(figsize = (10, 15))
    
tagged_labels = []
for i in range(2):
    
    alphas = plotting_alphas[i]
    evos_temp = plotting_evos[i]
    
    evo_steps = []
    for step in evos_temp:
        evo_steps.append(step + up_shift[i])
        
    ax.scatter(alphas, evo_steps, marker = ".", label = (i==0)*"Present" + (i==1)*"Absent")
        
fig.legend(
    title = "Hyperparasite",
    title_fontsize = 28,
   fontsize = 24,
   loc="upper right",
   bbox_to_anchor=(0.8,0.8)
)
ax.set_xlabel(r"Intrinsic virulence, $\alpha$", fontsize = 28)
ax.set_xlim([0,3])
ax.tick_params(axis='both', which='major', labelsize=24)
ax.set_ylabel("Evolutionary Time", fontsize = 28)

# ax.legend(loc='center left', bbox_to_anchor=(0.6, 0.85), fontsize = 14)
plt.savefig("../supplementary_figures/hyperparasite_die_out.png", bbox_inches='tight')
plt.savefig("../supplementary_figures/hyperparasite_die_out.pdf", bbox_inches='tight')
plt.close()
```
