## Supplementary material for "Hyperparasitism and the evolution of parasite virulence": Code_base: hyperparasite_eigenvalues.py

```
"""
This file is used to create heatmaps which demonstrate the invasion region of the
hyperparasite. Not used in the manuscript, held for completeness.
"""

import sys, os, shutil
import runner
import numpy as np
from math import sqrt, exp, dist, floor, ceil, isnan
import matplotlib.pyplot as plt
import imageio
import pandas as pd

#These local libraries are used to reduce the length of the file
sys.path.append("../src/Models/")

import singular_strategy as ss

sys.path.append("../src/zero_finder/")

import zero_finder_utilities as zf

sys.path.append("../src/figures/")

def Jacobian(S,I,H,b,beta,alpha,eta,d,q,gamma,rho,sigma,lam):
    
    Jac = np.array([
        [-2*q*(S+I+H) + b - beta*I - eta*beta*H - d, -2*q*(S+I+H) + b - beta*S + gamma, -2*q*(S+I+H) + b - eta*beta*S + gamma],
        [beta*I + eta*beta*(1-rho)*H, beta*S - sigma*H - d - alpha - gamma, -sigma*I + eta*(1-rho)*beta*H],
        [rho*eta*beta*H, sigma*H, rho*eta*beta*S + sigma*I - lam*alpha - d - gamma]
          ])
    
    return Jac

#This function calculates the gradient vector for a given parameter set
def calculate_alpha_gradient_vector(sol, resolution, beta_max, alpha_max, alpha_min_res, alpha_max_res, sigma_max, sigma, b, q, d, rho, eta, gamma, lam, c1, c2, hyper, seed):
    
    alpha_grad_vec = []
    alpha_val = []
    
    for i in range(resolution+1):
        
        #calculate the value of alpha
        alpha = alpha_min_res + (alpha_max_res - alpha_min_res) * (i / resolution)
        alpha_val.append(alpha)

        #and corresponding value of beta and it's derivatives
        beta = zf.beta_func(beta_max, alpha, alpha_max, sigma, sigma_max, c1, c2)
        
        dbetadalpha = zf.dbetadalpha_func(
            beta_max, alpha, alpha_max, sigma, sigma_max, c1, c2
        )

        #Solve the system to ecological steady state
        y = sol.eco_steady_state(
            beta, alpha, sigma, b, q, d, rho, eta, gamma, lam, c1, c2, hyper, seed
        )

        S = y[0]
        I = y[1]
        H = y[2]

        #Determine which version of the gradient we need. If the parasite is extinct just add an
        #arbitrary value
        if (I == 0 and H == 0):
            alpha_grad = 1*(alpha <= alpha_max/2) - 1*(alpha> alpha_max/2)
        elif (H == 0):
            alpha_grad = dbetadalpha*S/(d + alpha + gamma) - (beta*S)/((d + alpha + gamma)**2)
        elif (I == 0):
            alpha_grad = dbetadalpha*S/(d + lam*alpha + gamma) - (beta*S)/((d + lam*alpha + gamma)**2)
        else:
            alpha_grad = ss.fitness_gradient_alpha(
                beta, alpha, sigma, dbetadalpha, H, S, eta, lam, gamma, d, rho
            )

        #Convert the value into a sign
        alpha_grad = (alpha_grad > 0)*1 + (alpha_grad < 0)*-1
        alpha_grad_vec.append(alpha_grad)
        
    return alpha_grad_vec, alpha_val

#Iterate through the gradient vector to find any sign changes
def return_zeros(grad_vec, param_space):
    
    poss_attractors = []
    poss_repellers = []
    for i,val in enumerate(grad_vec[:-1]):
        if val == 1 and grad_vec[i+1] == -1:
            poss_attractors.append([param_space[i], param_space[i+1]])
        elif val == -1 and grad_vec[i+1] == 1:
            poss_repellers.append([param_space[i], param_space[i+1]])
        elif val == 0 and grad_vec[i+1] == -1:
            poss_attractors.append([param_space[i], param_space[i+1]])
            
    return poss_attractors, poss_repellers 
    
#This function calculates the singular strategy using the above functions to calculate the gradient and then find any sign changes
def recusive_sing_strat_function(sol, depth, iter_max, resolution, beta_max, alpha_max, alpha_min_res, alpha_max_res, sigma_max, sigma, b, q, d, rho, eta, gamma, lam, c1, c2, hyper, seed):
    
    
    sing_strats = []
    
    #We fill these lists with attractors and repellers
    attractors = []
    repellers = []
    
    #If we have resolved sufficiently we do one final resolution before exiting
    if depth == iter_max:
        #Calculating the alpha gradient vector
        alpha_grad_vec, alpha_val = calculate_alpha_gradient_vector(sol, resolution, beta_max, alpha_max, alpha_min_res, alpha_max_res, sigma_max, sigma, b, q, d, rho, eta, gamma, lam,c1, c2, hyper, seed)
        
        #Identifying singular strategies
        poss_attractors, poss_repellers = return_zeros(alpha_grad_vec,alpha_val)
#         if len(poss_attractors) > 2:
#             print(f"On run {eta} on rho {rho} with depth {depth}, and there are {len(poss_attractors)} possible singular strategies     ", end = "\r")
        #Allocating them to their respective places
        for zeros in poss_attractors:
            #We take the lower element as the midpoint can sometimes be outside the bounds of the hyperparasites existance
            attractors.append(zeros[0])
        for zeros in poss_repellers:
            repellers.append(zeros[0])
            
    #If we are not at our maximum depth we find possible positions of the singular strategy
    else:
        #First we calculate the alpha gradient
        alpha_grad_vec, alpha_val = calculate_alpha_gradient_vector(sol, resolution, beta_max, alpha_max, alpha_min_res, alpha_max_res, sigma_max, sigma, b, q, d, rho, eta, gamma, lam,c1, c2, hyper, seed)
        
        #Check for any sign changes using our function
        poss_attractors, poss_repellers = return_zeros(alpha_grad_vec,alpha_val)
#         if len(poss_attractors) > 2:
#             print(f"On run {eta} on rho {rho} with depth {depth}, and there are {len(poss_attractors)} possible singular strategies     ")
#             for pair in poss_attractors:
#                 print(pair[1] - pair[0])
        #Check bounds for any attractor regions
        if depth == 0:
            if alpha_grad_vec[0] == -1:
                attractors.append(alpha_val[0])
            if alpha_grad_vec[-1] == 1:
                attractors.append(alpha_val[-1])
                
        #Rerun the process of finding singular strategies for each of our possible attractors and repellers
        for pair in poss_attractors:
            temp_attractors, temp_repellers = recusive_sing_strat_function(sol, depth+1, iter_max, resolution, beta_max, alpha_max, pair[0], pair[1], sigma_max, sigma, b, q, d, rho, eta, gamma, lam, c1, c2, hyper, seed)
            
            for val in temp_attractors:
                attractors.append(val)
                
        for pair in poss_repellers:
            temp_attractors, temp_repellers = recusive_sing_strat_function(sol, depth+1, iter_max, resolution, beta_max, alpha_max, pair[0], pair[1], sigma_max, sigma, b, q, d, rho, eta, gamma, lam, c1, c2, hyper, seed)
                
            for val in temp_repellers:
                repellers.append(val)
                
    #We then clean up the attractors in case there are any singular strategies very close together
    if depth == 0:
        attractors = zf.clean_up_sing_strats(attractors)
        repellers = zf.clean_up_sing_strats(repellers)
                
    return attractors, repellers

rhos = [0.1, 0.5, 0.9]

rho = 0.9
lam = 1.0

lams = [0.5, 1, 2.0]

b = 2.0
q = 0.1
d = 0.5
gamma = 0.75

hyper = 1.0
c1 = 0.75
c2 = 2.25
b_base = 1
q_base = 1

alpha_max = 5
sigma_max = 4.0
beta_max = 5

sigma_min = 0.0
sigma_max_range = sigma_max

sigma = sigma_max

iter_max = 3
res_scalar = 4
depth_max = iter_max
resolution = 100
param_res = 200

seed = 10000

eta = 1.0

alpha_max_res = alpha_max
alpha_min_res = 0.0

beta_max = beta_max
alpha_max_param = alpha_max 
sigma_max_param = sigma_max
#Initiating the CPP solver
sol = runner.PySolver()

attractors_no_hyper, repellers_no_hyper = recusive_sing_strat_function(sol, 0, iter_max, resolution, beta_max, alpha_max, alpha_min_res, alpha_max_res, sigma_max, sigma, b, q, d, rho, eta, gamma, lam, c1, c2, 0.0, seed)

alpha = attractors_no_hyper[0]
beta = zf.beta_func(beta_max, alpha, alpha_max, sigma, sigma_max, c1, c2)

etas = [2.0*i/100 for i in range(101)]
lams = [2.0*i/100 for i in range(101)]

ticks = [0, 20, 40, 60, 80, 100]
eta_ticks = [etas[val] for val in ticks]
lam_ticks = [lams[val] for val in ticks]
#Solve the system to ecological steady state
y = sol.eco_steady_state(
    beta, alpha, sigma, b, q, d, rho, eta, gamma, lam, c1, c2, 0.0, seed
)

S = y[0]
I = y[1]
H = y[2]

for i, rho in enumerate(rhos):
    output_mat = []
    # output_mat_H = []
    for eta in etas:
        temp_row = []
        temp_row_H = []
        for lam in lams:
            #This function calculates the Jacobian
            Jac = Jacobian(S,I,H,b,beta,alpha,eta,d,q,gamma,rho,sigma,lam)

            #This function calculates the eigenvalues and eigenvectors of the Jacobian
            w, v = np.linalg.eig(Jac)

            eig_max = max(w.real)
            temp_row.append(eig_max)

        output_mat.append(temp_row)
    
    #This plotting code creates the Invasion heatmaps for the hyperparasite
    fig, ax = plt.subplots(figsize=(15, 15))
    im = ax.imshow(np.asarray(output_mat), origin = "lower")
    fig.colorbar(im, ax=ax)
    plt.xticks(ticks = ticks, labels = lam_ticks)
    plt.yticks(ticks = ticks, labels = eta_ticks)
    plt.xlabel(r"Hyperparasite Virulence Modifier, $\lambda$")
    plt.ylabel(r"Hyperparasite Infectivity Modifier, $\eta$")
    ax.set_title(fr"Parasite stability matrix with $\rho$ value {rho}")
    plt.savefig(f"../supplementary_figures/para_matrix{i}.png")
    plt.close()
```
