## Supplementary material for "Hyperparasitism and the evolution of parasite virulence": Code_base: linear_heatmap_res_scalar.py

```
"""
This file creates 3 csv files used by visualise_branching_matrices.py
to explore the possibility of branching being caused by the introduction
of hyperparasites. This file uses the multiprocessing library, the 
pandas library, the numpy library, the matplotlib library and the datetime 
library. It also invokes a custom built library called runner which is a 
cython compiled file. This file creates a low resolution heatmap and then 
refines one time on the boundary to reduce the time burden of the code.
"""

#Importing needed libraries
import pandas as pd
import numpy as np
import runner
import multiprocessing as mp
import matplotlib.pyplot as plt
from datetime import datetime

#This function creates a linear space of length resolution +  1
def make_linspace(val_min, val_max, resolution):
    
    """This function makes a list which contains a linear space between two points
    
    Args:
        val_min: float
            The minmum value we want the linear space to contain
        val_max: float
            The maximum value we want the linear space to contain
        resolution:
            The number of points between the minimum and maximum value in the linear space
    
    Returns:
        vals:
            The linear space specified above
    """
    
    vals = [val_min + (val_max - val_min)*(i/resolution) for i in range(resolution+1)]

    return vals

#This function sweeps across parameters using the multiprocessing library to reduce the time taken to run
def sweep_parameters(resolution, num_cpu, beta_max, beta_lin, alpha_max, alpha_min, sigma_max, b, q, d, rho, etas, gamma, lams, c1, c2, beta_scalar, hyper, seeds, alpha_init, sigma_init, evo_step_count, host_density, para_density, hyper_density, sol):
    
    """
    This function sweeps across the lists etas and lams to investigate which regions of the parameter space
    cause the hyperparasite to induce branching. This function uses the multiprocessing library to increase
    speed. After running a simulation the code creates a .csv file that can be accessed later to observe the
    evolutionary behaviour
    """
    #We create an empty list which we will add processes into as they are started
    processes = []
    
    #A counter we will use to track the number of processes we have begun
    count = 0
    for i in range(resolution+1):
        for j in range(resolution+1):
            #Provided there is a cpu we can allocate the process to we initate another process
            if count < num_cpu:
                p = mp.Process(target = sol.alpha_ad_dyn_v4, args = (beta_max, beta_lin, alpha_max, alpha_min, sigma_max, b, q, d, rho, etas[i], gamma, lams[j], c1, c2, beta_scalar, hyper, seeds[i][j], alpha_init, sigma_init, evo_step_count, host_density, para_density, hyper_density))
                p.start()
                processes.append(p)
                #We then increase the counter
                count += 1
            #If we have hit our number of cpus we collect the processes to be able to begin again
            else:
                for proc in processes:
                    proc.join()
                
                #We then create a new process and begin the loop again
                p = mp.Process(target = sol.alpha_ad_dyn_v4, args = (beta_max, beta_lin, alpha_max, alpha_min, sigma_max, b, q, d, rho, etas[i], gamma, lams[j], c1, c2, beta_scalar, hyper, seeds[i][j], alpha_init, sigma_init, evo_step_count, host_density, para_density, hyper_density))
                p.start()
                processes  = [p]
                count = 1    

    #Once we have finished our above loop we collect any outstanding files before
    #exiting the function
    for proc in processes:
        proc.join()
    return

#This function checks whether branching occured within a simulation
def check_branching_flag(seed):
    
    """This function reads the .csv file associated with a specific seed and
    then checks whether branching occured in this file. Branching is
    determined by a gap within the list of indices for the evolutionary trait
    values. This may over observe branching when it doesn't happen, but this is
    unlikely to consistently happen.
    """
    #We begin by setting our flag to be false
    branching_flag = False
    
    #We then read in the dataset
    df = pd.read_csv(f"../data/alpha_evo/data_set{seed}.csv")
    alpha_coords = df["alpha_val"].values
    evo_steps = df["Evolutionary_step"].unique()
    
    #We loop through the evolutionary steps, checking every 10 as it
    #is unlikely branching will have started and ended within 10 
    #evolutionary steps
    for step in evo_steps[::10]:
        dft = df[df["Evolutionary_step"]==step]
        alpha_coords = sorted(dft["Trait_index_1"].values)
        prev = alpha_coords[0]
        for val in alpha_coords[1:]:
            #If there is a gap greater than 1 index we assume branching
            #has happened
            if (val - prev) > 1:
                branching_flag = True
                break
            prev = val
        #If branching has happened we exit the code early
        if branching_flag:
            break
            
    #We also check whether hyperparasites are present at the end of the simulation
    dft = df[df["Evolutionary_step"]==evo_steps[-1]]
    hyperparasite_flag = 1 - dft["hyperparasites_present"].iloc[0]
    
    return branching_flag, hyperparasite_flag

#This function creates the heatmaps we will inspect to observe branching
def create_heatmap_matrice(resolution, res_scalar, num_cpu, max_depth, beta_max, beta_lin, alpha_max, alpha_min, sigma_max, b, q, d, rho, etas, gamma, lams, c1, c2, beta_scalar, hyper, seeds, alpha_init, sigma_init, evo_step_count, host_density, para_density, hyper_density, sol):
    
    """This function creates low resolution heatmaps showing branching regions within the host-parasite-hyperparasite
    system. This function then identifies the boundary of the branching region and reruns the analysis on that region
    at a higher resolution to refine the boundary of the branching region.
    """
    
    print("Doing sweep")
    #This creates the low resolution heatmap
    sweep_parameters(resolution, num_cpu, beta_max, beta_lin, alpha_max, alpha_min, sigma_max, b, q, d, rho, etas, gamma, lams, c1, c2, beta_scalar, hyper, seeds, alpha_init, sigma_init, evo_step_count, host_density, para_density, hyper_density, sol)
    
    heatmap_matrice_temp = []
    print("Making matrice")
    for i, row in enumerate(seeds):
        temp_row = []
        for j, seed in enumerate(row):
            branching_flag, hyperparasite_flag = check_branching_flag(seed)
            temp_row.append(branching_flag*1)
        heatmap_matrice_temp.append(temp_row)
        
    
    
    check_inds = []
    for i in range(resolution):
        for j in range(resolution):
            check_inds.append([i,j])
    
    for depth in range(1,max_depth+1):
        print(depth)
        iis = [0, 1]
        jjs = [0, 1]
        new_etas = []
        new_lams = []
        refine_inds = []
        
        
        #This section checks whether the index we are currently checking is on
        #the boundary of the branching region
        for ind in check_inds:
            val = heatmap_matrice_temp[ind[0]][ind[1]]
            flag = False
            for ii in iis:
                for jj in jjs:
                    if heatmap_matrice_temp[ind[0] + ii][ind[1] + jj] != val:
                        flag = True
            if flag:
                    new_etas.append([etas[ind[0]], etas[ind[0]+1]])
                    new_lams.append([lams[ind[1]], lams[ind[1]+1]])
                    refine_inds.append([ind[0], ind[1]])       

        #Here we replicate the branching matrix at a higher resolution
        heatmap_matrice_refined = []
        for i, row in enumerate(heatmap_matrice_temp[:-1]):
            for ii in range(res_scalar+1):
                temp_row = []
                for j, val in enumerate(row[:-1]):
                    for jj in range(res_scalar+1):
                        temp_row.append(val)
                temp_row.append(row[-1])
                heatmap_matrice_refined.append(temp_row)
                
                
        temp_row = []
        for j, val in enumerate(heatmap_matrice_temp[-1][:-1]):
            for jj in range(res_scalar+1):
                temp_row.append(val)
                
        temp_row.append(heatmap_matrice_refined[-1][-1])
#         print(len(temp_row))
        heatmap_matrice_refined.append(temp_row)
        
        print(len(heatmap_matrice_refined), len(heatmap_matrice_refined[0]))
        
        #Here we resweep over the region that needs refinement to create the more precise boundary region
        for k, ind in enumerate(refine_inds):
            print(f"Done {k+1} of {len(refine_inds)} runs       ", end = "\r")
            eta_pair = new_etas[k]
            lam_pair = new_lams[k]
            temp_etas = make_linspace(eta_pair[0], eta_pair[1], res_scalar)
            temp_lams = make_linspace(lam_pair[0], lam_pair[1], res_scalar)

            sweep_parameters(res_scalar, num_cpu, beta_max, beta_lin, alpha_max, alpha_min, sigma_max, b, q, d, rho, temp_etas, gamma, temp_lams, c1, c2, beta_scalar, hyper, seeds, alpha_init, sigma_init, evo_step_count, host_density, para_density, hyper_density, sol)
            for i in range(res_scalar + 1):
                for j in range(res_scalar + 1):
                    seed = seeds[i][j]
                    branching_flag, hyperparasite_flag = check_branching_flag(seed)
                    heatmap_matrice_refined[ind[0]*(res_scalar + 1) + i][ind[1]*(res_scalar + 1) + j] = 1*branching_flag
                    
        heatmap_matrice_temp = heatmap_matrice_refined
        etas = make_linspace(etas[0], etas[-1], len(heatmap_matrice_temp))
        lams = make_linspace(lams[0], lams[-1], len(heatmap_matrice_temp))
        
        check_inds = []
        for ind in refine_inds:
            for i in range(res_scalar + 1):
                for j in range(res_scalar + 1):
                    check_inds.append([ind[0]*res_scalar + i, ind[1]*res_scalar + j])
        
    heatmap_matrice = heatmap_matrice_temp
    
    return heatmap_matrice


t1 = datetime.now()

#Checking how many CPUs we have to work with
num_cpu = mp.cpu_count()

#Initiating the CPP based solver to perform our simulations
sol = runner.PySolver()

#Paramters of the system
b = 2.0
q = 0.1
d = 0.1
gamma = 0.1
sigma = 0.4
sigma_max = sigma
c1 = 1.0

#Value we initate the system at in the absence of the hyperparasite to derive
#the starting point in the presence of the hyperparasite
alpha_init = 10
sigma_init = 100
hyper = 1.0
beta_scalar = 1.0
rhos = [0.25, 0.5, 0.75]
lam_min = 0.0
lam_max = 5.0

eta_min = 0.0
eta_max = 1.0
seed_base = 100
evo_step_count = 2000

alpha_min = 0.0
alpha_max = 5.0
seeds = []

resolution = 30
res_scalar = 10

#Prepopulating the total seeds so we can reuse it later
for k in range(len(rhos)):
    tot_seeds = []
    for i in range(resolution+1):
        temp_seeds = []
        for j in range(resolution+1):
            temp_seeds.append(seed_base + k*((resolution+1)**2) + i*(resolution+1) + j)
        tot_seeds.append(temp_seeds)
    seeds.append(tot_seeds)
    
#Number of times we wish to refine our heatmaps by
max_depth = 1

#Making the initial linear spaces
lams = make_linspace(lam_min, lam_max, resolution)
etas = make_linspace(eta_min, eta_max, resolution)

processes = []

#Parameters for our trade-offs
beta_max = 0.5
beta_lin = 1.0
c2 = -60

#Performing the initial simulation
seed_for_baseline = 10

#As the hyperparasites are absent we can set any parameters associated with them to 0
rho_baseline = eta_baseline = lam_baseline = 0.0

sol.alpha_ad_dyn_v4(beta_max, beta_lin, alpha_max, alpha_min, sigma_max, b, q, d, rho_baseline, eta_baseline, gamma, lam_baseline, c1, c2, beta_scalar, 0.0, seed_for_baseline, alpha_init, sigma_init)

#We read in the data associated with the hyperparasite absent simulation
df_baseline = pd.read_csv(f"../data/alpha_evo/data_set{seed_for_baseline}.csv")
evo_steps = df_baseline["Evolutionary_step"].values

#We filter to only the last evolutionary step
dft = df_baseline[df_baseline["Evolutionary_step"]==evo_steps[-1]]

#Calculate the proportion of the population with each trait value
pops = []
for val in dft["Trait_index_1"].values:
    df_pop = dft[dft["Trait_index_1"]==val][["Density_of_parasite"]]
    pops.append((df_pop["Density_of_parasite"].iloc[0])/(sum(dft["Density_of_parasite"].values)))
    
#Calculating the average trait value
alpha_weights = []
for i, val in enumerate(pops):
    alpha_weights.append(val*dft["Trait_index_1"].values[i])
alpha_mean = sum(alpha_weights)

#Rounding this to the closest integer to use as our initial conditions
alpha_init = round(alpha_mean)

#The densities we initiate our populations with, taken from the end of the simulation
#without the hyperparasite present
host_density = dft["Density_of_Hosts"].iloc[0]
para_density = sum(dft["Density_of_parasite"].values)

#We initiate the hyperparasite with a low density
hyper_density = 0.1

#Here we loop through, calculating and then storing the heatmap at each point
for k, rho in enumerate(rhos):
    
    #Calculating the heatmap and refining it
    branching_mat = create_heatmap_matrice(resolution, res_scalar, num_cpu, max_depth, beta_max, beta_lin, alpha_max, alpha_min, sigma_max, b, q, d, rho, etas, gamma, lams, c1, c2, beta_scalar, hyper, seeds[k], alpha_init, sigma_init, evo_step_count, host_density, para_density, hyper_density, sol)

    #Getting our refined values of eta and lambda
    etas_refined = make_linspace(eta_min, eta_max, len(branching_mat) - 1)
    lams_refined = make_linspace(lam_min, lam_max, len(branching_mat) - 1)
    
    #Creating a temporary figure to check for issues
    fig, ax = plt.subplots(1, 1, figsize = (resolution, resolution))
    plt.imshow(branching_mat, origin = "lower")

    ticks_temp = []
    for i in range(len(branching_mat)):
        ticks_temp.append(i)

    ticks = ticks_temp[::resolution]
    ticks.append(ticks_temp[-1])
    lam_ticks = [round(lams_refined[i], 3) for i in ticks]
    eta_ticks = [round(etas_refined[i], 3) for i in ticks]
    plt.yticks(ticks = ticks, labels = eta_ticks, fontsize = max(0.4*resolution, 24))
    plt.xticks(ticks = ticks, labels = lam_ticks, fontsize = max(0.4*resolution, 24))
    plt.title(fr"Probability of cotransmission, $\rho$ = {rho}", fontsize = max(0.5*resolution, 30))
    plt.ylabel(r"$\eta$ value", fontsize = max(0.5*resolution, 30))
    plt.xlabel(r"$\lambda$ value", fontsize = max(0.5*resolution, 30))
    plt.savefig(f"../supplementary_figures/branching_heatmap_{k}_linear_test_comparison.png", bbox_inches = "tight")
    plt.close()
    print(f"The code took {datetime.now() - t1} seconds to run")
    
    #Here we create a pandas dataframe, within which we store our heatmap outputs
    array_for_df = []
    for i, row in enumerate(branching_mat):
        temp_row = [round(etas_refined[i], 3)]
        for val in row:
            temp_row.append(val)
        array_for_df.append(temp_row)
    df = pd.DataFrame(array_for_df)
    
    #The first column contains the eta_value
    df = df.rename(columns = {0: "eta_val"})
    
    #And the rest of the columns correspond to a value of lambda, meaning any
    #point within the .csv is described by a eta,lambda pair
    for i, val in enumerate(lams_refined):
        df = df.rename(columns = {i+1: str(round(val,3))})

    #We save the dataframe to a .csv file
    df.to_csv(f"../data/branching_mat_{k}.csv", index = False)
            
#We then delete the CPP solver
del sol
```
