## Supplementary material for "Hyperparasitism and the evolution of parasite virulence": Code_base: runner.cpp

```
/* Generated by Cython 0.29.32 */

/* BEGIN: Cython Metadata
{
    "distutils": {
        "depends": [
            "wrapper.cpp",
            "wrapper.h"
        ],
        "include_dirs": [
            "."
        ],
        "language": "c++",
        "name": "runner",
        "sources": [
            "runner.pyx"
        ]
    },
    "module_name": "runner"
}
END: Cython Metadata */

#ifndef PY_SSIZE_T_CLEAN
#define PY_SSIZE_T_CLEAN
#endif /* PY_SSIZE_T_CLEAN */
#include "Python.h"
#ifndef Py_PYTHON_H
    #error Python headers needed to compile C extensions, please install development version of Python.
#elif PY_VERSION_HEX < 0x02060000 || (0x03000000 <= PY_VERSION_HEX && PY_VERSION_HEX < 0x03030000)
    #error Cython requires Python 2.6+ or Python 3.3+.
#else
#define CYTHON_ABI "0_29_32"
#define CYTHON_HEX_VERSION 0x001D20F0
#define CYTHON_FUTURE_DIVISION 0
#include <stddef.h>
#ifndef offsetof
  #define offsetof(type, member) ( (size_t) & ((type*)0) -> member )
#endif
#if !defined(WIN32) && !defined(MS_WINDOWS)
  #ifndef __stdcall
    #define __stdcall
  #endif
  #ifndef __cdecl
    #define __cdecl
  #endif
  #ifndef __fastcall
    #define __fastcall
  #endif
#endif
#ifndef DL_IMPORT
  #define DL_IMPORT(t) t
#endif
#ifndef DL_EXPORT
  #define DL_EXPORT(t) t
#endif
#define __PYX_COMMA ,
#ifndef HAVE_LONG_LONG
  #if PY_VERSION_HEX >= 0x02070000
    #define HAVE_LONG_LONG
  #endif
#endif
#ifndef PY_LONG_LONG
  #define PY_LONG_LONG LONG_LONG
#endif
#ifndef Py_HUGE_VAL
  #define Py_HUGE_VAL HUGE_VAL
#endif
#ifdef PYPY_VERSION
  #define CYTHON_COMPILING_IN_PYPY 1
  #define CYTHON_COMPILING_IN_PYSTON 0
  #define CYTHON_COMPILING_IN_CPYTHON 0
  #define CYTHON_COMPILING_IN_NOGIL 0
  #undef CYTHON_USE_TYPE_SLOTS
  #define CYTHON_USE_TYPE_SLOTS 0
  #undef CYTHON_USE_PYTYPE_LOOKUP
  #define CYTHON_USE_PYTYPE_LOOKUP 0
  #if PY_VERSION_HEX < 0x03050000
    #undef CYTHON_USE_ASYNC_SLOTS
    #define CYTHON_USE_ASYNC_SLOTS 0
  #elif !defined(CYTHON_USE_ASYNC_SLOTS)
    #define CYTHON_USE_ASYNC_SLOTS 1
  #endif
  #undef CYTHON_USE_PYLIST_INTERNALS
  #define CYTHON_USE_PYLIST_INTERNALS 0
  #undef CYTHON_USE_UNICODE_INTERNALS
  #define CYTHON_USE_UNICODE_INTERNALS 0
  #undef CYTHON_USE_UNICODE_WRITER
  #define CYTHON_USE_UNICODE_WRITER 0
  #undef CYTHON_USE_PYLONG_INTERNALS
  #define CYTHON_USE_PYLONG_INTERNALS 0
  #undef CYTHON_AVOID_BORROWED_REFS
  #define CYTHON_AVOID_BORROWED_REFS 1
  #undef CYTHON_ASSUME_SAFE_MACROS
  #define CYTHON_ASSUME_SAFE_MACROS 0
  #undef CYTHON_UNPACK_METHODS
  #define CYTHON_UNPACK_METHODS 0
  #undef CYTHON_FAST_THREAD_STATE
  #define CYTHON_FAST_THREAD_STATE 0
  #undef CYTHON_FAST_PYCALL
  #define CYTHON_FAST_PYCALL 0
  #undef CYTHON_PEP489_MULTI_PHASE_INIT
  #define CYTHON_PEP489_MULTI_PHASE_INIT 0
  #undef CYTHON_USE_TP_FINALIZE
  #define CYTHON_USE_TP_FINALIZE 0
  #undef CYTHON_USE_DICT_VERSIONS
  #define CYTHON_USE_DICT_VERSIONS 0
  #undef CYTHON_USE_EXC_INFO_STACK
  #define CYTHON_USE_EXC_INFO_STACK 0
  #ifndef CYTHON_UPDATE_DESCRIPTOR_DOC
    #define CYTHON_UPDATE_DESCRIPTOR_DOC (PYPY_VERSION_HEX >= 0x07030900)
  #endif
#elif defined(PYSTON_VERSION)
  #define CYTHON_COMPILING_IN_PYPY 0
  #define CYTHON_COMPILING_IN_PYSTON 1
  #define CYTHON_COMPILING_IN_CPYTHON 0
  #define CYTHON_COMPILING_IN_NOGIL 0
  #ifndef CYTHON_USE_TYPE_SLOTS
    #define CYTHON_USE_TYPE_SLOTS 1
  #endif
  #undef CYTHON_USE_PYTYPE_LOOKUP
  #define CYTHON_USE_PYTYPE_LOOKUP 0
  #undef CYTHON_USE_ASYNC_SLOTS
  #define CYTHON_USE_ASYNC_SLOTS 0
  #undef CYTHON_USE_PYLIST_INTERNALS
  #define CYTHON_USE_PYLIST_INTERNALS 0
  #ifndef CYTHON_USE_UNICODE_INTERNALS
    #define CYTHON_USE_UNICODE_INTERNALS 1
  #endif
  #undef CYTHON_USE_UNICODE_WRITER
  #define CYTHON_USE_UNICODE_WRITER 0
  #undef CYTHON_USE_PYLONG_INTERNALS
  #define CYTHON_USE_PYLONG_INTERNALS 0
  #ifndef CYTHON_AVOID_BORROWED_REFS
    #define CYTHON_AVOID_BORROWED_REFS 0
  #endif
  #ifndef CYTHON_ASSUME_SAFE_MACROS
    #define CYTHON_ASSUME_SAFE_MACROS 1
  #endif
  #ifndef CYTHON_UNPACK_METHODS
    #define CYTHON_UNPACK_METHODS 1
  #endif
  #undef CYTHON_FAST_THREAD_STATE
  #define CYTHON_FAST_THREAD_STATE 0
  #undef CYTHON_FAST_PYCALL
  #define CYTHON_FAST_PYCALL 0
  #undef CYTHON_PEP489_MULTI_PHASE_INIT
  #define CYTHON_PEP489_MULTI_PHASE_INIT 0
  #undef CYTHON_USE_TP_FINALIZE
  #define CYTHON_USE_TP_FINALIZE 0
  #undef CYTHON_USE_DICT_VERSIONS
  #define CYTHON_USE_DICT_VERSIONS 0
  #undef CYTHON_USE_EXC_INFO_STACK
  #define CYTHON_USE_EXC_INFO_STACK 0
  #ifndef CYTHON_UPDATE_DESCRIPTOR_DOC
    #define CYTHON_UPDATE_DESCRIPTOR_DOC 0
  #endif
#elif defined(PY_NOGIL)
  #define CYTHON_COMPILING_IN_PYPY 0
  #define CYTHON_COMPILING_IN_PYSTON 0
  #define CYTHON_COMPILING_IN_CPYTHON 0
  #define CYTHON_COMPILING_IN_NOGIL 1
  #ifndef CYTHON_USE_TYPE_SLOTS
    #define CYTHON_USE_TYPE_SLOTS 1
  #endif
  #undef CYTHON_USE_PYTYPE_LOOKUP
  #define CYTHON_USE_PYTYPE_LOOKUP 0
  #ifndef CYTHON_USE_ASYNC_SLOTS
    #define CYTHON_USE_ASYNC_SLOTS 1
  #endif
  #undef CYTHON_USE_PYLIST_INTERNALS
  #define CYTHON_USE_PYLIST_INTERNALS 0
  #ifndef CYTHON_USE_UNICODE_INTERNALS
    #define CYTHON_USE_UNICODE_INTERNALS 1
  #endif
  #undef CYTHON_USE_UNICODE_WRITER
  #define CYTHON_USE_UNICODE_WRITER 0
  #undef CYTHON_USE_PYLONG_INTERNALS
  #define CYTHON_USE_PYLONG_INTERNALS 0
  #ifndef CYTHON_AVOID_BORROWED_REFS
    #define CYTHON_AVOID_BORROWED_REFS 0
  #endif
  #ifndef CYTHON_ASSUME_SAFE_MACROS
    #define CYTHON_ASSUME_SAFE_MACROS 1
  #endif
  #ifndef CYTHON_UNPACK_METHODS
    #define CYTHON_UNPACK_METHODS 1
  #endif
  #undef CYTHON_FAST_THREAD_STATE
  #define CYTHON_FAST_THREAD_STATE 0
  #undef CYTHON_FAST_PYCALL
  #define CYTHON_FAST_PYCALL 0
  #ifndef CYTHON_PEP489_MULTI_PHASE_INIT
    #define CYTHON_PEP489_MULTI_PHASE_INIT 1
  #endif
  #ifndef CYTHON_USE_TP_FINALIZE
    #define CYTHON_USE_TP_FINALIZE 1
  #endif
  #undef CYTHON_USE_DICT_VERSIONS
  #define CYTHON_USE_DICT_VERSIONS 0
  #undef CYTHON_USE_EXC_INFO_STACK
  #define CYTHON_USE_EXC_INFO_STACK 0
#else
  #define CYTHON_COMPILING_IN_PYPY 0
  #define CYTHON_COMPILING_IN_PYSTON 0
  #define CYTHON_COMPILING_IN_CPYTHON 1
  #define CYTHON_COMPILING_IN_NOGIL 0
  #ifndef CYTHON_USE_TYPE_SLOTS
    #define CYTHON_USE_TYPE_SLOTS 1
  #endif
  #if PY_VERSION_HEX < 0x02070000
    #undef CYTHON_USE_PYTYPE_LOOKUP
    #define CYTHON_USE_PYTYPE_LOOKUP 0
  #elif !defined(CYTHON_USE_PYTYPE_LOOKUP)
    #define CYTHON_USE_PYTYPE_LOOKUP 1
  #endif
  #if PY_MAJOR_VERSION < 3
    #undef CYTHON_USE_ASYNC_SLOTS
    #define CYTHON_USE_ASYNC_SLOTS 0
  #elif !defined(CYTHON_USE_ASYNC_SLOTS)
    #define CYTHON_USE_ASYNC_SLOTS 1
  #endif
  #if PY_VERSION_HEX < 0x02070000
    #undef CYTHON_USE_PYLONG_INTERNALS
    #define CYTHON_USE_PYLONG_INTERNALS 0
  #elif !defined(CYTHON_USE_PYLONG_INTERNALS)
    #define CYTHON_USE_PYLONG_INTERNALS 1
  #endif
  #ifndef CYTHON_USE_PYLIST_INTERNALS
    #define CYTHON_USE_PYLIST_INTERNALS 1
  #endif
  #ifndef CYTHON_USE_UNICODE_INTERNALS
    #define CYTHON_USE_UNICODE_INTERNALS 1
  #endif
  #if PY_VERSION_HEX < 0x030300F0 || PY_VERSION_HEX >= 0x030B00A2
    #undef CYTHON_USE_UNICODE_WRITER
    #define CYTHON_USE_UNICODE_WRITER 0
  #elif !defined(CYTHON_USE_UNICODE_WRITER)
    #define CYTHON_USE_UNICODE_WRITER 1
  #endif
  #ifndef CYTHON_AVOID_BORROWED_REFS
    #define CYTHON_AVOID_BORROWED_REFS 0
  #endif
  #ifndef CYTHON_ASSUME_SAFE_MACROS
    #define CYTHON_ASSUME_SAFE_MACROS 1
  #endif
  #ifndef CYTHON_UNPACK_METHODS
    #define CYTHON_UNPACK_METHODS 1
  #endif
  #if PY_VERSION_HEX >= 0x030B00A4
    #undef CYTHON_FAST_THREAD_STATE
    #define CYTHON_FAST_THREAD_STATE 0
  #elif !defined(CYTHON_FAST_THREAD_STATE)
    #define CYTHON_FAST_THREAD_STATE 1
  #endif
  #ifndef CYTHON_FAST_PYCALL
    #define CYTHON_FAST_PYCALL (PY_VERSION_HEX < 0x030A0000)
  #endif
  #ifndef CYTHON_PEP489_MULTI_PHASE_INIT
    #define CYTHON_PEP489_MULTI_PHASE_INIT (PY_VERSION_HEX >= 0x03050000)
  #endif
  #ifndef CYTHON_USE_TP_FINALIZE
    #define CYTHON_USE_TP_FINALIZE (PY_VERSION_HEX >= 0x030400a1)
  #endif
  #ifndef CYTHON_USE_DICT_VERSIONS
    #define CYTHON_USE_DICT_VERSIONS (PY_VERSION_HEX >= 0x030600B1)
  #endif
  #if PY_VERSION_HEX >= 0x030B00A4
    #undef CYTHON_USE_EXC_INFO_STACK
    #define CYTHON_USE_EXC_INFO_STACK 0
  #elif !defined(CYTHON_USE_EXC_INFO_STACK)
    #define CYTHON_USE_EXC_INFO_STACK (PY_VERSION_HEX >= 0x030700A3)
  #endif
  #ifndef CYTHON_UPDATE_DESCRIPTOR_DOC
    #define CYTHON_UPDATE_DESCRIPTOR_DOC 1
  #endif
#endif
#if !defined(CYTHON_FAST_PYCCALL)
#define CYTHON_FAST_PYCCALL  (CYTHON_FAST_PYCALL && PY_VERSION_HEX >= 0x030600B1)
#endif
#if CYTHON_USE_PYLONG_INTERNALS
  #if PY_MAJOR_VERSION < 3
    #include "longintrepr.h"
  #endif
  #undef SHIFT
  #undef BASE
  #undef MASK
  #ifdef SIZEOF_VOID_P
    enum { __pyx_check_sizeof_voidp = 1 / (int)(SIZEOF_VOID_P == sizeof(void*)) };
  #endif
#endif
#ifndef __has_attribute
  #define __has_attribute(x) 0
#endif
#ifndef __has_cpp_attribute
  #define __has_cpp_attribute(x) 0
#endif
#ifndef CYTHON_RESTRICT
  #if defined(__GNUC__)
    #define CYTHON_RESTRICT __restrict__
  #elif defined(_MSC_VER) && _MSC_VER >= 1400
    #define CYTHON_RESTRICT __restrict
  #elif defined (__STDC_VERSION__) && __STDC_VERSION__ >= 199901L
    #define CYTHON_RESTRICT restrict
  #else
    #define CYTHON_RESTRICT
  #endif
#endif
#ifndef CYTHON_UNUSED
# if defined(__GNUC__)
#   if !(defined(__cplusplus)) || (__GNUC__ > 3 || (__GNUC__ == 3 && __GNUC_MINOR__ >= 4))
#     define CYTHON_UNUSED __attribute__ ((__unused__))
#   else
#     define CYTHON_UNUSED
#   endif
# elif defined(__ICC) || (defined(__INTEL_COMPILER) && !defined(_MSC_VER))
#   define CYTHON_UNUSED __attribute__ ((__unused__))
# else
#   define CYTHON_UNUSED
# endif
#endif
#ifndef CYTHON_MAYBE_UNUSED_VAR
#  if defined(__cplusplus)
     template<class T> void CYTHON_MAYBE_UNUSED_VAR( const T& ) { }
#  else
#    define CYTHON_MAYBE_UNUSED_VAR(x) (void)(x)
#  endif
#endif
#ifndef CYTHON_NCP_UNUSED
# if CYTHON_COMPILING_IN_CPYTHON
#  define CYTHON_NCP_UNUSED
# else
#  define CYTHON_NCP_UNUSED CYTHON_UNUSED
# endif
#endif
#define __Pyx_void_to_None(void_result) ((void)(void_result), Py_INCREF(Py_None), Py_None)
#ifdef _MSC_VER
    #ifndef _MSC_STDINT_H_
        #if _MSC_VER < 1300
           typedef unsigned char     uint8_t;
           typedef unsigned int      uint32_t;
        #else
           typedef unsigned __int8   uint8_t;
           typedef unsigned __int32  uint32_t;
        #endif
    #endif
#else
   #include <stdint.h>
#endif
#ifndef CYTHON_FALLTHROUGH
  #if defined(__cplusplus) && __cplusplus >= 201103L
    #if __has_cpp_attribute(fallthrough)
      #define CYTHON_FALLTHROUGH [[fallthrough]]
    #elif __has_cpp_attribute(clang::fallthrough)
      #define CYTHON_FALLTHROUGH [[clang::fallthrough]]
    #elif __has_cpp_attribute(gnu::fallthrough)
      #define CYTHON_FALLTHROUGH [[gnu::fallthrough]]
    #endif
  #endif
  #ifndef CYTHON_FALLTHROUGH
    #if __has_attribute(fallthrough)
      #define CYTHON_FALLTHROUGH __attribute__((fallthrough))
    #else
      #define CYTHON_FALLTHROUGH
    #endif
  #endif
  #if defined(__clang__ ) && defined(__apple_build_version__)
    #if __apple_build_version__ < 7000000
      #undef  CYTHON_FALLTHROUGH
      #define CYTHON_FALLTHROUGH
    #endif
  #endif
#endif

#ifndef __cplusplus
  #error "Cython files generated with the C++ option must be compiled with a C++ compiler."
#endif
#ifndef CYTHON_INLINE
  #if defined(__clang__)
    #define CYTHON_INLINE __inline__ __attribute__ ((__unused__))
  #else
    #define CYTHON_INLINE inline
  #endif
#endif
template<typename T>
void __Pyx_call_destructor(T& x) {
    x.~T();
}
template<typename T>
class __Pyx_FakeReference {
  public:
    __Pyx_FakeReference() : ptr(NULL) { }
    __Pyx_FakeReference(const T& ref) : ptr(const_cast<T*>(&ref)) { }
    T *operator->() { return ptr; }
    T *operator&() { return ptr; }
    operator T&() { return *ptr; }
    template<typename U> bool operator ==(U other) { return *ptr == other; }
    template<typename U> bool operator !=(U other) { return *ptr != other; }
  private:
    T *ptr;
};

#if CYTHON_COMPILING_IN_PYPY && PY_VERSION_HEX < 0x02070600 && !defined(Py_OptimizeFlag)
  #define Py_OptimizeFlag 0
#endif
#define __PYX_BUILD_PY_SSIZE_T "n"
#define CYTHON_FORMAT_SSIZE_T "z"
#if PY_MAJOR_VERSION < 3
  #define __Pyx_BUILTIN_MODULE_NAME "__builtin__"
  #define __Pyx_PyCode_New(a, k, l, s, f, code, c, n, v, fv, cell, fn, name, fline, lnos)\
          PyCode_New(a+k, l, s, f, code, c, n, v, fv, cell, fn, name, fline, lnos)
  #define __Pyx_DefaultClassType PyClass_Type
#else
  #define __Pyx_BUILTIN_MODULE_NAME "builtins"
  #define __Pyx_DefaultClassType PyType_Type
#if PY_VERSION_HEX >= 0x030B00A1
    static CYTHON_INLINE PyCodeObject* __Pyx_PyCode_New(int a, int k, int l, int s, int f,
                                                    PyObject *code, PyObject *c, PyObject* n, PyObject *v,
                                                    PyObject *fv, PyObject *cell, PyObject* fn,
                                                    PyObject *name, int fline, PyObject *lnos) {
        PyObject *kwds=NULL, *argcount=NULL, *posonlyargcount=NULL, *kwonlyargcount=NULL;
        PyObject *nlocals=NULL, *stacksize=NULL, *flags=NULL, *replace=NULL, *call_result=NULL, *empty=NULL;
        const char *fn_cstr=NULL;
        const char *name_cstr=NULL;
        PyCodeObject* co=NULL;
        PyObject *type, *value, *traceback;
        PyErr_Fetch(&type, &value, &traceback);
        if (!(kwds=PyDict_New())) goto end;
        if (!(argcount=PyLong_FromLong(a))) goto end;
        if (PyDict_SetItemString(kwds, "co_argcount", argcount) != 0) goto end;
        if (!(posonlyargcount=PyLong_FromLong(0))) goto end;
        if (PyDict_SetItemString(kwds, "co_posonlyargcount", posonlyargcount) != 0) goto end;
        if (!(kwonlyargcount=PyLong_FromLong(k))) goto end;
        if (PyDict_SetItemString(kwds, "co_kwonlyargcount", kwonlyargcount) != 0) goto end;
        if (!(nlocals=PyLong_FromLong(l))) goto end;
        if (PyDict_SetItemString(kwds, "co_nlocals", nlocals) != 0) goto end;
        if (!(stacksize=PyLong_FromLong(s))) goto end;
        if (PyDict_SetItemString(kwds, "co_stacksize", stacksize) != 0) goto end;
        if (!(flags=PyLong_FromLong(f))) goto end;
        if (PyDict_SetItemString(kwds, "co_flags", flags) != 0) goto end;
        if (PyDict_SetItemString(kwds, "co_code", code) != 0) goto end;
        if (PyDict_SetItemString(kwds, "co_consts", c) != 0) goto end;
        if (PyDict_SetItemString(kwds, "co_names", n) != 0) goto end;
        if (PyDict_SetItemString(kwds, "co_varnames", v) != 0) goto end;
        if (PyDict_SetItemString(kwds, "co_freevars", fv) != 0) goto end;
        if (PyDict_SetItemString(kwds, "co_cellvars", cell) != 0) goto end;
        if (PyDict_SetItemString(kwds, "co_linetable", lnos) != 0) goto end;
        if (!(fn_cstr=PyUnicode_AsUTF8AndSize(fn, NULL))) goto end;
        if (!(name_cstr=PyUnicode_AsUTF8AndSize(name, NULL))) goto end;
        if (!(co = PyCode_NewEmpty(fn_cstr, name_cstr, fline))) goto end;
        if (!(replace = PyObject_GetAttrString((PyObject*)co, "replace"))) goto cleanup_code_too;
        if (!(empty = PyTuple_New(0))) goto cleanup_code_too; // unfortunately __pyx_empty_tuple isn't available here
        if (!(call_result = PyObject_Call(replace, empty, kwds))) goto cleanup_code_too;
        Py_XDECREF((PyObject*)co);
        co = (PyCodeObject*)call_result;
        call_result = NULL;
        if (0) {
            cleanup_code_too:
            Py_XDECREF((PyObject*)co);
            co = NULL;
        }
        end:
        Py_XDECREF(kwds);
        Py_XDECREF(argcount);
        Py_XDECREF(posonlyargcount);
        Py_XDECREF(kwonlyargcount);
        Py_XDECREF(nlocals);
        Py_XDECREF(stacksize);
        Py_XDECREF(replace);
        Py_XDECREF(call_result);
        Py_XDECREF(empty);
        if (type) {
            PyErr_Restore(type, value, traceback);
        }
        return co;
    }
#else
  #define __Pyx_PyCode_New(a, k, l, s, f, code, c, n, v, fv, cell, fn, name, fline, lnos)\
          PyCode_New(a, k, l, s, f, code, c, n, v, fv, cell, fn, name, fline, lnos)
#endif
  #define __Pyx_DefaultClassType PyType_Type
#endif
#ifndef Py_TPFLAGS_CHECKTYPES
  #define Py_TPFLAGS_CHECKTYPES 0
#endif
#ifndef Py_TPFLAGS_HAVE_INDEX
  #define Py_TPFLAGS_HAVE_INDEX 0
#endif
#ifndef Py_TPFLAGS_HAVE_NEWBUFFER
  #define Py_TPFLAGS_HAVE_NEWBUFFER 0
#endif
#ifndef Py_TPFLAGS_HAVE_FINALIZE
  #define Py_TPFLAGS_HAVE_FINALIZE 0
#endif
#ifndef METH_STACKLESS
  #define METH_STACKLESS 0
#endif
#if PY_VERSION_HEX <= 0x030700A3 || !defined(METH_FASTCALL)
  #ifndef METH_FASTCALL
     #define METH_FASTCALL 0x80
  #endif
  typedef PyObject *(*__Pyx_PyCFunctionFast) (PyObject *self, PyObject *const *args, Py_ssize_t nargs);
  typedef PyObject *(*__Pyx_PyCFunctionFastWithKeywords) (PyObject *self, PyObject *const *args,
                                                          Py_ssize_t nargs, PyObject *kwnames);
#else
  #define __Pyx_PyCFunctionFast _PyCFunctionFast
  #define __Pyx_PyCFunctionFastWithKeywords _PyCFunctionFastWithKeywords
#endif
#if CYTHON_FAST_PYCCALL
#define __Pyx_PyFastCFunction_Check(func)\
    ((PyCFunction_Check(func) && (METH_FASTCALL == (PyCFunction_GET_FLAGS(func) & ~(METH_CLASS | METH_STATIC | METH_COEXIST | METH_KEYWORDS | METH_STACKLESS)))))
#else
#define __Pyx_PyFastCFunction_Check(func) 0
#endif
#if CYTHON_COMPILING_IN_PYPY && !defined(PyObject_Malloc)
  #define PyObject_Malloc(s)   PyMem_Malloc(s)
  #define PyObject_Free(p)     PyMem_Free(p)
  #define PyObject_Realloc(p)  PyMem_Realloc(p)
#endif
#if CYTHON_COMPILING_IN_CPYTHON && PY_VERSION_HEX < 0x030400A1
  #define PyMem_RawMalloc(n)           PyMem_Malloc(n)
  #define PyMem_RawRealloc(p, n)       PyMem_Realloc(p, n)
  #define PyMem_RawFree(p)             PyMem_Free(p)
#endif
#if CYTHON_COMPILING_IN_PYSTON
  #define __Pyx_PyCode_HasFreeVars(co)  PyCode_HasFreeVars(co)
  #define __Pyx_PyFrame_SetLineNumber(frame, lineno) PyFrame_SetLineNumber(frame, lineno)
#else
  #define __Pyx_PyCode_HasFreeVars(co)  (PyCode_GetNumFree(co) > 0)
  #define __Pyx_PyFrame_SetLineNumber(frame, lineno)  (frame)->f_lineno = (lineno)
#endif
#if !CYTHON_FAST_THREAD_STATE || PY_VERSION_HEX < 0x02070000
  #define __Pyx_PyThreadState_Current PyThreadState_GET()
#elif PY_VERSION_HEX >= 0x03060000
  #define __Pyx_PyThreadState_Current _PyThreadState_UncheckedGet()
#elif PY_VERSION_HEX >= 0x03000000
  #define __Pyx_PyThreadState_Current PyThreadState_GET()
#else
  #define __Pyx_PyThreadState_Current _PyThreadState_Current
#endif
#if PY_VERSION_HEX < 0x030700A2 && !defined(PyThread_tss_create) && !defined(Py_tss_NEEDS_INIT)
#include "pythread.h"
#define Py_tss_NEEDS_INIT 0
typedef int Py_tss_t;
static CYTHON_INLINE int PyThread_tss_create(Py_tss_t *key) {
  *key = PyThread_create_key();
  return 0;
}
static CYTHON_INLINE Py_tss_t * PyThread_tss_alloc(void) {
  Py_tss_t *key = (Py_tss_t *)PyObject_Malloc(sizeof(Py_tss_t));
  *key = Py_tss_NEEDS_INIT;
  return key;
}
static CYTHON_INLINE void PyThread_tss_free(Py_tss_t *key) {
  PyObject_Free(key);
}
static CYTHON_INLINE int PyThread_tss_is_created(Py_tss_t *key) {
  return *key != Py_tss_NEEDS_INIT;
}
static CYTHON_INLINE void PyThread_tss_delete(Py_tss_t *key) {
  PyThread_delete_key(*key);
  *key = Py_tss_NEEDS_INIT;
}
static CYTHON_INLINE int PyThread_tss_set(Py_tss_t *key, void *value) {
  return PyThread_set_key_value(*key, value);
}
static CYTHON_INLINE void * PyThread_tss_get(Py_tss_t *key) {
  return PyThread_get_key_value(*key);
}
#endif
#if CYTHON_COMPILING_IN_CPYTHON || defined(_PyDict_NewPresized)
#define __Pyx_PyDict_NewPresized(n)  ((n <= 8) ? PyDict_New() : _PyDict_NewPresized(n))
#else
#define __Pyx_PyDict_NewPresized(n)  PyDict_New()
#endif
#if PY_MAJOR_VERSION >= 3 || CYTHON_FUTURE_DIVISION
  #define __Pyx_PyNumber_Divide(x,y)         PyNumber_TrueDivide(x,y)
  #define __Pyx_PyNumber_InPlaceDivide(x,y)  PyNumber_InPlaceTrueDivide(x,y)
#else
  #define __Pyx_PyNumber_Divide(x,y)         PyNumber_Divide(x,y)
  #define __Pyx_PyNumber_InPlaceDivide(x,y)  PyNumber_InPlaceDivide(x,y)
#endif
#if CYTHON_COMPILING_IN_CPYTHON && PY_VERSION_HEX >= 0x030500A1 && CYTHON_USE_UNICODE_INTERNALS
#define __Pyx_PyDict_GetItemStr(dict, name)  _PyDict_GetItem_KnownHash(dict, name, ((PyASCIIObject *) name)->hash)
#else
#define __Pyx_PyDict_GetItemStr(dict, name)  PyDict_GetItem(dict, name)
#endif
#if PY_VERSION_HEX > 0x03030000 && defined(PyUnicode_KIND)
  #define CYTHON_PEP393_ENABLED 1
  #if defined(PyUnicode_IS_READY)
  #define __Pyx_PyUnicode_READY(op)       (likely(PyUnicode_IS_READY(op)) ?\
                                              0 : _PyUnicode_Ready((PyObject *)(op)))
  #else
  #define __Pyx_PyUnicode_READY(op)       (0)
  #endif
  #define __Pyx_PyUnicode_GET_LENGTH(u)   PyUnicode_GET_LENGTH(u)
  #define __Pyx_PyUnicode_READ_CHAR(u, i) PyUnicode_READ_CHAR(u, i)
  #define __Pyx_PyUnicode_MAX_CHAR_VALUE(u)   PyUnicode_MAX_CHAR_VALUE(u)
  #define __Pyx_PyUnicode_KIND(u)         PyUnicode_KIND(u)
  #define __Pyx_PyUnicode_DATA(u)         PyUnicode_DATA(u)
  #define __Pyx_PyUnicode_READ(k, d, i)   PyUnicode_READ(k, d, i)
  #define __Pyx_PyUnicode_WRITE(k, d, i, ch)  PyUnicode_WRITE(k, d, i, ch)
  #if defined(PyUnicode_IS_READY) && defined(PyUnicode_GET_SIZE)
  #if CYTHON_COMPILING_IN_CPYTHON && PY_VERSION_HEX >= 0x03090000
  #define __Pyx_PyUnicode_IS_TRUE(u)      (0 != (likely(PyUnicode_IS_READY(u)) ? PyUnicode_GET_LENGTH(u) : ((PyCompactUnicodeObject *)(u))->wstr_length))
  #else
  #define __Pyx_PyUnicode_IS_TRUE(u)      (0 != (likely(PyUnicode_IS_READY(u)) ? PyUnicode_GET_LENGTH(u) : PyUnicode_GET_SIZE(u)))
  #endif
  #else
  #define __Pyx_PyUnicode_IS_TRUE(u)      (0 != PyUnicode_GET_LENGTH(u))
  #endif
#else
  #define CYTHON_PEP393_ENABLED 0
  #define PyUnicode_1BYTE_KIND  1
  #define PyUnicode_2BYTE_KIND  2
  #define PyUnicode_4BYTE_KIND  4
  #define __Pyx_PyUnicode_READY(op)       (0)
  #define __Pyx_PyUnicode_GET_LENGTH(u)   PyUnicode_GET_SIZE(u)
  #define __Pyx_PyUnicode_READ_CHAR(u, i) ((Py_UCS4)(PyUnicode_AS_UNICODE(u)[i]))
  #define __Pyx_PyUnicode_MAX_CHAR_VALUE(u)   ((sizeof(Py_UNICODE) == 2) ? 65535 : 1114111)
  #define __Pyx_PyUnicode_KIND(u)         (sizeof(Py_UNICODE))
  #define __Pyx_PyUnicode_DATA(u)         ((void*)PyUnicode_AS_UNICODE(u))
  #define __Pyx_PyUnicode_READ(k, d, i)   ((void)(k), (Py_UCS4)(((Py_UNICODE*)d)[i]))
  #define __Pyx_PyUnicode_WRITE(k, d, i, ch)  (((void)(k)), ((Py_UNICODE*)d)[i] = ch)
  #define __Pyx_PyUnicode_IS_TRUE(u)      (0 != PyUnicode_GET_SIZE(u))
#endif
#if CYTHON_COMPILING_IN_PYPY
  #define __Pyx_PyUnicode_Concat(a, b)      PyNumber_Add(a, b)
  #define __Pyx_PyUnicode_ConcatSafe(a, b)  PyNumber_Add(a, b)
#else
  #define __Pyx_PyUnicode_Concat(a, b)      PyUnicode_Concat(a, b)
  #define __Pyx_PyUnicode_ConcatSafe(a, b)  ((unlikely((a) == Py_None) || unlikely((b) == Py_None)) ?\
      PyNumber_Add(a, b) : __Pyx_PyUnicode_Concat(a, b))
#endif
#if CYTHON_COMPILING_IN_PYPY && !defined(PyUnicode_Contains)
  #define PyUnicode_Contains(u, s)  PySequence_Contains(u, s)
#endif
#if CYTHON_COMPILING_IN_PYPY && !defined(PyByteArray_Check)
  #define PyByteArray_Check(obj)  PyObject_TypeCheck(obj, &PyByteArray_Type)
#endif
#if CYTHON_COMPILING_IN_PYPY && !defined(PyObject_Format)
  #define PyObject_Format(obj, fmt)  PyObject_CallMethod(obj, "__format__", "O", fmt)
#endif
#define __Pyx_PyString_FormatSafe(a, b)   ((unlikely((a) == Py_None || (PyString_Check(b) && !PyString_CheckExact(b)))) ? PyNumber_Remainder(a, b) : __Pyx_PyString_Format(a, b))
#define __Pyx_PyUnicode_FormatSafe(a, b)  ((unlikely((a) == Py_None || (PyUnicode_Check(b) && !PyUnicode_CheckExact(b)))) ? PyNumber_Remainder(a, b) : PyUnicode_Format(a, b))
#if PY_MAJOR_VERSION >= 3
  #define __Pyx_PyString_Format(a, b)  PyUnicode_Format(a, b)
#else
  #define __Pyx_PyString_Format(a, b)  PyString_Format(a, b)
#endif
#if PY_MAJOR_VERSION < 3 && !defined(PyObject_ASCII)
  #define PyObject_ASCII(o)            PyObject_Repr(o)
#endif
#if PY_MAJOR_VERSION >= 3
  #define PyBaseString_Type            PyUnicode_Type
  #define PyStringObject               PyUnicodeObject
  #define PyString_Type                PyUnicode_Type
  #define PyString_Check               PyUnicode_Check
  #define PyString_CheckExact          PyUnicode_CheckExact
#ifndef PyObject_Unicode
  #define PyObject_Unicode             PyObject_Str
#endif
#endif
#if PY_MAJOR_VERSION >= 3
  #define __Pyx_PyBaseString_Check(obj) PyUnicode_Check(obj)
  #define __Pyx_PyBaseString_CheckExact(obj) PyUnicode_CheckExact(obj)
#else
  #define __Pyx_PyBaseString_Check(obj) (PyString_Check(obj) || PyUnicode_Check(obj))
  #define __Pyx_PyBaseString_CheckExact(obj) (PyString_CheckExact(obj) || PyUnicode_CheckExact(obj))
#endif
#ifndef PySet_CheckExact
  #define PySet_CheckExact(obj)        (Py_TYPE(obj) == &PySet_Type)
#endif
#if PY_VERSION_HEX >= 0x030900A4
  #define __Pyx_SET_REFCNT(obj, refcnt) Py_SET_REFCNT(obj, refcnt)
  #define __Pyx_SET_SIZE(obj, size) Py_SET_SIZE(obj, size)
#else
  #define __Pyx_SET_REFCNT(obj, refcnt) Py_REFCNT(obj) = (refcnt)
  #define __Pyx_SET_SIZE(obj, size) Py_SIZE(obj) = (size)
#endif
#if CYTHON_ASSUME_SAFE_MACROS
  #define __Pyx_PySequence_SIZE(seq)  Py_SIZE(seq)
#else
  #define __Pyx_PySequence_SIZE(seq)  PySequence_Size(seq)
#endif
#if PY_MAJOR_VERSION >= 3
  #define PyIntObject                  PyLongObject
  #define PyInt_Type                   PyLong_Type
  #define PyInt_Check(op)              PyLong_Check(op)
  #define PyInt_CheckExact(op)         PyLong_CheckExact(op)
  #define PyInt_FromString             PyLong_FromString
  #define PyInt_FromUnicode            PyLong_FromUnicode
  #define PyInt_FromLong               PyLong_FromLong
  #define PyInt_FromSize_t             PyLong_FromSize_t
  #define PyInt_FromSsize_t            PyLong_FromSsize_t
  #define PyInt_AsLong                 PyLong_AsLong
  #define PyInt_AS_LONG                PyLong_AS_LONG
  #define PyInt_AsSsize_t              PyLong_AsSsize_t
  #define PyInt_AsUnsignedLongMask     PyLong_AsUnsignedLongMask
  #define PyInt_AsUnsignedLongLongMask PyLong_AsUnsignedLongLongMask
  #define PyNumber_Int                 PyNumber_Long
#endif
#if PY_MAJOR_VERSION >= 3
  #define PyBoolObject                 PyLongObject
#endif
#if PY_MAJOR_VERSION >= 3 && CYTHON_COMPILING_IN_PYPY
  #ifndef PyUnicode_InternFromString
    #define PyUnicode_InternFromString(s) PyUnicode_FromString(s)
  #endif
#endif
#if PY_VERSION_HEX < 0x030200A4
  typedef long Py_hash_t;
  #define __Pyx_PyInt_FromHash_t PyInt_FromLong
  #define __Pyx_PyInt_AsHash_t   __Pyx_PyIndex_AsHash_t
#else
  #define __Pyx_PyInt_FromHash_t PyInt_FromSsize_t
  #define __Pyx_PyInt_AsHash_t   __Pyx_PyIndex_AsSsize_t
#endif
#if PY_MAJOR_VERSION >= 3
  #define __Pyx_PyMethod_New(func, self, klass) ((self) ? ((void)(klass), PyMethod_New(func, self)) : __Pyx_NewRef(func))
#else
  #define __Pyx_PyMethod_New(func, self, klass) PyMethod_New(func, self, klass)
#endif
#if CYTHON_USE_ASYNC_SLOTS
  #if PY_VERSION_HEX >= 0x030500B1
    #define __Pyx_PyAsyncMethodsStruct PyAsyncMethods
    #define __Pyx_PyType_AsAsync(obj) (Py_TYPE(obj)->tp_as_async)
  #else
    #define __Pyx_PyType_AsAsync(obj) ((__Pyx_PyAsyncMethodsStruct*) (Py_TYPE(obj)->tp_reserved))
  #endif
#else
  #define __Pyx_PyType_AsAsync(obj) NULL
#endif
#ifndef __Pyx_PyAsyncMethodsStruct
    typedef struct {
        unaryfunc am_await;
        unaryfunc am_aiter;
        unaryfunc am_anext;
    } __Pyx_PyAsyncMethodsStruct;
#endif

#if defined(_WIN32) || defined(WIN32) || defined(MS_WINDOWS)
  #if !defined(_USE_MATH_DEFINES)
    #define _USE_MATH_DEFINES
  #endif
#endif
#include <math.h>
#ifdef NAN
#define __PYX_NAN() ((float) NAN)
#else
static CYTHON_INLINE float __PYX_NAN() {
  float value;
  memset(&value, 0xFF, sizeof(value));
  return value;
}
#endif
#if defined(__CYGWIN__) && defined(_LDBL_EQ_DBL)
#define __Pyx_truncl trunc
#else
#define __Pyx_truncl truncl
#endif

#define __PYX_MARK_ERR_POS(f_index, lineno) \
    { __pyx_filename = __pyx_f[f_index]; (void)__pyx_filename; __pyx_lineno = lineno; (void)__pyx_lineno; __pyx_clineno = __LINE__; (void)__pyx_clineno; }
#define __PYX_ERR(f_index, lineno, Ln_error) \
    { __PYX_MARK_ERR_POS(f_index, lineno) goto Ln_error; }

#ifndef __PYX_EXTERN_C
  #ifdef __cplusplus
    #define __PYX_EXTERN_C extern "C"
  #else
    #define __PYX_EXTERN_C extern
  #endif
#endif

#define __PYX_HAVE__runner
#define __PYX_HAVE_API__runner
/* Early includes */
#include "wrapper.cpp"
#include "ios"
#include "new"
#include "stdexcept"
#include "typeinfo"
#include "wrapper.h"
#ifdef _OPENMP
#include <omp.h>
#endif /* _OPENMP */

#if defined(PYREX_WITHOUT_ASSERTIONS) && !defined(CYTHON_WITHOUT_ASSERTIONS)
#define CYTHON_WITHOUT_ASSERTIONS
#endif

typedef struct {PyObject **p; const char *s; const Py_ssize_t n; const char* encoding;
                const char is_unicode; const char is_str; const char intern; } __Pyx_StringTabEntry;

#define __PYX_DEFAULT_STRING_ENCODING_IS_ASCII 0
#define __PYX_DEFAULT_STRING_ENCODING_IS_UTF8 0
#define __PYX_DEFAULT_STRING_ENCODING_IS_DEFAULT (PY_MAJOR_VERSION >= 3 && __PYX_DEFAULT_STRING_ENCODING_IS_UTF8)
#define __PYX_DEFAULT_STRING_ENCODING ""
#define __Pyx_PyObject_FromString __Pyx_PyBytes_FromString
#define __Pyx_PyObject_FromStringAndSize __Pyx_PyBytes_FromStringAndSize
#define __Pyx_uchar_cast(c) ((unsigned char)c)
#define __Pyx_long_cast(x) ((long)x)
#define __Pyx_fits_Py_ssize_t(v, type, is_signed)  (\
    (sizeof(type) < sizeof(Py_ssize_t))  ||\
    (sizeof(type) > sizeof(Py_ssize_t) &&\
          likely(v < (type)PY_SSIZE_T_MAX ||\
                 v == (type)PY_SSIZE_T_MAX)  &&\
          (!is_signed || likely(v > (type)PY_SSIZE_T_MIN ||\
                                v == (type)PY_SSIZE_T_MIN)))  ||\
    (sizeof(type) == sizeof(Py_ssize_t) &&\
          (is_signed || likely(v < (type)PY_SSIZE_T_MAX ||\
                               v == (type)PY_SSIZE_T_MAX)))  )
static CYTHON_INLINE int __Pyx_is_valid_index(Py_ssize_t i, Py_ssize_t limit) {
    return (size_t) i < (size_t) limit;
}
#if defined (__cplusplus) && __cplusplus >= 201103L
    #include <cstdlib>
    #define __Pyx_sst_abs(value) std::abs(value)
#elif SIZEOF_INT >= SIZEOF_SIZE_T
    #define __Pyx_sst_abs(value) abs(value)
#elif SIZEOF_LONG >= SIZEOF_SIZE_T
    #define __Pyx_sst_abs(value) labs(value)
#elif defined (_MSC_VER)
    #define __Pyx_sst_abs(value) ((Py_ssize_t)_abs64(value))
#elif defined (__STDC_VERSION__) && __STDC_VERSION__ >= 199901L
    #define __Pyx_sst_abs(value) llabs(value)
#elif defined (__GNUC__)
    #define __Pyx_sst_abs(value) __builtin_llabs(value)
#else
    #define __Pyx_sst_abs(value) ((value<0) ? -value : value)
#endif
static CYTHON_INLINE const char* __Pyx_PyObject_AsString(PyObject*);
static CYTHON_INLINE const char* __Pyx_PyObject_AsStringAndSize(PyObject*, Py_ssize_t* length);
#define __Pyx_PyByteArray_FromString(s) PyByteArray_FromStringAndSize((const char*)s, strlen((const char*)s))
#define __Pyx_PyByteArray_FromStringAndSize(s, l) PyByteArray_FromStringAndSize((const char*)s, l)
#define __Pyx_PyBytes_FromString        PyBytes_FromString
#define __Pyx_PyBytes_FromStringAndSize PyBytes_FromStringAndSize
static CYTHON_INLINE PyObject* __Pyx_PyUnicode_FromString(const char*);
#if PY_MAJOR_VERSION < 3
    #define __Pyx_PyStr_FromString        __Pyx_PyBytes_FromString
    #define __Pyx_PyStr_FromStringAndSize __Pyx_PyBytes_FromStringAndSize
#else
    #define __Pyx_PyStr_FromString        __Pyx_PyUnicode_FromString
    #define __Pyx_PyStr_FromStringAndSize __Pyx_PyUnicode_FromStringAndSize
#endif
#define __Pyx_PyBytes_AsWritableString(s)     ((char*) PyBytes_AS_STRING(s))
#define __Pyx_PyBytes_AsWritableSString(s)    ((signed char*) PyBytes_AS_STRING(s))
#define __Pyx_PyBytes_AsWritableUString(s)    ((unsigned char*) PyBytes_AS_STRING(s))
#define __Pyx_PyBytes_AsString(s)     ((const char*) PyBytes_AS_STRING(s))
#define __Pyx_PyBytes_AsSString(s)    ((const signed char*) PyBytes_AS_STRING(s))
#define __Pyx_PyBytes_AsUString(s)    ((const unsigned char*) PyBytes_AS_STRING(s))
#define __Pyx_PyObject_AsWritableString(s)    ((char*) __Pyx_PyObject_AsString(s))
#define __Pyx_PyObject_AsWritableSString(s)    ((signed char*) __Pyx_PyObject_AsString(s))
#define __Pyx_PyObject_AsWritableUString(s)    ((unsigned char*) __Pyx_PyObject_AsString(s))
#define __Pyx_PyObject_AsSString(s)    ((const signed char*) __Pyx_PyObject_AsString(s))
#define __Pyx_PyObject_AsUString(s)    ((const unsigned char*) __Pyx_PyObject_AsString(s))
#define __Pyx_PyObject_FromCString(s)  __Pyx_PyObject_FromString((const char*)s)
#define __Pyx_PyBytes_FromCString(s)   __Pyx_PyBytes_FromString((const char*)s)
#define __Pyx_PyByteArray_FromCString(s)   __Pyx_PyByteArray_FromString((const char*)s)
#define __Pyx_PyStr_FromCString(s)     __Pyx_PyStr_FromString((const char*)s)
#define __Pyx_PyUnicode_FromCString(s) __Pyx_PyUnicode_FromString((const char*)s)
static CYTHON_INLINE size_t __Pyx_Py_UNICODE_strlen(const Py_UNICODE *u) {
    const Py_UNICODE *u_end = u;
    while (*u_end++) ;
    return (size_t)(u_end - u - 1);
}
#define __Pyx_PyUnicode_FromUnicode(u)       PyUnicode_FromUnicode(u, __Pyx_Py_UNICODE_strlen(u))
#define __Pyx_PyUnicode_FromUnicodeAndLength PyUnicode_FromUnicode
#define __Pyx_PyUnicode_AsUnicode            PyUnicode_AsUnicode
#define __Pyx_NewRef(obj) (Py_INCREF(obj), obj)
#define __Pyx_Owned_Py_None(b) __Pyx_NewRef(Py_None)
static CYTHON_INLINE PyObject * __Pyx_PyBool_FromLong(long b);
static CYTHON_INLINE int __Pyx_PyObject_IsTrue(PyObject*);
static CYTHON_INLINE int __Pyx_PyObject_IsTrueAndDecref(PyObject*);
static CYTHON_INLINE PyObject* __Pyx_PyNumber_IntOrLong(PyObject* x);
#define __Pyx_PySequence_Tuple(obj)\
    (likely(PyTuple_CheckExact(obj)) ? __Pyx_NewRef(obj) : PySequence_Tuple(obj))
static CYTHON_INLINE Py_ssize_t __Pyx_PyIndex_AsSsize_t(PyObject*);
static CYTHON_INLINE PyObject * __Pyx_PyInt_FromSize_t(size_t);
static CYTHON_INLINE Py_hash_t __Pyx_PyIndex_AsHash_t(PyObject*);
#if CYTHON_ASSUME_SAFE_MACROS
#define __pyx_PyFloat_AsDouble(x) (PyFloat_CheckExact(x) ? PyFloat_AS_DOUBLE(x) : PyFloat_AsDouble(x))
#else
#define __pyx_PyFloat_AsDouble(x) PyFloat_AsDouble(x)
#endif
#define __pyx_PyFloat_AsFloat(x) ((float) __pyx_PyFloat_AsDouble(x))
#if PY_MAJOR_VERSION >= 3
#define __Pyx_PyNumber_Int(x) (PyLong_CheckExact(x) ? __Pyx_NewRef(x) : PyNumber_Long(x))
#else
#define __Pyx_PyNumber_Int(x) (PyInt_CheckExact(x) ? __Pyx_NewRef(x) : PyNumber_Int(x))
#endif
#define __Pyx_PyNumber_Float(x) (PyFloat_CheckExact(x) ? __Pyx_NewRef(x) : PyNumber_Float(x))
#if PY_MAJOR_VERSION < 3 && __PYX_DEFAULT_STRING_ENCODING_IS_ASCII
static int __Pyx_sys_getdefaultencoding_not_ascii;
static int __Pyx_init_sys_getdefaultencoding_params(void) {
    PyObject* sys;
    PyObject* default_encoding = NULL;
    PyObject* ascii_chars_u = NULL;
    PyObject* ascii_chars_b = NULL;
    const char* default_encoding_c;
    sys = PyImport_ImportModule("sys");
    if (!sys) goto bad;
    default_encoding = PyObject_CallMethod(sys, (char*) "getdefaultencoding", NULL);
    Py_DECREF(sys);
    if (!default_encoding) goto bad;
    default_encoding_c = PyBytes_AsString(default_encoding);
    if (!default_encoding_c) goto bad;
    if (strcmp(default_encoding_c, "ascii") == 0) {
        __Pyx_sys_getdefaultencoding_not_ascii = 0;
    } else {
        char ascii_chars[128];
        int c;
        for (c = 0; c < 128; c++) {
            ascii_chars[c] = c;
        }
        __Pyx_sys_getdefaultencoding_not_ascii = 1;
        ascii_chars_u = PyUnicode_DecodeASCII(ascii_chars, 128, NULL);
        if (!ascii_chars_u) goto bad;
        ascii_chars_b = PyUnicode_AsEncodedString(ascii_chars_u, default_encoding_c, NULL);
        if (!ascii_chars_b || !PyBytes_Check(ascii_chars_b) || memcmp(ascii_chars, PyBytes_AS_STRING(ascii_chars_b), 128) != 0) {
            PyErr_Format(
                PyExc_ValueError,
                "This module compiled with c_string_encoding=ascii, but default encoding '%.200s' is not a superset of ascii.",
                default_encoding_c);
            goto bad;
        }
        Py_DECREF(ascii_chars_u);
        Py_DECREF(ascii_chars_b);
    }
    Py_DECREF(default_encoding);
    return 0;
bad:
    Py_XDECREF(default_encoding);
    Py_XDECREF(ascii_chars_u);
    Py_XDECREF(ascii_chars_b);
    return -1;
}
#endif
#if __PYX_DEFAULT_STRING_ENCODING_IS_DEFAULT && PY_MAJOR_VERSION >= 3
#define __Pyx_PyUnicode_FromStringAndSize(c_str, size) PyUnicode_DecodeUTF8(c_str, size, NULL)
#else
#define __Pyx_PyUnicode_FromStringAndSize(c_str, size) PyUnicode_Decode(c_str, size, __PYX_DEFAULT_STRING_ENCODING, NULL)
#if __PYX_DEFAULT_STRING_ENCODING_IS_DEFAULT
static char* __PYX_DEFAULT_STRING_ENCODING;
static int __Pyx_init_sys_getdefaultencoding_params(void) {
    PyObject* sys;
    PyObject* default_encoding = NULL;
    char* default_encoding_c;
    sys = PyImport_ImportModule("sys");
    if (!sys) goto bad;
    default_encoding = PyObject_CallMethod(sys, (char*) (const char*) "getdefaultencoding", NULL);
    Py_DECREF(sys);
    if (!default_encoding) goto bad;
    default_encoding_c = PyBytes_AsString(default_encoding);
    if (!default_encoding_c) goto bad;
    __PYX_DEFAULT_STRING_ENCODING = (char*) malloc(strlen(default_encoding_c) + 1);
    if (!__PYX_DEFAULT_STRING_ENCODING) goto bad;
    strcpy(__PYX_DEFAULT_STRING_ENCODING, default_encoding_c);
    Py_DECREF(default_encoding);
    return 0;
bad:
    Py_XDECREF(default_encoding);
    return -1;
}
#endif
#endif


/* Test for GCC > 2.95 */
#if defined(__GNUC__)     && (__GNUC__ > 2 || (__GNUC__ == 2 && (__GNUC_MINOR__ > 95)))
  #define likely(x)   __builtin_expect(!!(x), 1)
  #define unlikely(x) __builtin_expect(!!(x), 0)
#else /* !__GNUC__ or GCC < 2.95 */
  #define likely(x)   (x)
  #define unlikely(x) (x)
#endif /* __GNUC__ */
static CYTHON_INLINE void __Pyx_pretend_to_initialize(void* ptr) { (void)ptr; }

static PyObject *__pyx_m = NULL;
static PyObject *__pyx_d;
static PyObject *__pyx_b;
static PyObject *__pyx_cython_runtime = NULL;
static PyObject *__pyx_empty_tuple;
static PyObject *__pyx_empty_bytes;
static PyObject *__pyx_empty_unicode;
static int __pyx_lineno;
static int __pyx_clineno = 0;
static const char * __pyx_cfilenm= __FILE__;
static const char *__pyx_filename;


static const char *__pyx_f[] = {
  "stringsource",
  "runner.pyx",
};

/*--- Type declarations ---*/
struct __pyx_obj_6runner_PySolver;

/* "runner.pyx":4
 * from solvers cimport Quick_solver
 * 
 * cdef class PySolver:             # <<<<<<<<<<<<<<
 * 
 *     cdef Quick_solver cpp_solver
 */
struct __pyx_obj_6runner_PySolver {
  PyObject_HEAD
  solvers::Quick_solver cpp_solver;
};


/* --- Runtime support code (head) --- */
/* Refnanny.proto */
#ifndef CYTHON_REFNANNY
  #define CYTHON_REFNANNY 0
#endif
#if CYTHON_REFNANNY
  typedef struct {
    void (*INCREF)(void*, PyObject*, int);
    void (*DECREF)(void*, PyObject*, int);
    void (*GOTREF)(void*, PyObject*, int);
    void (*GIVEREF)(void*, PyObject*, int);
    void* (*SetupContext)(const char*, int, const char*);
    void (*FinishContext)(void**);
  } __Pyx_RefNannyAPIStruct;
  static __Pyx_RefNannyAPIStruct *__Pyx_RefNanny = NULL;
  static __Pyx_RefNannyAPIStruct *__Pyx_RefNannyImportAPI(const char *modname);
  #define __Pyx_RefNannyDeclarations void *__pyx_refnanny = NULL;
#ifdef WITH_THREAD
  #define __Pyx_RefNannySetupContext(name, acquire_gil)\
          if (acquire_gil) {\
              PyGILState_STATE __pyx_gilstate_save = PyGILState_Ensure();\
              __pyx_refnanny = __Pyx_RefNanny->SetupContext((name), __LINE__, __FILE__);\
              PyGILState_Release(__pyx_gilstate_save);\
          } else {\
              __pyx_refnanny = __Pyx_RefNanny->SetupContext((name), __LINE__, __FILE__);\
          }
#else
  #define __Pyx_RefNannySetupContext(name, acquire_gil)\
          __pyx_refnanny = __Pyx_RefNanny->SetupContext((name), __LINE__, __FILE__)
#endif
  #define __Pyx_RefNannyFinishContext()\
          __Pyx_RefNanny->FinishContext(&__pyx_refnanny)
  #define __Pyx_INCREF(r)  __Pyx_RefNanny->INCREF(__pyx_refnanny, (PyObject *)(r), __LINE__)
  #define __Pyx_DECREF(r)  __Pyx_RefNanny->DECREF(__pyx_refnanny, (PyObject *)(r), __LINE__)
  #define __Pyx_GOTREF(r)  __Pyx_RefNanny->GOTREF(__pyx_refnanny, (PyObject *)(r), __LINE__)
  #define __Pyx_GIVEREF(r) __Pyx_RefNanny->GIVEREF(__pyx_refnanny, (PyObject *)(r), __LINE__)
  #define __Pyx_XINCREF(r)  do { if((r) != NULL) {__Pyx_INCREF(r); }} while(0)
  #define __Pyx_XDECREF(r)  do { if((r) != NULL) {__Pyx_DECREF(r); }} while(0)
  #define __Pyx_XGOTREF(r)  do { if((r) != NULL) {__Pyx_GOTREF(r); }} while(0)
  #define __Pyx_XGIVEREF(r) do { if((r) != NULL) {__Pyx_GIVEREF(r);}} while(0)
#else
  #define __Pyx_RefNannyDeclarations
  #define __Pyx_RefNannySetupContext(name, acquire_gil)
  #define __Pyx_RefNannyFinishContext()
  #define __Pyx_INCREF(r) Py_INCREF(r)
  #define __Pyx_DECREF(r) Py_DECREF(r)
  #define __Pyx_GOTREF(r)
  #define __Pyx_GIVEREF(r)
  #define __Pyx_XINCREF(r) Py_XINCREF(r)
  #define __Pyx_XDECREF(r) Py_XDECREF(r)
  #define __Pyx_XGOTREF(r)
  #define __Pyx_XGIVEREF(r)
#endif
#define __Pyx_XDECREF_SET(r, v) do {\
        PyObject *tmp = (PyObject *) r;\
        r = v; __Pyx_XDECREF(tmp);\
    } while (0)
#define __Pyx_DECREF_SET(r, v) do {\
        PyObject *tmp = (PyObject *) r;\
        r = v; __Pyx_DECREF(tmp);\
    } while (0)
#define __Pyx_CLEAR(r)    do { PyObject* tmp = ((PyObject*)(r)); r = NULL; __Pyx_DECREF(tmp);} while(0)
#define __Pyx_XCLEAR(r)   do { if((r) != NULL) {PyObject* tmp = ((PyObject*)(r)); r = NULL; __Pyx_DECREF(tmp);}} while(0)

/* PyObjectGetAttrStr.proto */
#if CYTHON_USE_TYPE_SLOTS
static CYTHON_INLINE PyObject* __Pyx_PyObject_GetAttrStr(PyObject* obj, PyObject* attr_name);
#else
#define __Pyx_PyObject_GetAttrStr(o,n) PyObject_GetAttr(o,n)
#endif

/* GetBuiltinName.proto */
static PyObject *__Pyx_GetBuiltinName(PyObject *name);

/* RaiseArgTupleInvalid.proto */
static void __Pyx_RaiseArgtupleInvalid(const char* func_name, int exact,
    Py_ssize_t num_min, Py_ssize_t num_max, Py_ssize_t num_found);

/* KeywordStringCheck.proto */
static int __Pyx_CheckKeywordStrings(PyObject *kwdict, const char* function_name, int kw_allowed);

/* RaiseDoubleKeywords.proto */
static void __Pyx_RaiseDoubleKeywordsError(const char* func_name, PyObject* kw_name);

/* ParseKeywords.proto */
static int __Pyx_ParseOptionalKeywords(PyObject *kwds, PyObject **argnames[],\
    PyObject *kwds2, PyObject *values[], Py_ssize_t num_pos_args,\
    const char* function_name);

/* PyObjectCall.proto */
#if CYTHON_COMPILING_IN_CPYTHON
static CYTHON_INLINE PyObject* __Pyx_PyObject_Call(PyObject *func, PyObject *arg, PyObject *kw);
#else
#define __Pyx_PyObject_Call(func, arg, kw) PyObject_Call(func, arg, kw)
#endif

/* PyThreadStateGet.proto */
#if CYTHON_FAST_THREAD_STATE
#define __Pyx_PyThreadState_declare  PyThreadState *__pyx_tstate;
#define __Pyx_PyThreadState_assign  __pyx_tstate = __Pyx_PyThreadState_Current;
#define __Pyx_PyErr_Occurred()  __pyx_tstate->curexc_type
#else
#define __Pyx_PyThreadState_declare
#define __Pyx_PyThreadState_assign
#define __Pyx_PyErr_Occurred()  PyErr_Occurred()
#endif

/* PyErrFetchRestore.proto */
#if CYTHON_FAST_THREAD_STATE
#define __Pyx_PyErr_Clear() __Pyx_ErrRestore(NULL, NULL, NULL)
#define __Pyx_ErrRestoreWithState(type, value, tb)  __Pyx_ErrRestoreInState(PyThreadState_GET(), type, value, tb)
#define __Pyx_ErrFetchWithState(type, value, tb)    __Pyx_ErrFetchInState(PyThreadState_GET(), type, value, tb)
#define __Pyx_ErrRestore(type, value, tb)  __Pyx_ErrRestoreInState(__pyx_tstate, type, value, tb)
#define __Pyx_ErrFetch(type, value, tb)    __Pyx_ErrFetchInState(__pyx_tstate, type, value, tb)
static CYTHON_INLINE void __Pyx_ErrRestoreInState(PyThreadState *tstate, PyObject *type, PyObject *value, PyObject *tb);
static CYTHON_INLINE void __Pyx_ErrFetchInState(PyThreadState *tstate, PyObject **type, PyObject **value, PyObject **tb);
#if CYTHON_COMPILING_IN_CPYTHON
#define __Pyx_PyErr_SetNone(exc) (Py_INCREF(exc), __Pyx_ErrRestore((exc), NULL, NULL))
#else
#define __Pyx_PyErr_SetNone(exc) PyErr_SetNone(exc)
#endif
#else
#define __Pyx_PyErr_Clear() PyErr_Clear()
#define __Pyx_PyErr_SetNone(exc) PyErr_SetNone(exc)
#define __Pyx_ErrRestoreWithState(type, value, tb)  PyErr_Restore(type, value, tb)
#define __Pyx_ErrFetchWithState(type, value, tb)  PyErr_Fetch(type, value, tb)
#define __Pyx_ErrRestoreInState(tstate, type, value, tb)  PyErr_Restore(type, value, tb)
#define __Pyx_ErrFetchInState(tstate, type, value, tb)  PyErr_Fetch(type, value, tb)
#define __Pyx_ErrRestore(type, value, tb)  PyErr_Restore(type, value, tb)
#define __Pyx_ErrFetch(type, value, tb)  PyErr_Fetch(type, value, tb)
#endif

/* RaiseException.proto */
static void __Pyx_Raise(PyObject *type, PyObject *value, PyObject *tb, PyObject *cause);

/* PyObject_GenericGetAttrNoDict.proto */
#if CYTHON_USE_TYPE_SLOTS && CYTHON_USE_PYTYPE_LOOKUP && PY_VERSION_HEX < 0x03070000
static CYTHON_INLINE PyObject* __Pyx_PyObject_GenericGetAttrNoDict(PyObject* obj, PyObject* attr_name);
#else
#define __Pyx_PyObject_GenericGetAttrNoDict PyObject_GenericGetAttr
#endif

/* PyObject_GenericGetAttr.proto */
#if CYTHON_USE_TYPE_SLOTS && CYTHON_USE_PYTYPE_LOOKUP && PY_VERSION_HEX < 0x03070000
static PyObject* __Pyx_PyObject_GenericGetAttr(PyObject* obj, PyObject* attr_name);
#else
#define __Pyx_PyObject_GenericGetAttr PyObject_GenericGetAttr
#endif

/* PyErrExceptionMatches.proto */
#if CYTHON_FAST_THREAD_STATE
#define __Pyx_PyErr_ExceptionMatches(err) __Pyx_PyErr_ExceptionMatchesInState(__pyx_tstate, err)
static CYTHON_INLINE int __Pyx_PyErr_ExceptionMatchesInState(PyThreadState* tstate, PyObject* err);
#else
#define __Pyx_PyErr_ExceptionMatches(err)  PyErr_ExceptionMatches(err)
#endif

/* PyObjectGetAttrStrNoError.proto */
static CYTHON_INLINE PyObject* __Pyx_PyObject_GetAttrStrNoError(PyObject* obj, PyObject* attr_name);

/* SetupReduce.proto */
static int __Pyx_setup_reduce(PyObject* type_obj);

/* PyDictVersioning.proto */
#if CYTHON_USE_DICT_VERSIONS && CYTHON_USE_TYPE_SLOTS
#define __PYX_DICT_VERSION_INIT  ((PY_UINT64_T) -1)
#define __PYX_GET_DICT_VERSION(dict)  (((PyDictObject*)(dict))->ma_version_tag)
#define __PYX_UPDATE_DICT_CACHE(dict, value, cache_var, version_var)\
    (version_var) = __PYX_GET_DICT_VERSION(dict);\
    (cache_var) = (value);
#define __PYX_PY_DICT_LOOKUP_IF_MODIFIED(VAR, DICT, LOOKUP) {\
    static PY_UINT64_T __pyx_dict_version = 0;\
    static PyObject *__pyx_dict_cached_value = NULL;\
    if (likely(__PYX_GET_DICT_VERSION(DICT) == __pyx_dict_version)) {\
        (VAR) = __pyx_dict_cached_value;\
    } else {\
        (VAR) = __pyx_dict_cached_value = (LOOKUP);\
        __pyx_dict_version = __PYX_GET_DICT_VERSION(DICT);\
    }\
}
static CYTHON_INLINE PY_UINT64_T __Pyx_get_tp_dict_version(PyObject *obj);
static CYTHON_INLINE PY_UINT64_T __Pyx_get_object_dict_version(PyObject *obj);
static CYTHON_INLINE int __Pyx_object_dict_version_matches(PyObject* obj, PY_UINT64_T tp_dict_version, PY_UINT64_T obj_dict_version);
#else
#define __PYX_GET_DICT_VERSION(dict)  (0)
#define __PYX_UPDATE_DICT_CACHE(dict, value, cache_var, version_var)
#define __PYX_PY_DICT_LOOKUP_IF_MODIFIED(VAR, DICT, LOOKUP)  (VAR) = (LOOKUP);
#endif

/* CLineInTraceback.proto */
#ifdef CYTHON_CLINE_IN_TRACEBACK
#define __Pyx_CLineForTraceback(tstate, c_line)  (((CYTHON_CLINE_IN_TRACEBACK)) ? c_line : 0)
#else
static int __Pyx_CLineForTraceback(PyThreadState *tstate, int c_line);
#endif

/* CodeObjectCache.proto */
typedef struct {
    PyCodeObject* code_object;
    int code_line;
} __Pyx_CodeObjectCacheEntry;
struct __Pyx_CodeObjectCache {
    int count;
    int max_count;
    __Pyx_CodeObjectCacheEntry* entries;
};
static struct __Pyx_CodeObjectCache __pyx_code_cache = {0,0,NULL};
static int __pyx_bisect_code_objects(__Pyx_CodeObjectCacheEntry* entries, int count, int code_line);
static PyCodeObject *__pyx_find_code_object(int code_line);
static void __pyx_insert_code_object(int code_line, PyCodeObject* code_object);

/* AddTraceback.proto */
static void __Pyx_AddTraceback(const char *funcname, int c_line,
                               int py_line, const char *filename);

/* None.proto */
#include <new>

/* GCCDiagnostics.proto */
#if defined(__GNUC__) && (__GNUC__ > 4 || (__GNUC__ == 4 && __GNUC_MINOR__ >= 6))
#define __Pyx_HAS_GCC_DIAGNOSTIC
#endif

/* CppExceptionConversion.proto */
#ifndef __Pyx_CppExn2PyErr
#include <new>
#include <typeinfo>
#include <stdexcept>
#include <ios>
static void __Pyx_CppExn2PyErr() {
  try {
    if (PyErr_Occurred())
      ; // let the latest Python exn pass through and ignore the current one
    else
      throw;
  } catch (const std::bad_alloc& exn) {
    PyErr_SetString(PyExc_MemoryError, exn.what());
  } catch (const std::bad_cast& exn) {
    PyErr_SetString(PyExc_TypeError, exn.what());
  } catch (const std::bad_typeid& exn) {
    PyErr_SetString(PyExc_TypeError, exn.what());
  } catch (const std::domain_error& exn) {
    PyErr_SetString(PyExc_ValueError, exn.what());
  } catch (const std::invalid_argument& exn) {
    PyErr_SetString(PyExc_ValueError, exn.what());
  } catch (const std::ios_base::failure& exn) {
    PyErr_SetString(PyExc_IOError, exn.what());
  } catch (const std::out_of_range& exn) {
    PyErr_SetString(PyExc_IndexError, exn.what());
  } catch (const std::overflow_error& exn) {
    PyErr_SetString(PyExc_OverflowError, exn.what());
  } catch (const std::range_error& exn) {
    PyErr_SetString(PyExc_ArithmeticError, exn.what());
  } catch (const std::underflow_error& exn) {
    PyErr_SetString(PyExc_ArithmeticError, exn.what());
  } catch (const std::exception& exn) {
    PyErr_SetString(PyExc_RuntimeError, exn.what());
  }
  catch (...)
  {
    PyErr_SetString(PyExc_RuntimeError, "Unknown exception");
  }
}
#endif

/* CIntFromPy.proto */
static CYTHON_INLINE int __Pyx_PyInt_As_int(PyObject *);

/* CIntToPy.proto */
static CYTHON_INLINE PyObject* __Pyx_PyInt_From_int(int value);

/* CIntToPy.proto */
static CYTHON_INLINE PyObject* __Pyx_PyInt_From_long(long value);

/* CIntFromPy.proto */
static CYTHON_INLINE long __Pyx_PyInt_As_long(PyObject *);

/* FastTypeChecks.proto */
#if CYTHON_COMPILING_IN_CPYTHON
#define __Pyx_TypeCheck(obj, type) __Pyx_IsSubtype(Py_TYPE(obj), (PyTypeObject *)type)
static CYTHON_INLINE int __Pyx_IsSubtype(PyTypeObject *a, PyTypeObject *b);
static CYTHON_INLINE int __Pyx_PyErr_GivenExceptionMatches(PyObject *err, PyObject *type);
static CYTHON_INLINE int __Pyx_PyErr_GivenExceptionMatches2(PyObject *err, PyObject *type1, PyObject *type2);
#else
#define __Pyx_TypeCheck(obj, type) PyObject_TypeCheck(obj, (PyTypeObject *)type)
#define __Pyx_PyErr_GivenExceptionMatches(err, type) PyErr_GivenExceptionMatches(err, type)
#define __Pyx_PyErr_GivenExceptionMatches2(err, type1, type2) (PyErr_GivenExceptionMatches(err, type1) || PyErr_GivenExceptionMatches(err, type2))
#endif
#define __Pyx_PyException_Check(obj) __Pyx_TypeCheck(obj, PyExc_Exception)

/* CheckBinaryVersion.proto */
static int __Pyx_check_binary_version(void);

/* InitStrings.proto */
static int __Pyx_InitStrings(__Pyx_StringTabEntry *t);


/* Module declarations from 'solvers' */

/* Module declarations from 'runner' */
static PyTypeObject *__pyx_ptype_6runner_PySolver = 0;
#define __Pyx_MODULE_NAME "runner"
extern int __pyx_module_is_main_runner;
int __pyx_module_is_main_runner = 0;

/* Implementation of 'runner' */
static PyObject *__pyx_builtin_TypeError;
static const char __pyx_k_b[] = "b";
static const char __pyx_k_d[] = "d";
static const char __pyx_k_q[] = "q";
static const char __pyx_k_c1[] = "c1";
static const char __pyx_k_c2[] = "c2";
static const char __pyx_k_eta[] = "eta";
static const char __pyx_k_lam[] = "lam";
static const char __pyx_k_rho[] = "rho";
static const char __pyx_k_main[] = "__main__";
static const char __pyx_k_name[] = "__name__";
static const char __pyx_k_seed[] = "seed";
static const char __pyx_k_test[] = "__test__";
static const char __pyx_k_gamma[] = "gamma";
static const char __pyx_k_hyper[] = "hyper";
static const char __pyx_k_reduce[] = "__reduce__";
static const char __pyx_k_PySolver[] = "PySolver";
static const char __pyx_k_beta_lin[] = "beta_lin";
static const char __pyx_k_beta_max[] = "beta_max";
static const char __pyx_k_beta_min[] = "beta_min";
static const char __pyx_k_getstate[] = "__getstate__";
static const char __pyx_k_setstate[] = "__setstate__";
static const char __pyx_k_H_density[] = "H_density";
static const char __pyx_k_I_density[] = "I_density";
static const char __pyx_k_S_density[] = "S_density";
static const char __pyx_k_TypeError[] = "TypeError";
static const char __pyx_k_alpha_max[] = "alpha_max";
static const char __pyx_k_alpha_min[] = "alpha_min";
static const char __pyx_k_evo_steps[] = "evo_steps";
static const char __pyx_k_reduce_ex[] = "__reduce_ex__";
static const char __pyx_k_sigma_max[] = "sigma_max";
static const char __pyx_k_alpha_init[] = "alpha_init";
static const char __pyx_k_beta_value[] = "beta_value";
static const char __pyx_k_sigma_init[] = "sigma_init";
static const char __pyx_k_alpha_value[] = "alpha_value";
static const char __pyx_k_beta_scalar[] = "beta_scalar";
static const char __pyx_k_sigma_value[] = "sigma_value";
static const char __pyx_k_reduce_cython[] = "__reduce_cython__";
static const char __pyx_k_setstate_cython[] = "__setstate_cython__";
static const char __pyx_k_cline_in_traceback[] = "cline_in_traceback";
static const char __pyx_k_no_default___reduce___due_to_non[] = "no default __reduce__ due to non-trivial __cinit__";
static PyObject *__pyx_n_s_H_density;
static PyObject *__pyx_n_s_I_density;
static PyObject *__pyx_n_s_PySolver;
static PyObject *__pyx_n_s_S_density;
static PyObject *__pyx_n_s_TypeError;
static PyObject *__pyx_n_s_alpha_init;
static PyObject *__pyx_n_s_alpha_max;
static PyObject *__pyx_n_s_alpha_min;
static PyObject *__pyx_n_s_alpha_value;
static PyObject *__pyx_n_s_b;
static PyObject *__pyx_n_s_beta_lin;
static PyObject *__pyx_n_s_beta_max;
static PyObject *__pyx_n_s_beta_min;
static PyObject *__pyx_n_s_beta_scalar;
static PyObject *__pyx_n_s_beta_value;
static PyObject *__pyx_n_s_c1;
static PyObject *__pyx_n_s_c2;
static PyObject *__pyx_n_s_cline_in_traceback;
static PyObject *__pyx_n_s_d;
static PyObject *__pyx_n_s_eta;
static PyObject *__pyx_n_s_evo_steps;
static PyObject *__pyx_n_s_gamma;
static PyObject *__pyx_n_s_getstate;
static PyObject *__pyx_n_s_hyper;
static PyObject *__pyx_n_s_lam;
static PyObject *__pyx_n_s_main;
static PyObject *__pyx_n_s_name;
static PyObject *__pyx_kp_s_no_default___reduce___due_to_non;
static PyObject *__pyx_n_s_q;
static PyObject *__pyx_n_s_reduce;
static PyObject *__pyx_n_s_reduce_cython;
static PyObject *__pyx_n_s_reduce_ex;
static PyObject *__pyx_n_s_rho;
static PyObject *__pyx_n_s_seed;
static PyObject *__pyx_n_s_setstate;
static PyObject *__pyx_n_s_setstate_cython;
static PyObject *__pyx_n_s_sigma_init;
static PyObject *__pyx_n_s_sigma_max;
static PyObject *__pyx_n_s_sigma_value;
static PyObject *__pyx_n_s_test;
static int __pyx_pf_6runner_8PySolver___cinit__(struct __pyx_obj_6runner_PySolver *__pyx_v_self); /* proto */
static PyObject *__pyx_pf_6runner_8PySolver_2ad_dyn(struct __pyx_obj_6runner_PySolver *__pyx_v_self, float __pyx_v_beta_max, float __pyx_v_alpha_max, float __pyx_v_sigma_max, float __pyx_v_b, float __pyx_v_q, float __pyx_v_d, float __pyx_v_rho, float __pyx_v_eta, float __pyx_v_gamma, float __pyx_v_lam, float __pyx_v_c1, float __pyx_v_c2, float __pyx_v_hyper, int __pyx_v_seed, int __pyx_v_alpha_init, int __pyx_v_sigma_init); /* proto */
static PyObject *__pyx_pf_6runner_8PySolver_4alpha_ad_dyn(struct __pyx_obj_6runner_PySolver *__pyx_v_self, float __pyx_v_beta_max, float __pyx_v_alpha_max, float __pyx_v_sigma_max, float __pyx_v_b, float __pyx_v_q, float __pyx_v_d, float __pyx_v_rho, float __pyx_v_eta, float __pyx_v_gamma, float __pyx_v_lam, float __pyx_v_c1, float __pyx_v_c2, float __pyx_v_hyper, int __pyx_v_seed, int __pyx_v_alpha_init, int __pyx_v_sigma_init, float __pyx_v_S_density, float __pyx_v_I_density, float __pyx_v_H_density); /* proto */
static PyObject *__pyx_pf_6runner_8PySolver_6alpha_ad_dyn_v2(struct __pyx_obj_6runner_PySolver *__pyx_v_self, float __pyx_v_beta_max, float __pyx_v_beta_min, float __pyx_v_alpha_max, float __pyx_v_alpha_min, float __pyx_v_sigma_max, float __pyx_v_b, float __pyx_v_q, float __pyx_v_d, float __pyx_v_rho, float __pyx_v_eta, float __pyx_v_gamma, float __pyx_v_lam, float __pyx_v_c1, float __pyx_v_c2, float __pyx_v_beta_scalar, float __pyx_v_hyper, int __pyx_v_seed, int __pyx_v_alpha_init, int __pyx_v_sigma_init, float __pyx_v_S_density, float __pyx_v_I_density, float __pyx_v_H_density); /* proto */
static PyObject *__pyx_pf_6runner_8PySolver_8alpha_ad_dyn_v3(struct __pyx_obj_6runner_PySolver *__pyx_v_self, float __pyx_v_beta_max, float __pyx_v_alpha_max, float __pyx_v_sigma_max, float __pyx_v_b, float __pyx_v_q, float __pyx_v_d, float __pyx_v_rho, float __pyx_v_eta, float __pyx_v_gamma, float __pyx_v_lam, float __pyx_v_c1, float __pyx_v_c2, float __pyx_v_hyper, int __pyx_v_seed, int __pyx_v_alpha_init, int __pyx_v_sigma_init, float __pyx_v_S_density, float __pyx_v_I_density, float __pyx_v_H_density); /* proto */
static PyObject *__pyx_pf_6runner_8PySolver_10alpha_ad_dyn_v4(struct __pyx_obj_6runner_PySolver *__pyx_v_self, float __pyx_v_beta_max, float __pyx_v_beta_lin, float __pyx_v_alpha_max, float __pyx_v_alpha_min, float __pyx_v_sigma_max, float __pyx_v_b, float __pyx_v_q, float __pyx_v_d, float __pyx_v_rho, float __pyx_v_eta, float __pyx_v_gamma, float __pyx_v_lam, float __pyx_v_c1, float __pyx_v_c2, float __pyx_v_beta_scalar, float __pyx_v_hyper, int __pyx_v_seed, int __pyx_v_alpha_init, int __pyx_v_sigma_init, int __pyx_v_evo_steps, float __pyx_v_S_density, float __pyx_v_I_density, float __pyx_v_H_density); /* proto */
static PyObject *__pyx_pf_6runner_8PySolver_12eco_steady_state(struct __pyx_obj_6runner_PySolver *__pyx_v_self, float __pyx_v_beta_value, float __pyx_v_alpha_value, float __pyx_v_sigma_value, float __pyx_v_b, float __pyx_v_q, float __pyx_v_d, float __pyx_v_rho, float __pyx_v_eta, float __pyx_v_gamma, float __pyx_v_lam, float __pyx_v_c1, float __pyx_v_c2, float __pyx_v_hyper, int __pyx_v_seed, float __pyx_v_S_density, float __pyx_v_I_density, float __pyx_v_H_density); /* proto */
static PyObject *__pyx_pf_6runner_8PySolver_14__reduce_cython__(CYTHON_UNUSED struct __pyx_obj_6runner_PySolver *__pyx_v_self); /* proto */
static PyObject *__pyx_pf_6runner_8PySolver_16__setstate_cython__(CYTHON_UNUSED struct __pyx_obj_6runner_PySolver *__pyx_v_self, CYTHON_UNUSED PyObject *__pyx_v___pyx_state); /* proto */
static PyObject *__pyx_tp_new_6runner_PySolver(PyTypeObject *t, PyObject *a, PyObject *k); /*proto*/
static PyObject *__pyx_tuple_;
static PyObject *__pyx_tuple__2;
/* Late includes */

/* "runner.pyx":8
 *     cdef Quick_solver cpp_solver
 * 
 *     def __cinit__(self):             # <<<<<<<<<<<<<<
 *         self.cpp_solver = Quick_solver()
 *         return
 */

/* Python wrapper */
static int __pyx_pw_6runner_8PySolver_1__cinit__(PyObject *__pyx_v_self, PyObject *__pyx_args, PyObject *__pyx_kwds); /*proto*/
static int __pyx_pw_6runner_8PySolver_1__cinit__(PyObject *__pyx_v_self, PyObject *__pyx_args, PyObject *__pyx_kwds) {
  int __pyx_r;
  __Pyx_RefNannyDeclarations
  __Pyx_RefNannySetupContext("__cinit__ (wrapper)", 0);
  if (unlikely(PyTuple_GET_SIZE(__pyx_args) > 0)) {
    __Pyx_RaiseArgtupleInvalid("__cinit__", 1, 0, 0, PyTuple_GET_SIZE(__pyx_args)); return -1;}
  if (unlikely(__pyx_kwds) && unlikely(PyDict_Size(__pyx_kwds) > 0) && unlikely(!__Pyx_CheckKeywordStrings(__pyx_kwds, "__cinit__", 0))) return -1;
  __pyx_r = __pyx_pf_6runner_8PySolver___cinit__(((struct __pyx_obj_6runner_PySolver *)__pyx_v_self));

  /* function exit code */
  __Pyx_RefNannyFinishContext();
  return __pyx_r;
}

static int __pyx_pf_6runner_8PySolver___cinit__(struct __pyx_obj_6runner_PySolver *__pyx_v_self) {
  int __pyx_r;
  __Pyx_RefNannyDeclarations
  solvers::Quick_solver __pyx_t_1;
  int __pyx_lineno = 0;
  const char *__pyx_filename = NULL;
  int __pyx_clineno = 0;
  __Pyx_RefNannySetupContext("__cinit__", 0);

  /* "runner.pyx":9
 * 
 *     def __cinit__(self):
 *         self.cpp_solver = Quick_solver()             # <<<<<<<<<<<<<<
 *         return
 * 
 */
  try {
    __pyx_t_1 = solvers::Quick_solver();
  } catch(...) {
    __Pyx_CppExn2PyErr();
    __PYX_ERR(1, 9, __pyx_L1_error)
  }
  __pyx_v_self->cpp_solver = __pyx_t_1;

  /* "runner.pyx":10
 *     def __cinit__(self):
 *         self.cpp_solver = Quick_solver()
 *         return             # <<<<<<<<<<<<<<
 * 
 *     def ad_dyn(self, float beta_max, float alpha_max, float sigma_max, float b, float q, float d, float rho, float eta, float gamma, float lam, float c1, float c2, float hyper, int seed, int alpha_init, int sigma_init):
 */
  __pyx_r = 0;
  goto __pyx_L0;

  /* "runner.pyx":8
 *     cdef Quick_solver cpp_solver
 * 
 *     def __cinit__(self):             # <<<<<<<<<<<<<<
 *         self.cpp_solver = Quick_solver()
 *         return
 */

  /* function exit code */
  __pyx_L1_error:;
  __Pyx_AddTraceback("runner.PySolver.__cinit__", __pyx_clineno, __pyx_lineno, __pyx_filename);
  __pyx_r = -1;
  __pyx_L0:;
  __Pyx_RefNannyFinishContext();
  return __pyx_r;
}

/* "runner.pyx":12
 *         return
 * 
 *     def ad_dyn(self, float beta_max, float alpha_max, float sigma_max, float b, float q, float d, float rho, float eta, float gamma, float lam, float c1, float c2, float hyper, int seed, int alpha_init, int sigma_init):             # <<<<<<<<<<<<<<
 * 
 *         self.cpp_solver.ad_dyn(beta_max, alpha_max, sigma_max, b, q, d, rho, eta, gamma, lam, c1, c2, hyper, seed, alpha_init, sigma_init)
 */

/* Python wrapper */
static PyObject *__pyx_pw_6runner_8PySolver_3ad_dyn(PyObject *__pyx_v_self, PyObject *__pyx_args, PyObject *__pyx_kwds); /*proto*/
static PyObject *__pyx_pw_6runner_8PySolver_3ad_dyn(PyObject *__pyx_v_self, PyObject *__pyx_args, PyObject *__pyx_kwds) {
  float __pyx_v_beta_max;
  float __pyx_v_alpha_max;
  float __pyx_v_sigma_max;
  float __pyx_v_b;
  float __pyx_v_q;
  float __pyx_v_d;
  float __pyx_v_rho;
  float __pyx_v_eta;
  float __pyx_v_gamma;
  float __pyx_v_lam;
  float __pyx_v_c1;
  float __pyx_v_c2;
  float __pyx_v_hyper;
  int __pyx_v_seed;
  int __pyx_v_alpha_init;
  int __pyx_v_sigma_init;
  int __pyx_lineno = 0;
  const char *__pyx_filename = NULL;
  int __pyx_clineno = 0;
  PyObject *__pyx_r = 0;
  __Pyx_RefNannyDeclarations
  __Pyx_RefNannySetupContext("ad_dyn (wrapper)", 0);
  {
    static PyObject **__pyx_pyargnames[] = {&__pyx_n_s_beta_max,&__pyx_n_s_alpha_max,&__pyx_n_s_sigma_max,&__pyx_n_s_b,&__pyx_n_s_q,&__pyx_n_s_d,&__pyx_n_s_rho,&__pyx_n_s_eta,&__pyx_n_s_gamma,&__pyx_n_s_lam,&__pyx_n_s_c1,&__pyx_n_s_c2,&__pyx_n_s_hyper,&__pyx_n_s_seed,&__pyx_n_s_alpha_init,&__pyx_n_s_sigma_init,0};
    PyObject* values[16] = {0,0,0,0,0,0,0,0,0,0,0,0,0,0,0,0};
    if (unlikely(__pyx_kwds)) {
      Py_ssize_t kw_args;
      const Py_ssize_t pos_args = PyTuple_GET_SIZE(__pyx_args);
      switch (pos_args) {
        case 16: values[15] = PyTuple_GET_ITEM(__pyx_args, 15);
        CYTHON_FALLTHROUGH;
        case 15: values[14] = PyTuple_GET_ITEM(__pyx_args, 14);
        CYTHON_FALLTHROUGH;
        case 14: values[13] = PyTuple_GET_ITEM(__pyx_args, 13);
        CYTHON_FALLTHROUGH;
        case 13: values[12] = PyTuple_GET_ITEM(__pyx_args, 12);
        CYTHON_FALLTHROUGH;
        case 12: values[11] = PyTuple_GET_ITEM(__pyx_args, 11);
        CYTHON_FALLTHROUGH;
        case 11: values[10] = PyTuple_GET_ITEM(__pyx_args, 10);
        CYTHON_FALLTHROUGH;
        case 10: values[9] = PyTuple_GET_ITEM(__pyx_args, 9);
        CYTHON_FALLTHROUGH;
        case  9: values[8] = PyTuple_GET_ITEM(__pyx_args, 8);
        CYTHON_FALLTHROUGH;
        case  8: values[7] = PyTuple_GET_ITEM(__pyx_args, 7);
        CYTHON_FALLTHROUGH;
        case  7: values[6] = PyTuple_GET_ITEM(__pyx_args, 6);
        CYTHON_FALLTHROUGH;
        case  6: values[5] = PyTuple_GET_ITEM(__pyx_args, 5);
        CYTHON_FALLTHROUGH;
        case  5: values[4] = PyTuple_GET_ITEM(__pyx_args, 4);
        CYTHON_FALLTHROUGH;
        case  4: values[3] = PyTuple_GET_ITEM(__pyx_args, 3);
        CYTHON_FALLTHROUGH;
        case  3: values[2] = PyTuple_GET_ITEM(__pyx_args, 2);
        CYTHON_FALLTHROUGH;
        case  2: values[1] = PyTuple_GET_ITEM(__pyx_args, 1);
        CYTHON_FALLTHROUGH;
        case  1: values[0] = PyTuple_GET_ITEM(__pyx_args, 0);
        CYTHON_FALLTHROUGH;
        case  0: break;
        default: goto __pyx_L5_argtuple_error;
      }
      kw_args = PyDict_Size(__pyx_kwds);
      switch (pos_args) {
        case  0:
        if (likely((values[0] = __Pyx_PyDict_GetItemStr(__pyx_kwds, __pyx_n_s_beta_max)) != 0)) kw_args--;
        else goto __pyx_L5_argtuple_error;
        CYTHON_FALLTHROUGH;
        case  1:
        if (likely((values[1] = __Pyx_PyDict_GetItemStr(__pyx_kwds, __pyx_n_s_alpha_max)) != 0)) kw_args--;
        else {
          __Pyx_RaiseArgtupleInvalid("ad_dyn", 1, 16, 16, 1); __PYX_ERR(1, 12, __pyx_L3_error)
        }
        CYTHON_FALLTHROUGH;
        case  2:
        if (likely((values[2] = __Pyx_PyDict_GetItemStr(__pyx_kwds, __pyx_n_s_sigma_max)) != 0)) kw_args--;
        else {
          __Pyx_RaiseArgtupleInvalid("ad_dyn", 1, 16, 16, 2); __PYX_ERR(1, 12, __pyx_L3_error)
        }
        CYTHON_FALLTHROUGH;
        case  3:
        if (likely((values[3] = __Pyx_PyDict_GetItemStr(__pyx_kwds, __pyx_n_s_b)) != 0)) kw_args--;
        else {
          __Pyx_RaiseArgtupleInvalid("ad_dyn", 1, 16, 16, 3); __PYX_ERR(1, 12, __pyx_L3_error)
        }
        CYTHON_FALLTHROUGH;
        case  4:
        if (likely((values[4] = __Pyx_PyDict_GetItemStr(__pyx_kwds, __pyx_n_s_q)) != 0)) kw_args--;
        else {
          __Pyx_RaiseArgtupleInvalid("ad_dyn", 1, 16, 16, 4); __PYX_ERR(1, 12, __pyx_L3_error)
        }
        CYTHON_FALLTHROUGH;
        case  5:
        if (likely((values[5] = __Pyx_PyDict_GetItemStr(__pyx_kwds, __pyx_n_s_d)) != 0)) kw_args--;
        else {
          __Pyx_RaiseArgtupleInvalid("ad_dyn", 1, 16, 16, 5); __PYX_ERR(1, 12, __pyx_L3_error)
        }
        CYTHON_FALLTHROUGH;
        case  6:
        if (likely((values[6] = __Pyx_PyDict_GetItemStr(__pyx_kwds, __pyx_n_s_rho)) != 0)) kw_args--;
        else {
          __Pyx_RaiseArgtupleInvalid("ad_dyn", 1, 16, 16, 6); __PYX_ERR(1, 12, __pyx_L3_error)
        }
        CYTHON_FALLTHROUGH;
        case  7:
        if (likely((values[7] = __Pyx_PyDict_GetItemStr(__pyx_kwds, __pyx_n_s_eta)) != 0)) kw_args--;
        else {
          __Pyx_RaiseArgtupleInvalid("ad_dyn", 1, 16, 16, 7); __PYX_ERR(1, 12, __pyx_L3_error)
        }
        CYTHON_FALLTHROUGH;
        case  8:
        if (likely((values[8] = __Pyx_PyDict_GetItemStr(__pyx_kwds, __pyx_n_s_gamma)) != 0)) kw_args--;
        else {
          __Pyx_RaiseArgtupleInvalid("ad_dyn", 1, 16, 16, 8); __PYX_ERR(1, 12, __pyx_L3_error)
        }
        CYTHON_FALLTHROUGH;
        case  9:
        if (likely((values[9] = __Pyx_PyDict_GetItemStr(__pyx_kwds, __pyx_n_s_lam)) != 0)) kw_args--;
        else {
          __Pyx_RaiseArgtupleInvalid("ad_dyn", 1, 16, 16, 9); __PYX_ERR(1, 12, __pyx_L3_error)
        }
        CYTHON_FALLTHROUGH;
        case 10:
        if (likely((values[10] = __Pyx_PyDict_GetItemStr(__pyx_kwds, __pyx_n_s_c1)) != 0)) kw_args--;
        else {
          __Pyx_RaiseArgtupleInvalid("ad_dyn", 1, 16, 16, 10); __PYX_ERR(1, 12, __pyx_L3_error)
        }
        CYTHON_FALLTHROUGH;
        case 11:
        if (likely((values[11] = __Pyx_PyDict_GetItemStr(__pyx_kwds, __pyx_n_s_c2)) != 0)) kw_args--;
        else {
          __Pyx_RaiseArgtupleInvalid("ad_dyn", 1, 16, 16, 11); __PYX_ERR(1, 12, __pyx_L3_error)
        }
        CYTHON_FALLTHROUGH;
        case 12:
        if (likely((values[12] = __Pyx_PyDict_GetItemStr(__pyx_kwds, __pyx_n_s_hyper)) != 0)) kw_args--;
        else {
          __Pyx_RaiseArgtupleInvalid("ad_dyn", 1, 16, 16, 12); __PYX_ERR(1, 12, __pyx_L3_error)
        }
        CYTHON_FALLTHROUGH;
        case 13:
        if (likely((values[13] = __Pyx_PyDict_GetItemStr(__pyx_kwds, __pyx_n_s_seed)) != 0)) kw_args--;
        else {
          __Pyx_RaiseArgtupleInvalid("ad_dyn", 1, 16, 16, 13); __PYX_ERR(1, 12, __pyx_L3_error)
        }
        CYTHON_FALLTHROUGH;
        case 14:
        if (likely((values[14] = __Pyx_PyDict_GetItemStr(__pyx_kwds, __pyx_n_s_alpha_init)) != 0)) kw_args--;
        else {
          __Pyx_RaiseArgtupleInvalid("ad_dyn", 1, 16, 16, 14); __PYX_ERR(1, 12, __pyx_L3_error)
        }
        CYTHON_FALLTHROUGH;
        case 15:
        if (likely((values[15] = __Pyx_PyDict_GetItemStr(__pyx_kwds, __pyx_n_s_sigma_init)) != 0)) kw_args--;
        else {
          __Pyx_RaiseArgtupleInvalid("ad_dyn", 1, 16, 16, 15); __PYX_ERR(1, 12, __pyx_L3_error)
        }
      }
      if (unlikely(kw_args > 0)) {
        if (unlikely(__Pyx_ParseOptionalKeywords(__pyx_kwds, __pyx_pyargnames, 0, values, pos_args, "ad_dyn") < 0)) __PYX_ERR(1, 12, __pyx_L3_error)
      }
    } else if (PyTuple_GET_SIZE(__pyx_args) != 16) {
      goto __pyx_L5_argtuple_error;
    } else {
      values[0] = PyTuple_GET_ITEM(__pyx_args, 0);
      values[1] = PyTuple_GET_ITEM(__pyx_args, 1);
      values[2] = PyTuple_GET_ITEM(__pyx_args, 2);
      values[3] = PyTuple_GET_ITEM(__pyx_args, 3);
      values[4] = PyTuple_GET_ITEM(__pyx_args, 4);
      values[5] = PyTuple_GET_ITEM(__pyx_args, 5);
      values[6] = PyTuple_GET_ITEM(__pyx_args, 6);
      values[7] = PyTuple_GET_ITEM(__pyx_args, 7);
      values[8] = PyTuple_GET_ITEM(__pyx_args, 8);
      values[9] = PyTuple_GET_ITEM(__pyx_args, 9);
      values[10] = PyTuple_GET_ITEM(__pyx_args, 10);
      values[11] = PyTuple_GET_ITEM(__pyx_args, 11);
      values[12] = PyTuple_GET_ITEM(__pyx_args, 12);
      values[13] = PyTuple_GET_ITEM(__pyx_args, 13);
      values[14] = PyTuple_GET_ITEM(__pyx_args, 14);
      values[15] = PyTuple_GET_ITEM(__pyx_args, 15);
    }
    __pyx_v_beta_max = __pyx_PyFloat_AsFloat(values[0]); if (unlikely((__pyx_v_beta_max == (float)-1) && PyErr_Occurred())) __PYX_ERR(1, 12, __pyx_L3_error)
    __pyx_v_alpha_max = __pyx_PyFloat_AsFloat(values[1]); if (unlikely((__pyx_v_alpha_max == (float)-1) && PyErr_Occurred())) __PYX_ERR(1, 12, __pyx_L3_error)
    __pyx_v_sigma_max = __pyx_PyFloat_AsFloat(values[2]); if (unlikely((__pyx_v_sigma_max == (float)-1) && PyErr_Occurred())) __PYX_ERR(1, 12, __pyx_L3_error)
    __pyx_v_b = __pyx_PyFloat_AsFloat(values[3]); if (unlikely((__pyx_v_b == (float)-1) && PyErr_Occurred())) __PYX_ERR(1, 12, __pyx_L3_error)
    __pyx_v_q = __pyx_PyFloat_AsFloat(values[4]); if (unlikely((__pyx_v_q == (float)-1) && PyErr_Occurred())) __PYX_ERR(1, 12, __pyx_L3_error)
    __pyx_v_d = __pyx_PyFloat_AsFloat(values[5]); if (unlikely((__pyx_v_d == (float)-1) && PyErr_Occurred())) __PYX_ERR(1, 12, __pyx_L3_error)
    __pyx_v_rho = __pyx_PyFloat_AsFloat(values[6]); if (unlikely((__pyx_v_rho == (float)-1) && PyErr_Occurred())) __PYX_ERR(1, 12, __pyx_L3_error)
    __pyx_v_eta = __pyx_PyFloat_AsFloat(values[7]); if (unlikely((__pyx_v_eta == (float)-1) && PyErr_Occurred())) __PYX_ERR(1, 12, __pyx_L3_error)
    __pyx_v_gamma = __pyx_PyFloat_AsFloat(values[8]); if (unlikely((__pyx_v_gamma == (float)-1) && PyErr_Occurred())) __PYX_ERR(1, 12, __pyx_L3_error)
    __pyx_v_lam = __pyx_PyFloat_AsFloat(values[9]); if (unlikely((__pyx_v_lam == (float)-1) && PyErr_Occurred())) __PYX_ERR(1, 12, __pyx_L3_error)
    __pyx_v_c1 = __pyx_PyFloat_AsFloat(values[10]); if (unlikely((__pyx_v_c1 == (float)-1) && PyErr_Occurred())) __PYX_ERR(1, 12, __pyx_L3_error)
    __pyx_v_c2 = __pyx_PyFloat_AsFloat(values[11]); if (unlikely((__pyx_v_c2 == (float)-1) && PyErr_Occurred())) __PYX_ERR(1, 12, __pyx_L3_error)
    __pyx_v_hyper = __pyx_PyFloat_AsFloat(values[12]); if (unlikely((__pyx_v_hyper == (float)-1) && PyErr_Occurred())) __PYX_ERR(1, 12, __pyx_L3_error)
    __pyx_v_seed = __Pyx_PyInt_As_int(values[13]); if (unlikely((__pyx_v_seed == (int)-1) && PyErr_Occurred())) __PYX_ERR(1, 12, __pyx_L3_error)
    __pyx_v_alpha_init = __Pyx_PyInt_As_int(values[14]); if (unlikely((__pyx_v_alpha_init == (int)-1) && PyErr_Occurred())) __PYX_ERR(1, 12, __pyx_L3_error)
    __pyx_v_sigma_init = __Pyx_PyInt_As_int(values[15]); if (unlikely((__pyx_v_sigma_init == (int)-1) && PyErr_Occurred())) __PYX_ERR(1, 12, __pyx_L3_error)
  }
  goto __pyx_L4_argument_unpacking_done;
  __pyx_L5_argtuple_error:;
  __Pyx_RaiseArgtupleInvalid("ad_dyn", 1, 16, 16, PyTuple_GET_SIZE(__pyx_args)); __PYX_ERR(1, 12, __pyx_L3_error)
  __pyx_L3_error:;
  __Pyx_AddTraceback("runner.PySolver.ad_dyn", __pyx_clineno, __pyx_lineno, __pyx_filename);
  __Pyx_RefNannyFinishContext();
  return NULL;
  __pyx_L4_argument_unpacking_done:;
  __pyx_r = __pyx_pf_6runner_8PySolver_2ad_dyn(((struct __pyx_obj_6runner_PySolver *)__pyx_v_self), __pyx_v_beta_max, __pyx_v_alpha_max, __pyx_v_sigma_max, __pyx_v_b, __pyx_v_q, __pyx_v_d, __pyx_v_rho, __pyx_v_eta, __pyx_v_gamma, __pyx_v_lam, __pyx_v_c1, __pyx_v_c2, __pyx_v_hyper, __pyx_v_seed, __pyx_v_alpha_init, __pyx_v_sigma_init);

  /* function exit code */
  __Pyx_RefNannyFinishContext();
  return __pyx_r;
}

static PyObject *__pyx_pf_6runner_8PySolver_2ad_dyn(struct __pyx_obj_6runner_PySolver *__pyx_v_self, float __pyx_v_beta_max, float __pyx_v_alpha_max, float __pyx_v_sigma_max, float __pyx_v_b, float __pyx_v_q, float __pyx_v_d, float __pyx_v_rho, float __pyx_v_eta, float __pyx_v_gamma, float __pyx_v_lam, float __pyx_v_c1, float __pyx_v_c2, float __pyx_v_hyper, int __pyx_v_seed, int __pyx_v_alpha_init, int __pyx_v_sigma_init) {
  PyObject *__pyx_r = NULL;
  __Pyx_RefNannyDeclarations
  __Pyx_RefNannySetupContext("ad_dyn", 0);

  /* "runner.pyx":14
 *     def ad_dyn(self, float beta_max, float alpha_max, float sigma_max, float b, float q, float d, float rho, float eta, float gamma, float lam, float c1, float c2, float hyper, int seed, int alpha_init, int sigma_init):
 * 
 *         self.cpp_solver.ad_dyn(beta_max, alpha_max, sigma_max, b, q, d, rho, eta, gamma, lam, c1, c2, hyper, seed, alpha_init, sigma_init)             # <<<<<<<<<<<<<<
 * 
 *         return
 */
  (void)(__pyx_v_self->cpp_solver.ad_dyn(__pyx_v_beta_max, __pyx_v_alpha_max, __pyx_v_sigma_max, __pyx_v_b, __pyx_v_q, __pyx_v_d, __pyx_v_rho, __pyx_v_eta, __pyx_v_gamma, __pyx_v_lam, __pyx_v_c1, __pyx_v_c2, __pyx_v_hyper, __pyx_v_seed, __pyx_v_alpha_init, __pyx_v_sigma_init));

  /* "runner.pyx":16
 *         self.cpp_solver.ad_dyn(beta_max, alpha_max, sigma_max, b, q, d, rho, eta, gamma, lam, c1, c2, hyper, seed, alpha_init, sigma_init)
 * 
 *         return             # <<<<<<<<<<<<<<
 * 
 *     def alpha_ad_dyn(self, float beta_max, float alpha_max, float sigma_max, float b, float q, float d, float rho, float eta, float gamma, float lam, float c1, float c2, float hyper, int seed, int alpha_init, int sigma_init, float S_density = 4.0, float I_density = 4.0, float H_density = 4.0):
 */
  __Pyx_XDECREF(__pyx_r);
  __pyx_r = Py_None; __Pyx_INCREF(Py_None);
  goto __pyx_L0;

  /* "runner.pyx":12
 *         return
 * 
 *     def ad_dyn(self, float beta_max, float alpha_max, float sigma_max, float b, float q, float d, float rho, float eta, float gamma, float lam, float c1, float c2, float hyper, int seed, int alpha_init, int sigma_init):             # <<<<<<<<<<<<<<
 * 
 *         self.cpp_solver.ad_dyn(beta_max, alpha_max, sigma_max, b, q, d, rho, eta, gamma, lam, c1, c2, hyper, seed, alpha_init, sigma_init)
 */

  /* function exit code */
  __pyx_L0:;
  __Pyx_XGIVEREF(__pyx_r);
  __Pyx_RefNannyFinishContext();
  return __pyx_r;
}

/* "runner.pyx":18
 *         return
 * 
 *     def alpha_ad_dyn(self, float beta_max, float alpha_max, float sigma_max, float b, float q, float d, float rho, float eta, float gamma, float lam, float c1, float c2, float hyper, int seed, int alpha_init, int sigma_init, float S_density = 4.0, float I_density = 4.0, float H_density = 4.0):             # <<<<<<<<<<<<<<
 * 
 *         self.cpp_solver.alpha_evo_only(beta_max, alpha_max, sigma_max, b, q, d, rho, eta, gamma, lam, c1, c2, hyper, seed, alpha_init, sigma_init, S_density, I_density, H_density)
 */

/* Python wrapper */
static PyObject *__pyx_pw_6runner_8PySolver_5alpha_ad_dyn(PyObject *__pyx_v_self, PyObject *__pyx_args, PyObject *__pyx_kwds); /*proto*/
static PyObject *__pyx_pw_6runner_8PySolver_5alpha_ad_dyn(PyObject *__pyx_v_self, PyObject *__pyx_args, PyObject *__pyx_kwds) {
  float __pyx_v_beta_max;
  float __pyx_v_alpha_max;
  float __pyx_v_sigma_max;
  float __pyx_v_b;
  float __pyx_v_q;
  float __pyx_v_d;
  float __pyx_v_rho;
  float __pyx_v_eta;
  float __pyx_v_gamma;
  float __pyx_v_lam;
  float __pyx_v_c1;
  float __pyx_v_c2;
  float __pyx_v_hyper;
  int __pyx_v_seed;
  int __pyx_v_alpha_init;
  int __pyx_v_sigma_init;
  float __pyx_v_S_density;
  float __pyx_v_I_density;
  float __pyx_v_H_density;
  int __pyx_lineno = 0;
  const char *__pyx_filename = NULL;
  int __pyx_clineno = 0;
  PyObject *__pyx_r = 0;
  __Pyx_RefNannyDeclarations
  __Pyx_RefNannySetupContext("alpha_ad_dyn (wrapper)", 0);
  {
    static PyObject **__pyx_pyargnames[] = {&__pyx_n_s_beta_max,&__pyx_n_s_alpha_max,&__pyx_n_s_sigma_max,&__pyx_n_s_b,&__pyx_n_s_q,&__pyx_n_s_d,&__pyx_n_s_rho,&__pyx_n_s_eta,&__pyx_n_s_gamma,&__pyx_n_s_lam,&__pyx_n_s_c1,&__pyx_n_s_c2,&__pyx_n_s_hyper,&__pyx_n_s_seed,&__pyx_n_s_alpha_init,&__pyx_n_s_sigma_init,&__pyx_n_s_S_density,&__pyx_n_s_I_density,&__pyx_n_s_H_density,0};
    PyObject* values[19] = {0,0,0,0,0,0,0,0,0,0,0,0,0,0,0,0,0,0,0};
    if (unlikely(__pyx_kwds)) {
      Py_ssize_t kw_args;
      const Py_ssize_t pos_args = PyTuple_GET_SIZE(__pyx_args);
      switch (pos_args) {
        case 19: values[18] = PyTuple_GET_ITEM(__pyx_args, 18);
        CYTHON_FALLTHROUGH;
        case 18: values[17] = PyTuple_GET_ITEM(__pyx_args, 17);
        CYTHON_FALLTHROUGH;
        case 17: values[16] = PyTuple_GET_ITEM(__pyx_args, 16);
        CYTHON_FALLTHROUGH;
        case 16: values[15] = PyTuple_GET_ITEM(__pyx_args, 15);
        CYTHON_FALLTHROUGH;
        case 15: values[14] = PyTuple_GET_ITEM(__pyx_args, 14);
        CYTHON_FALLTHROUGH;
        case 14: values[13] = PyTuple_GET_ITEM(__pyx_args, 13);
        CYTHON_FALLTHROUGH;
        case 13: values[12] = PyTuple_GET_ITEM(__pyx_args, 12);
        CYTHON_FALLTHROUGH;
        case 12: values[11] = PyTuple_GET_ITEM(__pyx_args, 11);
        CYTHON_FALLTHROUGH;
        case 11: values[10] = PyTuple_GET_ITEM(__pyx_args, 10);
        CYTHON_FALLTHROUGH;
        case 10: values[9] = PyTuple_GET_ITEM(__pyx_args, 9);
        CYTHON_FALLTHROUGH;
        case  9: values[8] = PyTuple_GET_ITEM(__pyx_args, 8);
        CYTHON_FALLTHROUGH;
        case  8: values[7] = PyTuple_GET_ITEM(__pyx_args, 7);
        CYTHON_FALLTHROUGH;
        case  7: values[6] = PyTuple_GET_ITEM(__pyx_args, 6);
        CYTHON_FALLTHROUGH;
        case  6: values[5] = PyTuple_GET_ITEM(__pyx_args, 5);
        CYTHON_FALLTHROUGH;
        case  5: values[4] = PyTuple_GET_ITEM(__pyx_args, 4);
        CYTHON_FALLTHROUGH;
        case  4: values[3] = PyTuple_GET_ITEM(__pyx_args, 3);
        CYTHON_FALLTHROUGH;
        case  3: values[2] = PyTuple_GET_ITEM(__pyx_args, 2);
        CYTHON_FALLTHROUGH;
        case  2: values[1] = PyTuple_GET_ITEM(__pyx_args, 1);
        CYTHON_FALLTHROUGH;
        case  1: values[0] = PyTuple_GET_ITEM(__pyx_args, 0);
        CYTHON_FALLTHROUGH;
        case  0: break;
        default: goto __pyx_L5_argtuple_error;
      }
      kw_args = PyDict_Size(__pyx_kwds);
      switch (pos_args) {
        case  0:
        if (likely((values[0] = __Pyx_PyDict_GetItemStr(__pyx_kwds, __pyx_n_s_beta_max)) != 0)) kw_args--;
        else goto __pyx_L5_argtuple_error;
        CYTHON_FALLTHROUGH;
        case  1:
        if (likely((values[1] = __Pyx_PyDict_GetItemStr(__pyx_kwds, __pyx_n_s_alpha_max)) != 0)) kw_args--;
        else {
          __Pyx_RaiseArgtupleInvalid("alpha_ad_dyn", 0, 16, 19, 1); __PYX_ERR(1, 18, __pyx_L3_error)
        }
        CYTHON_FALLTHROUGH;
        case  2:
        if (likely((values[2] = __Pyx_PyDict_GetItemStr(__pyx_kwds, __pyx_n_s_sigma_max)) != 0)) kw_args--;
        else {
          __Pyx_RaiseArgtupleInvalid("alpha_ad_dyn", 0, 16, 19, 2); __PYX_ERR(1, 18, __pyx_L3_error)
        }
        CYTHON_FALLTHROUGH;
        case  3:
        if (likely((values[3] = __Pyx_PyDict_GetItemStr(__pyx_kwds, __pyx_n_s_b)) != 0)) kw_args--;
        else {
          __Pyx_RaiseArgtupleInvalid("alpha_ad_dyn", 0, 16, 19, 3); __PYX_ERR(1, 18, __pyx_L3_error)
        }
        CYTHON_FALLTHROUGH;
        case  4:
        if (likely((values[4] = __Pyx_PyDict_GetItemStr(__pyx_kwds, __pyx_n_s_q)) != 0)) kw_args--;
        else {
          __Pyx_RaiseArgtupleInvalid("alpha_ad_dyn", 0, 16, 19, 4); __PYX_ERR(1, 18, __pyx_L3_error)
        }
        CYTHON_FALLTHROUGH;
        case  5:
        if (likely((values[5] = __Pyx_PyDict_GetItemStr(__pyx_kwds, __pyx_n_s_d)) != 0)) kw_args--;
        else {
          __Pyx_RaiseArgtupleInvalid("alpha_ad_dyn", 0, 16, 19, 5); __PYX_ERR(1, 18, __pyx_L3_error)
        }
        CYTHON_FALLTHROUGH;
        case  6:
        if (likely((values[6] = __Pyx_PyDict_GetItemStr(__pyx_kwds, __pyx_n_s_rho)) != 0)) kw_args--;
        else {
          __Pyx_RaiseArgtupleInvalid("alpha_ad_dyn", 0, 16, 19, 6); __PYX_ERR(1, 18, __pyx_L3_error)
        }
        CYTHON_FALLTHROUGH;
        case  7:
        if (likely((values[7] = __Pyx_PyDict_GetItemStr(__pyx_kwds, __pyx_n_s_eta)) != 0)) kw_args--;
        else {
          __Pyx_RaiseArgtupleInvalid("alpha_ad_dyn", 0, 16, 19, 7); __PYX_ERR(1, 18, __pyx_L3_error)
        }
        CYTHON_FALLTHROUGH;
        case  8:
        if (likely((values[8] = __Pyx_PyDict_GetItemStr(__pyx_kwds, __pyx_n_s_gamma)) != 0)) kw_args--;
        else {
          __Pyx_RaiseArgtupleInvalid("alpha_ad_dyn", 0, 16, 19, 8); __PYX_ERR(1, 18, __pyx_L3_error)
        }
        CYTHON_FALLTHROUGH;
        case  9:
        if (likely((values[9] = __Pyx_PyDict_GetItemStr(__pyx_kwds, __pyx_n_s_lam)) != 0)) kw_args--;
        else {
          __Pyx_RaiseArgtupleInvalid("alpha_ad_dyn", 0, 16, 19, 9); __PYX_ERR(1, 18, __pyx_L3_error)
        }
        CYTHON_FALLTHROUGH;
        case 10:
        if (likely((values[10] = __Pyx_PyDict_GetItemStr(__pyx_kwds, __pyx_n_s_c1)) != 0)) kw_args--;
        else {
          __Pyx_RaiseArgtupleInvalid("alpha_ad_dyn", 0, 16, 19, 10); __PYX_ERR(1, 18, __pyx_L3_error)
        }
        CYTHON_FALLTHROUGH;
        case 11:
        if (likely((values[11] = __Pyx_PyDict_GetItemStr(__pyx_kwds, __pyx_n_s_c2)) != 0)) kw_args--;
        else {
          __Pyx_RaiseArgtupleInvalid("alpha_ad_dyn", 0, 16, 19, 11); __PYX_ERR(1, 18, __pyx_L3_error)
        }
        CYTHON_FALLTHROUGH;
        case 12:
        if (likely((values[12] = __Pyx_PyDict_GetItemStr(__pyx_kwds, __pyx_n_s_hyper)) != 0)) kw_args--;
        else {
          __Pyx_RaiseArgtupleInvalid("alpha_ad_dyn", 0, 16, 19, 12); __PYX_ERR(1, 18, __pyx_L3_error)
        }
        CYTHON_FALLTHROUGH;
        case 13:
        if (likely((values[13] = __Pyx_PyDict_GetItemStr(__pyx_kwds, __pyx_n_s_seed)) != 0)) kw_args--;
        else {
          __Pyx_RaiseArgtupleInvalid("alpha_ad_dyn", 0, 16, 19, 13); __PYX_ERR(1, 18, __pyx_L3_error)
        }
        CYTHON_FALLTHROUGH;
        case 14:
        if (likely((values[14] = __Pyx_PyDict_GetItemStr(__pyx_kwds, __pyx_n_s_alpha_init)) != 0)) kw_args--;
        else {
          __Pyx_RaiseArgtupleInvalid("alpha_ad_dyn", 0, 16, 19, 14); __PYX_ERR(1, 18, __pyx_L3_error)
        }
        CYTHON_FALLTHROUGH;
        case 15:
        if (likely((values[15] = __Pyx_PyDict_GetItemStr(__pyx_kwds, __pyx_n_s_sigma_init)) != 0)) kw_args--;
        else {
          __Pyx_RaiseArgtupleInvalid("alpha_ad_dyn", 0, 16, 19, 15); __PYX_ERR(1, 18, __pyx_L3_error)
        }
        CYTHON_FALLTHROUGH;
        case 16:
        if (kw_args > 0) {
          PyObject* value = __Pyx_PyDict_GetItemStr(__pyx_kwds, __pyx_n_s_S_density);
          if (value) { values[16] = value; kw_args--; }
        }
        CYTHON_FALLTHROUGH;
        case 17:
        if (kw_args > 0) {
          PyObject* value = __Pyx_PyDict_GetItemStr(__pyx_kwds, __pyx_n_s_I_density);
          if (value) { values[17] = value; kw_args--; }
        }
        CYTHON_FALLTHROUGH;
        case 18:
        if (kw_args > 0) {
          PyObject* value = __Pyx_PyDict_GetItemStr(__pyx_kwds, __pyx_n_s_H_density);
          if (value) { values[18] = value; kw_args--; }
        }
      }
      if (unlikely(kw_args > 0)) {
        if (unlikely(__Pyx_ParseOptionalKeywords(__pyx_kwds, __pyx_pyargnames, 0, values, pos_args, "alpha_ad_dyn") < 0)) __PYX_ERR(1, 18, __pyx_L3_error)
      }
    } else {
      switch (PyTuple_GET_SIZE(__pyx_args)) {
        case 19: values[18] = PyTuple_GET_ITEM(__pyx_args, 18);
        CYTHON_FALLTHROUGH;
        case 18: values[17] = PyTuple_GET_ITEM(__pyx_args, 17);
        CYTHON_FALLTHROUGH;
        case 17: values[16] = PyTuple_GET_ITEM(__pyx_args, 16);
        CYTHON_FALLTHROUGH;
        case 16: values[15] = PyTuple_GET_ITEM(__pyx_args, 15);
        values[14] = PyTuple_GET_ITEM(__pyx_args, 14);
        values[13] = PyTuple_GET_ITEM(__pyx_args, 13);
        values[12] = PyTuple_GET_ITEM(__pyx_args, 12);
        values[11] = PyTuple_GET_ITEM(__pyx_args, 11);
        values[10] = PyTuple_GET_ITEM(__pyx_args, 10);
        values[9] = PyTuple_GET_ITEM(__pyx_args, 9);
        values[8] = PyTuple_GET_ITEM(__pyx_args, 8);
        values[7] = PyTuple_GET_ITEM(__pyx_args, 7);
        values[6] = PyTuple_GET_ITEM(__pyx_args, 6);
        values[5] = PyTuple_GET_ITEM(__pyx_args, 5);
        values[4] = PyTuple_GET_ITEM(__pyx_args, 4);
        values[3] = PyTuple_GET_ITEM(__pyx_args, 3);
        values[2] = PyTuple_GET_ITEM(__pyx_args, 2);
        values[1] = PyTuple_GET_ITEM(__pyx_args, 1);
        values[0] = PyTuple_GET_ITEM(__pyx_args, 0);
        break;
        default: goto __pyx_L5_argtuple_error;
      }
    }
    __pyx_v_beta_max = __pyx_PyFloat_AsFloat(values[0]); if (unlikely((__pyx_v_beta_max == (float)-1) && PyErr_Occurred())) __PYX_ERR(1, 18, __pyx_L3_error)
    __pyx_v_alpha_max = __pyx_PyFloat_AsFloat(values[1]); if (unlikely((__pyx_v_alpha_max == (float)-1) && PyErr_Occurred())) __PYX_ERR(1, 18, __pyx_L3_error)
    __pyx_v_sigma_max = __pyx_PyFloat_AsFloat(values[2]); if (unlikely((__pyx_v_sigma_max == (float)-1) && PyErr_Occurred())) __PYX_ERR(1, 18, __pyx_L3_error)
    __pyx_v_b = __pyx_PyFloat_AsFloat(values[3]); if (unlikely((__pyx_v_b == (float)-1) && PyErr_Occurred())) __PYX_ERR(1, 18, __pyx_L3_error)
    __pyx_v_q = __pyx_PyFloat_AsFloat(values[4]); if (unlikely((__pyx_v_q == (float)-1) && PyErr_Occurred())) __PYX_ERR(1, 18, __pyx_L3_error)
    __pyx_v_d = __pyx_PyFloat_AsFloat(values[5]); if (unlikely((__pyx_v_d == (float)-1) && PyErr_Occurred())) __PYX_ERR(1, 18, __pyx_L3_error)
    __pyx_v_rho = __pyx_PyFloat_AsFloat(values[6]); if (unlikely((__pyx_v_rho == (float)-1) && PyErr_Occurred())) __PYX_ERR(1, 18, __pyx_L3_error)
    __pyx_v_eta = __pyx_PyFloat_AsFloat(values[7]); if (unlikely((__pyx_v_eta == (float)-1) && PyErr_Occurred())) __PYX_ERR(1, 18, __pyx_L3_error)
    __pyx_v_gamma = __pyx_PyFloat_AsFloat(values[8]); if (unlikely((__pyx_v_gamma == (float)-1) && PyErr_Occurred())) __PYX_ERR(1, 18, __pyx_L3_error)
    __pyx_v_lam = __pyx_PyFloat_AsFloat(values[9]); if (unlikely((__pyx_v_lam == (float)-1) && PyErr_Occurred())) __PYX_ERR(1, 18, __pyx_L3_error)
    __pyx_v_c1 = __pyx_PyFloat_AsFloat(values[10]); if (unlikely((__pyx_v_c1 == (float)-1) && PyErr_Occurred())) __PYX_ERR(1, 18, __pyx_L3_error)
    __pyx_v_c2 = __pyx_PyFloat_AsFloat(values[11]); if (unlikely((__pyx_v_c2 == (float)-1) && PyErr_Occurred())) __PYX_ERR(1, 18, __pyx_L3_error)
    __pyx_v_hyper = __pyx_PyFloat_AsFloat(values[12]); if (unlikely((__pyx_v_hyper == (float)-1) && PyErr_Occurred())) __PYX_ERR(1, 18, __pyx_L3_error)
    __pyx_v_seed = __Pyx_PyInt_As_int(values[13]); if (unlikely((__pyx_v_seed == (int)-1) && PyErr_Occurred())) __PYX_ERR(1, 18, __pyx_L3_error)
    __pyx_v_alpha_init = __Pyx_PyInt_As_int(values[14]); if (unlikely((__pyx_v_alpha_init == (int)-1) && PyErr_Occurred())) __PYX_ERR(1, 18, __pyx_L3_error)
    __pyx_v_sigma_init = __Pyx_PyInt_As_int(values[15]); if (unlikely((__pyx_v_sigma_init == (int)-1) && PyErr_Occurred())) __PYX_ERR(1, 18, __pyx_L3_error)
    if (values[16]) {
      __pyx_v_S_density = __pyx_PyFloat_AsFloat(values[16]); if (unlikely((__pyx_v_S_density == (float)-1) && PyErr_Occurred())) __PYX_ERR(1, 18, __pyx_L3_error)
    } else {
      __pyx_v_S_density = ((float)4.0);
    }
    if (values[17]) {
      __pyx_v_I_density = __pyx_PyFloat_AsFloat(values[17]); if (unlikely((__pyx_v_I_density == (float)-1) && PyErr_Occurred())) __PYX_ERR(1, 18, __pyx_L3_error)
    } else {
      __pyx_v_I_density = ((float)4.0);
    }
    if (values[18]) {
      __pyx_v_H_density = __pyx_PyFloat_AsFloat(values[18]); if (unlikely((__pyx_v_H_density == (float)-1) && PyErr_Occurred())) __PYX_ERR(1, 18, __pyx_L3_error)
    } else {
      __pyx_v_H_density = ((float)4.0);
    }
  }
  goto __pyx_L4_argument_unpacking_done;
  __pyx_L5_argtuple_error:;
  __Pyx_RaiseArgtupleInvalid("alpha_ad_dyn", 0, 16, 19, PyTuple_GET_SIZE(__pyx_args)); __PYX_ERR(1, 18, __pyx_L3_error)
  __pyx_L3_error:;
  __Pyx_AddTraceback("runner.PySolver.alpha_ad_dyn", __pyx_clineno, __pyx_lineno, __pyx_filename);
  __Pyx_RefNannyFinishContext();
  return NULL;
  __pyx_L4_argument_unpacking_done:;
  __pyx_r = __pyx_pf_6runner_8PySolver_4alpha_ad_dyn(((struct __pyx_obj_6runner_PySolver *)__pyx_v_self), __pyx_v_beta_max, __pyx_v_alpha_max, __pyx_v_sigma_max, __pyx_v_b, __pyx_v_q, __pyx_v_d, __pyx_v_rho, __pyx_v_eta, __pyx_v_gamma, __pyx_v_lam, __pyx_v_c1, __pyx_v_c2, __pyx_v_hyper, __pyx_v_seed, __pyx_v_alpha_init, __pyx_v_sigma_init, __pyx_v_S_density, __pyx_v_I_density, __pyx_v_H_density);

  /* function exit code */
  __Pyx_RefNannyFinishContext();
  return __pyx_r;
}

static PyObject *__pyx_pf_6runner_8PySolver_4alpha_ad_dyn(struct __pyx_obj_6runner_PySolver *__pyx_v_self, float __pyx_v_beta_max, float __pyx_v_alpha_max, float __pyx_v_sigma_max, float __pyx_v_b, float __pyx_v_q, float __pyx_v_d, float __pyx_v_rho, float __pyx_v_eta, float __pyx_v_gamma, float __pyx_v_lam, float __pyx_v_c1, float __pyx_v_c2, float __pyx_v_hyper, int __pyx_v_seed, int __pyx_v_alpha_init, int __pyx_v_sigma_init, float __pyx_v_S_density, float __pyx_v_I_density, float __pyx_v_H_density) {
  PyObject *__pyx_r = NULL;
  __Pyx_RefNannyDeclarations
  __Pyx_RefNannySetupContext("alpha_ad_dyn", 0);

  /* "runner.pyx":20
 *     def alpha_ad_dyn(self, float beta_max, float alpha_max, float sigma_max, float b, float q, float d, float rho, float eta, float gamma, float lam, float c1, float c2, float hyper, int seed, int alpha_init, int sigma_init, float S_density = 4.0, float I_density = 4.0, float H_density = 4.0):
 * 
 *         self.cpp_solver.alpha_evo_only(beta_max, alpha_max, sigma_max, b, q, d, rho, eta, gamma, lam, c1, c2, hyper, seed, alpha_init, sigma_init, S_density, I_density, H_density)             # <<<<<<<<<<<<<<
 * 
 *         return
 */
  __pyx_v_self->cpp_solver.alpha_evo_only(__pyx_v_beta_max, __pyx_v_alpha_max, __pyx_v_sigma_max, __pyx_v_b, __pyx_v_q, __pyx_v_d, __pyx_v_rho, __pyx_v_eta, __pyx_v_gamma, __pyx_v_lam, __pyx_v_c1, __pyx_v_c2, __pyx_v_hyper, __pyx_v_seed, __pyx_v_alpha_init, __pyx_v_sigma_init, __pyx_v_S_density, __pyx_v_I_density, __pyx_v_H_density);

  /* "runner.pyx":22
 *         self.cpp_solver.alpha_evo_only(beta_max, alpha_max, sigma_max, b, q, d, rho, eta, gamma, lam, c1, c2, hyper, seed, alpha_init, sigma_init, S_density, I_density, H_density)
 * 
 *         return             # <<<<<<<<<<<<<<
 * 
 *     def alpha_ad_dyn_v2(self, float beta_max, float beta_min, float alpha_max, float alpha_min, float sigma_max, float b, float q, float d, float rho, float eta, float gamma, float lam, float c1, float c2, float beta_scalar, float hyper, int seed, int alpha_init, int sigma_init, float S_density = 4.0, float I_density = 4.0, float H_density = 4.0):
 */
  __Pyx_XDECREF(__pyx_r);
  __pyx_r = Py_None; __Pyx_INCREF(Py_None);
  goto __pyx_L0;

  /* "runner.pyx":18
 *         return
 * 
 *     def alpha_ad_dyn(self, float beta_max, float alpha_max, float sigma_max, float b, float q, float d, float rho, float eta, float gamma, float lam, float c1, float c2, float hyper, int seed, int alpha_init, int sigma_init, float S_density = 4.0, float I_density = 4.0, float H_density = 4.0):             # <<<<<<<<<<<<<<
 * 
 *         self.cpp_solver.alpha_evo_only(beta_max, alpha_max, sigma_max, b, q, d, rho, eta, gamma, lam, c1, c2, hyper, seed, alpha_init, sigma_init, S_density, I_density, H_density)
 */

  /* function exit code */
  __pyx_L0:;
  __Pyx_XGIVEREF(__pyx_r);
  __Pyx_RefNannyFinishContext();
  return __pyx_r;
}

/* "runner.pyx":24
 *         return
 * 
 *     def alpha_ad_dyn_v2(self, float beta_max, float beta_min, float alpha_max, float alpha_min, float sigma_max, float b, float q, float d, float rho, float eta, float gamma, float lam, float c1, float c2, float beta_scalar, float hyper, int seed, int alpha_init, int sigma_init, float S_density = 4.0, float I_density = 4.0, float H_density = 4.0):             # <<<<<<<<<<<<<<
 * 
 * 
 */

/* Python wrapper */
static PyObject *__pyx_pw_6runner_8PySolver_7alpha_ad_dyn_v2(PyObject *__pyx_v_self, PyObject *__pyx_args, PyObject *__pyx_kwds); /*proto*/
static PyObject *__pyx_pw_6runner_8PySolver_7alpha_ad_dyn_v2(PyObject *__pyx_v_self, PyObject *__pyx_args, PyObject *__pyx_kwds) {
  float __pyx_v_beta_max;
  float __pyx_v_beta_min;
  float __pyx_v_alpha_max;
  float __pyx_v_alpha_min;
  float __pyx_v_sigma_max;
  float __pyx_v_b;
  float __pyx_v_q;
  float __pyx_v_d;
  float __pyx_v_rho;
  float __pyx_v_eta;
  float __pyx_v_gamma;
  float __pyx_v_lam;
  float __pyx_v_c1;
  float __pyx_v_c2;
  float __pyx_v_beta_scalar;
  float __pyx_v_hyper;
  int __pyx_v_seed;
  int __pyx_v_alpha_init;
  int __pyx_v_sigma_init;
  float __pyx_v_S_density;
  float __pyx_v_I_density;
  float __pyx_v_H_density;
  int __pyx_lineno = 0;
  const char *__pyx_filename = NULL;
  int __pyx_clineno = 0;
  PyObject *__pyx_r = 0;
  __Pyx_RefNannyDeclarations
  __Pyx_RefNannySetupContext("alpha_ad_dyn_v2 (wrapper)", 0);
  {
    static PyObject **__pyx_pyargnames[] = {&__pyx_n_s_beta_max,&__pyx_n_s_beta_min,&__pyx_n_s_alpha_max,&__pyx_n_s_alpha_min,&__pyx_n_s_sigma_max,&__pyx_n_s_b,&__pyx_n_s_q,&__pyx_n_s_d,&__pyx_n_s_rho,&__pyx_n_s_eta,&__pyx_n_s_gamma,&__pyx_n_s_lam,&__pyx_n_s_c1,&__pyx_n_s_c2,&__pyx_n_s_beta_scalar,&__pyx_n_s_hyper,&__pyx_n_s_seed,&__pyx_n_s_alpha_init,&__pyx_n_s_sigma_init,&__pyx_n_s_S_density,&__pyx_n_s_I_density,&__pyx_n_s_H_density,0};
    PyObject* values[22] = {0,0,0,0,0,0,0,0,0,0,0,0,0,0,0,0,0,0,0,0,0,0};
    if (unlikely(__pyx_kwds)) {
      Py_ssize_t kw_args;
      const Py_ssize_t pos_args = PyTuple_GET_SIZE(__pyx_args);
      switch (pos_args) {
        case 22: values[21] = PyTuple_GET_ITEM(__pyx_args, 21);
        CYTHON_FALLTHROUGH;
        case 21: values[20] = PyTuple_GET_ITEM(__pyx_args, 20);
        CYTHON_FALLTHROUGH;
        case 20: values[19] = PyTuple_GET_ITEM(__pyx_args, 19);
        CYTHON_FALLTHROUGH;
        case 19: values[18] = PyTuple_GET_ITEM(__pyx_args, 18);
        CYTHON_FALLTHROUGH;
        case 18: values[17] = PyTuple_GET_ITEM(__pyx_args, 17);
        CYTHON_FALLTHROUGH;
        case 17: values[16] = PyTuple_GET_ITEM(__pyx_args, 16);
        CYTHON_FALLTHROUGH;
        case 16: values[15] = PyTuple_GET_ITEM(__pyx_args, 15);
        CYTHON_FALLTHROUGH;
        case 15: values[14] = PyTuple_GET_ITEM(__pyx_args, 14);
        CYTHON_FALLTHROUGH;
        case 14: values[13] = PyTuple_GET_ITEM(__pyx_args, 13);
        CYTHON_FALLTHROUGH;
        case 13: values[12] = PyTuple_GET_ITEM(__pyx_args, 12);
        CYTHON_FALLTHROUGH;
        case 12: values[11] = PyTuple_GET_ITEM(__pyx_args, 11);
        CYTHON_FALLTHROUGH;
        case 11: values[10] = PyTuple_GET_ITEM(__pyx_args, 10);
        CYTHON_FALLTHROUGH;
        case 10: values[9] = PyTuple_GET_ITEM(__pyx_args, 9);
        CYTHON_FALLTHROUGH;
        case  9: values[8] = PyTuple_GET_ITEM(__pyx_args, 8);
        CYTHON_FALLTHROUGH;
        case  8: values[7] = PyTuple_GET_ITEM(__pyx_args, 7);
        CYTHON_FALLTHROUGH;
        case  7: values[6] = PyTuple_GET_ITEM(__pyx_args, 6);
        CYTHON_FALLTHROUGH;
        case  6: values[5] = PyTuple_GET_ITEM(__pyx_args, 5);
        CYTHON_FALLTHROUGH;
        case  5: values[4] = PyTuple_GET_ITEM(__pyx_args, 4);
        CYTHON_FALLTHROUGH;
        case  4: values[3] = PyTuple_GET_ITEM(__pyx_args, 3);
        CYTHON_FALLTHROUGH;
        case  3: values[2] = PyTuple_GET_ITEM(__pyx_args, 2);
        CYTHON_FALLTHROUGH;
        case  2: values[1] = PyTuple_GET_ITEM(__pyx_args, 1);
        CYTHON_FALLTHROUGH;
        case  1: values[0] = PyTuple_GET_ITEM(__pyx_args, 0);
        CYTHON_FALLTHROUGH;
        case  0: break;
        default: goto __pyx_L5_argtuple_error;
      }
      kw_args = PyDict_Size(__pyx_kwds);
      switch (pos_args) {
        case  0:
        if (likely((values[0] = __Pyx_PyDict_GetItemStr(__pyx_kwds, __pyx_n_s_beta_max)) != 0)) kw_args--;
        else goto __pyx_L5_argtuple_error;
        CYTHON_FALLTHROUGH;
        case  1:
        if (likely((values[1] = __Pyx_PyDict_GetItemStr(__pyx_kwds, __pyx_n_s_beta_min)) != 0)) kw_args--;
        else {
          __Pyx_RaiseArgtupleInvalid("alpha_ad_dyn_v2", 0, 19, 22, 1); __PYX_ERR(1, 24, __pyx_L3_error)
        }
        CYTHON_FALLTHROUGH;
        case  2:
        if (likely((values[2] = __Pyx_PyDict_GetItemStr(__pyx_kwds, __pyx_n_s_alpha_max)) != 0)) kw_args--;
        else {
          __Pyx_RaiseArgtupleInvalid("alpha_ad_dyn_v2", 0, 19, 22, 2); __PYX_ERR(1, 24, __pyx_L3_error)
        }
        CYTHON_FALLTHROUGH;
        case  3:
        if (likely((values[3] = __Pyx_PyDict_GetItemStr(__pyx_kwds, __pyx_n_s_alpha_min)) != 0)) kw_args--;
        else {
          __Pyx_RaiseArgtupleInvalid("alpha_ad_dyn_v2", 0, 19, 22, 3); __PYX_ERR(1, 24, __pyx_L3_error)
        }
        CYTHON_FALLTHROUGH;
        case  4:
        if (likely((values[4] = __Pyx_PyDict_GetItemStr(__pyx_kwds, __pyx_n_s_sigma_max)) != 0)) kw_args--;
        else {
          __Pyx_RaiseArgtupleInvalid("alpha_ad_dyn_v2", 0, 19, 22, 4); __PYX_ERR(1, 24, __pyx_L3_error)
        }
        CYTHON_FALLTHROUGH;
        case  5:
        if (likely((values[5] = __Pyx_PyDict_GetItemStr(__pyx_kwds, __pyx_n_s_b)) != 0)) kw_args--;
        else {
          __Pyx_RaiseArgtupleInvalid("alpha_ad_dyn_v2", 0, 19, 22, 5); __PYX_ERR(1, 24, __pyx_L3_error)
        }
        CYTHON_FALLTHROUGH;
        case  6:
        if (likely((values[6] = __Pyx_PyDict_GetItemStr(__pyx_kwds, __pyx_n_s_q)) != 0)) kw_args--;
        else {
          __Pyx_RaiseArgtupleInvalid("alpha_ad_dyn_v2", 0, 19, 22, 6); __PYX_ERR(1, 24, __pyx_L3_error)
        }
        CYTHON_FALLTHROUGH;
        case  7:
        if (likely((values[7] = __Pyx_PyDict_GetItemStr(__pyx_kwds, __pyx_n_s_d)) != 0)) kw_args--;
        else {
          __Pyx_RaiseArgtupleInvalid("alpha_ad_dyn_v2", 0, 19, 22, 7); __PYX_ERR(1, 24, __pyx_L3_error)
        }
        CYTHON_FALLTHROUGH;
        case  8:
        if (likely((values[8] = __Pyx_PyDict_GetItemStr(__pyx_kwds, __pyx_n_s_rho)) != 0)) kw_args--;
        else {
          __Pyx_RaiseArgtupleInvalid("alpha_ad_dyn_v2", 0, 19, 22, 8); __PYX_ERR(1, 24, __pyx_L3_error)
        }
        CYTHON_FALLTHROUGH;
        case  9:
        if (likely((values[9] = __Pyx_PyDict_GetItemStr(__pyx_kwds, __pyx_n_s_eta)) != 0)) kw_args--;
        else {
          __Pyx_RaiseArgtupleInvalid("alpha_ad_dyn_v2", 0, 19, 22, 9); __PYX_ERR(1, 24, __pyx_L3_error)
        }
        CYTHON_FALLTHROUGH;
        case 10:
        if (likely((values[10] = __Pyx_PyDict_GetItemStr(__pyx_kwds, __pyx_n_s_gamma)) != 0)) kw_args--;
        else {
          __Pyx_RaiseArgtupleInvalid("alpha_ad_dyn_v2", 0, 19, 22, 10); __PYX_ERR(1, 24, __pyx_L3_error)
        }
        CYTHON_FALLTHROUGH;
        case 11:
        if (likely((values[11] = __Pyx_PyDict_GetItemStr(__pyx_kwds, __pyx_n_s_lam)) != 0)) kw_args--;
        else {
          __Pyx_RaiseArgtupleInvalid("alpha_ad_dyn_v2", 0, 19, 22, 11); __PYX_ERR(1, 24, __pyx_L3_error)
        }
        CYTHON_FALLTHROUGH;
        case 12:
        if (likely((values[12] = __Pyx_PyDict_GetItemStr(__pyx_kwds, __pyx_n_s_c1)) != 0)) kw_args--;
        else {
          __Pyx_RaiseArgtupleInvalid("alpha_ad_dyn_v2", 0, 19, 22, 12); __PYX_ERR(1, 24, __pyx_L3_error)
        }
        CYTHON_FALLTHROUGH;
        case 13:
        if (likely((values[13] = __Pyx_PyDict_GetItemStr(__pyx_kwds, __pyx_n_s_c2)) != 0)) kw_args--;
        else {
          __Pyx_RaiseArgtupleInvalid("alpha_ad_dyn_v2", 0, 19, 22, 13); __PYX_ERR(1, 24, __pyx_L3_error)
        }
        CYTHON_FALLTHROUGH;
        case 14:
        if (likely((values[14] = __Pyx_PyDict_GetItemStr(__pyx_kwds, __pyx_n_s_beta_scalar)) != 0)) kw_args--;
        else {
          __Pyx_RaiseArgtupleInvalid("alpha_ad_dyn_v2", 0, 19, 22, 14); __PYX_ERR(1, 24, __pyx_L3_error)
        }
        CYTHON_FALLTHROUGH;
        case 15:
        if (likely((values[15] = __Pyx_PyDict_GetItemStr(__pyx_kwds, __pyx_n_s_hyper)) != 0)) kw_args--;
        else {
          __Pyx_RaiseArgtupleInvalid("alpha_ad_dyn_v2", 0, 19, 22, 15); __PYX_ERR(1, 24, __pyx_L3_error)
        }
        CYTHON_FALLTHROUGH;
        case 16:
        if (likely((values[16] = __Pyx_PyDict_GetItemStr(__pyx_kwds, __pyx_n_s_seed)) != 0)) kw_args--;
        else {
          __Pyx_RaiseArgtupleInvalid("alpha_ad_dyn_v2", 0, 19, 22, 16); __PYX_ERR(1, 24, __pyx_L3_error)
        }
        CYTHON_FALLTHROUGH;
        case 17:
        if (likely((values[17] = __Pyx_PyDict_GetItemStr(__pyx_kwds, __pyx_n_s_alpha_init)) != 0)) kw_args--;
        else {
          __Pyx_RaiseArgtupleInvalid("alpha_ad_dyn_v2", 0, 19, 22, 17); __PYX_ERR(1, 24, __pyx_L3_error)
        }
        CYTHON_FALLTHROUGH;
        case 18:
        if (likely((values[18] = __Pyx_PyDict_GetItemStr(__pyx_kwds, __pyx_n_s_sigma_init)) != 0)) kw_args--;
        else {
          __Pyx_RaiseArgtupleInvalid("alpha_ad_dyn_v2", 0, 19, 22, 18); __PYX_ERR(1, 24, __pyx_L3_error)
        }
        CYTHON_FALLTHROUGH;
        case 19:
        if (kw_args > 0) {
          PyObject* value = __Pyx_PyDict_GetItemStr(__pyx_kwds, __pyx_n_s_S_density);
          if (value) { values[19] = value; kw_args--; }
        }
        CYTHON_FALLTHROUGH;
        case 20:
        if (kw_args > 0) {
          PyObject* value = __Pyx_PyDict_GetItemStr(__pyx_kwds, __pyx_n_s_I_density);
          if (value) { values[20] = value; kw_args--; }
        }
        CYTHON_FALLTHROUGH;
        case 21:
        if (kw_args > 0) {
          PyObject* value = __Pyx_PyDict_GetItemStr(__pyx_kwds, __pyx_n_s_H_density);
          if (value) { values[21] = value; kw_args--; }
        }
      }
      if (unlikely(kw_args > 0)) {
        if (unlikely(__Pyx_ParseOptionalKeywords(__pyx_kwds, __pyx_pyargnames, 0, values, pos_args, "alpha_ad_dyn_v2") < 0)) __PYX_ERR(1, 24, __pyx_L3_error)
      }
    } else {
      switch (PyTuple_GET_SIZE(__pyx_args)) {
        case 22: values[21] = PyTuple_GET_ITEM(__pyx_args, 21);
        CYTHON_FALLTHROUGH;
        case 21: values[20] = PyTuple_GET_ITEM(__pyx_args, 20);
        CYTHON_FALLTHROUGH;
        case 20: values[19] = PyTuple_GET_ITEM(__pyx_args, 19);
        CYTHON_FALLTHROUGH;
        case 19: values[18] = PyTuple_GET_ITEM(__pyx_args, 18);
        values[17] = PyTuple_GET_ITEM(__pyx_args, 17);
        values[16] = PyTuple_GET_ITEM(__pyx_args, 16);
        values[15] = PyTuple_GET_ITEM(__pyx_args, 15);
        values[14] = PyTuple_GET_ITEM(__pyx_args, 14);
        values[13] = PyTuple_GET_ITEM(__pyx_args, 13);
        values[12] = PyTuple_GET_ITEM(__pyx_args, 12);
        values[11] = PyTuple_GET_ITEM(__pyx_args, 11);
        values[10] = PyTuple_GET_ITEM(__pyx_args, 10);
        values[9] = PyTuple_GET_ITEM(__pyx_args, 9);
        values[8] = PyTuple_GET_ITEM(__pyx_args, 8);
        values[7] = PyTuple_GET_ITEM(__pyx_args, 7);
        values[6] = PyTuple_GET_ITEM(__pyx_args, 6);
        values[5] = PyTuple_GET_ITEM(__pyx_args, 5);
        values[4] = PyTuple_GET_ITEM(__pyx_args, 4);
        values[3] = PyTuple_GET_ITEM(__pyx_args, 3);
        values[2] = PyTuple_GET_ITEM(__pyx_args, 2);
        values[1] = PyTuple_GET_ITEM(__pyx_args, 1);
        values[0] = PyTuple_GET_ITEM(__pyx_args, 0);
        break;
        default: goto __pyx_L5_argtuple_error;
      }
    }
    __pyx_v_beta_max = __pyx_PyFloat_AsFloat(values[0]); if (unlikely((__pyx_v_beta_max == (float)-1) && PyErr_Occurred())) __PYX_ERR(1, 24, __pyx_L3_error)
    __pyx_v_beta_min = __pyx_PyFloat_AsFloat(values[1]); if (unlikely((__pyx_v_beta_min == (float)-1) && PyErr_Occurred())) __PYX_ERR(1, 24, __pyx_L3_error)
    __pyx_v_alpha_max = __pyx_PyFloat_AsFloat(values[2]); if (unlikely((__pyx_v_alpha_max == (float)-1) && PyErr_Occurred())) __PYX_ERR(1, 24, __pyx_L3_error)
    __pyx_v_alpha_min = __pyx_PyFloat_AsFloat(values[3]); if (unlikely((__pyx_v_alpha_min == (float)-1) && PyErr_Occurred())) __PYX_ERR(1, 24, __pyx_L3_error)
    __pyx_v_sigma_max = __pyx_PyFloat_AsFloat(values[4]); if (unlikely((__pyx_v_sigma_max == (float)-1) && PyErr_Occurred())) __PYX_ERR(1, 24, __pyx_L3_error)
    __pyx_v_b = __pyx_PyFloat_AsFloat(values[5]); if (unlikely((__pyx_v_b == (float)-1) && PyErr_Occurred())) __PYX_ERR(1, 24, __pyx_L3_error)
    __pyx_v_q = __pyx_PyFloat_AsFloat(values[6]); if (unlikely((__pyx_v_q == (float)-1) && PyErr_Occurred())) __PYX_ERR(1, 24, __pyx_L3_error)
    __pyx_v_d = __pyx_PyFloat_AsFloat(values[7]); if (unlikely((__pyx_v_d == (float)-1) && PyErr_Occurred())) __PYX_ERR(1, 24, __pyx_L3_error)
    __pyx_v_rho = __pyx_PyFloat_AsFloat(values[8]); if (unlikely((__pyx_v_rho == (float)-1) && PyErr_Occurred())) __PYX_ERR(1, 24, __pyx_L3_error)
    __pyx_v_eta = __pyx_PyFloat_AsFloat(values[9]); if (unlikely((__pyx_v_eta == (float)-1) && PyErr_Occurred())) __PYX_ERR(1, 24, __pyx_L3_error)
    __pyx_v_gamma = __pyx_PyFloat_AsFloat(values[10]); if (unlikely((__pyx_v_gamma == (float)-1) && PyErr_Occurred())) __PYX_ERR(1, 24, __pyx_L3_error)
    __pyx_v_lam = __pyx_PyFloat_AsFloat(values[11]); if (unlikely((__pyx_v_lam == (float)-1) && PyErr_Occurred())) __PYX_ERR(1, 24, __pyx_L3_error)
    __pyx_v_c1 = __pyx_PyFloat_AsFloat(values[12]); if (unlikely((__pyx_v_c1 == (float)-1) && PyErr_Occurred())) __PYX_ERR(1, 24, __pyx_L3_error)
    __pyx_v_c2 = __pyx_PyFloat_AsFloat(values[13]); if (unlikely((__pyx_v_c2 == (float)-1) && PyErr_Occurred())) __PYX_ERR(1, 24, __pyx_L3_error)
    __pyx_v_beta_scalar = __pyx_PyFloat_AsFloat(values[14]); if (unlikely((__pyx_v_beta_scalar == (float)-1) && PyErr_Occurred())) __PYX_ERR(1, 24, __pyx_L3_error)
    __pyx_v_hyper = __pyx_PyFloat_AsFloat(values[15]); if (unlikely((__pyx_v_hyper == (float)-1) && PyErr_Occurred())) __PYX_ERR(1, 24, __pyx_L3_error)
    __pyx_v_seed = __Pyx_PyInt_As_int(values[16]); if (unlikely((__pyx_v_seed == (int)-1) && PyErr_Occurred())) __PYX_ERR(1, 24, __pyx_L3_error)
    __pyx_v_alpha_init = __Pyx_PyInt_As_int(values[17]); if (unlikely((__pyx_v_alpha_init == (int)-1) && PyErr_Occurred())) __PYX_ERR(1, 24, __pyx_L3_error)
    __pyx_v_sigma_init = __Pyx_PyInt_As_int(values[18]); if (unlikely((__pyx_v_sigma_init == (int)-1) && PyErr_Occurred())) __PYX_ERR(1, 24, __pyx_L3_error)
    if (values[19]) {
      __pyx_v_S_density = __pyx_PyFloat_AsFloat(values[19]); if (unlikely((__pyx_v_S_density == (float)-1) && PyErr_Occurred())) __PYX_ERR(1, 24, __pyx_L3_error)
    } else {
      __pyx_v_S_density = ((float)4.0);
    }
    if (values[20]) {
      __pyx_v_I_density = __pyx_PyFloat_AsFloat(values[20]); if (unlikely((__pyx_v_I_density == (float)-1) && PyErr_Occurred())) __PYX_ERR(1, 24, __pyx_L3_error)
    } else {
      __pyx_v_I_density = ((float)4.0);
    }
    if (values[21]) {
      __pyx_v_H_density = __pyx_PyFloat_AsFloat(values[21]); if (unlikely((__pyx_v_H_density == (float)-1) && PyErr_Occurred())) __PYX_ERR(1, 24, __pyx_L3_error)
    } else {
      __pyx_v_H_density = ((float)4.0);
    }
  }
  goto __pyx_L4_argument_unpacking_done;
  __pyx_L5_argtuple_error:;
  __Pyx_RaiseArgtupleInvalid("alpha_ad_dyn_v2", 0, 19, 22, PyTuple_GET_SIZE(__pyx_args)); __PYX_ERR(1, 24, __pyx_L3_error)
  __pyx_L3_error:;
  __Pyx_AddTraceback("runner.PySolver.alpha_ad_dyn_v2", __pyx_clineno, __pyx_lineno, __pyx_filename);
  __Pyx_RefNannyFinishContext();
  return NULL;
  __pyx_L4_argument_unpacking_done:;
  __pyx_r = __pyx_pf_6runner_8PySolver_6alpha_ad_dyn_v2(((struct __pyx_obj_6runner_PySolver *)__pyx_v_self), __pyx_v_beta_max, __pyx_v_beta_min, __pyx_v_alpha_max, __pyx_v_alpha_min, __pyx_v_sigma_max, __pyx_v_b, __pyx_v_q, __pyx_v_d, __pyx_v_rho, __pyx_v_eta, __pyx_v_gamma, __pyx_v_lam, __pyx_v_c1, __pyx_v_c2, __pyx_v_beta_scalar, __pyx_v_hyper, __pyx_v_seed, __pyx_v_alpha_init, __pyx_v_sigma_init, __pyx_v_S_density, __pyx_v_I_density, __pyx_v_H_density);

  /* function exit code */
  __Pyx_RefNannyFinishContext();
  return __pyx_r;
}

static PyObject *__pyx_pf_6runner_8PySolver_6alpha_ad_dyn_v2(struct __pyx_obj_6runner_PySolver *__pyx_v_self, float __pyx_v_beta_max, float __pyx_v_beta_min, float __pyx_v_alpha_max, float __pyx_v_alpha_min, float __pyx_v_sigma_max, float __pyx_v_b, float __pyx_v_q, float __pyx_v_d, float __pyx_v_rho, float __pyx_v_eta, float __pyx_v_gamma, float __pyx_v_lam, float __pyx_v_c1, float __pyx_v_c2, float __pyx_v_beta_scalar, float __pyx_v_hyper, int __pyx_v_seed, int __pyx_v_alpha_init, int __pyx_v_sigma_init, float __pyx_v_S_density, float __pyx_v_I_density, float __pyx_v_H_density) {
  PyObject *__pyx_r = NULL;
  __Pyx_RefNannyDeclarations
  __Pyx_RefNannySetupContext("alpha_ad_dyn_v2", 0);

  /* "runner.pyx":27
 * 
 * 
 *         self.cpp_solver.alpha_evo_only_v2( beta_max, beta_min, alpha_max, alpha_min, sigma_max, b, q, d, rho, eta, gamma, lam, c1, c2, beta_scalar, hyper, seed, alpha_init, sigma_init, S_density, I_density, H_density)             # <<<<<<<<<<<<<<
 * 
 *         return
 */
  __pyx_v_self->cpp_solver.alpha_evo_only_v2(__pyx_v_beta_max, __pyx_v_beta_min, __pyx_v_alpha_max, __pyx_v_alpha_min, __pyx_v_sigma_max, __pyx_v_b, __pyx_v_q, __pyx_v_d, __pyx_v_rho, __pyx_v_eta, __pyx_v_gamma, __pyx_v_lam, __pyx_v_c1, __pyx_v_c2, __pyx_v_beta_scalar, __pyx_v_hyper, __pyx_v_seed, __pyx_v_alpha_init, __pyx_v_sigma_init, __pyx_v_S_density, __pyx_v_I_density, __pyx_v_H_density);

  /* "runner.pyx":29
 *         self.cpp_solver.alpha_evo_only_v2( beta_max, beta_min, alpha_max, alpha_min, sigma_max, b, q, d, rho, eta, gamma, lam, c1, c2, beta_scalar, hyper, seed, alpha_init, sigma_init, S_density, I_density, H_density)
 * 
 *         return             # <<<<<<<<<<<<<<
 * 
 *     def alpha_ad_dyn_v3(self, float beta_max, float alpha_max, float sigma_max, float b, float q, float d, float rho, float eta, float gamma, float lam, float c1, float c2, float hyper, int seed, int alpha_init, int sigma_init, float S_density = 4.0, float I_density = 4.0, float H_density = 4.0):
 */
  __Pyx_XDECREF(__pyx_r);
  __pyx_r = Py_None; __Pyx_INCREF(Py_None);
  goto __pyx_L0;

  /* "runner.pyx":24
 *         return
 * 
 *     def alpha_ad_dyn_v2(self, float beta_max, float beta_min, float alpha_max, float alpha_min, float sigma_max, float b, float q, float d, float rho, float eta, float gamma, float lam, float c1, float c2, float beta_scalar, float hyper, int seed, int alpha_init, int sigma_init, float S_density = 4.0, float I_density = 4.0, float H_density = 4.0):             # <<<<<<<<<<<<<<
 * 
 * 
 */

  /* function exit code */
  __pyx_L0:;
  __Pyx_XGIVEREF(__pyx_r);
  __Pyx_RefNannyFinishContext();
  return __pyx_r;
}

/* "runner.pyx":31
 *         return
 * 
 *     def alpha_ad_dyn_v3(self, float beta_max, float alpha_max, float sigma_max, float b, float q, float d, float rho, float eta, float gamma, float lam, float c1, float c2, float hyper, int seed, int alpha_init, int sigma_init, float S_density = 4.0, float I_density = 4.0, float H_density = 4.0):             # <<<<<<<<<<<<<<
 * 
 * 
 */

/* Python wrapper */
static PyObject *__pyx_pw_6runner_8PySolver_9alpha_ad_dyn_v3(PyObject *__pyx_v_self, PyObject *__pyx_args, PyObject *__pyx_kwds); /*proto*/
static PyObject *__pyx_pw_6runner_8PySolver_9alpha_ad_dyn_v3(PyObject *__pyx_v_self, PyObject *__pyx_args, PyObject *__pyx_kwds) {
  float __pyx_v_beta_max;
  float __pyx_v_alpha_max;
  float __pyx_v_sigma_max;
  float __pyx_v_b;
  float __pyx_v_q;
  float __pyx_v_d;
  float __pyx_v_rho;
  float __pyx_v_eta;
  float __pyx_v_gamma;
  float __pyx_v_lam;
  float __pyx_v_c1;
  float __pyx_v_c2;
  float __pyx_v_hyper;
  int __pyx_v_seed;
  int __pyx_v_alpha_init;
  int __pyx_v_sigma_init;
  float __pyx_v_S_density;
  float __pyx_v_I_density;
  float __pyx_v_H_density;
  int __pyx_lineno = 0;
  const char *__pyx_filename = NULL;
  int __pyx_clineno = 0;
  PyObject *__pyx_r = 0;
  __Pyx_RefNannyDeclarations
  __Pyx_RefNannySetupContext("alpha_ad_dyn_v3 (wrapper)", 0);
  {
    static PyObject **__pyx_pyargnames[] = {&__pyx_n_s_beta_max,&__pyx_n_s_alpha_max,&__pyx_n_s_sigma_max,&__pyx_n_s_b,&__pyx_n_s_q,&__pyx_n_s_d,&__pyx_n_s_rho,&__pyx_n_s_eta,&__pyx_n_s_gamma,&__pyx_n_s_lam,&__pyx_n_s_c1,&__pyx_n_s_c2,&__pyx_n_s_hyper,&__pyx_n_s_seed,&__pyx_n_s_alpha_init,&__pyx_n_s_sigma_init,&__pyx_n_s_S_density,&__pyx_n_s_I_density,&__pyx_n_s_H_density,0};
    PyObject* values[19] = {0,0,0,0,0,0,0,0,0,0,0,0,0,0,0,0,0,0,0};
    if (unlikely(__pyx_kwds)) {
      Py_ssize_t kw_args;
      const Py_ssize_t pos_args = PyTuple_GET_SIZE(__pyx_args);
      switch (pos_args) {
        case 19: values[18] = PyTuple_GET_ITEM(__pyx_args, 18);
        CYTHON_FALLTHROUGH;
        case 18: values[17] = PyTuple_GET_ITEM(__pyx_args, 17);
        CYTHON_FALLTHROUGH;
        case 17: values[16] = PyTuple_GET_ITEM(__pyx_args, 16);
        CYTHON_FALLTHROUGH;
        case 16: values[15] = PyTuple_GET_ITEM(__pyx_args, 15);
        CYTHON_FALLTHROUGH;
        case 15: values[14] = PyTuple_GET_ITEM(__pyx_args, 14);
        CYTHON_FALLTHROUGH;
        case 14: values[13] = PyTuple_GET_ITEM(__pyx_args, 13);
        CYTHON_FALLTHROUGH;
        case 13: values[12] = PyTuple_GET_ITEM(__pyx_args, 12);
        CYTHON_FALLTHROUGH;
        case 12: values[11] = PyTuple_GET_ITEM(__pyx_args, 11);
        CYTHON_FALLTHROUGH;
        case 11: values[10] = PyTuple_GET_ITEM(__pyx_args, 10);
        CYTHON_FALLTHROUGH;
        case 10: values[9] = PyTuple_GET_ITEM(__pyx_args, 9);
        CYTHON_FALLTHROUGH;
        case  9: values[8] = PyTuple_GET_ITEM(__pyx_args, 8);
        CYTHON_FALLTHROUGH;
        case  8: values[7] = PyTuple_GET_ITEM(__pyx_args, 7);
        CYTHON_FALLTHROUGH;
        case  7: values[6] = PyTuple_GET_ITEM(__pyx_args, 6);
        CYTHON_FALLTHROUGH;
        case  6: values[5] = PyTuple_GET_ITEM(__pyx_args, 5);
        CYTHON_FALLTHROUGH;
        case  5: values[4] = PyTuple_GET_ITEM(__pyx_args, 4);
        CYTHON_FALLTHROUGH;
        case  4: values[3] = PyTuple_GET_ITEM(__pyx_args, 3);
        CYTHON_FALLTHROUGH;
        case  3: values[2] = PyTuple_GET_ITEM(__pyx_args, 2);
        CYTHON_FALLTHROUGH;
        case  2: values[1] = PyTuple_GET_ITEM(__pyx_args, 1);
        CYTHON_FALLTHROUGH;
        case  1: values[0] = PyTuple_GET_ITEM(__pyx_args, 0);
        CYTHON_FALLTHROUGH;
        case  0: break;
        default: goto __pyx_L5_argtuple_error;
      }
      kw_args = PyDict_Size(__pyx_kwds);
      switch (pos_args) {
        case  0:
        if (likely((values[0] = __Pyx_PyDict_GetItemStr(__pyx_kwds, __pyx_n_s_beta_max)) != 0)) kw_args--;
        else goto __pyx_L5_argtuple_error;
        CYTHON_FALLTHROUGH;
        case  1:
        if (likely((values[1] = __Pyx_PyDict_GetItemStr(__pyx_kwds, __pyx_n_s_alpha_max)) != 0)) kw_args--;
        else {
          __Pyx_RaiseArgtupleInvalid("alpha_ad_dyn_v3", 0, 16, 19, 1); __PYX_ERR(1, 31, __pyx_L3_error)
        }
        CYTHON_FALLTHROUGH;
        case  2:
        if (likely((values[2] = __Pyx_PyDict_GetItemStr(__pyx_kwds, __pyx_n_s_sigma_max)) != 0)) kw_args--;
        else {
          __Pyx_RaiseArgtupleInvalid("alpha_ad_dyn_v3", 0, 16, 19, 2); __PYX_ERR(1, 31, __pyx_L3_error)
        }
        CYTHON_FALLTHROUGH;
        case  3:
        if (likely((values[3] = __Pyx_PyDict_GetItemStr(__pyx_kwds, __pyx_n_s_b)) != 0)) kw_args--;
        else {
          __Pyx_RaiseArgtupleInvalid("alpha_ad_dyn_v3", 0, 16, 19, 3); __PYX_ERR(1, 31, __pyx_L3_error)
        }
        CYTHON_FALLTHROUGH;
        case  4:
        if (likely((values[4] = __Pyx_PyDict_GetItemStr(__pyx_kwds, __pyx_n_s_q)) != 0)) kw_args--;
        else {
          __Pyx_RaiseArgtupleInvalid("alpha_ad_dyn_v3", 0, 16, 19, 4); __PYX_ERR(1, 31, __pyx_L3_error)
        }
        CYTHON_FALLTHROUGH;
        case  5:
        if (likely((values[5] = __Pyx_PyDict_GetItemStr(__pyx_kwds, __pyx_n_s_d)) != 0)) kw_args--;
        else {
          __Pyx_RaiseArgtupleInvalid("alpha_ad_dyn_v3", 0, 16, 19, 5); __PYX_ERR(1, 31, __pyx_L3_error)
        }
        CYTHON_FALLTHROUGH;
        case  6:
        if (likely((values[6] = __Pyx_PyDict_GetItemStr(__pyx_kwds, __pyx_n_s_rho)) != 0)) kw_args--;
        else {
          __Pyx_RaiseArgtupleInvalid("alpha_ad_dyn_v3", 0, 16, 19, 6); __PYX_ERR(1, 31, __pyx_L3_error)
        }
        CYTHON_FALLTHROUGH;
        case  7:
        if (likely((values[7] = __Pyx_PyDict_GetItemStr(__pyx_kwds, __pyx_n_s_eta)) != 0)) kw_args--;
        else {
          __Pyx_RaiseArgtupleInvalid("alpha_ad_dyn_v3", 0, 16, 19, 7); __PYX_ERR(1, 31, __pyx_L3_error)
        }
        CYTHON_FALLTHROUGH;
        case  8:
        if (likely((values[8] = __Pyx_PyDict_GetItemStr(__pyx_kwds, __pyx_n_s_gamma)) != 0)) kw_args--;
        else {
          __Pyx_RaiseArgtupleInvalid("alpha_ad_dyn_v3", 0, 16, 19, 8); __PYX_ERR(1, 31, __pyx_L3_error)
        }
        CYTHON_FALLTHROUGH;
        case  9:
        if (likely((values[9] = __Pyx_PyDict_GetItemStr(__pyx_kwds, __pyx_n_s_lam)) != 0)) kw_args--;
        else {
          __Pyx_RaiseArgtupleInvalid("alpha_ad_dyn_v3", 0, 16, 19, 9); __PYX_ERR(1, 31, __pyx_L3_error)
        }
        CYTHON_FALLTHROUGH;
        case 10:
        if (likely((values[10] = __Pyx_PyDict_GetItemStr(__pyx_kwds, __pyx_n_s_c1)) != 0)) kw_args--;
        else {
          __Pyx_RaiseArgtupleInvalid("alpha_ad_dyn_v3", 0, 16, 19, 10); __PYX_ERR(1, 31, __pyx_L3_error)
        }
        CYTHON_FALLTHROUGH;
        case 11:
        if (likely((values[11] = __Pyx_PyDict_GetItemStr(__pyx_kwds, __pyx_n_s_c2)) != 0)) kw_args--;
        else {
          __Pyx_RaiseArgtupleInvalid("alpha_ad_dyn_v3", 0, 16, 19, 11); __PYX_ERR(1, 31, __pyx_L3_error)
        }
        CYTHON_FALLTHROUGH;
        case 12:
        if (likely((values[12] = __Pyx_PyDict_GetItemStr(__pyx_kwds, __pyx_n_s_hyper)) != 0)) kw_args--;
        else {
          __Pyx_RaiseArgtupleInvalid("alpha_ad_dyn_v3", 0, 16, 19, 12); __PYX_ERR(1, 31, __pyx_L3_error)
        }
        CYTHON_FALLTHROUGH;
        case 13:
        if (likely((values[13] = __Pyx_PyDict_GetItemStr(__pyx_kwds, __pyx_n_s_seed)) != 0)) kw_args--;
        else {
          __Pyx_RaiseArgtupleInvalid("alpha_ad_dyn_v3", 0, 16, 19, 13); __PYX_ERR(1, 31, __pyx_L3_error)
        }
        CYTHON_FALLTHROUGH;
        case 14:
        if (likely((values[14] = __Pyx_PyDict_GetItemStr(__pyx_kwds, __pyx_n_s_alpha_init)) != 0)) kw_args--;
        else {
          __Pyx_RaiseArgtupleInvalid("alpha_ad_dyn_v3", 0, 16, 19, 14); __PYX_ERR(1, 31, __pyx_L3_error)
        }
        CYTHON_FALLTHROUGH;
        case 15:
        if (likely((values[15] = __Pyx_PyDict_GetItemStr(__pyx_kwds, __pyx_n_s_sigma_init)) != 0)) kw_args--;
        else {
          __Pyx_RaiseArgtupleInvalid("alpha_ad_dyn_v3", 0, 16, 19, 15); __PYX_ERR(1, 31, __pyx_L3_error)
        }
        CYTHON_FALLTHROUGH;
        case 16:
        if (kw_args > 0) {
          PyObject* value = __Pyx_PyDict_GetItemStr(__pyx_kwds, __pyx_n_s_S_density);
          if (value) { values[16] = value; kw_args--; }
        }
        CYTHON_FALLTHROUGH;
        case 17:
        if (kw_args > 0) {
          PyObject* value = __Pyx_PyDict_GetItemStr(__pyx_kwds, __pyx_n_s_I_density);
          if (value) { values[17] = value; kw_args--; }
        }
        CYTHON_FALLTHROUGH;
        case 18:
        if (kw_args > 0) {
          PyObject* value = __Pyx_PyDict_GetItemStr(__pyx_kwds, __pyx_n_s_H_density);
          if (value) { values[18] = value; kw_args--; }
        }
      }
      if (unlikely(kw_args > 0)) {
        if (unlikely(__Pyx_ParseOptionalKeywords(__pyx_kwds, __pyx_pyargnames, 0, values, pos_args, "alpha_ad_dyn_v3") < 0)) __PYX_ERR(1, 31, __pyx_L3_error)
      }
    } else {
      switch (PyTuple_GET_SIZE(__pyx_args)) {
        case 19: values[18] = PyTuple_GET_ITEM(__pyx_args, 18);
        CYTHON_FALLTHROUGH;
        case 18: values[17] = PyTuple_GET_ITEM(__pyx_args, 17);
        CYTHON_FALLTHROUGH;
        case 17: values[16] = PyTuple_GET_ITEM(__pyx_args, 16);
        CYTHON_FALLTHROUGH;
        case 16: values[15] = PyTuple_GET_ITEM(__pyx_args, 15);
        values[14] = PyTuple_GET_ITEM(__pyx_args, 14);
        values[13] = PyTuple_GET_ITEM(__pyx_args, 13);
        values[12] = PyTuple_GET_ITEM(__pyx_args, 12);
        values[11] = PyTuple_GET_ITEM(__pyx_args, 11);
        values[10] = PyTuple_GET_ITEM(__pyx_args, 10);
        values[9] = PyTuple_GET_ITEM(__pyx_args, 9);
        values[8] = PyTuple_GET_ITEM(__pyx_args, 8);
        values[7] = PyTuple_GET_ITEM(__pyx_args, 7);
        values[6] = PyTuple_GET_ITEM(__pyx_args, 6);
        values[5] = PyTuple_GET_ITEM(__pyx_args, 5);
        values[4] = PyTuple_GET_ITEM(__pyx_args, 4);
        values[3] = PyTuple_GET_ITEM(__pyx_args, 3);
        values[2] = PyTuple_GET_ITEM(__pyx_args, 2);
        values[1] = PyTuple_GET_ITEM(__pyx_args, 1);
        values[0] = PyTuple_GET_ITEM(__pyx_args, 0);
        break;
        default: goto __pyx_L5_argtuple_error;
      }
    }
    __pyx_v_beta_max = __pyx_PyFloat_AsFloat(values[0]); if (unlikely((__pyx_v_beta_max == (float)-1) && PyErr_Occurred())) __PYX_ERR(1, 31, __pyx_L3_error)
    __pyx_v_alpha_max = __pyx_PyFloat_AsFloat(values[1]); if (unlikely((__pyx_v_alpha_max == (float)-1) && PyErr_Occurred())) __PYX_ERR(1, 31, __pyx_L3_error)
    __pyx_v_sigma_max = __pyx_PyFloat_AsFloat(values[2]); if (unlikely((__pyx_v_sigma_max == (float)-1) && PyErr_Occurred())) __PYX_ERR(1, 31, __pyx_L3_error)
    __pyx_v_b = __pyx_PyFloat_AsFloat(values[3]); if (unlikely((__pyx_v_b == (float)-1) && PyErr_Occurred())) __PYX_ERR(1, 31, __pyx_L3_error)
    __pyx_v_q = __pyx_PyFloat_AsFloat(values[4]); if (unlikely((__pyx_v_q == (float)-1) && PyErr_Occurred())) __PYX_ERR(1, 31, __pyx_L3_error)
    __pyx_v_d = __pyx_PyFloat_AsFloat(values[5]); if (unlikely((__pyx_v_d == (float)-1) && PyErr_Occurred())) __PYX_ERR(1, 31, __pyx_L3_error)
    __pyx_v_rho = __pyx_PyFloat_AsFloat(values[6]); if (unlikely((__pyx_v_rho == (float)-1) && PyErr_Occurred())) __PYX_ERR(1, 31, __pyx_L3_error)
    __pyx_v_eta = __pyx_PyFloat_AsFloat(values[7]); if (unlikely((__pyx_v_eta == (float)-1) && PyErr_Occurred())) __PYX_ERR(1, 31, __pyx_L3_error)
    __pyx_v_gamma = __pyx_PyFloat_AsFloat(values[8]); if (unlikely((__pyx_v_gamma == (float)-1) && PyErr_Occurred())) __PYX_ERR(1, 31, __pyx_L3_error)
    __pyx_v_lam = __pyx_PyFloat_AsFloat(values[9]); if (unlikely((__pyx_v_lam == (float)-1) && PyErr_Occurred())) __PYX_ERR(1, 31, __pyx_L3_error)
    __pyx_v_c1 = __pyx_PyFloat_AsFloat(values[10]); if (unlikely((__pyx_v_c1 == (float)-1) && PyErr_Occurred())) __PYX_ERR(1, 31, __pyx_L3_error)
    __pyx_v_c2 = __pyx_PyFloat_AsFloat(values[11]); if (unlikely((__pyx_v_c2 == (float)-1) && PyErr_Occurred())) __PYX_ERR(1, 31, __pyx_L3_error)
    __pyx_v_hyper = __pyx_PyFloat_AsFloat(values[12]); if (unlikely((__pyx_v_hyper == (float)-1) && PyErr_Occurred())) __PYX_ERR(1, 31, __pyx_L3_error)
    __pyx_v_seed = __Pyx_PyInt_As_int(values[13]); if (unlikely((__pyx_v_seed == (int)-1) && PyErr_Occurred())) __PYX_ERR(1, 31, __pyx_L3_error)
    __pyx_v_alpha_init = __Pyx_PyInt_As_int(values[14]); if (unlikely((__pyx_v_alpha_init == (int)-1) && PyErr_Occurred())) __PYX_ERR(1, 31, __pyx_L3_error)
    __pyx_v_sigma_init = __Pyx_PyInt_As_int(values[15]); if (unlikely((__pyx_v_sigma_init == (int)-1) && PyErr_Occurred())) __PYX_ERR(1, 31, __pyx_L3_error)
    if (values[16]) {
      __pyx_v_S_density = __pyx_PyFloat_AsFloat(values[16]); if (unlikely((__pyx_v_S_density == (float)-1) && PyErr_Occurred())) __PYX_ERR(1, 31, __pyx_L3_error)
    } else {
      __pyx_v_S_density = ((float)4.0);
    }
    if (values[17]) {
      __pyx_v_I_density = __pyx_PyFloat_AsFloat(values[17]); if (unlikely((__pyx_v_I_density == (float)-1) && PyErr_Occurred())) __PYX_ERR(1, 31, __pyx_L3_error)
    } else {
      __pyx_v_I_density = ((float)4.0);
    }
    if (values[18]) {
      __pyx_v_H_density = __pyx_PyFloat_AsFloat(values[18]); if (unlikely((__pyx_v_H_density == (float)-1) && PyErr_Occurred())) __PYX_ERR(1, 31, __pyx_L3_error)
    } else {
      __pyx_v_H_density = ((float)4.0);
    }
  }
  goto __pyx_L4_argument_unpacking_done;
  __pyx_L5_argtuple_error:;
  __Pyx_RaiseArgtupleInvalid("alpha_ad_dyn_v3", 0, 16, 19, PyTuple_GET_SIZE(__pyx_args)); __PYX_ERR(1, 31, __pyx_L3_error)
  __pyx_L3_error:;
  __Pyx_AddTraceback("runner.PySolver.alpha_ad_dyn_v3", __pyx_clineno, __pyx_lineno, __pyx_filename);
  __Pyx_RefNannyFinishContext();
  return NULL;
  __pyx_L4_argument_unpacking_done:;
  __pyx_r = __pyx_pf_6runner_8PySolver_8alpha_ad_dyn_v3(((struct __pyx_obj_6runner_PySolver *)__pyx_v_self), __pyx_v_beta_max, __pyx_v_alpha_max, __pyx_v_sigma_max, __pyx_v_b, __pyx_v_q, __pyx_v_d, __pyx_v_rho, __pyx_v_eta, __pyx_v_gamma, __pyx_v_lam, __pyx_v_c1, __pyx_v_c2, __pyx_v_hyper, __pyx_v_seed, __pyx_v_alpha_init, __pyx_v_sigma_init, __pyx_v_S_density, __pyx_v_I_density, __pyx_v_H_density);

  /* function exit code */
  __Pyx_RefNannyFinishContext();
  return __pyx_r;
}

static PyObject *__pyx_pf_6runner_8PySolver_8alpha_ad_dyn_v3(struct __pyx_obj_6runner_PySolver *__pyx_v_self, float __pyx_v_beta_max, float __pyx_v_alpha_max, float __pyx_v_sigma_max, float __pyx_v_b, float __pyx_v_q, float __pyx_v_d, float __pyx_v_rho, float __pyx_v_eta, float __pyx_v_gamma, float __pyx_v_lam, float __pyx_v_c1, float __pyx_v_c2, float __pyx_v_hyper, int __pyx_v_seed, int __pyx_v_alpha_init, int __pyx_v_sigma_init, float __pyx_v_S_density, float __pyx_v_I_density, float __pyx_v_H_density) {
  PyObject *__pyx_r = NULL;
  __Pyx_RefNannyDeclarations
  __Pyx_RefNannySetupContext("alpha_ad_dyn_v3", 0);

  /* "runner.pyx":34
 * 
 * 
 *         self.cpp_solver.alpha_evo_only_v3( beta_max, alpha_max, sigma_max, b, q, d, rho, eta, gamma, lam, c1, c2, hyper, seed, alpha_init, sigma_init, S_density, I_density, H_density)             # <<<<<<<<<<<<<<
 * 
 *         return
 */
  __pyx_v_self->cpp_solver.alpha_evo_only_v3(__pyx_v_beta_max, __pyx_v_alpha_max, __pyx_v_sigma_max, __pyx_v_b, __pyx_v_q, __pyx_v_d, __pyx_v_rho, __pyx_v_eta, __pyx_v_gamma, __pyx_v_lam, __pyx_v_c1, __pyx_v_c2, __pyx_v_hyper, __pyx_v_seed, __pyx_v_alpha_init, __pyx_v_sigma_init, __pyx_v_S_density, __pyx_v_I_density, __pyx_v_H_density);

  /* "runner.pyx":36
 *         self.cpp_solver.alpha_evo_only_v3( beta_max, alpha_max, sigma_max, b, q, d, rho, eta, gamma, lam, c1, c2, hyper, seed, alpha_init, sigma_init, S_density, I_density, H_density)
 * 
 *         return             # <<<<<<<<<<<<<<
 * 
 *     def alpha_ad_dyn_v4(self, float beta_max, float beta_lin, float alpha_max, float alpha_min, float sigma_max, float b, float q, float d, float rho, float eta, float gamma, float lam, float c1, float c2, float beta_scalar, float hyper, int seed, int alpha_init, int sigma_init, int evo_steps = 1000, float S_density = 4.0, float I_density = 4.0, float H_density = 4.0):
 */
  __Pyx_XDECREF(__pyx_r);
  __pyx_r = Py_None; __Pyx_INCREF(Py_None);
  goto __pyx_L0;

  /* "runner.pyx":31
 *         return
 * 
 *     def alpha_ad_dyn_v3(self, float beta_max, float alpha_max, float sigma_max, float b, float q, float d, float rho, float eta, float gamma, float lam, float c1, float c2, float hyper, int seed, int alpha_init, int sigma_init, float S_density = 4.0, float I_density = 4.0, float H_density = 4.0):             # <<<<<<<<<<<<<<
 * 
 * 
 */

  /* function exit code */
  __pyx_L0:;
  __Pyx_XGIVEREF(__pyx_r);
  __Pyx_RefNannyFinishContext();
  return __pyx_r;
}

/* "runner.pyx":38
 *         return
 * 
 *     def alpha_ad_dyn_v4(self, float beta_max, float beta_lin, float alpha_max, float alpha_min, float sigma_max, float b, float q, float d, float rho, float eta, float gamma, float lam, float c1, float c2, float beta_scalar, float hyper, int seed, int alpha_init, int sigma_init, int evo_steps = 1000, float S_density = 4.0, float I_density = 4.0, float H_density = 4.0):             # <<<<<<<<<<<<<<
 * 
 * 
 */

/* Python wrapper */
static PyObject *__pyx_pw_6runner_8PySolver_11alpha_ad_dyn_v4(PyObject *__pyx_v_self, PyObject *__pyx_args, PyObject *__pyx_kwds); /*proto*/
static PyObject *__pyx_pw_6runner_8PySolver_11alpha_ad_dyn_v4(PyObject *__pyx_v_self, PyObject *__pyx_args, PyObject *__pyx_kwds) {
  float __pyx_v_beta_max;
  float __pyx_v_beta_lin;
  float __pyx_v_alpha_max;
  float __pyx_v_alpha_min;
  float __pyx_v_sigma_max;
  float __pyx_v_b;
  float __pyx_v_q;
  float __pyx_v_d;
  float __pyx_v_rho;
  float __pyx_v_eta;
  float __pyx_v_gamma;
  float __pyx_v_lam;
  float __pyx_v_c1;
  float __pyx_v_c2;
  float __pyx_v_beta_scalar;
  float __pyx_v_hyper;
  int __pyx_v_seed;
  int __pyx_v_alpha_init;
  int __pyx_v_sigma_init;
  int __pyx_v_evo_steps;
  float __pyx_v_S_density;
  float __pyx_v_I_density;
  float __pyx_v_H_density;
  int __pyx_lineno = 0;
  const char *__pyx_filename = NULL;
  int __pyx_clineno = 0;
  PyObject *__pyx_r = 0;
  __Pyx_RefNannyDeclarations
  __Pyx_RefNannySetupContext("alpha_ad_dyn_v4 (wrapper)", 0);
  {
    static PyObject **__pyx_pyargnames[] = {&__pyx_n_s_beta_max,&__pyx_n_s_beta_lin,&__pyx_n_s_alpha_max,&__pyx_n_s_alpha_min,&__pyx_n_s_sigma_max,&__pyx_n_s_b,&__pyx_n_s_q,&__pyx_n_s_d,&__pyx_n_s_rho,&__pyx_n_s_eta,&__pyx_n_s_gamma,&__pyx_n_s_lam,&__pyx_n_s_c1,&__pyx_n_s_c2,&__pyx_n_s_beta_scalar,&__pyx_n_s_hyper,&__pyx_n_s_seed,&__pyx_n_s_alpha_init,&__pyx_n_s_sigma_init,&__pyx_n_s_evo_steps,&__pyx_n_s_S_density,&__pyx_n_s_I_density,&__pyx_n_s_H_density,0};
    PyObject* values[23] = {0,0,0,0,0,0,0,0,0,0,0,0,0,0,0,0,0,0,0,0,0,0,0};
    if (unlikely(__pyx_kwds)) {
      Py_ssize_t kw_args;
      const Py_ssize_t pos_args = PyTuple_GET_SIZE(__pyx_args);
      switch (pos_args) {
        case 23: values[22] = PyTuple_GET_ITEM(__pyx_args, 22);
        CYTHON_FALLTHROUGH;
        case 22: values[21] = PyTuple_GET_ITEM(__pyx_args, 21);
        CYTHON_FALLTHROUGH;
        case 21: values[20] = PyTuple_GET_ITEM(__pyx_args, 20);
        CYTHON_FALLTHROUGH;
        case 20: values[19] = PyTuple_GET_ITEM(__pyx_args, 19);
        CYTHON_FALLTHROUGH;
        case 19: values[18] = PyTuple_GET_ITEM(__pyx_args, 18);
        CYTHON_FALLTHROUGH;
        case 18: values[17] = PyTuple_GET_ITEM(__pyx_args, 17);
        CYTHON_FALLTHROUGH;
        case 17: values[16] = PyTuple_GET_ITEM(__pyx_args, 16);
        CYTHON_FALLTHROUGH;
        case 16: values[15] = PyTuple_GET_ITEM(__pyx_args, 15);
        CYTHON_FALLTHROUGH;
        case 15: values[14] = PyTuple_GET_ITEM(__pyx_args, 14);
        CYTHON_FALLTHROUGH;
        case 14: values[13] = PyTuple_GET_ITEM(__pyx_args, 13);
        CYTHON_FALLTHROUGH;
        case 13: values[12] = PyTuple_GET_ITEM(__pyx_args, 12);
        CYTHON_FALLTHROUGH;
        case 12: values[11] = PyTuple_GET_ITEM(__pyx_args, 11);
        CYTHON_FALLTHROUGH;
        case 11: values[10] = PyTuple_GET_ITEM(__pyx_args, 10);
        CYTHON_FALLTHROUGH;
        case 10: values[9] = PyTuple_GET_ITEM(__pyx_args, 9);
        CYTHON_FALLTHROUGH;
        case  9: values[8] = PyTuple_GET_ITEM(__pyx_args, 8);
        CYTHON_FALLTHROUGH;
        case  8: values[7] = PyTuple_GET_ITEM(__pyx_args, 7);
        CYTHON_FALLTHROUGH;
        case  7: values[6] = PyTuple_GET_ITEM(__pyx_args, 6);
        CYTHON_FALLTHROUGH;
        case  6: values[5] = PyTuple_GET_ITEM(__pyx_args, 5);
        CYTHON_FALLTHROUGH;
        case  5: values[4] = PyTuple_GET_ITEM(__pyx_args, 4);
        CYTHON_FALLTHROUGH;
        case  4: values[3] = PyTuple_GET_ITEM(__pyx_args, 3);
        CYTHON_FALLTHROUGH;
        case  3: values[2] = PyTuple_GET_ITEM(__pyx_args, 2);
        CYTHON_FALLTHROUGH;
        case  2: values[1] = PyTuple_GET_ITEM(__pyx_args, 1);
        CYTHON_FALLTHROUGH;
        case  1: values[0] = PyTuple_GET_ITEM(__pyx_args, 0);
        CYTHON_FALLTHROUGH;
        case  0: break;
        default: goto __pyx_L5_argtuple_error;
      }
      kw_args = PyDict_Size(__pyx_kwds);
      switch (pos_args) {
        case  0:
        if (likely((values[0] = __Pyx_PyDict_GetItemStr(__pyx_kwds, __pyx_n_s_beta_max)) != 0)) kw_args--;
        else goto __pyx_L5_argtuple_error;
        CYTHON_FALLTHROUGH;
        case  1:
        if (likely((values[1] = __Pyx_PyDict_GetItemStr(__pyx_kwds, __pyx_n_s_beta_lin)) != 0)) kw_args--;
        else {
          __Pyx_RaiseArgtupleInvalid("alpha_ad_dyn_v4", 0, 19, 23, 1); __PYX_ERR(1, 38, __pyx_L3_error)
        }
        CYTHON_FALLTHROUGH;
        case  2:
        if (likely((values[2] = __Pyx_PyDict_GetItemStr(__pyx_kwds, __pyx_n_s_alpha_max)) != 0)) kw_args--;
        else {
          __Pyx_RaiseArgtupleInvalid("alpha_ad_dyn_v4", 0, 19, 23, 2); __PYX_ERR(1, 38, __pyx_L3_error)
        }
        CYTHON_FALLTHROUGH;
        case  3:
        if (likely((values[3] = __Pyx_PyDict_GetItemStr(__pyx_kwds, __pyx_n_s_alpha_min)) != 0)) kw_args--;
        else {
          __Pyx_RaiseArgtupleInvalid("alpha_ad_dyn_v4", 0, 19, 23, 3); __PYX_ERR(1, 38, __pyx_L3_error)
        }
        CYTHON_FALLTHROUGH;
        case  4:
        if (likely((values[4] = __Pyx_PyDict_GetItemStr(__pyx_kwds, __pyx_n_s_sigma_max)) != 0)) kw_args--;
        else {
          __Pyx_RaiseArgtupleInvalid("alpha_ad_dyn_v4", 0, 19, 23, 4); __PYX_ERR(1, 38, __pyx_L3_error)
        }
        CYTHON_FALLTHROUGH;
        case  5:
        if (likely((values[5] = __Pyx_PyDict_GetItemStr(__pyx_kwds, __pyx_n_s_b)) != 0)) kw_args--;
        else {
          __Pyx_RaiseArgtupleInvalid("alpha_ad_dyn_v4", 0, 19, 23, 5); __PYX_ERR(1, 38, __pyx_L3_error)
        }
        CYTHON_FALLTHROUGH;
        case  6:
        if (likely((values[6] = __Pyx_PyDict_GetItemStr(__pyx_kwds, __pyx_n_s_q)) != 0)) kw_args--;
        else {
          __Pyx_RaiseArgtupleInvalid("alpha_ad_dyn_v4", 0, 19, 23, 6); __PYX_ERR(1, 38, __pyx_L3_error)
        }
        CYTHON_FALLTHROUGH;
        case  7:
        if (likely((values[7] = __Pyx_PyDict_GetItemStr(__pyx_kwds, __pyx_n_s_d)) != 0)) kw_args--;
        else {
          __Pyx_RaiseArgtupleInvalid("alpha_ad_dyn_v4", 0, 19, 23, 7); __PYX_ERR(1, 38, __pyx_L3_error)
        }
        CYTHON_FALLTHROUGH;
        case  8:
        if (likely((values[8] = __Pyx_PyDict_GetItemStr(__pyx_kwds, __pyx_n_s_rho)) != 0)) kw_args--;
        else {
          __Pyx_RaiseArgtupleInvalid("alpha_ad_dyn_v4", 0, 19, 23, 8); __PYX_ERR(1, 38, __pyx_L3_error)
        }
        CYTHON_FALLTHROUGH;
        case  9:
        if (likely((values[9] = __Pyx_PyDict_GetItemStr(__pyx_kwds, __pyx_n_s_eta)) != 0)) kw_args--;
        else {
          __Pyx_RaiseArgtupleInvalid("alpha_ad_dyn_v4", 0, 19, 23, 9); __PYX_ERR(1, 38, __pyx_L3_error)
        }
        CYTHON_FALLTHROUGH;
        case 10:
        if (likely((values[10] = __Pyx_PyDict_GetItemStr(__pyx_kwds, __pyx_n_s_gamma)) != 0)) kw_args--;
        else {
          __Pyx_RaiseArgtupleInvalid("alpha_ad_dyn_v4", 0, 19, 23, 10); __PYX_ERR(1, 38, __pyx_L3_error)
        }
        CYTHON_FALLTHROUGH;
        case 11:
        if (likely((values[11] = __Pyx_PyDict_GetItemStr(__pyx_kwds, __pyx_n_s_lam)) != 0)) kw_args--;
        else {
          __Pyx_RaiseArgtupleInvalid("alpha_ad_dyn_v4", 0, 19, 23, 11); __PYX_ERR(1, 38, __pyx_L3_error)
        }
        CYTHON_FALLTHROUGH;
        case 12:
        if (likely((values[12] = __Pyx_PyDict_GetItemStr(__pyx_kwds, __pyx_n_s_c1)) != 0)) kw_args--;
        else {
          __Pyx_RaiseArgtupleInvalid("alpha_ad_dyn_v4", 0, 19, 23, 12); __PYX_ERR(1, 38, __pyx_L3_error)
        }
        CYTHON_FALLTHROUGH;
        case 13:
        if (likely((values[13] = __Pyx_PyDict_GetItemStr(__pyx_kwds, __pyx_n_s_c2)) != 0)) kw_args--;
        else {
          __Pyx_RaiseArgtupleInvalid("alpha_ad_dyn_v4", 0, 19, 23, 13); __PYX_ERR(1, 38, __pyx_L3_error)
        }
        CYTHON_FALLTHROUGH;
        case 14:
        if (likely((values[14] = __Pyx_PyDict_GetItemStr(__pyx_kwds, __pyx_n_s_beta_scalar)) != 0)) kw_args--;
        else {
          __Pyx_RaiseArgtupleInvalid("alpha_ad_dyn_v4", 0, 19, 23, 14); __PYX_ERR(1, 38, __pyx_L3_error)
        }
        CYTHON_FALLTHROUGH;
        case 15:
        if (likely((values[15] = __Pyx_PyDict_GetItemStr(__pyx_kwds, __pyx_n_s_hyper)) != 0)) kw_args--;
        else {
          __Pyx_RaiseArgtupleInvalid("alpha_ad_dyn_v4", 0, 19, 23, 15); __PYX_ERR(1, 38, __pyx_L3_error)
        }
        CYTHON_FALLTHROUGH;
        case 16:
        if (likely((values[16] = __Pyx_PyDict_GetItemStr(__pyx_kwds, __pyx_n_s_seed)) != 0)) kw_args--;
        else {
          __Pyx_RaiseArgtupleInvalid("alpha_ad_dyn_v4", 0, 19, 23, 16); __PYX_ERR(1, 38, __pyx_L3_error)
        }
        CYTHON_FALLTHROUGH;
        case 17:
        if (likely((values[17] = __Pyx_PyDict_GetItemStr(__pyx_kwds, __pyx_n_s_alpha_init)) != 0)) kw_args--;
        else {
          __Pyx_RaiseArgtupleInvalid("alpha_ad_dyn_v4", 0, 19, 23, 17); __PYX_ERR(1, 38, __pyx_L3_error)
        }
        CYTHON_FALLTHROUGH;
        case 18:
        if (likely((values[18] = __Pyx_PyDict_GetItemStr(__pyx_kwds, __pyx_n_s_sigma_init)) != 0)) kw_args--;
        else {
          __Pyx_RaiseArgtupleInvalid("alpha_ad_dyn_v4", 0, 19, 23, 18); __PYX_ERR(1, 38, __pyx_L3_error)
        }
        CYTHON_FALLTHROUGH;
        case 19:
        if (kw_args > 0) {
          PyObject* value = __Pyx_PyDict_GetItemStr(__pyx_kwds, __pyx_n_s_evo_steps);
          if (value) { values[19] = value; kw_args--; }
        }
        CYTHON_FALLTHROUGH;
        case 20:
        if (kw_args > 0) {
          PyObject* value = __Pyx_PyDict_GetItemStr(__pyx_kwds, __pyx_n_s_S_density);
          if (value) { values[20] = value; kw_args--; }
        }
        CYTHON_FALLTHROUGH;
        case 21:
        if (kw_args > 0) {
          PyObject* value = __Pyx_PyDict_GetItemStr(__pyx_kwds, __pyx_n_s_I_density);
          if (value) { values[21] = value; kw_args--; }
        }
        CYTHON_FALLTHROUGH;
        case 22:
        if (kw_args > 0) {
          PyObject* value = __Pyx_PyDict_GetItemStr(__pyx_kwds, __pyx_n_s_H_density);
          if (value) { values[22] = value; kw_args--; }
        }
      }
      if (unlikely(kw_args > 0)) {
        if (unlikely(__Pyx_ParseOptionalKeywords(__pyx_kwds, __pyx_pyargnames, 0, values, pos_args, "alpha_ad_dyn_v4") < 0)) __PYX_ERR(1, 38, __pyx_L3_error)
      }
    } else {
      switch (PyTuple_GET_SIZE(__pyx_args)) {
        case 23: values[22] = PyTuple_GET_ITEM(__pyx_args, 22);
        CYTHON_FALLTHROUGH;
        case 22: values[21] = PyTuple_GET_ITEM(__pyx_args, 21);
        CYTHON_FALLTHROUGH;
        case 21: values[20] = PyTuple_GET_ITEM(__pyx_args, 20);
        CYTHON_FALLTHROUGH;
        case 20: values[19] = PyTuple_GET_ITEM(__pyx_args, 19);
        CYTHON_FALLTHROUGH;
        case 19: values[18] = PyTuple_GET_ITEM(__pyx_args, 18);
        values[17] = PyTuple_GET_ITEM(__pyx_args, 17);
        values[16] = PyTuple_GET_ITEM(__pyx_args, 16);
        values[15] = PyTuple_GET_ITEM(__pyx_args, 15);
        values[14] = PyTuple_GET_ITEM(__pyx_args, 14);
        values[13] = PyTuple_GET_ITEM(__pyx_args, 13);
        values[12] = PyTuple_GET_ITEM(__pyx_args, 12);
        values[11] = PyTuple_GET_ITEM(__pyx_args, 11);
        values[10] = PyTuple_GET_ITEM(__pyx_args, 10);
        values[9] = PyTuple_GET_ITEM(__pyx_args, 9);
        values[8] = PyTuple_GET_ITEM(__pyx_args, 8);
        values[7] = PyTuple_GET_ITEM(__pyx_args, 7);
        values[6] = PyTuple_GET_ITEM(__pyx_args, 6);
        values[5] = PyTuple_GET_ITEM(__pyx_args, 5);
        values[4] = PyTuple_GET_ITEM(__pyx_args, 4);
        values[3] = PyTuple_GET_ITEM(__pyx_args, 3);
        values[2] = PyTuple_GET_ITEM(__pyx_args, 2);
        values[1] = PyTuple_GET_ITEM(__pyx_args, 1);
        values[0] = PyTuple_GET_ITEM(__pyx_args, 0);
        break;
        default: goto __pyx_L5_argtuple_error;
      }
    }
    __pyx_v_beta_max = __pyx_PyFloat_AsFloat(values[0]); if (unlikely((__pyx_v_beta_max == (float)-1) && PyErr_Occurred())) __PYX_ERR(1, 38, __pyx_L3_error)
    __pyx_v_beta_lin = __pyx_PyFloat_AsFloat(values[1]); if (unlikely((__pyx_v_beta_lin == (float)-1) && PyErr_Occurred())) __PYX_ERR(1, 38, __pyx_L3_error)
    __pyx_v_alpha_max = __pyx_PyFloat_AsFloat(values[2]); if (unlikely((__pyx_v_alpha_max == (float)-1) && PyErr_Occurred())) __PYX_ERR(1, 38, __pyx_L3_error)
    __pyx_v_alpha_min = __pyx_PyFloat_AsFloat(values[3]); if (unlikely((__pyx_v_alpha_min == (float)-1) && PyErr_Occurred())) __PYX_ERR(1, 38, __pyx_L3_error)
    __pyx_v_sigma_max = __pyx_PyFloat_AsFloat(values[4]); if (unlikely((__pyx_v_sigma_max == (float)-1) && PyErr_Occurred())) __PYX_ERR(1, 38, __pyx_L3_error)
    __pyx_v_b = __pyx_PyFloat_AsFloat(values[5]); if (unlikely((__pyx_v_b == (float)-1) && PyErr_Occurred())) __PYX_ERR(1, 38, __pyx_L3_error)
    __pyx_v_q = __pyx_PyFloat_AsFloat(values[6]); if (unlikely((__pyx_v_q == (float)-1) && PyErr_Occurred())) __PYX_ERR(1, 38, __pyx_L3_error)
    __pyx_v_d = __pyx_PyFloat_AsFloat(values[7]); if (unlikely((__pyx_v_d == (float)-1) && PyErr_Occurred())) __PYX_ERR(1, 38, __pyx_L3_error)
    __pyx_v_rho = __pyx_PyFloat_AsFloat(values[8]); if (unlikely((__pyx_v_rho == (float)-1) && PyErr_Occurred())) __PYX_ERR(1, 38, __pyx_L3_error)
    __pyx_v_eta = __pyx_PyFloat_AsFloat(values[9]); if (unlikely((__pyx_v_eta == (float)-1) && PyErr_Occurred())) __PYX_ERR(1, 38, __pyx_L3_error)
    __pyx_v_gamma = __pyx_PyFloat_AsFloat(values[10]); if (unlikely((__pyx_v_gamma == (float)-1) && PyErr_Occurred())) __PYX_ERR(1, 38, __pyx_L3_error)
    __pyx_v_lam = __pyx_PyFloat_AsFloat(values[11]); if (unlikely((__pyx_v_lam == (float)-1) && PyErr_Occurred())) __PYX_ERR(1, 38, __pyx_L3_error)
    __pyx_v_c1 = __pyx_PyFloat_AsFloat(values[12]); if (unlikely((__pyx_v_c1 == (float)-1) && PyErr_Occurred())) __PYX_ERR(1, 38, __pyx_L3_error)
    __pyx_v_c2 = __pyx_PyFloat_AsFloat(values[13]); if (unlikely((__pyx_v_c2 == (float)-1) && PyErr_Occurred())) __PYX_ERR(1, 38, __pyx_L3_error)
    __pyx_v_beta_scalar = __pyx_PyFloat_AsFloat(values[14]); if (unlikely((__pyx_v_beta_scalar == (float)-1) && PyErr_Occurred())) __PYX_ERR(1, 38, __pyx_L3_error)
    __pyx_v_hyper = __pyx_PyFloat_AsFloat(values[15]); if (unlikely((__pyx_v_hyper == (float)-1) && PyErr_Occurred())) __PYX_ERR(1, 38, __pyx_L3_error)
    __pyx_v_seed = __Pyx_PyInt_As_int(values[16]); if (unlikely((__pyx_v_seed == (int)-1) && PyErr_Occurred())) __PYX_ERR(1, 38, __pyx_L3_error)
    __pyx_v_alpha_init = __Pyx_PyInt_As_int(values[17]); if (unlikely((__pyx_v_alpha_init == (int)-1) && PyErr_Occurred())) __PYX_ERR(1, 38, __pyx_L3_error)
    __pyx_v_sigma_init = __Pyx_PyInt_As_int(values[18]); if (unlikely((__pyx_v_sigma_init == (int)-1) && PyErr_Occurred())) __PYX_ERR(1, 38, __pyx_L3_error)
    if (values[19]) {
      __pyx_v_evo_steps = __Pyx_PyInt_As_int(values[19]); if (unlikely((__pyx_v_evo_steps == (int)-1) && PyErr_Occurred())) __PYX_ERR(1, 38, __pyx_L3_error)
    } else {
      __pyx_v_evo_steps = ((int)0x3E8);
    }
    if (values[20]) {
      __pyx_v_S_density = __pyx_PyFloat_AsFloat(values[20]); if (unlikely((__pyx_v_S_density == (float)-1) && PyErr_Occurred())) __PYX_ERR(1, 38, __pyx_L3_error)
    } else {
      __pyx_v_S_density = ((float)4.0);
    }
    if (values[21]) {
      __pyx_v_I_density = __pyx_PyFloat_AsFloat(values[21]); if (unlikely((__pyx_v_I_density == (float)-1) && PyErr_Occurred())) __PYX_ERR(1, 38, __pyx_L3_error)
    } else {
      __pyx_v_I_density = ((float)4.0);
    }
    if (values[22]) {
      __pyx_v_H_density = __pyx_PyFloat_AsFloat(values[22]); if (unlikely((__pyx_v_H_density == (float)-1) && PyErr_Occurred())) __PYX_ERR(1, 38, __pyx_L3_error)
    } else {
      __pyx_v_H_density = ((float)4.0);
    }
  }
  goto __pyx_L4_argument_unpacking_done;
  __pyx_L5_argtuple_error:;
  __Pyx_RaiseArgtupleInvalid("alpha_ad_dyn_v4", 0, 19, 23, PyTuple_GET_SIZE(__pyx_args)); __PYX_ERR(1, 38, __pyx_L3_error)
  __pyx_L3_error:;
  __Pyx_AddTraceback("runner.PySolver.alpha_ad_dyn_v4", __pyx_clineno, __pyx_lineno, __pyx_filename);
  __Pyx_RefNannyFinishContext();
  return NULL;
  __pyx_L4_argument_unpacking_done:;
  __pyx_r = __pyx_pf_6runner_8PySolver_10alpha_ad_dyn_v4(((struct __pyx_obj_6runner_PySolver *)__pyx_v_self), __pyx_v_beta_max, __pyx_v_beta_lin, __pyx_v_alpha_max, __pyx_v_alpha_min, __pyx_v_sigma_max, __pyx_v_b, __pyx_v_q, __pyx_v_d, __pyx_v_rho, __pyx_v_eta, __pyx_v_gamma, __pyx_v_lam, __pyx_v_c1, __pyx_v_c2, __pyx_v_beta_scalar, __pyx_v_hyper, __pyx_v_seed, __pyx_v_alpha_init, __pyx_v_sigma_init, __pyx_v_evo_steps, __pyx_v_S_density, __pyx_v_I_density, __pyx_v_H_density);

  /* function exit code */
  __Pyx_RefNannyFinishContext();
  return __pyx_r;
}

static PyObject *__pyx_pf_6runner_8PySolver_10alpha_ad_dyn_v4(struct __pyx_obj_6runner_PySolver *__pyx_v_self, float __pyx_v_beta_max, float __pyx_v_beta_lin, float __pyx_v_alpha_max, float __pyx_v_alpha_min, float __pyx_v_sigma_max, float __pyx_v_b, float __pyx_v_q, float __pyx_v_d, float __pyx_v_rho, float __pyx_v_eta, float __pyx_v_gamma, float __pyx_v_lam, float __pyx_v_c1, float __pyx_v_c2, float __pyx_v_beta_scalar, float __pyx_v_hyper, int __pyx_v_seed, int __pyx_v_alpha_init, int __pyx_v_sigma_init, int __pyx_v_evo_steps, float __pyx_v_S_density, float __pyx_v_I_density, float __pyx_v_H_density) {
  PyObject *__pyx_r = NULL;
  __Pyx_RefNannyDeclarations
  __Pyx_RefNannySetupContext("alpha_ad_dyn_v4", 0);

  /* "runner.pyx":41
 * 
 * 
 *         self.cpp_solver.alpha_evo_only_v4(beta_max, beta_lin, alpha_max, alpha_min, sigma_max, b, q, d, rho, eta, gamma, lam, c1, c2, beta_scalar, hyper, seed, alpha_init, sigma_init, evo_steps, S_density, I_density, H_density)             # <<<<<<<<<<<<<<
 * 
 *         return
 */
  __pyx_v_self->cpp_solver.alpha_evo_only_v4(__pyx_v_beta_max, __pyx_v_beta_lin, __pyx_v_alpha_max, __pyx_v_alpha_min, __pyx_v_sigma_max, __pyx_v_b, __pyx_v_q, __pyx_v_d, __pyx_v_rho, __pyx_v_eta, __pyx_v_gamma, __pyx_v_lam, __pyx_v_c1, __pyx_v_c2, __pyx_v_beta_scalar, __pyx_v_hyper, __pyx_v_seed, __pyx_v_alpha_init, __pyx_v_sigma_init, __pyx_v_evo_steps, __pyx_v_S_density, __pyx_v_I_density, __pyx_v_H_density);

  /* "runner.pyx":43
 *         self.cpp_solver.alpha_evo_only_v4(beta_max, beta_lin, alpha_max, alpha_min, sigma_max, b, q, d, rho, eta, gamma, lam, c1, c2, beta_scalar, hyper, seed, alpha_init, sigma_init, evo_steps, S_density, I_density, H_density)
 * 
 *         return             # <<<<<<<<<<<<<<
 * 
 *     def eco_steady_state(self, float beta_value, float alpha_value, float sigma_value, float b, float q, float d, float rho, float eta, float gamma, float lam, float c1, float c2, float hyper, int seed, float S_density = 4.0, float I_density = 4.0, float H_density = 4.0):
 */
  __Pyx_XDECREF(__pyx_r);
  __pyx_r = Py_None; __Pyx_INCREF(Py_None);
  goto __pyx_L0;

  /* "runner.pyx":38
 *         return
 * 
 *     def alpha_ad_dyn_v4(self, float beta_max, float beta_lin, float alpha_max, float alpha_min, float sigma_max, float b, float q, float d, float rho, float eta, float gamma, float lam, float c1, float c2, float beta_scalar, float hyper, int seed, int alpha_init, int sigma_init, int evo_steps = 1000, float S_density = 4.0, float I_density = 4.0, float H_density = 4.0):             # <<<<<<<<<<<<<<
 * 
 * 
 */

  /* function exit code */
  __pyx_L0:;
  __Pyx_XGIVEREF(__pyx_r);
  __Pyx_RefNannyFinishContext();
  return __pyx_r;
}

/* "runner.pyx":45
 *         return
 * 
 *     def eco_steady_state(self, float beta_value, float alpha_value, float sigma_value, float b, float q, float d, float rho, float eta, float gamma, float lam, float c1, float c2, float hyper, int seed, float S_density = 4.0, float I_density = 4.0, float H_density = 4.0):             # <<<<<<<<<<<<<<
 * 
 *         cdef float s, i, h,
 */

/* Python wrapper */
static PyObject *__pyx_pw_6runner_8PySolver_13eco_steady_state(PyObject *__pyx_v_self, PyObject *__pyx_args, PyObject *__pyx_kwds); /*proto*/
static PyObject *__pyx_pw_6runner_8PySolver_13eco_steady_state(PyObject *__pyx_v_self, PyObject *__pyx_args, PyObject *__pyx_kwds) {
  float __pyx_v_beta_value;
  float __pyx_v_alpha_value;
  float __pyx_v_sigma_value;
  float __pyx_v_b;
  float __pyx_v_q;
  float __pyx_v_d;
  float __pyx_v_rho;
  float __pyx_v_eta;
  float __pyx_v_gamma;
  float __pyx_v_lam;
  float __pyx_v_c1;
  float __pyx_v_c2;
  float __pyx_v_hyper;
  int __pyx_v_seed;
  float __pyx_v_S_density;
  float __pyx_v_I_density;
  float __pyx_v_H_density;
  int __pyx_lineno = 0;
  const char *__pyx_filename = NULL;
  int __pyx_clineno = 0;
  PyObject *__pyx_r = 0;
  __Pyx_RefNannyDeclarations
  __Pyx_RefNannySetupContext("eco_steady_state (wrapper)", 0);
  {
    static PyObject **__pyx_pyargnames[] = {&__pyx_n_s_beta_value,&__pyx_n_s_alpha_value,&__pyx_n_s_sigma_value,&__pyx_n_s_b,&__pyx_n_s_q,&__pyx_n_s_d,&__pyx_n_s_rho,&__pyx_n_s_eta,&__pyx_n_s_gamma,&__pyx_n_s_lam,&__pyx_n_s_c1,&__pyx_n_s_c2,&__pyx_n_s_hyper,&__pyx_n_s_seed,&__pyx_n_s_S_density,&__pyx_n_s_I_density,&__pyx_n_s_H_density,0};
    PyObject* values[17] = {0,0,0,0,0,0,0,0,0,0,0,0,0,0,0,0,0};
    if (unlikely(__pyx_kwds)) {
      Py_ssize_t kw_args;
      const Py_ssize_t pos_args = PyTuple_GET_SIZE(__pyx_args);
      switch (pos_args) {
        case 17: values[16] = PyTuple_GET_ITEM(__pyx_args, 16);
        CYTHON_FALLTHROUGH;
        case 16: values[15] = PyTuple_GET_ITEM(__pyx_args, 15);
        CYTHON_FALLTHROUGH;
        case 15: values[14] = PyTuple_GET_ITEM(__pyx_args, 14);
        CYTHON_FALLTHROUGH;
        case 14: values[13] = PyTuple_GET_ITEM(__pyx_args, 13);
        CYTHON_FALLTHROUGH;
        case 13: values[12] = PyTuple_GET_ITEM(__pyx_args, 12);
        CYTHON_FALLTHROUGH;
        case 12: values[11] = PyTuple_GET_ITEM(__pyx_args, 11);
        CYTHON_FALLTHROUGH;
        case 11: values[10] = PyTuple_GET_ITEM(__pyx_args, 10);
        CYTHON_FALLTHROUGH;
        case 10: values[9] = PyTuple_GET_ITEM(__pyx_args, 9);
        CYTHON_FALLTHROUGH;
        case  9: values[8] = PyTuple_GET_ITEM(__pyx_args, 8);
        CYTHON_FALLTHROUGH;
        case  8: values[7] = PyTuple_GET_ITEM(__pyx_args, 7);
        CYTHON_FALLTHROUGH;
        case  7: values[6] = PyTuple_GET_ITEM(__pyx_args, 6);
        CYTHON_FALLTHROUGH;
        case  6: values[5] = PyTuple_GET_ITEM(__pyx_args, 5);
        CYTHON_FALLTHROUGH;
        case  5: values[4] = PyTuple_GET_ITEM(__pyx_args, 4);
        CYTHON_FALLTHROUGH;
        case  4: values[3] = PyTuple_GET_ITEM(__pyx_args, 3);
        CYTHON_FALLTHROUGH;
        case  3: values[2] = PyTuple_GET_ITEM(__pyx_args, 2);
        CYTHON_FALLTHROUGH;
        case  2: values[1] = PyTuple_GET_ITEM(__pyx_args, 1);
        CYTHON_FALLTHROUGH;
        case  1: values[0] = PyTuple_GET_ITEM(__pyx_args, 0);
        CYTHON_FALLTHROUGH;
        case  0: break;
        default: goto __pyx_L5_argtuple_error;
      }
      kw_args = PyDict_Size(__pyx_kwds);
      switch (pos_args) {
        case  0:
        if (likely((values[0] = __Pyx_PyDict_GetItemStr(__pyx_kwds, __pyx_n_s_beta_value)) != 0)) kw_args--;
        else goto __pyx_L5_argtuple_error;
        CYTHON_FALLTHROUGH;
        case  1:
        if (likely((values[1] = __Pyx_PyDict_GetItemStr(__pyx_kwds, __pyx_n_s_alpha_value)) != 0)) kw_args--;
        else {
          __Pyx_RaiseArgtupleInvalid("eco_steady_state", 0, 14, 17, 1); __PYX_ERR(1, 45, __pyx_L3_error)
        }
        CYTHON_FALLTHROUGH;
        case  2:
        if (likely((values[2] = __Pyx_PyDict_GetItemStr(__pyx_kwds, __pyx_n_s_sigma_value)) != 0)) kw_args--;
        else {
          __Pyx_RaiseArgtupleInvalid("eco_steady_state", 0, 14, 17, 2); __PYX_ERR(1, 45, __pyx_L3_error)
        }
        CYTHON_FALLTHROUGH;
        case  3:
        if (likely((values[3] = __Pyx_PyDict_GetItemStr(__pyx_kwds, __pyx_n_s_b)) != 0)) kw_args--;
        else {
          __Pyx_RaiseArgtupleInvalid("eco_steady_state", 0, 14, 17, 3); __PYX_ERR(1, 45, __pyx_L3_error)
        }
        CYTHON_FALLTHROUGH;
        case  4:
        if (likely((values[4] = __Pyx_PyDict_GetItemStr(__pyx_kwds, __pyx_n_s_q)) != 0)) kw_args--;
        else {
          __Pyx_RaiseArgtupleInvalid("eco_steady_state", 0, 14, 17, 4); __PYX_ERR(1, 45, __pyx_L3_error)
        }
        CYTHON_FALLTHROUGH;
        case  5:
        if (likely((values[5] = __Pyx_PyDict_GetItemStr(__pyx_kwds, __pyx_n_s_d)) != 0)) kw_args--;
        else {
          __Pyx_RaiseArgtupleInvalid("eco_steady_state", 0, 14, 17, 5); __PYX_ERR(1, 45, __pyx_L3_error)
        }
        CYTHON_FALLTHROUGH;
        case  6:
        if (likely((values[6] = __Pyx_PyDict_GetItemStr(__pyx_kwds, __pyx_n_s_rho)) != 0)) kw_args--;
        else {
          __Pyx_RaiseArgtupleInvalid("eco_steady_state", 0, 14, 17, 6); __PYX_ERR(1, 45, __pyx_L3_error)
        }
        CYTHON_FALLTHROUGH;
        case  7:
        if (likely((values[7] = __Pyx_PyDict_GetItemStr(__pyx_kwds, __pyx_n_s_eta)) != 0)) kw_args--;
        else {
          __Pyx_RaiseArgtupleInvalid("eco_steady_state", 0, 14, 17, 7); __PYX_ERR(1, 45, __pyx_L3_error)
        }
        CYTHON_FALLTHROUGH;
        case  8:
        if (likely((values[8] = __Pyx_PyDict_GetItemStr(__pyx_kwds, __pyx_n_s_gamma)) != 0)) kw_args--;
        else {
          __Pyx_RaiseArgtupleInvalid("eco_steady_state", 0, 14, 17, 8); __PYX_ERR(1, 45, __pyx_L3_error)
        }
        CYTHON_FALLTHROUGH;
        case  9:
        if (likely((values[9] = __Pyx_PyDict_GetItemStr(__pyx_kwds, __pyx_n_s_lam)) != 0)) kw_args--;
        else {
          __Pyx_RaiseArgtupleInvalid("eco_steady_state", 0, 14, 17, 9); __PYX_ERR(1, 45, __pyx_L3_error)
        }
        CYTHON_FALLTHROUGH;
        case 10:
        if (likely((values[10] = __Pyx_PyDict_GetItemStr(__pyx_kwds, __pyx_n_s_c1)) != 0)) kw_args--;
        else {
          __Pyx_RaiseArgtupleInvalid("eco_steady_state", 0, 14, 17, 10); __PYX_ERR(1, 45, __pyx_L3_error)
        }
        CYTHON_FALLTHROUGH;
        case 11:
        if (likely((values[11] = __Pyx_PyDict_GetItemStr(__pyx_kwds, __pyx_n_s_c2)) != 0)) kw_args--;
        else {
          __Pyx_RaiseArgtupleInvalid("eco_steady_state", 0, 14, 17, 11); __PYX_ERR(1, 45, __pyx_L3_error)
        }
        CYTHON_FALLTHROUGH;
        case 12:
        if (likely((values[12] = __Pyx_PyDict_GetItemStr(__pyx_kwds, __pyx_n_s_hyper)) != 0)) kw_args--;
        else {
          __Pyx_RaiseArgtupleInvalid("eco_steady_state", 0, 14, 17, 12); __PYX_ERR(1, 45, __pyx_L3_error)
        }
        CYTHON_FALLTHROUGH;
        case 13:
        if (likely((values[13] = __Pyx_PyDict_GetItemStr(__pyx_kwds, __pyx_n_s_seed)) != 0)) kw_args--;
        else {
          __Pyx_RaiseArgtupleInvalid("eco_steady_state", 0, 14, 17, 13); __PYX_ERR(1, 45, __pyx_L3_error)
        }
        CYTHON_FALLTHROUGH;
        case 14:
        if (kw_args > 0) {
          PyObject* value = __Pyx_PyDict_GetItemStr(__pyx_kwds, __pyx_n_s_S_density);
          if (value) { values[14] = value; kw_args--; }
        }
        CYTHON_FALLTHROUGH;
        case 15:
        if (kw_args > 0) {
          PyObject* value = __Pyx_PyDict_GetItemStr(__pyx_kwds, __pyx_n_s_I_density);
          if (value) { values[15] = value; kw_args--; }
        }
        CYTHON_FALLTHROUGH;
        case 16:
        if (kw_args > 0) {
          PyObject* value = __Pyx_PyDict_GetItemStr(__pyx_kwds, __pyx_n_s_H_density);
          if (value) { values[16] = value; kw_args--; }
        }
      }
      if (unlikely(kw_args > 0)) {
        if (unlikely(__Pyx_ParseOptionalKeywords(__pyx_kwds, __pyx_pyargnames, 0, values, pos_args, "eco_steady_state") < 0)) __PYX_ERR(1, 45, __pyx_L3_error)
      }
    } else {
      switch (PyTuple_GET_SIZE(__pyx_args)) {
        case 17: values[16] = PyTuple_GET_ITEM(__pyx_args, 16);
        CYTHON_FALLTHROUGH;
        case 16: values[15] = PyTuple_GET_ITEM(__pyx_args, 15);
        CYTHON_FALLTHROUGH;
        case 15: values[14] = PyTuple_GET_ITEM(__pyx_args, 14);
        CYTHON_FALLTHROUGH;
        case 14: values[13] = PyTuple_GET_ITEM(__pyx_args, 13);
        values[12] = PyTuple_GET_ITEM(__pyx_args, 12);
        values[11] = PyTuple_GET_ITEM(__pyx_args, 11);
        values[10] = PyTuple_GET_ITEM(__pyx_args, 10);
        values[9] = PyTuple_GET_ITEM(__pyx_args, 9);
        values[8] = PyTuple_GET_ITEM(__pyx_args, 8);
        values[7] = PyTuple_GET_ITEM(__pyx_args, 7);
        values[6] = PyTuple_GET_ITEM(__pyx_args, 6);
        values[5] = PyTuple_GET_ITEM(__pyx_args, 5);
        values[4] = PyTuple_GET_ITEM(__pyx_args, 4);
        values[3] = PyTuple_GET_ITEM(__pyx_args, 3);
        values[2] = PyTuple_GET_ITEM(__pyx_args, 2);
        values[1] = PyTuple_GET_ITEM(__pyx_args, 1);
        values[0] = PyTuple_GET_ITEM(__pyx_args, 0);
        break;
        default: goto __pyx_L5_argtuple_error;
      }
    }
    __pyx_v_beta_value = __pyx_PyFloat_AsFloat(values[0]); if (unlikely((__pyx_v_beta_value == (float)-1) && PyErr_Occurred())) __PYX_ERR(1, 45, __pyx_L3_error)
    __pyx_v_alpha_value = __pyx_PyFloat_AsFloat(values[1]); if (unlikely((__pyx_v_alpha_value == (float)-1) && PyErr_Occurred())) __PYX_ERR(1, 45, __pyx_L3_error)
    __pyx_v_sigma_value = __pyx_PyFloat_AsFloat(values[2]); if (unlikely((__pyx_v_sigma_value == (float)-1) && PyErr_Occurred())) __PYX_ERR(1, 45, __pyx_L3_error)
    __pyx_v_b = __pyx_PyFloat_AsFloat(values[3]); if (unlikely((__pyx_v_b == (float)-1) && PyErr_Occurred())) __PYX_ERR(1, 45, __pyx_L3_error)
    __pyx_v_q = __pyx_PyFloat_AsFloat(values[4]); if (unlikely((__pyx_v_q == (float)-1) && PyErr_Occurred())) __PYX_ERR(1, 45, __pyx_L3_error)
    __pyx_v_d = __pyx_PyFloat_AsFloat(values[5]); if (unlikely((__pyx_v_d == (float)-1) && PyErr_Occurred())) __PYX_ERR(1, 45, __pyx_L3_error)
    __pyx_v_rho = __pyx_PyFloat_AsFloat(values[6]); if (unlikely((__pyx_v_rho == (float)-1) && PyErr_Occurred())) __PYX_ERR(1, 45, __pyx_L3_error)
    __pyx_v_eta = __pyx_PyFloat_AsFloat(values[7]); if (unlikely((__pyx_v_eta == (float)-1) && PyErr_Occurred())) __PYX_ERR(1, 45, __pyx_L3_error)
    __pyx_v_gamma = __pyx_PyFloat_AsFloat(values[8]); if (unlikely((__pyx_v_gamma == (float)-1) && PyErr_Occurred())) __PYX_ERR(1, 45, __pyx_L3_error)
    __pyx_v_lam = __pyx_PyFloat_AsFloat(values[9]); if (unlikely((__pyx_v_lam == (float)-1) && PyErr_Occurred())) __PYX_ERR(1, 45, __pyx_L3_error)
    __pyx_v_c1 = __pyx_PyFloat_AsFloat(values[10]); if (unlikely((__pyx_v_c1 == (float)-1) && PyErr_Occurred())) __PYX_ERR(1, 45, __pyx_L3_error)
    __pyx_v_c2 = __pyx_PyFloat_AsFloat(values[11]); if (unlikely((__pyx_v_c2 == (float)-1) && PyErr_Occurred())) __PYX_ERR(1, 45, __pyx_L3_error)
    __pyx_v_hyper = __pyx_PyFloat_AsFloat(values[12]); if (unlikely((__pyx_v_hyper == (float)-1) && PyErr_Occurred())) __PYX_ERR(1, 45, __pyx_L3_error)
    __pyx_v_seed = __Pyx_PyInt_As_int(values[13]); if (unlikely((__pyx_v_seed == (int)-1) && PyErr_Occurred())) __PYX_ERR(1, 45, __pyx_L3_error)
    if (values[14]) {
      __pyx_v_S_density = __pyx_PyFloat_AsFloat(values[14]); if (unlikely((__pyx_v_S_density == (float)-1) && PyErr_Occurred())) __PYX_ERR(1, 45, __pyx_L3_error)
    } else {
      __pyx_v_S_density = ((float)4.0);
    }
    if (values[15]) {
      __pyx_v_I_density = __pyx_PyFloat_AsFloat(values[15]); if (unlikely((__pyx_v_I_density == (float)-1) && PyErr_Occurred())) __PYX_ERR(1, 45, __pyx_L3_error)
    } else {
      __pyx_v_I_density = ((float)4.0);
    }
    if (values[16]) {
      __pyx_v_H_density = __pyx_PyFloat_AsFloat(values[16]); if (unlikely((__pyx_v_H_density == (float)-1) && PyErr_Occurred())) __PYX_ERR(1, 45, __pyx_L3_error)
    } else {
      __pyx_v_H_density = ((float)4.0);
    }
  }
  goto __pyx_L4_argument_unpacking_done;
  __pyx_L5_argtuple_error:;
  __Pyx_RaiseArgtupleInvalid("eco_steady_state", 0, 14, 17, PyTuple_GET_SIZE(__pyx_args)); __PYX_ERR(1, 45, __pyx_L3_error)
  __pyx_L3_error:;
  __Pyx_AddTraceback("runner.PySolver.eco_steady_state", __pyx_clineno, __pyx_lineno, __pyx_filename);
  __Pyx_RefNannyFinishContext();
  return NULL;
  __pyx_L4_argument_unpacking_done:;
  __pyx_r = __pyx_pf_6runner_8PySolver_12eco_steady_state(((struct __pyx_obj_6runner_PySolver *)__pyx_v_self), __pyx_v_beta_value, __pyx_v_alpha_value, __pyx_v_sigma_value, __pyx_v_b, __pyx_v_q, __pyx_v_d, __pyx_v_rho, __pyx_v_eta, __pyx_v_gamma, __pyx_v_lam, __pyx_v_c1, __pyx_v_c2, __pyx_v_hyper, __pyx_v_seed, __pyx_v_S_density, __pyx_v_I_density, __pyx_v_H_density);

  /* function exit code */
  __Pyx_RefNannyFinishContext();
  return __pyx_r;
}

static PyObject *__pyx_pf_6runner_8PySolver_12eco_steady_state(struct __pyx_obj_6runner_PySolver *__pyx_v_self, float __pyx_v_beta_value, float __pyx_v_alpha_value, float __pyx_v_sigma_value, float __pyx_v_b, float __pyx_v_q, float __pyx_v_d, float __pyx_v_rho, float __pyx_v_eta, float __pyx_v_gamma, float __pyx_v_lam, float __pyx_v_c1, float __pyx_v_c2, float __pyx_v_hyper, int __pyx_v_seed, float __pyx_v_S_density, float __pyx_v_I_density, float __pyx_v_H_density) {
  float __pyx_v_s;
  float __pyx_v_i;
  float __pyx_v_h;
  int __pyx_v_exit_flag;
  PyObject *__pyx_v_y = NULL;
  PyObject *__pyx_r = NULL;
  __Pyx_RefNannyDeclarations
  PyObject *__pyx_t_1 = NULL;
  PyObject *__pyx_t_2 = NULL;
  PyObject *__pyx_t_3 = NULL;
  PyObject *__pyx_t_4 = NULL;
  PyObject *__pyx_t_5 = NULL;
  int __pyx_lineno = 0;
  const char *__pyx_filename = NULL;
  int __pyx_clineno = 0;
  __Pyx_RefNannySetupContext("eco_steady_state", 0);

  /* "runner.pyx":49
 *         cdef float s, i, h,
 *         cdef int exit_flag
 *         self.cpp_solver.eco_dynamics(&s, &i, &h, &exit_flag, beta_value, alpha_value, sigma_value, b, q, d, rho, eta, gamma, lam, c1, c2, hyper, seed, S_density, I_density, H_density)             # <<<<<<<<<<<<<<
 * 
 *         y = [s,i,h, exit_flag]
 */
  __pyx_v_self->cpp_solver.eco_dynamics((&__pyx_v_s), (&__pyx_v_i), (&__pyx_v_h), (&__pyx_v_exit_flag), __pyx_v_beta_value, __pyx_v_alpha_value, __pyx_v_sigma_value, __pyx_v_b, __pyx_v_q, __pyx_v_d, __pyx_v_rho, __pyx_v_eta, __pyx_v_gamma, __pyx_v_lam, __pyx_v_c1, __pyx_v_c2, __pyx_v_hyper, __pyx_v_seed, __pyx_v_S_density, __pyx_v_I_density, __pyx_v_H_density);

  /* "runner.pyx":51
 *         self.cpp_solver.eco_dynamics(&s, &i, &h, &exit_flag, beta_value, alpha_value, sigma_value, b, q, d, rho, eta, gamma, lam, c1, c2, hyper, seed, S_density, I_density, H_density)
 * 
 *         y = [s,i,h, exit_flag]             # <<<<<<<<<<<<<<
 * 
 *         return y
 */
  __pyx_t_1 = PyFloat_FromDouble(__pyx_v_s); if (unlikely(!__pyx_t_1)) __PYX_ERR(1, 51, __pyx_L1_error)
  __Pyx_GOTREF(__pyx_t_1);
  __pyx_t_2 = PyFloat_FromDouble(__pyx_v_i); if (unlikely(!__pyx_t_2)) __PYX_ERR(1, 51, __pyx_L1_error)
  __Pyx_GOTREF(__pyx_t_2);
  __pyx_t_3 = PyFloat_FromDouble(__pyx_v_h); if (unlikely(!__pyx_t_3)) __PYX_ERR(1, 51, __pyx_L1_error)
  __Pyx_GOTREF(__pyx_t_3);
  __pyx_t_4 = __Pyx_PyInt_From_int(__pyx_v_exit_flag); if (unlikely(!__pyx_t_4)) __PYX_ERR(1, 51, __pyx_L1_error)
  __Pyx_GOTREF(__pyx_t_4);
  __pyx_t_5 = PyList_New(4); if (unlikely(!__pyx_t_5)) __PYX_ERR(1, 51, __pyx_L1_error)
  __Pyx_GOTREF(__pyx_t_5);
  __Pyx_GIVEREF(__pyx_t_1);
  PyList_SET_ITEM(__pyx_t_5, 0, __pyx_t_1);
  __Pyx_GIVEREF(__pyx_t_2);
  PyList_SET_ITEM(__pyx_t_5, 1, __pyx_t_2);
  __Pyx_GIVEREF(__pyx_t_3);
  PyList_SET_ITEM(__pyx_t_5, 2, __pyx_t_3);
  __Pyx_GIVEREF(__pyx_t_4);
  PyList_SET_ITEM(__pyx_t_5, 3, __pyx_t_4);
  __pyx_t_1 = 0;
  __pyx_t_2 = 0;
  __pyx_t_3 = 0;
  __pyx_t_4 = 0;
  __pyx_v_y = ((PyObject*)__pyx_t_5);
  __pyx_t_5 = 0;

  /* "runner.pyx":53
 *         y = [s,i,h, exit_flag]
 * 
 *         return y             # <<<<<<<<<<<<<<
 */
  __Pyx_XDECREF(__pyx_r);
  __Pyx_INCREF(__pyx_v_y);
  __pyx_r = __pyx_v_y;
  goto __pyx_L0;

  /* "runner.pyx":45
 *         return
 * 
 *     def eco_steady_state(self, float beta_value, float alpha_value, float sigma_value, float b, float q, float d, float rho, float eta, float gamma, float lam, float c1, float c2, float hyper, int seed, float S_density = 4.0, float I_density = 4.0, float H_density = 4.0):             # <<<<<<<<<<<<<<
 * 
 *         cdef float s, i, h,
 */

  /* function exit code */
  __pyx_L1_error:;
  __Pyx_XDECREF(__pyx_t_1);
  __Pyx_XDECREF(__pyx_t_2);
  __Pyx_XDECREF(__pyx_t_3);
  __Pyx_XDECREF(__pyx_t_4);
  __Pyx_XDECREF(__pyx_t_5);
  __Pyx_AddTraceback("runner.PySolver.eco_steady_state", __pyx_clineno, __pyx_lineno, __pyx_filename);
  __pyx_r = NULL;
  __pyx_L0:;
  __Pyx_XDECREF(__pyx_v_y);
  __Pyx_XGIVEREF(__pyx_r);
  __Pyx_RefNannyFinishContext();
  return __pyx_r;
}

/* "(tree fragment)":1
 * def __reduce_cython__(self):             # <<<<<<<<<<<<<<
 *     raise TypeError("no default __reduce__ due to non-trivial __cinit__")
 * def __setstate_cython__(self, __pyx_state):
 */

/* Python wrapper */
static PyObject *__pyx_pw_6runner_8PySolver_15__reduce_cython__(PyObject *__pyx_v_self, CYTHON_UNUSED PyObject *unused); /*proto*/
static PyObject *__pyx_pw_6runner_8PySolver_15__reduce_cython__(PyObject *__pyx_v_self, CYTHON_UNUSED PyObject *unused) {
  PyObject *__pyx_r = 0;
  __Pyx_RefNannyDeclarations
  __Pyx_RefNannySetupContext("__reduce_cython__ (wrapper)", 0);
  __pyx_r = __pyx_pf_6runner_8PySolver_14__reduce_cython__(((struct __pyx_obj_6runner_PySolver *)__pyx_v_self));

  /* function exit code */
  __Pyx_RefNannyFinishContext();
  return __pyx_r;
}

static PyObject *__pyx_pf_6runner_8PySolver_14__reduce_cython__(CYTHON_UNUSED struct __pyx_obj_6runner_PySolver *__pyx_v_self) {
  PyObject *__pyx_r = NULL;
  __Pyx_RefNannyDeclarations
  PyObject *__pyx_t_1 = NULL;
  int __pyx_lineno = 0;
  const char *__pyx_filename = NULL;
  int __pyx_clineno = 0;
  __Pyx_RefNannySetupContext("__reduce_cython__", 0);

  /* "(tree fragment)":2
 * def __reduce_cython__(self):
 *     raise TypeError("no default __reduce__ due to non-trivial __cinit__")             # <<<<<<<<<<<<<<
 * def __setstate_cython__(self, __pyx_state):
 *     raise TypeError("no default __reduce__ due to non-trivial __cinit__")
 */
  __pyx_t_1 = __Pyx_PyObject_Call(__pyx_builtin_TypeError, __pyx_tuple_, NULL); if (unlikely(!__pyx_t_1)) __PYX_ERR(0, 2, __pyx_L1_error)
  __Pyx_GOTREF(__pyx_t_1);
  __Pyx_Raise(__pyx_t_1, 0, 0, 0);
  __Pyx_DECREF(__pyx_t_1); __pyx_t_1 = 0;
  __PYX_ERR(0, 2, __pyx_L1_error)

  /* "(tree fragment)":1
 * def __reduce_cython__(self):             # <<<<<<<<<<<<<<
 *     raise TypeError("no default __reduce__ due to non-trivial __cinit__")
 * def __setstate_cython__(self, __pyx_state):
 */

  /* function exit code */
  __pyx_L1_error:;
  __Pyx_XDECREF(__pyx_t_1);
  __Pyx_AddTraceback("runner.PySolver.__reduce_cython__", __pyx_clineno, __pyx_lineno, __pyx_filename);
  __pyx_r = NULL;
  __Pyx_XGIVEREF(__pyx_r);
  __Pyx_RefNannyFinishContext();
  return __pyx_r;
}

/* "(tree fragment)":3
 * def __reduce_cython__(self):
 *     raise TypeError("no default __reduce__ due to non-trivial __cinit__")
 * def __setstate_cython__(self, __pyx_state):             # <<<<<<<<<<<<<<
 *     raise TypeError("no default __reduce__ due to non-trivial __cinit__")
 */

/* Python wrapper */
static PyObject *__pyx_pw_6runner_8PySolver_17__setstate_cython__(PyObject *__pyx_v_self, PyObject *__pyx_v___pyx_state); /*proto*/
static PyObject *__pyx_pw_6runner_8PySolver_17__setstate_cython__(PyObject *__pyx_v_self, PyObject *__pyx_v___pyx_state) {
  PyObject *__pyx_r = 0;
  __Pyx_RefNannyDeclarations
  __Pyx_RefNannySetupContext("__setstate_cython__ (wrapper)", 0);
  __pyx_r = __pyx_pf_6runner_8PySolver_16__setstate_cython__(((struct __pyx_obj_6runner_PySolver *)__pyx_v_self), ((PyObject *)__pyx_v___pyx_state));

  /* function exit code */
  __Pyx_RefNannyFinishContext();
  return __pyx_r;
}

static PyObject *__pyx_pf_6runner_8PySolver_16__setstate_cython__(CYTHON_UNUSED struct __pyx_obj_6runner_PySolver *__pyx_v_self, CYTHON_UNUSED PyObject *__pyx_v___pyx_state) {
  PyObject *__pyx_r = NULL;
  __Pyx_RefNannyDeclarations
  PyObject *__pyx_t_1 = NULL;
  int __pyx_lineno = 0;
  const char *__pyx_filename = NULL;
  int __pyx_clineno = 0;
  __Pyx_RefNannySetupContext("__setstate_cython__", 0);

  /* "(tree fragment)":4
 *     raise TypeError("no default __reduce__ due to non-trivial __cinit__")
 * def __setstate_cython__(self, __pyx_state):
 *     raise TypeError("no default __reduce__ due to non-trivial __cinit__")             # <<<<<<<<<<<<<<
 */
  __pyx_t_1 = __Pyx_PyObject_Call(__pyx_builtin_TypeError, __pyx_tuple__2, NULL); if (unlikely(!__pyx_t_1)) __PYX_ERR(0, 4, __pyx_L1_error)
  __Pyx_GOTREF(__pyx_t_1);
  __Pyx_Raise(__pyx_t_1, 0, 0, 0);
  __Pyx_DECREF(__pyx_t_1); __pyx_t_1 = 0;
  __PYX_ERR(0, 4, __pyx_L1_error)

  /* "(tree fragment)":3
 * def __reduce_cython__(self):
 *     raise TypeError("no default __reduce__ due to non-trivial __cinit__")
 * def __setstate_cython__(self, __pyx_state):             # <<<<<<<<<<<<<<
 *     raise TypeError("no default __reduce__ due to non-trivial __cinit__")
 */

  /* function exit code */
  __pyx_L1_error:;
  __Pyx_XDECREF(__pyx_t_1);
  __Pyx_AddTraceback("runner.PySolver.__setstate_cython__", __pyx_clineno, __pyx_lineno, __pyx_filename);
  __pyx_r = NULL;
  __Pyx_XGIVEREF(__pyx_r);
  __Pyx_RefNannyFinishContext();
  return __pyx_r;
}

static PyObject *__pyx_tp_new_6runner_PySolver(PyTypeObject *t, CYTHON_UNUSED PyObject *a, CYTHON_UNUSED PyObject *k) {
  struct __pyx_obj_6runner_PySolver *p;
  PyObject *o;
  if (likely((t->tp_flags & Py_TPFLAGS_IS_ABSTRACT) == 0)) {
    o = (*t->tp_alloc)(t, 0);
  } else {
    o = (PyObject *) PyBaseObject_Type.tp_new(t, __pyx_empty_tuple, 0);
  }
  if (unlikely(!o)) return 0;
  p = ((struct __pyx_obj_6runner_PySolver *)o);
  new((void*)&(p->cpp_solver)) solvers::Quick_solver();
  if (unlikely(__pyx_pw_6runner_8PySolver_1__cinit__(o, __pyx_empty_tuple, NULL) < 0)) goto bad;
  return o;
  bad:
  Py_DECREF(o); o = 0;
  return NULL;
}

static void __pyx_tp_dealloc_6runner_PySolver(PyObject *o) {
  struct __pyx_obj_6runner_PySolver *p = (struct __pyx_obj_6runner_PySolver *)o;
  #if CYTHON_USE_TP_FINALIZE
  if (unlikely(PyType_HasFeature(Py_TYPE(o), Py_TPFLAGS_HAVE_FINALIZE) && Py_TYPE(o)->tp_finalize) && (!PyType_IS_GC(Py_TYPE(o)) || !_PyGC_FINALIZED(o))) {
    if (PyObject_CallFinalizerFromDealloc(o)) return;
  }
  #endif
  __Pyx_call_destructor(p->cpp_solver);
  (*Py_TYPE(o)->tp_free)(o);
}

static PyMethodDef __pyx_methods_6runner_PySolver[] = {
  {"ad_dyn", (PyCFunction)(void*)(PyCFunctionWithKeywords)__pyx_pw_6runner_8PySolver_3ad_dyn, METH_VARARGS|METH_KEYWORDS, 0},
  {"alpha_ad_dyn", (PyCFunction)(void*)(PyCFunctionWithKeywords)__pyx_pw_6runner_8PySolver_5alpha_ad_dyn, METH_VARARGS|METH_KEYWORDS, 0},
  {"alpha_ad_dyn_v2", (PyCFunction)(void*)(PyCFunctionWithKeywords)__pyx_pw_6runner_8PySolver_7alpha_ad_dyn_v2, METH_VARARGS|METH_KEYWORDS, 0},
  {"alpha_ad_dyn_v3", (PyCFunction)(void*)(PyCFunctionWithKeywords)__pyx_pw_6runner_8PySolver_9alpha_ad_dyn_v3, METH_VARARGS|METH_KEYWORDS, 0},
  {"alpha_ad_dyn_v4", (PyCFunction)(void*)(PyCFunctionWithKeywords)__pyx_pw_6runner_8PySolver_11alpha_ad_dyn_v4, METH_VARARGS|METH_KEYWORDS, 0},
  {"eco_steady_state", (PyCFunction)(void*)(PyCFunctionWithKeywords)__pyx_pw_6runner_8PySolver_13eco_steady_state, METH_VARARGS|METH_KEYWORDS, 0},
  {"__reduce_cython__", (PyCFunction)__pyx_pw_6runner_8PySolver_15__reduce_cython__, METH_NOARGS, 0},
  {"__setstate_cython__", (PyCFunction)__pyx_pw_6runner_8PySolver_17__setstate_cython__, METH_O, 0},
  {0, 0, 0, 0}
};

static PyTypeObject __pyx_type_6runner_PySolver = {
  PyVarObject_HEAD_INIT(0, 0)
  "runner.PySolver", /*tp_name*/
  sizeof(struct __pyx_obj_6runner_PySolver), /*tp_basicsize*/
  0, /*tp_itemsize*/
  __pyx_tp_dealloc_6runner_PySolver, /*tp_dealloc*/
  #if PY_VERSION_HEX < 0x030800b4
  0, /*tp_print*/
  #endif
  #if PY_VERSION_HEX >= 0x030800b4
  0, /*tp_vectorcall_offset*/
  #endif
  0, /*tp_getattr*/
  0, /*tp_setattr*/
  #if PY_MAJOR_VERSION < 3
  0, /*tp_compare*/
  #endif
  #if PY_MAJOR_VERSION >= 3
  0, /*tp_as_async*/
  #endif
  0, /*tp_repr*/
  0, /*tp_as_number*/
  0, /*tp_as_sequence*/
  0, /*tp_as_mapping*/
  0, /*tp_hash*/
  0, /*tp_call*/
  0, /*tp_str*/
  0, /*tp_getattro*/
  0, /*tp_setattro*/
  0, /*tp_as_buffer*/
  Py_TPFLAGS_DEFAULT|Py_TPFLAGS_HAVE_VERSION_TAG|Py_TPFLAGS_CHECKTYPES|Py_TPFLAGS_HAVE_NEWBUFFER|Py_TPFLAGS_BASETYPE, /*tp_flags*/
  0, /*tp_doc*/
  0, /*tp_traverse*/
  0, /*tp_clear*/
  0, /*tp_richcompare*/
  0, /*tp_weaklistoffset*/
  0, /*tp_iter*/
  0, /*tp_iternext*/
  __pyx_methods_6runner_PySolver, /*tp_methods*/
  0, /*tp_members*/
  0, /*tp_getset*/
  0, /*tp_base*/
  0, /*tp_dict*/
  0, /*tp_descr_get*/
  0, /*tp_descr_set*/
  0, /*tp_dictoffset*/
  0, /*tp_init*/
  0, /*tp_alloc*/
  __pyx_tp_new_6runner_PySolver, /*tp_new*/
  0, /*tp_free*/
  0, /*tp_is_gc*/
  0, /*tp_bases*/
  0, /*tp_mro*/
  0, /*tp_cache*/
  0, /*tp_subclasses*/
  0, /*tp_weaklist*/
  0, /*tp_del*/
  0, /*tp_version_tag*/
  #if PY_VERSION_HEX >= 0x030400a1
  0, /*tp_finalize*/
  #endif
  #if PY_VERSION_HEX >= 0x030800b1 && (!CYTHON_COMPILING_IN_PYPY || PYPY_VERSION_NUM >= 0x07030800)
  0, /*tp_vectorcall*/
  #endif
  #if PY_VERSION_HEX >= 0x030800b4 && PY_VERSION_HEX < 0x03090000
  0, /*tp_print*/
  #endif
  #if CYTHON_COMPILING_IN_PYPY && PY_VERSION_HEX >= 0x03090000
  0, /*tp_pypy_flags*/
  #endif
};

static PyMethodDef __pyx_methods[] = {
  {0, 0, 0, 0}
};

#if PY_MAJOR_VERSION >= 3
#if CYTHON_PEP489_MULTI_PHASE_INIT
static PyObject* __pyx_pymod_create(PyObject *spec, PyModuleDef *def); /*proto*/
static int __pyx_pymod_exec_runner(PyObject* module); /*proto*/
static PyModuleDef_Slot __pyx_moduledef_slots[] = {
  {Py_mod_create, (void*)__pyx_pymod_create},
  {Py_mod_exec, (void*)__pyx_pymod_exec_runner},
  {0, NULL}
};
#endif

static struct PyModuleDef __pyx_moduledef = {
    PyModuleDef_HEAD_INIT,
    "runner",
    0, /* m_doc */
  #if CYTHON_PEP489_MULTI_PHASE_INIT
    0, /* m_size */
  #else
    -1, /* m_size */
  #endif
    __pyx_methods /* m_methods */,
  #if CYTHON_PEP489_MULTI_PHASE_INIT
    __pyx_moduledef_slots, /* m_slots */
  #else
    NULL, /* m_reload */
  #endif
    NULL, /* m_traverse */
    NULL, /* m_clear */
    NULL /* m_free */
};
#endif
#ifndef CYTHON_SMALL_CODE
#if defined(__clang__)
    #define CYTHON_SMALL_CODE
#elif defined(__GNUC__) && (__GNUC__ > 4 || (__GNUC__ == 4 && __GNUC_MINOR__ >= 3))
    #define CYTHON_SMALL_CODE __attribute__((cold))
#else
    #define CYTHON_SMALL_CODE
#endif
#endif

static __Pyx_StringTabEntry __pyx_string_tab[] = {
  {&__pyx_n_s_H_density, __pyx_k_H_density, sizeof(__pyx_k_H_density), 0, 0, 1, 1},
  {&__pyx_n_s_I_density, __pyx_k_I_density, sizeof(__pyx_k_I_density), 0, 0, 1, 1},
  {&__pyx_n_s_PySolver, __pyx_k_PySolver, sizeof(__pyx_k_PySolver), 0, 0, 1, 1},
  {&__pyx_n_s_S_density, __pyx_k_S_density, sizeof(__pyx_k_S_density), 0, 0, 1, 1},
  {&__pyx_n_s_TypeError, __pyx_k_TypeError, sizeof(__pyx_k_TypeError), 0, 0, 1, 1},
  {&__pyx_n_s_alpha_init, __pyx_k_alpha_init, sizeof(__pyx_k_alpha_init), 0, 0, 1, 1},
  {&__pyx_n_s_alpha_max, __pyx_k_alpha_max, sizeof(__pyx_k_alpha_max), 0, 0, 1, 1},
  {&__pyx_n_s_alpha_min, __pyx_k_alpha_min, sizeof(__pyx_k_alpha_min), 0, 0, 1, 1},
  {&__pyx_n_s_alpha_value, __pyx_k_alpha_value, sizeof(__pyx_k_alpha_value), 0, 0, 1, 1},
  {&__pyx_n_s_b, __pyx_k_b, sizeof(__pyx_k_b), 0, 0, 1, 1},
  {&__pyx_n_s_beta_lin, __pyx_k_beta_lin, sizeof(__pyx_k_beta_lin), 0, 0, 1, 1},
  {&__pyx_n_s_beta_max, __pyx_k_beta_max, sizeof(__pyx_k_beta_max), 0, 0, 1, 1},
  {&__pyx_n_s_beta_min, __pyx_k_beta_min, sizeof(__pyx_k_beta_min), 0, 0, 1, 1},
  {&__pyx_n_s_beta_scalar, __pyx_k_beta_scalar, sizeof(__pyx_k_beta_scalar), 0, 0, 1, 1},
  {&__pyx_n_s_beta_value, __pyx_k_beta_value, sizeof(__pyx_k_beta_value), 0, 0, 1, 1},
  {&__pyx_n_s_c1, __pyx_k_c1, sizeof(__pyx_k_c1), 0, 0, 1, 1},
  {&__pyx_n_s_c2, __pyx_k_c2, sizeof(__pyx_k_c2), 0, 0, 1, 1},
  {&__pyx_n_s_cline_in_traceback, __pyx_k_cline_in_traceback, sizeof(__pyx_k_cline_in_traceback), 0, 0, 1, 1},
  {&__pyx_n_s_d, __pyx_k_d, sizeof(__pyx_k_d), 0, 0, 1, 1},
  {&__pyx_n_s_eta, __pyx_k_eta, sizeof(__pyx_k_eta), 0, 0, 1, 1},
  {&__pyx_n_s_evo_steps, __pyx_k_evo_steps, sizeof(__pyx_k_evo_steps), 0, 0, 1, 1},
  {&__pyx_n_s_gamma, __pyx_k_gamma, sizeof(__pyx_k_gamma), 0, 0, 1, 1},
  {&__pyx_n_s_getstate, __pyx_k_getstate, sizeof(__pyx_k_getstate), 0, 0, 1, 1},
  {&__pyx_n_s_hyper, __pyx_k_hyper, sizeof(__pyx_k_hyper), 0, 0, 1, 1},
  {&__pyx_n_s_lam, __pyx_k_lam, sizeof(__pyx_k_lam), 0, 0, 1, 1},
  {&__pyx_n_s_main, __pyx_k_main, sizeof(__pyx_k_main), 0, 0, 1, 1},
  {&__pyx_n_s_name, __pyx_k_name, sizeof(__pyx_k_name), 0, 0, 1, 1},
  {&__pyx_kp_s_no_default___reduce___due_to_non, __pyx_k_no_default___reduce___due_to_non, sizeof(__pyx_k_no_default___reduce___due_to_non), 0, 0, 1, 0},
  {&__pyx_n_s_q, __pyx_k_q, sizeof(__pyx_k_q), 0, 0, 1, 1},
  {&__pyx_n_s_reduce, __pyx_k_reduce, sizeof(__pyx_k_reduce), 0, 0, 1, 1},
  {&__pyx_n_s_reduce_cython, __pyx_k_reduce_cython, sizeof(__pyx_k_reduce_cython), 0, 0, 1, 1},
  {&__pyx_n_s_reduce_ex, __pyx_k_reduce_ex, sizeof(__pyx_k_reduce_ex), 0, 0, 1, 1},
  {&__pyx_n_s_rho, __pyx_k_rho, sizeof(__pyx_k_rho), 0, 0, 1, 1},
  {&__pyx_n_s_seed, __pyx_k_seed, sizeof(__pyx_k_seed), 0, 0, 1, 1},
  {&__pyx_n_s_setstate, __pyx_k_setstate, sizeof(__pyx_k_setstate), 0, 0, 1, 1},
  {&__pyx_n_s_setstate_cython, __pyx_k_setstate_cython, sizeof(__pyx_k_setstate_cython), 0, 0, 1, 1},
  {&__pyx_n_s_sigma_init, __pyx_k_sigma_init, sizeof(__pyx_k_sigma_init), 0, 0, 1, 1},
  {&__pyx_n_s_sigma_max, __pyx_k_sigma_max, sizeof(__pyx_k_sigma_max), 0, 0, 1, 1},
  {&__pyx_n_s_sigma_value, __pyx_k_sigma_value, sizeof(__pyx_k_sigma_value), 0, 0, 1, 1},
  {&__pyx_n_s_test, __pyx_k_test, sizeof(__pyx_k_test), 0, 0, 1, 1},
  {0, 0, 0, 0, 0, 0, 0}
};
static CYTHON_SMALL_CODE int __Pyx_InitCachedBuiltins(void) {
  __pyx_builtin_TypeError = __Pyx_GetBuiltinName(__pyx_n_s_TypeError); if (!__pyx_builtin_TypeError) __PYX_ERR(0, 2, __pyx_L1_error)
  return 0;
  __pyx_L1_error:;
  return -1;
}

static CYTHON_SMALL_CODE int __Pyx_InitCachedConstants(void) {
  __Pyx_RefNannyDeclarations
  __Pyx_RefNannySetupContext("__Pyx_InitCachedConstants", 0);

  /* "(tree fragment)":2
 * def __reduce_cython__(self):
 *     raise TypeError("no default __reduce__ due to non-trivial __cinit__")             # <<<<<<<<<<<<<<
 * def __setstate_cython__(self, __pyx_state):
 *     raise TypeError("no default __reduce__ due to non-trivial __cinit__")
 */
  __pyx_tuple_ = PyTuple_Pack(1, __pyx_kp_s_no_default___reduce___due_to_non); if (unlikely(!__pyx_tuple_)) __PYX_ERR(0, 2, __pyx_L1_error)
  __Pyx_GOTREF(__pyx_tuple_);
  __Pyx_GIVEREF(__pyx_tuple_);

  /* "(tree fragment)":4
 *     raise TypeError("no default __reduce__ due to non-trivial __cinit__")
 * def __setstate_cython__(self, __pyx_state):
 *     raise TypeError("no default __reduce__ due to non-trivial __cinit__")             # <<<<<<<<<<<<<<
 */
  __pyx_tuple__2 = PyTuple_Pack(1, __pyx_kp_s_no_default___reduce___due_to_non); if (unlikely(!__pyx_tuple__2)) __PYX_ERR(0, 4, __pyx_L1_error)
  __Pyx_GOTREF(__pyx_tuple__2);
  __Pyx_GIVEREF(__pyx_tuple__2);
  __Pyx_RefNannyFinishContext();
  return 0;
  __pyx_L1_error:;
  __Pyx_RefNannyFinishContext();
  return -1;
}

static CYTHON_SMALL_CODE int __Pyx_InitGlobals(void) {
  if (__Pyx_InitStrings(__pyx_string_tab) < 0) __PYX_ERR(1, 1, __pyx_L1_error);
  return 0;
  __pyx_L1_error:;
  return -1;
}

static CYTHON_SMALL_CODE int __Pyx_modinit_global_init_code(void); /*proto*/
static CYTHON_SMALL_CODE int __Pyx_modinit_variable_export_code(void); /*proto*/
static CYTHON_SMALL_CODE int __Pyx_modinit_function_export_code(void); /*proto*/
static CYTHON_SMALL_CODE int __Pyx_modinit_type_init_code(void); /*proto*/
static CYTHON_SMALL_CODE int __Pyx_modinit_type_import_code(void); /*proto*/
static CYTHON_SMALL_CODE int __Pyx_modinit_variable_import_code(void); /*proto*/
static CYTHON_SMALL_CODE int __Pyx_modinit_function_import_code(void); /*proto*/

static int __Pyx_modinit_global_init_code(void) {
  __Pyx_RefNannyDeclarations
  __Pyx_RefNannySetupContext("__Pyx_modinit_global_init_code", 0);
  /*--- Global init code ---*/
  __Pyx_RefNannyFinishContext();
  return 0;
}

static int __Pyx_modinit_variable_export_code(void) {
  __Pyx_RefNannyDeclarations
  __Pyx_RefNannySetupContext("__Pyx_modinit_variable_export_code", 0);
  /*--- Variable export code ---*/
  __Pyx_RefNannyFinishContext();
  return 0;
}

static int __Pyx_modinit_function_export_code(void) {
  __Pyx_RefNannyDeclarations
  __Pyx_RefNannySetupContext("__Pyx_modinit_function_export_code", 0);
  /*--- Function export code ---*/
  __Pyx_RefNannyFinishContext();
  return 0;
}

static int __Pyx_modinit_type_init_code(void) {
  __Pyx_RefNannyDeclarations
  int __pyx_lineno = 0;
  const char *__pyx_filename = NULL;
  int __pyx_clineno = 0;
  __Pyx_RefNannySetupContext("__Pyx_modinit_type_init_code", 0);
  /*--- Type init code ---*/
  if (PyType_Ready(&__pyx_type_6runner_PySolver) < 0) __PYX_ERR(1, 4, __pyx_L1_error)
  #if PY_VERSION_HEX < 0x030800B1
  __pyx_type_6runner_PySolver.tp_print = 0;
  #endif
  if ((CYTHON_USE_TYPE_SLOTS && CYTHON_USE_PYTYPE_LOOKUP) && likely(!__pyx_type_6runner_PySolver.tp_dictoffset && __pyx_type_6runner_PySolver.tp_getattro == PyObject_GenericGetAttr)) {
    __pyx_type_6runner_PySolver.tp_getattro = __Pyx_PyObject_GenericGetAttr;
  }
  if (PyObject_SetAttr(__pyx_m, __pyx_n_s_PySolver, (PyObject *)&__pyx_type_6runner_PySolver) < 0) __PYX_ERR(1, 4, __pyx_L1_error)
  if (__Pyx_setup_reduce((PyObject*)&__pyx_type_6runner_PySolver) < 0) __PYX_ERR(1, 4, __pyx_L1_error)
  __pyx_ptype_6runner_PySolver = &__pyx_type_6runner_PySolver;
  __Pyx_RefNannyFinishContext();
  return 0;
  __pyx_L1_error:;
  __Pyx_RefNannyFinishContext();
  return -1;
}

static int __Pyx_modinit_type_import_code(void) {
  __Pyx_RefNannyDeclarations
  __Pyx_RefNannySetupContext("__Pyx_modinit_type_import_code", 0);
  /*--- Type import code ---*/
  __Pyx_RefNannyFinishContext();
  return 0;
}

static int __Pyx_modinit_variable_import_code(void) {
  __Pyx_RefNannyDeclarations
  __Pyx_RefNannySetupContext("__Pyx_modinit_variable_import_code", 0);
  /*--- Variable import code ---*/
  __Pyx_RefNannyFinishContext();
  return 0;
}

static int __Pyx_modinit_function_import_code(void) {
  __Pyx_RefNannyDeclarations
  __Pyx_RefNannySetupContext("__Pyx_modinit_function_import_code", 0);
  /*--- Function import code ---*/
  __Pyx_RefNannyFinishContext();
  return 0;
}


#ifndef CYTHON_NO_PYINIT_EXPORT
#define __Pyx_PyMODINIT_FUNC PyMODINIT_FUNC
#elif PY_MAJOR_VERSION < 3
#ifdef __cplusplus
#define __Pyx_PyMODINIT_FUNC extern "C" void
#else
#define __Pyx_PyMODINIT_FUNC void
#endif
#else
#ifdef __cplusplus
#define __Pyx_PyMODINIT_FUNC extern "C" PyObject *
#else
#define __Pyx_PyMODINIT_FUNC PyObject *
#endif
#endif


#if PY_MAJOR_VERSION < 3
__Pyx_PyMODINIT_FUNC initrunner(void) CYTHON_SMALL_CODE; /*proto*/
__Pyx_PyMODINIT_FUNC initrunner(void)
#else
__Pyx_PyMODINIT_FUNC PyInit_runner(void) CYTHON_SMALL_CODE; /*proto*/
__Pyx_PyMODINIT_FUNC PyInit_runner(void)
#if CYTHON_PEP489_MULTI_PHASE_INIT
{
  return PyModuleDef_Init(&__pyx_moduledef);
}
static CYTHON_SMALL_CODE int __Pyx_check_single_interpreter(void) {
    #if PY_VERSION_HEX >= 0x030700A1
    static PY_INT64_T main_interpreter_id = -1;
    PY_INT64_T current_id = PyInterpreterState_GetID(PyThreadState_Get()->interp);
    if (main_interpreter_id == -1) {
        main_interpreter_id = current_id;
        return (unlikely(current_id == -1)) ? -1 : 0;
    } else if (unlikely(main_interpreter_id != current_id))
    #else
    static PyInterpreterState *main_interpreter = NULL;
    PyInterpreterState *current_interpreter = PyThreadState_Get()->interp;
    if (!main_interpreter) {
        main_interpreter = current_interpreter;
    } else if (unlikely(main_interpreter != current_interpreter))
    #endif
    {
        PyErr_SetString(
            PyExc_ImportError,
            "Interpreter change detected - this module can only be loaded into one interpreter per process.");
        return -1;
    }
    return 0;
}
static CYTHON_SMALL_CODE int __Pyx_copy_spec_to_module(PyObject *spec, PyObject *moddict, const char* from_name, const char* to_name, int allow_none) {
    PyObject *value = PyObject_GetAttrString(spec, from_name);
    int result = 0;
    if (likely(value)) {
        if (allow_none || value != Py_None) {
            result = PyDict_SetItemString(moddict, to_name, value);
        }
        Py_DECREF(value);
    } else if (PyErr_ExceptionMatches(PyExc_AttributeError)) {
        PyErr_Clear();
    } else {
        result = -1;
    }
    return result;
}
static CYTHON_SMALL_CODE PyObject* __pyx_pymod_create(PyObject *spec, CYTHON_UNUSED PyModuleDef *def) {
    PyObject *module = NULL, *moddict, *modname;
    if (__Pyx_check_single_interpreter())
        return NULL;
    if (__pyx_m)
        return __Pyx_NewRef(__pyx_m);
    modname = PyObject_GetAttrString(spec, "name");
    if (unlikely(!modname)) goto bad;
    module = PyModule_NewObject(modname);
    Py_DECREF(modname);
    if (unlikely(!module)) goto bad;
    moddict = PyModule_GetDict(module);
    if (unlikely(!moddict)) goto bad;
    if (unlikely(__Pyx_copy_spec_to_module(spec, moddict, "loader", "__loader__", 1) < 0)) goto bad;
    if (unlikely(__Pyx_copy_spec_to_module(spec, moddict, "origin", "__file__", 1) < 0)) goto bad;
    if (unlikely(__Pyx_copy_spec_to_module(spec, moddict, "parent", "__package__", 1) < 0)) goto bad;
    if (unlikely(__Pyx_copy_spec_to_module(spec, moddict, "submodule_search_locations", "__path__", 0) < 0)) goto bad;
    return module;
bad:
    Py_XDECREF(module);
    return NULL;
}


static CYTHON_SMALL_CODE int __pyx_pymod_exec_runner(PyObject *__pyx_pyinit_module)
#endif
#endif
{
  PyObject *__pyx_t_1 = NULL;
  int __pyx_lineno = 0;
  const char *__pyx_filename = NULL;
  int __pyx_clineno = 0;
  __Pyx_RefNannyDeclarations
  #if CYTHON_PEP489_MULTI_PHASE_INIT
  if (__pyx_m) {
    if (__pyx_m == __pyx_pyinit_module) return 0;
    PyErr_SetString(PyExc_RuntimeError, "Module 'runner' has already been imported. Re-initialisation is not supported.");
    return -1;
  }
  #elif PY_MAJOR_VERSION >= 3
  if (__pyx_m) return __Pyx_NewRef(__pyx_m);
  #endif
  #if CYTHON_REFNANNY
__Pyx_RefNanny = __Pyx_RefNannyImportAPI("refnanny");
if (!__Pyx_RefNanny) {
  PyErr_Clear();
  __Pyx_RefNanny = __Pyx_RefNannyImportAPI("Cython.Runtime.refnanny");
  if (!__Pyx_RefNanny)
      Py_FatalError("failed to import 'refnanny' module");
}
#endif
  __Pyx_RefNannySetupContext("__Pyx_PyMODINIT_FUNC PyInit_runner(void)", 0);
  if (__Pyx_check_binary_version() < 0) __PYX_ERR(1, 1, __pyx_L1_error)
  #ifdef __Pxy_PyFrame_Initialize_Offsets
  __Pxy_PyFrame_Initialize_Offsets();
  #endif
  __pyx_empty_tuple = PyTuple_New(0); if (unlikely(!__pyx_empty_tuple)) __PYX_ERR(1, 1, __pyx_L1_error)
  __pyx_empty_bytes = PyBytes_FromStringAndSize("", 0); if (unlikely(!__pyx_empty_bytes)) __PYX_ERR(1, 1, __pyx_L1_error)
  __pyx_empty_unicode = PyUnicode_FromStringAndSize("", 0); if (unlikely(!__pyx_empty_unicode)) __PYX_ERR(1, 1, __pyx_L1_error)
  #ifdef __Pyx_CyFunction_USED
  if (__pyx_CyFunction_init() < 0) __PYX_ERR(1, 1, __pyx_L1_error)
  #endif
  #ifdef __Pyx_FusedFunction_USED
  if (__pyx_FusedFunction_init() < 0) __PYX_ERR(1, 1, __pyx_L1_error)
  #endif
  #ifdef __Pyx_Coroutine_USED
  if (__pyx_Coroutine_init() < 0) __PYX_ERR(1, 1, __pyx_L1_error)
  #endif
  #ifdef __Pyx_Generator_USED
  if (__pyx_Generator_init() < 0) __PYX_ERR(1, 1, __pyx_L1_error)
  #endif
  #ifdef __Pyx_AsyncGen_USED
  if (__pyx_AsyncGen_init() < 0) __PYX_ERR(1, 1, __pyx_L1_error)
  #endif
  #ifdef __Pyx_StopAsyncIteration_USED
  if (__pyx_StopAsyncIteration_init() < 0) __PYX_ERR(1, 1, __pyx_L1_error)
  #endif
  /*--- Library function declarations ---*/
  /*--- Threads initialization code ---*/
  #if defined(WITH_THREAD) && PY_VERSION_HEX < 0x030700F0 && defined(__PYX_FORCE_INIT_THREADS) && __PYX_FORCE_INIT_THREADS
  PyEval_InitThreads();
  #endif
  /*--- Module creation code ---*/
  #if CYTHON_PEP489_MULTI_PHASE_INIT
  __pyx_m = __pyx_pyinit_module;
  Py_INCREF(__pyx_m);
  #else
  #if PY_MAJOR_VERSION < 3
  __pyx_m = Py_InitModule4("runner", __pyx_methods, 0, 0, PYTHON_API_VERSION); Py_XINCREF(__pyx_m);
  #else
  __pyx_m = PyModule_Create(&__pyx_moduledef);
  #endif
  if (unlikely(!__pyx_m)) __PYX_ERR(1, 1, __pyx_L1_error)
  #endif
  __pyx_d = PyModule_GetDict(__pyx_m); if (unlikely(!__pyx_d)) __PYX_ERR(1, 1, __pyx_L1_error)
  Py_INCREF(__pyx_d);
  __pyx_b = PyImport_AddModule(__Pyx_BUILTIN_MODULE_NAME); if (unlikely(!__pyx_b)) __PYX_ERR(1, 1, __pyx_L1_error)
  Py_INCREF(__pyx_b);
  __pyx_cython_runtime = PyImport_AddModule((char *) "cython_runtime"); if (unlikely(!__pyx_cython_runtime)) __PYX_ERR(1, 1, __pyx_L1_error)
  Py_INCREF(__pyx_cython_runtime);
  if (PyObject_SetAttrString(__pyx_m, "__builtins__", __pyx_b) < 0) __PYX_ERR(1, 1, __pyx_L1_error);
  /*--- Initialize various global constants etc. ---*/
  if (__Pyx_InitGlobals() < 0) __PYX_ERR(1, 1, __pyx_L1_error)
  #if PY_MAJOR_VERSION < 3 && (__PYX_DEFAULT_STRING_ENCODING_IS_ASCII || __PYX_DEFAULT_STRING_ENCODING_IS_DEFAULT)
  if (__Pyx_init_sys_getdefaultencoding_params() < 0) __PYX_ERR(1, 1, __pyx_L1_error)
  #endif
  if (__pyx_module_is_main_runner) {
    if (PyObject_SetAttr(__pyx_m, __pyx_n_s_name, __pyx_n_s_main) < 0) __PYX_ERR(1, 1, __pyx_L1_error)
  }
  #if PY_MAJOR_VERSION >= 3
  {
    PyObject *modules = PyImport_GetModuleDict(); if (unlikely(!modules)) __PYX_ERR(1, 1, __pyx_L1_error)
    if (!PyDict_GetItemString(modules, "runner")) {
      if (unlikely(PyDict_SetItemString(modules, "runner", __pyx_m) < 0)) __PYX_ERR(1, 1, __pyx_L1_error)
    }
  }
  #endif
  /*--- Builtin init code ---*/
  if (__Pyx_InitCachedBuiltins() < 0) __PYX_ERR(1, 1, __pyx_L1_error)
  /*--- Constants init code ---*/
  if (__Pyx_InitCachedConstants() < 0) __PYX_ERR(1, 1, __pyx_L1_error)
  /*--- Global type/function init code ---*/
  (void)__Pyx_modinit_global_init_code();
  (void)__Pyx_modinit_variable_export_code();
  (void)__Pyx_modinit_function_export_code();
  if (unlikely(__Pyx_modinit_type_init_code() < 0)) __PYX_ERR(1, 1, __pyx_L1_error)
  (void)__Pyx_modinit_type_import_code();
  (void)__Pyx_modinit_variable_import_code();
  (void)__Pyx_modinit_function_import_code();
  /*--- Execution code ---*/
  #if defined(__Pyx_Generator_USED) || defined(__Pyx_Coroutine_USED)
  if (__Pyx_patch_abc() < 0) __PYX_ERR(1, 1, __pyx_L1_error)
  #endif

  /* "runner.pyx":1
 * # distutils: language = c++             # <<<<<<<<<<<<<<
 * from solvers cimport Quick_solver
 * 
 */
  __pyx_t_1 = __Pyx_PyDict_NewPresized(0); if (unlikely(!__pyx_t_1)) __PYX_ERR(1, 1, __pyx_L1_error)
  __Pyx_GOTREF(__pyx_t_1);
  if (PyDict_SetItem(__pyx_d, __pyx_n_s_test, __pyx_t_1) < 0) __PYX_ERR(1, 1, __pyx_L1_error)
  __Pyx_DECREF(__pyx_t_1); __pyx_t_1 = 0;

  /*--- Wrapped vars code ---*/

  goto __pyx_L0;
  __pyx_L1_error:;
  __Pyx_XDECREF(__pyx_t_1);
  if (__pyx_m) {
    if (__pyx_d) {
      __Pyx_AddTraceback("init runner", __pyx_clineno, __pyx_lineno, __pyx_filename);
    }
    Py_CLEAR(__pyx_m);
  } else if (!PyErr_Occurred()) {
    PyErr_SetString(PyExc_ImportError, "init runner");
  }
  __pyx_L0:;
  __Pyx_RefNannyFinishContext();
  #if CYTHON_PEP489_MULTI_PHASE_INIT
  return (__pyx_m != NULL) ? 0 : -1;
  #elif PY_MAJOR_VERSION >= 3
  return __pyx_m;
  #else
  return;
  #endif
}

/* --- Runtime support code --- */
/* Refnanny */
#if CYTHON_REFNANNY
static __Pyx_RefNannyAPIStruct *__Pyx_RefNannyImportAPI(const char *modname) {
    PyObject *m = NULL, *p = NULL;
    void *r = NULL;
    m = PyImport_ImportModule(modname);
    if (!m) goto end;
    p = PyObject_GetAttrString(m, "RefNannyAPI");
    if (!p) goto end;
    r = PyLong_AsVoidPtr(p);
end:
    Py_XDECREF(p);
    Py_XDECREF(m);
    return (__Pyx_RefNannyAPIStruct *)r;
}
#endif

/* PyObjectGetAttrStr */
#if CYTHON_USE_TYPE_SLOTS
static CYTHON_INLINE PyObject* __Pyx_PyObject_GetAttrStr(PyObject* obj, PyObject* attr_name) {
    PyTypeObject* tp = Py_TYPE(obj);
    if (likely(tp->tp_getattro))
        return tp->tp_getattro(obj, attr_name);
#if PY_MAJOR_VERSION < 3
    if (likely(tp->tp_getattr))
        return tp->tp_getattr(obj, PyString_AS_STRING(attr_name));
#endif
    return PyObject_GetAttr(obj, attr_name);
}
#endif

/* GetBuiltinName */
static PyObject *__Pyx_GetBuiltinName(PyObject *name) {
    PyObject* result = __Pyx_PyObject_GetAttrStr(__pyx_b, name);
    if (unlikely(!result)) {
        PyErr_Format(PyExc_NameError,
#if PY_MAJOR_VERSION >= 3
            "name '%U' is not defined", name);
#else
            "name '%.200s' is not defined", PyString_AS_STRING(name));
#endif
    }
    return result;
}

/* RaiseArgTupleInvalid */
static void __Pyx_RaiseArgtupleInvalid(
    const char* func_name,
    int exact,
    Py_ssize_t num_min,
    Py_ssize_t num_max,
    Py_ssize_t num_found)
{
    Py_ssize_t num_expected;
    const char *more_or_less;
    if (num_found < num_min) {
        num_expected = num_min;
        more_or_less = "at least";
    } else {
        num_expected = num_max;
        more_or_less = "at most";
    }
    if (exact) {
        more_or_less = "exactly";
    }
    PyErr_Format(PyExc_TypeError,
                 "%.200s() takes %.8s %" CYTHON_FORMAT_SSIZE_T "d positional argument%.1s (%" CYTHON_FORMAT_SSIZE_T "d given)",
                 func_name, more_or_less, num_expected,
                 (num_expected == 1) ? "" : "s", num_found);
}

/* KeywordStringCheck */
static int __Pyx_CheckKeywordStrings(
    PyObject *kwdict,
    const char* function_name,
    int kw_allowed)
{
    PyObject* key = 0;
    Py_ssize_t pos = 0;
#if CYTHON_COMPILING_IN_PYPY
    if (!kw_allowed && PyDict_Next(kwdict, &pos, &key, 0))
        goto invalid_keyword;
    return 1;
#else
    while (PyDict_Next(kwdict, &pos, &key, 0)) {
        #if PY_MAJOR_VERSION < 3
        if (unlikely(!PyString_Check(key)))
        #endif
            if (unlikely(!PyUnicode_Check(key)))
                goto invalid_keyword_type;
    }
    if ((!kw_allowed) && unlikely(key))
        goto invalid_keyword;
    return 1;
invalid_keyword_type:
    PyErr_Format(PyExc_TypeError,
        "%.200s() keywords must be strings", function_name);
    return 0;
#endif
invalid_keyword:
    PyErr_Format(PyExc_TypeError,
    #if PY_MAJOR_VERSION < 3
        "%.200s() got an unexpected keyword argument '%.200s'",
        function_name, PyString_AsString(key));
    #else
        "%s() got an unexpected keyword argument '%U'",
        function_name, key);
    #endif
    return 0;
}

/* RaiseDoubleKeywords */
static void __Pyx_RaiseDoubleKeywordsError(
    const char* func_name,
    PyObject* kw_name)
{
    PyErr_Format(PyExc_TypeError,
        #if PY_MAJOR_VERSION >= 3
        "%s() got multiple values for keyword argument '%U'", func_name, kw_name);
        #else
        "%s() got multiple values for keyword argument '%s'", func_name,
        PyString_AsString(kw_name));
        #endif
}

/* ParseKeywords */
static int __Pyx_ParseOptionalKeywords(
    PyObject *kwds,
    PyObject **argnames[],
    PyObject *kwds2,
    PyObject *values[],
    Py_ssize_t num_pos_args,
    const char* function_name)
{
    PyObject *key = 0, *value = 0;
    Py_ssize_t pos = 0;
    PyObject*** name;
    PyObject*** first_kw_arg = argnames + num_pos_args;
    while (PyDict_Next(kwds, &pos, &key, &value)) {
        name = first_kw_arg;
        while (*name && (**name != key)) name++;
        if (*name) {
            values[name-argnames] = value;
            continue;
        }
        name = first_kw_arg;
        #if PY_MAJOR_VERSION < 3
        if (likely(PyString_Check(key))) {
            while (*name) {
                if ((CYTHON_COMPILING_IN_PYPY || PyString_GET_SIZE(**name) == PyString_GET_SIZE(key))
                        && _PyString_Eq(**name, key)) {
                    values[name-argnames] = value;
                    break;
                }
                name++;
            }
            if (*name) continue;
            else {
                PyObject*** argname = argnames;
                while (argname != first_kw_arg) {
                    if ((**argname == key) || (
                            (CYTHON_COMPILING_IN_PYPY || PyString_GET_SIZE(**argname) == PyString_GET_SIZE(key))
                             && _PyString_Eq(**argname, key))) {
                        goto arg_passed_twice;
                    }
                    argname++;
                }
            }
        } else
        #endif
        if (likely(PyUnicode_Check(key))) {
            while (*name) {
                int cmp = (**name == key) ? 0 :
                #if !CYTHON_COMPILING_IN_PYPY && PY_MAJOR_VERSION >= 3
                    (__Pyx_PyUnicode_GET_LENGTH(**name) != __Pyx_PyUnicode_GET_LENGTH(key)) ? 1 :
                #endif
                    PyUnicode_Compare(**name, key);
                if (cmp < 0 && unlikely(PyErr_Occurred())) goto bad;
                if (cmp == 0) {
                    values[name-argnames] = value;
                    break;
                }
                name++;
            }
            if (*name) continue;
            else {
                PyObject*** argname = argnames;
                while (argname != first_kw_arg) {
                    int cmp = (**argname == key) ? 0 :
                    #if !CYTHON_COMPILING_IN_PYPY && PY_MAJOR_VERSION >= 3
                        (__Pyx_PyUnicode_GET_LENGTH(**argname) != __Pyx_PyUnicode_GET_LENGTH(key)) ? 1 :
                    #endif
                        PyUnicode_Compare(**argname, key);
                    if (cmp < 0 && unlikely(PyErr_Occurred())) goto bad;
                    if (cmp == 0) goto arg_passed_twice;
                    argname++;
                }
            }
        } else
            goto invalid_keyword_type;
        if (kwds2) {
            if (unlikely(PyDict_SetItem(kwds2, key, value))) goto bad;
        } else {
            goto invalid_keyword;
        }
    }
    return 0;
arg_passed_twice:
    __Pyx_RaiseDoubleKeywordsError(function_name, key);
    goto bad;
invalid_keyword_type:
    PyErr_Format(PyExc_TypeError,
        "%.200s() keywords must be strings", function_name);
    goto bad;
invalid_keyword:
    PyErr_Format(PyExc_TypeError,
    #if PY_MAJOR_VERSION < 3
        "%.200s() got an unexpected keyword argument '%.200s'",
        function_name, PyString_AsString(key));
    #else
        "%s() got an unexpected keyword argument '%U'",
        function_name, key);
    #endif
bad:
    return -1;
}

/* PyObjectCall */
#if CYTHON_COMPILING_IN_CPYTHON
static CYTHON_INLINE PyObject* __Pyx_PyObject_Call(PyObject *func, PyObject *arg, PyObject *kw) {
    PyObject *result;
    ternaryfunc call = Py_TYPE(func)->tp_call;
    if (unlikely(!call))
        return PyObject_Call(func, arg, kw);
    if (unlikely(Py_EnterRecursiveCall((char*)" while calling a Python object")))
        return NULL;
    result = (*call)(func, arg, kw);
    Py_LeaveRecursiveCall();
    if (unlikely(!result) && unlikely(!PyErr_Occurred())) {
        PyErr_SetString(
            PyExc_SystemError,
            "NULL result without error in PyObject_Call");
    }
    return result;
}
#endif

/* PyErrFetchRestore */
#if CYTHON_FAST_THREAD_STATE
static CYTHON_INLINE void __Pyx_ErrRestoreInState(PyThreadState *tstate, PyObject *type, PyObject *value, PyObject *tb) {
    PyObject *tmp_type, *tmp_value, *tmp_tb;
    tmp_type = tstate->curexc_type;
    tmp_value = tstate->curexc_value;
    tmp_tb = tstate->curexc_traceback;
    tstate->curexc_type = type;
    tstate->curexc_value = value;
    tstate->curexc_traceback = tb;
    Py_XDECREF(tmp_type);
    Py_XDECREF(tmp_value);
    Py_XDECREF(tmp_tb);
}
static CYTHON_INLINE void __Pyx_ErrFetchInState(PyThreadState *tstate, PyObject **type, PyObject **value, PyObject **tb) {
    *type = tstate->curexc_type;
    *value = tstate->curexc_value;
    *tb = tstate->curexc_traceback;
    tstate->curexc_type = 0;
    tstate->curexc_value = 0;
    tstate->curexc_traceback = 0;
}
#endif

/* RaiseException */
#if PY_MAJOR_VERSION < 3
static void __Pyx_Raise(PyObject *type, PyObject *value, PyObject *tb,
                        CYTHON_UNUSED PyObject *cause) {
    __Pyx_PyThreadState_declare
    Py_XINCREF(type);
    if (!value || value == Py_None)
        value = NULL;
    else
        Py_INCREF(value);
    if (!tb || tb == Py_None)
        tb = NULL;
    else {
        Py_INCREF(tb);
        if (!PyTraceBack_Check(tb)) {
            PyErr_SetString(PyExc_TypeError,
                "raise: arg 3 must be a traceback or None");
            goto raise_error;
        }
    }
    if (PyType_Check(type)) {
#if CYTHON_COMPILING_IN_PYPY
        if (!value) {
            Py_INCREF(Py_None);
            value = Py_None;
        }
#endif
        PyErr_NormalizeException(&type, &value, &tb);
    } else {
        if (value) {
            PyErr_SetString(PyExc_TypeError,
                "instance exception may not have a separate value");
            goto raise_error;
        }
        value = type;
        type = (PyObject*) Py_TYPE(type);
        Py_INCREF(type);
        if (!PyType_IsSubtype((PyTypeObject *)type, (PyTypeObject *)PyExc_BaseException)) {
            PyErr_SetString(PyExc_TypeError,
                "raise: exception class must be a subclass of BaseException");
            goto raise_error;
        }
    }
    __Pyx_PyThreadState_assign
    __Pyx_ErrRestore(type, value, tb);
    return;
raise_error:
    Py_XDECREF(value);
    Py_XDECREF(type);
    Py_XDECREF(tb);
    return;
}
#else
static void __Pyx_Raise(PyObject *type, PyObject *value, PyObject *tb, PyObject *cause) {
    PyObject* owned_instance = NULL;
    if (tb == Py_None) {
        tb = 0;
    } else if (tb && !PyTraceBack_Check(tb)) {
        PyErr_SetString(PyExc_TypeError,
            "raise: arg 3 must be a traceback or None");
        goto bad;
    }
    if (value == Py_None)
        value = 0;
    if (PyExceptionInstance_Check(type)) {
        if (value) {
            PyErr_SetString(PyExc_TypeError,
                "instance exception may not have a separate value");
            goto bad;
        }
        value = type;
        type = (PyObject*) Py_TYPE(value);
    } else if (PyExceptionClass_Check(type)) {
        PyObject *instance_class = NULL;
        if (value && PyExceptionInstance_Check(value)) {
            instance_class = (PyObject*) Py_TYPE(value);
            if (instance_class != type) {
                int is_subclass = PyObject_IsSubclass(instance_class, type);
                if (!is_subclass) {
                    instance_class = NULL;
                } else if (unlikely(is_subclass == -1)) {
                    goto bad;
                } else {
                    type = instance_class;
                }
            }
        }
        if (!instance_class) {
            PyObject *args;
            if (!value)
                args = PyTuple_New(0);
            else if (PyTuple_Check(value)) {
                Py_INCREF(value);
                args = value;
            } else
                args = PyTuple_Pack(1, value);
            if (!args)
                goto bad;
            owned_instance = PyObject_Call(type, args, NULL);
            Py_DECREF(args);
            if (!owned_instance)
                goto bad;
            value = owned_instance;
            if (!PyExceptionInstance_Check(value)) {
                PyErr_Format(PyExc_TypeError,
                             "calling %R should have returned an instance of "
                             "BaseException, not %R",
                             type, Py_TYPE(value));
                goto bad;
            }
        }
    } else {
        PyErr_SetString(PyExc_TypeError,
            "raise: exception class must be a subclass of BaseException");
        goto bad;
    }
    if (cause) {
        PyObject *fixed_cause;
        if (cause == Py_None) {
            fixed_cause = NULL;
        } else if (PyExceptionClass_Check(cause)) {
            fixed_cause = PyObject_CallObject(cause, NULL);
            if (fixed_cause == NULL)
                goto bad;
        } else if (PyExceptionInstance_Check(cause)) {
            fixed_cause = cause;
            Py_INCREF(fixed_cause);
        } else {
            PyErr_SetString(PyExc_TypeError,
                            "exception causes must derive from "
                            "BaseException");
            goto bad;
        }
        PyException_SetCause(value, fixed_cause);
    }
    PyErr_SetObject(type, value);
    if (tb) {
#if CYTHON_COMPILING_IN_PYPY
        PyObject *tmp_type, *tmp_value, *tmp_tb;
        PyErr_Fetch(&tmp_type, &tmp_value, &tmp_tb);
        Py_INCREF(tb);
        PyErr_Restore(tmp_type, tmp_value, tb);
        Py_XDECREF(tmp_tb);
#else
        PyThreadState *tstate = __Pyx_PyThreadState_Current;
        PyObject* tmp_tb = tstate->curexc_traceback;
        if (tb != tmp_tb) {
            Py_INCREF(tb);
            tstate->curexc_traceback = tb;
            Py_XDECREF(tmp_tb);
        }
#endif
    }
bad:
    Py_XDECREF(owned_instance);
    return;
}
#endif

/* PyObject_GenericGetAttrNoDict */
#if CYTHON_USE_TYPE_SLOTS && CYTHON_USE_PYTYPE_LOOKUP && PY_VERSION_HEX < 0x03070000
static PyObject *__Pyx_RaiseGenericGetAttributeError(PyTypeObject *tp, PyObject *attr_name) {
    PyErr_Format(PyExc_AttributeError,
#if PY_MAJOR_VERSION >= 3
                 "'%.50s' object has no attribute '%U'",
                 tp->tp_name, attr_name);
#else
                 "'%.50s' object has no attribute '%.400s'",
                 tp->tp_name, PyString_AS_STRING(attr_name));
#endif
    return NULL;
}
static CYTHON_INLINE PyObject* __Pyx_PyObject_GenericGetAttrNoDict(PyObject* obj, PyObject* attr_name) {
    PyObject *descr;
    PyTypeObject *tp = Py_TYPE(obj);
    if (unlikely(!PyString_Check(attr_name))) {
        return PyObject_GenericGetAttr(obj, attr_name);
    }
    assert(!tp->tp_dictoffset);
    descr = _PyType_Lookup(tp, attr_name);
    if (unlikely(!descr)) {
        return __Pyx_RaiseGenericGetAttributeError(tp, attr_name);
    }
    Py_INCREF(descr);
    #if PY_MAJOR_VERSION < 3
    if (likely(PyType_HasFeature(Py_TYPE(descr), Py_TPFLAGS_HAVE_CLASS)))
    #endif
    {
        descrgetfunc f = Py_TYPE(descr)->tp_descr_get;
        if (unlikely(f)) {
            PyObject *res = f(descr, obj, (PyObject *)tp);
            Py_DECREF(descr);
            return res;
        }
    }
    return descr;
}
#endif

/* PyObject_GenericGetAttr */
#if CYTHON_USE_TYPE_SLOTS && CYTHON_USE_PYTYPE_LOOKUP && PY_VERSION_HEX < 0x03070000
static PyObject* __Pyx_PyObject_GenericGetAttr(PyObject* obj, PyObject* attr_name) {
    if (unlikely(Py_TYPE(obj)->tp_dictoffset)) {
        return PyObject_GenericGetAttr(obj, attr_name);
    }
    return __Pyx_PyObject_GenericGetAttrNoDict(obj, attr_name);
}
#endif

/* PyErrExceptionMatches */
#if CYTHON_FAST_THREAD_STATE
static int __Pyx_PyErr_ExceptionMatchesTuple(PyObject *exc_type, PyObject *tuple) {
    Py_ssize_t i, n;
    n = PyTuple_GET_SIZE(tuple);
#if PY_MAJOR_VERSION >= 3
    for (i=0; i<n; i++) {
        if (exc_type == PyTuple_GET_ITEM(tuple, i)) return 1;
    }
#endif
    for (i=0; i<n; i++) {
        if (__Pyx_PyErr_GivenExceptionMatches(exc_type, PyTuple_GET_ITEM(tuple, i))) return 1;
    }
    return 0;
}
static CYTHON_INLINE int __Pyx_PyErr_ExceptionMatchesInState(PyThreadState* tstate, PyObject* err) {
    PyObject *exc_type = tstate->curexc_type;
    if (exc_type == err) return 1;
    if (unlikely(!exc_type)) return 0;
    if (unlikely(PyTuple_Check(err)))
        return __Pyx_PyErr_ExceptionMatchesTuple(exc_type, err);
    return __Pyx_PyErr_GivenExceptionMatches(exc_type, err);
}
#endif

/* PyObjectGetAttrStrNoError */
static void __Pyx_PyObject_GetAttrStr_ClearAttributeError(void) {
    __Pyx_PyThreadState_declare
    __Pyx_PyThreadState_assign
    if (likely(__Pyx_PyErr_ExceptionMatches(PyExc_AttributeError)))
        __Pyx_PyErr_Clear();
}
static CYTHON_INLINE PyObject* __Pyx_PyObject_GetAttrStrNoError(PyObject* obj, PyObject* attr_name) {
    PyObject *result;
#if CYTHON_COMPILING_IN_CPYTHON && CYTHON_USE_TYPE_SLOTS && PY_VERSION_HEX >= 0x030700B1
    PyTypeObject* tp = Py_TYPE(obj);
    if (likely(tp->tp_getattro == PyObject_GenericGetAttr)) {
        return _PyObject_GenericGetAttrWithDict(obj, attr_name, NULL, 1);
    }
#endif
    result = __Pyx_PyObject_GetAttrStr(obj, attr_name);
    if (unlikely(!result)) {
        __Pyx_PyObject_GetAttrStr_ClearAttributeError();
    }
    return result;
}

/* SetupReduce */
static int __Pyx_setup_reduce_is_named(PyObject* meth, PyObject* name) {
  int ret;
  PyObject *name_attr;
  name_attr = __Pyx_PyObject_GetAttrStr(meth, __pyx_n_s_name);
  if (likely(name_attr)) {
      ret = PyObject_RichCompareBool(name_attr, name, Py_EQ);
  } else {
      ret = -1;
  }
  if (unlikely(ret < 0)) {
      PyErr_Clear();
      ret = 0;
  }
  Py_XDECREF(name_attr);
  return ret;
}
static int __Pyx_setup_reduce(PyObject* type_obj) {
    int ret = 0;
    PyObject *object_reduce = NULL;
    PyObject *object_getstate = NULL;
    PyObject *object_reduce_ex = NULL;
    PyObject *reduce = NULL;
    PyObject *reduce_ex = NULL;
    PyObject *reduce_cython = NULL;
    PyObject *setstate = NULL;
    PyObject *setstate_cython = NULL;
    PyObject *getstate = NULL;
#if CYTHON_USE_PYTYPE_LOOKUP
    getstate = _PyType_Lookup((PyTypeObject*)type_obj, __pyx_n_s_getstate);
#else
    getstate = __Pyx_PyObject_GetAttrStrNoError(type_obj, __pyx_n_s_getstate);
    if (!getstate && PyErr_Occurred()) {
        goto __PYX_BAD;
    }
#endif
    if (getstate) {
#if CYTHON_USE_PYTYPE_LOOKUP
        object_getstate = _PyType_Lookup(&PyBaseObject_Type, __pyx_n_s_getstate);
#else
        object_getstate = __Pyx_PyObject_GetAttrStrNoError((PyObject*)&PyBaseObject_Type, __pyx_n_s_getstate);
        if (!object_getstate && PyErr_Occurred()) {
            goto __PYX_BAD;
        }
#endif
        if (object_getstate != getstate) {
            goto __PYX_GOOD;
        }
    }
#if CYTHON_USE_PYTYPE_LOOKUP
    object_reduce_ex = _PyType_Lookup(&PyBaseObject_Type, __pyx_n_s_reduce_ex); if (!object_reduce_ex) goto __PYX_BAD;
#else
    object_reduce_ex = __Pyx_PyObject_GetAttrStr((PyObject*)&PyBaseObject_Type, __pyx_n_s_reduce_ex); if (!object_reduce_ex) goto __PYX_BAD;
#endif
    reduce_ex = __Pyx_PyObject_GetAttrStr(type_obj, __pyx_n_s_reduce_ex); if (unlikely(!reduce_ex)) goto __PYX_BAD;
    if (reduce_ex == object_reduce_ex) {
#if CYTHON_USE_PYTYPE_LOOKUP
        object_reduce = _PyType_Lookup(&PyBaseObject_Type, __pyx_n_s_reduce); if (!object_reduce) goto __PYX_BAD;
#else
        object_reduce = __Pyx_PyObject_GetAttrStr((PyObject*)&PyBaseObject_Type, __pyx_n_s_reduce); if (!object_reduce) goto __PYX_BAD;
#endif
        reduce = __Pyx_PyObject_GetAttrStr(type_obj, __pyx_n_s_reduce); if (unlikely(!reduce)) goto __PYX_BAD;
        if (reduce == object_reduce || __Pyx_setup_reduce_is_named(reduce, __pyx_n_s_reduce_cython)) {
            reduce_cython = __Pyx_PyObject_GetAttrStrNoError(type_obj, __pyx_n_s_reduce_cython);
            if (likely(reduce_cython)) {
                ret = PyDict_SetItem(((PyTypeObject*)type_obj)->tp_dict, __pyx_n_s_reduce, reduce_cython); if (unlikely(ret < 0)) goto __PYX_BAD;
                ret = PyDict_DelItem(((PyTypeObject*)type_obj)->tp_dict, __pyx_n_s_reduce_cython); if (unlikely(ret < 0)) goto __PYX_BAD;
            } else if (reduce == object_reduce || PyErr_Occurred()) {
                goto __PYX_BAD;
            }
            setstate = __Pyx_PyObject_GetAttrStr(type_obj, __pyx_n_s_setstate);
            if (!setstate) PyErr_Clear();
            if (!setstate || __Pyx_setup_reduce_is_named(setstate, __pyx_n_s_setstate_cython)) {
                setstate_cython = __Pyx_PyObject_GetAttrStrNoError(type_obj, __pyx_n_s_setstate_cython);
                if (likely(setstate_cython)) {
                    ret = PyDict_SetItem(((PyTypeObject*)type_obj)->tp_dict, __pyx_n_s_setstate, setstate_cython); if (unlikely(ret < 0)) goto __PYX_BAD;
                    ret = PyDict_DelItem(((PyTypeObject*)type_obj)->tp_dict, __pyx_n_s_setstate_cython); if (unlikely(ret < 0)) goto __PYX_BAD;
                } else if (!setstate || PyErr_Occurred()) {
                    goto __PYX_BAD;
                }
            }
            PyType_Modified((PyTypeObject*)type_obj);
        }
    }
    goto __PYX_GOOD;
__PYX_BAD:
    if (!PyErr_Occurred())
        PyErr_Format(PyExc_RuntimeError, "Unable to initialize pickling for %s", ((PyTypeObject*)type_obj)->tp_name);
    ret = -1;
__PYX_GOOD:
#if !CYTHON_USE_PYTYPE_LOOKUP
    Py_XDECREF(object_reduce);
    Py_XDECREF(object_reduce_ex);
    Py_XDECREF(object_getstate);
    Py_XDECREF(getstate);
#endif
    Py_XDECREF(reduce);
    Py_XDECREF(reduce_ex);
    Py_XDECREF(reduce_cython);
    Py_XDECREF(setstate);
    Py_XDECREF(setstate_cython);
    return ret;
}

/* PyDictVersioning */
#if CYTHON_USE_DICT_VERSIONS && CYTHON_USE_TYPE_SLOTS
static CYTHON_INLINE PY_UINT64_T __Pyx_get_tp_dict_version(PyObject *obj) {
    PyObject *dict = Py_TYPE(obj)->tp_dict;
    return likely(dict) ? __PYX_GET_DICT_VERSION(dict) : 0;
}
static CYTHON_INLINE PY_UINT64_T __Pyx_get_object_dict_version(PyObject *obj) {
    PyObject **dictptr = NULL;
    Py_ssize_t offset = Py_TYPE(obj)->tp_dictoffset;
    if (offset) {
#if CYTHON_COMPILING_IN_CPYTHON
        dictptr = (likely(offset > 0)) ? (PyObject **) ((char *)obj + offset) : _PyObject_GetDictPtr(obj);
#else
        dictptr = _PyObject_GetDictPtr(obj);
#endif
    }
    return (dictptr && *dictptr) ? __PYX_GET_DICT_VERSION(*dictptr) : 0;
}
static CYTHON_INLINE int __Pyx_object_dict_version_matches(PyObject* obj, PY_UINT64_T tp_dict_version, PY_UINT64_T obj_dict_version) {
    PyObject *dict = Py_TYPE(obj)->tp_dict;
    if (unlikely(!dict) || unlikely(tp_dict_version != __PYX_GET_DICT_VERSION(dict)))
        return 0;
    return obj_dict_version == __Pyx_get_object_dict_version(obj);
}
#endif

/* CLineInTraceback */
#ifndef CYTHON_CLINE_IN_TRACEBACK
static int __Pyx_CLineForTraceback(CYTHON_NCP_UNUSED PyThreadState *tstate, int c_line) {
    PyObject *use_cline;
    PyObject *ptype, *pvalue, *ptraceback;
#if CYTHON_COMPILING_IN_CPYTHON
    PyObject **cython_runtime_dict;
#endif
    if (unlikely(!__pyx_cython_runtime)) {
        return c_line;
    }
    __Pyx_ErrFetchInState(tstate, &ptype, &pvalue, &ptraceback);
#if CYTHON_COMPILING_IN_CPYTHON
    cython_runtime_dict = _PyObject_GetDictPtr(__pyx_cython_runtime);
    if (likely(cython_runtime_dict)) {
        __PYX_PY_DICT_LOOKUP_IF_MODIFIED(
            use_cline, *cython_runtime_dict,
            __Pyx_PyDict_GetItemStr(*cython_runtime_dict, __pyx_n_s_cline_in_traceback))
    } else
#endif
    {
      PyObject *use_cline_obj = __Pyx_PyObject_GetAttrStr(__pyx_cython_runtime, __pyx_n_s_cline_in_traceback);
      if (use_cline_obj) {
        use_cline = PyObject_Not(use_cline_obj) ? Py_False : Py_True;
        Py_DECREF(use_cline_obj);
      } else {
        PyErr_Clear();
        use_cline = NULL;
      }
    }
    if (!use_cline) {
        c_line = 0;
        (void) PyObject_SetAttr(__pyx_cython_runtime, __pyx_n_s_cline_in_traceback, Py_False);
    }
    else if (use_cline == Py_False || (use_cline != Py_True && PyObject_Not(use_cline) != 0)) {
        c_line = 0;
    }
    __Pyx_ErrRestoreInState(tstate, ptype, pvalue, ptraceback);
    return c_line;
}
#endif

/* CodeObjectCache */
static int __pyx_bisect_code_objects(__Pyx_CodeObjectCacheEntry* entries, int count, int code_line) {
    int start = 0, mid = 0, end = count - 1;
    if (end >= 0 && code_line > entries[end].code_line) {
        return count;
    }
    while (start < end) {
        mid = start + (end - start) / 2;
        if (code_line < entries[mid].code_line) {
            end = mid;
        } else if (code_line > entries[mid].code_line) {
             start = mid + 1;
        } else {
            return mid;
        }
    }
    if (code_line <= entries[mid].code_line) {
        return mid;
    } else {
        return mid + 1;
    }
}
static PyCodeObject *__pyx_find_code_object(int code_line) {
    PyCodeObject* code_object;
    int pos;
    if (unlikely(!code_line) || unlikely(!__pyx_code_cache.entries)) {
        return NULL;
    }
    pos = __pyx_bisect_code_objects(__pyx_code_cache.entries, __pyx_code_cache.count, code_line);
    if (unlikely(pos >= __pyx_code_cache.count) || unlikely(__pyx_code_cache.entries[pos].code_line != code_line)) {
        return NULL;
    }
    code_object = __pyx_code_cache.entries[pos].code_object;
    Py_INCREF(code_object);
    return code_object;
}
static void __pyx_insert_code_object(int code_line, PyCodeObject* code_object) {
    int pos, i;
    __Pyx_CodeObjectCacheEntry* entries = __pyx_code_cache.entries;
    if (unlikely(!code_line)) {
        return;
    }
    if (unlikely(!entries)) {
        entries = (__Pyx_CodeObjectCacheEntry*)PyMem_Malloc(64*sizeof(__Pyx_CodeObjectCacheEntry));
        if (likely(entries)) {
            __pyx_code_cache.entries = entries;
            __pyx_code_cache.max_count = 64;
            __pyx_code_cache.count = 1;
            entries[0].code_line = code_line;
            entries[0].code_object = code_object;
            Py_INCREF(code_object);
        }
        return;
    }
    pos = __pyx_bisect_code_objects(__pyx_code_cache.entries, __pyx_code_cache.count, code_line);
    if ((pos < __pyx_code_cache.count) && unlikely(__pyx_code_cache.entries[pos].code_line == code_line)) {
        PyCodeObject* tmp = entries[pos].code_object;
        entries[pos].code_object = code_object;
        Py_DECREF(tmp);
        return;
    }
    if (__pyx_code_cache.count == __pyx_code_cache.max_count) {
        int new_max = __pyx_code_cache.max_count + 64;
        entries = (__Pyx_CodeObjectCacheEntry*)PyMem_Realloc(
            __pyx_code_cache.entries, ((size_t)new_max) * sizeof(__Pyx_CodeObjectCacheEntry));
        if (unlikely(!entries)) {
            return;
        }
        __pyx_code_cache.entries = entries;
        __pyx_code_cache.max_count = new_max;
    }
    for (i=__pyx_code_cache.count; i>pos; i--) {
        entries[i] = entries[i-1];
    }
    entries[pos].code_line = code_line;
    entries[pos].code_object = code_object;
    __pyx_code_cache.count++;
    Py_INCREF(code_object);
}

/* AddTraceback */
#include "compile.h"
#include "frameobject.h"
#include "traceback.h"
#if PY_VERSION_HEX >= 0x030b00a6
  #ifndef Py_BUILD_CORE
    #define Py_BUILD_CORE 1
  #endif
  #include "internal/pycore_frame.h"
#endif
static PyCodeObject* __Pyx_CreateCodeObjectForTraceback(
            const char *funcname, int c_line,
            int py_line, const char *filename) {
    PyCodeObject *py_code = NULL;
    PyObject *py_funcname = NULL;
    #if PY_MAJOR_VERSION < 3
    PyObject *py_srcfile = NULL;
    py_srcfile = PyString_FromString(filename);
    if (!py_srcfile) goto bad;
    #endif
    if (c_line) {
        #if PY_MAJOR_VERSION < 3
        py_funcname = PyString_FromFormat( "%s (%s:%d)", funcname, __pyx_cfilenm, c_line);
        if (!py_funcname) goto bad;
        #else
        py_funcname = PyUnicode_FromFormat( "%s (%s:%d)", funcname, __pyx_cfilenm, c_line);
        if (!py_funcname) goto bad;
        funcname = PyUnicode_AsUTF8(py_funcname);
        if (!funcname) goto bad;
        #endif
    }
    else {
        #if PY_MAJOR_VERSION < 3
        py_funcname = PyString_FromString(funcname);
        if (!py_funcname) goto bad;
        #endif
    }
    #if PY_MAJOR_VERSION < 3
    py_code = __Pyx_PyCode_New(
        0,
        0,
        0,
        0,
        0,
        __pyx_empty_bytes, /*PyObject *code,*/
        __pyx_empty_tuple, /*PyObject *consts,*/
        __pyx_empty_tuple, /*PyObject *names,*/
        __pyx_empty_tuple, /*PyObject *varnames,*/
        __pyx_empty_tuple, /*PyObject *freevars,*/
        __pyx_empty_tuple, /*PyObject *cellvars,*/
        py_srcfile,   /*PyObject *filename,*/
        py_funcname,  /*PyObject *name,*/
        py_line,
        __pyx_empty_bytes  /*PyObject *lnotab*/
    );
    Py_DECREF(py_srcfile);
    #else
    py_code = PyCode_NewEmpty(filename, funcname, py_line);
    #endif
    Py_XDECREF(py_funcname);  // XDECREF since it's only set on Py3 if cline
    return py_code;
bad:
    Py_XDECREF(py_funcname);
    #if PY_MAJOR_VERSION < 3
    Py_XDECREF(py_srcfile);
    #endif
    return NULL;
}
static void __Pyx_AddTraceback(const char *funcname, int c_line,
                               int py_line, const char *filename) {
    PyCodeObject *py_code = 0;
    PyFrameObject *py_frame = 0;
    PyThreadState *tstate = __Pyx_PyThreadState_Current;
    PyObject *ptype, *pvalue, *ptraceback;
    if (c_line) {
        c_line = __Pyx_CLineForTraceback(tstate, c_line);
    }
    py_code = __pyx_find_code_object(c_line ? -c_line : py_line);
    if (!py_code) {
        __Pyx_ErrFetchInState(tstate, &ptype, &pvalue, &ptraceback);
        py_code = __Pyx_CreateCodeObjectForTraceback(
            funcname, c_line, py_line, filename);
        if (!py_code) {
            /* If the code object creation fails, then we should clear the
               fetched exception references and propagate the new exception */
            Py_XDECREF(ptype);
            Py_XDECREF(pvalue);
            Py_XDECREF(ptraceback);
            goto bad;
        }
        __Pyx_ErrRestoreInState(tstate, ptype, pvalue, ptraceback);
        __pyx_insert_code_object(c_line ? -c_line : py_line, py_code);
    }
    py_frame = PyFrame_New(
        tstate,            /*PyThreadState *tstate,*/
        py_code,           /*PyCodeObject *code,*/
        __pyx_d,    /*PyObject *globals,*/
        0                  /*PyObject *locals*/
    );
    if (!py_frame) goto bad;
    __Pyx_PyFrame_SetLineNumber(py_frame, py_line);
    PyTraceBack_Here(py_frame);
bad:
    Py_XDECREF(py_code);
    Py_XDECREF(py_frame);
}

/* CIntFromPyVerify */
#define __PYX_VERIFY_RETURN_INT(target_type, func_type, func_value)\
    __PYX__VERIFY_RETURN_INT(target_type, func_type, func_value, 0)
#define __PYX_VERIFY_RETURN_INT_EXC(target_type, func_type, func_value)\
    __PYX__VERIFY_RETURN_INT(target_type, func_type, func_value, 1)
#define __PYX__VERIFY_RETURN_INT(target_type, func_type, func_value, exc)\
    {\
        func_type value = func_value;\
        if (sizeof(target_type) < sizeof(func_type)) {\
            if (unlikely(value != (func_type) (target_type) value)) {\
                func_type zero = 0;\
                if (exc && unlikely(value == (func_type)-1 && PyErr_Occurred()))\
                    return (target_type) -1;\
                if (is_unsigned && unlikely(value < zero))\
                    goto raise_neg_overflow;\
                else\
                    goto raise_overflow;\
            }\
        }\
        return (target_type) value;\
    }

/* CIntFromPy */
static CYTHON_INLINE int __Pyx_PyInt_As_int(PyObject *x) {
#ifdef __Pyx_HAS_GCC_DIAGNOSTIC
#pragma GCC diagnostic push
#pragma GCC diagnostic ignored "-Wconversion"
#endif
    const int neg_one = (int) -1, const_zero = (int) 0;
#ifdef __Pyx_HAS_GCC_DIAGNOSTIC
#pragma GCC diagnostic pop
#endif
    const int is_unsigned = neg_one > const_zero;
#if PY_MAJOR_VERSION < 3
    if (likely(PyInt_Check(x))) {
        if (sizeof(int) < sizeof(long)) {
            __PYX_VERIFY_RETURN_INT(int, long, PyInt_AS_LONG(x))
        } else {
            long val = PyInt_AS_LONG(x);
            if (is_unsigned && unlikely(val < 0)) {
                goto raise_neg_overflow;
            }
            return (int) val;
        }
    } else
#endif
    if (likely(PyLong_Check(x))) {
        if (is_unsigned) {
#if CYTHON_USE_PYLONG_INTERNALS
            const digit* digits = ((PyLongObject*)x)->ob_digit;
            switch (Py_SIZE(x)) {
                case  0: return (int) 0;
                case  1: __PYX_VERIFY_RETURN_INT(int, digit, digits[0])
                case 2:
                    if (8 * sizeof(int) > 1 * PyLong_SHIFT) {
                        if (8 * sizeof(unsigned long) > 2 * PyLong_SHIFT) {
                            __PYX_VERIFY_RETURN_INT(int, unsigned long, (((((unsigned long)digits[1]) << PyLong_SHIFT) | (unsigned long)digits[0])))
                        } else if (8 * sizeof(int) >= 2 * PyLong_SHIFT) {
                            return (int) (((((int)digits[1]) << PyLong_SHIFT) | (int)digits[0]));
                        }
                    }
                    break;
                case 3:
                    if (8 * sizeof(int) > 2 * PyLong_SHIFT) {
                        if (8 * sizeof(unsigned long) > 3 * PyLong_SHIFT) {
                            __PYX_VERIFY_RETURN_INT(int, unsigned long, (((((((unsigned long)digits[2]) << PyLong_SHIFT) | (unsigned long)digits[1]) << PyLong_SHIFT) | (unsigned long)digits[0])))
                        } else if (8 * sizeof(int) >= 3 * PyLong_SHIFT) {
                            return (int) (((((((int)digits[2]) << PyLong_SHIFT) | (int)digits[1]) << PyLong_SHIFT) | (int)digits[0]));
                        }
                    }
                    break;
                case 4:
                    if (8 * sizeof(int) > 3 * PyLong_SHIFT) {
                        if (8 * sizeof(unsigned long) > 4 * PyLong_SHIFT) {
                            __PYX_VERIFY_RETURN_INT(int, unsigned long, (((((((((unsigned long)digits[3]) << PyLong_SHIFT) | (unsigned long)digits[2]) << PyLong_SHIFT) | (unsigned long)digits[1]) << PyLong_SHIFT) | (unsigned long)digits[0])))
                        } else if (8 * sizeof(int) >= 4 * PyLong_SHIFT) {
                            return (int) (((((((((int)digits[3]) << PyLong_SHIFT) | (int)digits[2]) << PyLong_SHIFT) | (int)digits[1]) << PyLong_SHIFT) | (int)digits[0]));
                        }
                    }
                    break;
            }
#endif
#if CYTHON_COMPILING_IN_CPYTHON
            if (unlikely(Py_SIZE(x) < 0)) {
                goto raise_neg_overflow;
            }
#else
            {
                int result = PyObject_RichCompareBool(x, Py_False, Py_LT);
                if (unlikely(result < 0))
                    return (int) -1;
                if (unlikely(result == 1))
                    goto raise_neg_overflow;
            }
#endif
            if (sizeof(int) <= sizeof(unsigned long)) {
                __PYX_VERIFY_RETURN_INT_EXC(int, unsigned long, PyLong_AsUnsignedLong(x))
#ifdef HAVE_LONG_LONG
            } else if (sizeof(int) <= sizeof(unsigned PY_LONG_LONG)) {
                __PYX_VERIFY_RETURN_INT_EXC(int, unsigned PY_LONG_LONG, PyLong_AsUnsignedLongLong(x))
#endif
            }
        } else {
#if CYTHON_USE_PYLONG_INTERNALS
            const digit* digits = ((PyLongObject*)x)->ob_digit;
            switch (Py_SIZE(x)) {
                case  0: return (int) 0;
                case -1: __PYX_VERIFY_RETURN_INT(int, sdigit, (sdigit) (-(sdigit)digits[0]))
                case  1: __PYX_VERIFY_RETURN_INT(int,  digit, +digits[0])
                case -2:
                    if (8 * sizeof(int) - 1 > 1 * PyLong_SHIFT) {
                        if (8 * sizeof(unsigned long) > 2 * PyLong_SHIFT) {
                            __PYX_VERIFY_RETURN_INT(int, long, -(long) (((((unsigned long)digits[1]) << PyLong_SHIFT) | (unsigned long)digits[0])))
                        } else if (8 * sizeof(int) - 1 > 2 * PyLong_SHIFT) {
                            return (int) (((int)-1)*(((((int)digits[1]) << PyLong_SHIFT) | (int)digits[0])));
                        }
                    }
                    break;
                case 2:
                    if (8 * sizeof(int) > 1 * PyLong_SHIFT) {
                        if (8 * sizeof(unsigned long) > 2 * PyLong_SHIFT) {
                            __PYX_VERIFY_RETURN_INT(int, unsigned long, (((((unsigned long)digits[1]) << PyLong_SHIFT) | (unsigned long)digits[0])))
                        } else if (8 * sizeof(int) - 1 > 2 * PyLong_SHIFT) {
                            return (int) ((((((int)digits[1]) << PyLong_SHIFT) | (int)digits[0])));
                        }
                    }
                    break;
                case -3:
                    if (8 * sizeof(int) - 1 > 2 * PyLong_SHIFT) {
                        if (8 * sizeof(unsigned long) > 3 * PyLong_SHIFT) {
                            __PYX_VERIFY_RETURN_INT(int, long, -(long) (((((((unsigned long)digits[2]) << PyLong_SHIFT) | (unsigned long)digits[1]) << PyLong_SHIFT) | (unsigned long)digits[0])))
                        } else if (8 * sizeof(int) - 1 > 3 * PyLong_SHIFT) {
                            return (int) (((int)-1)*(((((((int)digits[2]) << PyLong_SHIFT) | (int)digits[1]) << PyLong_SHIFT) | (int)digits[0])));
                        }
                    }
                    break;
                case 3:
                    if (8 * sizeof(int) > 2 * PyLong_SHIFT) {
                        if (8 * sizeof(unsigned long) > 3 * PyLong_SHIFT) {
                            __PYX_VERIFY_RETURN_INT(int, unsigned long, (((((((unsigned long)digits[2]) << PyLong_SHIFT) | (unsigned long)digits[1]) << PyLong_SHIFT) | (unsigned long)digits[0])))
                        } else if (8 * sizeof(int) - 1 > 3 * PyLong_SHIFT) {
                            return (int) ((((((((int)digits[2]) << PyLong_SHIFT) | (int)digits[1]) << PyLong_SHIFT) | (int)digits[0])));
                        }
                    }
                    break;
                case -4:
                    if (8 * sizeof(int) - 1 > 3 * PyLong_SHIFT) {
                        if (8 * sizeof(unsigned long) > 4 * PyLong_SHIFT) {
                            __PYX_VERIFY_RETURN_INT(int, long, -(long) (((((((((unsigned long)digits[3]) << PyLong_SHIFT) | (unsigned long)digits[2]) << PyLong_SHIFT) | (unsigned long)digits[1]) << PyLong_SHIFT) | (unsigned long)digits[0])))
                        } else if (8 * sizeof(int) - 1 > 4 * PyLong_SHIFT) {
                            return (int) (((int)-1)*(((((((((int)digits[3]) << PyLong_SHIFT) | (int)digits[2]) << PyLong_SHIFT) | (int)digits[1]) << PyLong_SHIFT) | (int)digits[0])));
                        }
                    }
                    break;
                case 4:
                    if (8 * sizeof(int) > 3 * PyLong_SHIFT) {
                        if (8 * sizeof(unsigned long) > 4 * PyLong_SHIFT) {
                            __PYX_VERIFY_RETURN_INT(int, unsigned long, (((((((((unsigned long)digits[3]) << PyLong_SHIFT) | (unsigned long)digits[2]) << PyLong_SHIFT) | (unsigned long)digits[1]) << PyLong_SHIFT) | (unsigned long)digits[0])))
                        } else if (8 * sizeof(int) - 1 > 4 * PyLong_SHIFT) {
                            return (int) ((((((((((int)digits[3]) << PyLong_SHIFT) | (int)digits[2]) << PyLong_SHIFT) | (int)digits[1]) << PyLong_SHIFT) | (int)digits[0])));
                        }
                    }
                    break;
            }
#endif
            if (sizeof(int) <= sizeof(long)) {
                __PYX_VERIFY_RETURN_INT_EXC(int, long, PyLong_AsLong(x))
#ifdef HAVE_LONG_LONG
            } else if (sizeof(int) <= sizeof(PY_LONG_LONG)) {
                __PYX_VERIFY_RETURN_INT_EXC(int, PY_LONG_LONG, PyLong_AsLongLong(x))
#endif
            }
        }
        {
#if CYTHON_COMPILING_IN_PYPY && !defined(_PyLong_AsByteArray)
            PyErr_SetString(PyExc_RuntimeError,
                            "_PyLong_AsByteArray() not available in PyPy, cannot convert large numbers");
#else
            int val;
            PyObject *v = __Pyx_PyNumber_IntOrLong(x);
 #if PY_MAJOR_VERSION < 3
            if (likely(v) && !PyLong_Check(v)) {
                PyObject *tmp = v;
                v = PyNumber_Long(tmp);
                Py_DECREF(tmp);
            }
 #endif
            if (likely(v)) {
                int one = 1; int is_little = (int)*(unsigned char *)&one;
                unsigned char *bytes = (unsigned char *)&val;
                int ret = _PyLong_AsByteArray((PyLongObject *)v,
                                              bytes, sizeof(val),
                                              is_little, !is_unsigned);
                Py_DECREF(v);
                if (likely(!ret))
                    return val;
            }
#endif
            return (int) -1;
        }
    } else {
        int val;
        PyObject *tmp = __Pyx_PyNumber_IntOrLong(x);
        if (!tmp) return (int) -1;
        val = __Pyx_PyInt_As_int(tmp);
        Py_DECREF(tmp);
        return val;
    }
raise_overflow:
    PyErr_SetString(PyExc_OverflowError,
        "value too large to convert to int");
    return (int) -1;
raise_neg_overflow:
    PyErr_SetString(PyExc_OverflowError,
        "can't convert negative value to int");
    return (int) -1;
}

/* CIntToPy */
static CYTHON_INLINE PyObject* __Pyx_PyInt_From_int(int value) {
#ifdef __Pyx_HAS_GCC_DIAGNOSTIC
#pragma GCC diagnostic push
#pragma GCC diagnostic ignored "-Wconversion"
#endif
    const int neg_one = (int) -1, const_zero = (int) 0;
#ifdef __Pyx_HAS_GCC_DIAGNOSTIC
#pragma GCC diagnostic pop
#endif
    const int is_unsigned = neg_one > const_zero;
    if (is_unsigned) {
        if (sizeof(int) < sizeof(long)) {
            return PyInt_FromLong((long) value);
        } else if (sizeof(int) <= sizeof(unsigned long)) {
            return PyLong_FromUnsignedLong((unsigned long) value);
#ifdef HAVE_LONG_LONG
        } else if (sizeof(int) <= sizeof(unsigned PY_LONG_LONG)) {
            return PyLong_FromUnsignedLongLong((unsigned PY_LONG_LONG) value);
#endif
        }
    } else {
        if (sizeof(int) <= sizeof(long)) {
            return PyInt_FromLong((long) value);
#ifdef HAVE_LONG_LONG
        } else if (sizeof(int) <= sizeof(PY_LONG_LONG)) {
            return PyLong_FromLongLong((PY_LONG_LONG) value);
#endif
        }
    }
    {
        int one = 1; int little = (int)*(unsigned char *)&one;
        unsigned char *bytes = (unsigned char *)&value;
        return _PyLong_FromByteArray(bytes, sizeof(int),
                                     little, !is_unsigned);
    }
}

/* CIntToPy */
static CYTHON_INLINE PyObject* __Pyx_PyInt_From_long(long value) {
#ifdef __Pyx_HAS_GCC_DIAGNOSTIC
#pragma GCC diagnostic push
#pragma GCC diagnostic ignored "-Wconversion"
#endif
    const long neg_one = (long) -1, const_zero = (long) 0;
#ifdef __Pyx_HAS_GCC_DIAGNOSTIC
#pragma GCC diagnostic pop
#endif
    const int is_unsigned = neg_one > const_zero;
    if (is_unsigned) {
        if (sizeof(long) < sizeof(long)) {
            return PyInt_FromLong((long) value);
        } else if (sizeof(long) <= sizeof(unsigned long)) {
            return PyLong_FromUnsignedLong((unsigned long) value);
#ifdef HAVE_LONG_LONG
        } else if (sizeof(long) <= sizeof(unsigned PY_LONG_LONG)) {
            return PyLong_FromUnsignedLongLong((unsigned PY_LONG_LONG) value);
#endif
        }
    } else {
        if (sizeof(long) <= sizeof(long)) {
            return PyInt_FromLong((long) value);
#ifdef HAVE_LONG_LONG
        } else if (sizeof(long) <= sizeof(PY_LONG_LONG)) {
            return PyLong_FromLongLong((PY_LONG_LONG) value);
#endif
        }
    }
    {
        int one = 1; int little = (int)*(unsigned char *)&one;
        unsigned char *bytes = (unsigned char *)&value;
        return _PyLong_FromByteArray(bytes, sizeof(long),
                                     little, !is_unsigned);
    }
}

/* CIntFromPy */
static CYTHON_INLINE long __Pyx_PyInt_As_long(PyObject *x) {
#ifdef __Pyx_HAS_GCC_DIAGNOSTIC
#pragma GCC diagnostic push
#pragma GCC diagnostic ignored "-Wconversion"
#endif
    const long neg_one = (long) -1, const_zero = (long) 0;
#ifdef __Pyx_HAS_GCC_DIAGNOSTIC
#pragma GCC diagnostic pop
#endif
    const int is_unsigned = neg_one > const_zero;
#if PY_MAJOR_VERSION < 3
    if (likely(PyInt_Check(x))) {
        if (sizeof(long) < sizeof(long)) {
            __PYX_VERIFY_RETURN_INT(long, long, PyInt_AS_LONG(x))
        } else {
            long val = PyInt_AS_LONG(x);
            if (is_unsigned && unlikely(val < 0)) {
                goto raise_neg_overflow;
            }
            return (long) val;
        }
    } else
#endif
    if (likely(PyLong_Check(x))) {
        if (is_unsigned) {
#if CYTHON_USE_PYLONG_INTERNALS
            const digit* digits = ((PyLongObject*)x)->ob_digit;
            switch (Py_SIZE(x)) {
                case  0: return (long) 0;
                case  1: __PYX_VERIFY_RETURN_INT(long, digit, digits[0])
                case 2:
                    if (8 * sizeof(long) > 1 * PyLong_SHIFT) {
                        if (8 * sizeof(unsigned long) > 2 * PyLong_SHIFT) {
                            __PYX_VERIFY_RETURN_INT(long, unsigned long, (((((unsigned long)digits[1]) << PyLong_SHIFT) | (unsigned long)digits[0])))
                        } else if (8 * sizeof(long) >= 2 * PyLong_SHIFT) {
                            return (long) (((((long)digits[1]) << PyLong_SHIFT) | (long)digits[0]));
                        }
                    }
                    break;
                case 3:
                    if (8 * sizeof(long) > 2 * PyLong_SHIFT) {
                        if (8 * sizeof(unsigned long) > 3 * PyLong_SHIFT) {
                            __PYX_VERIFY_RETURN_INT(long, unsigned long, (((((((unsigned long)digits[2]) << PyLong_SHIFT) | (unsigned long)digits[1]) << PyLong_SHIFT) | (unsigned long)digits[0])))
                        } else if (8 * sizeof(long) >= 3 * PyLong_SHIFT) {
                            return (long) (((((((long)digits[2]) << PyLong_SHIFT) | (long)digits[1]) << PyLong_SHIFT) | (long)digits[0]));
                        }
                    }
                    break;
                case 4:
                    if (8 * sizeof(long) > 3 * PyLong_SHIFT) {
                        if (8 * sizeof(unsigned long) > 4 * PyLong_SHIFT) {
                            __PYX_VERIFY_RETURN_INT(long, unsigned long, (((((((((unsigned long)digits[3]) << PyLong_SHIFT) | (unsigned long)digits[2]) << PyLong_SHIFT) | (unsigned long)digits[1]) << PyLong_SHIFT) | (unsigned long)digits[0])))
                        } else if (8 * sizeof(long) >= 4 * PyLong_SHIFT) {
                            return (long) (((((((((long)digits[3]) << PyLong_SHIFT) | (long)digits[2]) << PyLong_SHIFT) | (long)digits[1]) << PyLong_SHIFT) | (long)digits[0]));
                        }
                    }
                    break;
            }
#endif
#if CYTHON_COMPILING_IN_CPYTHON
            if (unlikely(Py_SIZE(x) < 0)) {
                goto raise_neg_overflow;
            }
#else
            {
                int result = PyObject_RichCompareBool(x, Py_False, Py_LT);
                if (unlikely(result < 0))
                    return (long) -1;
                if (unlikely(result == 1))
                    goto raise_neg_overflow;
            }
#endif
            if (sizeof(long) <= sizeof(unsigned long)) {
                __PYX_VERIFY_RETURN_INT_EXC(long, unsigned long, PyLong_AsUnsignedLong(x))
#ifdef HAVE_LONG_LONG
            } else if (sizeof(long) <= sizeof(unsigned PY_LONG_LONG)) {
                __PYX_VERIFY_RETURN_INT_EXC(long, unsigned PY_LONG_LONG, PyLong_AsUnsignedLongLong(x))
#endif
            }
        } else {
#if CYTHON_USE_PYLONG_INTERNALS
            const digit* digits = ((PyLongObject*)x)->ob_digit;
            switch (Py_SIZE(x)) {
                case  0: return (long) 0;
                case -1: __PYX_VERIFY_RETURN_INT(long, sdigit, (sdigit) (-(sdigit)digits[0]))
                case  1: __PYX_VERIFY_RETURN_INT(long,  digit, +digits[0])
                case -2:
                    if (8 * sizeof(long) - 1 > 1 * PyLong_SHIFT) {
                        if (8 * sizeof(unsigned long) > 2 * PyLong_SHIFT) {
                            __PYX_VERIFY_RETURN_INT(long, long, -(long) (((((unsigned long)digits[1]) << PyLong_SHIFT) | (unsigned long)digits[0])))
                        } else if (8 * sizeof(long) - 1 > 2 * PyLong_SHIFT) {
                            return (long) (((long)-1)*(((((long)digits[1]) << PyLong_SHIFT) | (long)digits[0])));
                        }
                    }
                    break;
                case 2:
                    if (8 * sizeof(long) > 1 * PyLong_SHIFT) {
                        if (8 * sizeof(unsigned long) > 2 * PyLong_SHIFT) {
                            __PYX_VERIFY_RETURN_INT(long, unsigned long, (((((unsigned long)digits[1]) << PyLong_SHIFT) | (unsigned long)digits[0])))
                        } else if (8 * sizeof(long) - 1 > 2 * PyLong_SHIFT) {
                            return (long) ((((((long)digits[1]) << PyLong_SHIFT) | (long)digits[0])));
                        }
                    }
                    break;
                case -3:
                    if (8 * sizeof(long) - 1 > 2 * PyLong_SHIFT) {
                        if (8 * sizeof(unsigned long) > 3 * PyLong_SHIFT) {
                            __PYX_VERIFY_RETURN_INT(long, long, -(long) (((((((unsigned long)digits[2]) << PyLong_SHIFT) | (unsigned long)digits[1]) << PyLong_SHIFT) | (unsigned long)digits[0])))
                        } else if (8 * sizeof(long) - 1 > 3 * PyLong_SHIFT) {
                            return (long) (((long)-1)*(((((((long)digits[2]) << PyLong_SHIFT) | (long)digits[1]) << PyLong_SHIFT) | (long)digits[0])));
                        }
                    }
                    break;
                case 3:
                    if (8 * sizeof(long) > 2 * PyLong_SHIFT) {
                        if (8 * sizeof(unsigned long) > 3 * PyLong_SHIFT) {
                            __PYX_VERIFY_RETURN_INT(long, unsigned long, (((((((unsigned long)digits[2]) << PyLong_SHIFT) | (unsigned long)digits[1]) << PyLong_SHIFT) | (unsigned long)digits[0])))
                        } else if (8 * sizeof(long) - 1 > 3 * PyLong_SHIFT) {
                            return (long) ((((((((long)digits[2]) << PyLong_SHIFT) | (long)digits[1]) << PyLong_SHIFT) | (long)digits[0])));
                        }
                    }
                    break;
                case -4:
                    if (8 * sizeof(long) - 1 > 3 * PyLong_SHIFT) {
                        if (8 * sizeof(unsigned long) > 4 * PyLong_SHIFT) {
                            __PYX_VERIFY_RETURN_INT(long, long, -(long) (((((((((unsigned long)digits[3]) << PyLong_SHIFT) | (unsigned long)digits[2]) << PyLong_SHIFT) | (unsigned long)digits[1]) << PyLong_SHIFT) | (unsigned long)digits[0])))
                        } else if (8 * sizeof(long) - 1 > 4 * PyLong_SHIFT) {
                            return (long) (((long)-1)*(((((((((long)digits[3]) << PyLong_SHIFT) | (long)digits[2]) << PyLong_SHIFT) | (long)digits[1]) << PyLong_SHIFT) | (long)digits[0])));
                        }
                    }
                    break;
                case 4:
                    if (8 * sizeof(long) > 3 * PyLong_SHIFT) {
                        if (8 * sizeof(unsigned long) > 4 * PyLong_SHIFT) {
                            __PYX_VERIFY_RETURN_INT(long, unsigned long, (((((((((unsigned long)digits[3]) << PyLong_SHIFT) | (unsigned long)digits[2]) << PyLong_SHIFT) | (unsigned long)digits[1]) << PyLong_SHIFT) | (unsigned long)digits[0])))
                        } else if (8 * sizeof(long) - 1 > 4 * PyLong_SHIFT) {
                            return (long) ((((((((((long)digits[3]) << PyLong_SHIFT) | (long)digits[2]) << PyLong_SHIFT) | (long)digits[1]) << PyLong_SHIFT) | (long)digits[0])));
                        }
                    }
                    break;
            }
#endif
            if (sizeof(long) <= sizeof(long)) {
                __PYX_VERIFY_RETURN_INT_EXC(long, long, PyLong_AsLong(x))
#ifdef HAVE_LONG_LONG
            } else if (sizeof(long) <= sizeof(PY_LONG_LONG)) {
                __PYX_VERIFY_RETURN_INT_EXC(long, PY_LONG_LONG, PyLong_AsLongLong(x))
#endif
            }
        }
        {
#if CYTHON_COMPILING_IN_PYPY && !defined(_PyLong_AsByteArray)
            PyErr_SetString(PyExc_RuntimeError,
                            "_PyLong_AsByteArray() not available in PyPy, cannot convert large numbers");
#else
            long val;
            PyObject *v = __Pyx_PyNumber_IntOrLong(x);
 #if PY_MAJOR_VERSION < 3
            if (likely(v) && !PyLong_Check(v)) {
                PyObject *tmp = v;
                v = PyNumber_Long(tmp);
                Py_DECREF(tmp);
            }
 #endif
            if (likely(v)) {
                int one = 1; int is_little = (int)*(unsigned char *)&one;
                unsigned char *bytes = (unsigned char *)&val;
                int ret = _PyLong_AsByteArray((PyLongObject *)v,
                                              bytes, sizeof(val),
                                              is_little, !is_unsigned);
                Py_DECREF(v);
                if (likely(!ret))
                    return val;
            }
#endif
            return (long) -1;
        }
    } else {
        long val;
        PyObject *tmp = __Pyx_PyNumber_IntOrLong(x);
        if (!tmp) return (long) -1;
        val = __Pyx_PyInt_As_long(tmp);
        Py_DECREF(tmp);
        return val;
    }
raise_overflow:
    PyErr_SetString(PyExc_OverflowError,
        "value too large to convert to long");
    return (long) -1;
raise_neg_overflow:
    PyErr_SetString(PyExc_OverflowError,
        "can't convert negative value to long");
    return (long) -1;
}

/* FastTypeChecks */
#if CYTHON_COMPILING_IN_CPYTHON
static int __Pyx_InBases(PyTypeObject *a, PyTypeObject *b) {
    while (a) {
        a = a->tp_base;
        if (a == b)
            return 1;
    }
    return b == &PyBaseObject_Type;
}
static CYTHON_INLINE int __Pyx_IsSubtype(PyTypeObject *a, PyTypeObject *b) {
    PyObject *mro;
    if (a == b) return 1;
    mro = a->tp_mro;
    if (likely(mro)) {
        Py_ssize_t i, n;
        n = PyTuple_GET_SIZE(mro);
        for (i = 0; i < n; i++) {
            if (PyTuple_GET_ITEM(mro, i) == (PyObject *)b)
                return 1;
        }
        return 0;
    }
    return __Pyx_InBases(a, b);
}
#if PY_MAJOR_VERSION == 2
static int __Pyx_inner_PyErr_GivenExceptionMatches2(PyObject *err, PyObject* exc_type1, PyObject* exc_type2) {
    PyObject *exception, *value, *tb;
    int res;
    __Pyx_PyThreadState_declare
    __Pyx_PyThreadState_assign
    __Pyx_ErrFetch(&exception, &value, &tb);
    res = exc_type1 ? PyObject_IsSubclass(err, exc_type1) : 0;
    if (unlikely(res == -1)) {
        PyErr_WriteUnraisable(err);
        res = 0;
    }
    if (!res) {
        res = PyObject_IsSubclass(err, exc_type2);
        if (unlikely(res == -1)) {
            PyErr_WriteUnraisable(err);
            res = 0;
        }
    }
    __Pyx_ErrRestore(exception, value, tb);
    return res;
}
#else
static CYTHON_INLINE int __Pyx_inner_PyErr_GivenExceptionMatches2(PyObject *err, PyObject* exc_type1, PyObject *exc_type2) {
    int res = exc_type1 ? __Pyx_IsSubtype((PyTypeObject*)err, (PyTypeObject*)exc_type1) : 0;
    if (!res) {
        res = __Pyx_IsSubtype((PyTypeObject*)err, (PyTypeObject*)exc_type2);
    }
    return res;
}
#endif
static int __Pyx_PyErr_GivenExceptionMatchesTuple(PyObject *exc_type, PyObject *tuple) {
    Py_ssize_t i, n;
    assert(PyExceptionClass_Check(exc_type));
    n = PyTuple_GET_SIZE(tuple);
#if PY_MAJOR_VERSION >= 3
    for (i=0; i<n; i++) {
        if (exc_type == PyTuple_GET_ITEM(tuple, i)) return 1;
    }
#endif
    for (i=0; i<n; i++) {
        PyObject *t = PyTuple_GET_ITEM(tuple, i);
        #if PY_MAJOR_VERSION < 3
        if (likely(exc_type == t)) return 1;
        #endif
        if (likely(PyExceptionClass_Check(t))) {
            if (__Pyx_inner_PyErr_GivenExceptionMatches2(exc_type, NULL, t)) return 1;
        } else {
        }
    }
    return 0;
}
static CYTHON_INLINE int __Pyx_PyErr_GivenExceptionMatches(PyObject *err, PyObject* exc_type) {
    if (likely(err == exc_type)) return 1;
    if (likely(PyExceptionClass_Check(err))) {
        if (likely(PyExceptionClass_Check(exc_type))) {
            return __Pyx_inner_PyErr_GivenExceptionMatches2(err, NULL, exc_type);
        } else if (likely(PyTuple_Check(exc_type))) {
            return __Pyx_PyErr_GivenExceptionMatchesTuple(err, exc_type);
        } else {
        }
    }
    return PyErr_GivenExceptionMatches(err, exc_type);
}
static CYTHON_INLINE int __Pyx_PyErr_GivenExceptionMatches2(PyObject *err, PyObject *exc_type1, PyObject *exc_type2) {
    assert(PyExceptionClass_Check(exc_type1));
    assert(PyExceptionClass_Check(exc_type2));
    if (likely(err == exc_type1 || err == exc_type2)) return 1;
    if (likely(PyExceptionClass_Check(err))) {
        return __Pyx_inner_PyErr_GivenExceptionMatches2(err, exc_type1, exc_type2);
    }
    return (PyErr_GivenExceptionMatches(err, exc_type1) || PyErr_GivenExceptionMatches(err, exc_type2));
}
#endif

/* CheckBinaryVersion */
static int __Pyx_check_binary_version(void) {
    char ctversion[5];
    int same=1, i, found_dot;
    const char* rt_from_call = Py_GetVersion();
    PyOS_snprintf(ctversion, 5, "%d.%d", PY_MAJOR_VERSION, PY_MINOR_VERSION);
    found_dot = 0;
    for (i = 0; i < 4; i++) {
        if (!ctversion[i]) {
            same = (rt_from_call[i] < '0' || rt_from_call[i] > '9');
            break;
        }
        if (rt_from_call[i] != ctversion[i]) {
            same = 0;
            break;
        }
    }
    if (!same) {
        char rtversion[5] = {'\0'};
        char message[200];
        for (i=0; i<4; ++i) {
            if (rt_from_call[i] == '.') {
                if (found_dot) break;
                found_dot = 1;
            } else if (rt_from_call[i] < '0' || rt_from_call[i] > '9') {
                break;
            }
            rtversion[i] = rt_from_call[i];
        }
        PyOS_snprintf(message, sizeof(message),
                      "compiletime version %s of module '%.100s' "
                      "does not match runtime version %s",
                      ctversion, __Pyx_MODULE_NAME, rtversion);
        return PyErr_WarnEx(NULL, message, 1);
    }
    return 0;
}

/* InitStrings */
static int __Pyx_InitStrings(__Pyx_StringTabEntry *t) {
    while (t->p) {
        #if PY_MAJOR_VERSION < 3
        if (t->is_unicode) {
            *t->p = PyUnicode_DecodeUTF8(t->s, t->n - 1, NULL);
        } else if (t->intern) {
            *t->p = PyString_InternFromString(t->s);
        } else {
            *t->p = PyString_FromStringAndSize(t->s, t->n - 1);
        }
        #else
        if (t->is_unicode | t->is_str) {
            if (t->intern) {
                *t->p = PyUnicode_InternFromString(t->s);
            } else if (t->encoding) {
                *t->p = PyUnicode_Decode(t->s, t->n - 1, t->encoding, NULL);
            } else {
                *t->p = PyUnicode_FromStringAndSize(t->s, t->n - 1);
            }
        } else {
            *t->p = PyBytes_FromStringAndSize(t->s, t->n - 1);
        }
        #endif
        if (!*t->p)
            return -1;
        if (PyObject_Hash(*t->p) == -1)
            return -1;
        ++t;
    }
    return 0;
}

static CYTHON_INLINE PyObject* __Pyx_PyUnicode_FromString(const char* c_str) {
    return __Pyx_PyUnicode_FromStringAndSize(c_str, (Py_ssize_t)strlen(c_str));
}
static CYTHON_INLINE const char* __Pyx_PyObject_AsString(PyObject* o) {
    Py_ssize_t ignore;
    return __Pyx_PyObject_AsStringAndSize(o, &ignore);
}
#if __PYX_DEFAULT_STRING_ENCODING_IS_ASCII || __PYX_DEFAULT_STRING_ENCODING_IS_DEFAULT
#if !CYTHON_PEP393_ENABLED
static const char* __Pyx_PyUnicode_AsStringAndSize(PyObject* o, Py_ssize_t *length) {
    char* defenc_c;
    PyObject* defenc = _PyUnicode_AsDefaultEncodedString(o, NULL);
    if (!defenc) return NULL;
    defenc_c = PyBytes_AS_STRING(defenc);
#if __PYX_DEFAULT_STRING_ENCODING_IS_ASCII
    {
        char* end = defenc_c + PyBytes_GET_SIZE(defenc);
        char* c;
        for (c = defenc_c; c < end; c++) {
            if ((unsigned char) (*c) >= 128) {
                PyUnicode_AsASCIIString(o);
                return NULL;
            }
        }
    }
#endif
    *length = PyBytes_GET_SIZE(defenc);
    return defenc_c;
}
#else
static CYTHON_INLINE const char* __Pyx_PyUnicode_AsStringAndSize(PyObject* o, Py_ssize_t *length) {
    if (unlikely(__Pyx_PyUnicode_READY(o) == -1)) return NULL;
#if __PYX_DEFAULT_STRING_ENCODING_IS_ASCII
    if (likely(PyUnicode_IS_ASCII(o))) {
        *length = PyUnicode_GET_LENGTH(o);
        return PyUnicode_AsUTF8(o);
    } else {
        PyUnicode_AsASCIIString(o);
        return NULL;
    }
#else
    return PyUnicode_AsUTF8AndSize(o, length);
#endif
}
#endif
#endif
static CYTHON_INLINE const char* __Pyx_PyObject_AsStringAndSize(PyObject* o, Py_ssize_t *length) {
#if __PYX_DEFAULT_STRING_ENCODING_IS_ASCII || __PYX_DEFAULT_STRING_ENCODING_IS_DEFAULT
    if (
#if PY_MAJOR_VERSION < 3 && __PYX_DEFAULT_STRING_ENCODING_IS_ASCII
            __Pyx_sys_getdefaultencoding_not_ascii &&
#endif
            PyUnicode_Check(o)) {
        return __Pyx_PyUnicode_AsStringAndSize(o, length);
    } else
#endif
#if (!CYTHON_COMPILING_IN_PYPY) || (defined(PyByteArray_AS_STRING) && defined(PyByteArray_GET_SIZE))
    if (PyByteArray_Check(o)) {
        *length = PyByteArray_GET_SIZE(o);
        return PyByteArray_AS_STRING(o);
    } else
#endif
    {
        char* result;
        int r = PyBytes_AsStringAndSize(o, &result, length);
        if (unlikely(r < 0)) {
            return NULL;
        } else {
            return result;
        }
    }
}
static CYTHON_INLINE int __Pyx_PyObject_IsTrue(PyObject* x) {
   int is_true = x == Py_True;
   if (is_true | (x == Py_False) | (x == Py_None)) return is_true;
   else return PyObject_IsTrue(x);
}
static CYTHON_INLINE int __Pyx_PyObject_IsTrueAndDecref(PyObject* x) {
    int retval;
    if (unlikely(!x)) return -1;
    retval = __Pyx_PyObject_IsTrue(x);
    Py_DECREF(x);
    return retval;
}
static PyObject* __Pyx_PyNumber_IntOrLongWrongResultType(PyObject* result, const char* type_name) {
#if PY_MAJOR_VERSION >= 3
    if (PyLong_Check(result)) {
        if (PyErr_WarnFormat(PyExc_DeprecationWarning, 1,
                "__int__ returned non-int (type %.200s).  "
                "The ability to return an instance of a strict subclass of int "
                "is deprecated, and may be removed in a future version of Python.",
                Py_TYPE(result)->tp_name)) {
            Py_DECREF(result);
            return NULL;
        }
        return result;
    }
#endif
    PyErr_Format(PyExc_TypeError,
                 "__%.4s__ returned non-%.4s (type %.200s)",
                 type_name, type_name, Py_TYPE(result)->tp_name);
    Py_DECREF(result);
    return NULL;
}
static CYTHON_INLINE PyObject* __Pyx_PyNumber_IntOrLong(PyObject* x) {
#if CYTHON_USE_TYPE_SLOTS
  PyNumberMethods *m;
#endif
  const char *name = NULL;
  PyObject *res = NULL;
#if PY_MAJOR_VERSION < 3
  if (likely(PyInt_Check(x) || PyLong_Check(x)))
#else
  if (likely(PyLong_Check(x)))
#endif
    return __Pyx_NewRef(x);
#if CYTHON_USE_TYPE_SLOTS
  m = Py_TYPE(x)->tp_as_number;
  #if PY_MAJOR_VERSION < 3
  if (m && m->nb_int) {
    name = "int";
    res = m->nb_int(x);
  }
  else if (m && m->nb_long) {
    name = "long";
    res = m->nb_long(x);
  }
  #else
  if (likely(m && m->nb_int)) {
    name = "int";
    res = m->nb_int(x);
  }
  #endif
#else
  if (!PyBytes_CheckExact(x) && !PyUnicode_CheckExact(x)) {
    res = PyNumber_Int(x);
  }
#endif
  if (likely(res)) {
#if PY_MAJOR_VERSION < 3
    if (unlikely(!PyInt_Check(res) && !PyLong_Check(res))) {
#else
    if (unlikely(!PyLong_CheckExact(res))) {
#endif
        return __Pyx_PyNumber_IntOrLongWrongResultType(res, name);
    }
  }
  else if (!PyErr_Occurred()) {
    PyErr_SetString(PyExc_TypeError,
                    "an integer is required");
  }
  return res;
}
static CYTHON_INLINE Py_ssize_t __Pyx_PyIndex_AsSsize_t(PyObject* b) {
  Py_ssize_t ival;
  PyObject *x;
#if PY_MAJOR_VERSION < 3
  if (likely(PyInt_CheckExact(b))) {
    if (sizeof(Py_ssize_t) >= sizeof(long))
        return PyInt_AS_LONG(b);
    else
        return PyInt_AsSsize_t(b);
  }
#endif
  if (likely(PyLong_CheckExact(b))) {
    #if CYTHON_USE_PYLONG_INTERNALS
    const digit* digits = ((PyLongObject*)b)->ob_digit;
    const Py_ssize_t size = Py_SIZE(b);
    if (likely(__Pyx_sst_abs(size) <= 1)) {
        ival = likely(size) ? digits[0] : 0;
        if (size == -1) ival = -ival;
        return ival;
    } else {
      switch (size) {
         case 2:
           if (8 * sizeof(Py_ssize_t) > 2 * PyLong_SHIFT) {
             return (Py_ssize_t) (((((size_t)digits[1]) << PyLong_SHIFT) | (size_t)digits[0]));
           }
           break;
         case -2:
           if (8 * sizeof(Py_ssize_t) > 2 * PyLong_SHIFT) {
             return -(Py_ssize_t) (((((size_t)digits[1]) << PyLong_SHIFT) | (size_t)digits[0]));
           }
           break;
         case 3:
           if (8 * sizeof(Py_ssize_t) > 3 * PyLong_SHIFT) {
             return (Py_ssize_t) (((((((size_t)digits[2]) << PyLong_SHIFT) | (size_t)digits[1]) << PyLong_SHIFT) | (size_t)digits[0]));
           }
           break;
         case -3:
           if (8 * sizeof(Py_ssize_t) > 3 * PyLong_SHIFT) {
             return -(Py_ssize_t) (((((((size_t)digits[2]) << PyLong_SHIFT) | (size_t)digits[1]) << PyLong_SHIFT) | (size_t)digits[0]));
           }
           break;
         case 4:
           if (8 * sizeof(Py_ssize_t) > 4 * PyLong_SHIFT) {
             return (Py_ssize_t) (((((((((size_t)digits[3]) << PyLong_SHIFT) | (size_t)digits[2]) << PyLong_SHIFT) | (size_t)digits[1]) << PyLong_SHIFT) | (size_t)digits[0]));
           }
           break;
         case -4:
           if (8 * sizeof(Py_ssize_t) > 4 * PyLong_SHIFT) {
             return -(Py_ssize_t) (((((((((size_t)digits[3]) << PyLong_SHIFT) | (size_t)digits[2]) << PyLong_SHIFT) | (size_t)digits[1]) << PyLong_SHIFT) | (size_t)digits[0]));
           }
           break;
      }
    }
    #endif
    return PyLong_AsSsize_t(b);
  }
  x = PyNumber_Index(b);
  if (!x) return -1;
  ival = PyInt_AsSsize_t(x);
  Py_DECREF(x);
  return ival;
}
static CYTHON_INLINE Py_hash_t __Pyx_PyIndex_AsHash_t(PyObject* o) {
  if (sizeof(Py_hash_t) == sizeof(Py_ssize_t)) {
    return (Py_hash_t) __Pyx_PyIndex_AsSsize_t(o);
#if PY_MAJOR_VERSION < 3
  } else if (likely(PyInt_CheckExact(o))) {
    return PyInt_AS_LONG(o);
#endif
  } else {
    Py_ssize_t ival;
    PyObject *x;
    x = PyNumber_Index(o);
    if (!x) return -1;
    ival = PyInt_AsLong(x);
    Py_DECREF(x);
    return ival;
  }
}
static CYTHON_INLINE PyObject * __Pyx_PyBool_FromLong(long b) {
  return b ? __Pyx_NewRef(Py_True) : __Pyx_NewRef(Py_False);
}
static CYTHON_INLINE PyObject * __Pyx_PyInt_FromSize_t(size_t ival) {
    return PyInt_FromSize_t(ival);
}


#endif /* Py_PYTHON_H */
```
