## Supplementary material for "Hyperparasitism and the evolution of parasite virulence": Code_base: simple_sandhu_mimic.py

```
"""
This file is used to make Fig. S3. It is used to verify whether we see similar
results to Sandhu 2021, when our system is similar to theirs.
"""

import sys, os, shutil
import runner
import numpy as np
from math import sqrt, exp, dist, floor, ceil, isnan
import plotly.graph_objects as go
from time import sleep
import matplotlib.pyplot as plt
import seaborn as sn
import imageio
import multiprocessing as mp
import pandas as pd

#These local libraries are used to reduce the length of the file
sys.path.append("../src/Models/")

import singular_strategy as ss

sys.path.append("../src/zero_finder/")

import zero_finder_utilities as zf

sys.path.append("../src/figures/")

import figure_utilities as ff

from datetime import datetime

#This creates a list of words and cleans them for use later
with open("../data/words.txt", encoding='utf8') as f:
    lines = f.readlines()
    
lines_cleaned = []

for line in lines:
    if line[0] != "#" and "www" not in line and line[0] != " ":
        lines_cleaned.append(line[:-1])


#This function calculates the gradient vector for a given parameter set
def calculate_alpha_gradient_vector(sol, resolution, beta_max, alpha_max, alpha_min_res, alpha_max_res, sigma_max, sigma, b, q, d, rho, eta, gamma, lam, c1, c2, hyper, seed, depth):
    
    alpha_grad_vec = []
    alpha_val = []
    
    for i in range(resolution+1):
        
        #calculate the value of alpha
        alpha = alpha_min_res + (alpha_max_res - alpha_min_res) * (i / resolution)
        alpha_val.append(alpha)

        #and corresponding value of beta and it's derivatives
        beta = zf.beta_func(beta_max, alpha, alpha_max, sigma, sigma_max, c1, c2)
        
        dbetadalpha = zf.dbetadalpha_func(
            beta_max, alpha, alpha_max, sigma, sigma_max, c1, c2
        )

        #Solve the system to ecological steady state
        y = sol.eco_steady_state(
            beta, alpha, sigma, b, q, d, rho, eta, gamma, lam, c1, c2, hyper, seed
        )
        
        S = y[0]*(y[0] >= 1e-3)
        I = y[1]*(y[1] >= 1e-3)
        H = y[2]*(y[2] >= 1e-3)
        
#         if depth == 0 and eta == 1.0:
#             print(y, S, I, H, eta, alpha, beta)
        if y[3] != 1:
            print(S, I, H, rho, lam, eta, alpha)
            raise Exception("Solver did not exit early due to convergence")
        #Determine which version of the gradient we need. If the parasite is extinct just add an
        #arbitrary value
        if (I == 0 and H == 0):
            alpha_grad = 1*(alpha <= alpha_max/2) - 1*(alpha> alpha_max/2)
        elif (H == 0):
            alpha_grad = dbetadalpha*S/(d + alpha + gamma) - (beta*S)/((d + alpha + gamma)**2)
#         elif (I == 0):
#             alpha_grad = eta*dbetadalpha*S/(d + lam*alpha + gamma) - eta*beta*S/((d + lam*alpha + gamma)**2)
        else:
#             if rho != 1:
            alpha_grad = ss.fitness_gradient_alpha(
                beta, alpha, sigma, dbetadalpha, H, S, eta, lam, gamma, d, rho
            )
#             else:
#                 if (beta*S - sigma*H - (d + alpha)) > (eta*beta*S - (d + alpha*lam)):
#                     alpha_grad = dbetadalpha*S  - 1
#                 else:
#                     alpha_grad = eta*dbetadalpha*S/(lam*alpha + d + gamma) - eta*beta*S/((lam*alpha + d + gamma)**2)
        #Convert the value into a sign
        alpha_grad = (alpha_grad > 0)*1 + (alpha_grad < 0)*-1
        alpha_grad_vec.append(alpha_grad)
        
    return alpha_grad_vec, alpha_val

#Iterate through the gradient vector to find any sign changes
def return_zeros(grad_vec, param_space):
    
    poss_attractors = []
    poss_repellers = []
    for i,val in enumerate(grad_vec[:-1]):
        if val == 1 and grad_vec[i+1] == -1:
            poss_attractors.append([param_space[i], param_space[i+1]])
        elif val == -1 and grad_vec[i+1] == 1:
            poss_repellers.append([param_space[i], param_space[i+1]])
        elif val == 0 and grad_vec[i+1] == -1:
            poss_attractors.append([param_space[i], param_space[i+1]])
            
    return poss_attractors, poss_repellers 
    
#This function calculates the singular strategy using the above functions to calculate the gradient and then find any sign changes
def recusive_sing_strat_function(sol, depth, iter_max, resolution, beta_max, alpha_max, alpha_min_res, alpha_max_res, sigma_max, sigma, b, q, d, rho, eta, gamma, lam, c1, c2, hyper, seed):
    
    
    sing_strats = []
    
    #We fill these lists with attractors and repellers
    attractors = []
    repellers = []
    
    #If we have resolved sufficiently we do one final resolution before exiting
    if depth == iter_max:
        #Calculating the alpha gradient vector
        alpha_grad_vec, alpha_val = calculate_alpha_gradient_vector(sol, resolution, beta_max, alpha_max, alpha_min_res, alpha_max_res, sigma_max, sigma, b, q, d, rho, eta, gamma, lam,c1, c2, hyper, seed, depth)
        
        #Identifying singular strategies
        poss_attractors, poss_repellers = return_zeros(alpha_grad_vec,alpha_val)
#         if len(poss_attractors) > 2:
#             print(f"On run {eta} on rho {rho} with depth {depth}, and there are {len(poss_attractors)} possible singular strategies     ", end = "\r")
        #Allocating them to their respective places
        for zeros in poss_attractors:
            #We take the lower element as the midpoint can sometimes be outside the bounds of the hyperparasites existance
            attractors.append(zeros[0])
        for zeros in poss_repellers:
            repellers.append(zeros[0])
            
    #If we are not at our maximum depth we find possible positions of the singular strategy
    else:
        #First we calculate the alpha gradient
        alpha_grad_vec, alpha_val = calculate_alpha_gradient_vector(sol, resolution, beta_max, alpha_max, alpha_min_res, alpha_max_res, sigma_max, sigma, b, q, d, rho, eta, gamma, lam,c1, c2, hyper, seed, depth)
        
        #Check for any sign changes using our function
        poss_attractors, poss_repellers = return_zeros(alpha_grad_vec,alpha_val)
#         if len(poss_attractors) > 2:
#             print(f"On run {eta} on rho {rho} with depth {depth}, and there are {len(poss_attractors)} possible singular strategies     ")
#             for pair in poss_attractors:
#                 print(pair[1] - pair[0])
        #Check bounds for any attractor regions
        if depth == 0:
            if alpha_grad_vec[0] == -1:
                attractors.append(alpha_val[0])
            if alpha_grad_vec[-1] == 1:
                attractors.append(alpha_val[-1])
                
        #Rerun the process of finding singular strategies for each of our possible attractors and repellers
        for pair in poss_attractors:
            temp_attractors, temp_repellers = recusive_sing_strat_function(sol, depth+1, iter_max, resolution, beta_max, alpha_max, pair[0], pair[1], sigma_max, sigma, b, q, d, rho, eta, gamma, lam, c1, c2, hyper, seed)
            
            for val in temp_attractors:
                attractors.append(val)
                
        for pair in poss_repellers:
            temp_attractors, temp_repellers = recusive_sing_strat_function(sol, depth+1, iter_max, resolution, beta_max, alpha_max, pair[0], pair[1], sigma_max, sigma, b, q, d, rho, eta, gamma, lam, c1, c2, hyper, seed)
                
            for val in temp_repellers:
                repellers.append(val)
                
    #We then clean up the attractors in case there are any singular strategies very close together
    if depth == 0:
        attractors = zf.clean_up_sing_strats(attractors)
        repellers = zf.clean_up_sing_strats(repellers)
#         print(eta, attractors)
#         print(alpha_grad_vec)
    return attractors, repellers


sol = runner.PySolver()

rho = 1.0

lams = [0.5, 1.0, 2.0]
# lams = [2.0]
b = 2.0
q = 0.1
d = 0.1
gamma = 0.1

hyper = 1.0
c1 = 0.75
c2 = 2.25
b_base = 1
q_base = 1

alpha_max = 5
sigma_max = 0.4
beta_max = 5

sigma_min = 0.0
sigma_max_range = sigma_max

sigma = sigma_max

iter_max = 3
res_scalar = 4
depth_max = iter_max
resolution = 100
param_res = 200

seed = 10000

sigma_init = 100

beta_max = beta_max
alpha_max_param = alpha_max 
sigma_max_param = sigma_max

param_res = 100

eta_min = 0.0
eta_max = 2.0
etas = [eta_min + (eta_max - eta_min)*i/param_res for i in range(param_res + 1)]

texts = [["(A)", "(B)", "(C)"],["(D)", "(E)", "(F)"], ["(G)","(H)","(I)"], ["(J)", "(K)", "(L)"]]

titles = ["Hypovirulence ", "No direct effect on virulence ", "Hypervirulence "]

fig1 = plt.figure(figsize = (40, 44))
gs1 = fig1.add_gridspec(4,len(lams))
ax1 = []

for i in range(2):
    temp_ax = []
    for j in range(len(lams)):
        temp_ax.append(fig1.add_subplot(gs1[i,j]))
    ax1.append(temp_ax)
    
alpha_init = 10
#We initialise the system with the hyperparasites absent to find the natural ESS of the parasite
sol.alpha_ad_dyn(beta_max, alpha_max, sigma_max, b, q, d, rho, 0.0, gamma, 0.0, c1, c2, 0.0, seed, alpha_init, sigma_init, H_density = 0.0)
df = pd.read_csv(f"../data/alpha_evo/data_set{seed}.csv")

evo_steps = df["Evolutionary_step"].values
dft = df[df["Evolutionary_step"]==evo_steps[-1]]


host_density = dft["Density_of_Hosts"].iloc[0]
para_density = sum(dft["Density_of_parasite"].values)
hyper_density = 0.1

seed += 1

print(host_density, para_density, hyper_density)
for lam_ind, lam in enumerate(lams):
    
    temp_attractors_rec = []
    temp_attractors_no_rec = []
    
    temp_repellers_rec = []
    temp_repellers_no_rec = []
    
    temp_etas_rec = []
    temp_etas_no_rec = []
    
    temp_etas_rec_rep = []
    temp_etas_no_rec_rep = []
    
    temp_attractors_rec_no_hyper = []
    temp_attractors_no_rec_no_hyper = []
    
    temp_etas_rec_no_hyper = []
    temp_etas_no_rec_no_hyper = []
    
    simulated_alphas_rec = []
    simulated_etas_rec = []
    
    simulated_alphas_no_rec = []
    simulated_etas_no_rec = []
    for eta_t in etas:
            eta = round(eta_t,4)
            print(f"On run {eta} on rho {rho}, and lam {lam}  ", end = "\r")

            alpha_max_res = alpha_max
            alpha_min_res = 0.0


            #Calculating the singular strategies when the hyperparasite is present and absent
            attractors_rec, repellers_rec = recusive_sing_strat_function(sol, 0, iter_max, resolution, beta_max, alpha_max, alpha_min_res, alpha_max_res, sigma_max, sigma, b, q, d, rho, eta, gamma, lam, c1, c2, hyper, seed)

            attractors_no_rec, repellers_no_rec = recusive_sing_strat_function(sol, 0, iter_max, resolution, beta_max, alpha_max, alpha_min_res, alpha_max_res, sigma_max, sigma, b, q, d, rho, eta, 0.0, lam, c1, c2, hyper, seed)
            
            attractors_rec_no_hyper, repellers_rec_no_hyper = recusive_sing_strat_function(sol, 0, iter_max, resolution, beta_max, alpha_max, alpha_min_res, alpha_max_res, sigma_max, sigma, b, q, d, rho, eta, gamma, lam, c1, c2, 0, seed)

            attractors_no_rec_no_hyper, repellers_no_rec_no_hyper = recusive_sing_strat_function(sol, 0, iter_max, resolution, beta_max, alpha_max, alpha_min_res, alpha_max_res, sigma_max, sigma, b, q, d, rho, eta, 0.0, lam, c1, c2, 0, seed)
            
            for val in attractors_rec:
                temp_attractors_rec.append(val)
                temp_etas_rec.append(eta)
                
            for val in attractors_no_rec:
                temp_attractors_no_rec.append(val)
                temp_etas_no_rec.append(eta)
                
            for val in repellers_rec:
                temp_repellers_rec.append(val)
                temp_etas_rec_rep.append(eta)
                
            for val in repellers_no_rec:
                temp_repellers_no_rec.append(val)
                temp_etas_no_rec_rep.append(eta)
            
            for val in attractors_rec_no_hyper:
                temp_attractors_rec_no_hyper.append(val)
                temp_etas_rec_no_hyper.append(eta)
                
            for val in attractors_no_rec_no_hyper:
                temp_attractors_no_rec_no_hyper.append(val)
                temp_etas_no_rec_no_hyper.append(eta)
                
#             if lam == lams[-1]:
#             alpha_inits = [10, 20, 40]

#             for alpha_init in alpha_inits:
# #                 sol.alpha_ad_dyn(beta_max, alpha_max, sigma_max, b, q, d, rho, eta, gamma, lam, c1, c2, hyper, seed, alpha_init, sigma_init, S_density = host_density, I_density = para_density, H_density = hyper_density)

#                 df = pd.read_csv(f"../data/alpha_evo/data_set{seed}.csv")                    
#                 seed += 1
#                 try:
#                     evo_steps = df["Evolutionary_step"].values
#                     dft = df[df["Evolutionary_step"]==evo_steps[-1]]

#                     hyper_density_check = sum(dft["Density_of_hyperparasite"].values)
#                     if hyper_density_check != 0:
#                         pops = []
#                         for val in dft["alpha_val"].values:
#                             df_pop = dft[dft["alpha_val"]==val][["Density_of_parasite", "Density_of_hyperparasite"]]
#                             pops.append((df_pop["Density_of_parasite"].iloc[0] + df_pop["Density_of_hyperparasite"].iloc[0])/(sum(dft["Density_of_parasite"].values) + sum(dft["Density_of_hyperparasite"].values)))

#                         #Calculating the average trait value
#                         alpha_weights = []
#                         for i, val in enumerate(pops):
#                             alpha_weights.append(val*dft["alpha_val"].values[i])
#                         alpha_mean = sum(alpha_weights)

# #                         print(alpha_mean)
#                         simulated_alphas_rec.append(alpha_mean)
#                         simulated_etas_rec.append(eta)
# #                         flop
#                     else:
#                         continue
#                 except IndexError:
#                     pass

#                 for alpha_init in alpha_inits:
# #                     sol.alpha_ad_dyn(beta_max, alpha_max, sigma_max, b, q, d, rho, eta, 0.0, lam, c1, c2, hyper, seed, alpha_init, sigma_init, S_density = host_density, I_density = para_density, H_density = hyper_density)

#                     df = pd.read_csv(f"../data/alpha_evo/data_set{seed}.csv")                    
#                     seed += 1
#                     try:
#                         evo_steps = df["Evolutionary_step"].values
#                         dft = df[df["Evolutionary_step"]==evo_steps[-1]]

#                         hyper_density_check = sum(dft["Density_of_hyperparasite"].values)
#                         if hyper_density_check != 0:
#                             pops = []
#                             for val in dft["alpha_val"].values:
#                                 df_pop = dft[dft["alpha_val"]==val][["Density_of_parasite", "Density_of_hyperparasite"]]
#                                 pops.append((df_pop["Density_of_parasite"].iloc[0] + df_pop["Density_of_hyperparasite"].iloc[0])/(sum(dft["Density_of_parasite"].values) + sum(dft["Density_of_hyperparasite"].values)))

#                             #Calculating the average trait value
#                             alpha_weights = []
#                             for i, val in enumerate(pops):
#                                 alpha_weights.append(val*dft["alpha_val"].values[i])
#                             alpha_mean = sum(alpha_weights)

# #                             print(alpha_mean)
#                             simulated_alphas_no_rec.append(alpha_mean)
#                             simulated_etas_no_rec.append(eta)
# #                         flop
#                         else:
#                             continue
#                     except IndexError:
#                         pass
                        
    if len(temp_repellers_rec) > 0:
        paired_etas_alphas = [[temp_etas_rec[i], temp_attractors_rec[i]] for i in range(len(temp_attractors_rec))]
    
        sorted_pairs = sorted(paired_etas_alphas, key = lambda x: x[1])
        
        alpha_set_1 = [sorted_pairs[0][1]]
        eta_set_1 = [sorted_pairs[0][0]]
        
        alpha_set_2 = []
        eta_set_2 = []
        
        flag = True
        for pair in sorted_pairs[1:]:
            if (abs(pair[1] - alpha_set_1[-1]) < 1e-1) and flag:
                alpha_set_1.append(pair[1])
                eta_set_1.append(pair[0])
            else:
                flag = False
                alpha_set_2.append(pair[1])
                eta_set_2.append(pair[0])
        
        paired_set_1 = [[eta_set_1[i], alpha_set_1[i]] for i in range(len(alpha_set_1))]
        paired_set_2 = [[eta_set_2[i], alpha_set_2[i]] for i in range(len(alpha_set_2))]
        
        sorted_set_1 = sorted(paired_set_1, key = lambda x: x[0])
        sorted_set_2 = sorted(paired_set_2, key = lambda x: x[0])
#         print(sorted_set_1)
#         print(sorted_set_2)
#         flop
        temp_repellers_with_hist = [sorted_set_1[0][1]]
        temp_etas_with_hist = [sorted_set_1[0][0]]
        for i, val in enumerate(temp_repellers_rec):
            temp_repellers_with_hist.append(val)
            temp_etas_with_hist.append(temp_etas_rec_rep[i])
            
        temp_repellers_with_hist.append(sorted_set_2[-1][1])
        temp_etas_with_hist.append(sorted_set_2[-1][0])
        
        ax1[1][lam_ind].plot([x[0] for x in paired_set_1], [x[1] for x in paired_set_1], c = "tab:blue")
        ax1[1][lam_ind].plot([x[0] for x in paired_set_2], [x[1] for x in paired_set_2], c = "tab:blue")
        ax1[1][lam_ind].scatter(temp_etas_with_hist[0],temp_repellers_with_hist[0], c = "tab:blue")
        ax1[1][lam_ind].scatter(temp_etas_with_hist[-1],temp_repellers_with_hist[-1], c = "tab:blue")
        ax1[1][lam_ind].plot(temp_etas_with_hist, temp_repellers_with_hist, c = "tab:blue", linestyle='dashed')
        
    if len(temp_repellers_no_rec) > 0:
        paired_etas_alphas = [[temp_etas_no_rec[i], temp_attractors_no_rec[i]] for i in range(len(temp_attractors_no_rec))]
    
        sorted_pairs = sorted(paired_etas_alphas, key = lambda x: x[1])
        
        alpha_set_1 = [sorted_pairs[0][1]]
        eta_set_1 = [sorted_pairs[0][0]]
        
        alpha_set_2 = []
        eta_set_2 = []
        
        flag = True
        for pair in sorted_pairs[1:]:
            if (abs(pair[1] - alpha_set_1[-1]) < 1e-1) and flag:
                alpha_set_1.append(pair[1])
                eta_set_1.append(pair[0])
            else:
                flag = False
                alpha_set_2.append(pair[1])
                eta_set_2.append(pair[0])
                
        paired_set_1 = [[eta_set_1[i], alpha_set_1[i]] for i in range(len(alpha_set_1))]
        paired_set_2 = [[eta_set_2[i], alpha_set_2[i]] for i in range(len(alpha_set_2))]
        
        sorted_set_1 = sorted(paired_set_1, key = lambda x: x[0])
        sorted_set_2 = sorted(paired_set_2, key = lambda x: x[0])
#         print(sorted_set_1)
#         print(sorted_set_2)
#         flop
        temp_repellers_with_hist = [sorted_set_1[0][1]]
        temp_etas_with_hist = [sorted_set_1[0][0]]
        for i, val in enumerate(temp_repellers_rec):
            temp_repellers_with_hist.append(val)
            temp_etas_with_hist.append(temp_etas_rec_rep[i])
            
        temp_repellers_with_hist.append(sorted_set_2[-1][1])
        temp_etas_with_hist.append(sorted_set_2[-1][0])
        
        ax1[0][lam_ind].plot([x[0] for x in paired_set_1], [x[1] for x in paired_set_1], c = "tab:blue")
        ax1[0][lam_ind].plot([x[0] for x in paired_set_2], [x[1] for x in paired_set_2], c = "tab:blue")
        ax1[0][lam_ind].scatter(temp_etas_with_hist[0],temp_repellers_with_hist[0], c = "tab:blue")
        ax1[0][lam_ind].scatter(temp_etas_with_hist[-1],temp_repellers_with_hist[-1], c = "tab:blue")
        ax1[0][lam_ind].plot(temp_etas_with_hist, temp_repellers_with_hist, c = "tab:blue", linestyle='dashed')
        
    ax1[0][lam_ind].plot(temp_etas_no_rec_no_hyper,temp_attractors_no_rec_no_hyper, c = "k")
    ax1[1][lam_ind].plot(temp_etas_rec_no_hyper,temp_attractors_rec_no_hyper, c = "k")
#     if lam == lams[-1]:
#     ax1[0][lam_ind].scatter(simulated_etas_no_rec, simulated_alphas_no_rec)
#     ax1[1][lam_ind].scatter(simulated_etas_rec, simulated_alphas_rec)
#     else:
#     ax1[0][lam_ind].scatter(temp_etas_no_rec,temp_attractors_no_rec)
#     ax1[1][lam_ind].scatter(temp_etas_rec,temp_attractors_rec)
    
#     ax1[0][lam_ind].scatter(temp_etas_no_rec_rep,temp_repellers_no_rec)
#     ax1[1][lam_ind].scatter(temp_etas_rec_rep,temp_repellers_rec)
        
    ax1[0][lam_ind].set_ylim([0,2.5])
    ax1[1][lam_ind].set_ylim([0,2.5])
    ax1[0][lam_ind].text(0.03,1.01,texts[0][lam_ind], transform=ax1[0][lam_ind].transAxes, fontsize = 28)
    ax1[1][lam_ind].text(0.03,1.01,texts[1][lam_ind], transform=ax1[1][lam_ind].transAxes, fontsize = 28)
    ax1[0][lam_ind].set_title(fr"{titles[lam_ind]}($\lambda$ = {lam})", fontsize = 28)
    ax1[0][lam_ind].tick_params(axis='both', which='major', labelsize=34)
    ax1[1][lam_ind].tick_params(axis='both', which='major', labelsize=34)
    
ax1[1][1].set_xlabel(r"Hyperparasite Transmission Modifier, $\eta$", fontsize = 28)
ax1[0][0].set_ylabel(r"Intrinsic Virulence, $\alpha^*$, no recovery", fontsize = 28)
ax1[1][0].set_ylabel(r"Intrinsic Virulence, $\alpha^*$, with recovery", fontsize = 28)
plt.savefig("../supplementary_figures/sandhu_comparison_no_rec_vs_rec.png", bbox_inches = "tight")
plt.savefig("../supplementary_figures/sandhu_comparison_no_rec_vs_rec.pdf", bbox_inches = "tight")
plt.close()
```
