## Supplementary material for "Hyperparasitism and the evolution of parasite virulence": Code_base: validate_sandhu_results.py

```
"""
Depricated file, held here for completeness.
"""

#Importing libraries needed to perform the analysis
import pandas as pd
import numpy as np
from math import exp
import runner
import matplotlib.pyplot as plt


#Parameters used in the system
rho = 1.0

lams = [0.5, 1.0, 2.0]

b = 2.0
q = 0.1
d = 0.5
gamma = 0.0

seed = 100

sol = runner.PySolver()

plotting_alphas = []
plotting_evos = []
alpha_init = 20
sigma_init = 100

beta_max = 5.0
alpha_max = 5.0
sigma_max = 4.0

hyper = 1.0
c1 = 0.75
c2 = 2.25

#We initialise the system with the hyperparasites absent to find the natural ESS of the parasite
sol.alpha_ad_dyn(beta_max, alpha_max, sigma_max, b, q, d, rho, 0.0, gamma, 0.0, c1, c2, 0.0, seed, alpha_init, sigma_init, H_density = 0.0)
df = pd.read_csv(f"../data/alpha_evo/data_set{seed}.csv")

evo_steps = df["Evolutionary_step"].values
dft = df[df["Evolutionary_step"]==evo_steps[-1]]

pops = []
for val in dft["Trait_index_1"].values:
    df_pop = dft[dft["Trait_index_1"]==val][["Density_of_parasite"]]
    pops.append((df_pop["Density_of_parasite"].iloc[0])/(sum(dft["Density_of_parasite"].values)))
    
#Calculating the average trait value
alpha_weights = []
for i, val in enumerate(pops):
    alpha_weights.append(val*dft["Trait_index_1"].values[i])
alpha_mean = sum(alpha_weights)

#Rounding this to the closest integer to use as our initial conditions
alpha_init = round(alpha_mean)

alpha_mean_base = alpha_init*alpha_max/100

host_density = dft["Density_of_hyperparasite"].iloc[0]
para_density = sum(dft["Density_of_parasite"].values)
hyper_density = 0.1

etas = [2.0*i/100 for i in range(101)]

#We create two base figures, one for the evolutionary consequences
fig1 = plt.figure(figsize = (40, 22))
gs1 = fig1.add_gridspec(1,len(lams))
ax1 = []

#Flavour text to add to the readibility of the plots
texts = [["(A)", "(B)", "(C)"],["(D)", "(E)", "(F)"], ["(G)","(H)","(I)"]]

titles = ["Hypovirulence ", "No effect on virulence ", "Hypervirulence "]

#Structuring the figures so we can access subplots
for i in range(1):
    temp_ax = []
    for j in range(len(lams)):
        temp_ax.append(fig1.add_subplot(gs1[i,j]))
    ax1.append(temp_ax)
    
titles = ["Hypovirulence ", "No effect on virulence ", "Hypervirulence "]

for ind, lam in enumerate(lams):
    simulated_alphas = []
    for eta in etas:
#         print(eta, lam)
        sol.alpha_ad_dyn(beta_max, alpha_max, sigma_max, b, q, d, rho, eta, gamma, lam, c1, c2, hyper, seed, alpha_init, sigma_init, S_density = host_density, I_density = para_density, H_density = hyper_density)
        df = pd.read_csv(f"../data/alpha_evo/data_set{seed}.csv")

        seed += 1
        
        evo_steps = df["Evolutionary_step"].values
        
        dft = df[df["Evolutionary_step"]==evo_steps[-1]]
        total_pop = sum(sum(dft[["Density_of_parasite","Density_of_hyperparasite"]].values))
#         print(total_pop)
        pops = []
        for val in dft["alpha_val"].values:
            df_pop = dft[dft["alpha_val"]==val][["Density_of_parasite","Density_of_hyperparasite"]]
#             print(df_pop.values)
#             print(sum(df_pop.values[0]))
            pops.append(sum(df_pop.values[0])/total_pop)
            
#         print(pops)
        alpha_weights = []
        for i, val in enumerate(pops):
            alpha_weights.append(val*dft["alpha_val"].values[i])
#         print(alpha_weights)
        alpha_mean = sum(alpha_weights)
        simulated_alphas.append(alpha_mean)
#         print("alpha_mean")
#         print(alpha_mean)
#         print(dft)
#         flop
    ax1[0][ind].plot(etas, simulated_alphas)
    ax1[0][ind].plot(etas, [alpha_mean_base for eta in etas])
    ax1[0][ind].tick_params(axis='both', which='major', labelsize=34)
    ax1[0][ind].set_ylim([0,alpha_max])
    ax1[0][ind].set_title(fr"{titles[ind]}($\lambda$ = {lam})", fontsize = 34)

ax1[0][1].set_xlabel(r"Hyperparasite Infectivity Modifier, $\eta$", fontsize = 34)    
ax1[0][0].set_ylabel(r"Evolved Virulence, $\alpha$", fontsize = 34)
plt.savefig("../supplementary_figures/sandhu_validation.png", bbox_inches = "tight")
plt.close()
```
