## Supplementary material for "Hyperparasitism and the evolution of parasite virulence": Code_base: visualise_branching_matrices.py

```
""" This file creates a grouped image of the heatmaps showing the 
branching produced by the file linear_heatmap_res_scalar.py. This file
assumes that 3 csv files are stored in the data folder with the names 
branching_mat_0.csv, branching_mat_1.csv and branching_mat_2.csv. This
file then produces a .png file and .pdf file of these images, adding
various labels for clarity. This file requires the pandas, numpy and
matplotlib libraries. This file is used to create Fig. 4.
"""

#Importing needed libraries
import pandas as pd
import numpy as np
import matplotlib.pyplot as plt

#Importing the rho values used when creating the dataset
rhos = [0.25, 0.5, 0.75]

#Panel labels we will use to distinguish between subfigures
panel_labels = ["(A)", "(B)", "(C)"]

#We create the figure to be a large size as our data sets
#are reasonably big. We use gridspec to specify the 
#layout of the figures

resolution = 50

fig = plt.figure(figsize = (2.5*resolution, 2.5*resolution))
gs = fig.add_gridspec(1,3)
ax = []
for i in range(1):
    temp_ax = []
    for j in range(3):
        temp_ax.append(fig.add_subplot(gs[i,j]))
    ax.append(temp_ax)
    
#We now loop through the files to read in the datasets and
#extract the values from them
for i in range(3):
    rho = rhos[i]
    
    #Reading in the stored data
    df = pd.read_csv(f"../data/branching_mat_{i}.csv")
    
    etas = df["eta_val"].values
    lams = [float(val) for val in df.columns[1:]]
    value_matrix = df[df.columns[1:]].values
    
    #Plotting the heatmap
    ax[0][i].imshow(value_matrix, origin = "lower", cmap = "Set3")

    #Here we create the equidistant ticks used when plotting
    ticks_temp = []
    for k in range(len(value_matrix)):
        ticks_temp.append(k)
    
    eta_ticks = [0.0, 0.2, 0.4, 0.6, 0.8, 1.0]
    lam_ticks = [0.0, 1.0, 2.0, 3.0, 4.0, 5.0]
    ticks = [i for i in ticks_temp if lams[i] in lam_ticks]
    
    #Adding our ticks to the various subplots
    ax[0][i].set_yticks(ticks = ticks, labels = eta_ticks, fontsize = 2*resolution)
    ax[0][i].set_xticks(ticks = ticks, labels = lam_ticks, fontsize = 2*resolution, rotation=45)
    ax[0][i].set_title(fr"Probability of cotransmission, $\rho$ = {rho}", fontsize = 2*resolution)
    
    #Adding the panel label
    ax[0][i].text(0.05,0.9,panel_labels[i], transform=ax[0][i].transAxes, fontsize = 2*resolution)    
    
#Adding the axis labels that we want
ax[0][0].set_ylabel(r"Hyperparasite transmission modifier, $\eta$", fontsize = 2*resolution)
ax[0][1].set_xlabel(r"Hyperparasite virulence modifier, $\lambda$", fontsize = 2*resolution)

#Saving the file as both a pdf and a png
plt.savefig(f"../supplementary_figures/branching_heatmaps.pdf", bbox_inches = "tight") 
plt.savefig(f"../supplementary_figures/branching_heatmaps.png", bbox_inches = "tight")
plt.close()
```
