## Supplementary material for "Hyperparasitism and the evolution of parasite virulence": Code_base: wrapper.cpp

```
#include <iostream>
#include <cstdlib>
#include <ctime>
#include <fstream>
#include <time.h>
#include <cmath>
#include <unistd.h>
#include <iomanip> 
#include <string>
#include <sstream>
#include <vector>
#include <algorithm>
#include <wrapper.h>
using namespace std;

float TINY = 1e-3;
float EPS = 1e-6;

int trait_space_length = 101;

namespace solvers{
    
    Quick_solver::Quick_solver() {};
    Quick_solver::Quick_solver(float beta_max, float alpha_max, float sigma_max, float b, float q, float d, float rho, float eta, float gamma, float lambda, float c1, float c2, float hyper, int seed){
        this->beta_max = beta_max;
        this->alpha_max = alpha_max;
        this->sigma_max = sigma_max;
        this->b = b;
        this->q = q;
        this->d = d;
        this->rho = rho;
        this->eta = eta;
        this->gamma = gamma;
        this->lambda = lambda;
        this->c1 = c1;
        this->c2 = c2;
        this->hyper = hyper;
        this->seed = seed;

    };
    
    
    Quick_solver::~Quick_solver() {};

    
    float Quick_solver::fastmax(float a, float b){
        if (a > b){
            return a;
        }
        else{
            return b;
        }
    }

    float Quick_solver::fastmin(float a, float b){
        if (a < b){
            return a;
        }
        else{
            return b;
        }
    }

            // float fastmin(float a, float b){
            //     return (a<b)?a:b;
            // }

    int Quick_solver::dynamics(float* dydt, float* y, int num_strains, int* alpha_inds, int* sigma_inds, float b, float q, float d, float* beta, float* alpha, float eta, float gamma, float* sigma, float lambda, float rho){
        
        int i;
        float Isum = 0.0, Hsum = 0.0, N = 0.0;
        // float* I = new float[num_alpha];
        // float* H = new float[num_alpha];
        // float* Idot = new float[num_alpha];
        // float* Hdot = new float[num_alpha];
        float S, I[num_strains], H[num_strains];
        float Sdot, Idot[num_strains], Hdot[num_strains];
        float infection_force_para[num_strains], para_infection_force = 0.0;
        float infection_force_hyper[num_strains], hyper_infection_force = 0.0;
        
        // ofstream hdot_logs;
        S = y[0];

        for(i=0;i<num_strains;i++){
            I[i] = y[i + 1];
            H[i] = y[i + 1 + num_strains];
            Isum = Isum + I[i];
            Hsum = Hsum + H[i];
            infection_force_para[i] = beta[alpha_inds[i]*trait_space_length + sigma_inds[i]]*I[i];
            para_infection_force = para_infection_force + infection_force_para[i];
            infection_force_hyper[i] = eta*beta[alpha_inds[i]*trait_space_length + sigma_inds[i]]*H[i];
            hyper_infection_force = hyper_infection_force + infection_force_hyper[i];
        }

        N = Isum + Hsum + S;

        Sdot =  (b - q*N)*N - (para_infection_force + hyper_infection_force + d)*S + gamma*(Isum + Hsum);
        
        for(i = 0;i<num_strains;i++){
            Idot[i] = (beta[alpha_inds[i]*trait_space_length + sigma_inds[i]]*S - sigma[sigma_inds[i]]*Hsum - (d + alpha[alpha_inds[i]] + gamma))*I[i] + (1 - rho)*eta*beta[alpha_inds[i]*trait_space_length + sigma_inds[i]]*S*H[i];

            Hdot[i] = (rho*eta*beta[alpha_inds[i]*trait_space_length + sigma_inds[i]]*S - (d + lambda*alpha[alpha_inds[i]] + gamma))*H[i] + sigma[sigma_inds[i]]*I[i]*Hsum;

        }

        dydt[0] = Sdot;

        // if (abs(Sdot) > 1e-4){
        //     dydt[0] = Sdot;
        // }
        // else{
        //     dydt[0] = 0.0;
        // }

        for(i = 0;i<num_strains;i++){

            dydt[i + 1] = Idot[i];

            // if (abs(Idot[i]) > 1e-4){
            //     dydt[i + 1] = Idot[i];
            // }
            // else{
            //     dydt[i+1] = 0.0;
            // }

            dydt[i + 1 + num_strains] = Hdot[i];

            // if (abs(Hdot[i]) > 1e-4){
            //     dydt[i + 1 + num_strains] = Hdot[i];
            // }
            // else{
            //     dydt[i + 1 + num_strains] = 0.0;
            // }
        }

        // delete I, H, Idot, Hdot;
        return 0;
    }

    int Quick_solver::RK45(float* y_out, float* y, float* y_err, int num_strains, int* alpha_inds, int* sigma_inds, float b, float q, float d, float* beta, float* alpha, float eta, float gamma, float* sigma, float lambda, float rho, float* h){

        float* y_1 = new float[2*num_strains + 1];
        float* y_2 = new float[2*num_strains + 1];
        float* y_3 = new float[2*num_strains + 1];
        float* y_4 = new float[2*num_strains + 1];
        // float y_1[2*num_strains+1], y_2[2*num_strains+1], y_3[2*num_strains+1], y_4[2*num_strains+1];
        int i;
        
        for (i = 0; i<(2*(num_strains) + 1); i++){
            y_1[i] = 0.0;
            y_2[i] = 0.0;
            y_3[i] = 0.0;
            y_4[i] = 0.0;
        }

        Quick_solver::dynamics(y_1, y, num_strains, alpha_inds, sigma_inds, b, q, d, beta, alpha, eta, gamma, sigma, lambda, rho);

        for (i = 0; i<(2*(num_strains) + 1); i++){
            y_out[i] = y[i] + (y_1[i]*h[0]);
        }

        for (i = 0; i<(2*(num_strains) + 1); i++){
            if(isnan(y_1[i])){
                std::cout << "Ind " << i << " has created a NaN after 1 step\n";
            }
        }

        Quick_solver::dynamics(y_2, y_out, num_strains, alpha_inds, sigma_inds, b, q, d, beta, alpha, eta, gamma, sigma, lambda, rho);

        for (i = 0; i<(2*(num_strains) + 1); i++){
            y_out[i] = y[i] + (y_2[i]*h[0])/2;
        }

        for (i = 0; i<(2*(num_strains) + 1); i++){
            if(isnan(y_2[i])){
                std::cout << "Ind " << i << " has created a NaN after 2 steps\n";
            }
        }
        Quick_solver::dynamics(y_3, y_out, num_strains, alpha_inds, sigma_inds, b, q, d, beta, alpha, eta, gamma, sigma, lambda, rho);

        for (i = 0; i<(2*(num_strains) + 1); i++){
            y_out[i] = y[i] + (y_3[i]*h[0])/2;
        }
        
        for (i = 0; i<(2*(num_strains) + 1); i++){
            if(isnan(y_3[i])){
                std::cout << "Ind " << i << " has created a NaN after 3 steps\n";
            }
        }

        Quick_solver::dynamics(y_4, y_out, num_strains, alpha_inds, sigma_inds, b, q, d, beta, alpha, eta, gamma, sigma, lambda, rho);

        for (i = 0; i<(2*(num_strains) + 1); i++){
            y_out[i] = y[i] + h[0]*((y_1[i] + 2*(y_2[i] + y_3[i]) + y_4[i])/6);
            y_err[i]= h[0]*((y_1[i] + 2*(y_2[i] + y_3[i]) + y_4[i])/6);
        }

        for (i = 0; i<(2*(num_strains) + 1); i++){
            if(isnan(y_4[i])){
                std::cout << "Ind " << i << " has created a NaN after 4 steps\n";
            }

            if(isnan(y_err[i])){
                std::cout << "error calculater " << i << " has returned a NaN\n";
            }
        }
        delete y_1;
        delete y_2;
        delete y_3;
        delete y_4;
        return 0;
    }


    int Quick_solver::perform_RK45_step(float* y, float* t, int num_strains, int* alpha_inds, int* sigma_inds, float b, float q, float d, float* beta, float* alpha, float eta, float gamma, float* sigma, float lambda, float rho, float* h, float* discrepancy, int* counter, int* num_plus, int* num_down){

        // float discrepancy[1];
        // float* y_out = new float[2*num_alpha+1];
        float y_out[2*num_strains+1], y_err[2*num_strains + 1];
        bool flag = true;
        int i;
        // float htemp,errmax;

        // h[0] = 0.01;
        
        // counter = 0;
        // dynamics(dydt,y, num_strains, alpha_inds, sigma_inds, b, q, d, beta, alpha, eta, gamma, sigma, lambda, rho);
        // for (i = 0; i<(2*num_strains + 1); i++){
        //     y_scale[i] = y[i] + dydt[i]*h[0] + TINY;
        // }
        // while (t[0] < tmax){
        //     flag = true;
            while ((h[0] >= 1e-10) && (h[0] <= 1e8) && (flag))
            {  
                for (i=0;i<(2*(num_strains) + 1);i++){
                    y_out[i] = 0.0;
                }

                discrepancy[0] = 0.0;

                Quick_solver::RK45(y_out,y,y_err,num_strains,alpha_inds, sigma_inds, b,q,d,beta,alpha,eta,gamma,sigma,lambda,rho,h);
                // errmax= 0.0;
                float ysum = 0.0;
                for (i=0;i<(2*num_strains+1);i++){
                    ysum += y[i];
                }
                for (i=0;i<(2*num_strains+1);i++){
                    // discrepancy[0] = discrepancy[0] + abs(y_out[i] - y[i]);
                    discrepancy[0] = fastmax(discrepancy[0],abs((y_out[i] - y[i])/(y[i] + EPS)));            
                    // errmax= fastmax(errmax,abs(y_err[i]/y_scale[i]));
                }

                // discrepancy[0] = fastmax(0.0000000001, discrepancy[0]);
                // errmax/= TINY;
                // discrepancy[0] = errmax;
                // if(errmax<=1.0){ 
                    // std::cout << "Exit condition is working" << endl;
                    // flag = false;
                    // break;
                // }
                // htemp= 0.9*(*h)*pow(errmax,-0.25);
                // h[0]= (*h>=0.0 ? fmax(htemp,0.1*(*h)) : min(htemp,0.1*(*h)));

                // discrepancy[0] = discrepancy[0]/TINY;
                // if ((discrepancy[0] <=0.9) && (discrepancy[0] <= 1.1)){
                //     // std::cout << "discrepancy is " << discrepancy[0] << endl;
                //     flag = false;
                //     break;
                // }
                // else{
                //     std::cout << "discrepancy is " << discrepancy[0] << endl;
                //     // h[0] = h[0]/(discrepancy[0]*discrepancy[0]);
                //     // h[0] = h[0]/discrepancy[0];
                //     htemp= 0.9*(*h)*pow(discrepancy[0],-0.15);
                //     h[0]= (*h>=0.0 ? max(htemp,0.1*(*h)) : min(htemp,0.1*(*h)));
                //     // std::cout << "step size is " << h[0] << endl;
                // }
                // h[0]
                // if((discrepancy[0] >= 1e-2)){
                //     h[0] = h[0]*0.02;
                // } else if(discrepancy[0] <= 1e-5){
                //     h[0] = h[0]*1e4;   
                // }

                if (discrepancy[0]/(1e-2) < 1){
                    break;
                }
                discrepancy[0] = fastmax(0.0000000001, discrepancy[0]);

                h[0] = h[0]*pow(discrepancy[0]/(1e-2), -0.2);
                counter[0]++;
                // if (discrepancy[0] >= 1e-4){
                //     h[0] = h[0]*0.02*(1e-2/discrepancy[0]);
                //     h[0] = h[0]/5;
                //     counter[0]++;   
                //     num_down[0]++;
                // }
                // else if(discrepancy[0] <= 1e-6){
                //     h[0] = h[0]*1e4*(1e-5/discrepancy[0]); 
                //     h[0] = h[0]*10; 
                //     counter[0]++;
                //     num_plus[0]++;
                // }   
                // else{
                //     flag = false;
                // }
            

                if(counter[0] >= 10){
                    flag = false;
                }
                // std::cout << "Number of steps tried is " << counter << endl;
            }
            for (i = 0; i<(2*(num_strains) + 1); i++){
                // if (y_out[i] > TINY){
                //     y[i] = y_out[i];
                // }
                // else{
                //     y[i] = 0.0;
                // }
                y[i] = y_out[i];
            }
            t[0] = t[0] + h[0];
            // h[0] = fmin(10*h[0],10);
            // std::cout << "step size is " << h[0] << endl;
            // std::cout << "time step is " << t[0] << endl;
            // sleep(1);
        // }
        // std::cout << "Had to do " << counter << " number of steps " << endl;
        return 0;
    }

    int Quick_solver::rkck(float *y, float *dydt, float *yout, float *yerr, float h, int num_strains, int* alpha_inds, int* sigma_inds, float b, float q, float d, float* beta, float* alpha, float eta, float gamma, float* sigma, float lambda, float rho)
    {
        int i;
        // float y_1[2*num_strains + 1], y_2[2*num_strains + 1], y_3[2*num_strains + 1], y_4[2*num_strains + 1], y_5[2*num_strains + 1];
        float* y_1 = new float[2*num_strains + 1];
        float* y_2 = new float[2*num_strains + 1];
        float* y_3 = new float[2*num_strains + 1];
        float* y_4 = new float[2*num_strains + 1];
        float* y_5 = new float[2*num_strains + 1];
        float* y_temp = new float[2*num_strains + 1];
        // float y_temp[2*num_strains + 1];
        static float b21=0.2,b31=3.0/40.0,b32=9.0/40.0,b41=0.3,b42=-0.9,b43=1.2,
        b51=-11.0/54.0,b52=2.5,b53=-70.0/27.0,b54=35.0/27.0,b61=1631.0/55296,
        b62=175.0/512.0,b63=575.0/13824.0,b64=44275.0/110592,b65=253.0/4096.0,
        c1=37.0/378.0,c3=250.0/621.0,c4=125.0/594.0,c6=512.0/1771.0,dc5=-277.00/14336;
        float dc1=c1-2825.0/27648.0,dc3=c3-18575.0/48384.0,dc4=c4-13525.0/55296.0,
        dc6=c6-0.25;
        
        for(i=0;i<(2*num_strains + 1);i++){
            y_temp[i] = y[i] + b21*h*dydt[i];
        }
        Quick_solver::dynamics(y_1, y_temp, num_strains, alpha_inds, sigma_inds, b, q, d, beta, alpha, eta, gamma, sigma, lambda, rho);
        // dynamic(y_temp,Itemp,Htemp,u,v,E,b,y_2,Ik2,Hk2,(2*num_strains + 1),np,host_ind,par_ind);

        // for (i = 0; i<(2*(num_strains) + 1); i++){
        //     if(isnan(y_1[i])){
        //         std::cout << "Ind " << i << " has created a NaN after 1 step\n";
        //     }
        // }

        for(i=0;i<(2*num_strains + 1);i++){
            y_temp[i] = y[i]+h*(b31*dydt[i]+b32*y_1[i]);
        }
        Quick_solver::dynamics(y_2, y_temp, num_strains, alpha_inds, sigma_inds, b, q, d, beta, alpha, eta, gamma, sigma, lambda, rho);

        // for (i = 0; i<(2*(num_strains) + 1); i++){
        //     if(isnan(y_2[i])){
        //         std::cout << "Ind " << i << " has created a NaN after 2 steps\n";
        //     }
        // }

        for(i=0;i<(2*num_strains + 1);i++){
            y_temp[i]= y[i]+h*(b41*dydt[i]+b42*y_1[i]+b43*y_2[i]);
        }

        Quick_solver::dynamics(y_3, y_temp, num_strains, alpha_inds, sigma_inds, b, q, d, beta, alpha, eta, gamma, sigma, lambda, rho);
        
        // for (i = 0; i<(2*(num_strains) + 1); i++){
        //     if(isnan(y_3[i])){
        //         std::cout << "Ind " << i << " has created a NaN after 3 steps\n";
        //     }
        // }
        
        for(i=0;i<(2*num_strains + 1);i++){
            y_temp[i]= y[i]+h*(b51*dydt[i]+b52*y_2[i]+b53*y_2[i]+b54*y_3[i]);
        }

        Quick_solver::dynamics(y_4, y_temp, num_strains, alpha_inds, sigma_inds, b, q, d, beta, alpha, eta, gamma, sigma, lambda, rho);
        
        // for (i = 0; i<(2*(num_strains) + 1); i++){
        //     if(isnan(y_4[i])){
        //         std::cout << "Ind " << i << " has created a NaN after 4 steps\n";
        //     }
        // }
        
        for(i=0;i<(2*num_strains + 1);i++){
            y_temp[i]= y[i]+h*(b61*dydt[i]+b62*y_1[i]+b63*y_2[i]+b64*y_3[i]+b65*y_4[i]);
        }

        Quick_solver::dynamics(y_5, y_temp, num_strains, alpha_inds, sigma_inds, b, q, d, beta, alpha, eta, gamma, sigma, lambda, rho);
        
        // for (i = 0; i<(2*(num_strains) + 1); i++){
        //     if(isnan(y_5[i])){
        //         std::cout << "Ind " << i << " has created a NaN after 5 steps\n";
        //     }
        // }
        
        for(i=0;i<(2*num_strains + 1);i++){
            yout[i]= y[i]+h*(c1*dydt[i]+c3*y_2[i]+c4*y_3[i]+c6*y_5[i]);
            yerr[i]= h*(dc1*dydt[i]+dc3*y_2[i]+dc4*y_3[i]+dc5*y_4[i]+dc6*y_5[i]);
        }

        // for (i = 0; i<(2*(num_strains) + 1); i++){

        //     if(isnan(yerr[i])){
        //         std::cout << "error calculater " << i << " has returned a NaN\n";
        //     }
        // }
        delete y_1;
        delete y_2;
        delete y_3;
        delete y_4;
        delete y_5;
        delete y_temp;
        return 0;
    }

    int Quick_solver::rkqs(float *y, float *dydt, float *h,float *hnext,float *yscale, int num_strains, int* alpha_inds, int* sigma_inds, float b, float q, float d, float* beta, float* alpha, float eta, float gamma, float* sigma, float lambda, float rho, int* counter, int* num_up, int* num_down, float* discrepancy, int* int_tracker, float* t)
    {
        // float* y_temp = new float[2*num_strains + 1];
        // float* yerr = new float[2*num_strains + 1];
        float y_temp[2*num_strains + 1], yerr[2*num_strains + 1];
        float htemp,errmax;
        int i;//, counter_internal;
        
        htemp= *h;
        
        Quick_solver::rkck(y, dydt, y_temp, yerr, *h, num_strains, alpha_inds, sigma_inds, b, q, d, beta, alpha, eta, gamma, sigma, lambda, rho);
        *hnext= *h;
        
        // for(i=0;i<(2*num_strains + 1);i++){
        //     y_sum += y[i];
        // }
        // counter_internal = 0;
        for(;;)
        {
            Quick_solver::rkck(y, dydt, y_temp, yerr, *h, num_strains, alpha_inds, sigma_inds, b, q, d, beta, alpha, eta, gamma, sigma, lambda, rho);
            errmax= 0.0;
            // y_sum = 0.0;
            // for(i=0;i<(2*num_strains + 1);i++){
            //     y_sum += y_temp[i];
            // }
            for(i=0;i<(2*num_strains + 1);i++){
                if (errmax < abs(yerr[i]/yscale[i])){
                    errmax = abs(yerr[i]/yscale[i]);
                    int_tracker[0] = i;
                }
                // errmax= Quick_solver::fastmax(errmax,fabs(yerr[i]/yscale[i]));
                // errmax= Quick_solver::fastmax(errmax,fabs(yerr[i]/y_sum));
            }
            errmax/= 1e-4;
            discrepancy[0] = errmax;
            if(errmax<=1.0){ 
                break;
            }
            // if (counter_internal > 0){
            //     std::cout << counter_internal << "," << t[0] << "," << *h << "," << htemp << "," << errmax << "\n";
            //     std::cout << "Eco values input are: \n";
            //     for (i =0; i<(2*num_strains + 1); i++){
            //         std::cout << y[i] << "\n";
            //     } 
            //     std::cout << "Eco values output are: \n";
            //     for (i =0; i<(2*num_strains + 1); i++){
            //         std::cout << y_temp[i] << "\n";
            //     } 
            //     std::cout << "Input gradients are : \n";
            //     for (i =0; i<(2*num_strains + 1); i++){
            //         std::cout << dydt[i] << "\n";
            //     } 
            // }
            // if (counter_internal == 10){
            //     // std::cout << "Breaking cus counter\n";
            //     // std::cout << *h << "," << htemp << "\n";
            //     // sleep(5);
            //     break;
            //     // sleep(1);
            // }
            // counter_internal++;
            // htemp = *h * (errmax)^(.2)
            htemp= 0.9*(*h)*pow(errmax,-0.25);
            // if (htemp < *h){
            //     num_down[0]++;
            // }
            // else{
            //     num_up[0]++;
            // }
            // *h= 0.9*(*h)*pow(errmax,-0.25);
            *h= (*h>=0.0 ? Quick_solver::fastmax(htemp,0.1*(*h)) : Quick_solver::fastmin(htemp,0.1*(*h)));
            counter[0]++;
            // *h = fastmax(*h, 0.001);
            // break;
        }
        // *hnext= 0.9*(*h)*pow(errmax,-0.2)*(errmax > 1.89E-4) + 50.0*(*h)*(errmax <= 1.89E-4);
        if(errmax > 1.89E-4) {
            *hnext= 0.9*(*h)*pow(errmax,-0.2);
            // num_down[0]++;
        } 
        else {
            *hnext= 5.0*(*h);
            // num_up[0]++;
        }
        
        // *hnext = 5.0*(*h);
        for (i = 0; i<(2*(num_strains) + 1); i++){
            // y[i] = y_temp[i]*((y_temp[i] > y[i]) || (y[i] > 1e-2));
            // if ((y_temp[i] > y[i]) || (y[i] > 1e-2)){
            y[i] = y_temp[i];
            // }
            // else{
            //     y[i] = 0.0;
            // }
        }

        // delete y_temp;
        // delete yerr;
        return 0;
    }

    int Quick_solver::ad_dyn(float beta_max, float alpha_max, float sigma_max, float b, float q, float d, float rho, float eta, float gamma, float lambda, float c1, float c2, float hyper, int seed, int alpha_init, int sigma_init){
        time_t start,end;
        time (&start);

        srand(seed);
        
        int tmax = 1000, evo_steps = 1000, i, j;
        float beta[trait_space_length*trait_space_length], alpha[trait_space_length], sigma[trait_space_length];
        float t[1], tcheck[1], Hsum;
        int num_strains = 1, alpha_inds[trait_space_length], ind,num_strains_2;
        int sigma_inds[trait_space_length], num_strains_cleaned;
        float* y = new float[2*trait_space_length+1];
        float* y_check = new float[2*trait_space_length + 1];
        float* dydt = new float[2*trait_space_length + 1];
        float* yScale = new float[2*trait_space_length + 1];
        
        float* y_max = new float[2*trait_space_length + 1];
        float* y_min = new float[2*trait_space_length + 1];
        float* y_max_next = new float[2*trait_space_length + 1];
        float* y_min_next = new float[2*trait_space_length + 1];
        float discrep_check;
        int alpha_inds_cleaned[trait_space_length],counter=0;
        int sigma_inds_cleaned[trait_space_length];
        float I_temp[trait_space_length], H_temp[trait_space_length],total_density[trait_space_length];
        float cum_density[trait_space_length], cum_props[trait_space_length];
        int alpha_inds_2[trait_space_length];
        int sigma_inds_2[trait_space_length], int_tracker[1];
        float* y_temp = new float[2*trait_space_length+1];
        
        float h[1], hnext[1], discrepancy[1];

        int num_poss_outputs = 40000;
        int output_alpha_inds[num_poss_outputs], output_sigma_inds[num_poss_outputs], output_counter = 0, evo_step_tracker[num_poss_outputs];
        float output_parasite_density[num_poss_outputs], output_hyperparasite_density[num_poss_outputs], output_host_density[num_poss_outputs];
        bool output_hyperparasite_truths[num_poss_outputs];
        
        ind = 0;
        ofstream myfile, tracker_file;
        

        for(i=0;i<trait_space_length;i++){
            
            alpha[i] = (alpha_max*(i))/(trait_space_length - 1);
            sigma[i] = (sigma_max*(i))/(trait_space_length - 1);
            
        }

        for(i=0;i<trait_space_length;i++){
            for(j=0;j<trait_space_length;j++){
                
                if (c2 == 0.0){
                    beta[i*trait_space_length + j] = beta_max*(sqrt(alpha[i]/alpha_max)*((1-c1) + c1*(sigma[j]/sigma_max)));
                }
                else{
                    beta[i*trait_space_length+j] = fastmax(0,beta_max*(sqrt(alpha[i]/alpha_max)*((1-c1) + c1*(1 - exp((sigma[j]*c2)/sigma_max))/(1 - exp(c2)))));
                }
            }
        }

        ofstream beta_values,sigma_values,alpha_values;
        beta_values.open("../data/beta_vals.csv");
        sigma_values.open("../data/sigma_vals.csv");
        alpha_values.open("../data/alpha_vals.csv");

        for (i = 0; i<trait_space_length;i++){
            beta_values << i << ",";
            alpha_values << i << ",";
            sigma_values << i << ",";
        }
        beta_values << "\n";
        alpha_values << "\n";
        sigma_values << "\n";

        

        for(i=0;i<trait_space_length;i++){
            alpha_values << alpha[i] << ",";
            sigma_values << sigma[i] << ",";
            for(j=0;j<trait_space_length;j++){
                beta_values << beta[i*trait_space_length+j] << ",";
            }
            beta_values << "\n";
        }

        alpha_values << "\n";
        sigma_values << "\n";

        for (i = 0; i<trait_space_length;i++){
            sigma_values << ((1-c1) + c1*(1 - exp((sigma[i]*c2)/sigma_max))/(1 - exp(c2))) << ",";
            alpha_values << sqrt(alpha[i]/alpha_max) << ",";
        }

        alpha_values << "\n";
        sigma_values << "\n";

        beta_values.close();
        alpha_values.close();
        sigma_values.close();

        y[0] = 4.0;
        y[1] = 4.0;
        if (hyper > 0.0){
            y[2] = 4.0;
        }
        else{
            y[2] = 0.0;
        }

        alpha_inds[0] = alpha_init;
        sigma_inds[0] = sigma_init;
        
        
        num_strains = 1;
        for (int evo_counter = 0; evo_counter < evo_steps; evo_counter++){
            
            t[0] = 0.0;
            h[0] = 1e-1;
            hnext[0] = 1e-1;

            float y_check_sum = 0.0;
            
            tcheck[0] = 10.0;
            for(i=0;i<(2*num_strains + 1);i++){
                y_check[i] = y[i];
                y_max[i] = y[i];
                y_min[i] = y[i];
                y_max_next[i] = y[i];
                y_min_next[i] = y[i];
                y_check_sum += y[i];
                
            }
            int step_count = 0;
            int check_count = 0;
            while (t[0] <= tmax){   
                int counter[1],num_plus[1],num_down[1];
                
                counter[0] = 0;
                num_plus[0] = 0;
                num_down[0] = 0;

                /* This is where the equations are first solved */
                dynamics(dydt, y, num_strains, alpha_inds, sigma_inds, b, q, d, beta, alpha, eta, gamma, sigma, lambda, rho);

                /* Adjust the step size to maintain accuracy */
                for (i=0; i<(2*num_strains + 1); i++){
                    yScale[i]=fabs(y[i])+fabs(dydt[i]*(*h))+EPS;
                }
                
                rkqs(y, dydt, h,hnext,yScale, num_strains, alpha_inds, sigma_inds, b, q, d, beta, alpha, eta, gamma, sigma, lambda, rho,counter,num_plus,num_down,discrepancy, int_tracker, t);
                
                if (y[0] <= TINY){
                    y[0] = 0.0;
                }

                for(i=0;i<(num_strains);i++){
                    if (y[i+1] + y[i + num_strains + 1] <= TINY){
                        y[i+1] = 0.0;
                        y[i + num_strains + 1] = 0.0;
                    }
                    if(y[i + num_strains + 1] <= TINY){
                        y[i + num_strains + 1] = 0.0;
                    }
                }
                t[0]+=h[0];
                h[0] = hnext[0];
                
                
                for(i=0;i<(2*num_strains + 1);i++){
                    y_max[i] = fastmax(y[i], y_max[i]);
                    y_min[i] = fastmin(y[i], y_min[i]);
                }
                step_count += 1;
                
                if ((((t[0] - tcheck[0]) >= 10.0) && (step_count - check_count) >= 200) && (y[0] != 0.0)){
                    discrep_check = 0.0;
                    check_count = step_count;
                    for(i=0;i<(2*num_strains + 1);i++){

                        if (y[i] >= TINY){
                            if (discrep_check < fastmax(abs(y_max[i] - y_max_next[i]),abs(y_min[i] - y_min_next[i]))){
                                discrep_check = fastmax(abs(y_max[i] - y_max_next[i]),abs(y_min[i] - y_min_next[i]));
                            }
                        }
                    }

                    if (discrep_check >= 1e-3){
                        // std::cout << discrep_check << "\n";
                        // for(i=0;i<(2*num_strains + 1);i++){
                            // std::cout << abs((y_check[i] - y[i])/(y_check_sum + EPS)) << "\n";
                            // std::cout << "The difference between the max and the min is " << (y_max[i] - y_min[i])/(y_min[i] + EPS) << " for ind " << i <<  " on evolutionary step " << evo_counter << " with problem ind " << problem_ind <<  "\n";
                            // std::cout << "The difference between the check value and the base value for ind " << i << " is " << abs(y_check[i] - y[i]) << " this represents a discrepancy of " << abs((y_check[i] - y[i])/(y_check[i] + EPS)) << " relative to the system\n"; 
                            // discrep_check = fastmax(abs(y_max[i] - y_min[i]), discrep_check);
                        // }   

                        // std::cout << "For the problem ind " << problem_ind << " the max density since our last check was " << y_max_next[problem_ind] << " and the max density at the previous check was " << y_max[i] << "\n";
                        // std::cout << "and the min density was " << y_min_next[problem_ind] << " during the ecological steps  compared to " << y_min[problem_ind] << " at the previous time step, the current value is " << y[problem_ind] <<"\n";
                        // std::cout << "This is a difference of size " << (y_max[problem_ind] - y_min[problem_ind])/(y_min[problem_ind] + EPS) <<  " relative to the density fo the system + our small constant of size " << EPS << " :) \n";
                        y_check_sum = 0.0;
                        // std::cout << "Time we checked against was " << tcheck[0] << endl;
                        // discrep_check = 0.0;
                        // for(i=0;i<(2*num_strains + 1);i++){
                        //     std::cout << "At this time the value we are checking against was " << y_check[i] << " compared to " << y[i] << endl;
                        //     std::cout << "The difference is " << abs((y_check[i] - y[i])) << " compared to a baseline of " << (y_check[i] + EPS) << endl;
                        //     std::cout << "This gives a discrepancy of " << abs((y_check[i] - y[i])/(y_check[i] + EPS)) << endl;
                        //     discrep_check = fastmax(discrep_check,abs((y_check[i] - y[i])/(y_check[i] + EPS)));
                        //     std::cout << "Our new maximum discrepancy is " << discrep_check << endl;
                        
                        // }
                        // check_counter++;
                        tcheck[0] = t[0];
                        for(i=0;i<(2*num_strains + 1);i++){
                            // if (i != 0 && (y[i] < y_check[i]) && (dydt[i] < 0) && (y[i] < 1e-1)){
                            //     y_check[i] = y[i];
                            //     y_check_sum += y[i];
                            //     y[i] = 0.0;
                            // }
                            // else{
                            //     y_check[i] = y[i];
                            // }
                            y_check[i] = y[i];
                            y_max[i] = y[i];
                            y_min[i] = y[i];
                            // y_max[i] = y_max_next[i];
                            // y_min[i] = y_min_next[i];
                            y_max_next[i] = y[i];
                            y_min_next[i] = y[i];
                        }
                    }
                    else{
                        t[0] = tmax + 1.0;
                        // std::cout << "Skipping due to discrepancy\n";
                    }
                }
                // }
            // std::cout << t[0] << endl;
            }
            // std::cout << "For this run we checked the discrepancy " << check_counter << " times" << endl;
            // sleep(1);
            // std::cout << "Solver results " << endl;
            // for (i =0; i<(2*num_strains+1); i++){
            //     std::cout << y[i] << endl;
            // }
            
            // time (&end_eco);
            // float dif = difftime (end_eco,start_eco);
            // printf("Elasped time is %.2lf seconds.\n", dif ); 

            if (y[0] <= TINY){
                std::cout << "Healthy hosts extinct, breaking" << endl;
                break;
            }
            
            num_strains_cleaned = 0;
            // std::cout << "Number of strains pre counting " << num_strains_cleaned << endl;
            // std::cout << "Number of strains pre counting " << num_strains << endl;
            for(i=0;i<num_strains;i++){
                if ((y[i + 1] + y[i + 1 + (num_strains)]) > TINY){
                    // std::cout << "For index " << i << " the population densities are" <<  endl;
                    // std::cout << y[i + 1] << " for the parasite and" << endl;
                    // std::cout << y[i + 1 + num_strains] << " for the hyperparasite" << endl;
                    // std::cout << "The boolean is " << ((y[i + 1] + y[i + 1 +(num_strains)]) > 1e-3) << endl;
                    num_strains_cleaned = num_strains_cleaned + 1;
                }
            }

            if (num_strains_cleaned == 0){
                cout << "All parasites are dead\n";
                break;
            }
            // std::cout << "Number of strains post counting " << num_strains_cleaned << endl;

            float rand_stored1, rand_stored2;

            counter = 0;

            Hsum = 0.0;

            for (i=0;i<num_strains;i++){
                // std::cout << "I value is supposed to be " << y[i + 1] << endl;
                // std::cout << "H value is supposed to be " << y[i + 1 + num_strains] << endl;
                if((y[i + 1] + y[i + 1 + num_strains]) > TINY){
                    alpha_inds_cleaned[counter] = alpha_inds[i];
                    sigma_inds_cleaned[counter] = sigma_inds[i];
                    I_temp[counter] = y[i +1];
                    H_temp[counter] = y[i +1 + num_strains];
                    // std::cout << "I value written in is " << y[i + 1] << endl;
                    // std::cout << "H value written in is " << y[i + 1 + num_strains] << endl;
                    total_density[counter] = I_temp[counter] + H_temp[counter];
                    Hsum = Hsum + H_temp[counter];
                    counter++;
                }
            }
            // std::cout << "Hsum value is " << Hsum << endl;

            // std::cout << "Number of strains " << num_strains_cleaned << endl;
            // std::cout << "Counter, should be the same " << counter << endl;
            cum_density[0] = total_density[0];
            if (num_strains_cleaned > 1){
                for(i=1;i<num_strains_cleaned;i++){
                    cum_density[i] = total_density[i] + cum_density[i - 1];
                }
            }

            for(i=0;i<num_strains_cleaned;i++){
                // std::cout << "I density is " << I_temp[i] << endl;
                // std::cout << "H density is " << H_temp[i] << endl;
                cum_props[i] = cum_density[i]/cum_density[num_strains_cleaned-1];
            }

            rand_stored1 = (float(rand())/float((RAND_MAX)));
            rand_stored2 = (float(rand())/float((RAND_MAX)));

            for(i=0;i<num_strains_cleaned;i++){
                if(cum_props[i] >= rand_stored1){
                    ind = i;
                    break;
                }
            }
            
            float rand_stored_choice;
            rand_stored_choice = (float(rand())/float((RAND_MAX)));

            bool flag = true;
            int increment_ind;
            num_strains_2 = num_strains_cleaned;
            // std::cout << "\n num_strains_2 after createion" << endl;
            // std::cout << num_strains_2 << endl;
            if (rand_stored_choice >= 0.5){
                // Doing sigma mutation
                if (((rand_stored2 >=0.5) && (sigma_inds_cleaned[ind] != (trait_space_length-1))) || (sigma_inds_cleaned[ind] == 0)){
                    for(i=0;i<num_strains_cleaned;i++){
                        if (sigma_inds_cleaned[i] == (sigma_inds_cleaned[ind] + 1) && (alpha_inds_cleaned[i] == alpha_inds_cleaned[ind])){
                            flag = false;
                            increment_ind = i;
                            
                            break;
                        }
                    }
                }
                else{
                    for(i=0;i<num_strains_cleaned;i++){
                        if (sigma_inds_cleaned[i] == (sigma_inds_cleaned[ind] - 1) && (alpha_inds_cleaned[i] == alpha_inds_cleaned[ind])){
                            flag = false;
                            increment_ind = i;
                            break;
                        }
                    }
                }
                
                if(flag){
                    num_strains_2++;
                }
                // std::cout << num_strains_2 << endl;
                y_temp[0] = y[0];
                for(i = 0; i<num_strains_2;i++){
                    y_temp[i+1] = 0.0;
                    y_temp[i+1+num_strains_2] = 0.0;
                }
                for(i=0;i<num_strains_cleaned;i++){
                    alpha_inds_2[i] = alpha_inds_cleaned[i];
                    sigma_inds_2[i] = sigma_inds_cleaned[i];
                    y_temp[i+1] = I_temp[i];
                    y_temp[i+1+num_strains_2] = H_temp[i];
                }

                num_strains = num_strains_2;

                for(i = 0; i<(2*num_strains+1); i++){
                    y[i] = 0.0;
                }

                y[0] = y_temp[0];
            
                for(i=0;i<num_strains;i++){
                    alpha_inds[i] = alpha_inds_2[i];
                    sigma_inds[i] = sigma_inds_2[i];
                    y[i+1] = y_temp[i+1];
                    y[i+1+num_strains] = y_temp[i+1+num_strains];
                }

                if (((rand_stored2 >=0.5) && (sigma_inds_cleaned[ind] != trait_space_length-1)) || (sigma_inds_cleaned[ind] == 0)){
                    if(flag){
                        alpha_inds[num_strains-1] = alpha_inds_2[ind];
                        sigma_inds[num_strains-1] = sigma_inds_2[ind] + 1;
                        y[num_strains] = (y_temp[ind+1]/100);
                        y[2*num_strains] = (y_temp[ind+1+num_strains]/100);
                    }
                    else{
                        y[1+increment_ind] = y[1+increment_ind] + (y_temp[ind+1]/100);
                        y[1+increment_ind+num_strains] = y[1+increment_ind+num_strains] + (y_temp[ind+1+num_strains]/100);
                    }
                }
                else{
                    if(flag){
                        alpha_inds[num_strains-1] = alpha_inds_2[ind];
                        sigma_inds[num_strains-1] = sigma_inds_2[ind] - 1;
                        y[num_strains] = (y_temp[ind+1]/100);
                        y[2*num_strains] = (y_temp[ind+1+num_strains]/100);
                    }
                    else{
                        y[1+increment_ind] = y[1+increment_ind] + (y_temp[ind+1]/100);
                        y[1+increment_ind+num_strains] = y[1+increment_ind+num_strains] + (y_temp[ind+1+num_strains]/100);
                    }
                }
            }
            else{
                // Doing alpha mutation
                if (((rand_stored2 >=0.5) && (alpha_inds_cleaned[ind] != (trait_space_length-1))) || (alpha_inds_cleaned[ind] == 0)){
                    for(i=0;i<num_strains_cleaned;i++){
                        // Checking if a population already exists with the trait index we care about
                        if (alpha_inds_cleaned[i] == (alpha_inds_cleaned[ind] + 1) && (sigma_inds_cleaned[i] == sigma_inds_cleaned[ind])){
                            flag = false;
                            increment_ind = i;
                            break;
                        }
                    }
                }
                else{
                    for(i=0;i<num_strains_cleaned;i++){
                        if (alpha_inds_cleaned[i] == (alpha_inds_cleaned[ind] - 1) && (sigma_inds_cleaned[i] == sigma_inds_cleaned[ind])){
                            flag = false;
                            increment_ind = i;
                            break;
                        }
                    }
                }
                
                if(flag){
                    num_strains_2++;
                }

                y_temp[0] = y[0];
                for(i = 0; i<num_strains_2;i++){
                    y_temp[i+1] = 0.0;
                    y_temp[i+1+num_strains_2] = 0.0;
                }
                for(i=0;i<num_strains_cleaned;i++){
                    alpha_inds_2[i] = alpha_inds_cleaned[i];
                    sigma_inds_2[i] = sigma_inds_cleaned[i];
                    y_temp[i+1] = I_temp[i];
                    y_temp[i+1+num_strains_2] = H_temp[i];
                }

                num_strains = num_strains_2;

                for(i = 0; i<(2*num_strains+1); i++){
                    y[i] = 0.0;
                }

                y[0] = y_temp[0];
            
                for(i=0;i<num_strains;i++){
                    alpha_inds[i] = alpha_inds_2[i];
                    sigma_inds[i] = sigma_inds_2[i];
                    y[i+1] = y_temp[i+1];
                    y[i+1+num_strains] = y_temp[i+1+num_strains];
                }

                if (((rand_stored2 >=0.5) && (alpha_inds_cleaned[ind] != trait_space_length-1)) || (alpha_inds_cleaned[ind] == 0)){
                    if(flag){
                        alpha_inds[num_strains-1] = alpha_inds_2[ind] + 1;
                        sigma_inds[num_strains-1] = sigma_inds_2[ind];
                        y[num_strains] = (y_temp[ind+1]/100);
                        y[2*num_strains] = (y_temp[ind+1+num_strains]/100);
                    }
                    else{
                        y[1+increment_ind] = y[1+increment_ind] + (y_temp[ind+1]/100);
                        y[1+increment_ind+num_strains] = y[1+increment_ind+num_strains] + (y_temp[ind+1+num_strains]/100);
                    }
                }
                else{
                    if(flag){
                        alpha_inds[num_strains-1] = alpha_inds_2[ind] - 1;
                        sigma_inds[num_strains-1] = sigma_inds_2[ind];
                        y[num_strains] = (y_temp[ind+1]/100);
                        y[2*num_strains] = (y_temp[ind+1+num_strains]/100);
                    }
                    else{
                        y[1+increment_ind] = y[1+increment_ind] + (y_temp[ind+1]/100);
                        y[1+increment_ind+num_strains] = y[1+increment_ind+num_strains] + (y_temp[ind+1+num_strains]/100);
                    }
                }
            }

            for(i=0;i<num_strains;i++){
                output_alpha_inds[output_counter] = alpha_inds[i];
                output_sigma_inds[output_counter] = sigma_inds[i];
                output_parasite_density[output_counter] = y[i+1];
                output_hyperparasite_density[output_counter] = y[i+1+num_strains];
                output_hyperparasite_truths[output_counter] = (Hsum > 0.0);
                evo_step_tracker[output_counter] = evo_counter + 1;
                output_host_density[output_counter] = y[0];
                output_counter++;
            }

            if (evo_counter%100 == 0){
                std::cout << evo_counter << "\n";
            }
        }
        
        string test;

        string seed_str;
        seed_str = std::to_string(seed);

        //Updating the tracker file with our most recent run
        tracker_file.open("../data/tracker_file.csv", std::ios_base::app);
        
        tracker_file << "/coevo/data_set" + seed_str + ".csv," << b << "," << q << "," << d << "," << rho << "," << sigma_max << "," << gamma;
        tracker_file << "," << eta << "," << lambda << "," << c1 << "," << c2 << "," << beta_max << "," << (hyper > 0.0) << "," << alpha_init << "," << sigma_init  << "\n";
        tracker_file.close();


        // This creates and stores the data from our simulation
        myfile.open("../data/coevo/data_set"+seed_str+".csv");
        myfile << "Trait_index_1,Trait_index_2,Density_of_Hosts,Density_of_parasite,Density_of_hyperparasite,Evolutionary_step,hyperparasites_present,alpha_val,sigma_val,beta_val\n";
        
        for(i=0;i<output_counter;i++){
            myfile << output_alpha_inds[i] <<"," << output_sigma_inds[i] << "," << output_host_density[i] << "," << output_parasite_density[i] << ',' << output_hyperparasite_density[i] << ","<< evo_step_tracker[i] << "," << output_hyperparasite_truths[i] << "," << alpha[output_alpha_inds[i]] << "," << sigma[output_sigma_inds[i]] << "," << beta[output_alpha_inds[i]*trait_space_length + output_sigma_inds[i]] << "\n";
        }

        myfile.close();
        
        delete y_check;
        delete y;
        delete yScale;
        delete dydt;
        delete y_temp;
        time (&end);
        float dif = difftime (end,start);
        printf ("Elasped time is %.2lf seconds.\n", dif ); 
        return 0;
    }

    void Quick_solver::eco_dynamics(float* S, float* I, float* H,int* exit_flag, float beta_value, float alpha_value, float sigma_value, float b, float q, float d, float rho, float eta, float gamma, float lambda, float c1, float c2, float hyper, int seed, float S_density, float I_density, float H_density){

        float y[3], yScale[3], dydt[3], discrepancy[1], discrep_check;
        float t[1], tcheck[1], tmax = 4000, h[1], hnext[1];
        float y_max[3], y_min[3], y_max_next[3], y_min_next[3], y_check_sum;
        int alpha_inds[1], sigma_inds[1];
        int num_strains = 1;
        float beta[1], alpha[1], sigma[1];
        int i, int_tracker[0], step_count = 0;
        bool flag = true;
        alpha_inds[0] = 0;
        sigma_inds[0] = 0;

        y[0] = S_density/(b/q);
        y[1] = I_density/(b/q);
        if (hyper > 0.0){
            y[2] = H_density/(b/q);
        }
        else{
            y[2] = 0.0;
        }
        alpha[0] = alpha_value;
        sigma[0] = sigma_value;
        beta[0] = beta_value;

        alpha_inds[0] = 0;
        sigma_inds[0] = 0;
        
        t[0] = 0.0;
        h[0] = 1e-2;
        tcheck[0] = 10;
        for(i=0;i<(2*num_strains + 1);i++){

            y_max[i] = y[i];
            y_min[i] = y[i];
            y_max_next[i] = y[i];
            y_min_next[i] = y[i];
            y_check_sum += y[i];
        }
        hnext[0] = 1e-2;

        while (t[0] <= tmax){   
            int counter[1],num_plus[1],num_down[1];
            int check_count = 0;
            counter[0] = 0;
            num_plus[0] = 0;
            num_down[0] = 0;

            h[0] = hnext[0];
            
            /* This is where the equations are first solved */
            dynamics(dydt, y, num_strains, alpha_inds, sigma_inds, b, q, d, beta, alpha, eta, gamma, sigma, lambda, rho);
        
            /* Adjust the step size to maintain accuracy */
            for (i=0; i<(2*num_strains + 1); i++){
                yScale[i]=fabs(y[i])+fabs(dydt[i]*(*h))+TINY;
            }
            
            // Runga Kutta 4th order method to solve the equations
            rkqs(y, dydt, h,hnext,yScale, num_strains, alpha_inds, sigma_inds, b, q, d, beta, alpha, eta, gamma, sigma, lambda, rho,counter,num_plus,num_down,discrepancy, int_tracker, t);
            
            if (y[1] + y[2] <= TINY){
                y[1] = 0.0;
                y[2] = 0.0;
            }
            
//             if (y[2] <= TINY){
//                 y[2] = 0.0;
//             }
//             if (y[1] <= TINY && rho == 1.0){
//                 y[1] = 0.0;
//             }

            if (y[1] == 0.0 && y[2] == 0.0 && hyper > 0.0 && flag){
                flag = false;
                y[1] = TINY*10.0;
            }

            t[0]+=h[0];

            for(i=0;i<(2*num_strains + 1);i++){
                y_max[i] = fastmax(y[i], y_max[i]);
                y_min[i] = fastmin(y[i], y_min[i]);
            }
            step_count += 1;  
            
            //This section checks the amount of change in all of the populations, if all of the populations experience relatively small changes we exit
            if ((((t[0] - tcheck[0]) >= 10.0) || (step_count - check_count) >= 200) && (y[0] != 0.0)){
                discrep_check = 0.0;
                check_count = step_count;
                bool check_flag = false;
                
                for(i=0;i<(2*num_strains + 1);i++){
                    if (discrep_check < fastmax(abs(y_max[i] - y_max_next[i]),abs(y_min[i] - y_min_next[i]))){
                        discrep_check = fastmax(abs(y_max[i] - y_max_next[i]),abs(y_min[i] - y_min_next[i]));
                        if (y_max[i] != 0.0 && y[i] == 0.0){
                            check_flag = true;
                        }
                    }
                }

                if (discrep_check >= 10*TINY || check_flag){
                    y_check_sum = 0.0;

                    tcheck[0] = t[0];
                    for(i=0;i<(2*num_strains + 1);i++){
                        y_max[i] = y[i];
                        y_min[i] = y[i];
                        
                        y_max_next[i] = y[i];
                        y_min_next[i] = y[i];
                    }
                    exit_flag[0] = 0;
                }
                else{
                    exit_flag[0] = 1;
                    t[0] = tmax + 1.0;;
                }
            }
        }

        //Returning the values of the populations that we care about 
        S[0] = y[0];
        I[0] = y[1];
        H[0] = y[2];
//         exit_flag[0] = 1;
    }

    void Quick_solver::alpha_evo_only(float beta_max, float alpha_max, float sigma_max, float b, float q, float d, float rho, float eta, float gamma, float lambda, float c1, float c2, float hyper, int seed, int alpha_init, int sigma_init, float S_density, float I_density, float H_density){
        
        time_t start,end;
        time (&start);
        

        srand(seed);
        
        int tmax = 1000, evo_steps = 1000, i, j;
        float beta[trait_space_length*trait_space_length], alpha[trait_space_length], sigma[trait_space_length];
        float t[1], tcheck[1], Hsum;
        int num_strains = 1, alpha_inds[trait_space_length], ind,num_strains_2;
        int sigma_inds[trait_space_length], num_strains_cleaned;

        float* y = new float[2*trait_space_length+1];
        float* y_check = new float[2*trait_space_length + 1];
        float* dydt = new float[2*trait_space_length + 1];
        float* yScale = new float[2*trait_space_length + 1];
        
        float* y_max = new float[2*trait_space_length + 1];
        float* y_min = new float[2*trait_space_length + 1];
        float* y_max_next = new float[2*trait_space_length + 1];
        float* y_min_next = new float[2*trait_space_length + 1];

        float discrep_check;
        int alpha_inds_cleaned[trait_space_length],counter=0;
        int sigma_inds_cleaned[trait_space_length];
        float I_temp[trait_space_length], H_temp[trait_space_length],total_density[trait_space_length];
        float cum_density[trait_space_length], cum_props[trait_space_length];
        int alpha_inds_2[trait_space_length];
        int sigma_inds_2[trait_space_length], int_tracker[1];
        float* y_temp = new float[2*trait_space_length+1];
        
        float h[1], hnext[1], discrepancy[1];

        int num_poss_outputs = 30000;
        int output_alpha_inds[num_poss_outputs], output_sigma_inds[num_poss_outputs], output_counter = 0, evo_step_tracker[num_poss_outputs];
        float output_parasite_density[num_poss_outputs], output_hyperparasite_density[num_poss_outputs], output_host_density[num_poss_outputs];
        bool output_hyperparasite_truths[num_poss_outputs];
        
        ind = 0;
        ofstream myfile, tracker_file;

        for(i=0;i<trait_space_length;i++){
            alpha[i] = (alpha_max*(i))/(trait_space_length - 1);
            sigma[i] = (sigma_max*(i))/(trait_space_length - 1);
        }

        for(i=0;i<trait_space_length;i++){
            for(j=0;j<trait_space_length;j++){
                if (c2 == 0.0){
                    beta[i*trait_space_length + j] = beta_max*(sqrt(alpha[i]/alpha_max)*((1-c1) + c1*(sigma[j]/sigma_max)));
                }
                else{
                    beta[i*trait_space_length+j] = fastmax(0,beta_max*(sqrt(alpha[i]/alpha_max)*((1-c1) + c1*(1 - exp((sigma[j]*c2)/sigma_max))/(1 - exp(c2)))));
                }
            }
        }

        ofstream beta_values,sigma_values, alpha_values;
        beta_values.open("../data/beta_vals.csv");
        sigma_values.open("../data/sigma_vals.csv");
        alpha_values.open("../data/alpha_vals.csv");

        for (i = 0; i<trait_space_length;i++){
            beta_values << i << ",";
            alpha_values << i << ",";
            sigma_values << i << ",";
        }
        beta_values << "\n";
        alpha_values << "\n";
        sigma_values << "\n";

        

        for(i=0;i<trait_space_length;i++){
            sigma_values << sigma[i] << ",";
            for(j=0;j<trait_space_length;j++){
                beta_values << beta[i*trait_space_length+j] << ",";
            }
            beta_values << "\n";
        }

        alpha_values << "\n";
        sigma_values << "\n";

        for (i = 0; i<trait_space_length;i++){
            sigma_values << ((1-c1) + c1*(1 - exp((sigma[i]*c2)/sigma_max))/(1 - exp(c2))) << ",";
            alpha_values << sqrt(alpha[i]/alpha_max) << ",";
        }

        alpha_values << "\n";
        sigma_values << "\n";

        beta_values.close();
        sigma_values.close();
        alpha_values.close();

        // y[0] = (4.0)/(b/q);
        // y[1] = (4.0)/(b/q);
        // if (hyper > 0.0){
        //     y[2] = (4.0)/(b/q);
        // }
        // else{
        //     y[2] = 0.0;
        // }
        y[0] = S_density;
        y[1] = I_density;
        if (hyper > 0.0){
            y[2] = H_density;
        }
        else{
            y[2] = 0.0;
        }

        alpha_inds[0] = alpha_init;
        sigma_inds[0] = sigma_init;
        
        num_strains = 1;
        for (int evo_counter = 0; evo_counter < evo_steps; evo_counter++){
            
            t[0] = 0.0;
            h[0] = 1e-1;
            hnext[0] = 1e-1;

            float y_check_sum = 0.0;
            
            tcheck[0] = 10.0;
            for(i=0;i<(2*num_strains + 1);i++){
                y_check[i] = y[i];
                y_max[i] = y[i];
                y_min[i] = y[i];
                y_max_next[i] = y[i];
                y_min_next[i] = y[i];
                y_check_sum += y[i];
                // y_max[i] = y[i];
                // y_min[i] = y[i];
                // std::cout << "y value pre-solver is " << y[i] << endl;
            }
            int step_count = 0;
            int check_count = 0;
            while (t[0] <= tmax){   
                int counter[1],num_plus[1],num_down[1];
                
                counter[0] = 0;
                num_plus[0] = 0;
                num_down[0] = 0;

                // h[0] = 1.0;
                // grad_max = 0.0;
                /* This is where the equations are first solved */
                dynamics(dydt, y, num_strains, alpha_inds, sigma_inds, b, q, d, beta, alpha, eta, gamma, sigma, lambda, rho);

                /* Adjust the step size to maintain accuracy */
                for (i=0; i<(2*num_strains + 1); i++){
                    yScale[i]=fabs(y[i])+fabs(dydt[i]*(*h))+EPS;
                    // grad_max = Quick_solver::fastmax(grad_max, abs(dydt[i]));
                }
                
                rkqs(y, dydt, h,hnext,yScale, num_strains, alpha_inds, sigma_inds, b, q, d, beta, alpha, eta, gamma, sigma, lambda, rho,counter,num_plus,num_down,discrepancy, int_tracker, t);
                
                if (y[0] <= TINY){
                    y[0] = 0.0;
                }

                for(i=0;i<(num_strains);i++){
                    if (y[i+1] + y[i + num_strains + 1] <= TINY){
                        y[i+1] = 0.0;
                        y[i + num_strains + 1] = 0.0;
                    }
                    if(y[i + num_strains + 1] <= TINY){
                        y[i + num_strains + 1] = 0.0;
                    }
                }

                t[0]+=h[0];
                h[0] = hnext[0];
                
                for(i=0;i<(2*num_strains + 1);i++){
                    y_max[i] = fastmax(y[i], y_max[i]);
                    y_min[i] = fastmin(y[i], y_min[i]);
                }
                step_count += 1;
                
                if ((((t[0] - tcheck[0]) >= 10.0) && (step_count - check_count) >= 200) && (y[0] != 0.0)){
                    discrep_check = 0.0;
                    check_count = step_count;
                    
                    for(i=0;i<(2*num_strains + 1);i++){

                        if (y[i] >= TINY){
                            if (discrep_check < fastmax(abs(y_max[i] - y_max_next[i]),abs(y_min[i] - y_min_next[i]))){
                                
                                discrep_check = fastmax(abs(y_max[i] - y_max_next[i]),abs(y_min[i] - y_min_next[i]));
                            }
                        }
                    }

                    if (discrep_check >= 1e-3){

                        y_check_sum = 0.0;
                        tcheck[0] = t[0];

                        for(i=0;i<(2*num_strains + 1);i++){
                            y_check[i] = y[i];
                            y_max[i] = y[i];
                            y_min[i] = y[i];
                            y_max_next[i] = y[i];
                            y_min_next[i] = y[i];
                        }
                    }
                    else{
                        t[0] = tmax + 1.0;
                    }
                }
            }

            if (y[0] <= TINY){
                std::cout << "Healthy hosts extinct, breaking" << endl;
                break;
            }
            
            num_strains_cleaned = 0;
            for(i=0;i<num_strains;i++){
                if ((y[i + 1] + y[i + 1 + (num_strains)]) > TINY){
                    num_strains_cleaned = num_strains_cleaned + 1;
                }
            }

            if (num_strains_cleaned == 0){
                cout << "All parasites are dead\n";
                break;
            }

            float rand_stored1, rand_stored2;

            counter = 0;

            Hsum = 0.0;

            for (i=0;i<num_strains;i++){
                if((y[i + 1] + y[i + 1 + num_strains]) > TINY){
                    alpha_inds_cleaned[counter] = alpha_inds[i];
                    sigma_inds_cleaned[counter] = sigma_inds[i];
                    I_temp[counter] = y[i +1];
                    H_temp[counter] = y[i +1 + num_strains];
                    total_density[counter] = I_temp[counter] + H_temp[counter];
                    Hsum = Hsum + H_temp[counter];
                    counter++;
                }
            }
            
            bool HSum_flag = (Hsum >= TINY);

            cum_density[0] = total_density[0];
            if (num_strains_cleaned > 1){
                for(i=1;i<num_strains_cleaned;i++){
                    cum_density[i] = total_density[i] + cum_density[i - 1];
                }
            }

            for(i=0;i<num_strains_cleaned;i++){
                cum_props[i] = cum_density[i]/cum_density[num_strains_cleaned-1];
            }

            rand_stored1 = (float(rand())/float((RAND_MAX)));
            rand_stored2 = (float(rand())/float((RAND_MAX)));

            for(i=0;i<num_strains_cleaned;i++){
                if(cum_props[i] >= rand_stored1){
                    ind = i;
                    break;
                }
            }

            bool flag = true;
            int increment_ind;
            num_strains_2 = num_strains_cleaned;
            
            if (((rand_stored2 >=0.5) && (alpha_inds_cleaned[ind] != (trait_space_length-1))) || (alpha_inds_cleaned[ind] == 0)){
                for(i=0;i<num_strains_cleaned;i++){
                    if (alpha_inds_cleaned[i] == (alpha_inds_cleaned[ind] + 1)){
                        flag = false;
                        increment_ind = i;
                        // std::cout << "Alpha increment ind is" << increment_ind << endl;
                        break;
                    }
                }
            }
            else{
                for(i=0;i<num_strains_cleaned;i++){
                    if (alpha_inds_cleaned[i] == (alpha_inds_cleaned[ind] - 1) && (sigma_inds_cleaned[i] == sigma_inds_cleaned[ind])){
                        flag = false;
                        increment_ind = i;
                        break;
                    }
                }
            }
            // std::cout << "\n num_strains_2 after incrementing in alpha step" << endl;
            // int num_strains_2 = num_strains_cleaned;
            if(flag){
                num_strains_2++;
            }
            // std::cout << num_strains_2 << endl;

            y_temp[0] = y[0];
            for(i = 0; i<num_strains_2;i++){
                y_temp[i+1] = 0.0;
                y_temp[i+1+num_strains_2] = 0.0;
            }
            for(i=0;i<num_strains_cleaned;i++){
                alpha_inds_2[i] = alpha_inds_cleaned[i];
                sigma_inds_2[i] = sigma_inds_cleaned[i];
                y_temp[i+1] = I_temp[i];
                y_temp[i+1+num_strains_2] = H_temp[i]*HSum_flag;
            }

            // std::cout << "Reallocated number of strains" << endl;
            num_strains = num_strains_2;

            for(i = 0; i<(2*num_strains+1); i++){
                y[i] = 0.0;
            }

            y[0] = y_temp[0];
        
            for(i=0;i<num_strains;i++){
                alpha_inds[i] = alpha_inds_2[i];
                sigma_inds[i] = sigma_inds_2[i];
                y[i+1] = y_temp[i+1];
                y[i+1+num_strains] = y_temp[i+1+num_strains];
            }

            if (((rand_stored2 >=0.5) && (alpha_inds_cleaned[ind] != trait_space_length-1)) || (alpha_inds_cleaned[ind] == 0)){
                if(flag){
                    alpha_inds[num_strains-1] = alpha_inds_2[ind] + 1;
                    sigma_inds[num_strains-1] = sigma_inds_2[ind];
                    y[num_strains] = (y_temp[ind+1]/100);
                    y[2*num_strains] = (y_temp[ind+1+num_strains]/100);
                }
                else{
                    y[1+increment_ind] = y[1+increment_ind] + (y_temp[ind+1]/100);
                    y[1+increment_ind+num_strains] = y[1+increment_ind+num_strains] + (y_temp[ind+1+num_strains]/100);
                }
            }
            else{
                if(flag){
                    alpha_inds[num_strains-1] = alpha_inds_2[ind] - 1;
                    sigma_inds[num_strains-1] = sigma_inds_2[ind];
                    y[num_strains] = (y_temp[ind+1]/100);
                    y[2*num_strains] = (y_temp[ind+1+num_strains]/100);
                }
                else{
                    y[1+increment_ind] = y[1+increment_ind] + (y_temp[ind+1]/100);
                    y[1+increment_ind+num_strains] = y[1+increment_ind+num_strains] + (y_temp[ind+1+num_strains]/100);
                }
            }
            
            for(i=0;i<num_strains;i++){
                output_alpha_inds[output_counter] = alpha_inds[i];
                output_sigma_inds[output_counter] = sigma_inds[i];
                output_parasite_density[output_counter] = y[i+1];
                output_hyperparasite_density[output_counter] = y[i+1+num_strains];
                output_hyperparasite_truths[output_counter] = (Hsum > 0.0);
                evo_step_tracker[output_counter] = evo_counter + 1;
                output_host_density[output_counter] = y[0];
                output_counter++;
            }

            if (evo_counter%100 == 0){
                std::cout << evo_counter << "\r";
            }
        }
        
        string test;

        string seed_str;
        seed_str = std::to_string(seed);

        tracker_file.open("../data/tracker_file.csv", std::ios_base::app);
        tracker_file << "/alpha_evo/data_set" + seed_str + ".csv," << b << "," << q << "," << d << "," << rho << "," << sigma_max << "," << gamma;
        tracker_file << "," << eta << "," << lambda << "," << c1 << "," << c2 << "," << beta_max << "," << (hyper > 0.0) << "," << alpha_init << "," << sigma_init  << "\n";
        tracker_file.close();

        myfile.open("../data/alpha_evo/data_set"+seed_str+".csv");
        myfile << "Trait_index_1,Trait_index_2,Density_of_Hosts,Density_of_parasite,Density_of_hyperparasite,Evolutionary_step,hyperparasites_present,alpha_val,sigma_val,beta_val\n";
        
        for(i=0;i<output_counter;i++){
            myfile << output_alpha_inds[i] <<"," << output_sigma_inds[i] << "," << output_host_density[i] << "," << output_parasite_density[i] << ',' << output_hyperparasite_density[i] << ","<< evo_step_tracker[i] << "," << output_hyperparasite_truths[i] << "," << alpha[output_alpha_inds[i]] << "," << sigma[output_sigma_inds[i]] << "," << beta[output_alpha_inds[i]*trait_space_length + output_sigma_inds[i]] << "\n";
        }

        myfile.close();
        
        delete y_check;
        delete y;
        delete yScale;
        delete dydt;
        delete y_temp;
        time (&end);
        float dif = difftime (end,start);
        printf ("Elasped time is %.2lf seconds.\n", dif ); 
        return;
    }

    void Quick_solver::alpha_evo_only_v2(float beta_max, float beta_min, float alpha_max, float alpha_min, float sigma_max, float b, float q, float d, float rho, float eta, float gamma, float lambda, float c1, float c2, float beta_scalar, float hyper, int seed, int alpha_init, int sigma_init, float S_density, float I_density, float H_density){
        
        // time_t start,end;
        // time (&start);
        

        srand(seed);
        
        int tmax = 1000, evo_steps = 1000, i, j;
        float beta[trait_space_length*trait_space_length], alpha[trait_space_length], sigma[trait_space_length];
        float t[1], tcheck[1], Hsum;
        int num_strains = 1, alpha_inds[trait_space_length], ind,num_strains_2;
        int sigma_inds[trait_space_length], num_strains_cleaned;

        float* y = new float[2*trait_space_length+1];
        float* y_check = new float[2*trait_space_length + 1];
        float* dydt = new float[2*trait_space_length + 1];
        float* yScale = new float[2*trait_space_length + 1];
        
        float* y_max = new float[2*trait_space_length + 1];
        float* y_min = new float[2*trait_space_length + 1];
        float* y_max_next = new float[2*trait_space_length + 1];
        float* y_min_next = new float[2*trait_space_length + 1];
        
        float alpha_scalar, sigma_scalar;
        float discrep_check;
        int alpha_inds_cleaned[trait_space_length],counter=0;
        int sigma_inds_cleaned[trait_space_length];
        float I_temp[trait_space_length], H_temp[trait_space_length],total_density[trait_space_length];
        float cum_density[trait_space_length], cum_props[trait_space_length];
        int alpha_inds_2[trait_space_length];
        int sigma_inds_2[trait_space_length], int_tracker[1];
        float* y_temp = new float[2*trait_space_length+1];
        
        float h[1], hnext[1], discrepancy[1];

        int num_poss_outputs = 50000;
        int output_alpha_inds[num_poss_outputs], output_sigma_inds[num_poss_outputs], output_counter = 0, evo_step_tracker[num_poss_outputs];
        float output_parasite_density[num_poss_outputs], output_hyperparasite_density[num_poss_outputs], output_host_density[num_poss_outputs];
        bool output_hyperparasite_truths[num_poss_outputs];
        
        ind = 0;
        ofstream myfile, tracker_file;

        for(i=0;i<trait_space_length;i++){
            alpha[i] = alpha_min + ((alpha_max - alpha_min)*(i))/(trait_space_length - 1);
            sigma[i] = (sigma_max*(i))/(trait_space_length - 1);
        }

        for(i=0;i<trait_space_length;i++){
            if (c2 == 0.0){
                alpha_scalar = (1-c1) + c1*(alpha[i]/alpha_max);
            }
            else{
                alpha_scalar = (1-c1) + c1*(1 - exp(c2*(alpha[i]/alpha_max)))/(1 - exp(c2));
            }
            
            for(j=0;j<trait_space_length;j++){    
                sigma_scalar = beta_scalar;
                float beta_temp = beta_min + (beta_max - beta_min)*alpha_scalar;
                // float beta_temp = (beta_max - beta_min)*alpha_scalar;
                beta[i*trait_space_length+j] = fastmax(0,beta_temp);
            }
        }

        ofstream beta_values,sigma_values, alpha_values;
        beta_values.open("../data/beta_vals.csv");
        sigma_values.open("../data/sigma_vals.csv");
        alpha_values.open("../data/alpha_vals.csv");

        for (i = 0; i<trait_space_length;i++){
            beta_values << i << ",";
            alpha_values << i << ",";
            sigma_values << i << ",";
        }
        beta_values << "\n";
        alpha_values << "\n";
        sigma_values << "\n";

        

        for(i=0;i<trait_space_length;i++){
            sigma_values << sigma[i] << ",";
            alpha_values << alpha[i] << ",";
            for(j=0;j<trait_space_length;j++){
                beta_values << beta[i*trait_space_length+j] << ",";
            }
            beta_values << "\n";
        }

        alpha_values << "\n";
        sigma_values << "\n";

        for (i = 0; i<trait_space_length;i++){
            sigma_values << beta_scalar << ",";
            alpha_values << ((1-c1) + (c1*(1 - exp((alpha[i]*c2)/alpha_max))/(1 - exp(c2)))) << ",";
        }

        alpha_values << "\n";
        sigma_values << "\n";

        beta_values.close();
        sigma_values.close();
        alpha_values.close();

        // y[0] = (4.0)/(b/q);
        // y[1] = (4.0)/(b/q);
        // if (hyper > 0.0){
        //     y[2] = (4.0)/(b/q);
        // }
        // else{
        //     y[2] = 0.0;
        // }
        y[0] = S_density;
        y[1] = I_density;
        if (hyper > 0.0){
            y[2] = H_density;
        }
        else{
            y[2] = 0.0;
        }

        alpha_inds[0] = alpha_init;
        sigma_inds[0] = sigma_init;
        
        num_strains = 1;
        for (int evo_counter = 0; evo_counter < evo_steps; evo_counter++){
            
            t[0] = 0.0;
            h[0] = 1e-1;
            hnext[0] = 1e-1;

            float y_check_sum = 0.0;
            
            tcheck[0] = 10.0;
            for(i=0;i<(2*num_strains + 1);i++){
                y_check[i] = y[i];
                y_max[i] = y[i];
                y_min[i] = y[i];
                y_max_next[i] = y[i];
                y_min_next[i] = y[i];
                y_check_sum += y[i];
                // y_max[i] = y[i];
                // y_min[i] = y[i];
                // std::cout << "y value pre-solver is " << y[i] << endl;
            }
            int step_count = 0;
            int check_count = 0;
            while (t[0] <= tmax){   
                int counter[1],num_plus[1],num_down[1];
                
                counter[0] = 0;
                num_plus[0] = 0;
                num_down[0] = 0;

                // h[0] = 1.0;
                // grad_max = 0.0;
                /* This is where the equations are first solved */
                dynamics(dydt, y, num_strains, alpha_inds, sigma_inds, b, q, d, beta, alpha, eta, gamma, sigma, lambda, rho);

                /* Adjust the step size to maintain accuracy */
                for (i=0; i<(2*num_strains + 1); i++){
                    yScale[i]=fabs(y[i])+fabs(dydt[i]*(*h))+EPS;
                    // grad_max = Quick_solver::fastmax(grad_max, abs(dydt[i]));
                }
                
                rkqs(y, dydt, h,hnext,yScale, num_strains, alpha_inds, sigma_inds, b, q, d, beta, alpha, eta, gamma, sigma, lambda, rho,counter,num_plus,num_down,discrepancy, int_tracker, t);
                
                if (y[0] <= TINY){
                    y[0] = 0.0;
                }

                for(i=0;i<(num_strains);i++){
                    if (y[i+1] + y[i + num_strains + 1] <= TINY){
                        y[i+1] = 0.0;
                        y[i + num_strains + 1] = 0.0;
                    }
                    if(y[i + num_strains + 1] <= TINY){
                        y[i + num_strains + 1] = 0.0;
                    }
                }

                t[0]+=h[0];
                h[0] = hnext[0];
                
                for(i=0;i<(2*num_strains + 1);i++){
                    y_max[i] = fastmax(y[i], y_max[i]);
                    y_min[i] = fastmin(y[i], y_min[i]);
                }
                step_count += 1;
                
                if ((((t[0] - tcheck[0]) >= 10.0) && (step_count - check_count) >= 200) && (y[0] != 0.0)){
                    discrep_check = 0.0;
                    check_count = step_count;
                    
                    for(i=0;i<(2*num_strains + 1);i++){

                        if (discrep_check < fastmax(abs(y_max[i] - y_max_next[i]),abs(y_min[i] - y_min_next[i]))){
                            
                            discrep_check = fastmax(abs(y_max[i] - y_max_next[i]),abs(y_min[i] - y_min_next[i]));
                        }
                    }

                    if (discrep_check >= 1e-3){

                        y_check_sum = 0.0;
                        tcheck[0] = t[0];

                        for(i=0;i<(2*num_strains + 1);i++){
                            y_check[i] = y[i];
                            y_max[i] = y[i];
                            y_min[i] = y[i];
                            y_max_next[i] = y[i];
                            y_min_next[i] = y[i];
                        }
                    }
                    else{
                        t[0] = tmax + 1.0;
                    }
                }
            }

            if (y[0] <= TINY){
                std::cout << "Healthy hosts extinct, breaking" << endl;
                break;
            }
            
            num_strains_cleaned = 0;
            for(i=0;i<num_strains;i++){
                if ((y[i + 1] + y[i + 1 + (num_strains)]) > TINY){
                    num_strains_cleaned = num_strains_cleaned + 1;
                }
            }

            if (num_strains_cleaned == 0){
                cout << "All parasites are dead\n";
                break;
            }

            float rand_stored1, rand_stored2;

            counter = 0;

            Hsum = 0.0;

            for (i=0;i<num_strains;i++){
                if((y[i + 1] + y[i + 1 + num_strains]) > TINY){
                    alpha_inds_cleaned[counter] = alpha_inds[i];
                    sigma_inds_cleaned[counter] = sigma_inds[i];
                    I_temp[counter] = y[i +1];
                    H_temp[counter] = y[i +1 + num_strains];
                    total_density[counter] = I_temp[counter] + H_temp[counter];
                    Hsum = Hsum + H_temp[counter];
                    counter++;
                }
            }

            bool HSum_flag = (Hsum >= TINY);
            
            cum_density[0] = total_density[0];
            if (num_strains_cleaned > 1){
                for(i=1;i<num_strains_cleaned;i++){
                    cum_density[i] = total_density[i] + cum_density[i - 1];
                }
            }

            for(i=0;i<num_strains_cleaned;i++){
                cum_props[i] = cum_density[i]/cum_density[num_strains_cleaned-1];
            }

            rand_stored1 = (float(rand())/float((RAND_MAX)));
            rand_stored2 = (float(rand())/float((RAND_MAX)));

            for(i=0;i<num_strains_cleaned;i++){
                if(cum_props[i] >= rand_stored1){
                    ind = i;
                    break;
                }
            }

            bool flag = true;
            int increment_ind;
            num_strains_2 = num_strains_cleaned;
            
            if (((rand_stored2 >=0.5) && (alpha_inds_cleaned[ind] != (trait_space_length-1))) || (alpha_inds_cleaned[ind] == 0)){
                for(i=0;i<num_strains_cleaned;i++){
                    if (alpha_inds_cleaned[i] == (alpha_inds_cleaned[ind] + 1)){
                        flag = false;
                        increment_ind = i;
                        // std::cout << "Alpha increment ind is" << increment_ind << endl;
                        break;
                    }
                }
            }
            else{
                for(i=0;i<num_strains_cleaned;i++){
                    if (alpha_inds_cleaned[i] == (alpha_inds_cleaned[ind] - 1) && (sigma_inds_cleaned[i] == sigma_inds_cleaned[ind])){
                        flag = false;
                        increment_ind = i;
                        break;
                    }
                }
            }
            // std::cout << "\n num_strains_2 after incrementing in alpha step" << endl;
            // int num_strains_2 = num_strains_cleaned;
            if(flag){
                num_strains_2++;
            }
            // std::cout << num_strains_2 << endl;

            y_temp[0] = y[0];
            for(i = 0; i<num_strains_2;i++){
                y_temp[i+1] = 0.0;
                y_temp[i+1+num_strains_2] = 0.0;
            }
            for(i=0;i<num_strains_cleaned;i++){
                alpha_inds_2[i] = alpha_inds_cleaned[i];
                sigma_inds_2[i] = sigma_inds_cleaned[i];
                y_temp[i+1] = I_temp[i];
                y_temp[i+1+num_strains_2] = H_temp[i]*HSum_flag;
            }

            // std::cout << "Reallocated number of strains" << endl;
            num_strains = num_strains_2;

            for(i = 0; i<(2*num_strains+1); i++){
                y[i] = 0.0;
            }

            y[0] = y_temp[0];
        
            for(i=0;i<num_strains;i++){
                alpha_inds[i] = alpha_inds_2[i];
                sigma_inds[i] = sigma_inds_2[i];
                y[i+1] = y_temp[i+1];
                y[i+1+num_strains] = y_temp[i+1+num_strains];
            }

            if (((rand_stored2 >=0.5) && (alpha_inds_cleaned[ind] != trait_space_length-1)) || (alpha_inds_cleaned[ind] == 0)){
                if(flag){
                    alpha_inds[num_strains-1] = alpha_inds_2[ind] + 1;
                    sigma_inds[num_strains-1] = sigma_inds_2[ind];
                    y[num_strains] = (y_temp[ind+1]/100);
                    y[2*num_strains] = (y_temp[ind+1+num_strains]/100);
                }
                else{
                    y[1+increment_ind] = y[1+increment_ind] + (y_temp[ind+1]/100);
                    y[1+increment_ind+num_strains] = y[1+increment_ind+num_strains] + (y_temp[ind+1+num_strains]/100);
                }
            }
            else{
                if(flag){
                    alpha_inds[num_strains-1] = alpha_inds_2[ind] - 1;
                    sigma_inds[num_strains-1] = sigma_inds_2[ind];
                    y[num_strains] = (y_temp[ind+1]/100);
                    y[2*num_strains] = (y_temp[ind+1+num_strains]/100);
                }
                else{
                    y[1+increment_ind] = y[1+increment_ind] + (y_temp[ind+1]/100);
                    y[1+increment_ind+num_strains] = y[1+increment_ind+num_strains] + (y_temp[ind+1+num_strains]/100);
                }
            }
            
            for(i=0;i<num_strains;i++){
                output_alpha_inds[output_counter] = alpha_inds[i];
                output_sigma_inds[output_counter] = sigma_inds[i];
                output_parasite_density[output_counter] = y[i+1];
                output_hyperparasite_density[output_counter] = y[i+1+num_strains];
                output_hyperparasite_truths[output_counter] = (Hsum > 0.0);
                evo_step_tracker[output_counter] = evo_counter + 1;
                output_host_density[output_counter] = y[0];
                output_counter++;
            }

            // if (evo_counter%100 == 0){
            //     std::cout << evo_counter << "\n";
            // }
        }
        
        string test;

        string seed_str;
        seed_str = std::to_string(seed);

        tracker_file.open("../data/tracker_file.csv", std::ios_base::app);
        tracker_file << "/alpha_evo/data_set" + seed_str + ".csv," << b << "," << q << "," << d << "," << rho << "," << sigma_max << "," << gamma;
        tracker_file << "," << eta << "," << lambda << "," << c1 << "," << c2 << "," << beta_max << "," << (hyper > 0.0) << "," << alpha_init << "," << sigma_init  << "\n";
        tracker_file.close();

        myfile.open("../data/alpha_evo/data_set"+seed_str+".csv");
        myfile << "Trait_index_1,Trait_index_2,Density_of_Hosts,Density_of_parasite,Density_of_hyperparasite,Evolutionary_step,hyperparasites_present,alpha_val,sigma_val,beta_val\n";
        
        for(i=0;i<output_counter;i++){
            myfile << output_alpha_inds[i] <<"," << output_sigma_inds[i] << "," << output_host_density[i] << "," << output_parasite_density[i] << ',' << output_hyperparasite_density[i] << ","<< evo_step_tracker[i] << "," << output_hyperparasite_truths[i] << "," << alpha[output_alpha_inds[i]] << "," << sigma[output_sigma_inds[i]] << "," << beta[output_alpha_inds[i]*trait_space_length + output_sigma_inds[i]] << "\n";
        }

        myfile.close();
        
        delete y_check;
        delete y;
        delete yScale;
        delete dydt;
        delete y_temp;
        // time (&end);
        // float dif = difftime (end,start);
        // printf ("Elasped time is %.2lf seconds.\r", dif ); 
        return;
    }

    void Quick_solver::alpha_evo_only_v3(float beta_max, float alpha_max, float sigma_max, float b, float q, float d, float rho, float eta, float gamma, float lambda, float c1, float c2, float hyper, int seed, int alpha_init, int sigma_init, float S_density, float I_density, float H_density){
        
        time_t start,end;
        time (&start);
        

        srand(seed);
        
        int tmax = 1000, evo_steps = 1000, i, j;
        float beta[trait_space_length*trait_space_length], sigma[trait_space_length];
        float alpha[trait_space_length];
        float t[1], tcheck[1], Hsum;
        int num_strains = 1, alpha_inds[trait_space_length], ind,num_strains_2;
        int sigma_inds[trait_space_length], num_strains_cleaned;

        float* y = new float[2*trait_space_length+1];
        float* y_check = new float[2*trait_space_length + 1];
        float* dydt = new float[2*trait_space_length + 1];
        float* yScale = new float[2*trait_space_length + 1];
        
        float* y_max = new float[2*trait_space_length + 1];
        float* y_min = new float[2*trait_space_length + 1];
        float* y_max_next = new float[2*trait_space_length + 1];
        float* y_min_next = new float[2*trait_space_length + 1];

        float discrep_check;
        int alpha_inds_cleaned[trait_space_length],counter=0;
        int sigma_inds_cleaned[trait_space_length];
        float I_temp[trait_space_length], H_temp[trait_space_length],total_density[trait_space_length];
        float cum_density[trait_space_length], cum_props[trait_space_length];
        int alpha_inds_2[trait_space_length];
        int sigma_inds_2[trait_space_length], int_tracker[1];
        float* y_temp = new float[2*trait_space_length+1];
        
        float beta_0[trait_space_length];
        float h[1], hnext[1], discrepancy[1];

        int num_poss_outputs = 30000;
        int output_alpha_inds[num_poss_outputs], output_sigma_inds[num_poss_outputs], output_counter = 0, evo_step_tracker[num_poss_outputs];
        float output_parasite_density[num_poss_outputs], output_hyperparasite_density[num_poss_outputs], output_host_density[num_poss_outputs];
        bool output_hyperparasite_truths[num_poss_outputs];
        
        ind = 0;
        ofstream myfile, tracker_file;

        for(i=0;i<trait_space_length;i++){
            alpha[i] = (alpha_max*(i))/(trait_space_length - 1);
            beta_0[i] = beta_max*(1 - alpha[i]/alpha_max);
            sigma[i] = (sigma_max*(i))/(trait_space_length - 1);
        }

        // float alpha[trait_space_length] = {0,0.0700000000000000,0.140000000000000,0.210000000000000,0.280000000000000,0.350000000000000,0.420000000000000,0.490000000000000,0.560000000000000,0.630000000000000,0.700000000000000,0.770000000000000,0.840000000000000,0.910000000000000,0.980000000000000,1.05000000000000,1.12000000000000,1.19000000000000,1.26000000000000,1.33000000000000,1.40000000000000,1.47000000000000,1.54000000000000,1.61000000000000,1.68000000000000,1.75000000000000,1.82000000000000,1.89000000000000,1.96000000000000,2.03000000000000,2.10000000000000,2.17000000000000,2.24000000000000,2.31000000000000,2.38000000000000,2.45000000000000,2.52000000000000,2.59000000000000,2.66000000000000,2.73000000000000,2.80000000000000,2.87000000000000,2.94000000000000,3.01000000000000,3.08000000000000,3.15000000000000,3.22000000000000,3.29000000000000,3.36000000000000,3.43000000000000,3.50000000000000,3.57000000000000,3.64000000000000,3.71000000000000,3.78000000000000,3.85000000000000,3.92000000000000,3.99000000000000,4.06000000000000,4.13000000000000,4.20000000000000,4.27000000000000,4.34000000000000,4.41000000000000,4.48000000000000,4.55000000000000,4.62000000000000,4.69000000000000,4.76000000000000,4.83000000000000,4.90000000000000,4.97000000000000,5.04000000000000,5.11000000000000,5.18000000000000,5.25000000000000,5.32000000000000,5.39000000000000,5.46000000000000,5.53000000000000,5.60000000000000,5.67000000000000,5.74000000000000,5.81000000000000,5.88000000000000,5.95000000000000,6.02000000000000,6.09000000000000,6.16000000000000,6.23000000000000,6.30000000000000,6.37000000000000,6.44000000000000,6.51000000000000,6.58000000000000,6.65000000000000,6.72000000000000,6.79000000000000,6.86000000000000,6.93000000000000,7};
        // float beta_temp[trait_space_length] = {0,0.269866633728588,0.381649053450942,0.467422720885496,0.539733267457177,0.603440137876161,0.661035551237602,0.714000000000000,0.763298106901884,0.809599901185765,0.853393227064757,0.895046367513997,0.934845441770991,0.973017985445285,1.00974848353439,1.04518897812788,1.08174993001123,1.11398486442480,1.14638859171872,1.17896379156107,1.21171321528104,1.24463968858098,1.27774611438184,1.31103547581003,1.34451083933467,1.37817535806417,1.41203227521258,1.44608492774621,1.48033675022225,1.51479127883181,1.54945215566102,1.58432313318445,1.61940807900684,1.65471098086984,1.69023595194230,1.72598723641390,1.76196921541359,1.79818641327631,1.83464350418336,1.87134531920391,1.90829685376775,1.94550327560205,1.98296993316774,2.02070236463454,2.05870630743728,2.09698770846027,2.13555273490069,2.17440778586753,2.21355950477749,2.25301479261611,2.29278082213894,2.33286505309559,2.37327524856808,2.41401949252494,2.45510620870343,2.49654418094479,2.53834257512155,2.58051096281202,2.62305934689515,2.64035532222432,2.65750533212608,2.67451301091134,2.69138184453413,2.70811517893416,2.72471622778583,2.74118807970432,2.75753370495463,2.77375596170493,2.78985760186130,2.80584127651775,2.82170954105203,2.83746485989496,2.85310961099853,2.86864609002562,2.88407651428237,2.89940302641208,2.91462769786832,2.92975253218294,2.94477946804375,2.95971038219513,2.97454709217401,2.98929135889229,3.00394488907635,3.01850933757295,3.03298630953059,3.04737736246425,3.06168400821122,3.07590771478477,3.09004990813233,3.10411197380390,3.11809525853639,3.13200107175895,3.14583068702407,3.15958534336878,3.17326624661029,3.18687457057969,3.20041145829730,3.21387802309324,3.22727534967592,3.24060449515180,3.25386648999875};
        
        for(i=0;i<trait_space_length;i++){
            for(j=0;j<trait_space_length;j++){
                beta[i*trait_space_length + j] = beta_0[i]*(1 - 1/(1 + exp(-alpha[i]/alpha_max)));
                // if (c2 == 0.0){
                //     beta[i*trait_space_length + j] = beta_temp[i]*((1-c1) + c1*(sigma[j]/sigma_max));
                // }
                // else{
                //     beta[i*trait_space_length+j] = fastmax(0,beta_temp[i]*((1-c1) + c1*(1 - exp((sigma[j]*c2)/sigma_max))/(1 - exp(c2))));
                // }
            }
        }

        ofstream beta_values,sigma_values, alpha_values;
        beta_values.open("../data/beta_vals.csv");
        sigma_values.open("../data/sigma_vals.csv");
        alpha_values.open("../data/alpha_vals.csv");

        for (i = 0; i<trait_space_length;i++){
            beta_values << i << ",";
            alpha_values << i << ",";
            sigma_values << i << ",";
        }
        beta_values << "\n";
        alpha_values << "\n";
        sigma_values << "\n";

        

        for(i=0;i<trait_space_length;i++){
            alpha_values << alpha[i] << ",";
            sigma_values << sigma[i] << ",";
            for(j=0;j<trait_space_length;j++){
                beta_values << beta[i*trait_space_length+j] << ",";
            }
            beta_values << "\n";
        }

        alpha_values << "\n";
        sigma_values << "\n";

        for (i = 0; i<trait_space_length;i++){
            sigma_values << ((1-c1) + c1*(1 - exp((sigma[i]*c2)/sigma_max))/(1 - exp(c2))) << ",";
            alpha_values << sqrt(alpha[i]/alpha_max) << ",";
        }

        alpha_values << "\n";
        sigma_values << "\n";

        beta_values.close();
        sigma_values.close();
        alpha_values.close();

        // y[0] = (4.0)/(b/q);
        // y[1] = (4.0)/(b/q);
        // if (hyper > 0.0){
        //     y[2] = (4.0)/(b/q);
        // }
        // else{
        //     y[2] = 0.0;
        // }
        y[0] = S_density;
        y[1] = I_density;
        if (hyper > 0.0){
            y[2] = H_density;
        }
        else{
            y[2] = 0.0;
        }

        alpha_inds[0] = alpha_init;
        sigma_inds[0] = sigma_init;
        
        num_strains = 1;
        for (int evo_counter = 0; evo_counter < evo_steps; evo_counter++){
            
            t[0] = 0.0;
            h[0] = 1e-1;
            hnext[0] = 1e-1;

            float y_check_sum = 0.0;
            
            tcheck[0] = 10.0;
            for(i=0;i<(2*num_strains + 1);i++){
                y_check[i] = y[i];
                y_max[i] = y[i];
                y_min[i] = y[i];
                y_max_next[i] = y[i];
                y_min_next[i] = y[i];
                y_check_sum += y[i];
                // y_max[i] = y[i];
                // y_min[i] = y[i];
                // std::cout << "y value pre-solver is " << y[i] << endl;
            }
            int step_count = 0;
            int check_count = 0;
            while (t[0] <= tmax){   
                int counter[1],num_plus[1],num_down[1];
                
                counter[0] = 0;
                num_plus[0] = 0;
                num_down[0] = 0;

                // h[0] = 1.0;
                // grad_max = 0.0;
                /* This is where the equations are first solved */
                dynamics(dydt, y, num_strains, alpha_inds, sigma_inds, b, q, d, beta, alpha, eta, gamma, sigma, lambda, rho);

                /* Adjust the step size to maintain accuracy */
                for (i=0; i<(2*num_strains + 1); i++){
                    yScale[i]=fabs(y[i])+fabs(dydt[i]*(*h))+EPS;
                    // grad_max = Quick_solver::fastmax(grad_max, abs(dydt[i]));
                }
                
                rkqs(y, dydt, h,hnext,yScale, num_strains, alpha_inds, sigma_inds, b, q, d, beta, alpha, eta, gamma, sigma, lambda, rho,counter,num_plus,num_down,discrepancy, int_tracker, t);
                
                if (y[0] <= TINY){
                    y[0] = 0.0;
                }

                for(i=0;i<(num_strains);i++){
                    if (y[i+1] + y[i + num_strains + 1] <= TINY){
                        y[i+1] = 0.0;
                        y[i + num_strains + 1] = 0.0;
                    }
                    if(y[i + num_strains + 1] <= TINY){
                        y[i + num_strains + 1] = 0.0;
                    }
                }

                t[0]+=h[0];
                h[0] = hnext[0];
                
                for(i=0;i<(2*num_strains + 1);i++){
                    y_max[i] = fastmax(y[i], y_max[i]);
                    y_min[i] = fastmin(y[i], y_min[i]);
                }
                step_count += 1;
                
                if ((((t[0] - tcheck[0]) >= 10.0) && (step_count - check_count) >= 200) && (y[0] != 0.0)){
                    discrep_check = 0.0;
                    check_count = step_count;
                    
                    for(i=0;i<(2*num_strains + 1);i++){

                        if (discrep_check < fastmax(abs(y_max[i] - y_max_next[i]),abs(y_min[i] - y_min_next[i]))){
                            
                            discrep_check = fastmax(abs(y_max[i] - y_max_next[i]),abs(y_min[i] - y_min_next[i]));
                        }
                    }

                    if (discrep_check >= 1e-3){

                        y_check_sum = 0.0;
                        tcheck[0] = t[0];

                        for(i=0;i<(2*num_strains + 1);i++){
                            y_check[i] = y[i];
                            y_max[i] = y[i];
                            y_min[i] = y[i];
                            y_max_next[i] = y[i];
                            y_min_next[i] = y[i];
                        }
                    }
                    else{
                        t[0] = tmax + 1.0;
                    }
                }
            }

            if (y[0] <= TINY){
                std::cout << "Healthy hosts extinct, breaking" << endl;
                break;
            }
            
            num_strains_cleaned = 0;
            for(i=0;i<num_strains;i++){
                if ((y[i + 1] + y[i + 1 + (num_strains)]) > TINY){
                    num_strains_cleaned = num_strains_cleaned + 1;
                }
            }

            if (num_strains_cleaned == 0){
                cout << "All parasites are dead\n";
                break;
            }

            float rand_stored1, rand_stored2;

            counter = 0;

            Hsum = 0.0;

            for (i=0;i<num_strains;i++){
                if((y[i + 1] + y[i + 1 + num_strains]) > TINY){
                    alpha_inds_cleaned[counter] = alpha_inds[i];
                    sigma_inds_cleaned[counter] = sigma_inds[i];
                    I_temp[counter] = y[i +1];
                    H_temp[counter] = y[i +1 + num_strains];
                    total_density[counter] = I_temp[counter] + H_temp[counter];
                    Hsum = Hsum + H_temp[counter];
                    counter++;
                }
            }
            
            if (Hsum <= 1e-10){
                Hsum = 0.0;
                for(i=0;i<(num_strains);i++){
                    y[i + num_strains + 1] = 0.0;
                }
            }
            cum_density[0] = total_density[0];
            if (num_strains_cleaned > 1){
                for(i=1;i<num_strains_cleaned;i++){
                    cum_density[i] = total_density[i] + cum_density[i - 1];
                }
            }

            for(i=0;i<num_strains_cleaned;i++){
                cum_props[i] = cum_density[i]/cum_density[num_strains_cleaned-1];
            }

            rand_stored1 = (float(rand())/float((RAND_MAX)));
            rand_stored2 = (float(rand())/float((RAND_MAX)));

            for(i=0;i<num_strains_cleaned;i++){
                if(cum_props[i] >= rand_stored1){
                    ind = i;
                    break;
                }
            }

            bool flag = true;
            int increment_ind;
            num_strains_2 = num_strains_cleaned;
            
            if (((rand_stored2 >=0.5) && (alpha_inds_cleaned[ind] != (trait_space_length-1))) || (alpha_inds_cleaned[ind] == 0)){
                for(i=0;i<num_strains_cleaned;i++){
                    if (alpha_inds_cleaned[i] == (alpha_inds_cleaned[ind] + 1)){
                        flag = false;
                        increment_ind = i;
                        // std::cout << "Alpha increment ind is" << increment_ind << endl;
                        break;
                    }
                }
            }
            else{
                for(i=0;i<num_strains_cleaned;i++){
                    if (alpha_inds_cleaned[i] == (alpha_inds_cleaned[ind] - 1) && (sigma_inds_cleaned[i] == sigma_inds_cleaned[ind])){
                        flag = false;
                        increment_ind = i;
                        break;
                    }
                }
            }
            // std::cout << "\n num_strains_2 after incrementing in alpha step" << endl;
            // int num_strains_2 = num_strains_cleaned;
            if(flag){
                num_strains_2++;
            }
            // std::cout << num_strains_2 << endl;

            y_temp[0] = y[0];
            for(i = 0; i<num_strains_2;i++){
                y_temp[i+1] = 0.0;
                y_temp[i+1+num_strains_2] = 0.0;
            }
            for(i=0;i<num_strains_cleaned;i++){
                alpha_inds_2[i] = alpha_inds_cleaned[i];
                sigma_inds_2[i] = sigma_inds_cleaned[i];
                y_temp[i+1] = I_temp[i];
                y_temp[i+1+num_strains_2] = H_temp[i];
            }

            // std::cout << "Reallocated number of strains" << endl;
            num_strains = num_strains_2;

            for(i = 0; i<(2*num_strains+1); i++){
                y[i] = 0.0;
            }

            y[0] = y_temp[0];
        
            for(i=0;i<num_strains;i++){
                alpha_inds[i] = alpha_inds_2[i];
                sigma_inds[i] = sigma_inds_2[i];
                y[i+1] = y_temp[i+1];
                y[i+1+num_strains] = y_temp[i+1+num_strains];
            }

            if (((rand_stored2 >=0.5) && (alpha_inds_cleaned[ind] != trait_space_length-1)) || (alpha_inds_cleaned[ind] == 0)){
                if(flag){
                    alpha_inds[num_strains-1] = alpha_inds_2[ind] + 1;
                    sigma_inds[num_strains-1] = sigma_inds_2[ind];
                    y[num_strains] = (y_temp[ind+1]/100);
                    y[2*num_strains] = (y_temp[ind+1+num_strains]/100);
                }
                else{
                    y[1+increment_ind] = y[1+increment_ind] + (y_temp[ind+1]/100);
                    y[1+increment_ind+num_strains] = y[1+increment_ind+num_strains] + (y_temp[ind+1+num_strains]/100);
                }
            }
            else{
                if(flag){
                    alpha_inds[num_strains-1] = alpha_inds_2[ind] - 1;
                    sigma_inds[num_strains-1] = sigma_inds_2[ind];
                    y[num_strains] = (y_temp[ind+1]/100);
                    y[2*num_strains] = (y_temp[ind+1+num_strains]/100);
                }
                else{
                    y[1+increment_ind] = y[1+increment_ind] + (y_temp[ind+1]/100);
                    y[1+increment_ind+num_strains] = y[1+increment_ind+num_strains] + (y_temp[ind+1+num_strains]/100);
                }
            }
            
            for(i=0;i<num_strains;i++){
                output_alpha_inds[output_counter] = alpha_inds[i];
                output_sigma_inds[output_counter] = sigma_inds[i];
                output_parasite_density[output_counter] = y[i+1];
                output_hyperparasite_density[output_counter] = y[i+1+num_strains];
                output_hyperparasite_truths[output_counter] = (Hsum > 0.0);
                evo_step_tracker[output_counter] = evo_counter + 1;
                output_host_density[output_counter] = y[0];
                output_counter++;
            }

            if (evo_counter%100 == 0){
                std::cout << evo_counter << "\n";
            }
        }
        
        string test;

        string seed_str;
        seed_str = std::to_string(seed);

        tracker_file.open("../data/tracker_file.csv", std::ios_base::app);
        tracker_file << "/alpha_evo/data_set" + seed_str + ".csv," << b << "," << q << "," << d << "," << rho << "," << sigma_max << "," << gamma;
        tracker_file << "," << eta << "," << lambda << "," << c1 << "," << c2 << "," << beta_max << "," << (hyper > 0.0) << "," << alpha_init << "," << sigma_init  << "\n";
        tracker_file.close();

        myfile.open("../data/alpha_evo/data_set"+seed_str+".csv");
        myfile << "Trait_index_1,Trait_index_2,Density_of_Hosts,Density_of_parasite,Density_of_hyperparasite,Evolutionary_step,hyperparasites_present,alpha_val,sigma_val,beta_val\n";
        
        for(i=0;i<output_counter;i++){
            myfile << output_alpha_inds[i] <<"," << output_sigma_inds[i] << "," << output_host_density[i] << "," << output_parasite_density[i] << ',' << output_hyperparasite_density[i] << ","<< evo_step_tracker[i] << "," << output_hyperparasite_truths[i] << "," << alpha[output_alpha_inds[i]] << "," << sigma[output_sigma_inds[i]] << "," << beta[output_alpha_inds[i]*trait_space_length + output_sigma_inds[i]] << "\n";
        }

        myfile.close();
        
        delete y_check;
        delete y;
        delete yScale;
        delete dydt;
        delete y_temp;
        time (&end);
        float dif = difftime (end,start);
        printf ("Elasped time is %.2lf seconds.\n", dif ); 
        return;
    }
    
    void Quick_solver::alpha_evo_only_v4(float beta_max, float beta_lin, float alpha_max, float alpha_min, float sigma_max, float b, float q, float d, float rho, float eta, float gamma, float lambda, float c1, float c2, float beta_scalar, float hyper, int seed, int alpha_init, int sigma_init, int evo_steps, float S_density, float I_density, float H_density){
        
        // time_t start,end;
        // time (&start);
        

        srand(seed);
        
        int tmax = 1000, i, j;
        float beta[trait_space_length*trait_space_length], alpha[trait_space_length], sigma[trait_space_length];
        float t[1], tcheck[1], Hsum;
        int num_strains = 1, alpha_inds[trait_space_length], ind,num_strains_2;
        int sigma_inds[trait_space_length], num_strains_cleaned;

        float* y = new float[2*trait_space_length+1];
        float* y_check = new float[2*trait_space_length + 1];
        float* dydt = new float[2*trait_space_length + 1];
        float* yScale = new float[2*trait_space_length + 1];
        
        float* y_max = new float[2*trait_space_length + 1];
        float* y_min = new float[2*trait_space_length + 1];
        float* y_max_next = new float[2*trait_space_length + 1];
        float* y_min_next = new float[2*trait_space_length + 1];
        
        float alpha_scalar, sigma_scalar;
        float discrep_check;
        int alpha_inds_cleaned[trait_space_length],counter=0;
        int sigma_inds_cleaned[trait_space_length];
        float I_temp[trait_space_length], H_temp[trait_space_length],total_density[trait_space_length];
        float cum_density[trait_space_length], cum_props[trait_space_length];
        int alpha_inds_2[trait_space_length];
        int sigma_inds_2[trait_space_length], int_tracker[1];
        float* y_temp = new float[2*trait_space_length+1];
        
        float h[1], hnext[1], discrepancy[1];

        int num_poss_outputs = 50000;
        int output_alpha_inds[num_poss_outputs], output_sigma_inds[num_poss_outputs], output_counter = 0, evo_step_tracker[num_poss_outputs];
        float output_parasite_density[num_poss_outputs], output_hyperparasite_density[num_poss_outputs], output_host_density[num_poss_outputs];
        bool output_hyperparasite_truths[num_poss_outputs];
        
        ind = 0;
        ofstream myfile, tracker_file;

        for(i=0;i<trait_space_length;i++){
            alpha[i] = alpha_min + ((alpha_max - alpha_min)*(i))/(trait_space_length - 1);
            sigma[i] = (sigma_max*(i))/(trait_space_length - 1);
        }

        for(i=0;i<trait_space_length;i++){
            if (c2 == 0.0){
                alpha_scalar = (1-c1) + c1*(alpha[i]/alpha_max);
            }
            else{
                alpha_scalar = (1-c1) + c1*(1 - exp(c2*(alpha[i]/alpha_max)))/(1 - exp(c2));
            }
            
            for(j=0;j<trait_space_length;j++){    
                sigma_scalar = beta_scalar;
                float beta_temp = beta_max*alpha_scalar + beta_lin*alpha[i];
                // float beta_temp = (beta_max - beta_min)*alpha_scalar;
                beta[i*trait_space_length+j] = fastmax(0,beta_temp);
            }
        }

        ofstream beta_values,sigma_values, alpha_values;
        beta_values.open("../data/beta_vals.csv");
        sigma_values.open("../data/sigma_vals.csv");
        alpha_values.open("../data/alpha_vals.csv");

        for (i = 0; i<trait_space_length;i++){
            beta_values << i << ",";
            alpha_values << i << ",";
            sigma_values << i << ",";
        }
        beta_values << "\n";
        alpha_values << "\n";
        sigma_values << "\n";

        

        for(i=0;i<trait_space_length;i++){
            sigma_values << sigma[i] << ",";
            alpha_values << alpha[i] << ",";
            for(j=0;j<trait_space_length;j++){
                beta_values << beta[i*trait_space_length+j] << ",";
            }
            beta_values << "\n";
        }

        alpha_values << "\n";
        sigma_values << "\n";

        for (i = 0; i<trait_space_length;i++){
            sigma_values << beta_scalar << ",";
            alpha_values << ((1-c1) + (c1*(1 - exp((alpha[i]*c2)/alpha_max))/(1 - exp(c2)))) << ",";
        }

        alpha_values << "\n";
        sigma_values << "\n";

        beta_values.close();
        sigma_values.close();
        alpha_values.close();

        // y[0] = (4.0)/(b/q);
        // y[1] = (4.0)/(b/q);
        // if (hyper > 0.0){
        //     y[2] = (4.0)/(b/q);
        // }
        // else{
        //     y[2] = 0.0;
        // }
        y[0] = S_density;
        y[1] = I_density;
        if (hyper > 0.0){
            y[2] = H_density;
        }
        else{
            y[2] = 0.0;
        }

        alpha_inds[0] = alpha_init;
        sigma_inds[0] = sigma_init;
        
        num_strains = 1;
        for (int evo_counter = 0; evo_counter < evo_steps; evo_counter++){
            
            t[0] = 0.0;
            h[0] = 1e-1;
            hnext[0] = 1e-1;

            float y_check_sum = 0.0;
            
            tcheck[0] = 10.0;
            for(i=0;i<(2*num_strains + 1);i++){
                y_check[i] = y[i];
                y_max[i] = y[i];
                y_min[i] = y[i];
                y_max_next[i] = y[i];
                y_min_next[i] = y[i];
                y_check_sum += y[i];
                // y_max[i] = y[i];
                // y_min[i] = y[i];
                // std::cout << "y value pre-solver is " << y[i] << endl;
            }
            int step_count = 0;
            int check_count = 0;
            while (t[0] <= tmax){   
                int counter[1],num_plus[1],num_down[1];
                
                counter[0] = 0;
                num_plus[0] = 0;
                num_down[0] = 0;

                // h[0] = 1.0;
                // grad_max = 0.0;
                /* This is where the equations are first solved */
                dynamics(dydt, y, num_strains, alpha_inds, sigma_inds, b, q, d, beta, alpha, eta, gamma, sigma, lambda, rho);

                /* Adjust the step size to maintain accuracy */
                for (i=0; i<(2*num_strains + 1); i++){
                    yScale[i]=fabs(y[i])+fabs(dydt[i]*(*h))+EPS;
                    // grad_max = Quick_solver::fastmax(grad_max, abs(dydt[i]));
                }
                
                rkqs(y, dydt, h,hnext,yScale, num_strains, alpha_inds, sigma_inds, b, q, d, beta, alpha, eta, gamma, sigma, lambda, rho,counter,num_plus,num_down,discrepancy, int_tracker, t);
                
                if (y[0] <= TINY){
                    y[0] = 0.0;
                }

                for(i=0;i<(num_strains);i++){
                    if (y[i+1] + y[i + num_strains + 1] <= TINY){
                        y[i+1] = 0.0;
                        y[i + num_strains + 1] = 0.0;
                    }
                    if(y[i + num_strains + 1] <= TINY){
                        y[i + num_strains + 1] = 0.0;
                    }
                }

                t[0]+=h[0];
                h[0] = hnext[0];
                
                for(i=0;i<(2*num_strains + 1);i++){
                    y_max[i] = fastmax(y[i], y_max[i]);
                    y_min[i] = fastmin(y[i], y_min[i]);
                }
                step_count += 1;
                
                if ((((t[0] - tcheck[0]) >= 10.0) && (step_count - check_count) >= 200) && (y[0] != 0.0)){
                    discrep_check = 0.0;
                    check_count = step_count;
                    
                    for(i=0;i<(2*num_strains + 1);i++){

                        if (discrep_check < fastmax(abs(y_max[i] - y_max_next[i]),abs(y_min[i] - y_min_next[i]))){
                            
                            discrep_check = fastmax(abs(y_max[i] - y_max_next[i]),abs(y_min[i] - y_min_next[i]));
                        }
                    }

                    if (discrep_check >= 1e-3){

                        y_check_sum = 0.0;
                        tcheck[0] = t[0];

                        for(i=0;i<(2*num_strains + 1);i++){
                            y_check[i] = y[i];
                            y_max[i] = y[i];
                            y_min[i] = y[i];
                            y_max_next[i] = y[i];
                            y_min_next[i] = y[i];
                        }
                    }
                    else{
                        t[0] = tmax + 1.0;
                    }
                }
            }

            if (y[0] <= TINY){
                std::cout << "Healthy hosts extinct, breaking" << endl;
                break;
            }
            
            num_strains_cleaned = 0;
            for(i=0;i<num_strains;i++){
                if ((y[i + 1] + y[i + 1 + (num_strains)]) > TINY){
                    num_strains_cleaned = num_strains_cleaned + 1;
                }
            }

            if (num_strains_cleaned == 0){
                cout << "All parasites are dead\n";
                break;
            }

            float rand_stored1, rand_stored2;

            counter = 0;

            Hsum = 0.0;

            for (i=0;i<num_strains;i++){
                if((y[i + 1] + y[i + 1 + num_strains]) > TINY){
                    alpha_inds_cleaned[counter] = alpha_inds[i];
                    sigma_inds_cleaned[counter] = sigma_inds[i];
                    I_temp[counter] = y[i +1];
                    H_temp[counter] = y[i +1 + num_strains];
                    total_density[counter] = I_temp[counter] + H_temp[counter];
                    Hsum = Hsum + H_temp[counter];
                    counter++;
                }
            }

            bool HSum_flag = (Hsum >= TINY);
            
            cum_density[0] = total_density[0];
            if (num_strains_cleaned > 1){
                for(i=1;i<num_strains_cleaned;i++){
                    cum_density[i] = total_density[i] + cum_density[i - 1];
                }
            }

            for(i=0;i<num_strains_cleaned;i++){
                cum_props[i] = cum_density[i]/cum_density[num_strains_cleaned-1];
            }

            rand_stored1 = (float(rand())/float((RAND_MAX)));
            rand_stored2 = (float(rand())/float((RAND_MAX)));

            for(i=0;i<num_strains_cleaned;i++){
                if(cum_props[i] >= rand_stored1){
                    ind = i;
                    break;
                }
            }

            bool flag = true;
            int increment_ind;
            num_strains_2 = num_strains_cleaned;
            
            if (((rand_stored2 >=0.5) && (alpha_inds_cleaned[ind] != (trait_space_length-1))) || (alpha_inds_cleaned[ind] == 0)){
                for(i=0;i<num_strains_cleaned;i++){
                    if (alpha_inds_cleaned[i] == (alpha_inds_cleaned[ind] + 1)){
                        flag = false;
                        increment_ind = i;
                        // std::cout << "Alpha increment ind is" << increment_ind << endl;
                        break;
                    }
                }
            }
            else{
                for(i=0;i<num_strains_cleaned;i++){
                    if (alpha_inds_cleaned[i] == (alpha_inds_cleaned[ind] - 1) && (sigma_inds_cleaned[i] == sigma_inds_cleaned[ind])){
                        flag = false;
                        increment_ind = i;
                        break;
                    }
                }
            }
            // std::cout << "\n num_strains_2 after incrementing in alpha step" << endl;
            // int num_strains_2 = num_strains_cleaned;
            if(flag){
                num_strains_2++;
            }
            // std::cout << num_strains_2 << endl;

            y_temp[0] = y[0];
            for(i = 0; i<num_strains_2;i++){
                y_temp[i+1] = 0.0;
                y_temp[i+1+num_strains_2] = 0.0;
            }
            for(i=0;i<num_strains_cleaned;i++){
                alpha_inds_2[i] = alpha_inds_cleaned[i];
                sigma_inds_2[i] = sigma_inds_cleaned[i];
                y_temp[i+1] = I_temp[i];
                y_temp[i+1+num_strains_2] = H_temp[i]*HSum_flag;
            }

            // std::cout << "Reallocated number of strains" << endl;
            num_strains = num_strains_2;

            for(i = 0; i<(2*num_strains+1); i++){
                y[i] = 0.0;
            }

            y[0] = y_temp[0];
        
            for(i=0;i<num_strains;i++){
                alpha_inds[i] = alpha_inds_2[i];
                sigma_inds[i] = sigma_inds_2[i];
                y[i+1] = y_temp[i+1];
                y[i+1+num_strains] = y_temp[i+1+num_strains];
            }

            if (((rand_stored2 >=0.5) && (alpha_inds_cleaned[ind] != trait_space_length-1)) || (alpha_inds_cleaned[ind] == 0)){
                if(flag){
                    alpha_inds[num_strains-1] = alpha_inds_2[ind] + 1;
                    sigma_inds[num_strains-1] = sigma_inds_2[ind];
                    y[num_strains] = (y_temp[ind+1]/100);
                    y[2*num_strains] = (y_temp[ind+1+num_strains]/100);
                }
                else{
                    y[1+increment_ind] = y[1+increment_ind] + (y_temp[ind+1]/100);
                    y[1+increment_ind+num_strains] = y[1+increment_ind+num_strains] + (y_temp[ind+1+num_strains]/100);
                }
            }
            else{
                if(flag){
                    alpha_inds[num_strains-1] = alpha_inds_2[ind] - 1;
                    sigma_inds[num_strains-1] = sigma_inds_2[ind];
                    y[num_strains] = (y_temp[ind+1]/100);
                    y[2*num_strains] = (y_temp[ind+1+num_strains]/100);
                }
                else{
                    y[1+increment_ind] = y[1+increment_ind] + (y_temp[ind+1]/100);
                    y[1+increment_ind+num_strains] = y[1+increment_ind+num_strains] + (y_temp[ind+1+num_strains]/100);
                }
            }
            
            for(i=0;i<num_strains;i++){
                output_alpha_inds[output_counter] = alpha_inds[i];
                output_sigma_inds[output_counter] = sigma_inds[i];
                output_parasite_density[output_counter] = y[i+1];
                output_hyperparasite_density[output_counter] = y[i+1+num_strains];
                output_hyperparasite_truths[output_counter] = (Hsum > 0.0);
                evo_step_tracker[output_counter] = evo_counter + 1;
                output_host_density[output_counter] = y[0];
                output_counter++;
            }

            // if (evo_counter%100 == 0){
            //     std::cout << evo_counter << "\n";
            // }
        }
        
        string test;

        string seed_str;
        seed_str = std::to_string(seed);

        tracker_file.open("../data/tracker_file.csv", std::ios_base::app);
        tracker_file << "/alpha_evo/data_set" + seed_str + ".csv," << b << "," << q << "," << d << "," << rho << "," << sigma_max << "," << gamma;
        tracker_file << "," << eta << "," << lambda << "," << c1 << "," << c2 << "," << beta_max << "," << (hyper > 0.0) << "," << alpha_init << "," << sigma_init  << "\n";
        tracker_file.close();

        myfile.open("../data/alpha_evo/data_set"+seed_str+".csv");
        myfile << "Trait_index_1,Trait_index_2,Density_of_Hosts,Density_of_parasite,Density_of_hyperparasite,Evolutionary_step,hyperparasites_present,alpha_val,sigma_val,beta_val\n";
        
        for(i=0;i<output_counter;i++){
            myfile << output_alpha_inds[i] <<"," << output_sigma_inds[i] << "," << output_host_density[i] << "," << output_parasite_density[i] << ',' << output_hyperparasite_density[i] << ","<< evo_step_tracker[i] << "," << output_hyperparasite_truths[i] << "," << alpha[output_alpha_inds[i]] << "," << sigma[output_sigma_inds[i]] << "," << beta[output_alpha_inds[i]*trait_space_length + output_sigma_inds[i]] << "\n";
        }

        myfile.close();
        
        delete y_check;
        delete y;
        delete yScale;
        delete dydt;
        delete y_temp;
        // time (&end);
        // float dif = difftime (end,start);
        // printf ("Elasped time is %.2lf seconds.\r", dif ); 
        return;
    }
};
```
