## Supplementary material for "Hyperparasitism and the evolution of parasite virulence": Code_base: wrapper.h

```
    #ifndef WRAPPER_H
    #define WRAPPER_H

    namespace solvers{

        class Quick_solver{

            public:
                float beta_max, alpha_max, sigma_max, b, q, d, rho, eta, gamma, lambda, c1, c2, hyper;
                int seed;
                Quick_solver();
                Quick_solver(float beta_max, float alpha_max, float sigma_max, float b, float q, float d, float rho, float eta, float gamma, float lambda, float c1, float c2, float hyper, int seed);
                ~Quick_solver();
                // float TINY = 1e-3;
                // float EPS = 1e-5;
                float fastmax(float a, float b);
                float fastmin(float a, float b);
                int dynamics(float* dydt, float* y, int num_strains, int* alpha_inds, int* sigma_inds, float b, float q, float d, float* beta, float* alpha, float eta, float gamma, float* sigma, float lambda, float rho);
                int RK45(float* y_out, float* y, float* y_err, int num_strains, int* alpha_inds, int* sigma_inds, float b, float q, float d, float* beta, float* alpha, float eta, float gamma, float* sigma, float lambda, float rho, float* h);
                int perform_RK45_step(float* y, float* t, int num_strains, int* alpha_inds, int* sigma_inds, float b, float q, float d, float* beta, float* alpha, float eta, float gamma, float* sigma, float lambda, float rho, float* h, float* discrepancy, int* counter, int* num_plus, int* num_down);
                int rkck(float *y, float *dydt, float *yout, float *yerr, float h, int num_strains, int* alpha_inds, int* sigma_inds, float b, float q, float d, float* beta, float* alpha, float eta, float gamma, float* sigma, float lambda, float rho);
                int rkqs(float *y, float *dydt, float *h,float *hnext,float *yscale, int num_strains, int* alpha_inds, int* sigma_inds, float b, float q, float d, float* beta, float* alpha, float eta, float gamma, float* sigma, float lambda, float rho, int* counter, int* num_up, int* num_down, float* discrepancy, int* int_tracker, float* t);
                int ad_dyn(float beta_max, float alpha_max, float sigma_max, float b, float q, float d, float rho, float eta, float gamma, float lambda, float c1, float c2, float hyper, int seed, int alpha_init, int sigma_init);
                void eco_dynamics(float* S, float* I, float* H, int* exit_flag, float beta_max, float alpha_max, float sigma_max, float b, float q, float d, float rho, float eta, float gamma, float lambda, float c1, float c2, float hyper, int seed, float S_density, float I_density, float H_density);
                void alpha_evo_only(float beta_max, float alpha_max, float sigma_max, float b, float q, float d, float rho, float eta, float gamma, float lambda, float c1, float c2, float hyper, int seed, int alpha_init, int sigma_init, float S_density, float I_density, float H_density);
                void alpha_evo_only_v2(float beta_max, float beta_min, float alpha_max, float alpha_min, float sigma_max, float b, float q, float d, float rho, float eta, float gamma, float lambda, float c1, float c2, float beta_scalar,  float hyper, int seed, int alpha_init, int sigma_init, float S_density, float I_density, float H_density);
                void alpha_evo_only_v3(float beta_max, float alpha_max, float sigma_max, float b, float q, float d, float rho, float eta, float gamma, float lambda, float c1, float c2, float hyper, int seed, int alpha_init, int sigma_init, float S_density, float I_density, float H_density);
                void alpha_evo_only_v4(float beta_max, float beta_lin, float alpha_max, float alpha_min, float sigma_max, float b, float q, float d, float rho, float eta, float gamma, float lambda, float c1, float c2, float beta_scalar,  float hyper, int seed, int alpha_init, int sigma_init, int evo_steps, float S_density, float I_density, float H_density);
        };
    }


    #endif
```
