## Supplementary material for "Hyperparasitism and the evolution of parasite virulence": Code_base: figure_utilities.py

```
import plotly.graph_objects as go


def make_styled_figure(xtitle="", ytitle=""):

    fig = go.Figure()

    fig.update_layout(plot_bgcolor="White", showlegend=False)

    fig.update_xaxes(title="<b>" + xtitle + "<b>")
    fig.update_yaxes(title="<b>" + ytitle + "<b>")

    return fig
```
