## Supplementary material for "Hyperparasitism and the evolution of parasite virulence": Code_base: functions.py

```
from scipy.integrate import solve_ivp
import numpy as np


def H_P_HP_model(
    t,
    y,
    para_inds,
    b: float = 3,
    q: float = 0.05,
    d: float = 0.5,
    beta: float = [],
    alpha: float = [],
    eta: float = 0.9,
    gamma: float = 1.5,
    sigma: float = 2,
    lam: float = 0.75,
    rho: float = 0.5,
):

    S = y[0]
    Is = y[1 : len(para_inds) + 1]
    Hs = y[len(para_inds) + 1 :]

    Isum = sum(Is)
    Hsum = sum(Hs)
    y_dot = []

    N = S + Isum + Hsum

    S_dot = (
        (b - q * N) * N
        - (
            sum([beta[para_inds[i]] * I for i, I in enumerate(Is)])
            + sum([eta * beta[para_inds[i]] * H for i, H in enumerate(Hs)])
            - d
        )
        * S
        + gamma * (Isum + Hsum)
    )

    I_dots = []
    for i, I in enumerate(Is):
        I_dots.append(
            (beta[para_inds[i]] * S - sigma * Hsum - (d + alpha[para_inds[i]] + gamma))
            * I
            + (1 - rho) * eta * beta[para_inds[i]] * Hs[i] * S
        )

    H_dots = []
    for i, H in enumerate(Hs):
        H_dots.append(
            (
                rho * eta * beta[para_inds[i]] * S
                - (d + lam * alpha[para_inds[i]] + gamma)
            )
            * H
            + sigma * Is[i] * Hsum
        )

    y_dot = [S_dot]

    for val in I_dots:
        y_dot.append(val)

    for val in H_dots:
        y_dot.append(val)

    return y_dot


def ad_dyn_routine_HPHP(
    evo_steps,
    function,
    parameters,
    initial_conditions,
    trange: list = [0, 1000],
    verbose: bool = False,
):

    output = []

    y = initial_conditions

    parameters_temp = parameters

    for step in range(evo_steps):
        temp_output_row = []
        sol = solve_ivp(
            function, t_span=trange, y0=y, args=parameters_temp, method="RK45"
        )
        para_inds = parameters[0]
        temp = sol.y
        y_temp = []
        for row in temp:
            y_temp.append(row[-1])

        para_inds_temp = []
        y_cleaned = [y[0]]
        count = int(len(y_temp[1:]) / 2)

        i_temp = []
        h_temp = []
        aggregate_density = []
        for i in range(count):
            it = y_temp[i + 1]
            ht = y_temp[i + count + 1]
            if it >= 1e-3 or ht >= 1e-3:
                para_inds_temp.append(para_inds[i])
                i_temp.append(it)
                h_temp.append(ht)
                aggregate_density.append(it + ht)

        if len(aggregate_density) == 0:
            break

        for i, ind in enumerate(para_inds_temp):
            temp_output_row.append([ind, step + 1, aggregate_density[i]])

        proportions = [i / sum(aggregate_density) for i in np.cumsum(aggregate_density)]

        rand_stored = np.random.random()

        for j, val in enumerate(proportions):
            if val >= rand_stored:
                ind = j
                break

        if np.random.random() >= 0.5:
            if para_inds_temp[ind] + 1 not in para_inds_temp:
                para_inds_temp.append(para_inds_temp[ind] + 1)
                i_temp.append(i_temp[ind] / 100)
                h_temp.append(h_temp[ind] / 100)
            else:
                index = para_inds_temp.index(para_inds_temp[ind] + 1)
                i_temp[index] += i_temp[ind] / 100
                h_temp[index] += h_temp[ind] / 100
        else:
            if para_inds_temp[ind] - 1 not in para_inds_temp:
                para_inds_temp.append(para_inds_temp[ind] - 1)
                i_temp.append(i_temp[ind] / 100)
                h_temp.append(h_temp[ind] / 100)
            else:
                index = para_inds_temp.index(para_inds_temp[ind] - 1)
                i_temp[index] += i_temp[ind] / 100
                h_temp[index] += h_temp[ind] / 100

        for val in i_temp:
            y_cleaned.append(val)

        for val in h_temp:
            y_cleaned.append(val)

        y = y_cleaned
        parameters_temp[0] = para_inds_temp

        output.append(temp_output_row)
        if verbose:
            print(
                f"We have done {step} steps out of the desired {evo_steps}, this represents {round((step/evo_steps)*100,2)}% completed",
                end="\r",
            )

        if step % 50 == 0 and step >= 100:

            row_1 = output[step - 100]
            row_2 = output[step]
            flag = True

            for row in row_1:
                ind = row[0]
                check = True
                for target in row_2:
                    if target[0] == ind:
                        check = False
                        break
                if check and abs(row[2] - target[2]) > 1e-3:
                    flag = False

            if flag:
                if verbose:
                    print(
                        "Finishing simulation early as we appear to have reached an ESS"
                    )
                break
    return output
```
