## Supplementary material for "Hyperparasitism and the evolution of parasite virulence": Code_base: singular_strategy.py

```
"""
This file contains the expressions for the singular strategies calculated using
Maple. This file also contains the fitness gradients used to calculate the 
singular strategies in files such as eta_evo.py.
"""

from math import sqrt


def sing_strat(H, eta, a, beta, g, gh, l, sig, rho, d):

    singstrat = (
        (l * a + d + gh)
        * (sig * H + a + d + g)
        * (
            (H ** 2 * eta ** 2 * sig ** 2)
            + (d ** 2 * eta ** 2 * rho ** 2)
            + (eta ** 2 * rho ** 2 * g ** 2)
            - (2 * d ** 2 * eta ** 2 * rho)
            - (2 * eta ** 2 * rho * g ** 2)
            + (2 * d ** 2 * eta * rho)
            + (2 * d * eta ** 2 * g)
            - (2 * d * eta * gh)
            - (2 * d * eta * g)
            - (2 * eta * gh * g)
            + sqrt(
                max(
                    0.0,
                    (
                        H ** 2 * eta ** 2 * sig ** 2
                        + d ** 2 * eta ** 2 * rho ** 2
                        + eta ** 2 * rho ** 2 * g ** 2
                        - 2 * d ** 2 * eta ** 2 * rho
                        - 2 * eta ** 2 * rho * g ** 2
                        + 2 * d ** 2 * eta * rho
                        + 2 * d * eta ** 2 * g
                        - 2 * d * eta * gh
                        - 2 * d * eta * g
                        - 2 * eta * gh * g
                        - 2 * H * d * eta ** 2 * rho * sig
                        - 2 * H * eta ** 2 * rho * sig * g
                        + 4 * H * d * eta * rho * sig
                        + 4 * H * eta * rho * sig * gh
                        + 2 * d * l * a
                        - 2 * H * a * eta ** 2 * rho * sig
                        - 2 * H * a * eta * l * sig
                        + 2 * a * d * eta * l * rho
                        + 2 * a * eta * l * rho * g
                        + d ** 2 * eta ** 2
                        + eta ** 2 * g ** 2
                        - 2 * d ** 2 * eta
                        + 2 * d * gh
                        + gh ** 2
                        + d ** 2
                        + 2 * d * eta * rho * gh
                        + 2 * d * eta * rho * g
                        + 2 * eta * rho * gh * g
                        + 2 * d * eta ** 2 * rho ** 2 * g
                        + 2 * H * d * eta ** 2 * sig
                        + 2 * H * eta ** 2 * sig * g
                        - 4 * d * eta ** 2 * rho * g
                        - 2 * H * d * eta * sig
                        - 2 * H * eta * sig * gh
                        + a ** 2 * eta ** 2 * rho ** 2
                        - 2 * a ** 2 * eta ** 2 * rho
                        - 2 * a ** 2 * eta * l
                        + 2 * a * d * eta ** 2
                        + 2 * a * eta ** 2 * g
                        - 2 * a * d * eta
                        - 2 * a * eta * gh
                        + 2 * a * l * gh
                        + 2 * a * d * eta ** 2 * rho ** 2
                        + 2 * a * eta ** 2 * rho ** 2 * g
                        + 2 * H * a * eta ** 2 * sig
                        + 2 * a ** 2 * eta * l * rho
                        - 4 * a * d * eta ** 2 * rho
                        - 4 * a * eta ** 2 * rho * g
                        - 2 * a * d * eta * l
                        + 2 * a * d * eta * rho
                        - 2 * a * eta * l * g
                        + 2 * a * eta * rho * gh
                        + 4 * H * a * eta * l * rho * sig
                        + a ** 2 * eta ** 2
                        + a ** 2 * l ** 2
                    ),
                )
            )
            * gh
            - (2 * H * d * eta ** 2 * rho * sig)
            - (2 * H * eta ** 2 * rho * sig * g)
            + (4 * H * d * eta * rho * sig)
            + (4 * H * eta * rho * sig * gh)
            + H
            * sqrt(
                (
                    H ** 2 * eta ** 2 * sig ** 2
                    + d ** 2 * eta ** 2 * rho ** 2
                    + eta ** 2 * rho ** 2 * g ** 2
                    - 2 * d ** 2 * eta ** 2 * rho
                    - 2 * eta ** 2 * rho * g ** 2
                    + 2 * d ** 2 * eta * rho
                    + 2 * d * eta ** 2 * g
                    - 2 * d * eta * gh
                    - 2 * d * eta * g
                    - 2 * eta * gh * g
                    - 2 * H * d * eta ** 2 * rho * sig
                    - 2 * H * eta ** 2 * rho * sig * g
                    + 4 * H * d * eta * rho * sig
                    + 4 * H * eta * rho * sig * gh
                    + 2 * d * l * a
                    - 2 * H * a * eta ** 2 * rho * sig
                    - 2 * H * a * eta * l * sig
                    + 2 * a * d * eta * l * rho
                    + 2 * a * eta * l * rho * g
                    + d ** 2 * eta ** 2
                    + eta ** 2 * g ** 2
                    - 2 * d ** 2 * eta
                    + 2 * d * gh
                    + gh ** 2
                    + d ** 2
                    + 2 * d * eta * rho * gh
                    + 2 * d * eta * rho * g
                    + 2 * eta * rho * gh * g
                    + 2 * d * eta ** 2 * rho ** 2 * g
                    + 2 * H * d * eta ** 2 * sig
                    + 2 * H * eta ** 2 * sig * g
                    - 4 * d * eta ** 2 * rho * g
                    - 2 * H * d * eta * sig
                    - 2 * H * eta * sig * gh
                    + a ** 2 * eta ** 2 * rho ** 2
                    - 2 * a ** 2 * eta ** 2 * rho
                    - 2 * a ** 2 * eta * l
                    + 2 * a * d * eta ** 2
                    + 2 * a * eta ** 2 * g
                    - 2 * a * d * eta
                    - 2 * a * eta * gh
                    + 2 * a * l * gh
                    + 2 * a * d * eta ** 2 * rho ** 2
                    + 2 * a * eta ** 2 * rho ** 2 * g
                    + 2 * H * a * eta ** 2 * sig
                    + 2 * a ** 2 * eta * l * rho
                    - 4 * a * d * eta ** 2 * rho
                    - 4 * a * eta ** 2 * rho * g
                    - 2 * a * d * eta * l
                    + 2 * a * d * eta * rho
                    - 2 * a * eta * l * g
                    + 2 * a * eta * rho * gh
                    + 4 * H * a * eta * l * rho * sig
                    + a ** 2 * eta ** 2
                    + a ** 2 * l ** 2
                )
            )
            * eta
            * sig
            - a
            * sqrt(
                (
                    H ** 2 * eta ** 2 * sig ** 2
                    + d ** 2 * eta ** 2 * rho ** 2
                    + eta ** 2 * rho ** 2 * g ** 2
                    - 2 * d ** 2 * eta ** 2 * rho
                    - 2 * eta ** 2 * rho * g ** 2
                    + 2 * d ** 2 * eta * rho
                    + 2 * d * eta ** 2 * g
                    - 2 * d * eta * gh
                    - 2 * d * eta * g
                    - 2 * eta * gh * g
                    - 2 * H * d * eta ** 2 * rho * sig
                    - 2 * H * eta ** 2 * rho * sig * g
                    + 4 * H * d * eta * rho * sig
                    + 4 * H * eta * rho * sig * gh
                    + 2 * d * l * a
                    - 2 * H * a * eta ** 2 * rho * sig
                    - 2 * H * a * eta * l * sig
                    + 2 * a * d * eta * l * rho
                    + 2 * a * eta * l * rho * g
                    + d ** 2 * eta ** 2
                    + eta ** 2 * g ** 2
                    - 2 * d ** 2 * eta
                    + 2 * d * gh
                    + gh ** 2
                    + d ** 2
                    + 2 * d * eta * rho * gh
                    + 2 * d * eta * rho * g
                    + 2 * eta * rho * gh * g
                    + 2 * d * eta ** 2 * rho ** 2 * g
                    + 2 * H * d * eta ** 2 * sig
                    + 2 * H * eta ** 2 * sig * g
                    - 4 * d * eta ** 2 * rho * g
                    - 2 * H * d * eta * sig
                    - 2 * H * eta * sig * gh
                    + a ** 2 * eta ** 2 * rho ** 2
                    - 2 * a ** 2 * eta ** 2 * rho
                    - 2 * a ** 2 * eta * l
                    + 2 * a * d * eta ** 2
                    + 2 * a * eta ** 2 * g
                    - 2 * a * d * eta
                    - 2 * a * eta * gh
                    + 2 * a * l * gh
                    + 2 * a * d * eta ** 2 * rho ** 2
                    + 2 * a * eta ** 2 * rho ** 2 * g
                    + 2 * H * a * eta ** 2 * sig
                    + 2 * a ** 2 * eta * l * rho
                    - 4 * a * d * eta ** 2 * rho
                    - 4 * a * eta ** 2 * rho * g
                    - 2 * a * d * eta * l
                    + 2 * a * d * eta * rho
                    - 2 * a * eta * l * g
                    + 2 * a * eta * rho * gh
                    + 4 * H * a * eta * l * rho * sig
                    + a ** 2 * eta ** 2
                    + a ** 2 * l ** 2
                )
            )
            * eta
            * rho
            - sqrt(
                (
                    H ** 2 * eta ** 2 * sig ** 2
                    + d ** 2 * eta ** 2 * rho ** 2
                    + eta ** 2 * rho ** 2 * g ** 2
                    - 2 * d ** 2 * eta ** 2 * rho
                    - 2 * eta ** 2 * rho * g ** 2
                    + 2 * d ** 2 * eta * rho
                    + 2 * d * eta ** 2 * g
                    - 2 * d * eta * gh
                    - 2 * d * eta * g
                    - 2 * eta * gh * g
                    - 2 * H * d * eta ** 2 * rho * sig
                    - 2 * H * eta ** 2 * rho * sig * g
                    + 4 * H * d * eta * rho * sig
                    + 4 * H * eta * rho * sig * gh
                    + 2 * d * l * a
                    - 2 * H * a * eta ** 2 * rho * sig
                    - 2 * H * a * eta * l * sig
                    + 2 * a * d * eta * l * rho
                    + 2 * a * eta * l * rho * g
                    + d ** 2 * eta ** 2
                    + eta ** 2 * g ** 2
                    - 2 * d ** 2 * eta
                    + 2 * d * gh
                    + gh ** 2
                    + d ** 2
                    + 2 * d * eta * rho * gh
                    + 2 * d * eta * rho * g
                    + 2 * eta * rho * gh * g
                    + 2 * d * eta ** 2 * rho ** 2 * g
                    + 2 * H * d * eta ** 2 * sig
                    + 2 * H * eta ** 2 * sig * g
                    - 4 * d * eta ** 2 * rho * g
                    - 2 * H * d * eta * sig
                    - 2 * H * eta * sig * gh
                    + a ** 2 * eta ** 2 * rho ** 2
                    - 2 * a ** 2 * eta ** 2 * rho
                    - 2 * a ** 2 * eta * l
                    + 2 * a * d * eta ** 2
                    + 2 * a * eta ** 2 * g
                    - 2 * a * d * eta
                    - 2 * a * eta * gh
                    + 2 * a * l * gh
                    + 2 * a * d * eta ** 2 * rho ** 2
                    + 2 * a * eta ** 2 * rho ** 2 * g
                    + 2 * H * a * eta ** 2 * sig
                    + 2 * a ** 2 * eta * l * rho
                    - 4 * a * d * eta ** 2 * rho
                    - 4 * a * eta ** 2 * rho * g
                    - 2 * a * d * eta * l
                    + 2 * a * d * eta * rho
                    - 2 * a * eta * l * g
                    + 2 * a * eta * rho * gh
                    + 4 * H * a * eta * l * rho * sig
                    + a ** 2 * eta ** 2
                    + a ** 2 * l ** 2
                )
            )
            * d
            * eta
            * rho
            - sqrt(
                (
                    H ** 2 * eta ** 2 * sig ** 2
                    + d ** 2 * eta ** 2 * rho ** 2
                    + eta ** 2 * rho ** 2 * g ** 2
                    - 2 * d ** 2 * eta ** 2 * rho
                    - 2 * eta ** 2 * rho * g ** 2
                    + 2 * d ** 2 * eta * rho
                    + 2 * d * eta ** 2 * g
                    - 2 * d * eta * gh
                    - 2 * d * eta * g
                    - 2 * eta * gh * g
                    - 2 * H * d * eta ** 2 * rho * sig
                    - 2 * H * eta ** 2 * rho * sig * g
                    + 4 * H * d * eta * rho * sig
                    + 4 * H * eta * rho * sig * gh
                    + 2 * d * l * a
                    - 2 * H * a * eta ** 2 * rho * sig
                    - 2 * H * a * eta * l * sig
                    + 2 * a * d * eta * l * rho
                    + 2 * a * eta * l * rho * g
                    + d ** 2 * eta ** 2
                    + eta ** 2 * g ** 2
                    - 2 * d ** 2 * eta
                    + 2 * d * gh
                    + gh ** 2
                    + d ** 2
                    + 2 * d * eta * rho * gh
                    + 2 * d * eta * rho * g
                    + 2 * eta * rho * gh * g
                    + 2 * d * eta ** 2 * rho ** 2 * g
                    + 2 * H * d * eta ** 2 * sig
                    + 2 * H * eta ** 2 * sig * g
                    - 4 * d * eta ** 2 * rho * g
                    - 2 * H * d * eta * sig
                    - 2 * H * eta * sig * gh
                    + a ** 2 * eta ** 2 * rho ** 2
                    - 2 * a ** 2 * eta ** 2 * rho
                    - 2 * a ** 2 * eta * l
                    + 2 * a * d * eta ** 2
                    + 2 * a * eta ** 2 * g
                    - 2 * a * d * eta
                    - 2 * a * eta * gh
                    + 2 * a * l * gh
                    + 2 * a * d * eta ** 2 * rho ** 2
                    + 2 * a * eta ** 2 * rho ** 2 * g
                    + 2 * H * a * eta ** 2 * sig
                    + 2 * a ** 2 * eta * l * rho
                    - 4 * a * d * eta ** 2 * rho
                    - 4 * a * eta ** 2 * rho * g
                    - 2 * a * d * eta * l
                    + 2 * a * d * eta * rho
                    - 2 * a * eta * l * g
                    + 2 * a * eta * rho * gh
                    + 4 * H * a * eta * l * rho * sig
                    + a ** 2 * eta ** 2
                    + a ** 2 * l ** 2
                )
            )
            * eta
            * rho
            * g
            + (2 * d * l * a)
            - (2 * H * a * eta ** 2 * rho * sig)
            - (2 * H * a * eta * l * sig)
            + (2 * a * d * eta * l * rho)
            + (2 * a * eta * l * rho * g)
            + (d ** 2 * eta ** 2)
            + (eta ** 2 * g ** 2)
            - (2 * d ** 2 * eta)
            + (2 * d * gh)
            + (gh ** 2)
            + (d ** 2)
            + (2 * d * eta * rho * gh)
            + (2 * d * eta * rho * g)
            + (2 * eta * rho * gh * g)
            + (2 * d * eta ** 2 * rho ** 2 * g)
            + (2 * H * d * eta ** 2 * sig)
            + (2 * H * eta ** 2 * sig * g)
            - (4 * d * eta ** 2 * rho * g)
            - (2 * H * d * eta * sig)
            - (2 * H * eta * sig * gh)
            + a
            * sqrt(
                (
                    H ** 2 * eta ** 2 * sig ** 2
                    + d ** 2 * eta ** 2 * rho ** 2
                    + eta ** 2 * rho ** 2 * g ** 2
                    - 2 * d ** 2 * eta ** 2 * rho
                    - 2 * eta ** 2 * rho * g ** 2
                    + 2 * d ** 2 * eta * rho
                    + 2 * d * eta ** 2 * g
                    - 2 * d * eta * gh
                    - 2 * d * eta * g
                    - 2 * eta * gh * g
                    - 2 * H * d * eta ** 2 * rho * sig
                    - 2 * H * eta ** 2 * rho * sig * g
                    + 4 * H * d * eta * rho * sig
                    + 4 * H * eta * rho * sig * gh
                    + 2 * d * l * a
                    - 2 * H * a * eta ** 2 * rho * sig
                    - 2 * H * a * eta * l * sig
                    + 2 * a * d * eta * l * rho
                    + 2 * a * eta * l * rho * g
                    + d ** 2 * eta ** 2
                    + eta ** 2 * g ** 2
                    - 2 * d ** 2 * eta
                    + 2 * d * gh
                    + gh ** 2
                    + d ** 2
                    + 2 * d * eta * rho * gh
                    + 2 * d * eta * rho * g
                    + 2 * eta * rho * gh * g
                    + 2 * d * eta ** 2 * rho ** 2 * g
                    + 2 * H * d * eta ** 2 * sig
                    + 2 * H * eta ** 2 * sig * g
                    - 4 * d * eta ** 2 * rho * g
                    - 2 * H * d * eta * sig
                    - 2 * H * eta * sig * gh
                    + a ** 2 * eta ** 2 * rho ** 2
                    - 2 * a ** 2 * eta ** 2 * rho
                    - 2 * a ** 2 * eta * l
                    + 2 * a * d * eta ** 2
                    + 2 * a * eta ** 2 * g
                    - 2 * a * d * eta
                    - 2 * a * eta * gh
                    + 2 * a * l * gh
                    + 2 * a * d * eta ** 2 * rho ** 2
                    + 2 * a * eta ** 2 * rho ** 2 * g
                    + 2 * H * a * eta ** 2 * sig
                    + 2 * a ** 2 * eta * l * rho
                    - 4 * a * d * eta ** 2 * rho
                    - 4 * a * eta ** 2 * rho * g
                    - 2 * a * d * eta * l
                    + 2 * a * d * eta * rho
                    - 2 * a * eta * l * g
                    + 2 * a * eta * rho * gh
                    + 4 * H * a * eta * l * rho * sig
                    + a ** 2 * eta ** 2
                    + a ** 2 * l ** 2
                )
            )
            * l
            + sqrt(
                (
                    H ** 2 * eta ** 2 * sig ** 2
                    + d ** 2 * eta ** 2 * rho ** 2
                    + eta ** 2 * rho ** 2 * g ** 2
                    - 2 * d ** 2 * eta ** 2 * rho
                    - 2 * eta ** 2 * rho * g ** 2
                    + 2 * d ** 2 * eta * rho
                    + 2 * d * eta ** 2 * g
                    - 2 * d * eta * gh
                    - 2 * d * eta * g
                    - 2 * eta * gh * g
                    - 2 * H * d * eta ** 2 * rho * sig
                    - 2 * H * eta ** 2 * rho * sig * g
                    + 4 * H * d * eta * rho * sig
                    + 4 * H * eta * rho * sig * gh
                    + 2 * d * l * a
                    - 2 * H * a * eta ** 2 * rho * sig
                    - 2 * H * a * eta * l * sig
                    + 2 * a * d * eta * l * rho
                    + 2 * a * eta * l * rho * g
                    + d ** 2 * eta ** 2
                    + eta ** 2 * g ** 2
                    - 2 * d ** 2 * eta
                    + 2 * d * gh
                    + gh ** 2
                    + d ** 2
                    + 2 * d * eta * rho * gh
                    + 2 * d * eta * rho * g
                    + 2 * eta * rho * gh * g
                    + 2 * d * eta ** 2 * rho ** 2 * g
                    + 2 * H * d * eta ** 2 * sig
                    + 2 * H * eta ** 2 * sig * g
                    - 4 * d * eta ** 2 * rho * g
                    - 2 * H * d * eta * sig
                    - 2 * H * eta * sig * gh
                    + a ** 2 * eta ** 2 * rho ** 2
                    - 2 * a ** 2 * eta ** 2 * rho
                    - 2 * a ** 2 * eta * l
                    + 2 * a * d * eta ** 2
                    + 2 * a * eta ** 2 * g
                    - 2 * a * d * eta
                    - 2 * a * eta * gh
                    + 2 * a * l * gh
                    + 2 * a * d * eta ** 2 * rho ** 2
                    + 2 * a * eta ** 2 * rho ** 2 * g
                    + 2 * H * a * eta ** 2 * sig
                    + 2 * a ** 2 * eta * l * rho
                    - 4 * a * d * eta ** 2 * rho
                    - 4 * a * eta ** 2 * rho * g
                    - 2 * a * d * eta * l
                    + 2 * a * d * eta * rho
                    - 2 * a * eta * l * g
                    + 2 * a * eta * rho * gh
                    + 4 * H * a * eta * l * rho * sig
                    + a ** 2 * eta ** 2
                    + a ** 2 * l ** 2
                )
            )
            * d
            + a
            * sqrt(
                (
                    H ** 2 * eta ** 2 * sig ** 2
                    + d ** 2 * eta ** 2 * rho ** 2
                    + eta ** 2 * rho ** 2 * g ** 2
                    - 2 * d ** 2 * eta ** 2 * rho
                    - 2 * eta ** 2 * rho * g ** 2
                    + 2 * d ** 2 * eta * rho
                    + 2 * d * eta ** 2 * g
                    - 2 * d * eta * gh
                    - 2 * d * eta * g
                    - 2 * eta * gh * g
                    - 2 * H * d * eta ** 2 * rho * sig
                    - 2 * H * eta ** 2 * rho * sig * g
                    + 4 * H * d * eta * rho * sig
                    + 4 * H * eta * rho * sig * gh
                    + 2 * d * l * a
                    - 2 * H * a * eta ** 2 * rho * sig
                    - 2 * H * a * eta * l * sig
                    + 2 * a * d * eta * l * rho
                    + 2 * a * eta * l * rho * g
                    + d ** 2 * eta ** 2
                    + eta ** 2 * g ** 2
                    - 2 * d ** 2 * eta
                    + 2 * d * gh
                    + gh ** 2
                    + d ** 2
                    + 2 * d * eta * rho * gh
                    + 2 * d * eta * rho * g
                    + 2 * eta * rho * gh * g
                    + 2 * d * eta ** 2 * rho ** 2 * g
                    + 2 * H * d * eta ** 2 * sig
                    + 2 * H * eta ** 2 * sig * g
                    - 4 * d * eta ** 2 * rho * g
                    - 2 * H * d * eta * sig
                    - 2 * H * eta * sig * gh
                    + a ** 2 * eta ** 2 * rho ** 2
                    - 2 * a ** 2 * eta ** 2 * rho
                    - 2 * a ** 2 * eta * l
                    + 2 * a * d * eta ** 2
                    + 2 * a * eta ** 2 * g
                    - 2 * a * d * eta
                    - 2 * a * eta * gh
                    + 2 * a * l * gh
                    + 2 * a * d * eta ** 2 * rho ** 2
                    + 2 * a * eta ** 2 * rho ** 2 * g
                    + 2 * H * a * eta ** 2 * sig
                    + 2 * a ** 2 * eta * l * rho
                    - 4 * a * d * eta ** 2 * rho
                    - 4 * a * eta ** 2 * rho * g
                    - 2 * a * d * eta * l
                    + 2 * a * d * eta * rho
                    - 2 * a * eta * l * g
                    + 2 * a * eta * rho * gh
                    + 4 * H * a * eta * l * rho * sig
                    + a ** 2 * eta ** 2
                    + a ** 2 * l ** 2
                )
            )
            * eta
            + sqrt(
                (
                    H ** 2 * eta ** 2 * sig ** 2
                    + d ** 2 * eta ** 2 * rho ** 2
                    + eta ** 2 * rho ** 2 * g ** 2
                    - 2 * d ** 2 * eta ** 2 * rho
                    - 2 * eta ** 2 * rho * g ** 2
                    + 2 * d ** 2 * eta * rho
                    + 2 * d * eta ** 2 * g
                    - 2 * d * eta * gh
                    - 2 * d * eta * g
                    - 2 * eta * gh * g
                    - 2 * H * d * eta ** 2 * rho * sig
                    - 2 * H * eta ** 2 * rho * sig * g
                    + 4 * H * d * eta * rho * sig
                    + 4 * H * eta * rho * sig * gh
                    + 2 * d * l * a
                    - 2 * H * a * eta ** 2 * rho * sig
                    - 2 * H * a * eta * l * sig
                    + 2 * a * d * eta * l * rho
                    + 2 * a * eta * l * rho * g
                    + d ** 2 * eta ** 2
                    + eta ** 2 * g ** 2
                    - 2 * d ** 2 * eta
                    + 2 * d * gh
                    + gh ** 2
                    + d ** 2
                    + 2 * d * eta * rho * gh
                    + 2 * d * eta * rho * g
                    + 2 * eta * rho * gh * g
                    + 2 * d * eta ** 2 * rho ** 2 * g
                    + 2 * H * d * eta ** 2 * sig
                    + 2 * H * eta ** 2 * sig * g
                    - 4 * d * eta ** 2 * rho * g
                    - 2 * H * d * eta * sig
                    - 2 * H * eta * sig * gh
                    + a ** 2 * eta ** 2 * rho ** 2
                    - 2 * a ** 2 * eta ** 2 * rho
                    - 2 * a ** 2 * eta * l
                    + 2 * a * d * eta ** 2
                    + 2 * a * eta ** 2 * g
                    - 2 * a * d * eta
                    - 2 * a * eta * gh
                    + 2 * a * l * gh
                    + 2 * a * d * eta ** 2 * rho ** 2
                    + 2 * a * eta ** 2 * rho ** 2 * g
                    + 2 * H * a * eta ** 2 * sig
                    + 2 * a ** 2 * eta * l * rho
                    - 4 * a * d * eta ** 2 * rho
                    - 4 * a * eta ** 2 * rho * g
                    - 2 * a * d * eta * l
                    + 2 * a * d * eta * rho
                    - 2 * a * eta * l * g
                    + 2 * a * eta * rho * gh
                    + 4 * H * a * eta * l * rho * sig
                    + a ** 2 * eta ** 2
                    + a ** 2 * l ** 2
                )
            )
            * d
            * eta
            + sqrt(
                (
                    H ** 2 * eta ** 2 * sig ** 2
                    + d ** 2 * eta ** 2 * rho ** 2
                    + eta ** 2 * rho ** 2 * g ** 2
                    - 2 * d ** 2 * eta ** 2 * rho
                    - 2 * eta ** 2 * rho * g ** 2
                    + 2 * d ** 2 * eta * rho
                    + 2 * d * eta ** 2 * g
                    - 2 * d * eta * gh
                    - 2 * d * eta * g
                    - 2 * eta * gh * g
                    - 2 * H * d * eta ** 2 * rho * sig
                    - 2 * H * eta ** 2 * rho * sig * g
                    + 4 * H * d * eta * rho * sig
                    + 4 * H * eta * rho * sig * gh
                    + 2 * d * l * a
                    - 2 * H * a * eta ** 2 * rho * sig
                    - 2 * H * a * eta * l * sig
                    + 2 * a * d * eta * l * rho
                    + 2 * a * eta * l * rho * g
                    + d ** 2 * eta ** 2
                    + eta ** 2 * g ** 2
                    - 2 * d ** 2 * eta
                    + 2 * d * gh
                    + gh ** 2
                    + d ** 2
                    + 2 * d * eta * rho * gh
                    + 2 * d * eta * rho * g
                    + 2 * eta * rho * gh * g
                    + 2 * d * eta ** 2 * rho ** 2 * g
                    + 2 * H * d * eta ** 2 * sig
                    + 2 * H * eta ** 2 * sig * g
                    - 4 * d * eta ** 2 * rho * g
                    - 2 * H * d * eta * sig
                    - 2 * H * eta * sig * gh
                    + a ** 2 * eta ** 2 * rho ** 2
                    - 2 * a ** 2 * eta ** 2 * rho
                    - 2 * a ** 2 * eta * l
                    + 2 * a * d * eta ** 2
                    + 2 * a * eta ** 2 * g
                    - 2 * a * d * eta
                    - 2 * a * eta * gh
                    + 2 * a * l * gh
                    + 2 * a * d * eta ** 2 * rho ** 2
                    + 2 * a * eta ** 2 * rho ** 2 * g
                    + 2 * H * a * eta ** 2 * sig
                    + 2 * a ** 2 * eta * l * rho
                    - 4 * a * d * eta ** 2 * rho
                    - 4 * a * eta ** 2 * rho * g
                    - 2 * a * d * eta * l
                    + 2 * a * d * eta * rho
                    - 2 * a * eta * l * g
                    + 2 * a * eta * rho * gh
                    + 4 * H * a * eta * l * rho * sig
                    + a ** 2 * eta ** 2
                    + a ** 2 * l ** 2
                )
            )
            * eta
            * g
            + (a ** 2 * eta ** 2 * rho ** 2)
            - (2 * a ** 2 * eta ** 2 * rho)
            - (2 * a ** 2 * eta * l)
            + (2 * a * d * eta ** 2)
            + (2 * a * eta ** 2 * g)
            - (2 * a * d * eta)
            - (2 * a * eta * gh)
            + (2 * a * l * gh)
            + (2 * a * d * eta ** 2 * rho ** 2)
            + (2 * a * eta ** 2 * rho ** 2 * g)
            + (2 * H * a * eta ** 2 * sig)
            + (2 * a ** 2 * eta * l * rho)
            - (4 * a * d * eta ** 2 * rho)
            - (4 * a * eta ** 2 * rho * g)
            - (2 * a * d * eta * l)
            + (2 * a * d * eta * rho)
            - (2 * a * eta * l * g)
            + (2 * a * eta * rho * gh)
            + (4 * H * a * eta * l * rho * sig)
            + (a ** 2 * eta ** 2)
            + (a ** 2 * l ** 2)
        )
        / beta
        / (
            (6 * H * eta ** 2 * l * sig * g * a)
            + (H * eta ** 2 * rho * sig * gh * a)
            - (2 * H * eta * l ** 2 * sig * g * a)
            - (12 * d * eta ** 2 * l * rho * g * a)
            + (2 * d * eta * l ** 2 * rho * g * a)
            + 0.2e1
            * H
            * d
            * eta
            * l
            * sig
            * sqrt(
                (
                    H ** 2 * eta ** 2 * sig ** 2
                    + d ** 2 * eta ** 2 * rho ** 2
                    + eta ** 2 * rho ** 2 * g ** 2
                    - 2 * d ** 2 * eta ** 2 * rho
                    - 2 * eta ** 2 * rho * g ** 2
                    + 2 * d ** 2 * eta * rho
                    + 2 * d * eta ** 2 * g
                    - 2 * d * eta * gh
                    - 2 * d * eta * g
                    - 2 * eta * gh * g
                    - 2 * H * d * eta ** 2 * rho * sig
                    - 2 * H * eta ** 2 * rho * sig * g
                    + 4 * H * d * eta * rho * sig
                    + 4 * H * eta * rho * sig * gh
                    + 2 * d * l * a
                    - 2 * H * a * eta ** 2 * rho * sig
                    - 2 * H * a * eta * l * sig
                    + 2 * a * d * eta * l * rho
                    + 2 * a * eta * l * rho * g
                    + d ** 2 * eta ** 2
                    + eta ** 2 * g ** 2
                    - 2 * d ** 2 * eta
                    + 2 * d * gh
                    + gh ** 2
                    + d ** 2
                    + 2 * d * eta * rho * gh
                    + 2 * d * eta * rho * g
                    + 2 * eta * rho * gh * g
                    + 2 * d * eta ** 2 * rho ** 2 * g
                    + 2 * H * d * eta ** 2 * sig
                    + 2 * H * eta ** 2 * sig * g
                    - 4 * d * eta ** 2 * rho * g
                    - 2 * H * d * eta * sig
                    - 2 * H * eta * sig * gh
                    + a ** 2 * eta ** 2 * rho ** 2
                    - 2 * a ** 2 * eta ** 2 * rho
                    - 2 * a ** 2 * eta * l
                    + 2 * a * d * eta ** 2
                    + 2 * a * eta ** 2 * g
                    - 2 * a * d * eta
                    - 2 * a * eta * gh
                    + 2 * a * l * gh
                    + 2 * a * d * eta ** 2 * rho ** 2
                    + 2 * a * eta ** 2 * rho ** 2 * g
                    + 2 * H * a * eta ** 2 * sig
                    + 2 * a ** 2 * eta * l * rho
                    - 4 * a * d * eta ** 2 * rho
                    - 4 * a * eta ** 2 * rho * g
                    - 2 * a * d * eta * l
                    + 2 * a * d * eta * rho
                    - 2 * a * eta * l * g
                    + 2 * a * eta * rho * gh
                    + 4 * H * a * eta * l * rho * sig
                    + a ** 2 * eta ** 2
                    + a ** 2 * l ** 2
                )
            )
            - (4 * H * d * eta * l * sig * a)
            + H
            * d
            * eta
            * rho
            * sig
            * sqrt(
                (
                    H ** 2 * eta ** 2 * sig ** 2
                    + d ** 2 * eta ** 2 * rho ** 2
                    + eta ** 2 * rho ** 2 * g ** 2
                    - 2 * d ** 2 * eta ** 2 * rho
                    - 2 * eta ** 2 * rho * g ** 2
                    + 2 * d ** 2 * eta * rho
                    + 2 * d * eta ** 2 * g
                    - 2 * d * eta * gh
                    - 2 * d * eta * g
                    - 2 * eta * gh * g
                    - 2 * H * d * eta ** 2 * rho * sig
                    - 2 * H * eta ** 2 * rho * sig * g
                    + 4 * H * d * eta * rho * sig
                    + 4 * H * eta * rho * sig * gh
                    + 2 * d * l * a
                    - 2 * H * a * eta ** 2 * rho * sig
                    - 2 * H * a * eta * l * sig
                    + 2 * a * d * eta * l * rho
                    + 2 * a * eta * l * rho * g
                    + d ** 2 * eta ** 2
                    + eta ** 2 * g ** 2
                    - 2 * d ** 2 * eta
                    + 2 * d * gh
                    + gh ** 2
                    + d ** 2
                    + 2 * d * eta * rho * gh
                    + 2 * d * eta * rho * g
                    + 2 * eta * rho * gh * g
                    + 2 * d * eta ** 2 * rho ** 2 * g
                    + 2 * H * d * eta ** 2 * sig
                    + 2 * H * eta ** 2 * sig * g
                    - 4 * d * eta ** 2 * rho * g
                    - 2 * H * d * eta * sig
                    - 2 * H * eta * sig * gh
                    + a ** 2 * eta ** 2 * rho ** 2
                    - 2 * a ** 2 * eta ** 2 * rho
                    - 2 * a ** 2 * eta * l
                    + 2 * a * d * eta ** 2
                    + 2 * a * eta ** 2 * g
                    - 2 * a * d * eta
                    - 2 * a * eta * gh
                    + 2 * a * l * gh
                    + 2 * a * d * eta ** 2 * rho ** 2
                    + 2 * a * eta ** 2 * rho ** 2 * g
                    + 2 * H * a * eta ** 2 * sig
                    + 2 * a ** 2 * eta * l * rho
                    - 4 * a * d * eta ** 2 * rho
                    - 4 * a * eta ** 2 * rho * g
                    - 2 * a * d * eta * l
                    + 2 * a * d * eta * rho
                    - 2 * a * eta * l * g
                    + 2 * a * eta * rho * gh
                    + 4 * H * a * eta * l * rho * sig
                    + a ** 2 * eta ** 2
                    + a ** 2 * l ** 2
                )
            )
            - (4 * H * eta * l * sig * gh * a)
            + 0.2e1
            * H
            * eta
            * l
            * sig
            * g
            * sqrt(
                (
                    H ** 2 * eta ** 2 * sig ** 2
                    + d ** 2 * eta ** 2 * rho ** 2
                    + eta ** 2 * rho ** 2 * g ** 2
                    - 2 * d ** 2 * eta ** 2 * rho
                    - 2 * eta ** 2 * rho * g ** 2
                    + 2 * d ** 2 * eta * rho
                    + 2 * d * eta ** 2 * g
                    - 2 * d * eta * gh
                    - 2 * d * eta * g
                    - 2 * eta * gh * g
                    - 2 * H * d * eta ** 2 * rho * sig
                    - 2 * H * eta ** 2 * rho * sig * g
                    + 4 * H * d * eta * rho * sig
                    + 4 * H * eta * rho * sig * gh
                    + 2 * d * l * a
                    - 2 * H * a * eta ** 2 * rho * sig
                    - 2 * H * a * eta * l * sig
                    + 2 * a * d * eta * l * rho
                    + 2 * a * eta * l * rho * g
                    + d ** 2 * eta ** 2
                    + eta ** 2 * g ** 2
                    - 2 * d ** 2 * eta
                    + 2 * d * gh
                    + gh ** 2
                    + d ** 2
                    + 2 * d * eta * rho * gh
                    + 2 * d * eta * rho * g
                    + 2 * eta * rho * gh * g
                    + 2 * d * eta ** 2 * rho ** 2 * g
                    + 2 * H * d * eta ** 2 * sig
                    + 2 * H * eta ** 2 * sig * g
                    - 4 * d * eta ** 2 * rho * g
                    - 2 * H * d * eta * sig
                    - 2 * H * eta * sig * gh
                    + a ** 2 * eta ** 2 * rho ** 2
                    - 2 * a ** 2 * eta ** 2 * rho
                    - 2 * a ** 2 * eta * l
                    + 2 * a * d * eta ** 2
                    + 2 * a * eta ** 2 * g
                    - 2 * a * d * eta
                    - 2 * a * eta * gh
                    + 2 * a * l * gh
                    + 2 * a * d * eta ** 2 * rho ** 2
                    + 2 * a * eta ** 2 * rho ** 2 * g
                    + 2 * H * a * eta ** 2 * sig
                    + 2 * a ** 2 * eta * l * rho
                    - 4 * a * d * eta ** 2 * rho
                    - 4 * a * eta ** 2 * rho * g
                    - 2 * a * d * eta * l
                    + 2 * a * d * eta * rho
                    - 2 * a * eta * l * g
                    + 2 * a * eta * rho * gh
                    + 4 * H * a * eta * l * rho * sig
                    + a ** 2 * eta ** 2
                    + a ** 2 * l ** 2
                )
            )
            + 0.2e1
            * H
            * eta
            * l
            * sig
            * sqrt(
                (
                    H ** 2 * eta ** 2 * sig ** 2
                    + d ** 2 * eta ** 2 * rho ** 2
                    + eta ** 2 * rho ** 2 * g ** 2
                    - 2 * d ** 2 * eta ** 2 * rho
                    - 2 * eta ** 2 * rho * g ** 2
                    + 2 * d ** 2 * eta * rho
                    + 2 * d * eta ** 2 * g
                    - 2 * d * eta * gh
                    - 2 * d * eta * g
                    - 2 * eta * gh * g
                    - 2 * H * d * eta ** 2 * rho * sig
                    - 2 * H * eta ** 2 * rho * sig * g
                    + 4 * H * d * eta * rho * sig
                    + 4 * H * eta * rho * sig * gh
                    + 2 * d * l * a
                    - 2 * H * a * eta ** 2 * rho * sig
                    - 2 * H * a * eta * l * sig
                    + 2 * a * d * eta * l * rho
                    + 2 * a * eta * l * rho * g
                    + d ** 2 * eta ** 2
                    + eta ** 2 * g ** 2
                    - 2 * d ** 2 * eta
                    + 2 * d * gh
                    + gh ** 2
                    + d ** 2
                    + 2 * d * eta * rho * gh
                    + 2 * d * eta * rho * g
                    + 2 * eta * rho * gh * g
                    + 2 * d * eta ** 2 * rho ** 2 * g
                    + 2 * H * d * eta ** 2 * sig
                    + 2 * H * eta ** 2 * sig * g
                    - 4 * d * eta ** 2 * rho * g
                    - 2 * H * d * eta * sig
                    - 2 * H * eta * sig * gh
                    + a ** 2 * eta ** 2 * rho ** 2
                    - 2 * a ** 2 * eta ** 2 * rho
                    - 2 * a ** 2 * eta * l
                    + 2 * a * d * eta ** 2
                    + 2 * a * eta ** 2 * g
                    - 2 * a * d * eta
                    - 2 * a * eta * gh
                    + 2 * a * l * gh
                    + 2 * a * d * eta ** 2 * rho ** 2
                    + 2 * a * eta ** 2 * rho ** 2 * g
                    + 2 * H * a * eta ** 2 * sig
                    + 2 * a ** 2 * eta * l * rho
                    - 4 * a * d * eta ** 2 * rho
                    - 4 * a * eta ** 2 * rho * g
                    - 2 * a * d * eta * l
                    + 2 * a * d * eta * rho
                    - 2 * a * eta * l * g
                    + 2 * a * eta * rho * gh
                    + 4 * H * a * eta * l * rho * sig
                    + a ** 2 * eta ** 2
                    + a ** 2 * l ** 2
                )
            )
            * a
            + H
            * eta
            * rho
            * sig
            * gh
            * sqrt(
                (
                    H ** 2 * eta ** 2 * sig ** 2
                    + d ** 2 * eta ** 2 * rho ** 2
                    + eta ** 2 * rho ** 2 * g ** 2
                    - 2 * d ** 2 * eta ** 2 * rho
                    - 2 * eta ** 2 * rho * g ** 2
                    + 2 * d ** 2 * eta * rho
                    + 2 * d * eta ** 2 * g
                    - 2 * d * eta * gh
                    - 2 * d * eta * g
                    - 2 * eta * gh * g
                    - 2 * H * d * eta ** 2 * rho * sig
                    - 2 * H * eta ** 2 * rho * sig * g
                    + 4 * H * d * eta * rho * sig
                    + 4 * H * eta * rho * sig * gh
                    + 2 * d * l * a
                    - 2 * H * a * eta ** 2 * rho * sig
                    - 2 * H * a * eta * l * sig
                    + 2 * a * d * eta * l * rho
                    + 2 * a * eta * l * rho * g
                    + d ** 2 * eta ** 2
                    + eta ** 2 * g ** 2
                    - 2 * d ** 2 * eta
                    + 2 * d * gh
                    + gh ** 2
                    + d ** 2
                    + 2 * d * eta * rho * gh
                    + 2 * d * eta * rho * g
                    + 2 * eta * rho * gh * g
                    + 2 * d * eta ** 2 * rho ** 2 * g
                    + 2 * H * d * eta ** 2 * sig
                    + 2 * H * eta ** 2 * sig * g
                    - 4 * d * eta ** 2 * rho * g
                    - 2 * H * d * eta * sig
                    - 2 * H * eta * sig * gh
                    + a ** 2 * eta ** 2 * rho ** 2
                    - 2 * a ** 2 * eta ** 2 * rho
                    - 2 * a ** 2 * eta * l
                    + 2 * a * d * eta ** 2
                    + 2 * a * eta ** 2 * g
                    - 2 * a * d * eta
                    - 2 * a * eta * gh
                    + 2 * a * l * gh
                    + 2 * a * d * eta ** 2 * rho ** 2
                    + 2 * a * eta ** 2 * rho ** 2 * g
                    + 2 * H * a * eta ** 2 * sig
                    + 2 * a ** 2 * eta * l * rho
                    - 4 * a * d * eta ** 2 * rho
                    - 4 * a * eta ** 2 * rho * g
                    - 2 * a * d * eta * l
                    + 2 * a * d * eta * rho
                    - 2 * a * eta * l * g
                    + 2 * a * eta * rho * gh
                    + 4 * H * a * eta * l * rho * sig
                    + a ** 2 * eta ** 2
                    + a ** 2 * l ** 2
                )
            )
            + (4 * d * eta * l * rho * gh * a)
            - 0.2e1
            * d
            * eta
            * l
            * rho
            * g
            * sqrt(
                (
                    H ** 2 * eta ** 2 * sig ** 2
                    + d ** 2 * eta ** 2 * rho ** 2
                    + eta ** 2 * rho ** 2 * g ** 2
                    - 2 * d ** 2 * eta ** 2 * rho
                    - 2 * eta ** 2 * rho * g ** 2
                    + 2 * d ** 2 * eta * rho
                    + 2 * d * eta ** 2 * g
                    - 2 * d * eta * gh
                    - 2 * d * eta * g
                    - 2 * eta * gh * g
                    - 2 * H * d * eta ** 2 * rho * sig
                    - 2 * H * eta ** 2 * rho * sig * g
                    + 4 * H * d * eta * rho * sig
                    + 4 * H * eta * rho * sig * gh
                    + 2 * d * l * a
                    - 2 * H * a * eta ** 2 * rho * sig
                    - 2 * H * a * eta * l * sig
                    + 2 * a * d * eta * l * rho
                    + 2 * a * eta * l * rho * g
                    + d ** 2 * eta ** 2
                    + eta ** 2 * g ** 2
                    - 2 * d ** 2 * eta
                    + 2 * d * gh
                    + gh ** 2
                    + d ** 2
                    + 2 * d * eta * rho * gh
                    + 2 * d * eta * rho * g
                    + 2 * eta * rho * gh * g
                    + 2 * d * eta ** 2 * rho ** 2 * g
                    + 2 * H * d * eta ** 2 * sig
                    + 2 * H * eta ** 2 * sig * g
                    - 4 * d * eta ** 2 * rho * g
                    - 2 * H * d * eta * sig
                    - 2 * H * eta * sig * gh
                    + a ** 2 * eta ** 2 * rho ** 2
                    - 2 * a ** 2 * eta ** 2 * rho
                    - 2 * a ** 2 * eta * l
                    + 2 * a * d * eta ** 2
                    + 2 * a * eta ** 2 * g
                    - 2 * a * d * eta
                    - 2 * a * eta * gh
                    + 2 * a * l * gh
                    + 2 * a * d * eta ** 2 * rho ** 2
                    + 2 * a * eta ** 2 * rho ** 2 * g
                    + 2 * H * a * eta ** 2 * sig
                    + 2 * a ** 2 * eta * l * rho
                    - 4 * a * d * eta ** 2 * rho
                    - 4 * a * eta ** 2 * rho * g
                    - 2 * a * d * eta * l
                    + 2 * a * d * eta * rho
                    - 2 * a * eta * l * g
                    + 2 * a * eta * rho * gh
                    + 4 * H * a * eta * l * rho * sig
                    + a ** 2 * eta ** 2
                    + a ** 2 * l ** 2
                )
            )
            + (4 * d * eta * l * rho * g * a)
            - 0.2e1
            * d
            * eta
            * l
            * rho
            * sqrt(
                (
                    H ** 2 * eta ** 2 * sig ** 2
                    + d ** 2 * eta ** 2 * rho ** 2
                    + eta ** 2 * rho ** 2 * g ** 2
                    - 2 * d ** 2 * eta ** 2 * rho
                    - 2 * eta ** 2 * rho * g ** 2
                    + 2 * d ** 2 * eta * rho
                    + 2 * d * eta ** 2 * g
                    - 2 * d * eta * gh
                    - 2 * d * eta * g
                    - 2 * eta * gh * g
                    - 2 * H * d * eta ** 2 * rho * sig
                    - 2 * H * eta ** 2 * rho * sig * g
                    + 4 * H * d * eta * rho * sig
                    + 4 * H * eta * rho * sig * gh
                    + 2 * d * l * a
                    - 2 * H * a * eta ** 2 * rho * sig
                    - 2 * H * a * eta * l * sig
                    + 2 * a * d * eta * l * rho
                    + 2 * a * eta * l * rho * g
                    + d ** 2 * eta ** 2
                    + eta ** 2 * g ** 2
                    - 2 * d ** 2 * eta
                    + 2 * d * gh
                    + gh ** 2
                    + d ** 2
                    + 2 * d * eta * rho * gh
                    + 2 * d * eta * rho * g
                    + 2 * eta * rho * gh * g
                    + 2 * d * eta ** 2 * rho ** 2 * g
                    + 2 * H * d * eta ** 2 * sig
                    + 2 * H * eta ** 2 * sig * g
                    - 4 * d * eta ** 2 * rho * g
                    - 2 * H * d * eta * sig
                    - 2 * H * eta * sig * gh
                    + a ** 2 * eta ** 2 * rho ** 2
                    - 2 * a ** 2 * eta ** 2 * rho
                    - 2 * a ** 2 * eta * l
                    + 2 * a * d * eta ** 2
                    + 2 * a * eta ** 2 * g
                    - 2 * a * d * eta
                    - 2 * a * eta * gh
                    + 2 * a * l * gh
                    + 2 * a * d * eta ** 2 * rho ** 2
                    + 2 * a * eta ** 2 * rho ** 2 * g
                    + 2 * H * a * eta ** 2 * sig
                    + 2 * a ** 2 * eta * l * rho
                    - 4 * a * d * eta ** 2 * rho
                    - 4 * a * eta ** 2 * rho * g
                    - 2 * a * d * eta * l
                    + 2 * a * d * eta * rho
                    - 2 * a * eta * l * g
                    + 2 * a * eta * rho * gh
                    + 4 * H * a * eta * l * rho * sig
                    + a ** 2 * eta ** 2
                    + a ** 2 * l ** 2
                )
            )
            * a
            + (4 * eta * l * rho * gh * g * a)
            - 0.2e1
            * eta
            * l
            * rho
            * g
            * sqrt(
                (
                    H ** 2 * eta ** 2 * sig ** 2
                    + d ** 2 * eta ** 2 * rho ** 2
                    + eta ** 2 * rho ** 2 * g ** 2
                    - 2 * d ** 2 * eta ** 2 * rho
                    - 2 * eta ** 2 * rho * g ** 2
                    + 2 * d ** 2 * eta * rho
                    + 2 * d * eta ** 2 * g
                    - 2 * d * eta * gh
                    - 2 * d * eta * g
                    - 2 * eta * gh * g
                    - 2 * H * d * eta ** 2 * rho * sig
                    - 2 * H * eta ** 2 * rho * sig * g
                    + 4 * H * d * eta * rho * sig
                    + 4 * H * eta * rho * sig * gh
                    + 2 * d * l * a
                    - 2 * H * a * eta ** 2 * rho * sig
                    - 2 * H * a * eta * l * sig
                    + 2 * a * d * eta * l * rho
                    + 2 * a * eta * l * rho * g
                    + d ** 2 * eta ** 2
                    + eta ** 2 * g ** 2
                    - 2 * d ** 2 * eta
                    + 2 * d * gh
                    + gh ** 2
                    + d ** 2
                    + 2 * d * eta * rho * gh
                    + 2 * d * eta * rho * g
                    + 2 * eta * rho * gh * g
                    + 2 * d * eta ** 2 * rho ** 2 * g
                    + 2 * H * d * eta ** 2 * sig
                    + 2 * H * eta ** 2 * sig * g
                    - 4 * d * eta ** 2 * rho * g
                    - 2 * H * d * eta * sig
                    - 2 * H * eta * sig * gh
                    + a ** 2 * eta ** 2 * rho ** 2
                    - 2 * a ** 2 * eta ** 2 * rho
                    - 2 * a ** 2 * eta * l
                    + 2 * a * d * eta ** 2
                    + 2 * a * eta ** 2 * g
                    - 2 * a * d * eta
                    - 2 * a * eta * gh
                    + 2 * a * l * gh
                    + 2 * a * d * eta ** 2 * rho ** 2
                    + 2 * a * eta ** 2 * rho ** 2 * g
                    + 2 * H * a * eta ** 2 * sig
                    + 2 * a ** 2 * eta * l * rho
                    - 4 * a * d * eta ** 2 * rho
                    - 4 * a * eta ** 2 * rho * g
                    - 2 * a * d * eta * l
                    + 2 * a * d * eta * rho
                    - 2 * a * eta * l * g
                    + 2 * a * eta * rho * gh
                    + 4 * H * a * eta * l * rho * sig
                    + a ** 2 * eta ** 2
                    + a ** 2 * l ** 2
                )
            )
            * a
            + (d ** 3)
            - (2 * H ** 2 * d * eta ** 2 * l * rho * sig ** 2)
            - (2 * H ** 2 * eta ** 2 * l * rho * sig ** 2 * g)
            + (H * d ** 2 * eta ** 2 * l * rho ** 2 * sig)
            + (H * eta ** 2 * l * rho ** 2 * sig * g ** 2)
            + (2 * H ** 2 * d * eta * l * rho * sig ** 2)
            + (2 * H ** 2 * eta * l * rho * sig ** 2 * gh)
            - (4 * H * d ** 2 * eta ** 2 * l * rho * sig)
            - (H * d * eta ** 2 * rho ** 2 * sig * gh)
            - (H * d * eta ** 2 * rho ** 2 * sig * g)
            + (3 * H * d * eta * l * rho * sig * gh)
            + (3 * H * d * eta * l * rho * sig * g)
            + (3 * H * eta * l * rho * sig * gh * g)
            + (H * d * eta ** 2 * l * rho ** 2 * sig * a)
            + (H * eta ** 2 * l * rho ** 2 * sig * g * a)
            - (7 * H * d * eta ** 2 * l * rho * sig * a)
            + (2 * H * d * eta ** 2 * l * rho ** 2 * sig * g)
            - (8 * H * d * eta ** 2 * l * rho * sig * g)
            + 0.2e1
            * d
            * gh
            * sqrt(
                (
                    H ** 2 * eta ** 2 * sig ** 2
                    + d ** 2 * eta ** 2 * rho ** 2
                    + eta ** 2 * rho ** 2 * g ** 2
                    - 2 * d ** 2 * eta ** 2 * rho
                    - 2 * eta ** 2 * rho * g ** 2
                    + 2 * d ** 2 * eta * rho
                    + 2 * d * eta ** 2 * g
                    - 2 * d * eta * gh
                    - 2 * d * eta * g
                    - 2 * eta * gh * g
                    - 2 * H * d * eta ** 2 * rho * sig
                    - 2 * H * eta ** 2 * rho * sig * g
                    + 4 * H * d * eta * rho * sig
                    + 4 * H * eta * rho * sig * gh
                    + 2 * d * l * a
                    - 2 * H * a * eta ** 2 * rho * sig
                    - 2 * H * a * eta * l * sig
                    + 2 * a * d * eta * l * rho
                    + 2 * a * eta * l * rho * g
                    + d ** 2 * eta ** 2
                    + eta ** 2 * g ** 2
                    - 2 * d ** 2 * eta
                    + 2 * d * gh
                    + gh ** 2
                    + d ** 2
                    + 2 * d * eta * rho * gh
                    + 2 * d * eta * rho * g
                    + 2 * eta * rho * gh * g
                    + 2 * d * eta ** 2 * rho ** 2 * g
                    + 2 * H * d * eta ** 2 * sig
                    + 2 * H * eta ** 2 * sig * g
                    - 4 * d * eta ** 2 * rho * g
                    - 2 * H * d * eta * sig
                    - 2 * H * eta * sig * gh
                    + a ** 2 * eta ** 2 * rho ** 2
                    - 2 * a ** 2 * eta ** 2 * rho
                    - 2 * a ** 2 * eta * l
                    + 2 * a * d * eta ** 2
                    + 2 * a * eta ** 2 * g
                    - 2 * a * d * eta
                    - 2 * a * eta * gh
                    + 2 * a * l * gh
                    + 2 * a * d * eta ** 2 * rho ** 2
                    + 2 * a * eta ** 2 * rho ** 2 * g
                    + 2 * H * a * eta ** 2 * sig
                    + 2 * a ** 2 * eta * l * rho
                    - 4 * a * d * eta ** 2 * rho
                    - 4 * a * eta ** 2 * rho * g
                    - 2 * a * d * eta * l
                    + 2 * a * d * eta * rho
                    - 2 * a * eta * l * g
                    + 2 * a * eta * rho * gh
                    + 4 * H * a * eta * l * rho * sig
                    + a ** 2 * eta ** 2
                    + a ** 2 * l ** 2
                )
            )
            + (eta ** 2 * l * a ** 3)
            - (2 * eta * l ** 2 * a ** 3)
            + (3 * d * l ** 2 * a ** 2)
            + (3 * l ** 2 * gh * a ** 2)
            + (l ** 2)
            * sqrt(
                (
                    H ** 2 * eta ** 2 * sig ** 2
                    + d ** 2 * eta ** 2 * rho ** 2
                    + eta ** 2 * rho ** 2 * g ** 2
                    - 2 * d ** 2 * eta ** 2 * rho
                    - 2 * eta ** 2 * rho * g ** 2
                    + 2 * d ** 2 * eta * rho
                    + 2 * d * eta ** 2 * g
                    - 2 * d * eta * gh
                    - 2 * d * eta * g
                    - 2 * eta * gh * g
                    - 2 * H * d * eta ** 2 * rho * sig
                    - 2 * H * eta ** 2 * rho * sig * g
                    + 4 * H * d * eta * rho * sig
                    + 4 * H * eta * rho * sig * gh
                    + 2 * d * l * a
                    - 2 * H * a * eta ** 2 * rho * sig
                    - 2 * H * a * eta * l * sig
                    + 2 * a * d * eta * l * rho
                    + 2 * a * eta * l * rho * g
                    + d ** 2 * eta ** 2
                    + eta ** 2 * g ** 2
                    - 2 * d ** 2 * eta
                    + 2 * d * gh
                    + gh ** 2
                    + d ** 2
                    + 2 * d * eta * rho * gh
                    + 2 * d * eta * rho * g
                    + 2 * eta * rho * gh * g
                    + 2 * d * eta ** 2 * rho ** 2 * g
                    + 2 * H * d * eta ** 2 * sig
                    + 2 * H * eta ** 2 * sig * g
                    - 4 * d * eta ** 2 * rho * g
                    - 2 * H * d * eta * sig
                    - 2 * H * eta * sig * gh
                    + a ** 2 * eta ** 2 * rho ** 2
                    - 2 * a ** 2 * eta ** 2 * rho
                    - 2 * a ** 2 * eta * l
                    + 2 * a * d * eta ** 2
                    + 2 * a * eta ** 2 * g
                    - 2 * a * d * eta
                    - 2 * a * eta * gh
                    + 2 * a * l * gh
                    + 2 * a * d * eta ** 2 * rho ** 2
                    + 2 * a * eta ** 2 * rho ** 2 * g
                    + 2 * H * a * eta ** 2 * sig
                    + 2 * a ** 2 * eta * l * rho
                    - 4 * a * d * eta ** 2 * rho
                    - 4 * a * eta ** 2 * rho * g
                    - 2 * a * d * eta * l
                    + 2 * a * d * eta * rho
                    - 2 * a * eta * l * g
                    + 2 * a * eta * rho * gh
                    + 4 * H * a * eta * l * rho * sig
                    + a ** 2 * eta ** 2
                    + a ** 2 * l ** 2
                )
            )
            * (a ** 2)
            - (d ** 2 * eta * a)
            + (3 * d ** 2 * l * a)
            - (eta * gh ** 2 * a)
            + (3 * l * gh ** 2 * a)
            + (d ** 3 * eta ** 2 * l)
            + (eta ** 2 * l * g ** 3)
            - (d ** 3 * eta * l)
            + (d ** 3 * eta * rho)
            - (2 * d ** 2 * eta * gh)
            - (d ** 2 * eta * g)
            - (d * eta * gh ** 2)
            - (eta * gh ** 2 * g)
            + (l ** 3 * a ** 3)
            + (d ** 2)
            * sqrt(
                (
                    H ** 2 * eta ** 2 * sig ** 2
                    + d ** 2 * eta ** 2 * rho ** 2
                    + eta ** 2 * rho ** 2 * g ** 2
                    - 2 * d ** 2 * eta ** 2 * rho
                    - 2 * eta ** 2 * rho * g ** 2
                    + 2 * d ** 2 * eta * rho
                    + 2 * d * eta ** 2 * g
                    - 2 * d * eta * gh
                    - 2 * d * eta * g
                    - 2 * eta * gh * g
                    - 2 * H * d * eta ** 2 * rho * sig
                    - 2 * H * eta ** 2 * rho * sig * g
                    + 4 * H * d * eta * rho * sig
                    + 4 * H * eta * rho * sig * gh
                    + 2 * d * l * a
                    - 2 * H * a * eta ** 2 * rho * sig
                    - 2 * H * a * eta * l * sig
                    + 2 * a * d * eta * l * rho
                    + 2 * a * eta * l * rho * g
                    + d ** 2 * eta ** 2
                    + eta ** 2 * g ** 2
                    - 2 * d ** 2 * eta
                    + 2 * d * gh
                    + gh ** 2
                    + d ** 2
                    + 2 * d * eta * rho * gh
                    + 2 * d * eta * rho * g
                    + 2 * eta * rho * gh * g
                    + 2 * d * eta ** 2 * rho ** 2 * g
                    + 2 * H * d * eta ** 2 * sig
                    + 2 * H * eta ** 2 * sig * g
                    - 4 * d * eta ** 2 * rho * g
                    - 2 * H * d * eta * sig
                    - 2 * H * eta * sig * gh
                    + a ** 2 * eta ** 2 * rho ** 2
                    - 2 * a ** 2 * eta ** 2 * rho
                    - 2 * a ** 2 * eta * l
                    + 2 * a * d * eta ** 2
                    + 2 * a * eta ** 2 * g
                    - 2 * a * d * eta
                    - 2 * a * eta * gh
                    + 2 * a * l * gh
                    + 2 * a * d * eta ** 2 * rho ** 2
                    + 2 * a * eta ** 2 * rho ** 2 * g
                    + 2 * H * a * eta ** 2 * sig
                    + 2 * a ** 2 * eta * l * rho
                    - 4 * a * d * eta ** 2 * rho
                    - 4 * a * eta ** 2 * rho * g
                    - 2 * a * d * eta * l
                    + 2 * a * d * eta * rho
                    - 2 * a * eta * l * g
                    + 2 * a * eta * rho * gh
                    + 4 * H * a * eta * l * rho * sig
                    + a ** 2 * eta ** 2
                    + a ** 2 * l ** 2
                )
            )
            + (gh ** 2)
            * sqrt(
                (
                    H ** 2 * eta ** 2 * sig ** 2
                    + d ** 2 * eta ** 2 * rho ** 2
                    + eta ** 2 * rho ** 2 * g ** 2
                    - 2 * d ** 2 * eta ** 2 * rho
                    - 2 * eta ** 2 * rho * g ** 2
                    + 2 * d ** 2 * eta * rho
                    + 2 * d * eta ** 2 * g
                    - 2 * d * eta * gh
                    - 2 * d * eta * g
                    - 2 * eta * gh * g
                    - 2 * H * d * eta ** 2 * rho * sig
                    - 2 * H * eta ** 2 * rho * sig * g
                    + 4 * H * d * eta * rho * sig
                    + 4 * H * eta * rho * sig * gh
                    + 2 * d * l * a
                    - 2 * H * a * eta ** 2 * rho * sig
                    - 2 * H * a * eta * l * sig
                    + 2 * a * d * eta * l * rho
                    + 2 * a * eta * l * rho * g
                    + d ** 2 * eta ** 2
                    + eta ** 2 * g ** 2
                    - 2 * d ** 2 * eta
                    + 2 * d * gh
                    + gh ** 2
                    + d ** 2
                    + 2 * d * eta * rho * gh
                    + 2 * d * eta * rho * g
                    + 2 * eta * rho * gh * g
                    + 2 * d * eta ** 2 * rho ** 2 * g
                    + 2 * H * d * eta ** 2 * sig
                    + 2 * H * eta ** 2 * sig * g
                    - 4 * d * eta ** 2 * rho * g
                    - 2 * H * d * eta * sig
                    - 2 * H * eta * sig * gh
                    + a ** 2 * eta ** 2 * rho ** 2
                    - 2 * a ** 2 * eta ** 2 * rho
                    - 2 * a ** 2 * eta * l
                    + 2 * a * d * eta ** 2
                    + 2 * a * eta ** 2 * g
                    - 2 * a * d * eta
                    - 2 * a * eta * gh
                    + 2 * a * l * gh
                    + 2 * a * d * eta ** 2 * rho ** 2
                    + 2 * a * eta ** 2 * rho ** 2 * g
                    + 2 * H * a * eta ** 2 * sig
                    + 2 * a ** 2 * eta * l * rho
                    - 4 * a * d * eta ** 2 * rho
                    - 4 * a * eta ** 2 * rho * g
                    - 2 * a * d * eta * l
                    + 2 * a * d * eta * rho
                    - 2 * a * eta * l * g
                    + 2 * a * eta * rho * gh
                    + 4 * H * a * eta * l * rho * sig
                    + a ** 2 * eta ** 2
                    + a ** 2 * l ** 2
                )
            )
            - (d ** 3 * eta)
            + (3 * d ** 2 * gh)
            + (3 * d * gh ** 2)
            - (2 * d * eta * gh * a)
            + (6 * d * l * gh * a)
            + 0.2e1
            * d
            * l
            * sqrt(
                (
                    H ** 2 * eta ** 2 * sig ** 2
                    + d ** 2 * eta ** 2 * rho ** 2
                    + eta ** 2 * rho ** 2 * g ** 2
                    - 2 * d ** 2 * eta ** 2 * rho
                    - 2 * eta ** 2 * rho * g ** 2
                    + 2 * d ** 2 * eta * rho
                    + 2 * d * eta ** 2 * g
                    - 2 * d * eta * gh
                    - 2 * d * eta * g
                    - 2 * eta * gh * g
                    - 2 * H * d * eta ** 2 * rho * sig
                    - 2 * H * eta ** 2 * rho * sig * g
                    + 4 * H * d * eta * rho * sig
                    + 4 * H * eta * rho * sig * gh
                    + 2 * d * l * a
                    - 2 * H * a * eta ** 2 * rho * sig
                    - 2 * H * a * eta * l * sig
                    + 2 * a * d * eta * l * rho
                    + 2 * a * eta * l * rho * g
                    + d ** 2 * eta ** 2
                    + eta ** 2 * g ** 2
                    - 2 * d ** 2 * eta
                    + 2 * d * gh
                    + gh ** 2
                    + d ** 2
                    + 2 * d * eta * rho * gh
                    + 2 * d * eta * rho * g
                    + 2 * eta * rho * gh * g
                    + 2 * d * eta ** 2 * rho ** 2 * g
                    + 2 * H * d * eta ** 2 * sig
                    + 2 * H * eta ** 2 * sig * g
                    - 4 * d * eta ** 2 * rho * g
                    - 2 * H * d * eta * sig
                    - 2 * H * eta * sig * gh
                    + a ** 2 * eta ** 2 * rho ** 2
                    - 2 * a ** 2 * eta ** 2 * rho
                    - 2 * a ** 2 * eta * l
                    + 2 * a * d * eta ** 2
                    + 2 * a * eta ** 2 * g
                    - 2 * a * d * eta
                    - 2 * a * eta * gh
                    + 2 * a * l * gh
                    + 2 * a * d * eta ** 2 * rho ** 2
                    + 2 * a * eta ** 2 * rho ** 2 * g
                    + 2 * H * a * eta ** 2 * sig
                    + 2 * a ** 2 * eta * l * rho
                    - 4 * a * d * eta ** 2 * rho
                    - 4 * a * eta ** 2 * rho * g
                    - 2 * a * d * eta * l
                    + 2 * a * d * eta * rho
                    - 2 * a * eta * l * g
                    + 2 * a * eta * rho * gh
                    + 4 * H * a * eta * l * rho * sig
                    + a ** 2 * eta ** 2
                    + a ** 2 * l ** 2
                )
            )
            * a
            + 0.2e1
            * l
            * gh
            * sqrt(
                (
                    H ** 2 * eta ** 2 * sig ** 2
                    + d ** 2 * eta ** 2 * rho ** 2
                    + eta ** 2 * rho ** 2 * g ** 2
                    - 2 * d ** 2 * eta ** 2 * rho
                    - 2 * eta ** 2 * rho * g ** 2
                    + 2 * d ** 2 * eta * rho
                    + 2 * d * eta ** 2 * g
                    - 2 * d * eta * gh
                    - 2 * d * eta * g
                    - 2 * eta * gh * g
                    - 2 * H * d * eta ** 2 * rho * sig
                    - 2 * H * eta ** 2 * rho * sig * g
                    + 4 * H * d * eta * rho * sig
                    + 4 * H * eta * rho * sig * gh
                    + 2 * d * l * a
                    - 2 * H * a * eta ** 2 * rho * sig
                    - 2 * H * a * eta * l * sig
                    + 2 * a * d * eta * l * rho
                    + 2 * a * eta * l * rho * g
                    + d ** 2 * eta ** 2
                    + eta ** 2 * g ** 2
                    - 2 * d ** 2 * eta
                    + 2 * d * gh
                    + gh ** 2
                    + d ** 2
                    + 2 * d * eta * rho * gh
                    + 2 * d * eta * rho * g
                    + 2 * eta * rho * gh * g
                    + 2 * d * eta ** 2 * rho ** 2 * g
                    + 2 * H * d * eta ** 2 * sig
                    + 2 * H * eta ** 2 * sig * g
                    - 4 * d * eta ** 2 * rho * g
                    - 2 * H * d * eta * sig
                    - 2 * H * eta * sig * gh
                    + a ** 2 * eta ** 2 * rho ** 2
                    - 2 * a ** 2 * eta ** 2 * rho
                    - 2 * a ** 2 * eta * l
                    + 2 * a * d * eta ** 2
                    + 2 * a * eta ** 2 * g
                    - 2 * a * d * eta
                    - 2 * a * eta * gh
                    + 2 * a * l * gh
                    + 2 * a * d * eta ** 2 * rho ** 2
                    + 2 * a * eta ** 2 * rho ** 2 * g
                    + 2 * H * a * eta ** 2 * sig
                    + 2 * a ** 2 * eta * l * rho
                    - 4 * a * d * eta ** 2 * rho
                    - 4 * a * eta ** 2 * rho * g
                    - 2 * a * d * eta * l
                    + 2 * a * d * eta * rho
                    - 2 * a * eta * l * g
                    + 2 * a * eta * rho * gh
                    + 4 * H * a * eta * l * rho * sig
                    + a ** 2 * eta ** 2
                    + a ** 2 * l ** 2
                )
            )
            * a
            - (3 * d * eta * l ** 2 * a ** 2)
            + (3 * eta ** 2 * l * g ** 2 * a)
            + (3 * eta ** 2 * l * g * a ** 2)
            - (eta * l ** 2 * g ** 2 * a)
            - (3 * eta * l ** 2 * g * a ** 2)
            + (d ** 2)
            * eta
            * l
            * sqrt(
                (
                    H ** 2 * eta ** 2 * sig ** 2
                    + d ** 2 * eta ** 2 * rho ** 2
                    + eta ** 2 * rho ** 2 * g ** 2
                    - 2 * d ** 2 * eta ** 2 * rho
                    - 2 * eta ** 2 * rho * g ** 2
                    + 2 * d ** 2 * eta * rho
                    + 2 * d * eta ** 2 * g
                    - 2 * d * eta * gh
                    - 2 * d * eta * g
                    - 2 * eta * gh * g
                    - 2 * H * d * eta ** 2 * rho * sig
                    - 2 * H * eta ** 2 * rho * sig * g
                    + 4 * H * d * eta * rho * sig
                    + 4 * H * eta * rho * sig * gh
                    + 2 * d * l * a
                    - 2 * H * a * eta ** 2 * rho * sig
                    - 2 * H * a * eta * l * sig
                    + 2 * a * d * eta * l * rho
                    + 2 * a * eta * l * rho * g
                    + d ** 2 * eta ** 2
                    + eta ** 2 * g ** 2
                    - 2 * d ** 2 * eta
                    + 2 * d * gh
                    + gh ** 2
                    + d ** 2
                    + 2 * d * eta * rho * gh
                    + 2 * d * eta * rho * g
                    + 2 * eta * rho * gh * g
                    + 2 * d * eta ** 2 * rho ** 2 * g
                    + 2 * H * d * eta ** 2 * sig
                    + 2 * H * eta ** 2 * sig * g
                    - 4 * d * eta ** 2 * rho * g
                    - 2 * H * d * eta * sig
                    - 2 * H * eta * sig * gh
                    + a ** 2 * eta ** 2 * rho ** 2
                    - 2 * a ** 2 * eta ** 2 * rho
                    - 2 * a ** 2 * eta * l
                    + 2 * a * d * eta ** 2
                    + 2 * a * eta ** 2 * g
                    - 2 * a * d * eta
                    - 2 * a * eta * gh
                    + 2 * a * l * gh
                    + 2 * a * d * eta ** 2 * rho ** 2
                    + 2 * a * eta ** 2 * rho ** 2 * g
                    + 2 * H * a * eta ** 2 * sig
                    + 2 * a ** 2 * eta * l * rho
                    - 4 * a * d * eta ** 2 * rho
                    - 4 * a * eta ** 2 * rho * g
                    - 2 * a * d * eta * l
                    + 2 * a * d * eta * rho
                    - 2 * a * eta * l * g
                    + 2 * a * eta * rho * gh
                    + 4 * H * a * eta * l * rho * sig
                    + a ** 2 * eta ** 2
                    + a ** 2 * l ** 2
                )
            )
            - (4 * d ** 2 * eta * l * a)
            + (d ** 2 * eta * rho * a)
            - (3 * d * eta * l * a ** 2)
            - (3 * eta * l * gh * a ** 2)
            + eta
            * l
            * (g ** 2)
            * sqrt(
                (
                    H ** 2 * eta ** 2 * sig ** 2
                    + d ** 2 * eta ** 2 * rho ** 2
                    + eta ** 2 * rho ** 2 * g ** 2
                    - 2 * d ** 2 * eta ** 2 * rho
                    - 2 * eta ** 2 * rho * g ** 2
                    + 2 * d ** 2 * eta * rho
                    + 2 * d * eta ** 2 * g
                    - 2 * d * eta * gh
                    - 2 * d * eta * g
                    - 2 * eta * gh * g
                    - 2 * H * d * eta ** 2 * rho * sig
                    - 2 * H * eta ** 2 * rho * sig * g
                    + 4 * H * d * eta * rho * sig
                    + 4 * H * eta * rho * sig * gh
                    + 2 * d * l * a
                    - 2 * H * a * eta ** 2 * rho * sig
                    - 2 * H * a * eta * l * sig
                    + 2 * a * d * eta * l * rho
                    + 2 * a * eta * l * rho * g
                    + d ** 2 * eta ** 2
                    + eta ** 2 * g ** 2
                    - 2 * d ** 2 * eta
                    + 2 * d * gh
                    + gh ** 2
                    + d ** 2
                    + 2 * d * eta * rho * gh
                    + 2 * d * eta * rho * g
                    + 2 * eta * rho * gh * g
                    + 2 * d * eta ** 2 * rho ** 2 * g
                    + 2 * H * d * eta ** 2 * sig
                    + 2 * H * eta ** 2 * sig * g
                    - 4 * d * eta ** 2 * rho * g
                    - 2 * H * d * eta * sig
                    - 2 * H * eta * sig * gh
                    + a ** 2 * eta ** 2 * rho ** 2
                    - 2 * a ** 2 * eta ** 2 * rho
                    - 2 * a ** 2 * eta * l
                    + 2 * a * d * eta ** 2
                    + 2 * a * eta ** 2 * g
                    - 2 * a * d * eta
                    - 2 * a * eta * gh
                    + 2 * a * l * gh
                    + 2 * a * d * eta ** 2 * rho ** 2
                    + 2 * a * eta ** 2 * rho ** 2 * g
                    + 2 * H * a * eta ** 2 * sig
                    + 2 * a ** 2 * eta * l * rho
                    - 4 * a * d * eta ** 2 * rho
                    - 4 * a * eta ** 2 * rho * g
                    - 2 * a * d * eta * l
                    + 2 * a * d * eta * rho
                    - 2 * a * eta * l * g
                    + 2 * a * eta * rho * gh
                    + 4 * H * a * eta * l * rho * sig
                    + a ** 2 * eta ** 2
                    + a ** 2 * l ** 2
                )
            )
            + eta
            * l
            * sqrt(
                (
                    H ** 2 * eta ** 2 * sig ** 2
                    + d ** 2 * eta ** 2 * rho ** 2
                    + eta ** 2 * rho ** 2 * g ** 2
                    - 2 * d ** 2 * eta ** 2 * rho
                    - 2 * eta ** 2 * rho * g ** 2
                    + 2 * d ** 2 * eta * rho
                    + 2 * d * eta ** 2 * g
                    - 2 * d * eta * gh
                    - 2 * d * eta * g
                    - 2 * eta * gh * g
                    - 2 * H * d * eta ** 2 * rho * sig
                    - 2 * H * eta ** 2 * rho * sig * g
                    + 4 * H * d * eta * rho * sig
                    + 4 * H * eta * rho * sig * gh
                    + 2 * d * l * a
                    - 2 * H * a * eta ** 2 * rho * sig
                    - 2 * H * a * eta * l * sig
                    + 2 * a * d * eta * l * rho
                    + 2 * a * eta * l * rho * g
                    + d ** 2 * eta ** 2
                    + eta ** 2 * g ** 2
                    - 2 * d ** 2 * eta
                    + 2 * d * gh
                    + gh ** 2
                    + d ** 2
                    + 2 * d * eta * rho * gh
                    + 2 * d * eta * rho * g
                    + 2 * eta * rho * gh * g
                    + 2 * d * eta ** 2 * rho ** 2 * g
                    + 2 * H * d * eta ** 2 * sig
                    + 2 * H * eta ** 2 * sig * g
                    - 4 * d * eta ** 2 * rho * g
                    - 2 * H * d * eta * sig
                    - 2 * H * eta * sig * gh
                    + a ** 2 * eta ** 2 * rho ** 2
                    - 2 * a ** 2 * eta ** 2 * rho
                    - 2 * a ** 2 * eta * l
                    + 2 * a * d * eta ** 2
                    + 2 * a * eta ** 2 * g
                    - 2 * a * d * eta
                    - 2 * a * eta * gh
                    + 2 * a * l * gh
                    + 2 * a * d * eta ** 2 * rho ** 2
                    + 2 * a * eta ** 2 * rho ** 2 * g
                    + 2 * H * a * eta ** 2 * sig
                    + 2 * a ** 2 * eta * l * rho
                    - 4 * a * d * eta ** 2 * rho
                    - 4 * a * eta ** 2 * rho * g
                    - 2 * a * d * eta * l
                    + 2 * a * d * eta * rho
                    - 2 * a * eta * l * g
                    + 2 * a * eta * rho * gh
                    + 4 * H * a * eta * l * rho * sig
                    + a ** 2 * eta ** 2
                    + a ** 2 * l ** 2
                )
            )
            * (a ** 2)
            + (eta * rho * gh ** 2 * a)
            + (eta ** 2 * l * rho ** 2 * a ** 3)
            - (2 * eta ** 2 * l * rho * a ** 3)
            + (2 * eta * l ** 2 * rho * a ** 3)
            + (3 * d ** 2 * eta ** 2 * l * a)
            - (d ** 2 * eta * l ** 2 * a)
            + (3 * d * eta ** 2 * l * a ** 2)
            + (d ** 3 * eta * l * rho)
            + (3 * d ** 2 * eta ** 2 * l * g)
            + (3 * d * eta ** 2 * l * g ** 2)
            - (H * d ** 2 * eta * sig)
            - (H * eta * sig * gh ** 2)
            - (d ** 2 * eta * l * gh)
            - (2 * d ** 2 * eta * l * g)
            + (2 * d ** 2 * eta * rho * gh)
            + (d ** 2 * eta * rho * g)
            - (d * eta * l * g ** 2)
            + (d * eta * rho * gh ** 2)
            - (eta * l * gh * g ** 2)
            + (eta * rho * gh ** 2 * g)
            - (2 * d * eta * gh * g)
            + (H ** 3 * eta ** 2 * l * sig ** 3)
            + (d ** 3 * eta ** 2 * l * rho ** 2)
            + (eta ** 2 * l * rho ** 2 * g ** 3)
            - (2 * d ** 3 * eta ** 2 * l * rho)
            - (2 * eta ** 2 * l * rho * g ** 3)
            + (3 * eta * l * rho * gh * a ** 2)
            - eta
            * l
            * rho
            * (g ** 2)
            * sqrt(
                (
                    H ** 2 * eta ** 2 * sig ** 2
                    + d ** 2 * eta ** 2 * rho ** 2
                    + eta ** 2 * rho ** 2 * g ** 2
                    - 2 * d ** 2 * eta ** 2 * rho
                    - 2 * eta ** 2 * rho * g ** 2
                    + 2 * d ** 2 * eta * rho
                    + 2 * d * eta ** 2 * g
                    - 2 * d * eta * gh
                    - 2 * d * eta * g
                    - 2 * eta * gh * g
                    - 2 * H * d * eta ** 2 * rho * sig
                    - 2 * H * eta ** 2 * rho * sig * g
                    + 4 * H * d * eta * rho * sig
                    + 4 * H * eta * rho * sig * gh
                    + 2 * d * l * a
                    - 2 * H * a * eta ** 2 * rho * sig
                    - 2 * H * a * eta * l * sig
                    + 2 * a * d * eta * l * rho
                    + 2 * a * eta * l * rho * g
                    + d ** 2 * eta ** 2
                    + eta ** 2 * g ** 2
                    - 2 * d ** 2 * eta
                    + 2 * d * gh
                    + gh ** 2
                    + d ** 2
                    + 2 * d * eta * rho * gh
                    + 2 * d * eta * rho * g
                    + 2 * eta * rho * gh * g
                    + 2 * d * eta ** 2 * rho ** 2 * g
                    + 2 * H * d * eta ** 2 * sig
                    + 2 * H * eta ** 2 * sig * g
                    - 4 * d * eta ** 2 * rho * g
                    - 2 * H * d * eta * sig
                    - 2 * H * eta * sig * gh
                    + a ** 2 * eta ** 2 * rho ** 2
                    - 2 * a ** 2 * eta ** 2 * rho
                    - 2 * a ** 2 * eta * l
                    + 2 * a * d * eta ** 2
                    + 2 * a * eta ** 2 * g
                    - 2 * a * d * eta
                    - 2 * a * eta * gh
                    + 2 * a * l * gh
                    + 2 * a * d * eta ** 2 * rho ** 2
                    + 2 * a * eta ** 2 * rho ** 2 * g
                    + 2 * H * a * eta ** 2 * sig
                    + 2 * a ** 2 * eta * l * rho
                    - 4 * a * d * eta ** 2 * rho
                    - 4 * a * eta ** 2 * rho * g
                    - 2 * a * d * eta * l
                    + 2 * a * d * eta * rho
                    - 2 * a * eta * l * g
                    + 2 * a * eta * rho * gh
                    + 4 * H * a * eta * l * rho * sig
                    + a ** 2 * eta ** 2
                    + a ** 2 * l ** 2
                )
            )
            - eta
            * l
            * rho
            * sqrt(
                (
                    H ** 2 * eta ** 2 * sig ** 2
                    + d ** 2 * eta ** 2 * rho ** 2
                    + eta ** 2 * rho ** 2 * g ** 2
                    - 2 * d ** 2 * eta ** 2 * rho
                    - 2 * eta ** 2 * rho * g ** 2
                    + 2 * d ** 2 * eta * rho
                    + 2 * d * eta ** 2 * g
                    - 2 * d * eta * gh
                    - 2 * d * eta * g
                    - 2 * eta * gh * g
                    - 2 * H * d * eta ** 2 * rho * sig
                    - 2 * H * eta ** 2 * rho * sig * g
                    + 4 * H * d * eta * rho * sig
                    + 4 * H * eta * rho * sig * gh
                    + 2 * d * l * a
                    - 2 * H * a * eta ** 2 * rho * sig
                    - 2 * H * a * eta * l * sig
                    + 2 * a * d * eta * l * rho
                    + 2 * a * eta * l * rho * g
                    + d ** 2 * eta ** 2
                    + eta ** 2 * g ** 2
                    - 2 * d ** 2 * eta
                    + 2 * d * gh
                    + gh ** 2
                    + d ** 2
                    + 2 * d * eta * rho * gh
                    + 2 * d * eta * rho * g
                    + 2 * eta * rho * gh * g
                    + 2 * d * eta ** 2 * rho ** 2 * g
                    + 2 * H * d * eta ** 2 * sig
                    + 2 * H * eta ** 2 * sig * g
                    - 4 * d * eta ** 2 * rho * g
                    - 2 * H * d * eta * sig
                    - 2 * H * eta * sig * gh
                    + a ** 2 * eta ** 2 * rho ** 2
                    - 2 * a ** 2 * eta ** 2 * rho
                    - 2 * a ** 2 * eta * l
                    + 2 * a * d * eta ** 2
                    + 2 * a * eta ** 2 * g
                    - 2 * a * d * eta
                    - 2 * a * eta * gh
                    + 2 * a * l * gh
                    + 2 * a * d * eta ** 2 * rho ** 2
                    + 2 * a * eta ** 2 * rho ** 2 * g
                    + 2 * H * a * eta ** 2 * sig
                    + 2 * a ** 2 * eta * l * rho
                    - 4 * a * d * eta ** 2 * rho
                    - 4 * a * eta ** 2 * rho * g
                    - 2 * a * d * eta * l
                    + 2 * a * d * eta * rho
                    - 2 * a * eta * l * g
                    + 2 * a * eta * rho * gh
                    + 4 * H * a * eta * l * rho * sig
                    + a ** 2 * eta ** 2
                    + a ** 2 * l ** 2
                )
            )
            * (a ** 2)
            - (4 * d * eta * l * gh * a)
            + 0.2e1
            * d
            * eta
            * l
            * g
            * sqrt(
                (
                    H ** 2 * eta ** 2 * sig ** 2
                    + d ** 2 * eta ** 2 * rho ** 2
                    + eta ** 2 * rho ** 2 * g ** 2
                    - 2 * d ** 2 * eta ** 2 * rho
                    - 2 * eta ** 2 * rho * g ** 2
                    + 2 * d ** 2 * eta * rho
                    + 2 * d * eta ** 2 * g
                    - 2 * d * eta * gh
                    - 2 * d * eta * g
                    - 2 * eta * gh * g
                    - 2 * H * d * eta ** 2 * rho * sig
                    - 2 * H * eta ** 2 * rho * sig * g
                    + 4 * H * d * eta * rho * sig
                    + 4 * H * eta * rho * sig * gh
                    + 2 * d * l * a
                    - 2 * H * a * eta ** 2 * rho * sig
                    - 2 * H * a * eta * l * sig
                    + 2 * a * d * eta * l * rho
                    + 2 * a * eta * l * rho * g
                    + d ** 2 * eta ** 2
                    + eta ** 2 * g ** 2
                    - 2 * d ** 2 * eta
                    + 2 * d * gh
                    + gh ** 2
                    + d ** 2
                    + 2 * d * eta * rho * gh
                    + 2 * d * eta * rho * g
                    + 2 * eta * rho * gh * g
                    + 2 * d * eta ** 2 * rho ** 2 * g
                    + 2 * H * d * eta ** 2 * sig
                    + 2 * H * eta ** 2 * sig * g
                    - 4 * d * eta ** 2 * rho * g
                    - 2 * H * d * eta * sig
                    - 2 * H * eta * sig * gh
                    + a ** 2 * eta ** 2 * rho ** 2
                    - 2 * a ** 2 * eta ** 2 * rho
                    - 2 * a ** 2 * eta * l
                    + 2 * a * d * eta ** 2
                    + 2 * a * eta ** 2 * g
                    - 2 * a * d * eta
                    - 2 * a * eta * gh
                    + 2 * a * l * gh
                    + 2 * a * d * eta ** 2 * rho ** 2
                    + 2 * a * eta ** 2 * rho ** 2 * g
                    + 2 * H * a * eta ** 2 * sig
                    + 2 * a ** 2 * eta * l * rho
                    - 4 * a * d * eta ** 2 * rho
                    - 4 * a * eta ** 2 * rho * g
                    - 2 * a * d * eta * l
                    + 2 * a * d * eta * rho
                    - 2 * a * eta * l * g
                    + 2 * a * eta * rho * gh
                    + 4 * H * a * eta * l * rho * sig
                    + a ** 2 * eta ** 2
                    + a ** 2 * l ** 2
                )
            )
            - (4 * d * eta * l * g * a)
            + 0.2e1
            * d
            * eta
            * l
            * sqrt(
                (
                    H ** 2 * eta ** 2 * sig ** 2
                    + d ** 2 * eta ** 2 * rho ** 2
                    + eta ** 2 * rho ** 2 * g ** 2
                    - 2 * d ** 2 * eta ** 2 * rho
                    - 2 * eta ** 2 * rho * g ** 2
                    + 2 * d ** 2 * eta * rho
                    + 2 * d * eta ** 2 * g
                    - 2 * d * eta * gh
                    - 2 * d * eta * g
                    - 2 * eta * gh * g
                    - 2 * H * d * eta ** 2 * rho * sig
                    - 2 * H * eta ** 2 * rho * sig * g
                    + 4 * H * d * eta * rho * sig
                    + 4 * H * eta * rho * sig * gh
                    + 2 * d * l * a
                    - 2 * H * a * eta ** 2 * rho * sig
                    - 2 * H * a * eta * l * sig
                    + 2 * a * d * eta * l * rho
                    + 2 * a * eta * l * rho * g
                    + d ** 2 * eta ** 2
                    + eta ** 2 * g ** 2
                    - 2 * d ** 2 * eta
                    + 2 * d * gh
                    + gh ** 2
                    + d ** 2
                    + 2 * d * eta * rho * gh
                    + 2 * d * eta * rho * g
                    + 2 * eta * rho * gh * g
                    + 2 * d * eta ** 2 * rho ** 2 * g
                    + 2 * H * d * eta ** 2 * sig
                    + 2 * H * eta ** 2 * sig * g
                    - 4 * d * eta ** 2 * rho * g
                    - 2 * H * d * eta * sig
                    - 2 * H * eta * sig * gh
                    + a ** 2 * eta ** 2 * rho ** 2
                    - 2 * a ** 2 * eta ** 2 * rho
                    - 2 * a ** 2 * eta * l
                    + 2 * a * d * eta ** 2
                    + 2 * a * eta ** 2 * g
                    - 2 * a * d * eta
                    - 2 * a * eta * gh
                    + 2 * a * l * gh
                    + 2 * a * d * eta ** 2 * rho ** 2
                    + 2 * a * eta ** 2 * rho ** 2 * g
                    + 2 * H * a * eta ** 2 * sig
                    + 2 * a ** 2 * eta * l * rho
                    - 4 * a * d * eta ** 2 * rho
                    - 4 * a * eta ** 2 * rho * g
                    - 2 * a * d * eta * l
                    + 2 * a * d * eta * rho
                    - 2 * a * eta * l * g
                    + 2 * a * eta * rho * gh
                    + 4 * H * a * eta * l * rho * sig
                    + a ** 2 * eta ** 2
                    + a ** 2 * l ** 2
                )
            )
            * a
            + (2 * d * eta * rho * gh * a)
            - (4 * eta * l * gh * g * a)
            + 0.2e1
            * eta
            * l
            * g
            * sqrt(
                (
                    H ** 2 * eta ** 2 * sig ** 2
                    + d ** 2 * eta ** 2 * rho ** 2
                    + eta ** 2 * rho ** 2 * g ** 2
                    - 2 * d ** 2 * eta ** 2 * rho
                    - 2 * eta ** 2 * rho * g ** 2
                    + 2 * d ** 2 * eta * rho
                    + 2 * d * eta ** 2 * g
                    - 2 * d * eta * gh
                    - 2 * d * eta * g
                    - 2 * eta * gh * g
                    - 2 * H * d * eta ** 2 * rho * sig
                    - 2 * H * eta ** 2 * rho * sig * g
                    + 4 * H * d * eta * rho * sig
                    + 4 * H * eta * rho * sig * gh
                    + 2 * d * l * a
                    - 2 * H * a * eta ** 2 * rho * sig
                    - 2 * H * a * eta * l * sig
                    + 2 * a * d * eta * l * rho
                    + 2 * a * eta * l * rho * g
                    + d ** 2 * eta ** 2
                    + eta ** 2 * g ** 2
                    - 2 * d ** 2 * eta
                    + 2 * d * gh
                    + gh ** 2
                    + d ** 2
                    + 2 * d * eta * rho * gh
                    + 2 * d * eta * rho * g
                    + 2 * eta * rho * gh * g
                    + 2 * d * eta ** 2 * rho ** 2 * g
                    + 2 * H * d * eta ** 2 * sig
                    + 2 * H * eta ** 2 * sig * g
                    - 4 * d * eta ** 2 * rho * g
                    - 2 * H * d * eta * sig
                    - 2 * H * eta * sig * gh
                    + a ** 2 * eta ** 2 * rho ** 2
                    - 2 * a ** 2 * eta ** 2 * rho
                    - 2 * a ** 2 * eta * l
                    + 2 * a * d * eta ** 2
                    + 2 * a * eta ** 2 * g
                    - 2 * a * d * eta
                    - 2 * a * eta * gh
                    + 2 * a * l * gh
                    + 2 * a * d * eta ** 2 * rho ** 2
                    + 2 * a * eta ** 2 * rho ** 2 * g
                    + 2 * H * a * eta ** 2 * sig
                    + 2 * a ** 2 * eta * l * rho
                    - 4 * a * d * eta ** 2 * rho
                    - 4 * a * eta ** 2 * rho * g
                    - 2 * a * d * eta * l
                    + 2 * a * d * eta * rho
                    - 2 * a * eta * l * g
                    + 2 * a * eta * rho * gh
                    + 4 * H * a * eta * l * rho * sig
                    + a ** 2 * eta ** 2
                    + a ** 2 * l ** 2
                )
            )
            * a
            + (3 * H * eta ** 2 * l * sig * a ** 2)
            - (3 * H * eta * l ** 2 * sig * a ** 2)
            - (6 * d ** 2 * eta ** 2 * l * rho * a)
            + (d ** 2 * eta * l ** 2 * rho * a)
            - (6 * d * eta ** 2 * l * rho * a ** 2)
            + (3 * d * eta * l ** 2 * rho * a ** 2)
            - (6 * eta ** 2 * l * rho * g ** 2 * a)
            - (6 * eta ** 2 * l * rho * g * a ** 2)
            + (eta * l ** 2 * rho * g ** 2 * a)
            + (3 * eta * l ** 2 * rho * g * a ** 2)
            - (d ** 2)
            * eta
            * l
            * rho
            * sqrt(
                (
                    H ** 2 * eta ** 2 * sig ** 2
                    + d ** 2 * eta ** 2 * rho ** 2
                    + eta ** 2 * rho ** 2 * g ** 2
                    - 2 * d ** 2 * eta ** 2 * rho
                    - 2 * eta ** 2 * rho * g ** 2
                    + 2 * d ** 2 * eta * rho
                    + 2 * d * eta ** 2 * g
                    - 2 * d * eta * gh
                    - 2 * d * eta * g
                    - 2 * eta * gh * g
                    - 2 * H * d * eta ** 2 * rho * sig
                    - 2 * H * eta ** 2 * rho * sig * g
                    + 4 * H * d * eta * rho * sig
                    + 4 * H * eta * rho * sig * gh
                    + 2 * d * l * a
                    - 2 * H * a * eta ** 2 * rho * sig
                    - 2 * H * a * eta * l * sig
                    + 2 * a * d * eta * l * rho
                    + 2 * a * eta * l * rho * g
                    + d ** 2 * eta ** 2
                    + eta ** 2 * g ** 2
                    - 2 * d ** 2 * eta
                    + 2 * d * gh
                    + gh ** 2
                    + d ** 2
                    + 2 * d * eta * rho * gh
                    + 2 * d * eta * rho * g
                    + 2 * eta * rho * gh * g
                    + 2 * d * eta ** 2 * rho ** 2 * g
                    + 2 * H * d * eta ** 2 * sig
                    + 2 * H * eta ** 2 * sig * g
                    - 4 * d * eta ** 2 * rho * g
                    - 2 * H * d * eta * sig
                    - 2 * H * eta * sig * gh
                    + a ** 2 * eta ** 2 * rho ** 2
                    - 2 * a ** 2 * eta ** 2 * rho
                    - 2 * a ** 2 * eta * l
                    + 2 * a * d * eta ** 2
                    + 2 * a * eta ** 2 * g
                    - 2 * a * d * eta
                    - 2 * a * eta * gh
                    + 2 * a * l * gh
                    + 2 * a * d * eta ** 2 * rho ** 2
                    + 2 * a * eta ** 2 * rho ** 2 * g
                    + 2 * H * a * eta ** 2 * sig
                    + 2 * a ** 2 * eta * l * rho
                    - 4 * a * d * eta ** 2 * rho
                    - 4 * a * eta ** 2 * rho * g
                    - 2 * a * d * eta * l
                    + 2 * a * d * eta * rho
                    - 2 * a * eta * l * g
                    + 2 * a * eta * rho * gh
                    + 4 * H * a * eta * l * rho * sig
                    + a ** 2 * eta ** 2
                    + a ** 2 * l ** 2
                )
            )
            + (4 * d ** 2 * eta * l * rho * a)
            + (6 * d * eta ** 2 * l * g * a)
            - (2 * d * eta * l ** 2 * g * a)
            + (3 * d * eta * l * rho * a ** 2)
            + (2 * d ** 2 * eta * l * rho * g)
            + (d * eta * l * rho * g ** 2)
            + (eta * l * rho * gh * g ** 2)
            - (2 * H * d * eta * sig * gh)
            - (2 * d * eta * l * gh * g)
            + (2 * d * eta * rho * gh * g)
            + (3 * H ** 2 * eta ** 2 * l * sig ** 2 * a)
            - (H ** 2 * eta * l ** 2 * sig ** 2 * a)
            + (3 * d ** 2 * eta ** 2 * l * rho ** 2 * a)
            + (3 * d * eta ** 2 * l * rho ** 2 * a ** 2)
            + (3 * eta ** 2 * l * rho ** 2 * g ** 2 * a)
            + (3 * eta ** 2 * l * rho ** 2 * g * a ** 2)
            + (H ** 2)
            * eta
            * l
            * (sig ** 2)
            * sqrt(
                (
                    H ** 2 * eta ** 2 * sig ** 2
                    + d ** 2 * eta ** 2 * rho ** 2
                    + eta ** 2 * rho ** 2 * g ** 2
                    - 2 * d ** 2 * eta ** 2 * rho
                    - 2 * eta ** 2 * rho * g ** 2
                    + 2 * d ** 2 * eta * rho
                    + 2 * d * eta ** 2 * g
                    - 2 * d * eta * gh
                    - 2 * d * eta * g
                    - 2 * eta * gh * g
                    - 2 * H * d * eta ** 2 * rho * sig
                    - 2 * H * eta ** 2 * rho * sig * g
                    + 4 * H * d * eta * rho * sig
                    + 4 * H * eta * rho * sig * gh
                    + 2 * d * l * a
                    - 2 * H * a * eta ** 2 * rho * sig
                    - 2 * H * a * eta * l * sig
                    + 2 * a * d * eta * l * rho
                    + 2 * a * eta * l * rho * g
                    + d ** 2 * eta ** 2
                    + eta ** 2 * g ** 2
                    - 2 * d ** 2 * eta
                    + 2 * d * gh
                    + gh ** 2
                    + d ** 2
                    + 2 * d * eta * rho * gh
                    + 2 * d * eta * rho * g
                    + 2 * eta * rho * gh * g
                    + 2 * d * eta ** 2 * rho ** 2 * g
                    + 2 * H * d * eta ** 2 * sig
                    + 2 * H * eta ** 2 * sig * g
                    - 4 * d * eta ** 2 * rho * g
                    - 2 * H * d * eta * sig
                    - 2 * H * eta * sig * gh
                    + a ** 2 * eta ** 2 * rho ** 2
                    - 2 * a ** 2 * eta ** 2 * rho
                    - 2 * a ** 2 * eta * l
                    + 2 * a * d * eta ** 2
                    + 2 * a * eta ** 2 * g
                    - 2 * a * d * eta
                    - 2 * a * eta * gh
                    + 2 * a * l * gh
                    + 2 * a * d * eta ** 2 * rho ** 2
                    + 2 * a * eta ** 2 * rho ** 2 * g
                    + 2 * H * a * eta ** 2 * sig
                    + 2 * a ** 2 * eta * l * rho
                    - 4 * a * d * eta ** 2 * rho
                    - 4 * a * eta ** 2 * rho * g
                    - 2 * a * d * eta * l
                    + 2 * a * d * eta * rho
                    - 2 * a * eta * l * g
                    + 2 * a * eta * rho * gh
                    + 4 * H * a * eta * l * rho * sig
                    + a ** 2 * eta ** 2
                    + a ** 2 * l ** 2
                )
            )
            + (3 * H ** 2 * d * eta ** 2 * l * sig ** 2)
            + (H ** 2 * d * eta ** 2 * rho * sig ** 2)
            + (3 * H ** 2 * eta ** 2 * l * sig ** 2 * g)
            + (H ** 2 * eta ** 2 * rho * sig ** 2 * gh)
            - (H * d ** 2 * eta ** 2 * rho ** 2 * sig)
            + (3 * d ** 2 * eta ** 2 * l * rho ** 2 * g)
            + (3 * d * eta ** 2 * l * rho ** 2 * g ** 2)
            - (H ** 2 * d * eta * l * sig ** 2)
            - (H ** 2 * eta * l * sig ** 2 * gh)
            + (3 * H * d ** 2 * eta ** 2 * l * sig)
            + (H * d ** 2 * eta ** 2 * rho * sig)
            + (3 * H * eta ** 2 * l * sig * g ** 2)
            - (6 * d ** 2 * eta ** 2 * l * rho * g)
            - (6 * d * eta ** 2 * l * rho * g ** 2)
            - (2 * H * d ** 2 * eta * l * sig)
            + (3 * H * d ** 2 * eta * rho * sig)
            + (3 * H * eta * rho * sig * gh ** 2)
            + (d ** 2 * eta * l * rho * gh)
            - (4 * H * eta ** 2 * l * rho * sig * g ** 2)
            - (H * eta ** 2 * rho ** 2 * sig * gh * g)
            + (3 * H * d ** 2 * eta * l * rho * sig)
            + (6 * H * d * eta ** 2 * l * sig * g)
            + (H * d * eta ** 2 * rho * sig * gh)
            + (H * d * eta ** 2 * rho * sig * g)
            + (H * eta ** 2 * rho * sig * gh * g)
            - (2 * H * d * eta * l * sig * gh)
            - (2 * H * d * eta * l * sig * g)
            + (6 * H * d * eta * rho * sig * gh)
            - (2 * H * eta * l * sig * gh * g)
            + (2 * d * eta * l * rho * gh * g)
            - (H ** 2 * eta ** 2 * l * rho * sig ** 2 * a)
            + (2 * H ** 2 * eta * l ** 2 * rho * sig ** 2 * a)
            + (gh ** 3)
            - H
            * eta
            * l
            * rho
            * sig
            * g
            * sqrt(
                (
                    H ** 2 * eta ** 2 * sig ** 2
                    + d ** 2 * eta ** 2 * rho ** 2
                    + eta ** 2 * rho ** 2 * g ** 2
                    - 2 * d ** 2 * eta ** 2 * rho
                    - 2 * eta ** 2 * rho * g ** 2
                    + 2 * d ** 2 * eta * rho
                    + 2 * d * eta ** 2 * g
                    - 2 * d * eta * gh
                    - 2 * d * eta * g
                    - 2 * eta * gh * g
                    - 2 * H * d * eta ** 2 * rho * sig
                    - 2 * H * eta ** 2 * rho * sig * g
                    + 4 * H * d * eta * rho * sig
                    + 4 * H * eta * rho * sig * gh
                    + 2 * d * l * a
                    - 2 * H * a * eta ** 2 * rho * sig
                    - 2 * H * a * eta * l * sig
                    + 2 * a * d * eta * l * rho
                    + 2 * a * eta * l * rho * g
                    + d ** 2 * eta ** 2
                    + eta ** 2 * g ** 2
                    - 2 * d ** 2 * eta
                    + 2 * d * gh
                    + gh ** 2
                    + d ** 2
                    + 2 * d * eta * rho * gh
                    + 2 * d * eta * rho * g
                    + 2 * eta * rho * gh * g
                    + 2 * d * eta ** 2 * rho ** 2 * g
                    + 2 * H * d * eta ** 2 * sig
                    + 2 * H * eta ** 2 * sig * g
                    - 4 * d * eta ** 2 * rho * g
                    - 2 * H * d * eta * sig
                    - 2 * H * eta * sig * gh
                    + a ** 2 * eta ** 2 * rho ** 2
                    - 2 * a ** 2 * eta ** 2 * rho
                    - 2 * a ** 2 * eta * l
                    + 2 * a * d * eta ** 2
                    + 2 * a * eta ** 2 * g
                    - 2 * a * d * eta
                    - 2 * a * eta * gh
                    + 2 * a * l * gh
                    + 2 * a * d * eta ** 2 * rho ** 2
                    + 2 * a * eta ** 2 * rho ** 2 * g
                    + 2 * H * a * eta ** 2 * sig
                    + 2 * a ** 2 * eta * l * rho
                    - 4 * a * d * eta ** 2 * rho
                    - 4 * a * eta ** 2 * rho * g
                    - 2 * a * d * eta * l
                    + 2 * a * d * eta * rho
                    - 2 * a * eta * l * g
                    + 2 * a * eta * rho * gh
                    + 4 * H * a * eta * l * rho * sig
                    + a ** 2 * eta ** 2
                    + a ** 2 * l ** 2
                )
            )
            + (3 * H * d * eta * l ** 2 * rho * sig * a)
            - (7 * H * eta ** 2 * l * rho * sig * g * a)
            + (3 * H * eta * l ** 2 * rho * sig * g * a)
            - H
            * d
            * eta
            * l
            * rho
            * sig
            * sqrt(
                (
                    H ** 2 * eta ** 2 * sig ** 2
                    + d ** 2 * eta ** 2 * rho ** 2
                    + eta ** 2 * rho ** 2 * g ** 2
                    - 2 * d ** 2 * eta ** 2 * rho
                    - 2 * eta ** 2 * rho * g ** 2
                    + 2 * d ** 2 * eta * rho
                    + 2 * d * eta ** 2 * g
                    - 2 * d * eta * gh
                    - 2 * d * eta * g
                    - 2 * eta * gh * g
                    - 2 * H * d * eta ** 2 * rho * sig
                    - 2 * H * eta ** 2 * rho * sig * g
                    + 4 * H * d * eta * rho * sig
                    + 4 * H * eta * rho * sig * gh
                    + 2 * d * l * a
                    - 2 * H * a * eta ** 2 * rho * sig
                    - 2 * H * a * eta * l * sig
                    + 2 * a * d * eta * l * rho
                    + 2 * a * eta * l * rho * g
                    + d ** 2 * eta ** 2
                    + eta ** 2 * g ** 2
                    - 2 * d ** 2 * eta
                    + 2 * d * gh
                    + gh ** 2
                    + d ** 2
                    + 2 * d * eta * rho * gh
                    + 2 * d * eta * rho * g
                    + 2 * eta * rho * gh * g
                    + 2 * d * eta ** 2 * rho ** 2 * g
                    + 2 * H * d * eta ** 2 * sig
                    + 2 * H * eta ** 2 * sig * g
                    - 4 * d * eta ** 2 * rho * g
                    - 2 * H * d * eta * sig
                    - 2 * H * eta * sig * gh
                    + a ** 2 * eta ** 2 * rho ** 2
                    - 2 * a ** 2 * eta ** 2 * rho
                    - 2 * a ** 2 * eta * l
                    + 2 * a * d * eta ** 2
                    + 2 * a * eta ** 2 * g
                    - 2 * a * d * eta
                    - 2 * a * eta * gh
                    + 2 * a * l * gh
                    + 2 * a * d * eta ** 2 * rho ** 2
                    + 2 * a * eta ** 2 * rho ** 2 * g
                    + 2 * H * a * eta ** 2 * sig
                    + 2 * a ** 2 * eta * l * rho
                    - 4 * a * d * eta ** 2 * rho
                    - 4 * a * eta ** 2 * rho * g
                    - 2 * a * d * eta * l
                    + 2 * a * d * eta * rho
                    - 2 * a * eta * l * g
                    + 2 * a * eta * rho * gh
                    + 4 * H * a * eta * l * rho * sig
                    + a ** 2 * eta ** 2
                    + a ** 2 * l ** 2
                )
            )
            + (9 * H * d * eta * l * rho * sig * a)
            + (9 * H * eta * l * rho * sig * gh * a)
            - (H * d * eta ** 2 * rho ** 2 * sig * a)
            - (3 * H * eta ** 2 * l * rho * sig * a ** 2)
            - (H * eta ** 2 * rho ** 2 * sig * gh * a)
            + (6 * H * eta * l ** 2 * rho * sig * a ** 2)
            + (6 * d * eta ** 2 * l * rho ** 2 * g * a)
            + (6 * H * d * eta ** 2 * l * sig * a)
            + (H * d * eta ** 2 * rho * sig * a)
            - (2 * H * d * eta * l ** 2 * sig * a)
        )
    )
    return sing_strat


def sing_strat_alpha(beta, alpha, sigma, H, eta, lam, gamma, d, rho):
    alpha_sing_strat = (
        beta
        * (
            H ** 3 * eta ** 2 * lam * sigma ** 3
            + alpha ** 3 * eta ** 2 * lam * rho ** 2
            + d ** 3 * eta ** 2 * lam * rho ** 2
            + H ** 2 * d * eta ** 2 * sigma ** 2
            - 2 * alpha ** 3 * eta * lam ** 2 * rho
            - d ** 3 * eta * lam * rho
            - 3 * H * d ** 2 * eta * rho * sigma
            + 2 * H * d ** 2 * eta * sigma
            - alpha * d ** 2 * eta * rho
            + 2 * H ** 2 * alpha * eta ** 2 * lam * sigma ** 2
            + H ** 2 * alpha * eta * lam ** 2 * sigma ** 2
            + H ** 2 * d * eta ** 2 * lam * sigma ** 2
            - H ** 2 * d * eta ** 2 * rho * sigma ** 2
            + 5 * H * alpha * d * eta * lam * sigma
            + gamma ** 2
            * sqrt(
                H ** 2 * eta ** 2 * sigma ** 2
                + 2 * H * alpha * eta ** 2 * rho * sigma
                - 4 * H * alpha * eta * lam * rho * sigma
                + 2 * H * d * eta ** 2 * rho * sigma
                + 2 * H * eta ** 2 * gamma * rho * sigma
                + alpha ** 2 * eta ** 2 * rho ** 2
                + 2 * alpha * d * eta ** 2 * rho ** 2
                + 2 * alpha * eta ** 2 * gamma * rho ** 2
                + d ** 2 * eta ** 2 * rho ** 2
                + 2 * d * eta ** 2 * gamma * rho ** 2
                + eta ** 2 * gamma ** 2 * rho ** 2
                + 2 * H * alpha * eta * lam * sigma
                - 4 * H * d * eta * rho * sigma
                - 4 * H * eta * gamma * rho * sigma
                - 2 * alpha ** 2 * eta * lam * rho
                - 2 * alpha * d * eta * lam * rho
                - 2 * alpha * eta * gamma * lam * rho
                + 2 * H * d * eta * sigma
                + 2 * H * eta * gamma * sigma
                + alpha ** 2 * lam ** 2
                - 2 * alpha * d * eta * rho
                - 2 * alpha * eta * gamma * rho
                - 2 * d ** 2 * eta * rho
                - 4 * d * eta * gamma * rho
                - 2 * eta * gamma ** 2 * rho
                + 2 * alpha * d * lam
                + 2 * alpha * gamma * lam
                + d ** 2
                + 2 * d * gamma
                + gamma ** 2
            )
            + 3 * gamma ** 2 * d
            + 3 * gamma * d ** 2
            + sqrt(
                H ** 2 * eta ** 2 * sigma ** 2
                + 2 * H * alpha * eta ** 2 * rho * sigma
                - 4 * H * alpha * eta * lam * rho * sigma
                + 2 * H * d * eta ** 2 * rho * sigma
                + 2 * H * eta ** 2 * gamma * rho * sigma
                + alpha ** 2 * eta ** 2 * rho ** 2
                + 2 * alpha * d * eta ** 2 * rho ** 2
                + 2 * alpha * eta ** 2 * gamma * rho ** 2
                + d ** 2 * eta ** 2 * rho ** 2
                + 2 * d * eta ** 2 * gamma * rho ** 2
                + eta ** 2 * gamma ** 2 * rho ** 2
                + 2 * H * alpha * eta * lam * sigma
                - 4 * H * d * eta * rho * sigma
                - 4 * H * eta * gamma * rho * sigma
                - 2 * alpha ** 2 * eta * lam * rho
                - 2 * alpha * d * eta * lam * rho
                - 2 * alpha * eta * gamma * lam * rho
                + 2 * H * d * eta * sigma
                + 2 * H * eta * gamma * sigma
                + alpha ** 2 * lam ** 2
                - 2 * alpha * d * eta * rho
                - 2 * alpha * eta * gamma * rho
                - 2 * d ** 2 * eta * rho
                - 4 * d * eta * gamma * rho
                - 2 * eta * gamma ** 2 * rho
                + 2 * alpha * d * lam
                + 2 * alpha * gamma * lam
                + d ** 2
                + 2 * d * gamma
                + gamma ** 2
            )
            * d ** 2
            + d ** 3
            + gamma ** 3
            + alpha ** 3 * lam ** 3
            - d ** 3 * eta * rho
            + 3 * alpha ** 2 * d * lam ** 2
            + 3 * alpha * d ** 2 * lam
            - 3 * alpha ** 2 * d * eta * lam * rho
            - 4 * alpha * d ** 2 * eta * lam * rho
            - 9 * H * alpha * d * eta * lam * rho * sigma
            + H * gamma ** 2 * eta ** 2 * rho * sigma
            - gamma ** 2 * alpha * eta * lam ** 2 * rho
            - 3 * gamma * alpha ** 2 * eta * lam ** 2 * rho
            + H * gamma ** 2 * eta * lam * sigma
            - 3 * H * gamma ** 2 * eta * rho * sigma
            + gamma ** 2
            * sqrt(
                H ** 2 * eta ** 2 * sigma ** 2
                + 2 * H * alpha * eta ** 2 * rho * sigma
                - 4 * H * alpha * eta * lam * rho * sigma
                + 2 * H * d * eta ** 2 * rho * sigma
                + 2 * H * eta ** 2 * gamma * rho * sigma
                + alpha ** 2 * eta ** 2 * rho ** 2
                + 2 * alpha * d * eta ** 2 * rho ** 2
                + 2 * alpha * eta ** 2 * gamma * rho ** 2
                + d ** 2 * eta ** 2 * rho ** 2
                + 2 * d * eta ** 2 * gamma * rho ** 2
                + eta ** 2 * gamma ** 2 * rho ** 2
                + 2 * H * alpha * eta * lam * sigma
                - 4 * H * d * eta * rho * sigma
                - 4 * H * eta * gamma * rho * sigma
                - 2 * alpha ** 2 * eta * lam * rho
                - 2 * alpha * d * eta * lam * rho
                - 2 * alpha * eta * gamma * lam * rho
                + 2 * H * d * eta * sigma
                + 2 * H * eta * gamma * sigma
                + alpha ** 2 * lam ** 2
                - 2 * alpha * d * eta * rho
                - 2 * alpha * eta * gamma * rho
                - 2 * d ** 2 * eta * rho
                - 4 * d * eta * gamma * rho
                - 2 * eta * gamma ** 2 * rho
                + 2 * alpha * d * lam
                + 2 * alpha * gamma * lam
                + d ** 2
                + 2 * d * gamma
                + gamma ** 2
            )
            * eta
            * lam
            * rho
            - 4 * gamma ** 2 * alpha * eta * lam * rho
            - 3 * gamma ** 2 * d * eta * lam * rho
            - 3 * gamma * alpha ** 2 * eta * lam * rho
            - 3 * gamma * d ** 2 * eta * lam * rho
            + sqrt(
                H ** 2 * eta ** 2 * sigma ** 2
                + 2 * H * alpha * eta ** 2 * rho * sigma
                - 4 * H * alpha * eta * lam * rho * sigma
                + 2 * H * d * eta ** 2 * rho * sigma
                + 2 * H * eta ** 2 * gamma * rho * sigma
                + alpha ** 2 * eta ** 2 * rho ** 2
                + 2 * alpha * d * eta ** 2 * rho ** 2
                + 2 * alpha * eta ** 2 * gamma * rho ** 2
                + d ** 2 * eta ** 2 * rho ** 2
                + 2 * d * eta ** 2 * gamma * rho ** 2
                + eta ** 2 * gamma ** 2 * rho ** 2
                + 2 * H * alpha * eta * lam * sigma
                - 4 * H * d * eta * rho * sigma
                - 4 * H * eta * gamma * rho * sigma
                - 2 * alpha ** 2 * eta * lam * rho
                - 2 * alpha * d * eta * lam * rho
                - 2 * alpha * eta * gamma * lam * rho
                + 2 * H * d * eta * sigma
                + 2 * H * eta * gamma * sigma
                + alpha ** 2 * lam ** 2
                - 2 * alpha * d * eta * rho
                - 2 * alpha * eta * gamma * rho
                - 2 * d ** 2 * eta * rho
                - 4 * d * eta * gamma * rho
                - 2 * eta * gamma ** 2 * rho
                + 2 * alpha * d * lam
                + 2 * alpha * gamma * lam
                + d ** 2
                + 2 * d * gamma
                + gamma ** 2
            )
            * alpha ** 2
            * eta
            * lam
            * rho
            + sqrt(
                H ** 2 * eta ** 2 * sigma ** 2
                + 2 * H * alpha * eta ** 2 * rho * sigma
                - 4 * H * alpha * eta * lam * rho * sigma
                + 2 * H * d * eta ** 2 * rho * sigma
                + 2 * H * eta ** 2 * gamma * rho * sigma
                + alpha ** 2 * eta ** 2 * rho ** 2
                + 2 * alpha * d * eta ** 2 * rho ** 2
                + 2 * alpha * eta ** 2 * gamma * rho ** 2
                + d ** 2 * eta ** 2 * rho ** 2
                + 2 * d * eta ** 2 * gamma * rho ** 2
                + eta ** 2 * gamma ** 2 * rho ** 2
                + 2 * H * alpha * eta * lam * sigma
                - 4 * H * d * eta * rho * sigma
                - 4 * H * eta * gamma * rho * sigma
                - 2 * alpha ** 2 * eta * lam * rho
                - 2 * alpha * d * eta * lam * rho
                - 2 * alpha * eta * gamma * lam * rho
                + 2 * H * d * eta * sigma
                + 2 * H * eta * gamma * sigma
                + alpha ** 2 * lam ** 2
                - 2 * alpha * d * eta * rho
                - 2 * alpha * eta * gamma * rho
                - 2 * d ** 2 * eta * rho
                - 4 * d * eta * gamma * rho
                - 2 * eta * gamma ** 2 * rho
                + 2 * alpha * d * lam
                + 2 * alpha * gamma * lam
                + d ** 2
                + 2 * d * gamma
                + gamma ** 2
            )
            * d ** 2
            * eta
            * lam
            * rho
            + H
            * gamma
            * sqrt(
                H ** 2 * eta ** 2 * sigma ** 2
                + 2 * H * alpha * eta ** 2 * rho * sigma
                - 4 * H * alpha * eta * lam * rho * sigma
                + 2 * H * d * eta ** 2 * rho * sigma
                + 2 * H * eta ** 2 * gamma * rho * sigma
                + alpha ** 2 * eta ** 2 * rho ** 2
                + 2 * alpha * d * eta ** 2 * rho ** 2
                + 2 * alpha * eta ** 2 * gamma * rho ** 2
                + d ** 2 * eta ** 2 * rho ** 2
                + 2 * d * eta ** 2 * gamma * rho ** 2
                + eta ** 2 * gamma ** 2 * rho ** 2
                + 2 * H * alpha * eta * lam * sigma
                - 4 * H * d * eta * rho * sigma
                - 4 * H * eta * gamma * rho * sigma
                - 2 * alpha ** 2 * eta * lam * rho
                - 2 * alpha * d * eta * lam * rho
                - 2 * alpha * eta * gamma * lam * rho
                + 2 * H * d * eta * sigma
                + 2 * H * eta * gamma * sigma
                + alpha ** 2 * lam ** 2
                - 2 * alpha * d * eta * rho
                - 2 * alpha * eta * gamma * rho
                - 2 * d ** 2 * eta * rho
                - 4 * d * eta * gamma * rho
                - 2 * eta * gamma ** 2 * rho
                + 2 * alpha * d * lam
                + 2 * alpha * gamma * lam
                + d ** 2
                + 2 * d * gamma
                + gamma ** 2
            )
            * eta
            * sigma
            + 4 * H * gamma * d * eta * sigma
            + H
            * sqrt(
                H ** 2 * eta ** 2 * sigma ** 2
                + 2 * H * alpha * eta ** 2 * rho * sigma
                - 4 * H * alpha * eta * lam * rho * sigma
                + 2 * H * d * eta ** 2 * rho * sigma
                + 2 * H * eta ** 2 * gamma * rho * sigma
                + alpha ** 2 * eta ** 2 * rho ** 2
                + 2 * alpha * d * eta ** 2 * rho ** 2
                + 2 * alpha * eta ** 2 * gamma * rho ** 2
                + d ** 2 * eta ** 2 * rho ** 2
                + 2 * d * eta ** 2 * gamma * rho ** 2
                + eta ** 2 * gamma ** 2 * rho ** 2
                + 2 * H * alpha * eta * lam * sigma
                - 4 * H * d * eta * rho * sigma
                - 4 * H * eta * gamma * rho * sigma
                - 2 * alpha ** 2 * eta * lam * rho
                - 2 * alpha * d * eta * lam * rho
                - 2 * alpha * eta * gamma * lam * rho
                + 2 * H * d * eta * sigma
                + 2 * H * eta * gamma * sigma
                + alpha ** 2 * lam ** 2
                - 2 * alpha * d * eta * rho
                - 2 * alpha * eta * gamma * rho
                - 2 * d ** 2 * eta * rho
                - 4 * d * eta * gamma * rho
                - 2 * eta * gamma ** 2 * rho
                + 2 * alpha * d * lam
                + 2 * alpha * gamma * lam
                + d ** 2
                + 2 * d * gamma
                + gamma ** 2
            )
            * d
            * eta
            * sigma
            - 2 * gamma * alpha * d * eta * rho
            - H * d ** 2 * eta ** 2 * rho ** 2 * sigma
            + 3 * alpha ** 2 * d * eta ** 2 * lam * rho ** 2
            + 3 * alpha * d ** 2 * eta ** 2 * lam * rho ** 2
            + H ** 2 * d * eta * lam * sigma ** 2
            + 3 * H * alpha ** 2 * eta * lam ** 2 * sigma
            + H * d ** 2 * eta ** 2 * rho * sigma
            - 3 * alpha ** 2 * d * eta * lam ** 2 * rho
            - alpha * d ** 2 * eta * lam ** 2 * rho
            + H * d ** 2 * eta * lam * sigma
            + H ** 2 * gamma * eta ** 2 * lam * sigma ** 2
            - H ** 2 * gamma * eta ** 2 * rho * sigma ** 2
            - H * gamma ** 2 * eta ** 2 * rho ** 2 * sigma
            + 3 * gamma ** 2 * alpha * eta ** 2 * lam * rho ** 2
            + 3 * gamma ** 2 * d * eta ** 2 * lam * rho ** 2
            + 3 * gamma * alpha ** 2 * eta ** 2 * lam * rho ** 2
            + 3 * gamma * d ** 2 * eta ** 2 * lam * rho ** 2
            + H ** 2 * gamma * eta * lam * sigma ** 2
            + H ** 2
            * sqrt(
                H ** 2 * eta ** 2 * sigma ** 2
                + 2 * H * alpha * eta ** 2 * rho * sigma
                - 4 * H * alpha * eta * lam * rho * sigma
                + 2 * H * d * eta ** 2 * rho * sigma
                + 2 * H * eta ** 2 * gamma * rho * sigma
                + alpha ** 2 * eta ** 2 * rho ** 2
                + 2 * alpha * d * eta ** 2 * rho ** 2
                + 2 * alpha * eta ** 2 * gamma * rho ** 2
                + d ** 2 * eta ** 2 * rho ** 2
                + 2 * d * eta ** 2 * gamma * rho ** 2
                + eta ** 2 * gamma ** 2 * rho ** 2
                + 2 * H * alpha * eta * lam * sigma
                - 4 * H * d * eta * rho * sigma
                - 4 * H * eta * gamma * rho * sigma
                - 2 * alpha ** 2 * eta * lam * rho
                - 2 * alpha * d * eta * lam * rho
                - 2 * alpha * eta * gamma * lam * rho
                + 2 * H * d * eta * sigma
                + 2 * H * eta * gamma * sigma
                + alpha ** 2 * lam ** 2
                - 2 * alpha * d * eta * rho
                - 2 * alpha * eta * gamma * rho
                - 2 * d ** 2 * eta * rho
                - 4 * d * eta * gamma * rho
                - 2 * eta * gamma ** 2 * rho
                + 2 * alpha * d * lam
                + 2 * alpha * gamma * lam
                + d ** 2
                + 2 * d * gamma
                + gamma ** 2
            )
            * eta
            * lam
            * sigma ** 2
            + gamma ** 3 * eta ** 2 * lam * rho ** 2
            + H ** 2 * gamma * eta ** 2 * sigma ** 2
            - gamma ** 3 * eta * lam * rho
            + 2 * H * gamma ** 2 * eta * sigma
            - gamma ** 2 * alpha * eta * rho
            - 3 * gamma ** 2 * d * eta * rho
            - 3 * gamma * d ** 2 * eta * rho
            + 2
            * gamma
            * sqrt(
                H ** 2 * eta ** 2 * sigma ** 2
                + 2 * H * alpha * eta ** 2 * rho * sigma
                - 4 * H * alpha * eta * lam * rho * sigma
                + 2 * H * d * eta ** 2 * rho * sigma
                + 2 * H * eta ** 2 * gamma * rho * sigma
                + alpha ** 2 * eta ** 2 * rho ** 2
                + 2 * alpha * d * eta ** 2 * rho ** 2
                + 2 * alpha * eta ** 2 * gamma * rho ** 2
                + d ** 2 * eta ** 2 * rho ** 2
                + 2 * d * eta ** 2 * gamma * rho ** 2
                + eta ** 2 * gamma ** 2 * rho ** 2
                + 2 * H * alpha * eta * lam * sigma
                - 4 * H * d * eta * rho * sigma
                - 4 * H * eta * gamma * rho * sigma
                - 2 * alpha ** 2 * eta * lam * rho
                - 2 * alpha * d * eta * lam * rho
                - 2 * alpha * eta * gamma * lam * rho
                + 2 * H * d * eta * sigma
                + 2 * H * eta * gamma * sigma
                + alpha ** 2 * lam ** 2
                - 2 * alpha * d * eta * rho
                - 2 * alpha * eta * gamma * rho
                - 2 * d ** 2 * eta * rho
                - 4 * d * eta * gamma * rho
                - 2 * eta * gamma ** 2 * rho
                + 2 * alpha * d * lam
                + 2 * alpha * gamma * lam
                + d ** 2
                + 2 * d * gamma
                + gamma ** 2
            )
            * alpha
            * lam
            + 6 * gamma * alpha * d * lam
            + 2
            * sqrt(
                H ** 2 * eta ** 2 * sigma ** 2
                + 2 * H * alpha * eta ** 2 * rho * sigma
                - 4 * H * alpha * eta * lam * rho * sigma
                + 2 * H * d * eta ** 2 * rho * sigma
                + 2 * H * eta ** 2 * gamma * rho * sigma
                + alpha ** 2 * eta ** 2 * rho ** 2
                + 2 * alpha * d * eta ** 2 * rho ** 2
                + 2 * alpha * eta ** 2 * gamma * rho ** 2
                + d ** 2 * eta ** 2 * rho ** 2
                + 2 * d * eta ** 2 * gamma * rho ** 2
                + eta ** 2 * gamma ** 2 * rho ** 2
                + 2 * H * alpha * eta * lam * sigma
                - 4 * H * d * eta * rho * sigma
                - 4 * H * eta * gamma * rho * sigma
                - 2 * alpha ** 2 * eta * lam * rho
                - 2 * alpha * d * eta * lam * rho
                - 2 * alpha * eta * gamma * lam * rho
                + 2 * H * d * eta * sigma
                + 2 * H * eta * gamma * sigma
                + alpha ** 2 * lam ** 2
                - 2 * alpha * d * eta * rho
                - 2 * alpha * eta * gamma * rho
                - 2 * d ** 2 * eta * rho
                - 4 * d * eta * gamma * rho
                - 2 * eta * gamma ** 2 * rho
                + 2 * alpha * d * lam
                + 2 * alpha * gamma * lam
                + d ** 2
                + 2 * d * gamma
                + gamma ** 2
            )
            * alpha
            * d
            * lam
            + H ** 2 * alpha * eta ** 2 * lam * rho * sigma ** 2
            - 2 * H ** 2 * alpha * eta * lam ** 2 * rho * sigma ** 2
            + 2 * H ** 2 * d * eta ** 2 * lam * rho * sigma ** 2
            + H * d ** 2 * eta ** 2 * lam * rho ** 2 * sigma
            - 2 * H ** 2 * d * eta * lam * rho * sigma ** 2
            + 3 * H * alpha ** 2 * eta ** 2 * lam * rho * sigma
            - 6 * H * alpha ** 2 * eta * lam ** 2 * rho * sigma
            - H * alpha * d * eta ** 2 * rho ** 2 * sigma
            + 2 * H * d ** 2 * eta ** 2 * lam * rho * sigma
            + H * alpha * d * eta ** 2 * rho * sigma
            + H * alpha * d * eta * lam ** 2 * sigma
            - 3 * H * d ** 2 * eta * lam * rho * sigma
            + H * gamma * alpha * eta ** 2 * lam * rho ** 2 * sigma
            + 2 * H * gamma * d * eta ** 2 * lam * rho ** 2 * sigma
            + 5 * H * gamma * alpha * eta ** 2 * lam * rho * sigma
            - 3 * H * gamma * alpha * eta * lam ** 2 * rho * sigma
            + 4 * H * gamma * d * eta ** 2 * lam * rho * sigma
            + H
            * gamma
            * sqrt(
                H ** 2 * eta ** 2 * sigma ** 2
                + 2 * H * alpha * eta ** 2 * rho * sigma
                - 4 * H * alpha * eta * lam * rho * sigma
                + 2 * H * d * eta ** 2 * rho * sigma
                + 2 * H * eta ** 2 * gamma * rho * sigma
                + alpha ** 2 * eta ** 2 * rho ** 2
                + 2 * alpha * d * eta ** 2 * rho ** 2
                + 2 * alpha * eta ** 2 * gamma * rho ** 2
                + d ** 2 * eta ** 2 * rho ** 2
                + 2 * d * eta ** 2 * gamma * rho ** 2
                + eta ** 2 * gamma ** 2 * rho ** 2
                + 2 * H * alpha * eta * lam * sigma
                - 4 * H * d * eta * rho * sigma
                - 4 * H * eta * gamma * rho * sigma
                - 2 * alpha ** 2 * eta * lam * rho
                - 2 * alpha * d * eta * lam * rho
                - 2 * alpha * eta * gamma * lam * rho
                + 2 * H * d * eta * sigma
                + 2 * H * eta * gamma * sigma
                + alpha ** 2 * lam ** 2
                - 2 * alpha * d * eta * rho
                - 2 * alpha * eta * gamma * rho
                - 2 * d ** 2 * eta * rho
                - 4 * d * eta * gamma * rho
                - 2 * eta * gamma ** 2 * rho
                + 2 * alpha * d * lam
                + 2 * alpha * gamma * lam
                + d ** 2
                + 2 * d * gamma
                + gamma ** 2
            )
            * eta
            * lam
            * rho
            * sigma
            - 9 * H * gamma * alpha * eta * lam * rho * sigma
            - 6 * H * gamma * d * eta * lam * rho * sigma
            + H
            * sqrt(
                H ** 2 * eta ** 2 * sigma ** 2
                + 2 * H * alpha * eta ** 2 * rho * sigma
                - 4 * H * alpha * eta * lam * rho * sigma
                + 2 * H * d * eta ** 2 * rho * sigma
                + 2 * H * eta ** 2 * gamma * rho * sigma
                + alpha ** 2 * eta ** 2 * rho ** 2
                + 2 * alpha * d * eta ** 2 * rho ** 2
                + 2 * alpha * eta ** 2 * gamma * rho ** 2
                + d ** 2 * eta ** 2 * rho ** 2
                + 2 * d * eta ** 2 * gamma * rho ** 2
                + eta ** 2 * gamma ** 2 * rho ** 2
                + 2 * H * alpha * eta * lam * sigma
                - 4 * H * d * eta * rho * sigma
                - 4 * H * eta * gamma * rho * sigma
                - 2 * alpha ** 2 * eta * lam * rho
                - 2 * alpha * d * eta * lam * rho
                - 2 * alpha * eta * gamma * lam * rho
                + 2 * H * d * eta * sigma
                + 2 * H * eta * gamma * sigma
                + alpha ** 2 * lam ** 2
                - 2 * alpha * d * eta * rho
                - 2 * alpha * eta * gamma * rho
                - 2 * d ** 2 * eta * rho
                - 4 * d * eta * gamma * rho
                - 2 * eta * gamma ** 2 * rho
                + 2 * alpha * d * lam
                + 2 * alpha * gamma * lam
                + d ** 2
                + 2 * d * gamma
                + gamma ** 2
            )
            * d
            * eta
            * lam
            * rho
            * sigma
            + 2 * H ** 2 * gamma * eta ** 2 * lam * rho * sigma ** 2
            + H * gamma ** 2 * eta ** 2 * lam * rho ** 2 * sigma
            - 2 * H ** 2 * gamma * eta * lam * rho * sigma ** 2
            + 2 * H * gamma ** 2 * eta ** 2 * lam * rho * sigma
            - H * gamma * alpha * eta ** 2 * rho ** 2 * sigma
            - 2 * H * gamma * d * eta ** 2 * rho ** 2 * sigma
            + 6 * gamma * alpha * d * eta ** 2 * lam * rho ** 2
            - 3 * H * gamma ** 2 * eta * lam * rho * sigma
            + H * gamma * alpha * eta ** 2 * rho * sigma
            + H * gamma * alpha * eta * lam ** 2 * sigma
            + 2 * H * gamma * d * eta ** 2 * rho * sigma
            - 2 * gamma * alpha * d * eta * lam ** 2 * rho
            + H
            * gamma
            * sqrt(
                H ** 2 * eta ** 2 * sigma ** 2
                + 2 * H * alpha * eta ** 2 * rho * sigma
                - 4 * H * alpha * eta * lam * rho * sigma
                + 2 * H * d * eta ** 2 * rho * sigma
                + 2 * H * eta ** 2 * gamma * rho * sigma
                + alpha ** 2 * eta ** 2 * rho ** 2
                + 2 * alpha * d * eta ** 2 * rho ** 2
                + 2 * alpha * eta ** 2 * gamma * rho ** 2
                + d ** 2 * eta ** 2 * rho ** 2
                + 2 * d * eta ** 2 * gamma * rho ** 2
                + eta ** 2 * gamma ** 2 * rho ** 2
                + 2 * H * alpha * eta * lam * sigma
                - 4 * H * d * eta * rho * sigma
                - 4 * H * eta * gamma * rho * sigma
                - 2 * alpha ** 2 * eta * lam * rho
                - 2 * alpha * d * eta * lam * rho
                - 2 * alpha * eta * gamma * lam * rho
                + 2 * H * d * eta * sigma
                + 2 * H * eta * gamma * sigma
                + alpha ** 2 * lam ** 2
                - 2 * alpha * d * eta * rho
                - 2 * alpha * eta * gamma * rho
                - 2 * d ** 2 * eta * rho
                - 4 * d * eta * gamma * rho
                - 2 * eta * gamma ** 2 * rho
                + 2 * alpha * d * lam
                + 2 * alpha * gamma * lam
                + d ** 2
                + 2 * d * gamma
                + gamma ** 2
            )
            * eta
            * lam
            * sigma
            - H
            * gamma
            * sqrt(
                H ** 2 * eta ** 2 * sigma ** 2
                + 2 * H * alpha * eta ** 2 * rho * sigma
                - 4 * H * alpha * eta * lam * rho * sigma
                + 2 * H * d * eta ** 2 * rho * sigma
                + 2 * H * eta ** 2 * gamma * rho * sigma
                + alpha ** 2 * eta ** 2 * rho ** 2
                + 2 * alpha * d * eta ** 2 * rho ** 2
                + 2 * alpha * eta ** 2 * gamma * rho ** 2
                + d ** 2 * eta ** 2 * rho ** 2
                + 2 * d * eta ** 2 * gamma * rho ** 2
                + eta ** 2 * gamma ** 2 * rho ** 2
                + 2 * H * alpha * eta * lam * sigma
                - 4 * H * d * eta * rho * sigma
                - 4 * H * eta * gamma * rho * sigma
                - 2 * alpha ** 2 * eta * lam * rho
                - 2 * alpha * d * eta * lam * rho
                - 2 * alpha * eta * gamma * lam * rho
                + 2 * H * d * eta * sigma
                + 2 * H * eta * gamma * sigma
                + alpha ** 2 * lam ** 2
                - 2 * alpha * d * eta * rho
                - 2 * alpha * eta * gamma * rho
                - 2 * d ** 2 * eta * rho
                - 4 * d * eta * gamma * rho
                - 2 * eta * gamma ** 2 * rho
                + 2 * alpha * d * lam
                + 2 * alpha * gamma * lam
                + d ** 2
                + 2 * d * gamma
                + gamma ** 2
            )
            * eta
            * rho
            * sigma
            + 5 * H * gamma * alpha * eta * lam * sigma
            + 2 * H * gamma * d * eta * lam * sigma
            - 6 * H * gamma * d * eta * rho * sigma
            + 2
            * H
            * sqrt(
                H ** 2 * eta ** 2 * sigma ** 2
                + 2 * H * alpha * eta ** 2 * rho * sigma
                - 4 * H * alpha * eta * lam * rho * sigma
                + 2 * H * d * eta ** 2 * rho * sigma
                + 2 * H * eta ** 2 * gamma * rho * sigma
                + alpha ** 2 * eta ** 2 * rho ** 2
                + 2 * alpha * d * eta ** 2 * rho ** 2
                + 2 * alpha * eta ** 2 * gamma * rho ** 2
                + d ** 2 * eta ** 2 * rho ** 2
                + 2 * d * eta ** 2 * gamma * rho ** 2
                + eta ** 2 * gamma ** 2 * rho ** 2
                + 2 * H * alpha * eta * lam * sigma
                - 4 * H * d * eta * rho * sigma
                - 4 * H * eta * gamma * rho * sigma
                - 2 * alpha ** 2 * eta * lam * rho
                - 2 * alpha * d * eta * lam * rho
                - 2 * alpha * eta * gamma * lam * rho
                + 2 * H * d * eta * sigma
                + 2 * H * eta * gamma * sigma
                + alpha ** 2 * lam ** 2
                - 2 * alpha * d * eta * rho
                - 2 * alpha * eta * gamma * rho
                - 2 * d ** 2 * eta * rho
                - 4 * d * eta * gamma * rho
                - 2 * eta * gamma ** 2 * rho
                + 2 * alpha * d * lam
                + 2 * alpha * gamma * lam
                + d ** 2
                + 2 * d * gamma
                + gamma ** 2
            )
            * alpha
            * eta
            * lam
            * sigma
            + H
            * sqrt(
                H ** 2 * eta ** 2 * sigma ** 2
                + 2 * H * alpha * eta ** 2 * rho * sigma
                - 4 * H * alpha * eta * lam * rho * sigma
                + 2 * H * d * eta ** 2 * rho * sigma
                + 2 * H * eta ** 2 * gamma * rho * sigma
                + alpha ** 2 * eta ** 2 * rho ** 2
                + 2 * alpha * d * eta ** 2 * rho ** 2
                + 2 * alpha * eta ** 2 * gamma * rho ** 2
                + d ** 2 * eta ** 2 * rho ** 2
                + 2 * d * eta ** 2 * gamma * rho ** 2
                + eta ** 2 * gamma ** 2 * rho ** 2
                + 2 * H * alpha * eta * lam * sigma
                - 4 * H * d * eta * rho * sigma
                - 4 * H * eta * gamma * rho * sigma
                - 2 * alpha ** 2 * eta * lam * rho
                - 2 * alpha * d * eta * lam * rho
                - 2 * alpha * eta * gamma * lam * rho
                + 2 * H * d * eta * sigma
                + 2 * H * eta * gamma * sigma
                + alpha ** 2 * lam ** 2
                - 2 * alpha * d * eta * rho
                - 2 * alpha * eta * gamma * rho
                - 2 * d ** 2 * eta * rho
                - 4 * d * eta * gamma * rho
                - 2 * eta * gamma ** 2 * rho
                + 2 * alpha * d * lam
                + 2 * alpha * gamma * lam
                + d ** 2
                + 2 * d * gamma
                + gamma ** 2
            )
            * d
            * eta
            * lam
            * sigma
            - H
            * sqrt(
                H ** 2 * eta ** 2 * sigma ** 2
                + 2 * H * alpha * eta ** 2 * rho * sigma
                - 4 * H * alpha * eta * lam * rho * sigma
                + 2 * H * d * eta ** 2 * rho * sigma
                + 2 * H * eta ** 2 * gamma * rho * sigma
                + alpha ** 2 * eta ** 2 * rho ** 2
                + 2 * alpha * d * eta ** 2 * rho ** 2
                + 2 * alpha * eta ** 2 * gamma * rho ** 2
                + d ** 2 * eta ** 2 * rho ** 2
                + 2 * d * eta ** 2 * gamma * rho ** 2
                + eta ** 2 * gamma ** 2 * rho ** 2
                + 2 * H * alpha * eta * lam * sigma
                - 4 * H * d * eta * rho * sigma
                - 4 * H * eta * gamma * rho * sigma
                - 2 * alpha ** 2 * eta * lam * rho
                - 2 * alpha * d * eta * lam * rho
                - 2 * alpha * eta * gamma * lam * rho
                + 2 * H * d * eta * sigma
                + 2 * H * eta * gamma * sigma
                + alpha ** 2 * lam ** 2
                - 2 * alpha * d * eta * rho
                - 2 * alpha * eta * gamma * rho
                - 2 * d ** 2 * eta * rho
                - 4 * d * eta * gamma * rho
                - 2 * eta * gamma ** 2 * rho
                + 2 * alpha * d * lam
                + 2 * alpha * gamma * lam
                + d ** 2
                + 2 * d * gamma
                + gamma ** 2
            )
            * d
            * eta
            * rho
            * sigma
            + 2
            * gamma
            * sqrt(
                H ** 2 * eta ** 2 * sigma ** 2
                + 2 * H * alpha * eta ** 2 * rho * sigma
                - 4 * H * alpha * eta * lam * rho * sigma
                + 2 * H * d * eta ** 2 * rho * sigma
                + 2 * H * eta ** 2 * gamma * rho * sigma
                + alpha ** 2 * eta ** 2 * rho ** 2
                + 2 * alpha * d * eta ** 2 * rho ** 2
                + 2 * alpha * eta ** 2 * gamma * rho ** 2
                + d ** 2 * eta ** 2 * rho ** 2
                + 2 * d * eta ** 2 * gamma * rho ** 2
                + eta ** 2 * gamma ** 2 * rho ** 2
                + 2 * H * alpha * eta * lam * sigma
                - 4 * H * d * eta * rho * sigma
                - 4 * H * eta * gamma * rho * sigma
                - 2 * alpha ** 2 * eta * lam * rho
                - 2 * alpha * d * eta * lam * rho
                - 2 * alpha * eta * gamma * lam * rho
                + 2 * H * d * eta * sigma
                + 2 * H * eta * gamma * sigma
                + alpha ** 2 * lam ** 2
                - 2 * alpha * d * eta * rho
                - 2 * alpha * eta * gamma * rho
                - 2 * d ** 2 * eta * rho
                - 4 * d * eta * gamma * rho
                - 2 * eta * gamma ** 2 * rho
                + 2 * alpha * d * lam
                + 2 * alpha * gamma * lam
                + d ** 2
                + 2 * d * gamma
                + gamma ** 2
            )
            * alpha
            * eta
            * lam
            * rho
            + 2
            * gamma
            * sqrt(
                H ** 2 * eta ** 2 * sigma ** 2
                + 2 * H * alpha * eta ** 2 * rho * sigma
                - 4 * H * alpha * eta * lam * rho * sigma
                + 2 * H * d * eta ** 2 * rho * sigma
                + 2 * H * eta ** 2 * gamma * rho * sigma
                + alpha ** 2 * eta ** 2 * rho ** 2
                + 2 * alpha * d * eta ** 2 * rho ** 2
                + 2 * alpha * eta ** 2 * gamma * rho ** 2
                + d ** 2 * eta ** 2 * rho ** 2
                + 2 * d * eta ** 2 * gamma * rho ** 2
                + eta ** 2 * gamma ** 2 * rho ** 2
                + 2 * H * alpha * eta * lam * sigma
                - 4 * H * d * eta * rho * sigma
                - 4 * H * eta * gamma * rho * sigma
                - 2 * alpha ** 2 * eta * lam * rho
                - 2 * alpha * d * eta * lam * rho
                - 2 * alpha * eta * gamma * lam * rho
                + 2 * H * d * eta * sigma
                + 2 * H * eta * gamma * sigma
                + alpha ** 2 * lam ** 2
                - 2 * alpha * d * eta * rho
                - 2 * alpha * eta * gamma * rho
                - 2 * d ** 2 * eta * rho
                - 4 * d * eta * gamma * rho
                - 2 * eta * gamma ** 2 * rho
                + 2 * alpha * d * lam
                + 2 * alpha * gamma * lam
                + d ** 2
                + 2 * d * gamma
                + gamma ** 2
            )
            * d
            * eta
            * lam
            * rho
            - 8 * gamma * alpha * d * eta * lam * rho
            + 2
            * sqrt(
                H ** 2 * eta ** 2 * sigma ** 2
                + 2 * H * alpha * eta ** 2 * rho * sigma
                - 4 * H * alpha * eta * lam * rho * sigma
                + 2 * H * d * eta ** 2 * rho * sigma
                + 2 * H * eta ** 2 * gamma * rho * sigma
                + alpha ** 2 * eta ** 2 * rho ** 2
                + 2 * alpha * d * eta ** 2 * rho ** 2
                + 2 * alpha * eta ** 2 * gamma * rho ** 2
                + d ** 2 * eta ** 2 * rho ** 2
                + 2 * d * eta ** 2 * gamma * rho ** 2
                + eta ** 2 * gamma ** 2 * rho ** 2
                + 2 * H * alpha * eta * lam * sigma
                - 4 * H * d * eta * rho * sigma
                - 4 * H * eta * gamma * rho * sigma
                - 2 * alpha ** 2 * eta * lam * rho
                - 2 * alpha * d * eta * lam * rho
                - 2 * alpha * eta * gamma * lam * rho
                + 2 * H * d * eta * sigma
                + 2 * H * eta * gamma * sigma
                + alpha ** 2 * lam ** 2
                - 2 * alpha * d * eta * rho
                - 2 * alpha * eta * gamma * rho
                - 2 * d ** 2 * eta * rho
                - 4 * d * eta * gamma * rho
                - 2 * eta * gamma ** 2 * rho
                + 2 * alpha * d * lam
                + 2 * alpha * gamma * lam
                + d ** 2
                + 2 * d * gamma
                + gamma ** 2
            )
            * alpha
            * d
            * eta
            * lam
            * rho
            - gamma ** 3 * eta * rho
            + 3 * gamma * alpha ** 2 * lam ** 2
            + sqrt(
                H ** 2 * eta ** 2 * sigma ** 2
                + 2 * H * alpha * eta ** 2 * rho * sigma
                - 4 * H * alpha * eta * lam * rho * sigma
                + 2 * H * d * eta ** 2 * rho * sigma
                + 2 * H * eta ** 2 * gamma * rho * sigma
                + alpha ** 2 * eta ** 2 * rho ** 2
                + 2 * alpha * d * eta ** 2 * rho ** 2
                + 2 * alpha * eta ** 2 * gamma * rho ** 2
                + d ** 2 * eta ** 2 * rho ** 2
                + 2 * d * eta ** 2 * gamma * rho ** 2
                + eta ** 2 * gamma ** 2 * rho ** 2
                + 2 * H * alpha * eta * lam * sigma
                - 4 * H * d * eta * rho * sigma
                - 4 * H * eta * gamma * rho * sigma
                - 2 * alpha ** 2 * eta * lam * rho
                - 2 * alpha * d * eta * lam * rho
                - 2 * alpha * eta * gamma * lam * rho
                + 2 * H * d * eta * sigma
                + 2 * H * eta * gamma * sigma
                + alpha ** 2 * lam ** 2
                - 2 * alpha * d * eta * rho
                - 2 * alpha * eta * gamma * rho
                - 2 * d ** 2 * eta * rho
                - 4 * d * eta * gamma * rho
                - 2 * eta * gamma ** 2 * rho
                + 2 * alpha * d * lam
                + 2 * alpha * gamma * lam
                + d ** 2
                + 2 * d * gamma
                + gamma ** 2
            )
            * alpha ** 2
            * lam ** 2
            + 3 * gamma ** 2 * alpha * lam
            + 2
            * gamma
            * sqrt(
                H ** 2 * eta ** 2 * sigma ** 2
                + 2 * H * alpha * eta ** 2 * rho * sigma
                - 4 * H * alpha * eta * lam * rho * sigma
                + 2 * H * d * eta ** 2 * rho * sigma
                + 2 * H * eta ** 2 * gamma * rho * sigma
                + alpha ** 2 * eta ** 2 * rho ** 2
                + 2 * alpha * d * eta ** 2 * rho ** 2
                + 2 * alpha * eta ** 2 * gamma * rho ** 2
                + d ** 2 * eta ** 2 * rho ** 2
                + 2 * d * eta ** 2 * gamma * rho ** 2
                + eta ** 2 * gamma ** 2 * rho ** 2
                + 2 * H * alpha * eta * lam * sigma
                - 4 * H * d * eta * rho * sigma
                - 4 * H * eta * gamma * rho * sigma
                - 2 * alpha ** 2 * eta * lam * rho
                - 2 * alpha * d * eta * lam * rho
                - 2 * alpha * eta * gamma * lam * rho
                + 2 * H * d * eta * sigma
                + 2 * H * eta * gamma * sigma
                + alpha ** 2 * lam ** 2
                - 2 * alpha * d * eta * rho
                - 2 * alpha * eta * gamma * rho
                - 2 * d ** 2 * eta * rho
                - 4 * d * eta * gamma * rho
                - 2 * eta * gamma ** 2 * rho
                + 2 * alpha * d * lam
                + 2 * alpha * gamma * lam
                + d ** 2
                + 2 * d * gamma
                + gamma ** 2
            )
            * d
            + H * alpha * d * eta ** 2 * lam * rho ** 2 * sigma
            + 5 * H * alpha * d * eta ** 2 * lam * rho * sigma
            - 3 * H * alpha * d * eta * lam ** 2 * rho * sigma
        )
        / (lam * alpha + d + gamma)
        / (sigma * H + alpha + d + gamma)
        / (
            gamma ** 2
            + 2 * H * alpha * eta ** 2 * rho * sigma
            + 2 * H * alpha * eta * lam * sigma
            - 2 * alpha * d * eta * lam * rho
            - 2 * alpha * eta * gamma * lam * rho
            + d ** 2
            + alpha ** 2 * eta ** 2 * rho ** 2
            + 2 * H * d * eta * sigma
            - 2 * alpha ** 2 * eta * lam * rho
            + 2 * H * d * eta ** 2 * rho * sigma
            - 4 * H * d * eta * rho * sigma
            + 2 * H * eta ** 2 * gamma * rho * sigma
            - 4 * H * eta * gamma * rho * sigma
            + sqrt(
                H ** 2 * eta ** 2 * sigma ** 2
                + 2 * H * alpha * eta ** 2 * rho * sigma
                - 4 * H * alpha * eta * lam * rho * sigma
                + 2 * H * d * eta ** 2 * rho * sigma
                + 2 * H * eta ** 2 * gamma * rho * sigma
                + alpha ** 2 * eta ** 2 * rho ** 2
                + 2 * alpha * d * eta ** 2 * rho ** 2
                + 2 * alpha * eta ** 2 * gamma * rho ** 2
                + d ** 2 * eta ** 2 * rho ** 2
                + 2 * d * eta ** 2 * gamma * rho ** 2
                + eta ** 2 * gamma ** 2 * rho ** 2
                + 2 * H * alpha * eta * lam * sigma
                - 4 * H * d * eta * rho * sigma
                - 4 * H * eta * gamma * rho * sigma
                - 2 * alpha ** 2 * eta * lam * rho
                - 2 * alpha * d * eta * lam * rho
                - 2 * alpha * eta * gamma * lam * rho
                + 2 * H * d * eta * sigma
                + 2 * H * eta * gamma * sigma
                + alpha ** 2 * lam ** 2
                - 2 * alpha * d * eta * rho
                - 2 * alpha * eta * gamma * rho
                - 2 * d ** 2 * eta * rho
                - 4 * d * eta * gamma * rho
                - 2 * eta * gamma ** 2 * rho
                + 2 * alpha * d * lam
                + 2 * alpha * gamma * lam
                + d ** 2
                + 2 * d * gamma
                + gamma ** 2
            )
            * alpha
            * lam
            + 2 * d * eta ** 2 * gamma * rho ** 2
            + 2 * H * eta * gamma * sigma
            - 4 * d * eta * gamma * rho
            + 2 * alpha * d * lam
            + 2 * alpha * gamma * lam
            - 4 * H * alpha * eta * lam * rho * sigma
            + gamma
            * sqrt(
                H ** 2 * eta ** 2 * sigma ** 2
                + 2 * H * alpha * eta ** 2 * rho * sigma
                - 4 * H * alpha * eta * lam * rho * sigma
                + 2 * H * d * eta ** 2 * rho * sigma
                + 2 * H * eta ** 2 * gamma * rho * sigma
                + alpha ** 2 * eta ** 2 * rho ** 2
                + 2 * alpha * d * eta ** 2 * rho ** 2
                + 2 * alpha * eta ** 2 * gamma * rho ** 2
                + d ** 2 * eta ** 2 * rho ** 2
                + 2 * d * eta ** 2 * gamma * rho ** 2
                + eta ** 2 * gamma ** 2 * rho ** 2
                + 2 * H * alpha * eta * lam * sigma
                - 4 * H * d * eta * rho * sigma
                - 4 * H * eta * gamma * rho * sigma
                - 2 * alpha ** 2 * eta * lam * rho
                - 2 * alpha * d * eta * lam * rho
                - 2 * alpha * eta * gamma * lam * rho
                + 2 * H * d * eta * sigma
                + 2 * H * eta * gamma * sigma
                + alpha ** 2 * lam ** 2
                - 2 * alpha * d * eta * rho
                - 2 * alpha * eta * gamma * rho
                - 2 * d ** 2 * eta * rho
                - 4 * d * eta * gamma * rho
                - 2 * eta * gamma ** 2 * rho
                + 2 * alpha * d * lam
                + 2 * alpha * gamma * lam
                + d ** 2
                + 2 * d * gamma
                + gamma ** 2
            )
            + sqrt(
                H ** 2 * eta ** 2 * sigma ** 2
                + 2 * H * alpha * eta ** 2 * rho * sigma
                - 4 * H * alpha * eta * lam * rho * sigma
                + 2 * H * d * eta ** 2 * rho * sigma
                + 2 * H * eta ** 2 * gamma * rho * sigma
                + alpha ** 2 * eta ** 2 * rho ** 2
                + 2 * alpha * d * eta ** 2 * rho ** 2
                + 2 * alpha * eta ** 2 * gamma * rho ** 2
                + d ** 2 * eta ** 2 * rho ** 2
                + 2 * d * eta ** 2 * gamma * rho ** 2
                + eta ** 2 * gamma ** 2 * rho ** 2
                + 2 * H * alpha * eta * lam * sigma
                - 4 * H * d * eta * rho * sigma
                - 4 * H * eta * gamma * rho * sigma
                - 2 * alpha ** 2 * eta * lam * rho
                - 2 * alpha * d * eta * lam * rho
                - 2 * alpha * eta * gamma * lam * rho
                + 2 * H * d * eta * sigma
                + 2 * H * eta * gamma * sigma
                + alpha ** 2 * lam ** 2
                - 2 * alpha * d * eta * rho
                - 2 * alpha * eta * gamma * rho
                - 2 * d ** 2 * eta * rho
                - 4 * d * eta * gamma * rho
                - 2 * eta * gamma ** 2 * rho
                + 2 * alpha * d * lam
                + 2 * alpha * gamma * lam
                + d ** 2
                + 2 * d * gamma
                + gamma ** 2
            )
            * d
            + alpha ** 2 * lam ** 2
            + 2 * alpha * d * eta ** 2 * rho ** 2
            + 2 * alpha * eta ** 2 * gamma * rho ** 2
            - 2 * alpha * d * eta * rho
            - 2 * alpha * eta * gamma * rho
            + 2 * d * gamma
            + H
            * sqrt(
                H ** 2 * eta ** 2 * sigma ** 2
                + 2 * H * alpha * eta ** 2 * rho * sigma
                - 4 * H * alpha * eta * lam * rho * sigma
                + 2 * H * d * eta ** 2 * rho * sigma
                + 2 * H * eta ** 2 * gamma * rho * sigma
                + alpha ** 2 * eta ** 2 * rho ** 2
                + 2 * alpha * d * eta ** 2 * rho ** 2
                + 2 * alpha * eta ** 2 * gamma * rho ** 2
                + d ** 2 * eta ** 2 * rho ** 2
                + 2 * d * eta ** 2 * gamma * rho ** 2
                + eta ** 2 * gamma ** 2 * rho ** 2
                + 2 * H * alpha * eta * lam * sigma
                - 4 * H * d * eta * rho * sigma
                - 4 * H * eta * gamma * rho * sigma
                - 2 * alpha ** 2 * eta * lam * rho
                - 2 * alpha * d * eta * lam * rho
                - 2 * alpha * eta * gamma * lam * rho
                + 2 * H * d * eta * sigma
                + 2 * H * eta * gamma * sigma
                + alpha ** 2 * lam ** 2
                - 2 * alpha * d * eta * rho
                - 2 * alpha * eta * gamma * rho
                - 2 * d ** 2 * eta * rho
                - 4 * d * eta * gamma * rho
                - 2 * eta * gamma ** 2 * rho
                + 2 * alpha * d * lam
                + 2 * alpha * gamma * lam
                + d ** 2
                + 2 * d * gamma
                + gamma ** 2
            )
            * eta
            * sigma
            + gamma
            * sqrt(
                H ** 2 * eta ** 2 * sigma ** 2
                + 2 * H * alpha * eta ** 2 * rho * sigma
                - 4 * H * alpha * eta * lam * rho * sigma
                + 2 * H * d * eta ** 2 * rho * sigma
                + 2 * H * eta ** 2 * gamma * rho * sigma
                + alpha ** 2 * eta ** 2 * rho ** 2
                + 2 * alpha * d * eta ** 2 * rho ** 2
                + 2 * alpha * eta ** 2 * gamma * rho ** 2
                + d ** 2 * eta ** 2 * rho ** 2
                + 2 * d * eta ** 2 * gamma * rho ** 2
                + eta ** 2 * gamma ** 2 * rho ** 2
                + 2 * H * alpha * eta * lam * sigma
                - 4 * H * d * eta * rho * sigma
                - 4 * H * eta * gamma * rho * sigma
                - 2 * alpha ** 2 * eta * lam * rho
                - 2 * alpha * d * eta * lam * rho
                - 2 * alpha * eta * gamma * lam * rho
                + 2 * H * d * eta * sigma
                + 2 * H * eta * gamma * sigma
                + alpha ** 2 * lam ** 2
                - 2 * alpha * d * eta * rho
                - 2 * alpha * eta * gamma * rho
                - 2 * d ** 2 * eta * rho
                - 4 * d * eta * gamma * rho
                - 2 * eta * gamma ** 2 * rho
                + 2 * alpha * d * lam
                + 2 * alpha * gamma * lam
                + d ** 2
                + 2 * d * gamma
                + gamma ** 2
            )
            * eta
            * rho
            + sqrt(
                H ** 2 * eta ** 2 * sigma ** 2
                + 2 * H * alpha * eta ** 2 * rho * sigma
                - 4 * H * alpha * eta * lam * rho * sigma
                + 2 * H * d * eta ** 2 * rho * sigma
                + 2 * H * eta ** 2 * gamma * rho * sigma
                + alpha ** 2 * eta ** 2 * rho ** 2
                + 2 * alpha * d * eta ** 2 * rho ** 2
                + 2 * alpha * eta ** 2 * gamma * rho ** 2
                + d ** 2 * eta ** 2 * rho ** 2
                + 2 * d * eta ** 2 * gamma * rho ** 2
                + eta ** 2 * gamma ** 2 * rho ** 2
                + 2 * H * alpha * eta * lam * sigma
                - 4 * H * d * eta * rho * sigma
                - 4 * H * eta * gamma * rho * sigma
                - 2 * alpha ** 2 * eta * lam * rho
                - 2 * alpha * d * eta * lam * rho
                - 2 * alpha * eta * gamma * lam * rho
                + 2 * H * d * eta * sigma
                + 2 * H * eta * gamma * sigma
                + alpha ** 2 * lam ** 2
                - 2 * alpha * d * eta * rho
                - 2 * alpha * eta * gamma * rho
                - 2 * d ** 2 * eta * rho
                - 4 * d * eta * gamma * rho
                - 2 * eta * gamma ** 2 * rho
                + 2 * alpha * d * lam
                + 2 * alpha * gamma * lam
                + d ** 2
                + 2 * d * gamma
                + gamma ** 2
            )
            * alpha
            * eta
            * rho
            + sqrt(
                H ** 2 * eta ** 2 * sigma ** 2
                + 2 * H * alpha * eta ** 2 * rho * sigma
                - 4 * H * alpha * eta * lam * rho * sigma
                + 2 * H * d * eta ** 2 * rho * sigma
                + 2 * H * eta ** 2 * gamma * rho * sigma
                + alpha ** 2 * eta ** 2 * rho ** 2
                + 2 * alpha * d * eta ** 2 * rho ** 2
                + 2 * alpha * eta ** 2 * gamma * rho ** 2
                + d ** 2 * eta ** 2 * rho ** 2
                + 2 * d * eta ** 2 * gamma * rho ** 2
                + eta ** 2 * gamma ** 2 * rho ** 2
                + 2 * H * alpha * eta * lam * sigma
                - 4 * H * d * eta * rho * sigma
                - 4 * H * eta * gamma * rho * sigma
                - 2 * alpha ** 2 * eta * lam * rho
                - 2 * alpha * d * eta * lam * rho
                - 2 * alpha * eta * gamma * lam * rho
                + 2 * H * d * eta * sigma
                + 2 * H * eta * gamma * sigma
                + alpha ** 2 * lam ** 2
                - 2 * alpha * d * eta * rho
                - 2 * alpha * eta * gamma * rho
                - 2 * d ** 2 * eta * rho
                - 4 * d * eta * gamma * rho
                - 2 * eta * gamma ** 2 * rho
                + 2 * alpha * d * lam
                + 2 * alpha * gamma * lam
                + d ** 2
                + 2 * d * gamma
                + gamma ** 2
            )
            * d
            * eta
            * rho
            + eta ** 2 * gamma ** 2 * rho ** 2
            - 2 * eta * gamma ** 2 * rho
            + H ** 2 * eta ** 2 * sigma ** 2
            + d ** 2 * eta ** 2 * rho ** 2
            - 2 * d ** 2 * eta * rho
        )
    )

    return alpha_sing_strat


def sing_strat_sigma(beta, alpha, sigma, H, eta, lam, gamma, d, rho):

    sigma_sing_strat = (
        (
            -2 * gamma * d * eta
            - alpha ** 2 * eta * lam
            - alpha ** 2 * eta ** 2 * rho
            - sqrt(
                H ** 2 * eta ** 2 * sigma ** 2
                + 2 * H * alpha * eta ** 2 * rho * sigma
                - 4 * H * alpha * eta * lam * rho * sigma
                + 2 * H * d * eta ** 2 * rho * sigma
                + 2 * H * eta ** 2 * gamma * rho * sigma
                + alpha ** 2 * eta ** 2 * rho ** 2
                + 2 * alpha * d * eta ** 2 * rho ** 2
                + 2 * alpha * eta ** 2 * gamma * rho ** 2
                + d ** 2 * eta ** 2 * rho ** 2
                + 2 * d * eta ** 2 * gamma * rho ** 2
                + eta ** 2 * gamma ** 2 * rho ** 2
                + 2 * H * alpha * eta * lam * sigma
                - 4 * H * d * eta * rho * sigma
                - 4 * H * eta * gamma * rho * sigma
                - 2 * alpha ** 2 * eta * lam * rho
                - 2 * alpha * d * eta * lam * rho
                - 2 * alpha * eta * gamma * lam * rho
                + 2 * H * d * eta * sigma
                + 2 * H * eta * gamma * sigma
                + alpha ** 2 * lam ** 2
                - 2 * alpha * d * eta * rho
                - 2 * alpha * eta * gamma * rho
                - 2 * d ** 2 * eta * rho
                - 4 * d * eta * gamma * rho
                - 2 * eta * gamma ** 2 * rho
                + 2 * alpha * d * lam
                + 2 * alpha * gamma * lam
                + d ** 2
                + 2 * d * gamma
                + gamma ** 2
            )
            * alpha
            * eta
            - gamma ** 2 * eta ** 2 * rho
            - gamma
            * sqrt(
                H ** 2 * eta ** 2 * sigma ** 2
                + 2 * H * alpha * eta ** 2 * rho * sigma
                - 4 * H * alpha * eta * lam * rho * sigma
                + 2 * H * d * eta ** 2 * rho * sigma
                + 2 * H * eta ** 2 * gamma * rho * sigma
                + alpha ** 2 * eta ** 2 * rho ** 2
                + 2 * alpha * d * eta ** 2 * rho ** 2
                + 2 * alpha * eta ** 2 * gamma * rho ** 2
                + d ** 2 * eta ** 2 * rho ** 2
                + 2 * d * eta ** 2 * gamma * rho ** 2
                + eta ** 2 * gamma ** 2 * rho ** 2
                + 2 * H * alpha * eta * lam * sigma
                - 4 * H * d * eta * rho * sigma
                - 4 * H * eta * gamma * rho * sigma
                - 2 * alpha ** 2 * eta * lam * rho
                - 2 * alpha * d * eta * lam * rho
                - 2 * alpha * eta * gamma * lam * rho
                + 2 * H * d * eta * sigma
                + 2 * H * eta * gamma * sigma
                + alpha ** 2 * lam ** 2
                - 2 * alpha * d * eta * rho
                - 2 * alpha * eta * gamma * rho
                - 2 * d ** 2 * eta * rho
                - 4 * d * eta * gamma * rho
                - 2 * eta * gamma ** 2 * rho
                + 2 * alpha * d * lam
                + 2 * alpha * gamma * lam
                + d ** 2
                + 2 * d * gamma
                + gamma ** 2
            )
            * eta
            - gamma * alpha * eta
            - d ** 2 * eta ** 2 * rho
            - sqrt(
                H ** 2 * eta ** 2 * sigma ** 2
                + 2 * H * alpha * eta ** 2 * rho * sigma
                - 4 * H * alpha * eta * lam * rho * sigma
                + 2 * H * d * eta ** 2 * rho * sigma
                + 2 * H * eta ** 2 * gamma * rho * sigma
                + alpha ** 2 * eta ** 2 * rho ** 2
                + 2 * alpha * d * eta ** 2 * rho ** 2
                + 2 * alpha * eta ** 2 * gamma * rho ** 2
                + d ** 2 * eta ** 2 * rho ** 2
                + 2 * d * eta ** 2 * gamma * rho ** 2
                + eta ** 2 * gamma ** 2 * rho ** 2
                + 2 * H * alpha * eta * lam * sigma
                - 4 * H * d * eta * rho * sigma
                - 4 * H * eta * gamma * rho * sigma
                - 2 * alpha ** 2 * eta * lam * rho
                - 2 * alpha * d * eta * lam * rho
                - 2 * alpha * eta * gamma * lam * rho
                + 2 * H * d * eta * sigma
                + 2 * H * eta * gamma * sigma
                + alpha ** 2 * lam ** 2
                - 2 * alpha * d * eta * rho
                - 2 * alpha * eta * gamma * rho
                - 2 * d ** 2 * eta * rho
                - 4 * d * eta * gamma * rho
                - 2 * eta * gamma ** 2 * rho
                + 2 * alpha * d * lam
                + 2 * alpha * gamma * lam
                + d ** 2
                + 2 * d * gamma
                + gamma ** 2
            )
            * d
            * eta
            - alpha * d * eta
            + d ** 2 * eta ** 2 * rho ** 2
            + eta ** 2 * gamma ** 2 * rho ** 2
            + alpha ** 2 * eta ** 2 * rho ** 2
            + H * alpha * eta ** 2 * rho * sigma
            + H * alpha * eta * lam * sigma
            + alpha ** 2 * lam ** 2
            - 2 * H * eta * gamma * rho * sigma
            + H * eta ** 2 * gamma * rho * sigma
            + H * d * eta ** 2 * rho * sigma
            - 2 * H * d * eta * rho * sigma
            - 2 * gamma * alpha * eta ** 2 * rho
            - 2 * gamma * d * eta ** 2 * rho
            - gamma * alpha * eta * lam
            - H * alpha * eta ** 2 * sigma
            - H * d * eta ** 2 * sigma
            - 2 * alpha * d * eta ** 2 * rho
            - alpha * d * eta * lam
            - H * gamma * eta ** 2 * sigma
            + 2 * d * eta ** 2 * gamma * rho ** 2
            + H * eta * gamma * sigma
            + H * d * eta * sigma
            + 2 * d * gamma
            + 2 * alpha * d * lam
            + 2 * alpha * gamma * lam
            + gamma ** 2
            + d ** 2
            - 2 * H * alpha * eta * lam * rho * sigma
            + 2 * alpha * d * eta ** 2 * rho ** 2
            + 2 * alpha * eta ** 2 * gamma * rho ** 2
            + sqrt(
                H ** 2 * eta ** 2 * sigma ** 2
                + 2 * H * alpha * eta ** 2 * rho * sigma
                - 4 * H * alpha * eta * lam * rho * sigma
                + 2 * H * d * eta ** 2 * rho * sigma
                + 2 * H * eta ** 2 * gamma * rho * sigma
                + alpha ** 2 * eta ** 2 * rho ** 2
                + 2 * alpha * d * eta ** 2 * rho ** 2
                + 2 * alpha * eta ** 2 * gamma * rho ** 2
                + d ** 2 * eta ** 2 * rho ** 2
                + 2 * d * eta ** 2 * gamma * rho ** 2
                + eta ** 2 * gamma ** 2 * rho ** 2
                + 2 * H * alpha * eta * lam * sigma
                - 4 * H * d * eta * rho * sigma
                - 4 * H * eta * gamma * rho * sigma
                - 2 * alpha ** 2 * eta * lam * rho
                - 2 * alpha * d * eta * lam * rho
                - 2 * alpha * eta * gamma * lam * rho
                + 2 * H * d * eta * sigma
                + 2 * H * eta * gamma * sigma
                + alpha ** 2 * lam ** 2
                - 2 * alpha * d * eta * rho
                - 2 * alpha * eta * gamma * rho
                - 2 * d ** 2 * eta * rho
                - 4 * d * eta * gamma * rho
                - 2 * eta * gamma ** 2 * rho
                + 2 * alpha * d * lam
                + 2 * alpha * gamma * lam
                + d ** 2
                + 2 * d * gamma
                + gamma ** 2
            )
            * alpha
            * lam
            + gamma
            * sqrt(
                H ** 2 * eta ** 2 * sigma ** 2
                + 2 * H * alpha * eta ** 2 * rho * sigma
                - 4 * H * alpha * eta * lam * rho * sigma
                + 2 * H * d * eta ** 2 * rho * sigma
                + 2 * H * eta ** 2 * gamma * rho * sigma
                + alpha ** 2 * eta ** 2 * rho ** 2
                + 2 * alpha * d * eta ** 2 * rho ** 2
                + 2 * alpha * eta ** 2 * gamma * rho ** 2
                + d ** 2 * eta ** 2 * rho ** 2
                + 2 * d * eta ** 2 * gamma * rho ** 2
                + eta ** 2 * gamma ** 2 * rho ** 2
                + 2 * H * alpha * eta * lam * sigma
                - 4 * H * d * eta * rho * sigma
                - 4 * H * eta * gamma * rho * sigma
                - 2 * alpha ** 2 * eta * lam * rho
                - 2 * alpha * d * eta * lam * rho
                - 2 * alpha * eta * gamma * lam * rho
                + 2 * H * d * eta * sigma
                + 2 * H * eta * gamma * sigma
                + alpha ** 2 * lam ** 2
                - 2 * alpha * d * eta * rho
                - 2 * alpha * eta * gamma * rho
                - 2 * d ** 2 * eta * rho
                - 4 * d * eta * gamma * rho
                - 2 * eta * gamma ** 2 * rho
                + 2 * alpha * d * lam
                + 2 * alpha * gamma * lam
                + d ** 2
                + 2 * d * gamma
                + gamma ** 2
            )
            + sqrt(
                H ** 2 * eta ** 2 * sigma ** 2
                + 2 * H * alpha * eta ** 2 * rho * sigma
                - 4 * H * alpha * eta * lam * rho * sigma
                + 2 * H * d * eta ** 2 * rho * sigma
                + 2 * H * eta ** 2 * gamma * rho * sigma
                + alpha ** 2 * eta ** 2 * rho ** 2
                + 2 * alpha * d * eta ** 2 * rho ** 2
                + 2 * alpha * eta ** 2 * gamma * rho ** 2
                + d ** 2 * eta ** 2 * rho ** 2
                + 2 * d * eta ** 2 * gamma * rho ** 2
                + eta ** 2 * gamma ** 2 * rho ** 2
                + 2 * H * alpha * eta * lam * sigma
                - 4 * H * d * eta * rho * sigma
                - 4 * H * eta * gamma * rho * sigma
                - 2 * alpha ** 2 * eta * lam * rho
                - 2 * alpha * d * eta * lam * rho
                - 2 * alpha * eta * gamma * lam * rho
                + 2 * H * d * eta * sigma
                + 2 * H * eta * gamma * sigma
                + alpha ** 2 * lam ** 2
                - 2 * alpha * d * eta * rho
                - 2 * alpha * eta * gamma * rho
                - 2 * d ** 2 * eta * rho
                - 4 * d * eta * gamma * rho
                - 2 * eta * gamma ** 2 * rho
                + 2 * alpha * d * lam
                + 2 * alpha * gamma * lam
                + d ** 2
                + 2 * d * gamma
                + gamma ** 2
            )
            * d
            + gamma
            * sqrt(
                H ** 2 * eta ** 2 * sigma ** 2
                + 2 * H * alpha * eta ** 2 * rho * sigma
                - 4 * H * alpha * eta * lam * rho * sigma
                + 2 * H * d * eta ** 2 * rho * sigma
                + 2 * H * eta ** 2 * gamma * rho * sigma
                + alpha ** 2 * eta ** 2 * rho ** 2
                + 2 * alpha * d * eta ** 2 * rho ** 2
                + 2 * alpha * eta ** 2 * gamma * rho ** 2
                + d ** 2 * eta ** 2 * rho ** 2
                + 2 * d * eta ** 2 * gamma * rho ** 2
                + eta ** 2 * gamma ** 2 * rho ** 2
                + 2 * H * alpha * eta * lam * sigma
                - 4 * H * d * eta * rho * sigma
                - 4 * H * eta * gamma * rho * sigma
                - 2 * alpha ** 2 * eta * lam * rho
                - 2 * alpha * d * eta * lam * rho
                - 2 * alpha * eta * gamma * lam * rho
                + 2 * H * d * eta * sigma
                + 2 * H * eta * gamma * sigma
                + alpha ** 2 * lam ** 2
                - 2 * alpha * d * eta * rho
                - 2 * alpha * eta * gamma * rho
                - 2 * d ** 2 * eta * rho
                - 4 * d * eta * gamma * rho
                - 2 * eta * gamma ** 2 * rho
                + 2 * alpha * d * lam
                + 2 * alpha * gamma * lam
                + d ** 2
                + 2 * d * gamma
                + gamma ** 2
            )
            * eta
            * rho
            + sqrt(
                H ** 2 * eta ** 2 * sigma ** 2
                + 2 * H * alpha * eta ** 2 * rho * sigma
                - 4 * H * alpha * eta * lam * rho * sigma
                + 2 * H * d * eta ** 2 * rho * sigma
                + 2 * H * eta ** 2 * gamma * rho * sigma
                + alpha ** 2 * eta ** 2 * rho ** 2
                + 2 * alpha * d * eta ** 2 * rho ** 2
                + 2 * alpha * eta ** 2 * gamma * rho ** 2
                + d ** 2 * eta ** 2 * rho ** 2
                + 2 * d * eta ** 2 * gamma * rho ** 2
                + eta ** 2 * gamma ** 2 * rho ** 2
                + 2 * H * alpha * eta * lam * sigma
                - 4 * H * d * eta * rho * sigma
                - 4 * H * eta * gamma * rho * sigma
                - 2 * alpha ** 2 * eta * lam * rho
                - 2 * alpha * d * eta * lam * rho
                - 2 * alpha * eta * gamma * lam * rho
                + 2 * H * d * eta * sigma
                + 2 * H * eta * gamma * sigma
                + alpha ** 2 * lam ** 2
                - 2 * alpha * d * eta * rho
                - 2 * alpha * eta * gamma * rho
                - 2 * d ** 2 * eta * rho
                - 4 * d * eta * gamma * rho
                - 2 * eta * gamma ** 2 * rho
                + 2 * alpha * d * lam
                + 2 * alpha * gamma * lam
                + d ** 2
                + 2 * d * gamma
                + gamma ** 2
            )
            * alpha
            * eta
            * rho
            + sqrt(
                H ** 2 * eta ** 2 * sigma ** 2
                + 2 * H * alpha * eta ** 2 * rho * sigma
                - 4 * H * alpha * eta * lam * rho * sigma
                + 2 * H * d * eta ** 2 * rho * sigma
                + 2 * H * eta ** 2 * gamma * rho * sigma
                + alpha ** 2 * eta ** 2 * rho ** 2
                + 2 * alpha * d * eta ** 2 * rho ** 2
                + 2 * alpha * eta ** 2 * gamma * rho ** 2
                + d ** 2 * eta ** 2 * rho ** 2
                + 2 * d * eta ** 2 * gamma * rho ** 2
                + eta ** 2 * gamma ** 2 * rho ** 2
                + 2 * H * alpha * eta * lam * sigma
                - 4 * H * d * eta * rho * sigma
                - 4 * H * eta * gamma * rho * sigma
                - 2 * alpha ** 2 * eta * lam * rho
                - 2 * alpha * d * eta * lam * rho
                - 2 * alpha * eta * gamma * lam * rho
                + 2 * H * d * eta * sigma
                + 2 * H * eta * gamma * sigma
                + alpha ** 2 * lam ** 2
                - 2 * alpha * d * eta * rho
                - 2 * alpha * eta * gamma * rho
                - 2 * d ** 2 * eta * rho
                - 4 * d * eta * gamma * rho
                - 2 * eta * gamma ** 2 * rho
                + 2 * alpha * d * lam
                + 2 * alpha * gamma * lam
                + d ** 2
                + 2 * d * gamma
                + gamma ** 2
            )
            * d
            * eta
            * rho
            - d ** 2 * eta
            - gamma ** 2 * eta
        )
        * H
        * beta
        / (sigma * H + alpha + d + gamma)
        / (
            H ** 2 * eta ** 2 * sigma ** 2
            + d ** 2 * eta ** 2 * rho ** 2
            - 2 * d ** 2 * eta * rho
            + eta ** 2 * gamma ** 2 * rho ** 2
            - 2 * eta * gamma ** 2 * rho
            + alpha ** 2 * eta ** 2 * rho ** 2
            + 2 * H * alpha * eta ** 2 * rho * sigma
            + 2 * H * alpha * eta * lam * sigma
            - 2 * alpha * d * eta * lam * rho
            - 2 * alpha * eta * gamma * lam * rho
            + alpha ** 2 * lam ** 2
            - 4 * H * eta * gamma * rho * sigma
            + 2 * H * eta ** 2 * gamma * rho * sigma
            + 2 * H * d * eta ** 2 * rho * sigma
            - 4 * H * d * eta * rho * sigma
            + 2 * d * eta ** 2 * gamma * rho ** 2
            + 2 * H * eta * gamma * sigma
            - 4 * d * eta * gamma * rho
            + 2 * H * d * eta * sigma
            + 2 * d * gamma
            + 2 * alpha * d * lam
            + 2 * alpha * gamma * lam
            + gamma ** 2
            + d ** 2
            - 4 * H * alpha * eta * lam * rho * sigma
            + 2 * alpha * d * eta ** 2 * rho ** 2
            + 2 * alpha * eta ** 2 * gamma * rho ** 2
            - 2 * alpha ** 2 * eta * lam * rho
            - 2 * alpha * d * eta * rho
            - 2 * alpha * eta * gamma * rho
            + sqrt(
                H ** 2 * eta ** 2 * sigma ** 2
                + 2 * H * alpha * eta ** 2 * rho * sigma
                - 4 * H * alpha * eta * lam * rho * sigma
                + 2 * H * d * eta ** 2 * rho * sigma
                + 2 * H * eta ** 2 * gamma * rho * sigma
                + alpha ** 2 * eta ** 2 * rho ** 2
                + 2 * alpha * d * eta ** 2 * rho ** 2
                + 2 * alpha * eta ** 2 * gamma * rho ** 2
                + d ** 2 * eta ** 2 * rho ** 2
                + 2 * d * eta ** 2 * gamma * rho ** 2
                + eta ** 2 * gamma ** 2 * rho ** 2
                + 2 * H * alpha * eta * lam * sigma
                - 4 * H * d * eta * rho * sigma
                - 4 * H * eta * gamma * rho * sigma
                - 2 * alpha ** 2 * eta * lam * rho
                - 2 * alpha * d * eta * lam * rho
                - 2 * alpha * eta * gamma * lam * rho
                + 2 * H * d * eta * sigma
                + 2 * H * eta * gamma * sigma
                + alpha ** 2 * lam ** 2
                - 2 * alpha * d * eta * rho
                - 2 * alpha * eta * gamma * rho
                - 2 * d ** 2 * eta * rho
                - 4 * d * eta * gamma * rho
                - 2 * eta * gamma ** 2 * rho
                + 2 * alpha * d * lam
                + 2 * alpha * gamma * lam
                + d ** 2
                + 2 * d * gamma
                + gamma ** 2
            )
            * alpha
            * lam
            + gamma
            * sqrt(
                H ** 2 * eta ** 2 * sigma ** 2
                + 2 * H * alpha * eta ** 2 * rho * sigma
                - 4 * H * alpha * eta * lam * rho * sigma
                + 2 * H * d * eta ** 2 * rho * sigma
                + 2 * H * eta ** 2 * gamma * rho * sigma
                + alpha ** 2 * eta ** 2 * rho ** 2
                + 2 * alpha * d * eta ** 2 * rho ** 2
                + 2 * alpha * eta ** 2 * gamma * rho ** 2
                + d ** 2 * eta ** 2 * rho ** 2
                + 2 * d * eta ** 2 * gamma * rho ** 2
                + eta ** 2 * gamma ** 2 * rho ** 2
                + 2 * H * alpha * eta * lam * sigma
                - 4 * H * d * eta * rho * sigma
                - 4 * H * eta * gamma * rho * sigma
                - 2 * alpha ** 2 * eta * lam * rho
                - 2 * alpha * d * eta * lam * rho
                - 2 * alpha * eta * gamma * lam * rho
                + 2 * H * d * eta * sigma
                + 2 * H * eta * gamma * sigma
                + alpha ** 2 * lam ** 2
                - 2 * alpha * d * eta * rho
                - 2 * alpha * eta * gamma * rho
                - 2 * d ** 2 * eta * rho
                - 4 * d * eta * gamma * rho
                - 2 * eta * gamma ** 2 * rho
                + 2 * alpha * d * lam
                + 2 * alpha * gamma * lam
                + d ** 2
                + 2 * d * gamma
                + gamma ** 2
            )
            + sqrt(
                H ** 2 * eta ** 2 * sigma ** 2
                + 2 * H * alpha * eta ** 2 * rho * sigma
                - 4 * H * alpha * eta * lam * rho * sigma
                + 2 * H * d * eta ** 2 * rho * sigma
                + 2 * H * eta ** 2 * gamma * rho * sigma
                + alpha ** 2 * eta ** 2 * rho ** 2
                + 2 * alpha * d * eta ** 2 * rho ** 2
                + 2 * alpha * eta ** 2 * gamma * rho ** 2
                + d ** 2 * eta ** 2 * rho ** 2
                + 2 * d * eta ** 2 * gamma * rho ** 2
                + eta ** 2 * gamma ** 2 * rho ** 2
                + 2 * H * alpha * eta * lam * sigma
                - 4 * H * d * eta * rho * sigma
                - 4 * H * eta * gamma * rho * sigma
                - 2 * alpha ** 2 * eta * lam * rho
                - 2 * alpha * d * eta * lam * rho
                - 2 * alpha * eta * gamma * lam * rho
                + 2 * H * d * eta * sigma
                + 2 * H * eta * gamma * sigma
                + alpha ** 2 * lam ** 2
                - 2 * alpha * d * eta * rho
                - 2 * alpha * eta * gamma * rho
                - 2 * d ** 2 * eta * rho
                - 4 * d * eta * gamma * rho
                - 2 * eta * gamma ** 2 * rho
                + 2 * alpha * d * lam
                + 2 * alpha * gamma * lam
                + d ** 2
                + 2 * d * gamma
                + gamma ** 2
            )
            * d
            + H
            * sqrt(
                H ** 2 * eta ** 2 * sigma ** 2
                + 2 * H * alpha * eta ** 2 * rho * sigma
                - 4 * H * alpha * eta * lam * rho * sigma
                + 2 * H * d * eta ** 2 * rho * sigma
                + 2 * H * eta ** 2 * gamma * rho * sigma
                + alpha ** 2 * eta ** 2 * rho ** 2
                + 2 * alpha * d * eta ** 2 * rho ** 2
                + 2 * alpha * eta ** 2 * gamma * rho ** 2
                + d ** 2 * eta ** 2 * rho ** 2
                + 2 * d * eta ** 2 * gamma * rho ** 2
                + eta ** 2 * gamma ** 2 * rho ** 2
                + 2 * H * alpha * eta * lam * sigma
                - 4 * H * d * eta * rho * sigma
                - 4 * H * eta * gamma * rho * sigma
                - 2 * alpha ** 2 * eta * lam * rho
                - 2 * alpha * d * eta * lam * rho
                - 2 * alpha * eta * gamma * lam * rho
                + 2 * H * d * eta * sigma
                + 2 * H * eta * gamma * sigma
                + alpha ** 2 * lam ** 2
                - 2 * alpha * d * eta * rho
                - 2 * alpha * eta * gamma * rho
                - 2 * d ** 2 * eta * rho
                - 4 * d * eta * gamma * rho
                - 2 * eta * gamma ** 2 * rho
                + 2 * alpha * d * lam
                + 2 * alpha * gamma * lam
                + d ** 2
                + 2 * d * gamma
                + gamma ** 2
            )
            * eta
            * sigma
            + gamma
            * sqrt(
                H ** 2 * eta ** 2 * sigma ** 2
                + 2 * H * alpha * eta ** 2 * rho * sigma
                - 4 * H * alpha * eta * lam * rho * sigma
                + 2 * H * d * eta ** 2 * rho * sigma
                + 2 * H * eta ** 2 * gamma * rho * sigma
                + alpha ** 2 * eta ** 2 * rho ** 2
                + 2 * alpha * d * eta ** 2 * rho ** 2
                + 2 * alpha * eta ** 2 * gamma * rho ** 2
                + d ** 2 * eta ** 2 * rho ** 2
                + 2 * d * eta ** 2 * gamma * rho ** 2
                + eta ** 2 * gamma ** 2 * rho ** 2
                + 2 * H * alpha * eta * lam * sigma
                - 4 * H * d * eta * rho * sigma
                - 4 * H * eta * gamma * rho * sigma
                - 2 * alpha ** 2 * eta * lam * rho
                - 2 * alpha * d * eta * lam * rho
                - 2 * alpha * eta * gamma * lam * rho
                + 2 * H * d * eta * sigma
                + 2 * H * eta * gamma * sigma
                + alpha ** 2 * lam ** 2
                - 2 * alpha * d * eta * rho
                - 2 * alpha * eta * gamma * rho
                - 2 * d ** 2 * eta * rho
                - 4 * d * eta * gamma * rho
                - 2 * eta * gamma ** 2 * rho
                + 2 * alpha * d * lam
                + 2 * alpha * gamma * lam
                + d ** 2
                + 2 * d * gamma
                + gamma ** 2
            )
            * eta
            * rho
            + sqrt(
                H ** 2 * eta ** 2 * sigma ** 2
                + 2 * H * alpha * eta ** 2 * rho * sigma
                - 4 * H * alpha * eta * lam * rho * sigma
                + 2 * H * d * eta ** 2 * rho * sigma
                + 2 * H * eta ** 2 * gamma * rho * sigma
                + alpha ** 2 * eta ** 2 * rho ** 2
                + 2 * alpha * d * eta ** 2 * rho ** 2
                + 2 * alpha * eta ** 2 * gamma * rho ** 2
                + d ** 2 * eta ** 2 * rho ** 2
                + 2 * d * eta ** 2 * gamma * rho ** 2
                + eta ** 2 * gamma ** 2 * rho ** 2
                + 2 * H * alpha * eta * lam * sigma
                - 4 * H * d * eta * rho * sigma
                - 4 * H * eta * gamma * rho * sigma
                - 2 * alpha ** 2 * eta * lam * rho
                - 2 * alpha * d * eta * lam * rho
                - 2 * alpha * eta * gamma * lam * rho
                + 2 * H * d * eta * sigma
                + 2 * H * eta * gamma * sigma
                + alpha ** 2 * lam ** 2
                - 2 * alpha * d * eta * rho
                - 2 * alpha * eta * gamma * rho
                - 2 * d ** 2 * eta * rho
                - 4 * d * eta * gamma * rho
                - 2 * eta * gamma ** 2 * rho
                + 2 * alpha * d * lam
                + 2 * alpha * gamma * lam
                + d ** 2
                + 2 * d * gamma
                + gamma ** 2
            )
            * alpha
            * eta
            * rho
            + sqrt(
                H ** 2 * eta ** 2 * sigma ** 2
                + 2 * H * alpha * eta ** 2 * rho * sigma
                - 4 * H * alpha * eta * lam * rho * sigma
                + 2 * H * d * eta ** 2 * rho * sigma
                + 2 * H * eta ** 2 * gamma * rho * sigma
                + alpha ** 2 * eta ** 2 * rho ** 2
                + 2 * alpha * d * eta ** 2 * rho ** 2
                + 2 * alpha * eta ** 2 * gamma * rho ** 2
                + d ** 2 * eta ** 2 * rho ** 2
                + 2 * d * eta ** 2 * gamma * rho ** 2
                + eta ** 2 * gamma ** 2 * rho ** 2
                + 2 * H * alpha * eta * lam * sigma
                - 4 * H * d * eta * rho * sigma
                - 4 * H * eta * gamma * rho * sigma
                - 2 * alpha ** 2 * eta * lam * rho
                - 2 * alpha * d * eta * lam * rho
                - 2 * alpha * eta * gamma * lam * rho
                + 2 * H * d * eta * sigma
                + 2 * H * eta * gamma * sigma
                + alpha ** 2 * lam ** 2
                - 2 * alpha * d * eta * rho
                - 2 * alpha * eta * gamma * rho
                - 2 * d ** 2 * eta * rho
                - 4 * d * eta * gamma * rho
                - 2 * eta * gamma ** 2 * rho
                + 2 * alpha * d * lam
                + 2 * alpha * gamma * lam
                + d ** 2
                + 2 * d * gamma
                + gamma ** 2
            )
            * d
            * eta
            * rho
        )
    )
    return sigma_sing_strat


def fitness_gradient_alpha(
    beta, alpha, sigma, dbetadalpha, H, S, eta, lam, gamma, d, rho
):

    alpha_fitness_gradient = (
        -0.1e1
        / (
            H * alpha * lam * sigma
            + H * d * sigma
            + H * gamma * sigma
            + alpha ** 2 * lam
            + alpha * d * lam
            + alpha * gamma * lam
            + d * alpha
            + alpha * gamma
            + d ** 2
            + 2 * d * gamma
            + gamma ** 2
        )
        ** 2
        * (
            eta * sigma * H
            + d * eta * rho
            + eta * gamma * rho
            + eta * rho * alpha
            + lam * alpha
            + d
            + gamma
            + sqrt(
                max(
                    0.0,
                    H ** 2 * eta ** 2 * sigma ** 2
                    + 2 * H * alpha * eta ** 2 * rho * sigma
                    - 4 * H * alpha * eta * lam * rho * sigma
                    + 2 * H * d * eta ** 2 * rho * sigma
                    + 2 * H * eta ** 2 * gamma * rho * sigma
                    + alpha ** 2 * eta ** 2 * rho ** 2
                    + 2 * alpha * d * eta ** 2 * rho ** 2
                    + 2 * alpha * eta ** 2 * gamma * rho ** 2
                    + d ** 2 * eta ** 2 * rho ** 2
                    + 2 * d * eta ** 2 * gamma * rho ** 2
                    + eta ** 2 * gamma ** 2 * rho ** 2
                    + 2 * H * alpha * eta * lam * sigma
                    - 4 * H * d * eta * rho * sigma
                    - 4 * H * eta * gamma * rho * sigma
                    - 2 * alpha ** 2 * eta * lam * rho
                    - 2 * alpha * d * eta * lam * rho
                    - 2 * alpha * eta * gamma * lam * rho
                    + 2 * H * d * eta * sigma
                    + 2 * H * eta * gamma * sigma
                    + alpha ** 2 * lam ** 2
                    - 2 * alpha * d * eta * rho
                    - 2 * alpha * eta * gamma * rho
                    - 2 * d ** 2 * eta * rho
                    - 4 * d * eta * gamma * rho
                    - 2 * eta * gamma ** 2 * rho
                    + 2 * alpha * d * lam
                    + 2 * alpha * gamma * lam
                    + d ** 2
                    + 2 * d * gamma
                    + gamma ** 2,
                )
            )
        )
        * S
        * beta
        * (H * lam * sigma + 2 * lam * alpha + d * lam + gamma * lam + d + gamma)
        / 2
        + 0.1e1
        / (
            H * alpha * lam * sigma
            + H * d * sigma
            + H * gamma * sigma
            + alpha ** 2 * lam
            + alpha * d * lam
            + alpha * gamma * lam
            + d * alpha
            + alpha * gamma
            + d ** 2
            + 2 * d * gamma
            + gamma ** 2
        )
        * (
            rho * eta
            + lam
            + (
                max(
                    H ** 2 * eta ** 2 * sigma ** 2
                    + 2 * H * alpha * eta ** 2 * rho * sigma
                    - 4 * H * alpha * eta * lam * rho * sigma
                    + 2 * H * d * eta ** 2 * rho * sigma
                    + 2 * H * eta ** 2 * gamma * rho * sigma
                    + alpha ** 2 * eta ** 2 * rho ** 2
                    + 2 * alpha * d * eta ** 2 * rho ** 2
                    + 2 * alpha * eta ** 2 * gamma * rho ** 2
                    + d ** 2 * eta ** 2 * rho ** 2
                    + 2 * d * eta ** 2 * gamma * rho ** 2
                    + eta ** 2 * gamma ** 2 * rho ** 2
                    + 2 * H * alpha * eta * lam * sigma
                    - 4 * H * d * eta * rho * sigma
                    - 4 * H * eta * gamma * rho * sigma
                    - 2 * alpha ** 2 * eta * lam * rho
                    - 2 * alpha * d * eta * lam * rho
                    - 2 * alpha * eta * gamma * lam * rho
                    + 2 * H * d * eta * sigma
                    + 2 * H * eta * gamma * sigma
                    + alpha ** 2 * lam ** 2
                    - 2 * alpha * d * eta * rho
                    - 2 * alpha * eta * gamma * rho
                    - 2 * d ** 2 * eta * rho
                    - 4 * d * eta * gamma * rho
                    - 2 * eta * gamma ** 2 * rho
                    + 2 * alpha * d * lam
                    + 2 * alpha * gamma * lam
                    + d ** 2
                    + 2 * d * gamma
                    + gamma ** 2,
                    1e-6,
                )
            )
            ** (-0.1e1 / 0.2e1)
            * (
                2 * H * eta ** 2 * rho * sigma
                - 4 * H * eta * lam * rho * sigma
                + 2 * alpha * eta ** 2 * rho ** 2
                + 2 * d * eta ** 2 * rho ** 2
                + 2 * eta ** 2 * gamma * rho ** 2
                + 2 * H * eta * lam * sigma
                - 4 * alpha * eta * lam * rho
                - 2 * d * eta * lam * rho
                - 2 * eta * gamma * lam * rho
                + 2 * alpha * lam ** 2
                - 2 * d * eta * rho
                - 2 * eta * gamma * rho
                + 2 * d * lam
                + 2 * gamma * lam
            )
            / 2
        )
        * S
        * beta
        / 2
        + 0.1e1
        / (
            H * alpha * lam * sigma
            + H * d * sigma
            + H * gamma * sigma
            + alpha ** 2 * lam
            + alpha * d * lam
            + alpha * gamma * lam
            + d * alpha
            + alpha * gamma
            + d ** 2
            + 2 * d * gamma
            + gamma ** 2
        )
        * (
            eta * sigma * H
            + d * eta * rho
            + eta * gamma * rho
            + eta * rho * alpha
            + lam * alpha
            + d
            + gamma
            + sqrt(
                max(
                    0.0,
                    H ** 2 * eta ** 2 * sigma ** 2
                    + 2 * H * alpha * eta ** 2 * rho * sigma
                    - 4 * H * alpha * eta * lam * rho * sigma
                    + 2 * H * d * eta ** 2 * rho * sigma
                    + 2 * H * eta ** 2 * gamma * rho * sigma
                    + alpha ** 2 * eta ** 2 * rho ** 2
                    + 2 * alpha * d * eta ** 2 * rho ** 2
                    + 2 * alpha * eta ** 2 * gamma * rho ** 2
                    + d ** 2 * eta ** 2 * rho ** 2
                    + 2 * d * eta ** 2 * gamma * rho ** 2
                    + eta ** 2 * gamma ** 2 * rho ** 2
                    + 2 * H * alpha * eta * lam * sigma
                    - 4 * H * d * eta * rho * sigma
                    - 4 * H * eta * gamma * rho * sigma
                    - 2 * alpha ** 2 * eta * lam * rho
                    - 2 * alpha * d * eta * lam * rho
                    - 2 * alpha * eta * gamma * lam * rho
                    + 2 * H * d * eta * sigma
                    + 2 * H * eta * gamma * sigma
                    + alpha ** 2 * lam ** 2
                    - 2 * alpha * d * eta * rho
                    - 2 * alpha * eta * gamma * rho
                    - 2 * d ** 2 * eta * rho
                    - 4 * d * eta * gamma * rho
                    - 2 * eta * gamma ** 2 * rho
                    + 2 * alpha * d * lam
                    + 2 * alpha * gamma * lam
                    + d ** 2
                    + 2 * d * gamma
                    + gamma ** 2,
                )
            )
        )
        * S
        * dbetadalpha
        / 2
    )

    return alpha_fitness_gradient


def fitness_gradient_sigma(
    beta, alpha, sigma, dbetadsigma, H, S, eta, lam, gamma, d, rho
):

    sigma_fitness_gradient = (
        -0.1e1
        / (
            H * alpha * lam * sigma
            + H * d * sigma
            + H * gamma * sigma
            + alpha ** 2 * lam
            + alpha * d * lam
            + alpha * gamma * lam
            + d * alpha
            + alpha * gamma
            + d ** 2
            + 2 * d * gamma
            + gamma ** 2
        )
        ** 2
        * (
            eta * sigma * H
            + d * eta * rho
            + eta * gamma * rho
            + eta * rho * alpha
            + lam * alpha
            + d
            + gamma
            + sqrt(
                max(
                    H ** 2 * eta ** 2 * sigma ** 2
                    + 2 * H * alpha * eta ** 2 * rho * sigma
                    - 4 * H * alpha * eta * lam * rho * sigma
                    + 2 * H * d * eta ** 2 * rho * sigma
                    + 2 * H * eta ** 2 * gamma * rho * sigma
                    + alpha ** 2 * eta ** 2 * rho ** 2
                    + 2 * alpha * d * eta ** 2 * rho ** 2
                    + 2 * alpha * eta ** 2 * gamma * rho ** 2
                    + d ** 2 * eta ** 2 * rho ** 2
                    + 2 * d * eta ** 2 * gamma * rho ** 2
                    + eta ** 2 * gamma ** 2 * rho ** 2
                    + 2 * H * alpha * eta * lam * sigma
                    - 4 * H * d * eta * rho * sigma
                    - 4 * H * eta * gamma * rho * sigma
                    - 2 * alpha ** 2 * eta * lam * rho
                    - 2 * alpha * d * eta * lam * rho
                    - 2 * alpha * eta * gamma * lam * rho
                    + 2 * H * d * eta * sigma
                    + 2 * H * eta * gamma * sigma
                    + alpha ** 2 * lam ** 2
                    - 2 * alpha * d * eta * rho
                    - 2 * alpha * eta * gamma * rho
                    - 2 * d ** 2 * eta * rho
                    - 4 * d * eta * gamma * rho
                    - 2 * eta * gamma ** 2 * rho
                    + 2 * alpha * d * lam
                    + 2 * alpha * gamma * lam
                    + d ** 2
                    + 2 * d * gamma
                    + gamma ** 2,
                    1e-6,
                )
            )
        )
        * S
        * beta
        * (H * lam * alpha + H * d + gamma * H)
        / 2
        + 0.1e1
        / (
            H * alpha * lam * sigma
            + H * d * sigma
            + H * gamma * sigma
            + alpha ** 2 * lam
            + alpha * d * lam
            + alpha * gamma * lam
            + d * alpha
            + alpha * gamma
            + d ** 2
            + 2 * d * gamma
            + gamma ** 2
        )
        * (
            eta * H
            + (
                max(
                    H ** 2 * eta ** 2 * sigma ** 2
                    + 2 * H * alpha * eta ** 2 * rho * sigma
                    - 4 * H * alpha * eta * lam * rho * sigma
                    + 2 * H * d * eta ** 2 * rho * sigma
                    + 2 * H * eta ** 2 * gamma * rho * sigma
                    + alpha ** 2 * eta ** 2 * rho ** 2
                    + 2 * alpha * d * eta ** 2 * rho ** 2
                    + 2 * alpha * eta ** 2 * gamma * rho ** 2
                    + d ** 2 * eta ** 2 * rho ** 2
                    + 2 * d * eta ** 2 * gamma * rho ** 2
                    + eta ** 2 * gamma ** 2 * rho ** 2
                    + 2 * H * alpha * eta * lam * sigma
                    - 4 * H * d * eta * rho * sigma
                    - 4 * H * eta * gamma * rho * sigma
                    - 2 * alpha ** 2 * eta * lam * rho
                    - 2 * alpha * d * eta * lam * rho
                    - 2 * alpha * eta * gamma * lam * rho
                    + 2 * H * d * eta * sigma
                    + 2 * H * eta * gamma * sigma
                    + alpha ** 2 * lam ** 2
                    - 2 * alpha * d * eta * rho
                    - 2 * alpha * eta * gamma * rho
                    - 2 * d ** 2 * eta * rho
                    - 4 * d * eta * gamma * rho
                    - 2 * eta * gamma ** 2 * rho
                    + 2 * alpha * d * lam
                    + 2 * alpha * gamma * lam
                    + d ** 2
                    + 2 * d * gamma
                    + gamma ** 2,
                    1e-6,
                )
            )
            ** (-0.1e1 / 0.2e1)
            * (
                2 * H ** 2 * eta ** 2 * sigma
                + 2 * H * alpha * eta ** 2 * rho
                - 4 * H * alpha * eta * lam * rho
                + 2 * H * d * eta ** 2 * rho
                + 2 * H * eta ** 2 * gamma * rho
                + 2 * H * alpha * eta * lam
                - 4 * H * d * eta * rho
                - 4 * H * eta * gamma * rho
                + 2 * H * d * eta
                + 2 * H * eta * gamma
            )
            / 2
        )
        * S
        * beta
        / 2
        + 0.1e1
        / (
            H * alpha * lam * sigma
            + H * d * sigma
            + H * gamma * sigma
            + alpha ** 2 * lam
            + alpha * d * lam
            + alpha * gamma * lam
            + d * alpha
            + alpha * gamma
            + d ** 2
            + 2 * d * gamma
            + gamma ** 2
        )
        * (
            eta * sigma * H
            + d * eta * rho
            + eta * gamma * rho
            + eta * rho * alpha
            + lam * alpha
            + d
            + gamma
            + sqrt(
                max(
                    1e-6,
                    H ** 2 * eta ** 2 * sigma ** 2
                    + 2 * H * alpha * eta ** 2 * rho * sigma
                    - 4 * H * alpha * eta * lam * rho * sigma
                    + 2 * H * d * eta ** 2 * rho * sigma
                    + 2 * H * eta ** 2 * gamma * rho * sigma
                    + alpha ** 2 * eta ** 2 * rho ** 2
                    + 2 * alpha * d * eta ** 2 * rho ** 2
                    + 2 * alpha * eta ** 2 * gamma * rho ** 2
                    + d ** 2 * eta ** 2 * rho ** 2
                    + 2 * d * eta ** 2 * gamma * rho ** 2
                    + eta ** 2 * gamma ** 2 * rho ** 2
                    + 2 * H * alpha * eta * lam * sigma
                    - 4 * H * d * eta * rho * sigma
                    - 4 * H * eta * gamma * rho * sigma
                    - 2 * alpha ** 2 * eta * lam * rho
                    - 2 * alpha * d * eta * lam * rho
                    - 2 * alpha * eta * gamma * lam * rho
                    + 2 * H * d * eta * sigma
                    + 2 * H * eta * gamma * sigma
                    + alpha ** 2 * lam ** 2
                    - 2 * alpha * d * eta * rho
                    - 2 * alpha * eta * gamma * rho
                    - 2 * d ** 2 * eta * rho
                    - 4 * d * eta * gamma * rho
                    - 2 * eta * gamma ** 2 * rho
                    + 2 * alpha * d * lam
                    + 2 * alpha * gamma * lam
                    + d ** 2
                    + 2 * d * gamma
                    + gamma ** 2,
                )
            )
        )
        * S
        * dbetadsigma
        / 2
    )

    return sigma_fitness_gradient


def fit_grad_alpha_no_sub(
    beta_m, alpha_m, sigma, dbetadalpha_m, H, S, eta, lam, gamma, d, rho
):

    fit_grad_no_sub = (
        -0.1e1
        / (
            H * lam * sigma * alpha_m
            + H * d * sigma
            + H * gamma * sigma
            + d * lam * alpha_m
            + gamma * lam * alpha_m
            + lam * alpha_m ** 2
            + d ** 2
            + 2 * d * gamma
            + d * alpha_m
            + gamma ** 2
            + gamma * alpha_m
        )
        ** 2
        * (
            eta * sigma * H
            + d * eta * rho
            + eta * gamma * rho
            + eta * rho * alpha_m
            + lam * alpha_m
            + d
            + gamma
            + sqrt(
                H ** 2 * eta ** 2 * sigma ** 2
                + 2 * H * d * eta ** 2 * rho * sigma
                + 2 * H * eta ** 2 * gamma * rho * sigma
                + 2 * H * eta ** 2 * rho * sigma * alpha_m
                - 4 * H * eta * lam * rho * sigma * alpha_m
                + d ** 2 * eta ** 2 * rho ** 2
                + 2 * d * eta ** 2 * gamma * rho ** 2
                + 2 * d * eta ** 2 * rho ** 2 * alpha_m
                + eta ** 2 * gamma ** 2 * rho ** 2
                + 2 * eta ** 2 * gamma * rho ** 2 * alpha_m
                + eta ** 2 * rho ** 2 * alpha_m ** 2
                - 4 * H * d * eta * rho * sigma
                - 4 * H * eta * gamma * rho * sigma
                + 2 * H * eta * lam * sigma * alpha_m
                - 2 * d * eta * lam * rho * alpha_m
                - 2 * eta * gamma * lam * rho * alpha_m
                - 2 * eta * lam * rho * alpha_m ** 2
                + 2 * H * d * eta * sigma
                + 2 * H * eta * gamma * sigma
                - 2 * d ** 2 * eta * rho
                - 4 * d * eta * gamma * rho
                - 2 * d * eta * rho * alpha_m
                - 2 * eta * gamma ** 2 * rho
                - 2 * eta * gamma * rho * alpha_m
                + lam ** 2 * alpha_m ** 2
                + 2 * d * lam * alpha_m
                + 2 * gamma * lam * alpha_m
                + d ** 2
                + 2 * d * gamma
                + gamma ** 2
            )
        )
        * S
        * beta_m
        * (H * lam * sigma + d * lam + gamma * lam + 2 * lam * alpha_m + d + gamma)
        / 2
        + 0.1e1
        / (
            H * lam * sigma * alpha_m
            + H * d * sigma
            + H * gamma * sigma
            + d * lam * alpha_m
            + gamma * lam * alpha_m
            + lam * alpha_m ** 2
            + d ** 2
            + 2 * d * gamma
            + d * alpha_m
            + gamma ** 2
            + gamma * alpha_m
        )
        * (
            rho * eta
            + lam
            + (
                H ** 2 * eta ** 2 * sigma ** 2
                + 2 * H * d * eta ** 2 * rho * sigma
                + 2 * H * eta ** 2 * gamma * rho * sigma
                + 2 * H * eta ** 2 * rho * sigma * alpha_m
                - 4 * H * eta * lam * rho * sigma * alpha_m
                + d ** 2 * eta ** 2 * rho ** 2
                + 2 * d * eta ** 2 * gamma * rho ** 2
                + 2 * d * eta ** 2 * rho ** 2 * alpha_m
                + eta ** 2 * gamma ** 2 * rho ** 2
                + 2 * eta ** 2 * gamma * rho ** 2 * alpha_m
                + eta ** 2 * rho ** 2 * alpha_m ** 2
                - 4 * H * d * eta * rho * sigma
                - 4 * H * eta * gamma * rho * sigma
                + 2 * H * eta * lam * sigma * alpha_m
                - 2 * d * eta * lam * rho * alpha_m
                - 2 * eta * gamma * lam * rho * alpha_m
                - 2 * eta * lam * rho * alpha_m ** 2
                + 2 * H * d * eta * sigma
                + 2 * H * eta * gamma * sigma
                - 2 * d ** 2 * eta * rho
                - 4 * d * eta * gamma * rho
                - 2 * d * eta * rho * alpha_m
                - 2 * eta * gamma ** 2 * rho
                - 2 * eta * gamma * rho * alpha_m
                + lam ** 2 * alpha_m ** 2
                + 2 * d * lam * alpha_m
                + 2 * gamma * lam * alpha_m
                + d ** 2
                + 2 * d * gamma
                + gamma ** 2
            )
            ** (-0.1e1 / 0.2e1)
            * (
                2 * H * eta ** 2 * rho * sigma
                - 4 * H * eta * lam * rho * sigma
                + 2 * d * eta ** 2 * rho ** 2
                + 2 * eta ** 2 * gamma * rho ** 2
                + 2 * eta ** 2 * rho ** 2 * alpha_m
                + 2 * H * eta * lam * sigma
                - 2 * d * eta * lam * rho
                - 2 * eta * gamma * lam * rho
                - 4 * eta * lam * rho * alpha_m
                - 2 * d * eta * rho
                - 2 * eta * gamma * rho
                + 2 * lam ** 2 * alpha_m
                + 2 * d * lam
                + 2 * gamma * lam
            )
            / 2
        )
        * S
        * beta_m
        / 2
        + 0.1e1
        / (
            H * lam * sigma * alpha_m
            + H * d * sigma
            + H * gamma * sigma
            + d * lam * alpha_m
            + gamma * lam * alpha_m
            + lam * alpha_m ** 2
            + d ** 2
            + 2 * d * gamma
            + d * alpha_m
            + gamma ** 2
            + gamma * alpha_m
        )
        * (
            eta * sigma * H
            + d * eta * rho
            + eta * gamma * rho
            + eta * rho * alpha_m
            + lam * alpha_m
            + d
            + gamma
            + sqrt(
                H ** 2 * eta ** 2 * sigma ** 2
                + 2 * H * d * eta ** 2 * rho * sigma
                + 2 * H * eta ** 2 * gamma * rho * sigma
                + 2 * H * eta ** 2 * rho * sigma * alpha_m
                - 4 * H * eta * lam * rho * sigma * alpha_m
                + d ** 2 * eta ** 2 * rho ** 2
                + 2 * d * eta ** 2 * gamma * rho ** 2
                + 2 * d * eta ** 2 * rho ** 2 * alpha_m
                + eta ** 2 * gamma ** 2 * rho ** 2
                + 2 * eta ** 2 * gamma * rho ** 2 * alpha_m
                + eta ** 2 * rho ** 2 * alpha_m ** 2
                - 4 * H * d * eta * rho * sigma
                - 4 * H * eta * gamma * rho * sigma
                + 2 * H * eta * lam * sigma * alpha_m
                - 2 * d * eta * lam * rho * alpha_m
                - 2 * eta * gamma * lam * rho * alpha_m
                - 2 * eta * lam * rho * alpha_m ** 2
                + 2 * H * d * eta * sigma
                + 2 * H * eta * gamma * sigma
                - 2 * d ** 2 * eta * rho
                - 4 * d * eta * gamma * rho
                - 2 * d * eta * rho * alpha_m
                - 2 * eta * gamma ** 2 * rho
                - 2 * eta * gamma * rho * alpha_m
                + lam ** 2 * alpha_m ** 2
                + 2 * d * lam * alpha_m
                + 2 * gamma * lam * alpha_m
                + d ** 2
                + 2 * d * gamma
                + gamma ** 2
            )
        )
        * S
        * dbetadalpha_m
        / 2
    )

    return fit_grad_no_sub


def calculate_alpha_E(
    beta, alpha, sigma, d_beta_d_alpha, d_beta_d_alpha_2, H, S, eta, lam, g, d, rho
):

    E = (
        (
            1
            / (
                lam * sigma * alpha * H
                + alpha * d * lam
                + d * sigma * H
                + alpha * g * lam
                + g * sigma * H
                + alpha ** 2 * lam
                + d ** 2
                + 2 * d * g
                + d * alpha
                + g ** 2
                + alpha * g
            )
            ** 3
            * (
                eta * sigma * H
                + d * eta * rho
                + eta * g * rho
                + eta * rho * alpha
                + lam * alpha
                + d
                + g
                + sqrt(
                    2 * d * eta ** 2 * g * rho ** 2
                    - 4 * d * eta * g * rho
                    + 2 * H * d * eta * sigma
                    + 2 * H * eta * g * sigma
                    + 2 * d * eta ** 2 * rho ** 2 * alpha
                    + 2 * eta ** 2 * g * rho ** 2 * alpha
                    - 2 * eta * lam * rho * alpha ** 2
                    - 2 * d * eta * rho * alpha
                    - 2 * eta * g * rho * alpha
                    + d ** 2 * eta ** 2 * rho ** 2
                    + eta ** 2 * g ** 2 * rho ** 2
                    - 2 * d ** 2 * eta * rho
                    - 2 * eta * g ** 2 * rho
                    + H ** 2 * eta ** 2 * sigma ** 2
                    - 4 * H * eta * g * rho * sigma
                    + 2 * H * d * eta ** 2 * rho * sigma
                    + 2 * H * eta ** 2 * g * rho * sigma
                    - 4 * H * d * eta * rho * sigma
                    + 2 * alpha * d * lam
                    + 2 * alpha * g * lam
                    + d ** 2
                    + 2 * d * g
                    + g ** 2
                    + 2 * H * eta * lam * sigma * alpha
                    - 2 * d * eta * lam * rho * alpha
                    - 2 * eta * g * lam * rho * alpha
                    + 2 * H * eta ** 2 * rho * sigma * alpha
                    - 4 * H * eta * lam * rho * sigma * alpha
                    + lam ** 2 * alpha ** 2
                    + eta ** 2 * rho ** 2 * alpha ** 2
                )
            )
            * S
            * beta
            * (lam * sigma * H + d * lam + g * lam + 2 * lam * alpha + d + g) ** 2
        )
        - 1
        / (
            lam * sigma * alpha * H
            + alpha * d * lam
            + d * sigma * H
            + alpha * g * lam
            + g * sigma * H
            + alpha ** 2 * lam
            + d ** 2
            + 2 * d * g
            + d * alpha
            + g ** 2
            + alpha * g
        )
        ** 2
        * (
            rho * eta
            + lam
            + (
                2 * d * eta ** 2 * g * rho ** 2
                - 4 * d * eta * g * rho
                + 2 * H * d * eta * sigma
                + 2 * H * eta * g * sigma
                + 2 * d * eta ** 2 * rho ** 2 * alpha
                + 2 * eta ** 2 * g * rho ** 2 * alpha
                - 2 * eta * lam * rho * alpha ** 2
                - 2 * d * eta * rho * alpha
                - 2 * eta * g * rho * alpha
                + d ** 2 * eta ** 2 * rho ** 2
                + eta ** 2 * g ** 2 * rho ** 2
                - 2 * d ** 2 * eta * rho
                - 2 * eta * g ** 2 * rho
                + H ** 2 * eta ** 2 * sigma ** 2
                - 4 * H * eta * g * rho * sigma
                + 2 * H * d * eta ** 2 * rho * sigma
                + 2 * H * eta ** 2 * g * rho * sigma
                - 4 * H * d * eta * rho * sigma
                + 2 * alpha * d * lam
                + 2 * alpha * g * lam
                + d ** 2
                + 2 * d * g
                + g ** 2
                + 2 * H * eta * lam * sigma * alpha
                - 2 * d * eta * lam * rho * alpha
                - 2 * eta * g * lam * rho * alpha
                + 2 * H * eta ** 2 * rho * sigma * alpha
                - 4 * H * eta * lam * rho * sigma * alpha
                + lam ** 2 * alpha ** 2
                + eta ** 2 * rho ** 2 * alpha ** 2
            )
            ** (-0.1e1 / 0.2e1)
            * (
                2 * d * eta ** 2 * rho ** 2
                + 2 * eta ** 2 * g * rho ** 2
                - 4 * eta * lam * rho * alpha
                - 2 * d * eta * rho
                - 2 * eta * g * rho
                + 2 * d * lam
                + 2 * g * lam
                + 2 * lam ** 2 * alpha
                + 2 * eta ** 2 * rho ** 2 * alpha
                + 2 * H * eta * lam * sigma
                - 2 * d * eta * lam * rho
                - 2 * eta * g * lam * rho
                + 2 * H * eta ** 2 * rho * sigma
                - 4 * H * eta * lam * rho * sigma
            )
            / 2
        )
        * S
        * beta
        * (lam * sigma * H + d * lam + g * lam + 2 * lam * alpha + d + g)
        - 1
        / (
            lam * sigma * alpha * H
            + alpha * d * lam
            + d * sigma * H
            + alpha * g * lam
            + g * sigma * H
            + alpha ** 2 * lam
            + d ** 2
            + 2 * d * g
            + d * alpha
            + g ** 2
            + alpha * g
        )
        ** 2
        * (
            eta * sigma * H
            + d * eta * rho
            + eta * g * rho
            + eta * rho * alpha
            + lam * alpha
            + d
            + g
            + sqrt(
                2 * d * eta ** 2 * g * rho ** 2
                - 4 * d * eta * g * rho
                + 2 * H * d * eta * sigma
                + 2 * H * eta * g * sigma
                + 2 * d * eta ** 2 * rho ** 2 * alpha
                + 2 * eta ** 2 * g * rho ** 2 * alpha
                - 2 * eta * lam * rho * alpha ** 2
                - 2 * d * eta * rho * alpha
                - 2 * eta * g * rho * alpha
                + d ** 2 * eta ** 2 * rho ** 2
                + eta ** 2 * g ** 2 * rho ** 2
                - 2 * d ** 2 * eta * rho
                - 2 * eta * g ** 2 * rho
                + H ** 2 * eta ** 2 * sigma ** 2
                - 4 * H * eta * g * rho * sigma
                + 2 * H * d * eta ** 2 * rho * sigma
                + 2 * H * eta ** 2 * g * rho * sigma
                - 4 * H * d * eta * rho * sigma
                + 2 * alpha * d * lam
                + 2 * alpha * g * lam
                + d ** 2
                + 2 * d * g
                + g ** 2
                + 2 * H * eta * lam * sigma * alpha
                - 2 * d * eta * lam * rho * alpha
                - 2 * eta * g * lam * rho * alpha
                + 2 * H * eta ** 2 * rho * sigma * alpha
                - 4 * H * eta * lam * rho * sigma * alpha
                + lam ** 2 * alpha ** 2
                + eta ** 2 * rho ** 2 * alpha ** 2
            )
        )
        * S
        * d_beta_d_alpha
        * (lam * sigma * H + d * lam + g * lam + 2 * lam * alpha + d + g)
        - 1
        / (
            lam * sigma * alpha * H
            + alpha * d * lam
            + d * sigma * H
            + alpha * g * lam
            + g * sigma * H
            + alpha ** 2 * lam
            + d ** 2
            + 2 * d * g
            + d * alpha
            + g ** 2
            + alpha * g
        )
        ** 2
        * (
            eta * sigma * H
            + d * eta * rho
            + eta * g * rho
            + eta * rho * alpha
            + lam * alpha
            + d
            + g
            + sqrt(
                2 * d * eta ** 2 * g * rho ** 2
                - 4 * d * eta * g * rho
                + 2 * H * d * eta * sigma
                + 2 * H * eta * g * sigma
                + 2 * d * eta ** 2 * rho ** 2 * alpha
                + 2 * eta ** 2 * g * rho ** 2 * alpha
                - 2 * eta * lam * rho * alpha ** 2
                - 2 * d * eta * rho * alpha
                - 2 * eta * g * rho * alpha
                + d ** 2 * eta ** 2 * rho ** 2
                + eta ** 2 * g ** 2 * rho ** 2
                - 2 * d ** 2 * eta * rho
                - 2 * eta * g ** 2 * rho
                + H ** 2 * eta ** 2 * sigma ** 2
                - 4 * H * eta * g * rho * sigma
                + 2 * H * d * eta ** 2 * rho * sigma
                + 2 * H * eta ** 2 * g * rho * sigma
                - 4 * H * d * eta * rho * sigma
                + 2 * alpha * d * lam
                + 2 * alpha * g * lam
                + d ** 2
                + 2 * d * g
                + g ** 2
                + 2 * H * eta * lam * sigma * alpha
                - 2 * d * eta * lam * rho * alpha
                - 2 * eta * g * lam * rho * alpha
                + 2 * H * eta ** 2 * rho * sigma * alpha
                - 4 * H * eta * lam * rho * sigma * alpha
                + lam ** 2 * alpha ** 2
                + eta ** 2 * rho ** 2 * alpha ** 2
            )
        )
        * S
        * beta
        * lam
        + 1
        / (
            lam * sigma * alpha * H
            + alpha * d * lam
            + d * sigma * H
            + alpha * g * lam
            + g * sigma * H
            + alpha ** 2 * lam
            + d ** 2
            + 2 * d * g
            + d * alpha
            + g ** 2
            + alpha * g
        )
        * (
            -(
                (
                    2 * d * eta ** 2 * g * rho ** 2
                    - 4 * d * eta * g * rho
                    + 2 * H * d * eta * sigma
                    + 2 * H * eta * g * sigma
                    + 2 * d * eta ** 2 * rho ** 2 * alpha
                    + 2 * eta ** 2 * g * rho ** 2 * alpha
                    - 2 * eta * lam * rho * alpha ** 2
                    - 2 * d * eta * rho * alpha
                    - 2 * eta * g * rho * alpha
                    + d ** 2 * eta ** 2 * rho ** 2
                    + eta ** 2 * g ** 2 * rho ** 2
                    - 2 * d ** 2 * eta * rho
                    - 2 * eta * g ** 2 * rho
                    + H ** 2 * eta ** 2 * sigma ** 2
                    - 4 * H * eta * g * rho * sigma
                    + 2 * H * d * eta ** 2 * rho * sigma
                    + 2 * H * eta ** 2 * g * rho * sigma
                    - 4 * H * d * eta * rho * sigma
                    + 2 * alpha * d * lam
                    + 2 * alpha * g * lam
                    + d ** 2
                    + 2 * d * g
                    + g ** 2
                    + 2 * H * eta * lam * sigma * alpha
                    - 2 * d * eta * lam * rho * alpha
                    - 2 * eta * g * lam * rho * alpha
                    + 2 * H * eta ** 2 * rho * sigma * alpha
                    - 4 * H * eta * lam * rho * sigma * alpha
                    + lam ** 2 * alpha ** 2
                    + eta ** 2 * rho ** 2 * alpha ** 2
                )
                ** (-0.3e1 / 0.2e1)
            )
            * (
                2 * d * eta ** 2 * rho ** 2
                + 2 * eta ** 2 * g * rho ** 2
                - 4 * eta * lam * rho * alpha
                - 2 * d * eta * rho
                - 2 * eta * g * rho
                + 2 * d * lam
                + 2 * g * lam
                + 2 * lam ** 2 * alpha
                + 2 * eta ** 2 * rho ** 2 * alpha
                + 2 * H * eta * lam * sigma
                - 2 * d * eta * lam * rho
                - 2 * eta * g * lam * rho
                + 2 * H * eta ** 2 * rho * sigma
                - 4 * H * eta * lam * rho * sigma
            )
            ** 2
            / 4
            + (
                2 * d * eta ** 2 * g * rho ** 2
                - 4 * d * eta * g * rho
                + 2 * H * d * eta * sigma
                + 2 * H * eta * g * sigma
                + 2 * d * eta ** 2 * rho ** 2 * alpha
                + 2 * eta ** 2 * g * rho ** 2 * alpha
                - 2 * eta * lam * rho * alpha ** 2
                - 2 * d * eta * rho * alpha
                - 2 * eta * g * rho * alpha
                + d ** 2 * eta ** 2 * rho ** 2
                + eta ** 2 * g ** 2 * rho ** 2
                - 2 * d ** 2 * eta * rho
                - 2 * eta * g ** 2 * rho
                + H ** 2 * eta ** 2 * sigma ** 2
                - 4 * H * eta * g * rho * sigma
                + 2 * H * d * eta ** 2 * rho * sigma
                + 2 * H * eta ** 2 * g * rho * sigma
                - 4 * H * d * eta * rho * sigma
                + 2 * alpha * d * lam
                + 2 * alpha * g * lam
                + d ** 2
                + 2 * d * g
                + g ** 2
                + 2 * H * eta * lam * sigma * alpha
                - 2 * d * eta * lam * rho * alpha
                - 2 * eta * g * lam * rho * alpha
                + 2 * H * eta ** 2 * rho * sigma * alpha
                - 4 * H * eta * lam * rho * sigma * alpha
                + lam ** 2 * alpha ** 2
                + eta ** 2 * rho ** 2 * alpha ** 2
            )
            ** (-0.1e1 / 0.2e1)
            * (2 * eta ** 2 * rho ** 2 - 4 * eta * lam * rho + 2 * lam ** 2)
            / 2
        )
        * S
        * beta
        / 2
        + 1
        / (
            lam * sigma * alpha * H
            + alpha * d * lam
            + d * sigma * H
            + alpha * g * lam
            + g * sigma * H
            + alpha ** 2 * lam
            + d ** 2
            + 2 * d * g
            + d * alpha
            + g ** 2
            + alpha * g
        )
        * (
            rho * eta
            + lam
            + (
                2 * d * eta ** 2 * g * rho ** 2
                - 4 * d * eta * g * rho
                + 2 * H * d * eta * sigma
                + 2 * H * eta * g * sigma
                + 2 * d * eta ** 2 * rho ** 2 * alpha
                + 2 * eta ** 2 * g * rho ** 2 * alpha
                - 2 * eta * lam * rho * alpha ** 2
                - 2 * d * eta * rho * alpha
                - 2 * eta * g * rho * alpha
                + d ** 2 * eta ** 2 * rho ** 2
                + eta ** 2 * g ** 2 * rho ** 2
                - 2 * d ** 2 * eta * rho
                - 2 * eta * g ** 2 * rho
                + H ** 2 * eta ** 2 * sigma ** 2
                - 4 * H * eta * g * rho * sigma
                + 2 * H * d * eta ** 2 * rho * sigma
                + 2 * H * eta ** 2 * g * rho * sigma
                - 4 * H * d * eta * rho * sigma
                + 2 * alpha * d * lam
                + 2 * alpha * g * lam
                + d ** 2
                + 2 * d * g
                + g ** 2
                + 2 * H * eta * lam * sigma * alpha
                - 2 * d * eta * lam * rho * alpha
                - 2 * eta * g * lam * rho * alpha
                + 2 * H * eta ** 2 * rho * sigma * alpha
                - 4 * H * eta * lam * rho * sigma * alpha
                + lam ** 2 * alpha ** 2
                + eta ** 2 * rho ** 2 * alpha ** 2
            )
            ** (-0.1e1 / 0.2e1)
            * (
                2 * d * eta ** 2 * rho ** 2
                + 2 * eta ** 2 * g * rho ** 2
                - 4 * eta * lam * rho * alpha
                - 2 * d * eta * rho
                - 2 * eta * g * rho
                + 2 * d * lam
                + 2 * g * lam
                + 2 * lam ** 2 * alpha
                + 2 * eta ** 2 * rho ** 2 * alpha
                + 2 * H * eta * lam * sigma
                - 2 * d * eta * lam * rho
                - 2 * eta * g * lam * rho
                + 2 * H * eta ** 2 * rho * sigma
                - 4 * H * eta * lam * rho * sigma
            )
            / 2
        )
        * S
        * d_beta_d_alpha
        + 1
        / (
            lam * sigma * alpha * H
            + alpha * d * lam
            + d * sigma * H
            + alpha * g * lam
            + g * sigma * H
            + alpha ** 2 * lam
            + d ** 2
            + 2 * d * g
            + d * alpha
            + g ** 2
            + alpha * g
        )
        * (
            eta * sigma * H
            + d * eta * rho
            + eta * g * rho
            + eta * rho * alpha
            + lam * alpha
            + d
            + g
            + sqrt(
                2 * d * eta ** 2 * g * rho ** 2
                - 4 * d * eta * g * rho
                + 2 * H * d * eta * sigma
                + 2 * H * eta * g * sigma
                + 2 * d * eta ** 2 * rho ** 2 * alpha
                + 2 * eta ** 2 * g * rho ** 2 * alpha
                - 2 * eta * lam * rho * alpha ** 2
                - 2 * d * eta * rho * alpha
                - 2 * eta * g * rho * alpha
                + d ** 2 * eta ** 2 * rho ** 2
                + eta ** 2 * g ** 2 * rho ** 2
                - 2 * d ** 2 * eta * rho
                - 2 * eta * g ** 2 * rho
                + H ** 2 * eta ** 2 * sigma ** 2
                - 4 * H * eta * g * rho * sigma
                + 2 * H * d * eta ** 2 * rho * sigma
                + 2 * H * eta ** 2 * g * rho * sigma
                - 4 * H * d * eta * rho * sigma
                + 2 * alpha * d * lam
                + 2 * alpha * g * lam
                + d ** 2
                + 2 * d * g
                + g ** 2
                + 2 * H * eta * lam * sigma * alpha
                - 2 * d * eta * lam * rho * alpha
                - 2 * eta * g * lam * rho * alpha
                + 2 * H * eta ** 2 * rho * sigma * alpha
                - 4 * H * eta * lam * rho * sigma * alpha
                + lam ** 2 * alpha ** 2
                + eta ** 2 * rho ** 2 * alpha ** 2
            )
        )
        * S
        * d_beta_d_alpha_2
        / 2
    )

    return E


def calculate_alpha_M(
    beta,
    alpha,
    sigma,
    d_beta_d_alpha,
    H,
    S,
    d_H_d_alpha,
    d_S_d_alpha,
    eta,
    lam,
    g,
    d,
    rho,
):

    M = (
        1
        / (
            lam * sigma * alpha * H
            + alpha * d * lam
            + d * sigma * H
            + alpha * g * lam
            + g * sigma * H
            + alpha ** 2 * lam
            + d ** 2
            + 2 * d * g
            + d * alpha
            + g ** 2
            + alpha * g
        )
        ** 3
        * (
            eta * sigma * H
            + d * eta * rho
            + eta * g * rho
            + eta * rho * alpha
            + lam * alpha
            + d
            + g
            + sqrt(
                2 * d * eta ** 2 * g * rho ** 2
                - 4 * d * eta * g * rho
                + 2 * H * d * eta * sigma
                + 2 * H * eta * g * sigma
                - 4 * H * eta * lam * rho * sigma * alpha
                + lam ** 2 * alpha ** 2
                + d ** 2 * eta ** 2 * rho ** 2
                + eta ** 2 * g ** 2 * rho ** 2
                - 2 * d ** 2 * eta * rho
                - 2 * eta * g ** 2 * rho
                + H ** 2 * eta ** 2 * sigma ** 2
                + 2 * H * d * eta ** 2 * rho * sigma
                + 2 * H * eta ** 2 * g * rho * sigma
                - 4 * H * d * eta * rho * sigma
                - 4 * H * eta * g * rho * sigma
                - 2 * d * eta * lam * rho * alpha
                - 2 * eta * g * lam * rho * alpha
                + 2 * H * eta ** 2 * rho * sigma * alpha
                + 2 * H * eta * lam * sigma * alpha
                + 2 * alpha * d * lam
                + 2 * alpha * g * lam
                + d ** 2
                + 2 * d * g
                + g ** 2
                + eta ** 2 * rho ** 2 * alpha ** 2
                + 2 * d * eta ** 2 * rho ** 2 * alpha
                + 2 * eta ** 2 * g * rho ** 2 * alpha
                - 2 * eta * lam * rho * alpha ** 2
                - 2 * d * eta * rho * alpha
                - 2 * eta * g * rho * alpha
            )
        )
        * S
        * beta
        * (lam * sigma * H + d * lam + g * lam + 2 * lam * alpha + d + g)
        * (
            lam * sigma * alpha * d_H_d_alpha
            + d * sigma * d_H_d_alpha
            + g * sigma * d_H_d_alpha
        )
        - 1
        / (
            lam * sigma * alpha * H
            + alpha * d * lam
            + d * sigma * H
            + alpha * g * lam
            + g * sigma * H
            + alpha ** 2 * lam
            + d ** 2
            + 2 * d * g
            + d * alpha
            + g ** 2
            + alpha * g
        )
        ** 2
        * (
            eta * sigma * d_H_d_alpha
            + (
                2 * d * eta ** 2 * g * rho ** 2
                - 4 * d * eta * g * rho
                + 2 * H * d * eta * sigma
                + 2 * H * eta * g * sigma
                - 4 * H * eta * lam * rho * sigma * alpha
                + lam ** 2 * alpha ** 2
                + d ** 2 * eta ** 2 * rho ** 2
                + eta ** 2 * g ** 2 * rho ** 2
                - 2 * d ** 2 * eta * rho
                - 2 * eta * g ** 2 * rho
                + H ** 2 * eta ** 2 * sigma ** 2
                + 2 * H * d * eta ** 2 * rho * sigma
                + 2 * H * eta ** 2 * g * rho * sigma
                - 4 * H * d * eta * rho * sigma
                - 4 * H * eta * g * rho * sigma
                - 2 * d * eta * lam * rho * alpha
                - 2 * eta * g * lam * rho * alpha
                + 2 * H * eta ** 2 * rho * sigma * alpha
                + 2 * H * eta * lam * sigma * alpha
                + 2 * alpha * d * lam
                + 2 * alpha * g * lam
                + d ** 2
                + 2 * d * g
                + g ** 2
                + eta ** 2 * rho ** 2 * alpha ** 2
                + 2 * d * eta ** 2 * rho ** 2 * alpha
                + 2 * eta ** 2 * g * rho ** 2 * alpha
                - 2 * eta * lam * rho * alpha ** 2
                - 2 * d * eta * rho * alpha
                - 2 * eta * g * rho * alpha
            )
            ** (-0.1e1 / 0.2e1)
            * (
                2 * d_H_d_alpha * d * eta * sigma
                + 2 * d_H_d_alpha * eta * g * sigma
                - 4 * d_H_d_alpha * eta * lam * rho * sigma * alpha
                + 2 * H * eta ** 2 * sigma ** 2 * d_H_d_alpha
                + 2 * d_H_d_alpha * d * eta ** 2 * rho * sigma
                + 2 * d_H_d_alpha * eta ** 2 * g * rho * sigma
                + 2 * d_H_d_alpha * eta ** 2 * rho * sigma * alpha
                - 4 * d_H_d_alpha * d * eta * rho * sigma
                - 4 * d_H_d_alpha * eta * g * rho * sigma
                + 2 * d_H_d_alpha * eta * lam * sigma * alpha
            )
            / 2
        )
        * S
        * beta
        * (lam * sigma * H + d * lam + g * lam + 2 * lam * alpha + d + g)
        / 2
        - 1
        / (
            lam * sigma * alpha * H
            + alpha * d * lam
            + d * sigma * H
            + alpha * g * lam
            + g * sigma * H
            + alpha ** 2 * lam
            + d ** 2
            + 2 * d * g
            + d * alpha
            + g ** 2
            + alpha * g
        )
        ** 2
        * (
            eta * sigma * H
            + d * eta * rho
            + eta * g * rho
            + eta * rho * alpha
            + lam * alpha
            + d
            + g
            + sqrt(
                2 * d * eta ** 2 * g * rho ** 2
                - 4 * d * eta * g * rho
                + 2 * H * d * eta * sigma
                + 2 * H * eta * g * sigma
                - 4 * H * eta * lam * rho * sigma * alpha
                + lam ** 2 * alpha ** 2
                + d ** 2 * eta ** 2 * rho ** 2
                + eta ** 2 * g ** 2 * rho ** 2
                - 2 * d ** 2 * eta * rho
                - 2 * eta * g ** 2 * rho
                + H ** 2 * eta ** 2 * sigma ** 2
                + 2 * H * d * eta ** 2 * rho * sigma
                + 2 * H * eta ** 2 * g * rho * sigma
                - 4 * H * d * eta * rho * sigma
                - 4 * H * eta * g * rho * sigma
                - 2 * d * eta * lam * rho * alpha
                - 2 * eta * g * lam * rho * alpha
                + 2 * H * eta ** 2 * rho * sigma * alpha
                + 2 * H * eta * lam * sigma * alpha
                + 2 * alpha * d * lam
                + 2 * alpha * g * lam
                + d ** 2
                + 2 * d * g
                + g ** 2
                + eta ** 2 * rho ** 2 * alpha ** 2
                + 2 * d * eta ** 2 * rho ** 2 * alpha
                + 2 * eta ** 2 * g * rho ** 2 * alpha
                - 2 * eta * lam * rho * alpha ** 2
                - 2 * d * eta * rho * alpha
                - 2 * eta * g * rho * alpha
            )
        )
        * d_S_d_alpha
        * beta
        * (lam * sigma * H + d * lam + g * lam + 2 * lam * alpha + d + g)
        / 2
        - 1
        / (
            lam * sigma * alpha * H
            + alpha * d * lam
            + d * sigma * H
            + alpha * g * lam
            + g * sigma * H
            + alpha ** 2 * lam
            + d ** 2
            + 2 * d * g
            + d * alpha
            + g ** 2
            + alpha * g
        )
        ** 2
        * (
            eta * sigma * H
            + d * eta * rho
            + eta * g * rho
            + eta * rho * alpha
            + lam * alpha
            + d
            + g
            + sqrt(
                2 * d * eta ** 2 * g * rho ** 2
                - 4 * d * eta * g * rho
                + 2 * H * d * eta * sigma
                + 2 * H * eta * g * sigma
                - 4 * H * eta * lam * rho * sigma * alpha
                + lam ** 2 * alpha ** 2
                + d ** 2 * eta ** 2 * rho ** 2
                + eta ** 2 * g ** 2 * rho ** 2
                - 2 * d ** 2 * eta * rho
                - 2 * eta * g ** 2 * rho
                + H ** 2 * eta ** 2 * sigma ** 2
                + 2 * H * d * eta ** 2 * rho * sigma
                + 2 * H * eta ** 2 * g * rho * sigma
                - 4 * H * d * eta * rho * sigma
                - 4 * H * eta * g * rho * sigma
                - 2 * d * eta * lam * rho * alpha
                - 2 * eta * g * lam * rho * alpha
                + 2 * H * eta ** 2 * rho * sigma * alpha
                + 2 * H * eta * lam * sigma * alpha
                + 2 * alpha * d * lam
                + 2 * alpha * g * lam
                + d ** 2
                + 2 * d * g
                + g ** 2
                + eta ** 2 * rho ** 2 * alpha ** 2
                + 2 * d * eta ** 2 * rho ** 2 * alpha
                + 2 * eta ** 2 * g * rho ** 2 * alpha
                - 2 * eta * lam * rho * alpha ** 2
                - 2 * d * eta * rho * alpha
                - 2 * eta * g * rho * alpha
            )
        )
        * S
        * beta
        * lam
        * sigma
        * d_H_d_alpha
        / 2
        - 1
        / (
            lam * sigma * alpha * H
            + alpha * d * lam
            + d * sigma * H
            + alpha * g * lam
            + g * sigma * H
            + alpha ** 2 * lam
            + d ** 2
            + 2 * d * g
            + d * alpha
            + g ** 2
            + alpha * g
        )
        ** 2
        * (
            rho * eta
            + lam
            + (
                2 * d * eta ** 2 * g * rho ** 2
                - 4 * d * eta * g * rho
                + 2 * H * d * eta * sigma
                + 2 * H * eta * g * sigma
                - 4 * H * eta * lam * rho * sigma * alpha
                + lam ** 2 * alpha ** 2
                + d ** 2 * eta ** 2 * rho ** 2
                + eta ** 2 * g ** 2 * rho ** 2
                - 2 * d ** 2 * eta * rho
                - 2 * eta * g ** 2 * rho
                + H ** 2 * eta ** 2 * sigma ** 2
                + 2 * H * d * eta ** 2 * rho * sigma
                + 2 * H * eta ** 2 * g * rho * sigma
                - 4 * H * d * eta * rho * sigma
                - 4 * H * eta * g * rho * sigma
                - 2 * d * eta * lam * rho * alpha
                - 2 * eta * g * lam * rho * alpha
                + 2 * H * eta ** 2 * rho * sigma * alpha
                + 2 * H * eta * lam * sigma * alpha
                + 2 * alpha * d * lam
                + 2 * alpha * g * lam
                + d ** 2
                + 2 * d * g
                + g ** 2
                + eta ** 2 * rho ** 2 * alpha ** 2
                + 2 * d * eta ** 2 * rho ** 2 * alpha
                + 2 * eta ** 2 * g * rho ** 2 * alpha
                - 2 * eta * lam * rho * alpha ** 2
                - 2 * d * eta * rho * alpha
                - 2 * eta * g * rho * alpha
            )
            ** (-0.1e1 / 0.2e1)
            * (
                2 * d * eta ** 2 * rho ** 2
                + 2 * g * rho ** 2 * eta ** 2
                - 4 * eta * lam * rho * alpha
                - 2 * d * eta * rho
                - 2 * eta * g * rho
                - 4 * H * eta * lam * rho * sigma
                + 2 * d * lam
                + 2 * g * lam
                + 2 * rho ** 2 * alpha * eta ** 2
                + 2 * lam ** 2 * alpha
                - 2 * d * eta * lam * rho
                - 2 * eta * g * lam * rho
                + 2 * H * eta ** 2 * rho * sigma
                + 2 * H * eta * lam * sigma
            )
            / 2
        )
        * S
        * beta
        * (
            lam * sigma * alpha * d_H_d_alpha
            + d * sigma * d_H_d_alpha
            + g * sigma * d_H_d_alpha
        )
        / 2
        + 1
        / (
            lam * sigma * alpha * H
            + alpha * d * lam
            + d * sigma * H
            + alpha * g * lam
            + g * sigma * H
            + alpha ** 2 * lam
            + d ** 2
            + 2 * d * g
            + d * alpha
            + g ** 2
            + alpha * g
        )
        * (
            -(
                (
                    2 * d * eta ** 2 * g * rho ** 2
                    - 4 * d * eta * g * rho
                    + 2 * H * d * eta * sigma
                    + 2 * H * eta * g * sigma
                    - 4 * H * eta * lam * rho * sigma * alpha
                    + lam ** 2 * alpha ** 2
                    + d ** 2 * eta ** 2 * rho ** 2
                    + eta ** 2 * g ** 2 * rho ** 2
                    - 2 * d ** 2 * eta * rho
                    - 2 * eta * g ** 2 * rho
                    + H ** 2 * eta ** 2 * sigma ** 2
                    + 2 * H * d * eta ** 2 * rho * sigma
                    + 2 * H * eta ** 2 * g * rho * sigma
                    - 4 * H * d * eta * rho * sigma
                    - 4 * H * eta * g * rho * sigma
                    - 2 * d * eta * lam * rho * alpha
                    - 2 * eta * g * lam * rho * alpha
                    + 2 * H * eta ** 2 * rho * sigma * alpha
                    + 2 * H * eta * lam * sigma * alpha
                    + 2 * alpha * d * lam
                    + 2 * alpha * g * lam
                    + d ** 2
                    + 2 * d * g
                    + g ** 2
                    + eta ** 2 * rho ** 2 * alpha ** 2
                    + 2 * d * eta ** 2 * rho ** 2 * alpha
                    + 2 * eta ** 2 * g * rho ** 2 * alpha
                    - 2 * eta * lam * rho * alpha ** 2
                    - 2 * d * eta * rho * alpha
                    - 2 * eta * g * rho * alpha
                )
                ** (-0.3e1 / 0.2e1)
            )
            * (
                2 * d * eta ** 2 * rho ** 2
                + 2 * g * rho ** 2 * eta ** 2
                - 4 * eta * lam * rho * alpha
                - 2 * d * eta * rho
                - 2 * eta * g * rho
                - 4 * H * eta * lam * rho * sigma
                + 2 * d * lam
                + 2 * g * lam
                + 2 * rho ** 2 * alpha * eta ** 2
                + 2 * lam ** 2 * alpha
                - 2 * d * eta * lam * rho
                - 2 * eta * g * lam * rho
                + 2 * H * eta ** 2 * rho * sigma
                + 2 * H * eta * lam * sigma
            )
            * (
                2 * d_H_d_alpha * d * eta * sigma
                + 2 * d_H_d_alpha * eta * g * sigma
                - 4 * d_H_d_alpha * eta * lam * rho * sigma * alpha
                + 2 * H * eta ** 2 * sigma ** 2 * d_H_d_alpha
                + 2 * d_H_d_alpha * d * eta ** 2 * rho * sigma
                + 2 * d_H_d_alpha * eta ** 2 * g * rho * sigma
                + 2 * d_H_d_alpha * eta ** 2 * rho * sigma * alpha
                - 4 * d_H_d_alpha * d * eta * rho * sigma
                - 4 * d_H_d_alpha * eta * g * rho * sigma
                + 2 * d_H_d_alpha * eta * lam * sigma * alpha
            )
            / 4
            + (
                2 * d * eta ** 2 * g * rho ** 2
                - 4 * d * eta * g * rho
                + 2 * H * d * eta * sigma
                + 2 * H * eta * g * sigma
                - 4 * H * eta * lam * rho * sigma * alpha
                + lam ** 2 * alpha ** 2
                + d ** 2 * eta ** 2 * rho ** 2
                + eta ** 2 * g ** 2 * rho ** 2
                - 2 * d ** 2 * eta * rho
                - 2 * eta * g ** 2 * rho
                + H ** 2 * eta ** 2 * sigma ** 2
                + 2 * H * d * eta ** 2 * rho * sigma
                + 2 * H * eta ** 2 * g * rho * sigma
                - 4 * H * d * eta * rho * sigma
                - 4 * H * eta * g * rho * sigma
                - 2 * d * eta * lam * rho * alpha
                - 2 * eta * g * lam * rho * alpha
                + 2 * H * eta ** 2 * rho * sigma * alpha
                + 2 * H * eta * lam * sigma * alpha
                + 2 * alpha * d * lam
                + 2 * alpha * g * lam
                + d ** 2
                + 2 * d * g
                + g ** 2
                + eta ** 2 * rho ** 2 * alpha ** 2
                + 2 * d * eta ** 2 * rho ** 2 * alpha
                + 2 * eta ** 2 * g * rho ** 2 * alpha
                - 2 * eta * lam * rho * alpha ** 2
                - 2 * d * eta * rho * alpha
                - 2 * eta * g * rho * alpha
            )
            ** (-0.1e1 / 0.2e1)
            * (
                -4 * d_H_d_alpha * eta * lam * rho * sigma
                + 2 * d_H_d_alpha * eta ** 2 * rho * sigma
                + 2 * d_H_d_alpha * eta * lam * sigma
            )
            / 2
        )
        * S
        * beta
        / 2
        + 1
        / (
            lam * sigma * alpha * H
            + alpha * d * lam
            + d * sigma * H
            + alpha * g * lam
            + g * sigma * H
            + alpha ** 2 * lam
            + d ** 2
            + 2 * d * g
            + d * alpha
            + g ** 2
            + alpha * g
        )
        * (
            rho * eta
            + lam
            + (
                2 * d * eta ** 2 * g * rho ** 2
                - 4 * d * eta * g * rho
                + 2 * H * d * eta * sigma
                + 2 * H * eta * g * sigma
                - 4 * H * eta * lam * rho * sigma * alpha
                + lam ** 2 * alpha ** 2
                + d ** 2 * eta ** 2 * rho ** 2
                + eta ** 2 * g ** 2 * rho ** 2
                - 2 * d ** 2 * eta * rho
                - 2 * eta * g ** 2 * rho
                + H ** 2 * eta ** 2 * sigma ** 2
                + 2 * H * d * eta ** 2 * rho * sigma
                + 2 * H * eta ** 2 * g * rho * sigma
                - 4 * H * d * eta * rho * sigma
                - 4 * H * eta * g * rho * sigma
                - 2 * d * eta * lam * rho * alpha
                - 2 * eta * g * lam * rho * alpha
                + 2 * H * eta ** 2 * rho * sigma * alpha
                + 2 * H * eta * lam * sigma * alpha
                + 2 * alpha * d * lam
                + 2 * alpha * g * lam
                + d ** 2
                + 2 * d * g
                + g ** 2
                + eta ** 2 * rho ** 2 * alpha ** 2
                + 2 * d * eta ** 2 * rho ** 2 * alpha
                + 2 * eta ** 2 * g * rho ** 2 * alpha
                - 2 * eta * lam * rho * alpha ** 2
                - 2 * d * eta * rho * alpha
                - 2 * eta * g * rho * alpha
            )
            ** (-0.1e1 / 0.2e1)
            * (
                2 * d * eta ** 2 * rho ** 2
                + 2 * g * rho ** 2 * eta ** 2
                - 4 * eta * lam * rho * alpha
                - 2 * d * eta * rho
                - 2 * eta * g * rho
                - 4 * H * eta * lam * rho * sigma
                + 2 * d * lam
                + 2 * g * lam
                + 2 * rho ** 2 * alpha * eta ** 2
                + 2 * lam ** 2 * alpha
                - 2 * d * eta * lam * rho
                - 2 * eta * g * lam * rho
                + 2 * H * eta ** 2 * rho * sigma
                + 2 * H * eta * lam * sigma
            )
            / 2
        )
        * d_S_d_alpha
        * beta
        / 2
        - 1
        / (
            lam * sigma * alpha * H
            + alpha * d * lam
            + d * sigma * H
            + alpha * g * lam
            + g * sigma * H
            + alpha ** 2 * lam
            + d ** 2
            + 2 * d * g
            + d * alpha
            + g ** 2
            + alpha * g
        )
        ** 2
        * (
            eta * sigma * H
            + d * eta * rho
            + eta * g * rho
            + eta * rho * alpha
            + lam * alpha
            + d
            + g
            + sqrt(
                2 * d * eta ** 2 * g * rho ** 2
                - 4 * d * eta * g * rho
                + 2 * H * d * eta * sigma
                + 2 * H * eta * g * sigma
                - 4 * H * eta * lam * rho * sigma * alpha
                + lam ** 2 * alpha ** 2
                + d ** 2 * eta ** 2 * rho ** 2
                + eta ** 2 * g ** 2 * rho ** 2
                - 2 * d ** 2 * eta * rho
                - 2 * eta * g ** 2 * rho
                + H ** 2 * eta ** 2 * sigma ** 2
                + 2 * H * d * eta ** 2 * rho * sigma
                + 2 * H * eta ** 2 * g * rho * sigma
                - 4 * H * d * eta * rho * sigma
                - 4 * H * eta * g * rho * sigma
                - 2 * d * eta * lam * rho * alpha
                - 2 * eta * g * lam * rho * alpha
                + 2 * H * eta ** 2 * rho * sigma * alpha
                + 2 * H * eta * lam * sigma * alpha
                + 2 * alpha * d * lam
                + 2 * alpha * g * lam
                + d ** 2
                + 2 * d * g
                + g ** 2
                + eta ** 2 * rho ** 2 * alpha ** 2
                + 2 * d * eta ** 2 * rho ** 2 * alpha
                + 2 * eta ** 2 * g * rho ** 2 * alpha
                - 2 * eta * lam * rho * alpha ** 2
                - 2 * d * eta * rho * alpha
                - 2 * eta * g * rho * alpha
            )
        )
        * S
        * d_beta_d_alpha
        * (
            lam * sigma * alpha * d_H_d_alpha
            + d * sigma * d_H_d_alpha
            + g * sigma * d_H_d_alpha
        )
        / 2
        + 1
        / (
            lam * sigma * alpha * H
            + alpha * d * lam
            + d * sigma * H
            + alpha * g * lam
            + g * sigma * H
            + alpha ** 2 * lam
            + d ** 2
            + 2 * d * g
            + d * alpha
            + g ** 2
            + alpha * g
        )
        * (
            eta * sigma * d_H_d_alpha
            + (
                2 * d * eta ** 2 * g * rho ** 2
                - 4 * d * eta * g * rho
                + 2 * H * d * eta * sigma
                + 2 * H * eta * g * sigma
                - 4 * H * eta * lam * rho * sigma * alpha
                + lam ** 2 * alpha ** 2
                + d ** 2 * eta ** 2 * rho ** 2
                + eta ** 2 * g ** 2 * rho ** 2
                - 2 * d ** 2 * eta * rho
                - 2 * eta * g ** 2 * rho
                + H ** 2 * eta ** 2 * sigma ** 2
                + 2 * H * d * eta ** 2 * rho * sigma
                + 2 * H * eta ** 2 * g * rho * sigma
                - 4 * H * d * eta * rho * sigma
                - 4 * H * eta * g * rho * sigma
                - 2 * d * eta * lam * rho * alpha
                - 2 * eta * g * lam * rho * alpha
                + 2 * H * eta ** 2 * rho * sigma * alpha
                + 2 * H * eta * lam * sigma * alpha
                + 2 * alpha * d * lam
                + 2 * alpha * g * lam
                + d ** 2
                + 2 * d * g
                + g ** 2
                + eta ** 2 * rho ** 2 * alpha ** 2
                + 2 * d * eta ** 2 * rho ** 2 * alpha
                + 2 * eta ** 2 * g * rho ** 2 * alpha
                - 2 * eta * lam * rho * alpha ** 2
                - 2 * d * eta * rho * alpha
                - 2 * eta * g * rho * alpha
            )
            ** (-0.1e1 / 0.2e1)
            * (
                2 * d_H_d_alpha * d * eta * sigma
                + 2 * d_H_d_alpha * eta * g * sigma
                - 4 * d_H_d_alpha * eta * lam * rho * sigma * alpha
                + 2 * H * eta ** 2 * sigma ** 2 * d_H_d_alpha
                + 2 * d_H_d_alpha * d * eta ** 2 * rho * sigma
                + 2 * d_H_d_alpha * eta ** 2 * g * rho * sigma
                + 2 * d_H_d_alpha * eta ** 2 * rho * sigma * alpha
                - 4 * d_H_d_alpha * d * eta * rho * sigma
                - 4 * d_H_d_alpha * eta * g * rho * sigma
                + 2 * d_H_d_alpha * eta * lam * sigma * alpha
            )
            / 2
        )
        * S
        * d_beta_d_alpha
        / 2
        + 1
        / (
            lam * sigma * alpha * H
            + alpha * d * lam
            + d * sigma * H
            + alpha * g * lam
            + g * sigma * H
            + alpha ** 2 * lam
            + d ** 2
            + 2 * d * g
            + d * alpha
            + g ** 2
            + alpha * g
        )
        * (
            eta * sigma * H
            + d * eta * rho
            + eta * g * rho
            + eta * rho * alpha
            + lam * alpha
            + d
            + g
            + sqrt(
                2 * d * eta ** 2 * g * rho ** 2
                - 4 * d * eta * g * rho
                + 2 * H * d * eta * sigma
                + 2 * H * eta * g * sigma
                - 4 * H * eta * lam * rho * sigma * alpha
                + lam ** 2 * alpha ** 2
                + d ** 2 * eta ** 2 * rho ** 2
                + eta ** 2 * g ** 2 * rho ** 2
                - 2 * d ** 2 * eta * rho
                - 2 * eta * g ** 2 * rho
                + H ** 2 * eta ** 2 * sigma ** 2
                + 2 * H * d * eta ** 2 * rho * sigma
                + 2 * H * eta ** 2 * g * rho * sigma
                - 4 * H * d * eta * rho * sigma
                - 4 * H * eta * g * rho * sigma
                - 2 * d * eta * lam * rho * alpha
                - 2 * eta * g * lam * rho * alpha
                + 2 * H * eta ** 2 * rho * sigma * alpha
                + 2 * H * eta * lam * sigma * alpha
                + 2 * alpha * d * lam
                + 2 * alpha * g * lam
                + d ** 2
                + 2 * d * g
                + g ** 2
                + eta ** 2 * rho ** 2 * alpha ** 2
                + 2 * d * eta ** 2 * rho ** 2 * alpha
                + 2 * eta ** 2 * g * rho ** 2 * alpha
                - 2 * eta * lam * rho * alpha ** 2
                - 2 * d * eta * rho * alpha
                - 2 * eta * g * rho * alpha
            )
        )
        * d_S_d_alpha
        * d_beta_d_alpha
        / 2
    )
    return M
```
