## Supplementary material for "Hyperparasitism and the evolution of parasite virulence": Code_base: zero_finder_utilities.py

```
"""
These functions are used to assist in the calculating of singular strategies.
"""
import sys
import runner
import numpy as np
from math import sqrt, exp, dist, floor, ceil, isnan
import plotly.graph_objects as go
from time import sleep
import matplotlib.pyplot as plt

import singular_strategy as ss

# sol = runner.PySolver()


def alpha_sigma_evaluater(
    sol,
    alpha,
    sigma,
    alpha_max_res=5,
    alpha_min_res=0.01,
    sigma_min_res=0,
    sigma_max_res=4,
    alpha_max=5,
    sigma_max=4,
    beta_max=5,
    resolution=100,
    rho=0.25,
    b=2.0,
    q=0.01,
    d=0.01,
    eta=0.25,
    lam=1.85,
    hyper=1.0,
    seed=31,
    c1=0.65,
    c2=-1.75,
    gamma=0.5,
):
    beta = beta_func(beta_max, alpha, alpha_max, sigma, sigma_max, c1, c2)
    dbetadalpha = dbetadalpha_func(beta_max, alpha, alpha_max, sigma, sigma_max, c1, c2)
    dbetadsigma = dbetadsigma_func(beta_max, alpha, alpha_max, sigma, sigma_max, c1, c2)

    y = sol.eco_steady_state(
        beta, alpha, sigma, b, q, d, rho, eta, gamma, lam, c1, c2, hyper, seed
    )

    S = y[0]
    I = y[1]
    H = y[2]
    #     temp_H.append(H)
    #             print(f"Run with alpha value {alpha} and sigma value {sigma}, with susceptible density {S}, parasitised density {I} and hyperparasitisted density {H}")
    if I == 0 and H == 0:
        alpha_grad = -1 * (alpha <= alpha_max / 2) + 1 * (alpha > alpha_max / 2)
        sigma_grad = -1 * (sigma <= sigma_max / 2) + 1 * (sigma > sigma_max / 2)
    elif H == 0:
        sigma_grad = dbetadsigma * S / (d + alpha + gamma)
        alpha_grad = dbetadalpha * S / (d + alpha + gamma) - (beta * S) / (
            (d + alpha + gamma) ** 2
        )
    #             #print(alpha,sigma,H)
    elif I == 0:
        sigma_grad = dbetadsigma * S / (d + lam * alpha + gamma)
        alpha_grad = dbetadalpha * S / (d + lam * alpha + gamma) - (beta * S) / (
            (d + lam * alpha + gamma) ** 2
        )
    else:
        sigma_grad = ss.fitness_gradient_sigma(
            beta, alpha, sigma, dbetadsigma, H, S, eta, lam, gamma, d, rho
        )
        alpha_grad = ss.fitness_gradient_alpha(
            beta, alpha, sigma, dbetadalpha, H, S, eta, lam, gamma, d, rho
        )

    alpha_grad = (alpha_grad > 0) * 1 + (alpha_grad < 0) * -1
    sigma_grad = (sigma_grad > 0) * 1 + (sigma_grad < 0) * -1

    return alpha_grad, sigma_grad, S, I, H


def beta_func(beta_max, alpha, alpha_max, sigma, sigma_max, c1, c2):

    if c2 == 0.0:
        beta = beta_max * (
            sqrt(alpha / alpha_max) * ((1 - c1) + c1 * (sigma / sigma_max))
        )
    else:
        beta = beta_max * (
            sqrt(alpha / alpha_max)
            * ((1 - c1) + c1 * ((1 - exp(c2 * (sigma / sigma_max))) / (1 - exp(c2))))
        )

    return beta


def dbetadalpha_func(beta_max, alpha, alpha_max, sigma, sigma_max, c1, c2):

    if c2 == 0.0:
        alpha_grad = (
            beta_max
            * ((alpha + 1e-10) / alpha_max) ** (-0.1e1 / 0.2e1)
            * (1 - c1 + c1 * sigma / sigma_max)
            / alpha_max
            / 2
        )
    else:
        alpha_grad = (
            beta_max
            * ((alpha + 1e-10) / alpha_max) ** (-0.1e1 / 0.2e1)
            * (1 - c1 + c1 * (1 - exp(sigma * c2 / sigma_max)) / (1 - exp(c2)))
            / alpha_max
            / 2
        )

    return alpha_grad


def dbetadsigma_func(beta_max, alpha, alpha_max, sigma, sigma_max, c1, c2):

    if c2 == 0.0:
        sigma_grad = beta_max * sqrt(alpha / alpha_max) * c1 / sigma_max
    else:
        sigma_grad = (
            -beta_max
            * sqrt(alpha / alpha_max)
            * c1
            * c2
            / sigma_max
            * exp(sigma * c2 / sigma_max)
            / (1 - exp(c2))
        )

    return sigma_grad


def get_zeros(grad_mat):

    poss_zeros = []

    for i, alpha_row in enumerate(grad_mat[:-1]):
        #         #print(len(alpha_row))
        alpha_up_row = grad_mat[i + 1]
        #         #print(len(alpha_up_row))
        for j, alpha_val_centre in enumerate(alpha_row[:-1]):
            sigma_val_centre = alpha_row[j + 1]
            alpha_up_val = alpha_up_row[j]
            sigma_up_val = alpha_up_row[j + 1]
            score = alpha_val_centre + sigma_val_centre + alpha_up_val + sigma_up_val
            if abs(score) != 4:
                poss_zeros.append([i, j])
    #             #print(score)
    return poss_zeros


def calculate_gradient_matrices(
    sol,
    alpha_max_res=5,
    alpha_min_res=0.01,
    sigma_min_res=0,
    sigma_max_res=4,
    alpha_max=5,
    sigma_max=4,
    beta_max=5,
    resolution=100,
    rho=0.25,
    b=2.0,
    q=0.01,
    d=0.01,
    eta=0.25,
    lam=1.85,
    hyper=1.0,
    seed=31,
    c1=0.65,
    c2=-1.75,
    gamma=0.5,
):

    # grad_mat_alpha = [[1*float("inf") for i in range(resolution+2)]]
    grad_mat_alpha = []
    grad_mat_sigma = []
    grad_mat_both = []
    H_mat = []
    # grad_mat_sigma = [[1*float("inf") for i in range(resolution+2)]]
    # grad_mat_both = [[1*float("inf") for i in range(resolution+2)]]
    # alpha_val = [0]
    # sigma_val = [0]
    alpha_val = []
    sigma_val = []

    alpha_zeros_x = []
    sigma_zeros_x = []
    alpha_zeros_y = []
    sigma_zeros_y = []

    for i in range(resolution + 1):
        alpha = alpha_min_res + (alpha_max_res - alpha_min_res) * (i / resolution)
        alpha_val.append(alpha)
        sigma_val.append(
            sigma_min_res + (sigma_max_res - sigma_min_res) * (i / resolution)
        )

        #     temp_alpha = [1*float("inf")]
        #     temp_sigma = [1*float("inf")]
        #     temp_both = [1*float("inf")]

        temp_alpha = []
        temp_sigma = []
        temp_both = []
        temp_H = []
        for j in range(resolution + 1):
            #         #print(i,j)
            sigma = sigma_min_res + (sigma_max_res - sigma_min_res) * (j / resolution)
            #             print(f"Run with alpha value {alpha} and sigma value {sigma}")
            #             beta = beta_func(beta_max, alpha, alpha_max, sigma, sigma_max, c1, c2)
            #             dbetadalpha = dbetadalpha_func(
            #                 beta_max, alpha, alpha_max, sigma, sigma_max, c1, c2
            #             )
            #             dbetadsigma = dbetadsigma_func(
            #                 beta_max, alpha, alpha_max, sigma, sigma_max, c1, c2
            #             )

            #             y = sol.eco_steady_state(
            #                 beta, alpha, sigma, b, q, d, rho, eta, gamma, lam, c1, c2, hyper, seed
            #             )

            #             S = y[0]
            #             I = y[1]
            #             H = y[2]
            #             temp_H.append(H)
            # #             print(f"Run with alpha value {alpha} and sigma value {sigma}, with susceptible density {S}, parasitised density {I} and hyperparasitisted density {H}")
            #             if (I == 0 and H == 0):
            #                 alpha_grad = -1*(alpha <= alpha_max/2) + 1*(alpha> alpha_max/2)
            #                 sigma_grad = -1*(sigma <= sigma_max/2) + 1*(sigma> sigma_max/2)
            #             elif (H == 0):
            #                 sigma_grad = dbetadsigma*S/(d + alpha + gamma)
            #                 alpha_grad = dbetadalpha*S/(d + alpha + gamma) - (beta*S)/((d + alpha + gamma)**2)
            #     #             #print(alpha,sigma,H)
            #             elif (I==0):
            #                 sigma_grad = dbetadsigma*S/(d + lam*alpha + gamma)
            #                 alpha_grad = dbetadalpha*S/(d + lam*alpha + gamma) - (beta*S)/((d + lam*alpha + gamma)**2)
            #             else:
            #                 sigma_grad = ss.fitness_gradient_sigma(
            #                     beta, alpha, sigma, dbetadsigma, H, S, eta, lam, gamma, d, rho
            #                 )
            #                 alpha_grad = ss.fitness_gradient_alpha(
            #                     beta, alpha, sigma, dbetadalpha, H, S, eta, lam, gamma, d, rho
            #                 )

            #             alpha_grad = (alpha_grad > 0)*1 + (alpha_grad < 0)*-1
            #             sigma_grad = (sigma_grad > 0)*1 + (sigma_grad < 0)*-1
            #         alpha_grad = max(-0.3,min(alpha_grad,0.3))
            #         sigma_grad = max(-0.3,min(sigma_grad,0.3))

            alpha_grad, sigma_grad, S, I, H = alpha_sigma_evaluater(
                sol,
                alpha,
                sigma,
                alpha_max_res=alpha_max_res,
                alpha_min_res=alpha_min_res,
                sigma_min_res=sigma_min_res,
                sigma_max_res=sigma_max_res,
                alpha_max=alpha_max,
                sigma_max=sigma_max,
                beta_max=beta_max,
                resolution=resolution,
                rho=rho,
                b=b,
                q=q,
                d=d,
                eta=eta,
                lam=lam,
                hyper=hyper,
                seed=seed,
                c1=c1,
                c2=c2,
                gamma=gamma,
            )
            pair_grad = alpha_grad + sigma_grad

            if abs(alpha_grad) < 1e-1:
                alpha_zeros_x.append(alpha)
                alpha_zeros_y.append(sigma)
            if abs(sigma_grad) < 1e-1:
                sigma_zeros_x.append(alpha)
                sigma_zeros_y.append(sigma)
            temp_alpha.append(alpha_grad)
            temp_sigma.append(sigma_grad)
            temp_both.append(pair_grad)
        #         H_mat.append(temp_H)
        #     temp_alpha.append(-1*float("inf"))
        #     temp_sigma.append(-1*float("inf"))
        #     temp_both.append(-1*float("inf"))

        grad_mat_alpha.append(temp_alpha)
        grad_mat_sigma.append(temp_sigma)
        grad_mat_both.append(temp_both)

    return grad_mat_alpha, grad_mat_sigma, grad_mat_both, alpha_val, sigma_val


def pairs_to_coords(poss_zeros_both, alpha_val, sigma_val):

    poss_alphas = []
    poss_sigmas = []
    for ind_pair in poss_zeros_both:
        poss_alphas.append([alpha_val[ind_pair[0]], alpha_val[ind_pair[0] + 1]])
        poss_sigmas.append([sigma_val[ind_pair[1]], sigma_val[ind_pair[1] + 1]])

    return poss_alphas, poss_sigmas


def poss_zeros_finder(
    sol,
    poss_alphas,
    poss_sigmas,
    alpha_max=5,
    sigma_max=4,
    beta_max=5,
    resolution=4,
    rho=0.25,
    b=2.0,
    q=0.01,
    d=0.01,
    eta=0.25,
    lam=1.85,
    hyper=1.0,
    seed=31,
    c1=0.65,
    c2=0.25,
    gamma=0.5,
):

    refined_alpha = []
    refined_sigma = []

    for i, alphas in enumerate(poss_alphas):
        alpha_lower = alphas[0]
        alpha_upper = alphas[1]
        sigma_lower = poss_sigmas[i][0]
        sigma_upper = poss_sigmas[i][1]

        (
            grad_mat_alpha_t,
            grad_mat_sigma_t,
            grad_mat_both_t,
            alpha_val_t,
            sigma_val_t,
        ) = calculate_gradient_matrices(
            sol=sol,
            alpha_max_res=alpha_upper,
            alpha_min_res=alpha_lower,
            sigma_min_res=sigma_lower,
            sigma_max_res=sigma_upper,
            beta_max=beta_max,
            alpha_max=alpha_max,
            sigma_max=sigma_max,
            resolution=resolution,
            rho=rho,
            b=b,
            q=q,
            d=d,
            eta=eta,
            lam=lam,
            hyper=hyper,
            seed=seed,
            c1=c1,
            c2=c2,
            gamma=gamma,
        )

        poss_zeros_alpha_t = get_zeros(grad_mat_alpha_t)
        poss_zeros_sigma_t = get_zeros(grad_mat_sigma_t)

        #         #print(alpha_lower, alpha_upper)
        #         #print(alpha_val_t)

        #         #print(sigma_lower, sigma_upper)
        #         #print(sigma_val_t)

        poss_zeros_both_t = []
        for inds in poss_zeros_alpha_t:
            if inds in poss_zeros_sigma_t:
                #                 print(inds)

                alpha_ind_check = inds[0]
                sigma_ind_check = inds[1]

                grad_alpha_row_1 = grad_mat_alpha_t[alpha_ind_check]
                grad_alpha_row_2 = grad_mat_alpha_t[alpha_ind_check + 1]
                grad_sigma_row_1 = grad_mat_sigma_t[alpha_ind_check]
                grad_sigma_row_2 = grad_mat_sigma_t[alpha_ind_check + 1]

                alpha_row_1_val_1 = grad_alpha_row_1[sigma_ind_check]
                alpha_row_1_val_2 = grad_alpha_row_1[sigma_ind_check + 1]
                sigma_row_1_val_1 = grad_sigma_row_1[sigma_ind_check]
                sigma_row_1_val_2 = grad_sigma_row_1[sigma_ind_check + 1]

                alpha_row_2_val_1 = grad_alpha_row_2[sigma_ind_check]
                alpha_row_2_val_2 = grad_alpha_row_2[sigma_ind_check + 1]
                sigma_row_2_val_1 = grad_sigma_row_2[sigma_ind_check]
                sigma_row_2_val_2 = grad_sigma_row_2[sigma_ind_check + 1]

                #                 print(alpha_row_1_val_1)
                #                 print(alpha_row_1_val_2)
                #                 print(sigma_row_1_val_1)
                #                 print(sigma_row_1_val_2)

                #                 print(alpha_row_2_val_1)
                #                 print(alpha_row_2_val_2)
                #                 print(sigma_row_2_val_1)
                #                 print(sigma_row_2_val_2)
                #                 if ((alpha_row_1_val_1 > 0  and alpha_row_2_val_1 < 0) and (sigma_row_1_val_1 > 0  and sigma_row_1_val_2 < 0)):
                poss_zeros_both_t.append(inds)

        #         for ind_pair in poss_zeros_both_t:
        #             for ii in [0,1]:
        #                 for jj in [0,1]:
        #                     poss_alphas.append(alpha_val_t[ind_pair[0] + ii])
        #                     poss_sigmas.append(sigma_val_t[ind_pair[1] + jj])

        poss_alpha_t, poss_sigma_t = pairs_to_coords(
            poss_zeros_both_t, alpha_val_t, sigma_val_t
        )

        for i, alpha_inds in enumerate(poss_alpha_t):
            refined_alpha.append(alpha_inds)
            refined_sigma.append(poss_sigma_t[i])

    return refined_alpha, refined_sigma


def get_edge_zeros(grad_mat_alpha, grad_mat_sigma, alpha_val, sigma_val):

    #     sigma_changes = [[i,sigma_val[-1]] for i in alpha_val]
    #     alpha_changes = [[alpha_val[-1], i] for i in sigma_val]

    #     for i in range(len(alpha_val)):
    #         sigma_changes.append([alpha_val[i],sigma_val[0]])
    #         alpha_changes.append([alpha_val[0],sigma_val[i]])

    edges = []
    for i, row in enumerate(grad_mat_alpha[:-1]):
        if grad_mat_alpha[i + 1][-1] != row[-1] and grad_mat_sigma[i][-1] > 0:
            edges.append([(alpha_val[i] + alpha_val[i + 1]) / 2, sigma_val[-1]])
        if grad_mat_alpha[i + 1][0] != row[0] and grad_mat_sigma[i][-1] < 0:
            edges.append([(alpha_val[i] + alpha_val[i + 1]) / 2, sigma_val[0]])

    sigma_row_1 = grad_mat_sigma[0]
    sigma_row_2 = grad_mat_sigma[-1]

    for j, val in enumerate(sigma_row_1[:-1]):
        if sigma_row_1[j + 1] != val and grad_mat_alpha[0][j] < 0:
            edges.append([alpha_val[0], (sigma_val[j] + sigma_val[j + 1]) / 2])
        if sigma_row_2[j + 1] != sigma_row_2[j] and grad_mat_alpha[-1][j] > 0:
            edges.append([alpha_val[-1], (sigma_val[j] + sigma_val[j + 1]) / 2])

    if grad_mat_alpha[0][0] < 0 and grad_mat_sigma[0][0] < 0:
        edges.append([alpha_val[0], sigma_val[0]])

    if grad_mat_alpha[-1][-1] > 0 and grad_mat_sigma[-1][-1] > 0:
        edges.append([alpha_val[-1], sigma_val[-1]])

    if grad_mat_alpha[-1][-1] > 0 and grad_mat_sigma[0][0] < 0:
        edges.append([alpha_val[-1], sigma_val[0]])

    if grad_mat_alpha[0][0] < 0 and grad_mat_sigma[-1][-1] > 0:
        edges.append([alpha_val[0], sigma_val[-1]])
    return edges


def get_edge_zeros_refined(
    sol,
    grad_mat_alpha,
    grad_mat_sigma,
    alpha_val,
    sigma_val,
    depth,
    depth_max=5,
    alpha_max=5,
    sigma_max=4,
    beta_max=5,
    resolution=4,
    rho=0.25,
    b=2.0,
    q=0.01,
    d=0.01,
    eta=0.25,
    lam=1.85,
    hyper=1.0,
    seed=31,
    c1=0.65,
    c2=0.25,
    gamma=0.5,
):
    edges = []
    if depth != depth_max:
        for i, row in enumerate(grad_mat_alpha[:-1]):
            if (
                row[-1] > 0
                and grad_mat_alpha[i + 1][-1] < 0
                and grad_mat_sigma[i][-1] > 0
            ):
                alpha_lower = alpha_val[i]
                alpha_upper = alpha_val[i + 1]
                sigma_lower = sigma_upper = sigma_val[-1]
                (
                    grad_mat_alpha_t,
                    grad_mat_sigma_t,
                    grad_mat_both_t,
                    alpha_val_t,
                    sigma_val_t,
                ) = calculate_gradient_matrices(
                    sol=sol,
                    alpha_max_res=alpha_upper,
                    alpha_min_res=alpha_lower,
                    sigma_min_res=sigma_lower,
                    sigma_max_res=sigma_upper,
                    beta_max=beta_max,
                    alpha_max=alpha_max,
                    sigma_max=sigma_max,
                    resolution=resolution,
                    rho=rho,
                    b=b,
                    q=q,
                    d=d,
                    eta=eta,
                    lam=lam,
                    hyper=hyper,
                    seed=seed,
                    c1=c1,
                    c2=c2,
                    gamma=gamma,
                )
                edges_temp = get_edge_zeros_refined(
                    sol=sol,
                    grad_mat_alpha=grad_mat_alpha_t,
                    grad_mat_sigma=grad_mat_sigma_t,
                    alpha_val=alpha_val_t,
                    sigma_val=sigma_val_t,
                    depth=depth + 1,
                    depth_max=depth_max,
                    alpha_max=alpha_max,
                    sigma_max=sigma_max,
                    beta_max=beta_max,
                    resolution=resolution,
                    rho=rho,
                    b=b,
                    q=q,
                    d=d,
                    eta=eta,
                    lam=lam,
                    hyper=hyper,
                    seed=seed,
                    c1=c1,
                    c2=c2,
                    gamma=gamma,
                )
                for edge in edges_temp:
                    edges.append(edge)
            if row[0] > 0 and grad_mat_alpha[i + 1][0] < 0 and grad_mat_sigma[i][0] < 0:
                alpha_lower = alpha_val[i]
                alpha_upper = alpha_val[i + 1]
                sigma_lower = sigma_upper = sigma_val[0]
                (
                    grad_mat_alpha_t,
                    grad_mat_sigma_t,
                    grad_mat_both_t,
                    alpha_val_t,
                    sigma_val_t,
                ) = calculate_gradient_matrices(
                    sol=sol,
                    alpha_max_res=alpha_upper,
                    alpha_min_res=alpha_lower,
                    sigma_min_res=sigma_lower,
                    sigma_max_res=sigma_upper,
                    beta_max=beta_max,
                    alpha_max=alpha_max,
                    sigma_max=sigma_max,
                    resolution=resolution,
                    rho=rho,
                    b=b,
                    q=q,
                    d=d,
                    eta=eta,
                    lam=lam,
                    hyper=hyper,
                    seed=seed,
                    c1=c1,
                    c2=c2,
                    gamma=gamma,
                )
                edges_temp = get_edge_zeros_refined(
                    sol=sol,
                    grad_mat_alpha=grad_mat_alpha_t,
                    grad_mat_sigma=grad_mat_sigma_t,
                    alpha_val=alpha_val_t,
                    sigma_val=sigma_val_t,
                    depth=depth + 1,
                    depth_max=depth_max,
                    alpha_max=alpha_max,
                    sigma_max=sigma_max,
                    beta_max=beta_max,
                    resolution=resolution,
                    rho=rho,
                    b=b,
                    q=q,
                    d=d,
                    eta=eta,
                    lam=lam,
                    hyper=hyper,
                    seed=seed,
                    c1=c1,
                    c2=c2,
                    gamma=gamma,
                )
                for edge in edges_temp:
                    edges.append(edge)

        sigma_row_1 = grad_mat_sigma[0]
        sigma_row_2 = grad_mat_sigma[-1]

        for j, val in enumerate(sigma_row_1[:-1]):
            if sigma_row_1[j + 1] < 0 and val > 0 and grad_mat_alpha[0][j] < 0:
                alpha_lower = alpha_val[0]
                alpha_upper = alpha_val[0]
                sigma_lower = sigma_val[j]
                sigma_upper = sigma_val[j + 1]
                (
                    grad_mat_alpha_t,
                    grad_mat_sigma_t,
                    grad_mat_both_t,
                    alpha_val_t,
                    sigma_val_t,
                ) = calculate_gradient_matrices(
                    sol=sol,
                    alpha_max_res=alpha_upper,
                    alpha_min_res=alpha_lower,
                    sigma_min_res=sigma_lower,
                    sigma_max_res=sigma_upper,
                    beta_max=beta_max,
                    alpha_max=alpha_max,
                    sigma_max=sigma_max,
                    resolution=resolution,
                    rho=rho,
                    b=b,
                    q=q,
                    d=d,
                    eta=eta,
                    lam=lam,
                    hyper=hyper,
                    seed=seed,
                    c1=c1,
                    c2=c2,
                    gamma=gamma,
                )
                edges_temp = get_edge_zeros_refined(
                    sol=sol,
                    grad_mat_alpha=grad_mat_alpha_t,
                    grad_mat_sigma=grad_mat_sigma_t,
                    alpha_val=alpha_val_t,
                    sigma_val=sigma_val_t,
                    depth=depth + 1,
                    depth_max=depth_max,
                    alpha_max=alpha_max,
                    sigma_max=sigma_max,
                    beta_max=beta_max,
                    resolution=resolution,
                    rho=rho,
                    b=b,
                    q=q,
                    d=d,
                    eta=eta,
                    lam=lam,
                    hyper=hyper,
                    seed=seed,
                    c1=c1,
                    c2=c2,
                    gamma=gamma,
                )
                for edge in edges_temp:
                    edges.append(edge)
            #                 edges.append([alpha_val[0],(sigma_val[j] + sigma_val[j+1])/2])
            if (
                sigma_row_2[j + 1] < 0
                and sigma_row_2[j] > 0
                and grad_mat_alpha[-1][j] > 0
            ):
                alpha_lower = alpha_val[-1]
                alpha_upper = alpha_val[-1]
                sigma_lower = sigma_val[j]
                sigma_upper = sigma_val[j + 1]
                (
                    grad_mat_alpha_t,
                    grad_mat_sigma_t,
                    grad_mat_both_t,
                    alpha_val_t,
                    sigma_val_t,
                ) = calculate_gradient_matrices(
                    sol=sol,
                    alpha_max_res=alpha_upper,
                    alpha_min_res=alpha_lower,
                    sigma_min_res=sigma_lower,
                    sigma_max_res=sigma_upper,
                    beta_max=beta_max,
                    alpha_max=alpha_max,
                    sigma_max=sigma_max,
                    resolution=resolution,
                    rho=rho,
                    b=b,
                    q=q,
                    d=d,
                    eta=eta,
                    lam=lam,
                    hyper=hyper,
                    seed=seed,
                    c1=c1,
                    c2=c2,
                    gamma=gamma,
                )
                edges_temp = get_edge_zeros_refined(
                    sol=sol,
                    grad_mat_alpha=grad_mat_alpha_t,
                    grad_mat_sigma=grad_mat_sigma_t,
                    alpha_val=alpha_val_t,
                    sigma_val=sigma_val_t,
                    depth=depth + 1,
                    depth_max=depth_max,
                    alpha_max=alpha_max,
                    sigma_max=sigma_max,
                    beta_max=beta_max,
                    resolution=resolution,
                    rho=rho,
                    b=b,
                    q=q,
                    d=d,
                    eta=eta,
                    lam=lam,
                    hyper=hyper,
                    seed=seed,
                    c1=c1,
                    c2=c2,
                    gamma=gamma,
                )
                for edge in edges_temp:
                    edges.append(edge)
    #                 edges.append([alpha_val[-1],(sigma_val[j] + sigma_val[j+1])/2])
    else:
        edges_temp = get_edge_zeros(
            grad_mat_alpha, grad_mat_sigma, alpha_val, sigma_val
        )
        for edge in edges_temp:
            edges.append(edge)

    if depth == 0:
        if grad_mat_alpha[0][0] < 0 and grad_mat_sigma[0][0] < 0:
            edges.append([alpha_val[0], sigma_val[0]])

        if grad_mat_alpha[-1][-1] > 0 and grad_mat_sigma[-1][-1] > 0:
            edges.append([alpha_val[-1], sigma_val[-1]])

        if grad_mat_alpha[-1][-1] > 0 and grad_mat_sigma[0][0] < 0:
            edges.append([alpha_val[-1], sigma_val[0]])

        if grad_mat_alpha[0][0] < 0 and grad_mat_sigma[-1][-1] > 0:
            edges.append([alpha_val[0], sigma_val[-1]])

    return edges


def population_boundaries(H_mat):

    H_plotting_sigma = []
    H_plotting_alpha = []
    for i, row in enumerate(H_mat):
        down = -1 * (i != 0)
        up = 1 * (i != (len(H_mat) - 1))
        i_vec = [down, 0, up]
        for j, val in enumerate(row):
            left = -1 * (j != 0)
            right = 1 * (j != (len(row) - 1))
            j_vec = [left, 0, right]
            H_sum = 0
            for ii in i_vec:
                for jj in j_vec:
                    if ii != 0 and jj != 0:
                        H_sum += H_mat[i + ii][j + jj]

            if val == 0 and H_sum != 0:
                H_plotting_alpha.append(i)
                H_plotting_sigma.append(j)

    return H_plotting_alpha, H_plotting_sigma


def plotting_vectors_to_inds(
    plotting_alphas, plotting_sigmas, alpha_val, sigma_val, resolution
):

    alpha_scatter_cords = []
    for val in plotting_alphas:
        if val == alpha_val[-1]:
            alpha_scatter_cords.append(resolution - 1)
        elif val == alpha_val[0]:
            alpha_scatter_cords.append(0)
        else:
            i_temp = 0
            while alpha_val[i_temp] < val:
                i_temp += 1
            alpha_diff = alpha_val[i_temp] - alpha_val[i_temp - 1]
            shift = val - alpha_val[i_temp - 1]
            approximate_index = i_temp - 1 + (shift / alpha_diff)
            alpha_scatter_cords.append(approximate_index)

    sigma_scatter_cords = []
    for val in plotting_sigmas:
        if val == sigma_val[-1]:
            sigma_scatter_cords.append(resolution - 1)
        elif val == sigma_val[0]:
            sigma_scatter_cords.append(0)
        else:
            i_temp = 0
            while sigma_val[i_temp] < val:
                i_temp += 1
            sigma_diff = sigma_val[i_temp] - sigma_val[i_temp - 1]
            shift = val - sigma_val[i_temp - 1]
            approximate_index = i_temp - 1 + (shift / sigma_diff)
            sigma_scatter_cords.append(approximate_index)

    return alpha_scatter_cords, sigma_scatter_cords

def clean_up_sing_strats(attractors_hyper_temp):
    
    attractors_hyper = []
    for val in attractors_hyper_temp:
        flag = True
        for i,check_val in enumerate(attractors_hyper):
            if abs(val - check_val) < 1e-1:
                flag = False
                break
        if flag:
            attractors_hyper.append(val)
            
    return attractors_hyper
```
