## Supplementary_Material for "Hyperparasitism and the evolution of parasite virulence"

### Supplementary Material:
### Hyperparasitism and the evolution of parasite virulence

#### S1. Parasite evolution in the absence of the hyperparasite

In the absence of the hyperparasite, the dynamics of a rare mutant parasite in a resident population at equilibrium (denoted by asterisks) are,

$$\dot{I_m} = \big(\beta(\alpha_m)S^* - (d + \alpha_m + \gamma)\big)I_m. \tag{S1}$$

The invasion fitness is then simply the per-capita growth rate,

$$w(\alpha_m, \alpha) = \beta(\alpha_m)S^* - (d + \alpha_m + \gamma) \tag{S2}$$

with fitness gradient,

$$\left.\frac{\partial w(\alpha_m, \alpha)}{\partial \alpha_m}\right|_{\alpha_m=\alpha} = \frac{d\beta}{d\alpha}S^* - 1. \tag{S3}$$

The singular strategy is defined when the fitness gradient vanishes, and so the singular strategy, $\alpha_0$, satisfies

$$\left.\frac{d\beta}{d\alpha}\right|_{\alpha=\alpha_0} = \frac{1}{S^*} = \frac{\beta(\alpha_0)}{d + \alpha_0 + \gamma}. \tag{S4}$$

The singular strategy is evolutionarily stable when

$$E = \left.\frac{\partial^2 w(\alpha_m, \alpha)}{\partial \alpha_m^2}\right|_{\alpha_m=\alpha=\alpha_0} = \left.\frac{d^2\beta}{d\alpha^2}\right|_{\alpha=\alpha_0} \frac{d + \alpha_0 + \gamma}{\beta(\alpha_0)} < 0 \tag{S5}$$

which requires $\left.\frac{d^2\beta}{d\alpha^2}\right|_{\alpha=\alpha_0} < 0$ (i.e. there must be diminishing returns in the trade-off between transmission and virulence). Convergence stability is guaranteed if $E + M < 0$, where,

$$M = \left.\frac{\partial^2 w(\alpha_m, \alpha)}{\partial \alpha_m \partial \alpha}\right|_{\alpha_m=\alpha=\alpha_0} = \left.\frac{d\beta}{d\alpha}\right|_{\alpha=\alpha_0} \left.\frac{dS^*}{d\alpha}\right|_{\alpha=\alpha_0} = \left.\frac{d\beta}{d\alpha}\right|_{\alpha=\alpha_0} \left(\frac{d + \alpha_0 + \gamma}{\beta(\alpha_0)^2}\left.\frac{d\beta}{d\alpha}\right|_{\alpha=\alpha_0} - \frac{1}{\beta(\alpha_0)}\right) \tag{S6}$$

By substituting in $\left.\frac{d\beta}{d\alpha}\right|_{\alpha=\alpha_0} = \frac{\beta(\alpha_0)}{d+\alpha_0+\gamma}$, we have

$$M = \left.\frac{d\beta}{d\alpha}\right|_{\alpha=\alpha_0} \left(\frac{d+\alpha_0+\gamma}{\beta(\alpha_0)^2} \left.\frac{d\beta}{d\alpha}\right|_{\alpha=\alpha_0} - \frac{1}{\beta(\alpha_0)}\right) = \left.\frac{d\beta}{d\alpha}\right|_{\alpha=\alpha_0} \left(\frac{1}{\beta(\alpha_0)} - \frac{1}{\beta(\alpha_0)}\right) = 0. \tag{S7}$$

Hence the singular strategy is convergence stable if it is evolutionarily stable, which for any trade-off with diminishing returns $\left(\frac{d^2\beta}{d\alpha^2} < 0\right)$ is true.

#### S2. Ecological dynamics of the host-parasite-hyperparasite system

We investigate the ecology of the system to understand when a hyperparasite can invade and to determine the stability of the ecological equilibria. The ecological dynamics in the absence of the hyperparasite are,

$$\dot{S} = (b - qN)N - (\beta I + d)S + \gamma I \tag{S8}$$
$$\dot{I} = \big(\beta S - (d + \alpha + \gamma)\big)I \tag{S9}$$

where $N = S + I$. The endemic steady state for the susceptible population is then

$$S^* = \frac{d + \alpha + \gamma}{\beta}. \tag{S10}$$

and for the infected population is the positive solution to the quadratic

$$I^{*2} - \frac{b - 2qS^* - \beta S^* + \gamma}{q} I^* - \frac{bS^* - dS^* - qS^{*2}}{q} = 0 \tag{S11}$$

which we omit here for brevity. The ecological dynamics in the presence of the hyperparasite are,

$$\dot{S} = (b - qN)N - (\beta I + \eta\beta H + d)S + \gamma(I + H) \tag{S12}$$
$$\dot{I} = \big(\beta S - \sigma H - (d + \alpha + \gamma)\big)I + (1 - \rho)\eta\beta S H \tag{S13}$$
$$\dot{H} = \big(\rho\eta\beta S + \sigma I - (d + \lambda\alpha + \gamma)\big)H \tag{S14}$$

with Jacobian

$$J = \begin{pmatrix} -2qN + b - \beta I - \eta\beta H - d & -2qN + b - \beta S + \gamma & -2qN + b - \eta\beta S + \gamma \\ \beta I + (1-\rho)\eta\beta H & \beta S - \sigma H - (d + \alpha + \gamma) & -\sigma I + (1-\rho)\eta\beta S \\ \rho\eta\beta H & \sigma H & \rho\eta\beta S + \sigma I - (d + \lambda\alpha + \gamma) \end{pmatrix} \tag{S15}$$

where now, $N = S + I + H$.

Substituting in the steady state in the absence of the hyperparasite (from equations S10 and S11), we can calculate the eigenvalues of the Jacobian above to check whether the hyperparasite can invade (i.e., is the hyperparasite-free equilibrium unstable?). For instance, for the parameter set in

Table 1, with $\eta = 0.5, \lambda = 0.5, \alpha = 1.5, \beta = 2.0, \rho = 0.5$, the real part of the dominant eigenvalue is $\omega = \sim 11.20 > 0$ and so the hyperparasite can invade.

This shows when the hyperparasite can invade, but not the stability of the non-trivial coexistence equilibrium with all species present. We therefore numerically explore the ecological dynamics of the system by initialising a population of hyperparasites at an arbitrarily small density, with the host and parasites at the steady state in the absence of the hyperparasite. We then use a fourth-order Runge-Kutta method to solve the system of differential equations to approximate the steady state and record the species that are present above a threshold of $10^{-3}$ (Fig. S1). No limit cycles were observed in our numerical analysis.

#### S3. Derivation of parasite invasion fitness

The dynamics of rare mutant parasites with virulence $\alpha_m$ and transmission $\beta_m = \beta(\alpha_m)$ in a resident population at equilibrium (denoted by asterisks) are:

$$\dot{I_m} = \big(\beta_m S^* - \sigma H^* - (d + \alpha_m + \gamma)\big) I_m + (1-\rho)\eta\beta_m S^* H_m, \quad \text{(S16)}$$

$$\dot{H_m} = \big(\rho\eta\beta_m S^* - (d + \lambda\alpha_m + \gamma)\big) H_m + \sigma I_m H^*. \quad \text{(S17)}$$

The Jacobian is then:

$$J = \begin{bmatrix} \beta_m S^* - \sigma H^* - (d + \alpha_m + \gamma) & (1-\rho)\eta\beta_m S^* \\ \sigma H^* & \rho\eta\beta_m S^* - (d + \lambda\alpha_m + \gamma) \end{bmatrix} \quad \text{(S18)}$$

We separate the Jacobian into two sub-matrices, $F$ and $V$, such that $J = F - V$. We follow the convention that $F$ corresponds to all newly created parasites, and $V$ contains movement between classes and loss of parasites:

$$F = \begin{bmatrix} \beta_m S^* & (1-\rho)\eta\beta_m S^* \\ 0 & \rho\eta\beta_m S^* \end{bmatrix} \quad \text{(S19)}$$

$$V = \begin{bmatrix} \sigma H^* + (d + \alpha_m + \gamma) & 0 \\ -\sigma H^* & (d + \lambda\alpha_m + \gamma) \end{bmatrix} \quad \text{(S20)}$$

Taking the inverse of $V$ we obtain,

$$V^{-1} = \begin{bmatrix} \dfrac{1}{\sigma H^* + d + \alpha + \gamma} & 0 \\ \dfrac{\sigma H}{(\sigma H^* + d + \alpha + \gamma)(d + \lambda\alpha + \gamma)} & \dfrac{1}{d + \lambda\alpha + \gamma} \end{bmatrix} \quad \text{(S21)}$$

The next generation matrix $NG$ is calculated as $NG = FV^{-1}$

$$NG = \begin{bmatrix} \frac{\beta_m S^*}{\sigma H^* + d + \alpha_m + \gamma} + \frac{(1-\rho)\eta\sigma\beta_m S^* H^*}{(\sigma H^* + d + \alpha_m + \gamma)(d + \lambda\alpha_m + \gamma)} & \frac{(1-\rho)\eta\beta_m S^*}{d + \lambda\alpha + \gamma} \\ \frac{\rho\eta\sigma\beta_m S^* H^*}{(\sigma H^* + d + \alpha_m + \gamma)(d + \lambda\alpha_m + \gamma)} & \frac{\rho\eta\beta_m S^*}{d + \lambda\alpha + \gamma} \end{bmatrix} \quad \text{(S22)}$$

Invasion fitness is sign-equivalent to the largest eigenvalue of this matrix minus 1, which we omit here for brevity.

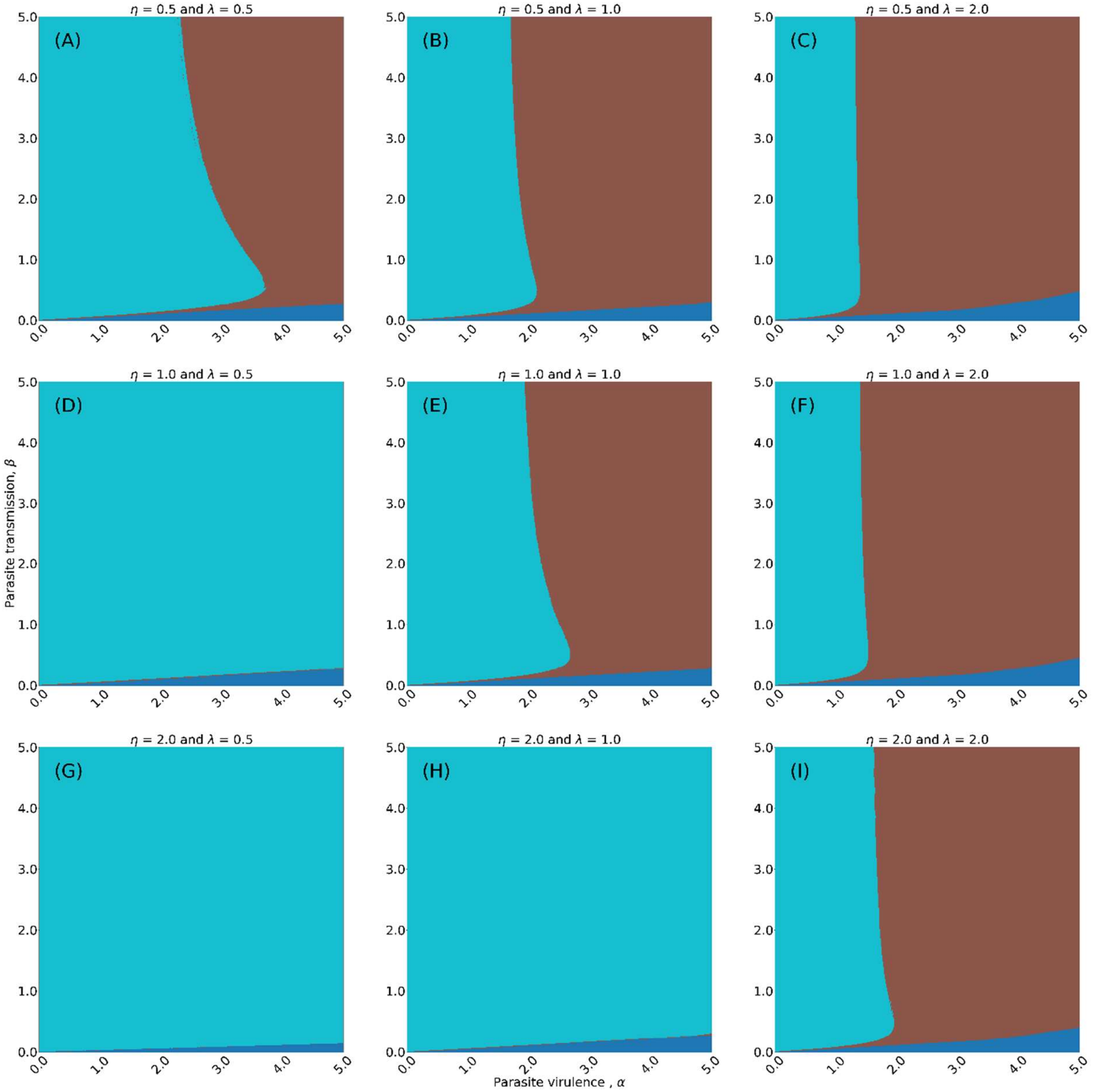


*Figure S1: Ecological states at steady state for different combinations of Parasite virulence $\alpha$, parasite transmission $\beta$, hyperparasite transmission modifier $\eta$, and hyperparasite virulence modifier $\lambda$. Here parameters are as in Table 1 except for $d = 0.1, \gamma = 0.1, \sigma = 0.4, \rho = 0.5$. Light blue colour denotes the presence of hosts, parasites and hyperparasites, brown denote only hosts and parasites present, and dark blue denotes only hosts present.*

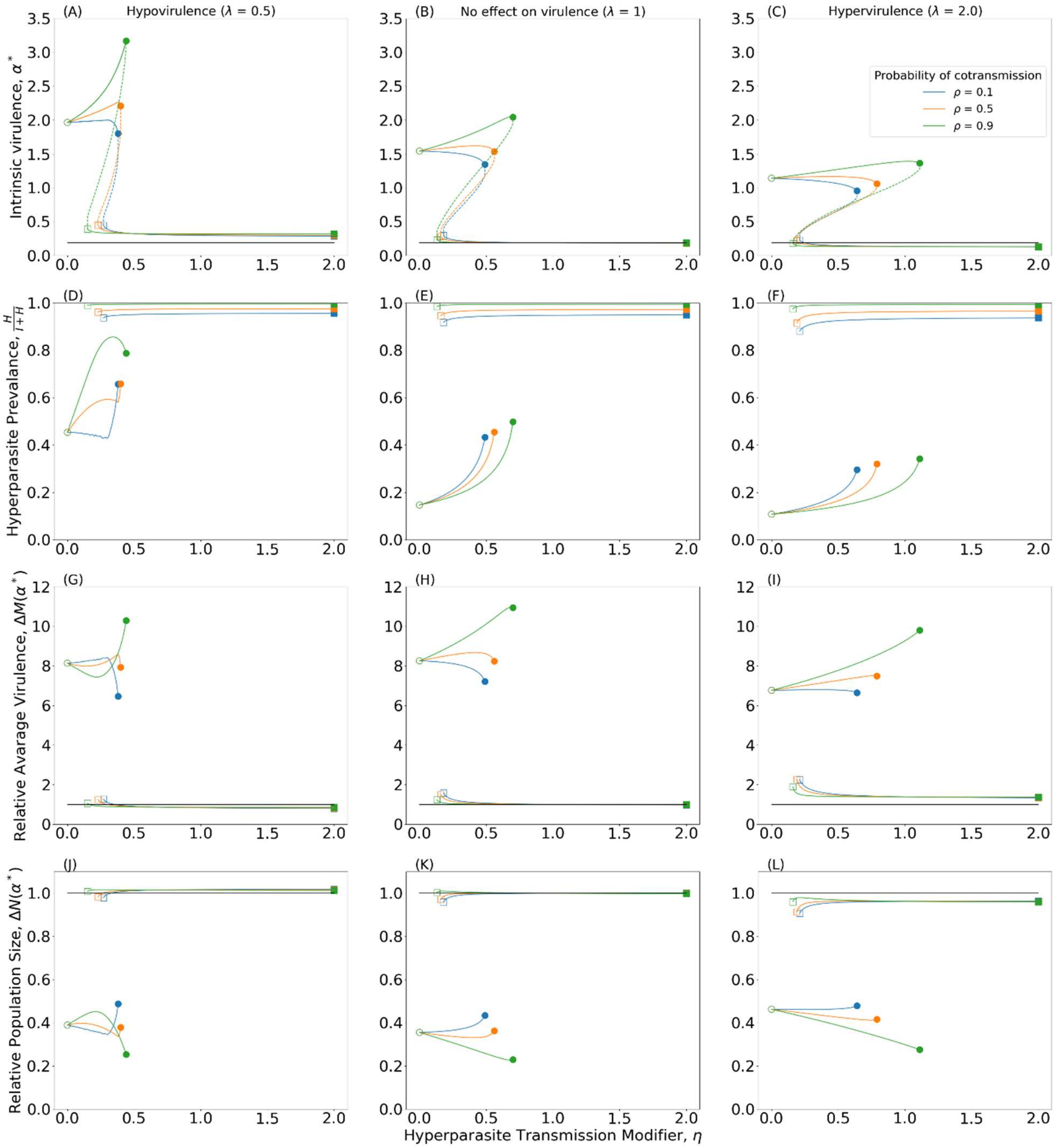


*Figure S2: Evolutionary consequences for parasite virulence (A-C) (solid: evolutionary attractors, dashed: evolutionary repellers), the prevalence of the hyperparasite (D-E), relative average mortality (G-I) and relative population size (J-L), as the direct effects of the hyperparasite on parasite transmission (hypotransmission: $\eta < 1$; hypertransmission: $\eta > 1$) and virulence ($\lambda$) vary. Evolutionary attractors are both convergence stable and evolutionarily stable and are therefore continuously stable strategies (CSSs). Left column - hypovirulence ($\lambda < 1$); Central column - no effect on virulence ($\lambda = 1$); Right column - hypervirulence ($\lambda > 1$). All panels contain three sets of curves showing the evolutionary endpoint as the probability of hyperparasite co-transmission varies: $\rho = 0.1$ (blue), $\rho = 0.5$ (orange), $\rho = 0.9$ (green). The start and end of each continuous set of evolutionary attractors are shown with empty and filled shapes. The black line indicates the ancestral state prior to the introduction of the hyperparasite. Parameters as in Table 1 except d=0.1, σ=0.4, γ=0.1. Trade-off used described in Equation 3.*

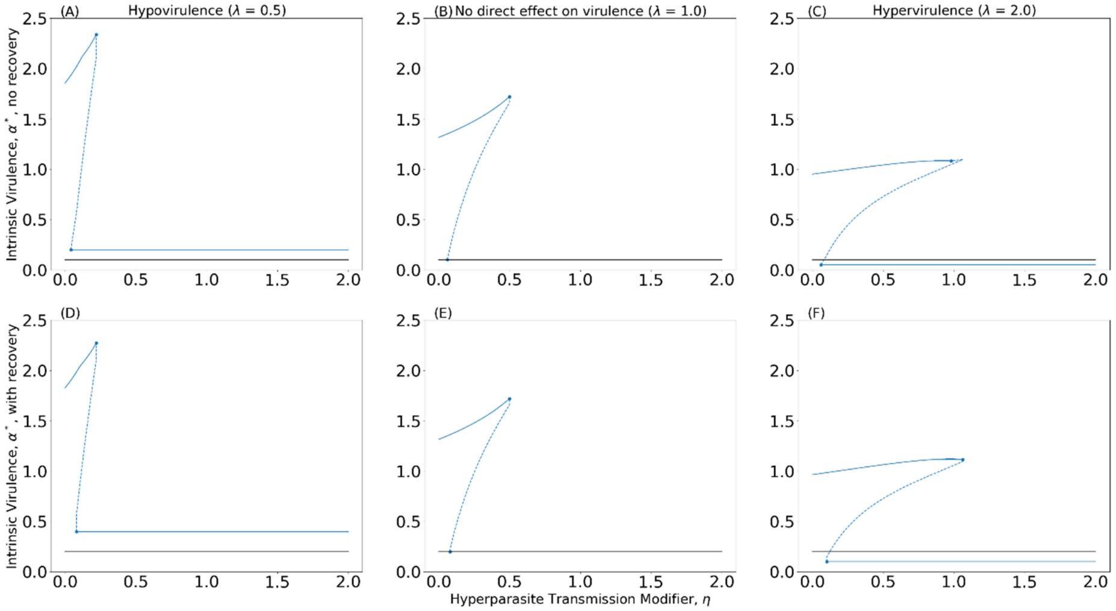


*Figure S3: Evolutionary consequences for parasite virulence without recovery (A-C) and with recovery (D-F), when $\rho = 1$, as the direct effects of the hyperparasite on parasite transmission (hypotransmission: η<1; hypertransmission: η>1) and virulence (λ) vary. Evolutionary attractors are both convergence stable and evolutionarily stable and are therefore continuously stable strategies (CSSs). The start and end of each continuous set of evolutionary attractors are shown with empty and filled shapes. The black line indicates the ancestral state prior to the introduction of the hyperparasite. Parameters as in Table 1 except d=0.1, σ=0.4, γ=0.1. Trade-off used described in Equation 2.*

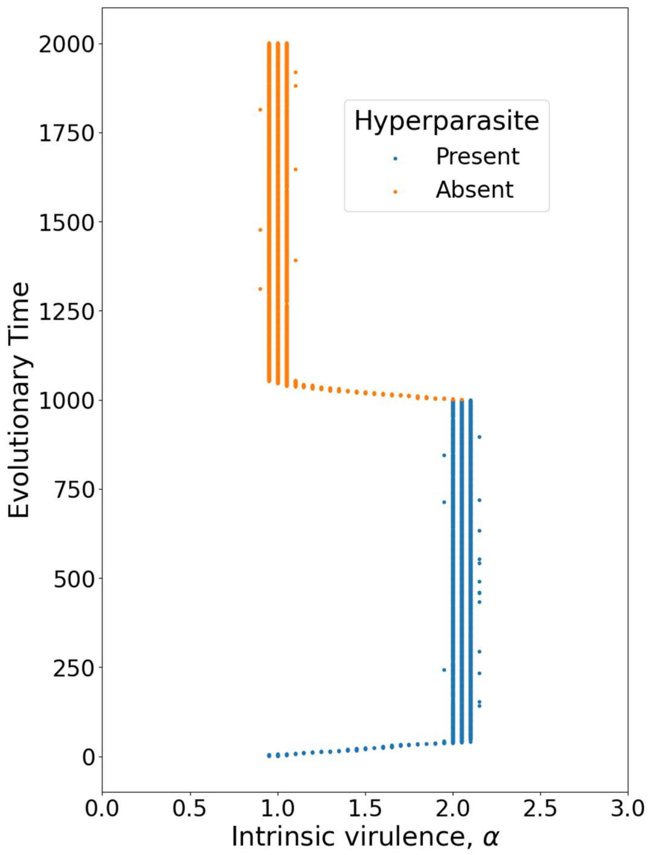


*Figure S4: Simulation showing that after the artificial removal of the hyperparasite, the parasite returns to its ancestral level of virulence (equation S4). Trade-off used described in Equation 2. Parameters as in Table 1, except* $\eta = 0.3, \lambda = 2.0, \rho = 0.9$.

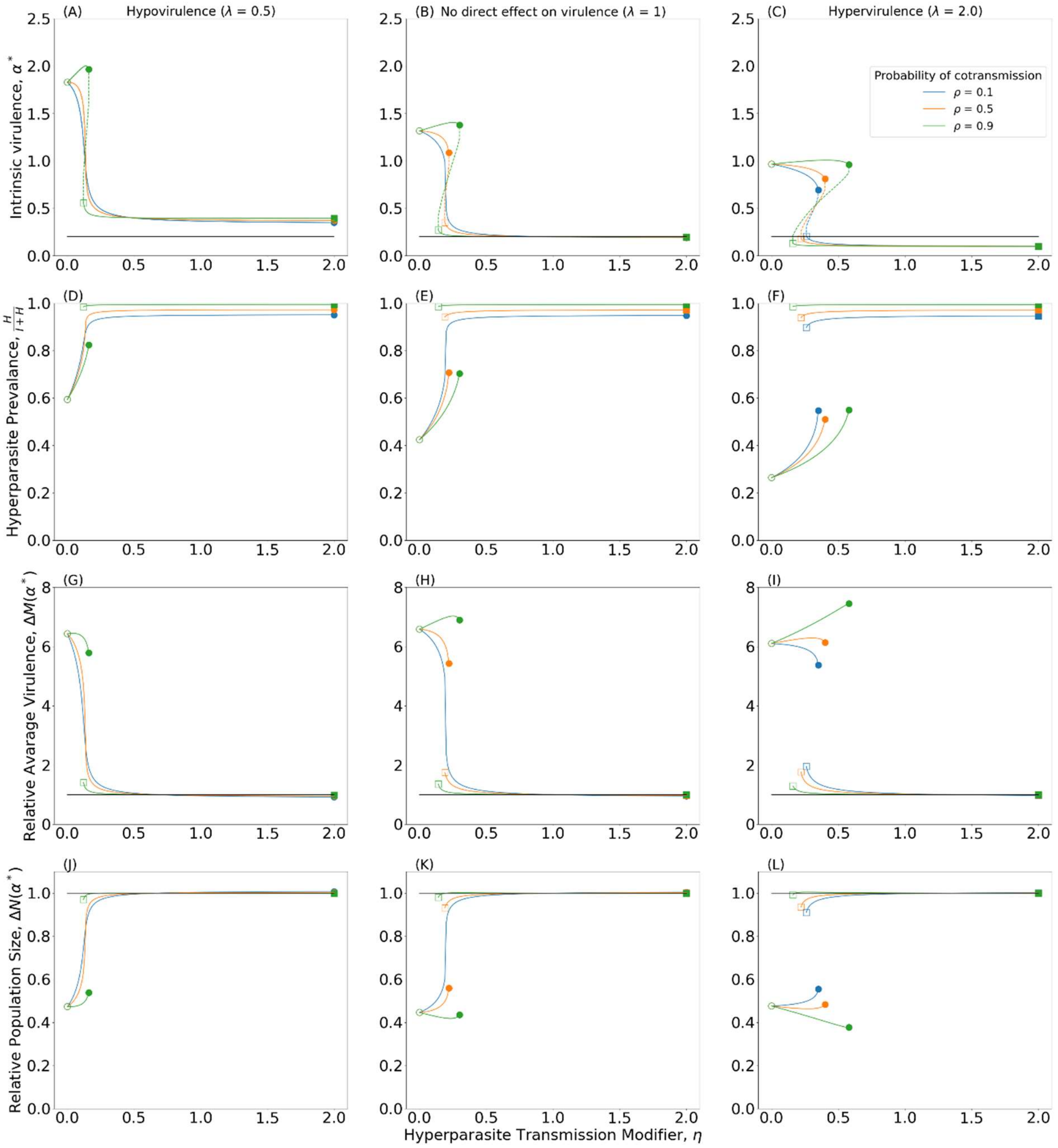


*Figure S5: Evolutionary consequences for parasite virulence (A-C) (solid: evolutionary attractors, dashed: evolutionary repellers), the prevalence of the hyperparasite (D-E), relative average mortality (G-I) and relative population size (J-L), as the direct effects of the hyperparasite on parasite transmission (hypotransmission: $\eta < 1$; hypertransmission: $\eta > 1$) and virulence ($\lambda$) vary. Evolutionary attractors are both convergence stable and evolutionarily stable and are therefore continuously stable strategies (CSSs). Left column - hypovirulence (λ<1); Central column - no effect on virulence (λ=1); Right column - hypervirulence (λ>1). All panels contain three sets of curves showing the evolutionary endpoint as the probability of hyperparasite co-transmission varies: $\rho = 0.1$ (blue), $\rho = 0.5$ (orange), $\rho = 0.9$ (green). The start and end of each continuous set of evolutionary attractors are shown with empty and filled shapes. The black line indicates the ancestral state prior to the introduction of the hyperparasite. Parameters as in Table 1 except d=0.1, σ=0.4, γ=0.1. Trade-off here described in Equation 2.*
